# Falsifiable substitution tests reveal task-structured neural evidence for auditory attention

**DOI:** 10.64898/2026.08.03.742580

**Authors:** Yu Ding

## Abstract

A neural decoder can predict a mental-state label without using information specific to that state. We made auditory-attention attribution falsifiable by requiring candidate evidence to persist in disjoint data, respond to capacity-matched substitutions of physical organization or listener/population template, and remain testable after target-event exclusion or command-identity residualization; two event-related datasets also permitted electrooculography (EOG)-only comparisons. The design drew on Wang and Zahl’s three-dimensional Kakeya proof strategy: examine the organized family and its concentration, not only the strongest member. Across six EEG datasets, averaging four neural–speech margins improved 5-s decoding relative to the leading margin in three evaluation sets whose rules were fixed before their results were computed (41 participants; study-equal gain, 0.0201; 95% interval, 0.0125–0.0279). A 16-cell scalp-direction–delay representation replicated in a participant-disjoint cohort and exceeded the mean of 15 capacity-matched remappings. Across three continuous-speech datasets (43 participants; 86 directed transfers), listener-matched weights outranked other-listener weights by 0.0969 and wrong mappings by 0.1371, although accuracy did not improve universally. In two hierarchical interfaces, a parent-stream error score retained AUCs of 0.968 and 0.965 after oracle-label exclusion of all target-command events. It depended on the physical command–stream mapping, exceeded an EOG-only comparator, and generalized within listeners after training-only removal of command identity. Eight electrodes retained 59–77% of binding specificity, but one listener-consistency criterion failed. The main contribution is a transferable standard for testing what information supports a decoded psychological construct.

**Significance Statement:** Inspired by the proof strategy of the three-dimensional Kakeya theorem, we turn “a neural decoder reads auditory attention” from an interpretation of accuracy into a falsifiable test of evidence attribution. Engineering can exploit any stable predictor; science of latent mental constructs must ask whether the proposed construct remains necessary after plausible alternatives are removed or substituted. Across six electroencephalography (EEG) datasets, task-organized scores survived disjoint data and were challenged by matched substitutions of physical mapping or listener template, target-event exclusion, command-identity residualization, and EOG-only comparison. This framework does not prove that attention is the only cause. It offers neuroscience and brain–computer interfaces (BCIs) a standard: evidence should transport, its proposed organization should matter, and credible shortcuts should fail.

---

At a cocktail party, the ears receive several voices at once, yet perception follows one talker. Electroencephalography (EEG) and intracranial recordings show that cortical activity follows attended speech more closely than competing speech (1–4). Auditory-attention decoding (AAD) uses this difference to infer the selected talker or command, and the field now includes wearable sensors, cross-listener models, and closed-loop hearing systems (5–10); a recent intracranial system improved speech intelligibility and listening effort in real time (8). Engineering and cognitive neuroscience, however, ask different questions of such a decoder. An engineering system may profitably use every stable cue that improves prediction. Attention and consciousness are latent constructs: they are inferred rather than directly observed in a voltage trace. For construct inference, accuracy alone is therefore insufficient. The claimed construct should remain necessary after plausible alternative sources of prediction have been removed or substituted. The aim is not to erase every signal, but to discover which evidence survives controlled removal and which interpretation fails when its proposed source is changed.

The three-dimensional Kakeya problem offers an unusually apt strategy for this identifiability problem. A Kakeya set contains a unit line segment in every direction; Wang and Zahl proved that in three dimensions it must have full Hausdorff and Minkowski dimension (11–12). The obstacle is overlap: many thin tubes can pass through the same region, so a compact union can conceal the breadth of the directional family that produced it. The proof therefore controls the family rather than one extremal tube, quantifies concentration and overlap multiplicity inside convex regions, organizes the family across scales, and reduces the general problem to structured “sticky” configurations (11– 13). Its methodological lesson is that the same large or compact summary can be compatible with two explanations—a genuinely organized whole or concentrated overlap—and that the explanations become distinguishable only when organization, alternatives, and scale are tested. This is a statement about identifiability, not a claim about the ontology of attention or consciousness.

The mathematical and neural uses of “direction” are not identical. In Kakeya geometry, direction is the orientation of a line segment or tube. Borrowing the proof’s family-level indexing idea, we define a **neural direction family** as a set of projections or contrasts indexed by an experimentally meaningful coordinate—for example, scalp sector and response delay, or auditory stream and command (Fig. 1). We use **physical organization** to mean that this coordinate relation is fixed by the experimental arrangement rather than by the observed outcome: which sector is paired with which delay, or which command belongs to which pitch-defined stream. Four tests follow. Does the family retain information lost by its leading member? Is apparent breadth concentrated in a few responses? Does the proposed organization beat alternatives with the same number of elements and fitted quantities? Does the evidence transport to disjoint material or another sensor scale?

**Figure 1.**
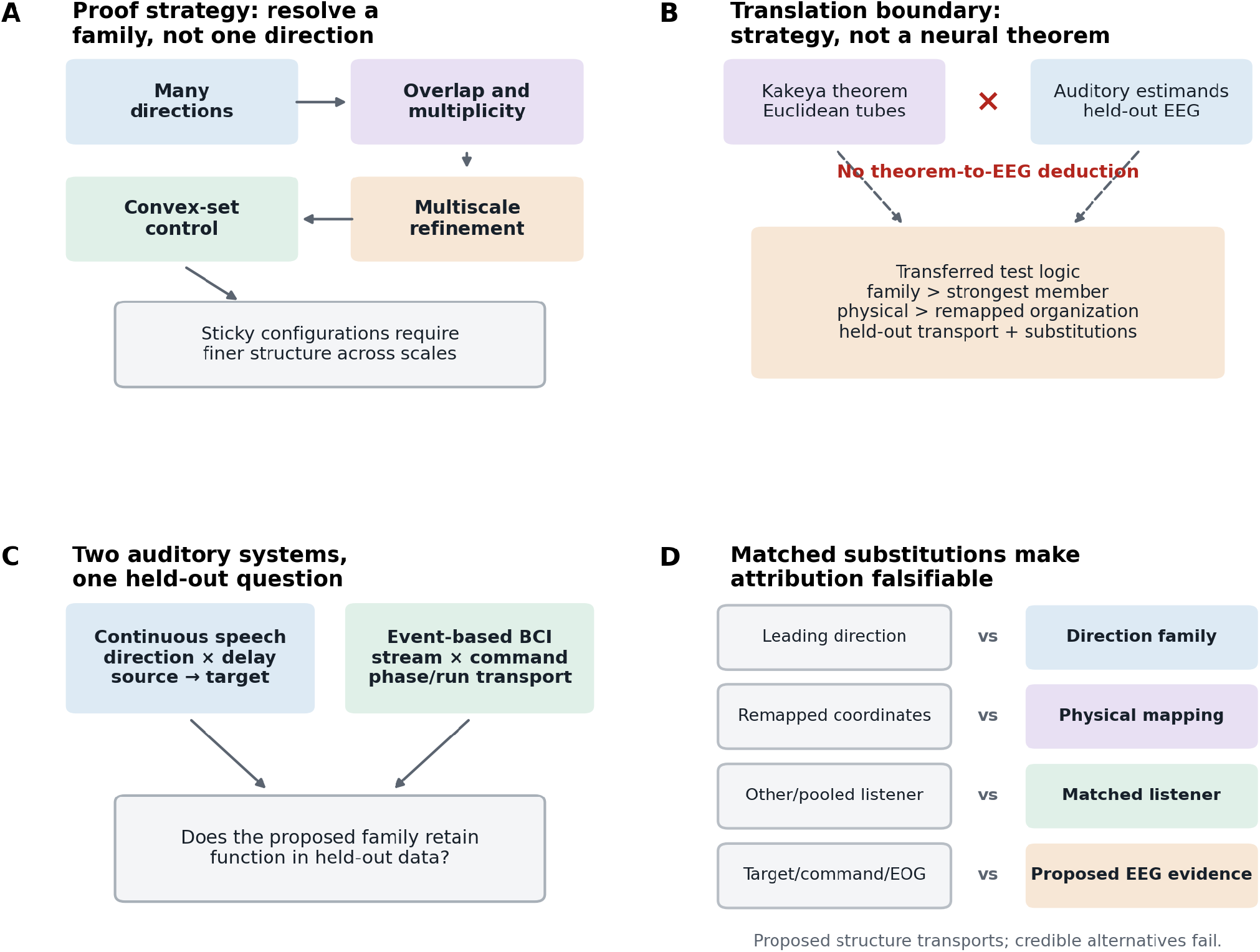
From Kakeya proof architecture to falsifiable attribution of auditory-attention evidence. (A) Wang–Zahl and Guth–Wang–Zahl control a family of thin tubes through non-clustering and multiplicity bounds, organization within convex regions, and decomposition across scales (11–13). (B) The theorem concerns Euclidean sets and does not imply a neural result; the transfer is a strategy for distinguishing organized structure from concentrated overlap. (C) Continuous-speech direction–delay cells and event-based parent–child directions instantiate one common held-out-function test. (D) The interpretation survives only if the proposed task structure transports while capacity-matched substitutions and shortcut explanations fail; the ocular comparison establishes that EEG evidence exceeds an EOG-only comparator, not that ocular activity has been removed from EEG.

These tests address known ambiguities in auditory decoding. Canonical correlation analysis (CCA), which finds paired linear projections of neural and speech signals with maximal correlation, may assign the largest training correlation to one component even when later components remain necessary to distinguish attended from unattended speech in new data (6). Spatial AAD can exploit gaze or other lateralized artifacts (14). Event-related interfaces can rely on the P300, a positive voltage deflection evoked by a rare target, while appearing to recover the task hierarchy. Cross-listener alignment can improve prediction by mapping people into a common space, but it can also discard individual structure (9, 15). Likewise, recurrent speech-tracking responses and intracranial speaker patterns show that neural patterns can reappear (16–17); they do not test the stronger counterfactual question of whether held-out evidence changes when the listener’s own weights are replaced while everything else is kept fixed.

We applied this logic to continuous speech and hierarchical auditory brain–computer interfaces (BCIs). For continuous speech, we compared ordered CCA projections and a fixed grid of eight scalp sectors crossed with early and late response delays. In the interfaces, commands (children) belonged to task-defined auditory streams (parents). A conventional **flat decoder** chose among all commands; a separate parent-stream score asked whether the remaining events supported the stream containing that choice. We then substituted listener weights, pooled templates, or physical mappings while keeping receiver data and model capacity fixed. Event analyses additionally removed target responses and command means and compared EEG with EOG-only models. Together these operations define a falsifiable attribution rule:

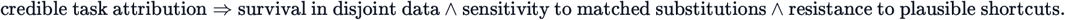

No single accuracy gain establishes this conjunction. The scientific claim lies in the predicted pattern: evidence survives a change of data, depends on the proposed organization, and weakens when a credible alternative source is substituted.

## Results

### Information beyond the leading direction survives held-out evaluations

We first asked whether the strongest neural–speech projection was an adequate summary of the family. Regularized CCA was fitted separately within each training fold, with all transformations estimated without the held-out stimulus. In each 5-s test window, projection pair *k* yielded a margin *M*_*k*_: its correlation with the attended speech minus its correlation with the unattended speech. The conventional readout used only *M*_1_, the largest training correlation. The breadth readout averaged *M*_1_, … , *M*_4_. Because both readouts used the same fitted CCA, they differed only in how the held-out projections were summarized.

The four-margin rule was selected in odd-numbered participants from the KU Leuven Auditory Attention Dataset (KUL; 18–19). It was then applied without retuning to the Technical University of Denmark dataset (DTU), the untouched even-numbered KUL participants, and Ear-SAAD, a dataset with simultaneous scalp, around-ear, and in-ear EEG. The three evaluation rules were fixed before their results were computed. Among the 41 participants, 32 improved (Fig. 2A). Giving each study equal weight, the accuracy gain was 0.0201 (stratified-bootstrap 95% interval, 0.0125–0.0279). The conservative replicability value—the largest of the three one-sided *P* values—was 0.0117. Weighting the four projections by their training canonical correlations also outperformed the leading projection in every study, but equal and canonical-correlation weighting changed order across studies. What replicated was the contribution of projections beyond the first, not a single optimal weighting rule.

**Figure 2.**
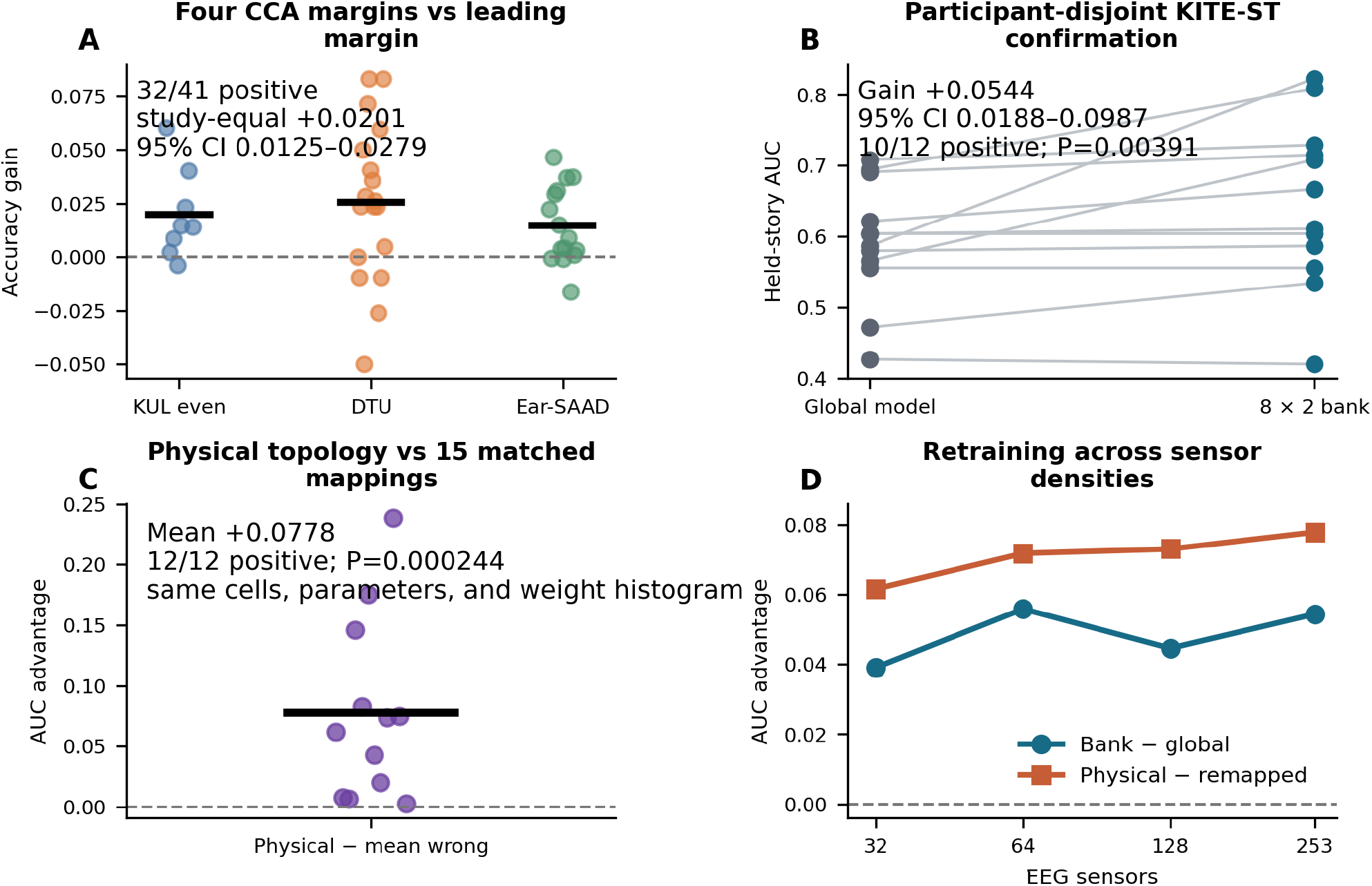
Continuous-speech information extends beyond the leading projection and depends on direction–delay structure. (A) Participant gains from using four canonical-correlation-analysis (CCA) margins instead of only the leading margin in three tests whose rules were fixed before their results were computed. Horizontal bars are study means. (B) Paired held-story areas under the receiver-operating-characteristic curve (AUCs) in the participant-disjoint KITE-ST confirmation. (C) Participant AUC advantage of the physical 16-cell topology over 15 capacity-matched mappings. (D) Independent retraining across sensor densities. A separate prediction that the complete cell-by-cell reliability pattern would correlate across stories was not supported; physical organization remained functional without a universally stable profile fingerprint.

We next asked whether the identities of those directions mattered. The public ultra-high-density EEG-AAD dataset (referred to here as KITE-ST) records two-talker continuous speech with a nominal 255-channel montage (20–21). A development cohort defined 16 cells: eight fixed scalp sectors crossed with early and late response delays. In a participant-disjoint complete-case cohort (*n* = 12), weighting these cells improved the held-story area under the receiver-operating-characteristic curve (AUC) by 0.0544 relative to a comparator that pooled across cells (95% interval, 0.0188–0.0987; 10/12 positive; exact one-sided *P* = 0.00391; Fig. 2B). AUC is the probability that a randomly chosen positive observation receives a higher score than a negative observation. The physical correspondence between training and test cells exceeded the mean of 15 prespecified remappings by 0.0778 AUC (12/12 positive; *P* = 0.000244; Fig. 2C). Each remapping preserved the same 16 cells, number of fitted quantities, and fold-specific weight distribution; it changed only which training weight was assigned to which test cell. Models retrained independently with 32, 64, 128, and 253 sensors retained positive bank and mapping advantages, although the prespecified gain-by-density slope was not significant (Fig. 2D). We also predicted that the complete cell-by-cell reliability pattern would correlate across stories; it did not. Thus physical organization remained functional without forming a universally stable profile fingerprint.

Thus, two forms of compression lost different information. Keeping only the leading CCA margin discarded reproducible discriminative signal; pooling or relabeling the 16 cells discarded their physical direction–delay organization. Neither result, on its own, showed that this organization was specific to the listener.

### Structure-dependent held-speech scores across nonoverlapping speech

To determine whether the 16-cell organization carried listener-specific function, we next transported it across nonoverlapping speech. Before calculating the four transport endpoints, we recorded one analysis specification for all three datasets: the partitions, transformations, endpoints, and decision criteria (SI Appendix, Table S1). KITE-ST (*n* = 12, 253 retained scalp channels), KUL (*n* = 16, 64 channels), and Ear-SAAD (*n* = 15, 29 scalp channels) were each divided into nonoverlapping source and target data (18, 22–24). The two partitions shared no EEG window, trial, story, or audio-part identity, as appropriate to the dataset. Source-only cross-validation produced a **reliability profile**: 16 dimensionless values describing how consistently each direction–delay cell contributed. These values were transformed into normalized positive weights. In a donor substitution, only those 16 values came from another listener; the receiver’s decoder coefficients, feature normalization, EEG, and target data were unchanged. Each participant contributed both A-to-B and B-to-A evaluations, yielding 43 participant endpoints and 86 directed source-to-target comparisons.

The four endpoints separated two questions: does the profile itself recur, and do source-derived weights order held-out speech evidence as predicted (Fig. 3A–D)? For each participant, a **rank advantage** was the probability that the matched value exceeded a substitute, with half credit for ties, minus 0.5; zero therefore denotes chance ordering. Across datasets, own-listener profiles exceeded donor profiles by 0.1676 (95% interval, 0.0879–0.2431; randomization *P* = 0.000324), and the physical profile correspondence exceeded the mean of the 15 prespecified remappings by 0.2586 (0.1820– 0.3334; *P* = 0.000006). This average requires qualification. KITE-ST alone was null for both static endpoints (listener mean, −0.0227; mapping mean, 0.0167; both intervals included zero). The data therefore do not support a stable profile fingerprint in every dataset.

**Figure 3.**
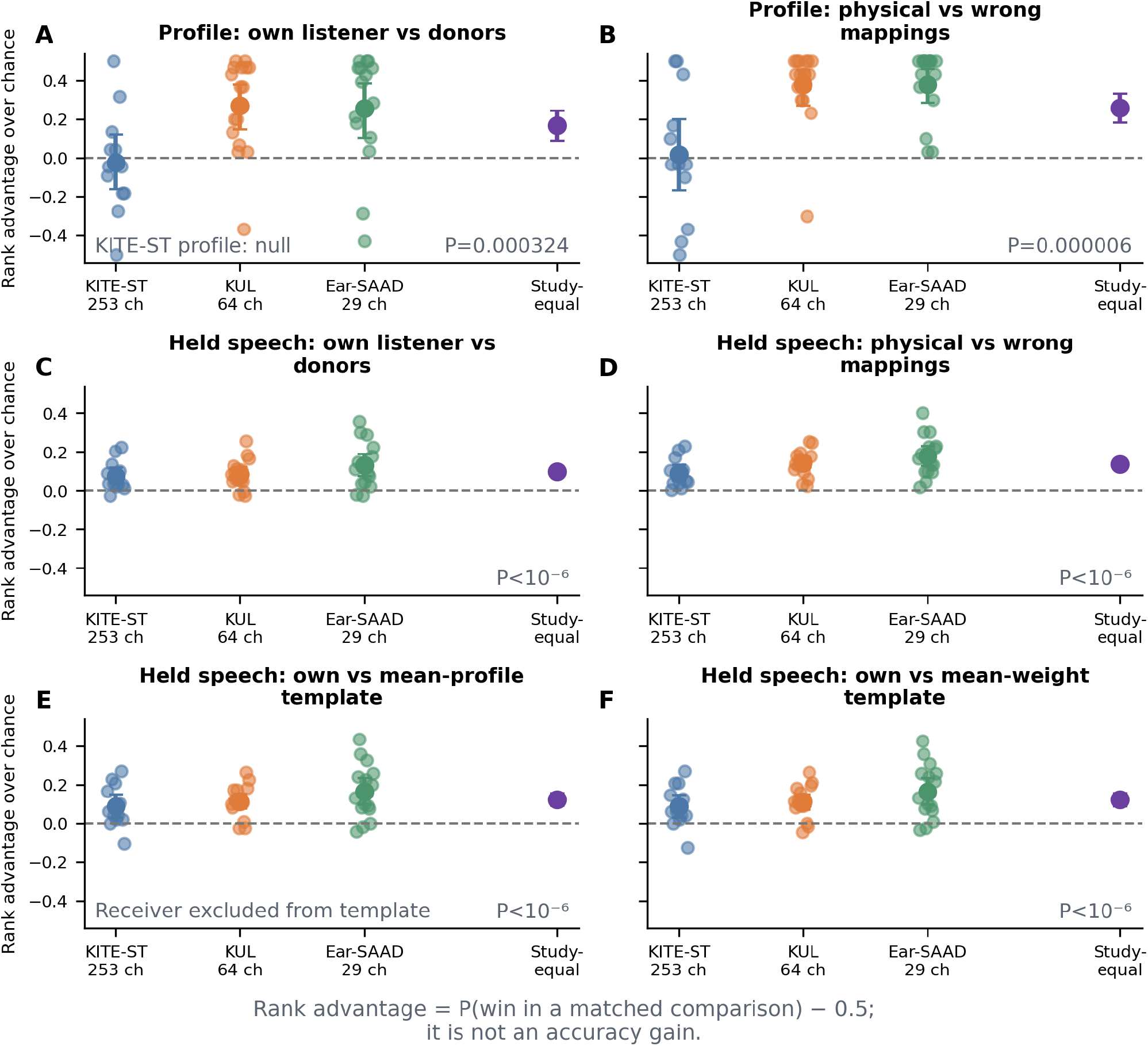
Structure-dependent held-speech evidence across nonoverlapping continuous speech. (A,B) Static profile rank advantages for own versus donor listeners and physical versus wrong mappings. KITE-ST is null for both static endpoints. (C,D) Held-speech score ranks are positive in all three datasets. (E,F) Receiver-matched weights exceed two leave-one-listener pooled templates. Points are participants, large circles are dataset means with participant-bootstrap 95% intervals, and purple points are study-equal means with stratified intervals. Rank advantage is winning probability minus 0.5; accuracy and calibration are separate endpoints.

Held-speech score ordering was more consistent. With the receiver model and target contributions fixed, the receiver’s source weights outranked every other listener’s weights by 0.0969 (0.0710–0.1241; *P* < 10^−6^); all three dataset means were positive. This value corresponds to a winning probability of approximately 0.597; classification accuracy is reported separately below. The physical direction–delay mapping outranked the mean of the 15 prespecified remappings by 0.1371 (0.1132–0.1620; *P* < 10^−6^). This effect was positive in all 43 participants and corresponds to a winning probability of approximately 0.637.

Averaging donors provides a less noisy population comparator. Two additional templates excluded the receiver: one averaged the other listeners’ reliability profiles before weight transformation, and the other averaged their transformed weights. Receiver-matched weights exceeded these templates by 0.1225 and 0.1219, respectively (both stratified intervals excluded zero; both *P* < 10^−6^; Fig. 3E,F). Final classification AUC, however, increased in KITE-ST, decreased slightly in KUL, and was nearly unchanged in Ear-SAAD. Listener matching therefore altered the ordering and scaling of held-speech evidence but did not improve classification accuracy universally; calibration and practical confidence were not tested.

The contrast between static and held-speech score endpoints is informative. A 16-cell profile can correlate poorly across stories when noisy or redundant cells dominate the correlation, even though the source-derived weights still emphasize the target cells that carry useful attention evidence. The reproducible quantity was therefore not always the complete profile itself, but the way source weights combined with held-out target contributions.

### Non-target sibling events detect parent-stream errors

Continuous speech has no discrete target event that can be removed, so we next turned to two public interfaces based on event-related potentials (ERPs), voltage responses time-locked to individual auditory events. Both use the Auditory Stream Segregation Multiclass ERP (ASME) paradigm. The four-command dataset of Kojima and Kanoh (hereafter Kojima) places two child commands in each of two streams. The ASME30 speller places 30 spoken commands in three pitch-defined streams corresponding to the three rows of the familiar QWERTY keyboard layout (25–27). The flat decoder selected one child command. A separate equal-prior linear discriminant was trained on **non-target siblings** from the attended parent stream (*S*) versus commands from unattended parents (*U*). For a selected child, the parent score was the mean *S*-versus-*U* evidence in its predicted parent minus the largest mean in a competing parent. This score asked whether the child prediction had crossed a stream boundary; it did not replace the child decoder (Fig. 4A).

**Figure 4.**
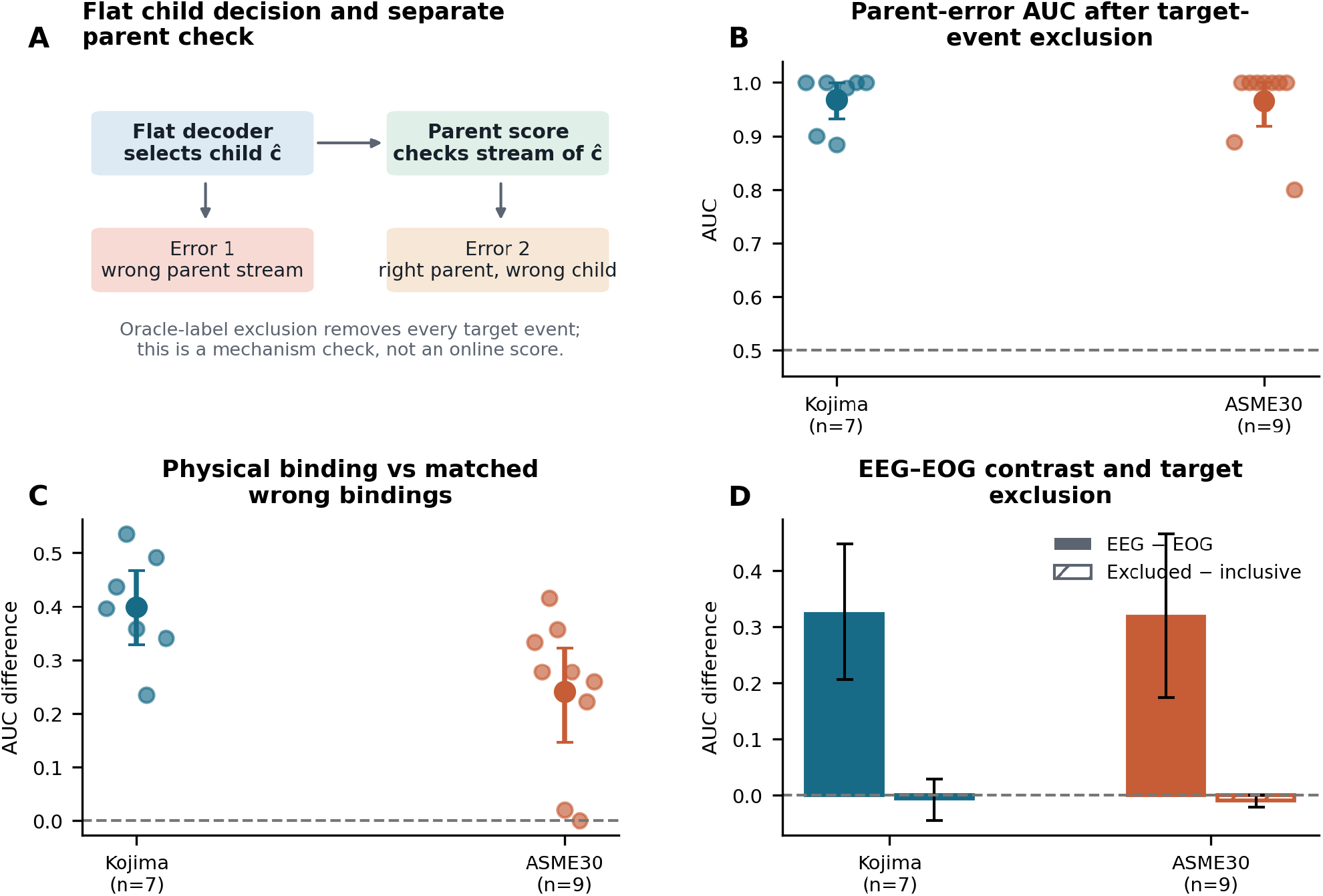
Non-target sibling events detect parent-stream errors hidden by a flat auditory brain–computer interface. (A) The flat decoder selects a child command; a separate sibling-versus-unattended (*S* − *U*) score evaluates its parent stream and distinguishes wrong-parent from within-parent errors. (B) Target-event-excluded parent-error AUC after deletion of every true-target-command event using the known label. (C) Physical parent binding versus capacity-matched wrong bindings. (D) EEG exceeds an EOG-only comparator, whereas target-event exclusion changes the inclusive AUC minimally. The EOG contrast does not establish absence of ocular contamination. Error bars are participant-bootstrap 95% intervals.

The critical analysis removed every event belonging to the true target command before parent evidence was pooled. The flat prediction, trained model, physical mapping, event horizon, and analysis windows were otherwise unchanged. Because removal requires the known target label, this **target-event-excluded** score is an oracle-label, mechanism-oriented exclusion check rather than an online confidence measure. It asks whether the parent score can be reduced to the target P300 under this specific deletion.

Target-event-excluded parent-error AUC was 0.9678 in the seven held-out Kojima participants (95% interval, 0.9324– 0.9986; 7/7 above 0.5; exact one-sided *P* = 0.00781) and 0.9654 in nine evaluable ASME30 participants (0.9185– 1.0000; 9/9; *P* = 0.00195; Fig. 4B). The published physical command–stream binding exceeded capacity-matched wrong parent assignments by 0.3996 AUC in Kojima and 0.2405 in ASME30 (Fig. 4C). EEG also exceeded an identically pooled EOG-only comparator by 0.3221 and 0.3179 AUC; this contrast does not prove that the EEG was free of ocular contamination. Removing target events changed the inclusive AUC by only −0.0052 and −0.0084 (Fig. 4D). Both datasets met the criteria, recorded before these values were computed, for absolute performance, physical-binding specificity, and an EEG–EOG contrast.

The two scores describe different errors. Flat confidence identifies the child with the largest accumulated response. Sibling evidence asks whether the rest of the auditory scene supports that child’s parent stream. The parent score remained informative without the most conspicuous target response and depended on the task’s actual stream–command assignment.

### Task-valid event directions transport within listeners

A high target-event-excluded AUC could still arise from a scalp pattern shared by all participants. We therefore tested whether each participant’s *S* − *U* sensor–time direction transported from offline to later online phases in ASME30 and across held-out runs in Kojima (Fig. 5). Target-command events were excluded from model fitting, testing, and direction estimation. For each participant and each physical or wrong mapping, Ledoit–Wolf linear discriminant analysis—a regularized linear classifier suitable for correlated features—estimated a direction from 64 channels × ten 100-ms bins. Test scores were centered and scaled with training data only. The physical effect was the held-out mean standardized *S* score minus the *U* score. Alignment was the cosine similarity between the training and held-out *S* − *U* vectors.

**Figure 5.**
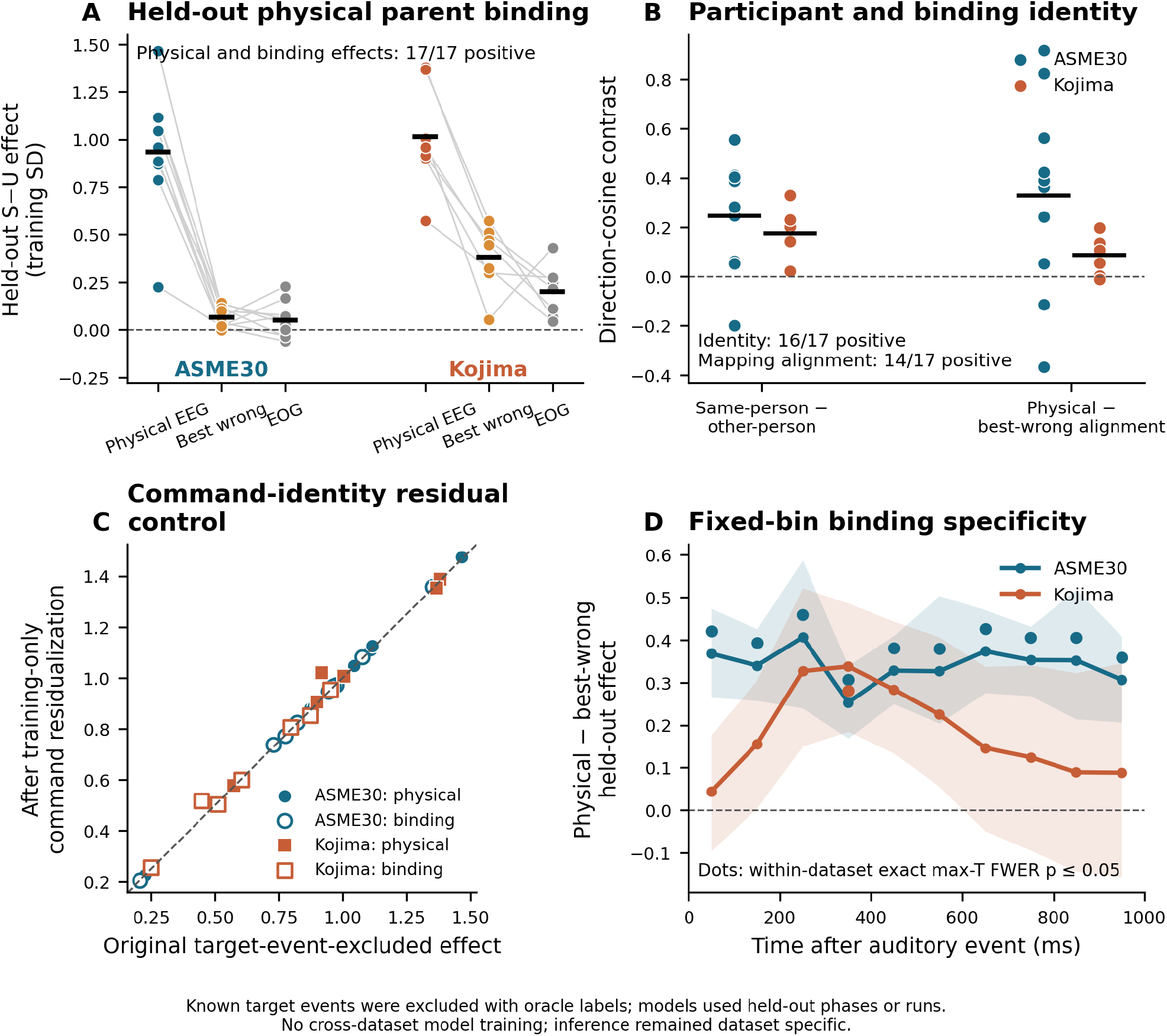
Task-defined event directions transport within listeners across phases or runs. (A) Held-out physical, wrong-binding, and EOG-only effects in ASME30 and Kojima. (B) Same-listener directions exceed other-listener substitutions, and physical alignment exceeds wrong-binding alignment. (C) Training-only command residualization preserves physical and binding effects. (D) For fixed-bin physical-minus-wrong effects, lines are participant means, shaded bands are participant-bootstrap 95% intervals, and dots mark bins significant under exact within-dataset maximum-*t* familywise control. The two datasets are not trained or tested across one another; inference remains dataset-specific.

Before computing these results, we had recorded seven criteria: a positive physical effect, specificity to the physical binding, an EEG effect greater than the EOG-only comparator, stronger alignment within than between listeners, stronger alignment for the physical than for a wrong mapping, a familywise-controlled temporal effect, and complete data coverage. Both datasets met all seven. ASME30 physical and binding effects were 0.9351 and 0.8711 training SD, respectively; the corresponding Kojima effects were 1.0141 and 0.6323. Same-listener alignment exceeded other-listener alignment by 0.2483 in ASME30 and 0.1762 in Kojima. The ten fixed 100-ms bins were tested together with an exact maximum-*t* procedure, which controls the familywise error rate across bins and avoids choosing a favorable latency after inspection.

Descriptively, the physical, binding, and EEG-minus-EOG effects were positive in 17/17 participants, same-listener transport in 16/17, and physical-versus-wrong mapping alignment in 14/17. Statistical inference remained separate for the two datasets because their tasks and train–test structures were not exchangeable.

A stable acoustic or command-identity pattern could also generate an *S* − *U* direction. In a separate control, each command mean was estimated from non-target events in the training partition and removed from both training and test features; held-out data and target events never contributed to that mean. All nine prespecified coverage, physical, binding, listener, and mapping criteria were met. The residualized physical/binding effects were 0.9394/0.8725 in ASME30 and 1.0288/0.6414 in Kojima, closely matching the original effects (Fig. 5C). A stable child-command mean therefore could not replace the listener-matched, task-bound direction.

### Physical binding survives wearable reduction, but listener specificity is less robust

The source studies used conventional 64-channel recordings and, for KITE-ST, an ultra-high-density montage with 253 retained scalp signals. Neither density nor a full calibration session is realistic for many practical BCIs. We therefore retrained every reduced montage and calibration subset rather than deleting sensors after fitting. With eight channels, both event datasets retained positive physical-binding effects that exceeded EOG. Binding specificity preserved 59.1% of the 64-channel ASME30 effect and 76.9% of the Kojima effect (Fig. 6A,C). Three training runs were sufficient to meet the prespecified physical and binding criteria in both datasets.

**Figure 6.**
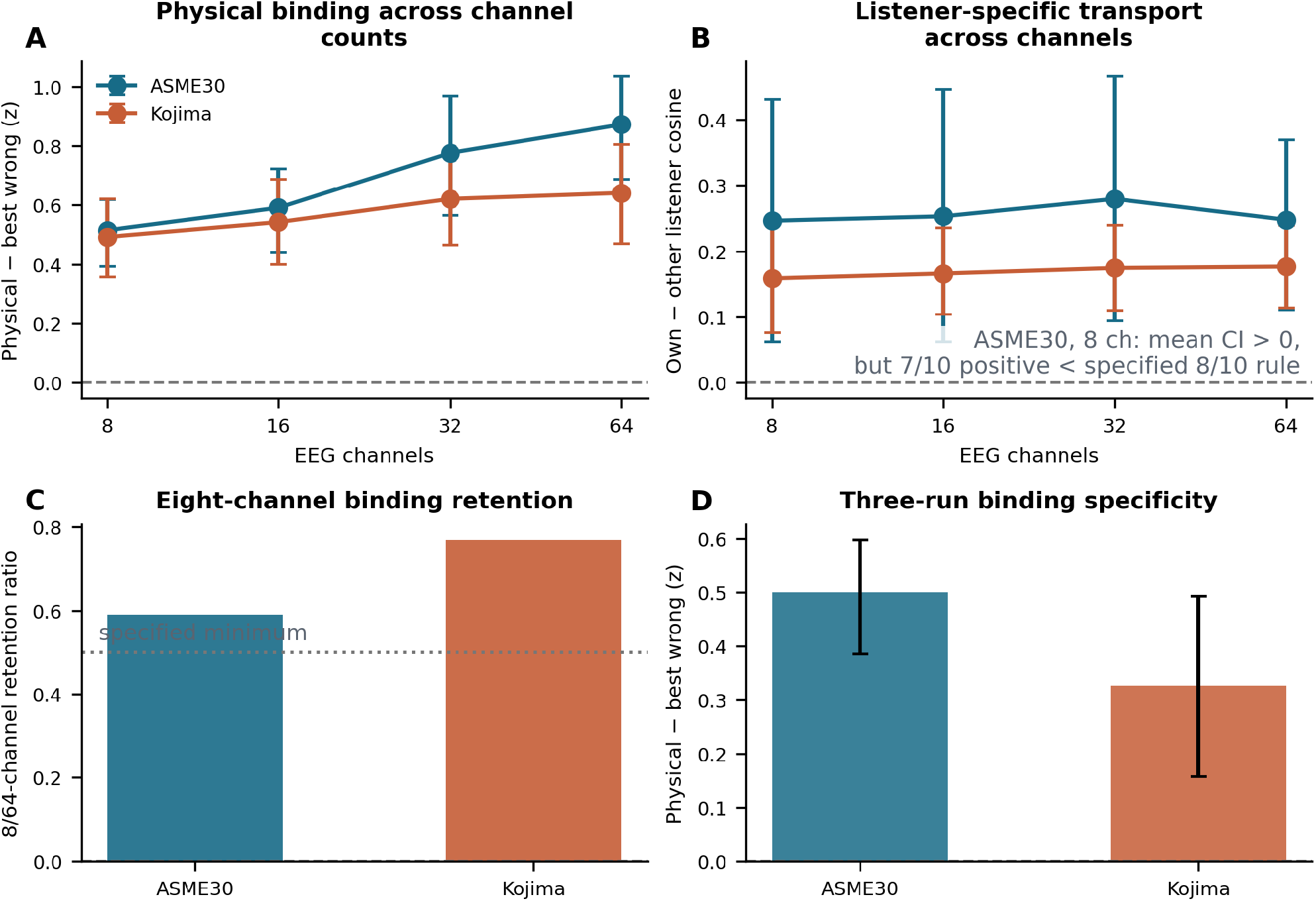
Physical binding survives wearable reduction, but listener specificity is less robust. (A) Physical binding across 8–64 channels. (B) Listener-specific transport; the eight-channel ASME30 mean remains positive, but only 7/10 participants show a positive effect, short of the criterion of 8/10 recorded before this reduction was evaluated. (C) Eight-channel binding retention, shown as a dimensionless proportion of the 64-channel effect. (D) Three-run binding specificity, shown in training-score SD units. Error bars in (A), (B), and (D) are participant-bootstrap 95% intervals. Every montage and calibration subset was retrained.

Listener specificity was less robust. The eight-channel ASME30 mean remained positive (*P* = 0.0195), but only 7/10 participants were positive, short of the prespecified 8/10 consistency criterion; Kojima met its criterion (Fig. 6B). The joint wearable/calibration test therefore missed one of eight requirements. Low-density EEG preserved the group-level physical parent binding more reliably than it preserved each listener’s spatial direction.

The present event datasets do not causally separate task hierarchy from acoustic organization: in ASME30, for example, parent streams are also pitch-defined. Training-only command residualization weakens a stable command-mean explanation but cannot remove every parent-level acoustic, adaptation, or sequence cue. A decisive prospective experiment would hold stimuli and low-level acoustics fixed while randomizing whether listeners must first identify the parent stream.

## Discussion

The experiments provide diagnostic tests for auditory decoders rather than a single attribution claim. The leading CCA component missed reproducible information. More importantly, held-out scores changed when we substituted another listener, pooled donor templates, or a wrong physical mapping. In the event datasets, the parent-stream score remained informative after target responses and stable command means were removed, and EEG outperformed an EOG-only model. These results rule out specific shortcuts that each dataset allowed us to construct. They do not show that attention is the only remaining cause.

Kakeya supplied the experimental discipline behind these tests. The Wang–Zahl proof and its streamlined formulation centrally use family-level volume and multiplicity bounds, non-clustering conditions, organization through convex sets, induction across scales, and reduction to the sticky case (11–13). We translated that architecture rather than any formula: family-level analysis replaced reliance on one extremal component; concentration motivated deletion tests; alternative organization motivated capacity-matched remappings; and multiscale reasoning motivated transport across held material and sensor density. The remappings are an empirical analogue of challenging a proposed factorization, not an operation deduced from the theorem. This translation generated experiments that accuracy optimization alone would not require— and produced informative failures when the full KITE-ST profile did not recur, the high-density bank did not improve every external dataset, and the eight-channel listener criterion failed in ASME30.

More broadly, the framework addresses a recurring gap in cognitive neuroscience: a classifier may predict an experimental label without identifying the psychological construct used to interpret that label. This gap is especially visible for latent concepts such as attention and consciousness, where multiple processes can support the same report or neural contrast. Recent adversarial tests of consciousness theories have advanced the field by specifying divergent predictions and conditions under which each theory would be challenged (28). The present study does not test a theory of consciousness. It contributes a compatible methodological standard for decoding studies: competing explanations should be converted into matched substitutions with different predicted failures. Such tests can turn rapid gains in prediction into cumulative evidence about what a decoded construct requires.

The findings address three persistent difficulties in AAD. First, neural responses change with the speech material, so a static template can appear unreliable. KITE-ST illustrates why a held-material ordering test can be more informative: the complete profile did not recur across stories, but listener-matched weights still changed the ordering of held-story evidence. Second, cross-listener variation is commonly treated as noise to remove. Population alignment and listener-independent networks remain valuable (9, 15), but donor and pooled-template comparisons show that some individual structure changes held-out scores. A future model could combine a shared physical prior with a small, calibrated listener residual; the present study did not test calibration benefit. Third, ordinary decoder margins do not identify the kind of error. The parent score distinguished “wrong stream” from “right stream, wrong command” using sibling events discarded by a winner-take-all decoder.

These results complement recent reports of reliable speech tracking across natural stimuli (16), stable intracranial patterns for attended-speaker identity (17), and increasingly accurate spatiotemporal networks (9). The distinction is experimental: we combined nonoverlapping material, a common physical coordinate, substitutions by individual and pooled donors, 15 capacity-matched mappings, separate static and held-score endpoints, target-event exclusion, command residualization, and cross-phase or cross-run transport. The individualized object was the listener’s weighting of task directions, not the identity of the attended speaker.

The same substitution logic can be used outside EEG, although the present empirical conclusions cannot simply be transferred. In magnetoencephalography (MEG), functional magnetic resonance imaging (fMRI), intracranial electrophysiology, or another BCI modality, the indexed family might consist of sensors or sources, voxels or networks, time–frequency components, or levels of a task hierarchy. A valid extension would need its own physically or experimentally defined coordinate, disjoint evaluation data, equal-capacity substitutions, and modality-specific artifact controls.

For brain-controlled hearing, these tests suggest a layered controller. A low-density parent score could ask whether a predicted command or talker remains in a supported stream, while a small listener-specific calibration refines the score. Wrong mappings would serve as internal references: if they perform as well as the physical mapping, the device should withhold its decision. A closed-loop trial must determine whether this improves calibration, abstention, speech benefit, or safety.

Embodied listening adds an action to the same problem. A head turn or microphone movement changes source separation and can create a better observation. A future system could act when the physical mapping clearly beats its remapped controls and sample again after an informative movement when it does not. The present data contain no robot or active head movement. Public attention-switching EEG and recent cortical measurements make such transition experiments feasible (29–30), while generative augmentation may help characterize short-window uncertainty (31). The decisive comparison is prospective: ordinary confidence versus structure-sensitive controls in a closed-loop task.

The study has important limits. All analyses used public data, and some mechanism and robustness questions were specified after related results in the same datasets were known; the evidence chronology is reported explicitly in SI Appendix. The study was not preregistered. The local protocol files and hashes that preserve the internal analysis chronology are included in the source-code archive supplied with this submission. Continuous speech and event-based interfaces support a common testing logic but need not share a neural mechanism. Rank advantage and classification accuracy are different endpoints, and listener matching did not improve classification AUC in every dataset. EOG-only comparisons were available only for Kojima and ASME30; they reduce but do not eliminate ocular explanations, and the continuous-speech analyses did not include an equivalent ocular substitution. The KUL release has a documented gaze bias, which further limits spatial interpretations. Target-event exclusion uses the known target label and cannot run online in its present form. The event datasets do not causally separate task hierarchy from acoustic stream organization. Sensor-space directions are not cortical sources. Participants were healthy, and the study measured neither calibration benefit, clinical benefit, nor closed-loop behavior. Finally, the low-density listener criterion failed in ASME30.

A leading projection, winning child label, and population template each left useful information behind. The remaining held-out scores depended on the physical task mapping and, in part, on listener-specific weights. The primary contribution is therefore not six EEG analyses in isolation, but a transferable standard for construct attribution: preserve candidate evidence in disjoint data, replace its proposed source at matched capacity, and measure which interpretation breaks.

## Materials and Methods

### Study design and evidence classes

All analyses used deidentified public datasets collected under the original studies’ ethics approvals. We distinguished analyses used to select a method from tests whose rules were recorded before their endpoint was computed, participant-disjoint within-site confirmation, mechanism analyses proposed after a related primary result, and post-outcome robustness analyses. For each new endpoint, protocol files recorded participants, partitions, mappings, statistical criteria, seeds, and stopping rules before evaluation. Separate scripts audited source-level and aggregate results. SI Appendix reports the chronology, implementation deviations, negative analyses, and data licenses.

### Projection-breadth experiment

KUL, DTU, and Ear-SAAD preprocessing and cross-validation followed the specifications in SI Appendix. EEG and speech envelopes were band limited and resampled. Principal-component, lag, covariance, and CCA transformations were estimated from the training data in each outer fold, where “outer” denotes the partition held aside for final evaluation. All samples derived from the same story identity remained in the same fold. For canonical pair *k*,

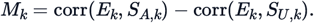

The baseline classified by *M*_1_; the breadth readout used *B*_4_ = (*M*_1_ + *M*_2_ + *M*_3_ + *M*_4_)/4. Ear-SAAD modalities were averaged within participant before inference. Bootstrap intervals and exact sign-flip tests treated the participant as the unit of analysis. The secondary synthesis resampled participants within each study and then averaged the study means equally.

### Continuous direction–delay bank and transport

The common coordinate comprised eight fixed scalp sectors crossed with early and late response delays. Source-only cross-validation estimated a 16-cell reliability profile: the aligned mean held-out contribution of each cell divided by its root-mean-square, bounded in [−1, 1]. Exponentiating twice each reliability and normalizing the resulting values produced positive weights with sum 16. The physical mapping used zero angular shift and preserved the delay class. The 15 alternatives comprised the remaining circular shifts, with or without exchanging early and late delays.

KITE-ST partitions were its two stories. KUL used balanced, nonoverlapping base-part groups 1, 4 and 2, 3. Ear-SAAD used a deterministic, balanced 3-of-6 trial split. Source profiles and the receiver’s ridge-regression model used source data only. Target data supplied held-out contributions and labels for the fixed score. A donor supplied only the 16 dimensionless profile values.

For static endpoints, the source-to-target correlation of the receiver’s own profiles was ranked against donor profiles (F1) or the 15 remapped own profiles (F2). For held-speech score endpoints, attended-side target scores obtained with the receiver’s own weights were ranked against donor weights (P1) or remapped weights (P2). Ties contributed 0.5. Participant rank advantage was the mean winning probability minus 0.5, averaged over both transport directions. Leave-one-listener templates either averaged donor profiles before the weight transformation or averaged donor weights after transformation.

Cross-dataset summaries weighted the three dataset means equally. Confidence intervals used 100,000 participant resamples stratified by dataset. One million fixed-seed sign flips were applied to the study-equal statistic; an add-one correction prevented a zero Monte Carlo *P* value. Support for transport required all four F1–P2 criteria.

### Hierarchical parent-error score

Kojima used seven two-stream participants with leave-one-complete-run-out validation. ASME30 used offline runs for training and later online runs for testing. Nine of ten participants had both parent-correct and parent-error trials and were evaluable for AUC; participant 6 had 15/15 parent-correct predictions and no parent-error trial, making AUC undefined rather than missing. This evaluability rule was determined by the definition of AUC, not by the observed parent-score values. For each physical or wrong mapping, *T* denoted the target command and was excluded, *S* the non-target siblings in the target’s parent, and *U* commands in other parents. Equal-prior Ledoit–Wolf linear discriminant analysis fitted *S* versus *U* separately to EEG and EOG.

The flat prediction averaged the first three accepted event scores for each command and selected the maximum. The target-event-excluded analysis then deleted all three events belonging to the known target before parent aggregation. The signed parent score was the mean outer-fold *S* − *U* score in the predicted parent minus the largest mean in a competing parent. The published auditory-stream/QWERTY mapping defined physical correctness. Participant AUC distinguished parent-correct from parent-error flat predictions. Inference used 20,000 participant-bootstrap samples and exhaustive one-sided sign flips. After the present reporting audit, an explicitly exploratory all-participant endpoint averaged each participant’s target-event-excluded physical score across all 15 trials and subtracted the better of two identically averaged wrong-mapping scores; it includes participant 6 and is reported in SI Appendix without upgrading the AUC claim.

### Cross-phase and cross-run direction geometry

ASME30 EEG was represented by 64 channels × ten fixed 100-ms bins from 0 to 1 s; offline data trained the model and later online data tested it. Kojima used the same feature dimension with leave-one-run-out validation. Separate *S* − *U* models were trained for the physical and each wrong mapping. Training scores determined centering and scaling. The physical held-out effect was the trial-balanced mean standardized *S* score minus the *U* score. Binding specificity subtracted the larger effect from the two wrong mappings, each fitted separately; the EEG–EOG contrast subtracted the EOG-only effect.

Within each participant, pooled training *S*/*U* data standardized every feature. The training direction was mean(*S*) −mean(*U*); the held-out direction was its trial-balanced analogue. Listener-specific transport was the cosine similarity of matched training and held-out directions minus the mean similarity obtained after substituting other participants’ training directions. Mapping specificity subtracted the larger within-person cosine from the two wrong mappings. Exhaustive sign flips and a two-sided maximum-*t* statistic controlled familywise error across the ten time bins.

For command residualization, each command mean was estimated independently within the training partition from non-target events only. That mean was removed from the training and held-out features before the geometry analysis was repeated. No held-out observation or target event contributed to a residual mean.

### Sensor and calibration reductions

Montages of 8, 16, 32, and 64 electrodes were selected from sensor geometry without reference to outcomes. Training-run subsets were enumerated within participant, and repeated subsets of the same size were averaged before group inference. The wearable analysis jointly required complete coverage, eight-channel physical and binding effects, an EEG effect greater than its EOG-only comparator, a listener effect, minimum binding retention, and three-run physical and binding effects.

### Statistical reporting and auditability

Participants were the inferential unit throughout. Unless stated otherwise, scalar effects are reported with participant percentile-bootstrap 95% intervals and exhaustive exact one-sided sign-flip tests. Participant counts across nonexchangeable task structures are descriptive. Text values are rounded from machine-readable outputs. A claim ledger records the source JSON path and SHA-256 digest for 33 ledger-indexed quantitative claims. A separately implemented source audit reconstructed the continuous endpoints with maximum participant and direction errors of 1.11 × 10^−16^ and 2.22 × 10^−16^, respectively; aggregate recomputation error was zero. Separate audits covered the event, command-residual, and wearable analyses.

### Use of generative AI

Y.D. formulated the research questions and experimental design, selected and checked the literature and public datasets, designed the analysis logic and code architecture, wrote the initial manuscript, interpreted the results, and approved the final text and files. OpenAI Codex assisted interactively with code editing and refactoring, prose editing, organization of references and project materials, and technical explanations. Y.D. checked cited sources against publisher or repository records, reviewed the code changes, quantitative outputs, and independent audits, and accepts responsibility for the work. Codex is not an author. No generative-AI image system was used; scientific figures were produced by deterministic plotting code from reported results and were reviewed by Y.D.

## Supporting information

Article_Analysis_Source_Code

Derived_Result_Verification_Archive

## Acknowledgments

We thank the investigators who made the KUL, DTU, Ear-SAAD, ultra-high-density EEG-AAD, Kojima ASME, and ASME30 datasets publicly available. This study involved no new participant recruitment or data collection.

## Author Contributions

Y.D. was responsible for conceptualization, methodology, literature and data curation, software design, formal analysis, validation, writing the original draft, revision, and final approval.

## Data, Materials, and Software Availability

The six source datasets used for the primary analyses are publicly available under their original licenses: KUL (19), DTU (32), ultra-high-density EEG-AAD (21), Ear-SAAD (24), Kojima ASME (33), and ASME30 (34). Raw EEG, audio, and third-party publications are not redistributed. Two versioned archives are supplied with this submission: `Article_Analysis_Source_Code.zip` contains the preprocessing, feature-extraction, model-fitting, statistical, source-audit, and figure programs needed to rerun the reported workflows after the source datasets are downloaded; `Derived_Result_Verification_Archive.zip` checks 33 ledger-indexed quantitative claims against their referenced result files, recomputes participant-level summaries supported by deidentified tables, and renders all six figures. Each archive includes a README, environment and license records, and a member-level SHA-256 manifest. The archives are being prepared for permanent public deposition. Original code and documentation are licensed separately; derived data remain subject to the applicable source-dataset terms.

