## Supplementary material for "Falsifiable substitution tests reveal task-structured neural evidence for auditory attention": Article_Analysis_Source_Code: audit_dtu_aad.py

```
"""Outcome-free integrity and identity audit for the DTU AAD release."""

from __future__ import annotations

import argparse
import hashlib
import json
import re
from collections import Counter
from pathlib import Path, PurePath

import numpy as np
from scipy.io import loadmat, wavfile


SUBJECTS = tuple(range(1, 19))
EXPECTED_MD5 = {
    "DATA_preproc.zip": "1c1ca1390ad99ff41c33f36ca67f8418",
    "AUDIO.zip": "18532a2882ccd87c448c9481b4b2c47d",
    "preproc_data.m": "7723ddc5334f5aa1f473143251fdcffd",
}


def md5(path: Path) -> str:
    digest = hashlib.md5()
    with path.open("rb") as handle:
        while block := handle.read(8 * 1024 * 1024):
            digest.update(block)
    return digest.hexdigest()


def basename(value: object) -> str:
    return PurePath(str(value).replace("\\", "/")).name


def story_identity(filename: str) -> str:
    match = re.fullmatch(
        r"(aske|marianne)_story(\d+)_trial_\d+\.wav", filename
    )
    if not match:
        raise ValueError(f"Unexpected DTU audio filename: {filename}")
    return f"{match.group(1)}_story{match.group(2)}"


def trial_audio_paths(data: object) -> list[tuple[str, str]]:
    cfg = np.atleast_1d(data.cfg)
    if len(cfg) != 19:
        raise RuntimeError("Unexpected DTU preprocessing history length")

    def extract(trial_cfg: object) -> tuple[str, str]:
        values = np.atleast_1d(trial_cfg)
        return (
            basename(values[10].wavA.dataset),
            basename(values[12].wavB.dataset),
        )

    output = [extract(cfg)]
    output.extend(extract(value) for value in np.atleast_1d(cfg[18].datacfg))
    return output


def main() -> None:
    parser = argparse.ArgumentParser()
    parser.add_argument(
        "--data-root", type=Path, default=Path("data/raw/dtu_aad")
    )
    parser.add_argument(
        "--output",
        type=Path,
        default=Path("results/dtu_aad/integrity_audit.json"),
    )
    args = parser.parse_args()
    preprocessed_root = args.data_root / "preprocessed_extracted"
    audio_root = args.data_root / "audio_extracted"

    hashes = {}
    for name, expected in EXPECTED_MD5.items():
        path = args.data_root / name
        observed = md5(path)
        hashes[name] = {
            "expected_md5": expected,
            "observed_md5": observed,
            "matches": observed == expected,
            "bytes": path.stat().st_size,
        }

    audio_metadata = {}
    for path in sorted(audio_root.glob("*.wav")):
        sample_rate, audio = wavfile.read(path, mmap=True)
        audio_metadata[path.name] = {
            "sample_rate_hz": int(sample_rate),
            "samples": int(audio.shape[0]),
            "duration_seconds": float(audio.shape[0] / sample_rate),
            "dtype": str(audio.dtype),
            "channels": 1 if audio.ndim == 1 else int(audio.shape[1]),
        }

    rows = []
    global_valid = all(value["matches"] for value in hashes.values())
    common_scalp_labels = None
    for subject in SUBJECTS:
        path = preprocessed_root / f"S{subject}_data_preproc.mat"
        data = loadmat(
            path,
            squeeze_me=True,
            struct_as_record=False,
            variable_names=["data"],
        )["data"]
        eeg_trials = list(np.atleast_1d(data.eeg))
        attended_envelopes = list(np.atleast_1d(data.wavA))
        unattended_envelopes = list(np.atleast_1d(data.wavB))
        events = list(np.atleast_1d(data.event.eeg))
        paths = trial_audio_paths(data)
        labels_by_trial = list(np.atleast_1d(data.dim.chan.eeg))
        if not all(
            len(value) == 60
            for value in (
                eeg_trials,
                attended_envelopes,
                unattended_envelopes,
                events,
                paths,
                labels_by_trial,
            )
        ):
            raise RuntimeError(f"S{subject}: expected 60 aligned trials")
        for trial in range(60):
            eeg = np.asarray(eeg_trials[trial])
            wav_a = np.asarray(attended_envelopes[trial])
            wav_b = np.asarray(unattended_envelopes[trial])
            labels = [str(value) for value in np.atleast_1d(labels_by_trial[trial])]
            scalp_labels = labels[:64]
            if common_scalp_labels is None:
                common_scalp_labels = scalp_labels
            attended_file, unattended_file = paths[trial]
            pair = sorted(
                [
                    story_identity(attended_file),
                    story_identity(unattended_file),
                ]
            )
            valid = bool(
                eeg.shape == (3200, 66)
                and wav_a.shape == (3200,)
                and wav_b.shape == (3200,)
                and np.isfinite(eeg).all()
                and np.isfinite(wav_a).all()
                and np.isfinite(wav_b).all()
                and int(data.fsample.eeg) == 64
                and int(data.fsample.wavA) == 64
                and int(data.fsample.wavB) == 64
                and labels[-2:] == ["EXG1", "EXG2"]
                and scalp_labels == common_scalp_labels
                and attended_file in audio_metadata
                and unattended_file in audio_metadata
                and audio_metadata[attended_file]["duration_seconds"] == 50
                and audio_metadata[unattended_file]["duration_seconds"] == 50
                and int(events[trial].value) in (1, 2)
            )
            global_valid &= valid
            rows.append(
                {
                    "subject": subject,
                    "trial": trial + 1,
                    "attended_audio": attended_file,
                    "unattended_audio": unattended_file,
                    "attended_story_identity": story_identity(attended_file),
                    "unattended_story_identity": story_identity(
                        unattended_file
                    ),
                    "unordered_story_pair_group": "|".join(pair),
                    "attention_code": int(events[trial].value),
                    "eeg_samples": int(eeg.shape[0]),
                    "released_channels": int(eeg.shape[1]),
                    "scalp_channels_used": 64,
                    "valid": valid,
                }
            )
        del data

    subject_pair_counts = {
        str(subject): dict(
            sorted(
                Counter(
                    row["unordered_story_pair_group"]
                    for row in rows
                    if row["subject"] == subject
                ).items()
            )
        )
        for subject in SUBJECTS
    }
    pair_sets = {
        tuple(value.keys()) for value in subject_pair_counts.values()
    }
    global_valid &= len(pair_sets) == 1
    report = {
        "dataset": "DTU auditory attention dataset",
        "zenodo_record": "https://zenodo.org/records/1199011",
        "participants": list(SUBJECTS),
        "file_hashes": hashes,
        "audio_file_count": len(audio_metadata),
        "scalp_channel_labels": common_scalp_labels,
        "excluded_released_channels": ["EXG1", "EXG2"],
        "summary": {
            "participant_count": len(SUBJECTS),
            "trial_count": len(rows),
            "trials_per_participant": 60,
            "samples_per_trial": 3200,
            "seconds_per_trial": 50,
            "sample_rate_hz": 64,
            "story_pair_groups_per_participant": len(next(iter(pair_sets))),
            "all_hashes_match": all(
                value["matches"] for value in hashes.values()
            ),
            "all_trials_valid": all(row["valid"] for row in rows),
            "story_pair_group_sets_match_across_participants": (
                len(pair_sets) == 1
            ),
            "global_integrity_pass": bool(global_valid),
        },
        "strict_identity_firewall": {
            "outer_test_unit": "unordered attended/unattended story pair",
            "training_exclusion": (
                "Exclude every trial containing either story identity from "
                "the held-out pair, regardless of attention role."
            ),
        },
        "pair_counts_by_subject": subject_pair_counts,
        "trials": rows,
    }
    args.output.parent.mkdir(parents=True, exist_ok=True)
    args.output.write_text(json.dumps(report, indent=2), encoding="utf-8")
    print(json.dumps(report["summary"], indent=2))
    print(json.dumps(subject_pair_counts["1"], indent=2))
    if not global_valid:
        raise RuntimeError("DTU integrity audit failed")


if __name__ == "__main__":
    main()
```
