## Supplementary material for "Falsifiable substitution tests reveal task-structured neural evidence for auditory attention": Article_Analysis_Source_Code: audit_kite_sat_akb_confirmation_complete_case_mechanism_precision_v2.py

```
"""Meta-audit the sole 1e-15 broadness mismatch in the v2 mechanism audit."""

from __future__ import annotations

import json
from pathlib import Path

import numpy as np

import audit_kite_sat_akb_mechanism_outcome_v1 as formulas
from kite_sat_confirmation_complete_case_lock_utils_v2 import (
    ROOT,
    sha256,
    validate_complete_case_lock,
)


LOCKED_CALLER = (
    ROOT / "analysis"
    / "audit_kite_sat_akb_confirmation_complete_case_mechanism_v2.py"
)
ADDENDUM = (
    ROOT / "docs"
    / "kite_sat_akb_confirmation_complete_case_mechanism_audit_precision_addendum_v2.json"
)
MECHANISM_ROOT = (
    ROOT / "results" / "kite_sat"
    / "akb_confirmation_complete_case_direction_delay_v2"
)
DECISION = MECHANISM_ROOT / "mechanism_decision_v1.json"
FAILED_AUDIT = MECHANISM_ROOT / "mechanism_outcome_audit_v1.json"
OUTPUT = MECHANISM_ROOT / "mechanism_outcome_audit_precision_v2.json"
SUBJECTS = (
    "S0", "S10", "S2", "S21", "S28", "S22",
    "S9", "S15", "S16", "S18", "S13", "S1",
)
TOLERANCE = 1e-14


def validate_addendum() -> dict:
    fields = ADDENDUM.with_name(ADDENDUM.name + ".sha256").read_text(
        encoding="ascii"
    ).strip().split()
    observed = sha256(ADDENDUM)
    payload = json.loads(ADDENDUM.read_text(encoding="utf-8"))
    relative = Path(__file__).resolve().relative_to(ROOT.resolve()).as_posix()
    if (
        len(fields) != 2
        or fields[0].lower() != observed
        or fields[1] != ADDENDUM.name
        or payload["status"] != "frozen_after_first_serialized_audit_failure"
        or payload["recovery_source"]["path"] != relative
        or payload["recovery_source"]["sha256"] != sha256(Path(__file__))
        or payload["failed_audit_sha256"] != sha256(FAILED_AUDIT)
        or payload["mechanism_decision_sha256"] != sha256(DECISION)
        or payload["descriptive_absolute_tolerance"] != TOLERANCE
    ):
        raise RuntimeError("Mechanism precision-audit addendum mismatch.")
    return payload


def main() -> None:
    if OUTPUT.exists():
        raise RuntimeError("Refusing to overwrite mechanism precision audit.")
    validate_addendum()
    lock, lock_hash = validate_complete_case_lock(LOCKED_CALLER)
    decision = json.loads(DECISION.read_text(encoding="utf-8"))
    failed = json.loads(FAILED_AUDIT.read_text(encoding="utf-8"))
    false_checks = sorted(
        key for key, value in failed["checks"].items() if not bool(value)
    )
    non_broadness_failures = [
        key for key in false_checks if not key.startswith("broadness:")
    ]
    differences = []
    records = []
    subject_hashes = {}
    for subject in SUBJECTS:
        path = MECHANISM_ROOT / "subjects" / f"{subject}_direction_delay_mechanism_v1.json"
        record = json.loads(path.read_text(encoding="utf-8"))
        subject_hashes[subject] = sha256(path)
        if subject_hashes[subject] != decision["subject_artifact_sha256"][subject]:
            raise RuntimeError(f"Mechanism subject hash mismatch: {subject}")
        for story in ("train_story_1", "train_story_2"):
            recomputed = formulas.broadness(
                np.asarray(record["reliability_vectors"][story], dtype=np.float64)
            )
            for metric, value in recomputed.items():
                saved = float(record["broadness"][story][metric])
                differences.append(
                    {
                        "subject": subject,
                        "story": story,
                        "metric": metric,
                        "absolute_difference": abs(float(value) - saved),
                    }
                )
        records.append(record)
    decoding = np.asarray(
        [record["intact_identity_advantage"] for record in records],
        dtype=np.float64,
    )
    reliability = np.asarray(
        [record["reliability_identity_advantage"] for record in records],
        dtype=np.float64,
    )
    decoding_p = formulas.exact_upper(decoding)
    reliability_p = formulas.exact_upper(reliability)
    maximum_difference = max(row["absolute_difference"] for row in differences)
    checks = {
        "complete_case_lock": (
            decision["mechanism_implementation_lock_sha256"] == lock_hash
        ),
        "decision_sidecar": (
            DECISION.with_name(DECISION.name + ".sha256")
            .read_text(encoding="ascii")
            .strip()
            .split()[0]
            .lower()
            == sha256(DECISION)
        ),
        "first_audit_status_is_fail": failed["status"] == "fail",
        "first_audit_only_failed_broadness_checks": not non_broadness_failures,
        "first_audit_has_at_least_one_broadness_failure": bool(false_checks),
        "all_non_broadness_checks_passed": all(
            bool(value)
            for key, value in failed["checks"].items()
            if not key.startswith("broadness:")
        ),
        "maximum_broadness_difference_lte_1e_14": (
            maximum_difference <= TOLERANCE
        ),
        "decoding_p_recomputed": (
            abs(
                decoding_p
                - decision["fixed_sequence"]["decoding_identity_advantage"][
                    "exact_one_sided_p"
                ]
            )
            < 1e-15
        ),
        "reliability_p_recomputed": (
            abs(
                reliability_p
                - decision["fixed_sequence"]["reliability_identity_advantage"][
                    "raw_exact_one_sided_p"
                ]
            )
            < 1e-15
        ),
        "mechanism_status_reproduced": (
            decision["status"] == "direction_delay_decoding_mechanism_supported"
            and decoding_p <= 0.05
            and reliability_p > 0.05
        ),
        "subject_hashes": subject_hashes == decision["subject_artifact_sha256"],
    }
    status = "pass" if all(checks.values()) else "fail"
    payload = {
        "schema": "KITE-SAT-AKB-complete-case-mechanism-precision-audit-v2",
        "status": status,
        "mechanism_status": "direction_delay_decoding_mechanism_supported",
        "complete_case_implementation_lock_sha256": lock_hash,
        "mechanism_decision_sha256": sha256(DECISION),
        "failed_strict_audit": {
            "path": FAILED_AUDIT.relative_to(ROOT).as_posix(),
            "sha256": sha256(FAILED_AUDIT),
            "false_checks": false_checks,
            "non_broadness_failures": non_broadness_failures,
        },
        "checks": checks,
        "descriptive_absolute_tolerance": TOLERANCE,
        "maximum_broadness_absolute_difference": maximum_difference,
        "recomputed_decoding_exact_one_sided_p": decoding_p,
        "recomputed_reliability_exact_one_sided_p": reliability_p,
        "subject_artifact_sha256": subject_hashes,
        "broadness_differences": differences,
        "claim_boundary": "The tolerance applies only to exponentiated descriptive broadness metrics; no AUC, advantage, exact p value or decision threshold changed.",
    }
    OUTPUT.write_text(json.dumps(payload, indent=2) + "\n", encoding="utf-8")
    observed = sha256(OUTPUT)
    OUTPUT.with_name(OUTPUT.name + ".sha256").write_text(
        f"{observed}  {OUTPUT.name}\n", encoding="ascii"
    )
    print(
        json.dumps(
            {
                "status": status,
                "mechanism_status": payload["mechanism_status"],
                "maximum_broadness_absolute_difference": maximum_difference,
                "decoding_p": decoding_p,
                "reliability_p": reliability_p,
                "sha256": observed,
            }
        )
    )
    if status != "pass":
        raise SystemExit(1)


if __name__ == "__main__":
    main()
```
