## Supplementary material for "Falsifiable substitution tests reveal task-structured neural evidence for auditory attention": Article_Analysis_Source_Code: audit_kite_sat_akb_confirmation_complete_case_mechanism_serialization_recovery_v2.py

```
"""Serialization-only recovery for the independently computed v2 mechanism audit."""

from __future__ import annotations

import json
from pathlib import Path

import numpy as np

import audit_kite_sat_akb_mechanism_outcome_v1 as audit
from kite_sat_confirmation_complete_case_lock_utils_v2 import (
    ROOT,
    sha256,
    validate_complete_case_lock,
)


LOCKED_CALLER = (
    ROOT / "analysis"
    / "audit_kite_sat_akb_confirmation_complete_case_mechanism_v2.py"
)
ADDENDUM = (
    ROOT / "docs"
    / "kite_sat_akb_confirmation_complete_case_mechanism_audit_serialization_addendum_v2.json"
)
SUBJECTS = (
    "S0", "S10", "S2", "S21", "S28", "S22",
    "S9", "S15", "S16", "S18", "S13", "S1",
)
MECHANISM_ROOT = (
    ROOT / "results" / "kite_sat"
    / "akb_confirmation_complete_case_direction_delay_v2"
)


def validate_addendum() -> None:
    fields = ADDENDUM.with_name(ADDENDUM.name + ".sha256").read_text(
        encoding="ascii"
    ).strip().split()
    observed = sha256(ADDENDUM)
    payload = json.loads(ADDENDUM.read_text(encoding="utf-8"))
    relative = Path(__file__).resolve().relative_to(ROOT.resolve()).as_posix()
    if (
        len(fields) != 2
        or fields[0].lower() != observed
        or fields[1] != ADDENDUM.name
        or payload["status"]
        != "frozen_after_audit_computation_and_before_audit_JSON_output"
        or payload["recovery_source"]["path"] != relative
        or payload["recovery_source"]["sha256"] != sha256(Path(__file__))
        or payload["mechanism_decision_sha256"]
        != sha256(MECHANISM_ROOT / "mechanism_decision_v1.json")
    ):
        raise RuntimeError("Mechanism audit serialization addendum mismatch.")


def json_default(value):
    if isinstance(value, np.bool_):
        return bool(value)
    if isinstance(value, np.integer):
        return int(value)
    if isinstance(value, np.floating):
        return float(value)
    raise TypeError(f"Object of type {type(value).__name__} is not JSON serializable")


def main() -> None:
    validate_addendum()
    lock, lock_hash = validate_complete_case_lock(LOCKED_CALLER)
    output = MECHANISM_ROOT / "mechanism_outcome_audit_v1.json"
    if output.exists():
        raise RuntimeError("Refusing to overwrite recovered mechanism audit.")
    original_dumps = audit.json.dumps

    def safe_dumps(value, *args, **kwargs):
        kwargs.setdefault("default", json_default)
        return original_dumps(value, *args, **kwargs)

    def validate_adapter(_: Path):
        return lock, lock_hash

    audit.SUBJECTS = SUBJECTS
    audit.DECISION = MECHANISM_ROOT / "mechanism_decision_v1.json"
    audit.OUTPUT = output
    audit.validate_mechanism_lock = validate_adapter
    audit.json.dumps = safe_dumps
    audit.main()


if __name__ == "__main__":
    main()
```
