## Supplementary material for "Falsifiable substitution tests reveal task-structured neural evidence for auditory attention": Article_Analysis_Source_Code: audit_kite_sat_akb_confirmation_lock_v1.py

```
"""Preoutcome audit of the KITE-SAT AKB independent-confirmation lock."""

from __future__ import annotations

import json
from pathlib import Path

from download_kite_st_hd_confirmation_v1 import SUBJECTS
from kite_sat_confirmation_lock_utils_v1 import ROOT, sha256, validate_confirmation_lock


OUTPUT = (
    ROOT / "results" / "kite_sat" / "akb_confirmation_v1"
    / "implementation_lock_audit_v1.json"
)


def existing_files(path: Path) -> list[str]:
    if not path.exists():
        return []
    return sorted(
        item.relative_to(ROOT).as_posix()
        for item in path.rglob("*")
        if item.is_file()
    )


def main() -> None:
    if OUTPUT.exists():
        raise RuntimeError("Refusing to overwrite confirmation lock audit.")
    lock, lock_hash = validate_confirmation_lock(Path(__file__))
    checks = {
        "subject_order": tuple(lock["confirmation_subject_order"]) == SUBJECTS,
        "protocol_status": lock["protocol_status_verified_preoutcome"] is True,
    }
    raw_hits = existing_files(
        ROOT / "data" / "raw" / "kite_st_hd_zenodo4518754"
        / "confirmation_archives"
    )
    derived_hits = (
        existing_files(
            ROOT / "data" / "derived" / "kite_st_hd_zenodo4518754"
            / "confirmation_preprocessed_v1"
        )
        + existing_files(
            ROOT / "data" / "derived" / "kite_st_hd_zenodo4518754"
            / "confirmation_features_v1"
        )
        + existing_files(
            ROOT / "data" / "derived" / "kite_st_hd_zenodo4518754"
            / "kite_sat_confirmation_models_v1"
        )
    )
    preprocessing_hits = existing_files(
        ROOT / "results" / "kite_st_hd" / "confirmation_v1"
    )
    science_hits = existing_files(
        ROOT / "results" / "kite_sat" / "akb_confirmation_v1"
    )
    checks["zero_confirmation_archives"] = not raw_hits
    checks["zero_confirmation_derivatives"] = not derived_hits
    checks["zero_confirmation_preprocessing_results"] = not preprocessing_hits
    checks["zero_confirmation_science_results"] = not science_hits
    status = "pass" if all(checks.values()) else "fail"
    payload = {
        "schema": "KITE-SAT-AKB-confirmation-implementation-lock-audit-v1",
        "status": status,
        "implementation_lock_sha256": lock_hash,
        "checks": checks,
        "existing_raw_hits": raw_hits,
        "existing_derived_hits": derived_hits,
        "existing_preprocessing_hits": preprocessing_hits,
        "existing_science_hits": science_hits,
    }
    OUTPUT.parent.mkdir(parents=True, exist_ok=True)
    OUTPUT.write_text(json.dumps(payload, indent=2) + "\n", encoding="utf-8")
    observed = sha256(OUTPUT)
    OUTPUT.with_name(OUTPUT.name + ".sha256").write_text(
        f"{observed}  {OUTPUT.name}\n", encoding="ascii"
    )
    print(json.dumps({"status": status, "sha256": observed, "checks": checks}))
    if status != "pass":
        raise SystemExit(1)


if __name__ == "__main__":
    main()
```
