## Supplementary material for "Falsifiable substitution tests reveal task-structured neural evidence for auditory attention": Article_Analysis_Source_Code: audit_kite_sat_akb_confirmation_outcome_v1.py

```
"""Independent endpoint/statistical audit of the AKB confirmation decision."""

from __future__ import annotations

import hashlib
import itertools
import json
from pathlib import Path

import numpy as np

from download_kite_st_hd_confirmation_v1 import SUBJECTS
from kite_sat_confirmation_lock_utils_v1 import (
    ROOT,
    parse_sidecar,
    sha256,
    validate_confirmation_lock,
)


ANCHORS = ("TP7H", "PO3", "PO4", "TP8H", "FT8H", "AF4", "AF3", "FT7H")
DECISION = (
    ROOT / "results" / "kite_sat" / "akb_confirmation_v1"
    / "confirmation_decision_v1.json"
)
OUTPUT = DECISION.parent / "confirmation_outcome_audit_v1.json"


def auc(left: list[float], right: list[float]) -> float:
    a = np.asarray(left, dtype=np.float64)
    b = np.asarray(right, dtype=np.float64)
    difference = a[:, None] - b[None, :]
    return float(np.mean(difference > 0) + 0.5 * np.mean(difference == 0))


def exact_upper(values: np.ndarray) -> float:
    observed = float(np.mean(values))
    exceed = sum(
        float(np.mean(values * np.asarray(signs))) >= observed - 1e-15
        for signs in itertools.product((-1.0, 1.0), repeat=len(values))
    )
    return exceed / (2 ** len(values))


def stable_seed(text: str) -> int:
    return int.from_bytes(hashlib.sha256(text.encode("utf-8")).digest()[:8], "little")


def bootstrap_ci(values: np.ndarray, endpoint: str) -> list[float]:
    rng = np.random.default_rng(
        stable_seed(f"KITE-SAT-AKB-confirmation-bootstrap-v1|{endpoint}")
    )
    indices = rng.integers(0, len(values), size=(100000, len(values)))
    means = np.mean(values[indices], axis=1)
    return [
        float(np.quantile(means, 0.025)),
        float(np.quantile(means, 0.975)),
    ]


def main() -> None:
    if OUTPUT.exists():
        raise RuntimeError("Refusing to overwrite AKB confirmation outcome audit.")
    lock, lock_hash = validate_confirmation_lock(Path(__file__))
    sidecar_hash, sidecar_name = parse_sidecar(
        DECISION.with_name(DECISION.name + ".sha256")
    )
    decision_hash = sha256(DECISION)
    decision = json.loads(DECISION.read_text(encoding="utf-8"))
    checks: dict[str, bool] = {
        "decision_sidecar": (
            sidecar_name == DECISION.name and sidecar_hash == decision_hash
        ),
        "decision_lock": (
            decision["confirmation_implementation_lock_sha256"] == lock_hash
        ),
        "subject_order": decision["subject_order"] == list(SUBJECTS),
        "decision_firewall": (
            decision["firewall"]["alternative_candidate_selected"] is False
            and decision["firewall"]["alternative_patch_selected"] is False
            and decision["firewall"]["paper_mutated"] is False
        ),
    }
    records = []
    subject_hashes, model_hashes = {}, {}
    for subject in SUBJECTS:
        path = DECISION.parent / "subjects" / f"{subject}_AKB_confirmation_v1.json"
        record = json.loads(path.read_text(encoding="utf-8"))
        subject_hashes[subject] = sha256(path)
        checks[f"subject_hash:{subject}"] = (
            subject_hashes[subject] == decision["subject_artifact_sha256"][subject]
        )
        checks[f"subject_identity:{subject}"] = (
            record["status"] == "pass"
            and record["subject"] == subject
            and record["confirmation_implementation_lock_sha256"] == lock_hash
        )
        checks[f"subject_firewall:{subject}"] = (
            record["firewall"]["alternative_AKB_candidate_computed"] is False
            and record["firewall"]["alternative_patch_family_computed"] is False
            and record["firewall"]["paper_mutated"] is False
        )
        model_path = ROOT / record["model_archive"]["path"]
        model_hashes[subject] = sha256(model_path)
        checks[f"model_hash:{subject}"] = (
            model_hashes[subject]
            == record["model_archive"]["sha256"]
            == decision["model_archive_sha256"][subject]
        )
        for estimator in ("global", "AKB"):
            scores = record["intact_condition_scores"][estimator]
            story1 = auc(scores["1L"], scores["1R"])
            story2 = auc(scores["2L"], scores["2R"])
            saved = record["intact_endpoints"][estimator]
            checks[f"intact_AUC:{subject}:{estimator}"] = (
                abs(story1 - saved["story1_auc"]) < 1e-15
                and abs(story2 - saved["story2_auc"]) < 1e-15
                and abs((story1 + story2) / 2 - saved["mean_auc"]) < 1e-15
            )
            patch_means = []
            for patch_index, anchor in enumerate(ANCHORS):
                patch_scores = record["patch_condition_scores"][estimator]
                patch_story1 = auc(
                    patch_scores["1L"][patch_index],
                    patch_scores["1R"][patch_index],
                )
                patch_story2 = auc(
                    patch_scores["2L"][patch_index],
                    patch_scores["2R"][patch_index],
                )
                patch_mean = (patch_story1 + patch_story2) / 2
                patch_saved = record["patch_endpoints"][estimator]["patches"][
                    patch_index
                ]
                checks[f"patch_AUC:{subject}:{estimator}:{anchor}"] = (
                    patch_saved["anchor"] == anchor
                    and abs(patch_story1 - patch_saved["story1_auc"]) < 1e-15
                    and abs(patch_story2 - patch_saved["story2_auc"]) < 1e-15
                    and abs(patch_mean - patch_saved["mean_auc"]) < 1e-15
                )
                patch_means.append(patch_mean)
            checks[f"patch_average:{subject}:{estimator}"] = (
                abs(
                    float(np.mean(patch_means))
                    - record["patch_endpoints"][estimator]["patch_average_auc"]
                )
                < 1e-15
            )
        checks[f"intact_gain:{subject}"] = (
            abs(
                record["intact_endpoints"]["AKB"]["mean_auc"]
                - record["intact_endpoints"]["global"]["mean_auc"]
                - record["intact_gain_auc"]
            )
            < 1e-15
        )
        checks[f"patch_gain:{subject}"] = (
            abs(
                record["patch_endpoints"]["AKB"]["patch_average_auc"]
                - record["patch_endpoints"]["global"]["patch_average_auc"]
                - record["patch_average_gain_auc"]
            )
            < 1e-15
        )
        records.append(record)
    intact_global = np.asarray(
        [record["intact_endpoints"]["global"]["mean_auc"] for record in records]
    )
    intact_akb = np.asarray(
        [record["intact_endpoints"]["AKB"]["mean_auc"] for record in records]
    )
    patch_global = np.asarray(
        [record["patch_endpoints"]["global"]["patch_average_auc"] for record in records]
    )
    patch_akb = np.asarray(
        [record["patch_endpoints"]["AKB"]["patch_average_auc"] for record in records]
    )
    intact_gain = intact_akb - intact_global
    patch_gain = patch_akb - patch_global
    intact_p = exact_upper(intact_gain)
    patch_p = exact_upper(patch_gain)
    test1 = intact_p <= 0.05
    test2 = test1 and patch_p <= 0.05
    expected_status = (
        "intact_and_patch_confirmed"
        if test2
        else "intact_only_confirmed"
        if test1
        else "not_confirmed"
    )
    checks["decision_status"] = decision["status"] == expected_status
    fixed = decision["fixed_sequence_decision"]
    checks["fixed_sequence_test1"] = (
        abs(fixed["test_1_intact_gain"]["exact_one_sided_p"] - intact_p) < 1e-15
        and fixed["test_1_intact_gain"]["rejected"] is test1
    )
    checks["fixed_sequence_test2"] = (
        fixed["test_2_patch_gain"]["eligible_after_test_1"] is test1
        and abs(
            fixed["test_2_patch_gain"]["raw_exact_one_sided_p"] - patch_p
        )
        < 1e-15
        and fixed["test_2_patch_gain"]["rejected_with_confirmatory_standing"]
        is test2
    )
    patch_rows = []
    for patch_index, anchor in enumerate(ANCHORS):
        local_global = np.asarray(
            [
                record["patch_endpoints"]["global"]["patches"][patch_index]["mean_auc"]
                for record in records
            ]
        )
        local_akb = np.asarray(
            [
                record["patch_endpoints"]["AKB"]["patches"][patch_index]["mean_auc"]
                for record in records
            ]
        )
        patch_rows.append(
            {
                "anchor": anchor,
                "mean_global_auc": float(np.mean(local_global)),
                "mean_AKB_auc": float(np.mean(local_akb)),
                "mean_gain_auc": float(np.mean(local_akb - local_global)),
            }
        )
    metrics = {
        "n": len(records),
        "mean_intact_global_auc": float(np.mean(intact_global)),
        "mean_intact_AKB_auc": float(np.mean(intact_akb)),
        "mean_intact_gain_auc": float(np.mean(intact_gain)),
        "median_intact_gain_auc": float(np.median(intact_gain)),
        "intact_gain_bootstrap_95_percent_ci": bootstrap_ci(
            intact_gain, "intact_gain"
        ),
        "intact_AKB_vs_chance_exact_one_sided_p": exact_upper(intact_akb - 0.5),
        "participants_positive_intact_gain": int(np.sum(intact_gain > 0)),
        "story1_intact_AKB_group_mean": float(
            np.mean(
                [
                    record["intact_endpoints"]["AKB"]["story1_auc"]
                    for record in records
                ]
            )
        ),
        "story2_intact_AKB_group_mean": float(
            np.mean(
                [
                    record["intact_endpoints"]["AKB"]["story2_auc"]
                    for record in records
                ]
            )
        ),
        "mean_patch_average_global_auc": float(np.mean(patch_global)),
        "mean_patch_average_AKB_auc": float(np.mean(patch_akb)),
        "mean_patch_average_gain_auc": float(np.mean(patch_gain)),
        "median_patch_average_gain_auc": float(np.median(patch_gain)),
        "patch_gain_bootstrap_95_percent_ci": bootstrap_ci(
            patch_gain, "patch_gain"
        ),
        "patch_AKB_vs_chance_exact_one_sided_p": exact_upper(patch_akb - 0.5),
        "participants_positive_patch_gain": int(np.sum(patch_gain > 0)),
        "minimum_patch_specific_group_mean_AKB_auc": float(
            min(row["mean_AKB_auc"] for row in patch_rows)
        ),
        "minimum_patch_specific_group_mean_gain_auc": float(
            min(row["mean_gain_auc"] for row in patch_rows)
        ),
    }
    for key, value in metrics.items():
        saved = decision["metrics"][key]
        checks[f"group_metric:{key}"] = (
            bool(np.allclose(value, saved, atol=1e-15, rtol=0.0))
            if isinstance(value, list)
            else abs(float(value) - float(saved)) < 1e-15
        )
    checks["patch_group_metrics"] = all(
        saved["anchor"] == observed["anchor"]
        and abs(saved["mean_global_auc"] - observed["mean_global_auc"]) < 1e-15
        and abs(saved["mean_AKB_auc"] - observed["mean_AKB_auc"]) < 1e-15
        and abs(saved["mean_gain_auc"] - observed["mean_gain_auc"]) < 1e-15
        for saved, observed in zip(
            decision["patch_specific_group_metrics"], patch_rows
        )
    )
    status = "pass" if all(checks.values()) else "fail"
    payload = {
        "schema": "KITE-SAT-AKB-independent-confirmation-outcome-audit-v1",
        "status": status,
        "confirmation_decision_status": expected_status,
        "confirmation_implementation_lock_sha256": lock_hash,
        "confirmation_decision_sha256": decision_hash,
        "checks": checks,
        "recomputed_fixed_sequence": {
            "intact_gain_exact_one_sided_p": intact_p,
            "intact_rejected": test1,
            "patch_gain_raw_exact_one_sided_p": patch_p,
            "patch_eligible": test1,
            "patch_rejected_with_confirmatory_standing": test2,
        },
        "recomputed_metrics": metrics,
        "recomputed_patch_specific_group_metrics": patch_rows,
        "subject_artifact_sha256": subject_hashes,
        "model_archive_sha256": model_hashes,
    }
    OUTPUT.write_text(json.dumps(payload, indent=2) + "\n", encoding="utf-8")
    observed = sha256(OUTPUT)
    OUTPUT.with_name(OUTPUT.name + ".sha256").write_text(
        f"{observed}  {OUTPUT.name}\n", encoding="ascii"
    )
    print(
        json.dumps(
            {
                "status": status,
                "confirmation_status": expected_status,
                "fixed_sequence": payload["recomputed_fixed_sequence"],
                "metrics": metrics,
                "sha256": observed,
            }
        )
    )
    if status != "pass":
        print(json.dumps({"failed": [key for key, value in checks.items() if not value]}))
        raise SystemExit(1)


if __name__ == "__main__":
    main()
```
