## Supplementary material for "Falsifiable substitution tests reveal task-structured neural evidence for auditory attention": Article_Analysis_Source_Code: audit_kite_sat_akb_mechanism_lock_v1.py

```
"""Preoutcome audit of the AKB direction-delay mechanism lock."""

from __future__ import annotations

import json
from pathlib import Path

from download_kite_st_hd_confirmation_v1 import SUBJECTS
from kite_sat_mechanism_lock_utils_v1 import ROOT, sha256, validate_mechanism_lock


OUTPUT = (
    ROOT / "results" / "kite_sat" / "akb_direction_delay_mechanism_v1"
    / "implementation_lock_audit_v1.json"
)


def files_under(path: Path) -> list[str]:
    if not path.exists():
        return []
    return sorted(
        item.relative_to(ROOT).as_posix()
        for item in path.rglob("*")
        if item.is_file()
    )


def main() -> None:
    if OUTPUT.exists():
        raise RuntimeError("Refusing to overwrite mechanism lock audit.")
    lock, lock_hash = validate_mechanism_lock(Path(__file__))
    primary_root = ROOT / "results" / "kite_sat" / "akb_confirmation_v1"
    primary_science = []
    for path in (
        primary_root / "subjects",
        primary_root / "confirmation_decision_v1.json",
        primary_root / "confirmation_outcome_audit_v1.json",
    ):
        if path.is_file():
            primary_science.append(path.relative_to(ROOT).as_posix())
        elif path.is_dir():
            primary_science.extend(files_under(path))
    mechanism_hits = files_under(OUTPUT.parent)
    checks = {
        "subject_order": tuple(lock["confirmation_subject_order"]) == SUBJECTS,
        "no_primary_confirmation_score": not primary_science,
        "no_mechanism_output": not mechanism_hits,
        "partial_archive_allowed_but_unopened": lock[
            "acquisition_state_at_lock"
        ]["partial_archive_download_in_progress"]
        is True,
    }
    status = "pass" if all(checks.values()) else "fail"
    payload = {
        "schema": "KITE-SAT-AKB-direction-delay-mechanism-lock-audit-v1",
        "status": status,
        "mechanism_implementation_lock_sha256": lock_hash,
        "checks": checks,
        "primary_science_hits": primary_science,
        "mechanism_hits": mechanism_hits,
    }
    OUTPUT.parent.mkdir(parents=True, exist_ok=True)
    OUTPUT.write_text(json.dumps(payload, indent=2) + "\n", encoding="utf-8")
    observed = sha256(OUTPUT)
    OUTPUT.with_name(OUTPUT.name + ".sha256").write_text(
        f"{observed}  {OUTPUT.name}\n", encoding="ascii"
    )
    print(json.dumps({"status": status, "sha256": observed, "checks": checks}))
    if status != "pass":
        raise SystemExit(1)


if __name__ == "__main__":
    main()
```
