## Supplementary material for "Falsifiable substitution tests reveal task-structured neural evidence for auditory attention": Article_Analysis_Source_Code: audit_kite_sat_akb_mechanism_outcome_v1.py

```
"""Independent outcome audit of AKB direction-delay mechanism controls."""

from __future__ import annotations

import itertools
import json
from pathlib import Path

import numpy as np

from download_kite_st_hd_confirmation_v1 import SUBJECTS
from kite_sat_mechanism_lock_utils_v1 import ROOT, sha256, validate_mechanism_lock


ANCHORS = ("TP7H", "PO3", "PO4", "TP8H", "FT8H", "AF4", "AF3", "FT7H")
TRANSFORMS = tuple(
    (angular_shift, delay_swap)
    for delay_swap in range(2)
    for angular_shift in range(8)
)
DECISION = (
    ROOT / "results" / "kite_sat" / "akb_direction_delay_mechanism_v1"
    / "mechanism_decision_v1.json"
)
OUTPUT = DECISION.parent / "mechanism_outcome_audit_v1.json"


def auc(left: list[float], right: list[float]) -> float:
    a = np.asarray(left, dtype=np.float64)
    b = np.asarray(right, dtype=np.float64)
    difference = a[:, None] - b[None, :]
    return float(np.mean(difference > 0) + 0.5 * np.mean(difference == 0))


def exact_upper(values: np.ndarray) -> float:
    observed = float(np.mean(values))
    exceed = sum(
        float(np.mean(values * np.asarray(signs))) >= observed - 1e-15
        for signs in itertools.product((-1.0, 1.0), repeat=len(values))
    )
    return exceed / (2 ** len(values))


def transform(values: np.ndarray, shift: int, swap: int) -> np.ndarray:
    output = np.empty(16, dtype=np.float64)
    for delay in range(2):
        for sector in range(8):
            output[sector + 8 * delay] = values[
                (sector - shift) % 8 + 8 * (delay ^ swap)
            ]
    return output


def correlation(first: np.ndarray, second: np.ndarray) -> float:
    a = first - np.mean(first)
    b = second - np.mean(second)
    return float(
        np.sum(a * b)
        / max(float(np.sqrt(np.sum(a * a) * np.sum(b * b))), 1e-12)
    )


def broadness(reliability: np.ndarray) -> dict[str, float]:
    shifted = 2 * reliability
    weights = np.exp(shifted - np.max(shifted))
    probability = weights / np.sum(weights)
    entropy = float(-np.sum(probability * np.log(probability + 1e-300)))
    angular = probability[:8] + probability[8:]
    angular_entropy = float(-np.sum(angular * np.log(angular + 1e-300)))
    return {
        "effective_cell_number": float(np.exp(entropy)),
        "normalized_cell_entropy": float(entropy / np.log(16)),
        "maximum_cell_mass": float(np.max(probability)),
        "effective_angular_sector_number": float(np.exp(angular_entropy)),
    }


def main() -> None:
    if OUTPUT.exists():
        raise RuntimeError("Refusing to overwrite mechanism outcome audit.")
    lock, lock_hash = validate_mechanism_lock(Path(__file__))
    decision = json.loads(DECISION.read_text(encoding="utf-8"))
    sidecar = DECISION.with_name(DECISION.name + ".sha256").read_text(
        encoding="ascii"
    ).strip().split()
    checks: dict[str, bool] = {
        "decision_sidecar": (
            len(sidecar) == 2
            and sidecar[0].lower() == sha256(DECISION)
            and sidecar[1] == DECISION.name
        ),
        "decision_lock": decision["mechanism_implementation_lock_sha256"]
        == lock_hash,
        "subject_order": decision["subject_order"] == list(SUBJECTS),
    }
    records, hashes = [], {}
    for subject in SUBJECTS:
        path = DECISION.parent / "subjects" / f"{subject}_direction_delay_mechanism_v1.json"
        record = json.loads(path.read_text(encoding="utf-8"))
        hashes[subject] = sha256(path)
        checks[f"subject_hash:{subject}"] = (
            hashes[subject] == decision["subject_artifact_sha256"][subject]
        )
        checks[f"subject_identity:{subject}"] = (
            record["status"] == "pass"
            and record["subject"] == subject
            and record["mechanism_implementation_lock_sha256"] == lock_hash
        )
        reconstructed = []
        for transform_index, (shift, swap) in enumerate(TRANSFORMS):
            intact = record["intact_condition_scores"]
            story1 = auc(
                intact["1L"][transform_index], intact["1R"][transform_index]
            )
            story2 = auc(
                intact["2L"][transform_index], intact["2R"][transform_index]
            )
            patch_values = []
            patch = record["patch_condition_scores"]
            for patch_index in range(len(ANCHORS)):
                patch_story1 = auc(
                    patch["1L"][patch_index][transform_index],
                    patch["1R"][patch_index][transform_index],
                )
                patch_story2 = auc(
                    patch["2L"][patch_index][transform_index],
                    patch["2R"][patch_index][transform_index],
                )
                patch_values.append((patch_story1 + patch_story2) / 2)
            reconstructed.append(
                {
                    "angular_shift": shift,
                    "delay_swap": bool(swap),
                    "story1_auc": story1,
                    "story2_auc": story2,
                    "intact_mean_auc": (story1 + story2) / 2,
                    "patch_average_auc": float(np.mean(patch_values)),
                }
            )
        checks[f"transform_AUCs:{subject}"] = all(
            all(
                (
                    saved["angular_shift"] == observed["angular_shift"],
                    saved["delay_swap"] is observed["delay_swap"],
                    abs(saved["story1_auc"] - observed["story1_auc"]) < 1e-15,
                    abs(saved["story2_auc"] - observed["story2_auc"]) < 1e-15,
                    abs(saved["intact_mean_auc"] - observed["intact_mean_auc"]) < 1e-15,
                    abs(saved["patch_average_auc"] - observed["patch_average_auc"])
                    < 1e-15,
                )
            )
            for saved, observed in zip(record["transform_results"], reconstructed)
        )
        identity = reconstructed[0]
        intact_control = np.mean(
            [row["intact_mean_auc"] for row in reconstructed[1:]]
        )
        patch_control = np.mean(
            [row["patch_average_auc"] for row in reconstructed[1:]]
        )
        checks[f"advantages:{subject}"] = (
            abs(
                identity["intact_mean_auc"]
                - intact_control
                - record["intact_identity_advantage"]
            )
            < 1e-15
            and abs(
                identity["patch_average_auc"]
                - patch_control
                - record["patch_identity_advantage"]
            )
            < 1e-15
        )
        r1 = np.asarray(record["reliability_vectors"]["train_story_1"])
        r2 = np.asarray(record["reliability_vectors"]["train_story_2"])
        correlations = np.asarray(
            [correlation(r1, transform(r2, shift, swap)) for shift, swap in TRANSFORMS]
        )
        checks[f"reliability:{subject}"] = (
            bool(
                np.allclose(
                    correlations,
                    record["reliability_correlations"],
                    atol=1e-15,
                    rtol=0.0,
                )
            )
            and abs(
                correlations[0]
                - np.mean(correlations[1:])
                - record["reliability_identity_advantage"]
            )
            < 1e-15
        )
        for key, vector in (
            ("train_story_1", r1),
            ("train_story_2", r2),
        ):
            observed_broadness = broadness(vector)
            checks[f"broadness:{subject}:{key}"] = all(
                abs(value - record["broadness"][key][name]) < 1e-15
                for name, value in observed_broadness.items()
            )
        records.append(record)
    decoding = np.asarray(
        [record["intact_identity_advantage"] for record in records]
    )
    reliability = np.asarray(
        [record["reliability_identity_advantage"] for record in records]
    )
    patch = np.asarray(
        [record["patch_identity_advantage"] for record in records]
    )
    decoding_p = exact_upper(decoding)
    reliability_p = exact_upper(reliability)
    primary_confirmed = decision["primary_confirmation_status"] in (
        "intact_and_patch_confirmed",
        "intact_only_confirmed",
    )
    decoding_rejected = decoding_p <= 0.05
    reliability_rejected = decoding_rejected and reliability_p <= 0.05
    mechanism_supported = primary_confirmed and decoding_rejected
    expected_status = (
        "full_direction_delay_stability_supported"
        if mechanism_supported and reliability_rejected
        else "direction_delay_decoding_mechanism_supported"
        if mechanism_supported
        else "mechanism_not_supported"
    )
    checks["decision_status"] = decision["status"] == expected_status
    fixed = decision["fixed_sequence"]
    checks["fixed_sequence"] = (
        fixed["primary_intact_AKB_confirmed"] is primary_confirmed
        and abs(
            fixed["decoding_identity_advantage"]["exact_one_sided_p"] - decoding_p
        )
        < 1e-15
        and fixed["decoding_identity_advantage"]["rejected"] is decoding_rejected
        and fixed["decoding_identity_advantage"]["mechanistic_standing"]
        is mechanism_supported
        and fixed["reliability_identity_advantage"]["eligible_after_decoding_test"]
        is decoding_rejected
        and abs(
            fixed["reliability_identity_advantage"]["raw_exact_one_sided_p"]
            - reliability_p
        )
        < 1e-15
        and fixed["reliability_identity_advantage"][
            "rejected_with_mechanistic_standing"
        ]
        is reliability_rejected
    )
    broadness_rows = [
        record["broadness"][story]
        for record in records
        for story in ("train_story_1", "train_story_2")
    ]
    metrics = {
        "n": len(records),
        "mean_intact_identity_advantage_auc": float(np.mean(decoding)),
        "median_intact_identity_advantage_auc": float(np.median(decoding)),
        "participants_positive_intact_identity_advantage": int(np.sum(decoding > 0)),
        "mean_angular_only_advantage_auc": float(
            np.mean(
                [record["intact_angular_only_advantage"] for record in records]
            )
        ),
        "mean_delay_swapped_advantage_auc": float(
            np.mean(
                [record["intact_delay_swapped_advantage"] for record in records]
            )
        ),
        "mean_patch_identity_advantage_auc": float(np.mean(patch)),
        "mean_reliability_identity_advantage": float(np.mean(reliability)),
        "participants_positive_reliability_identity_advantage": int(
            np.sum(reliability > 0)
        ),
        "mean_effective_cell_number": float(
            np.mean([row["effective_cell_number"] for row in broadness_rows])
        ),
        "mean_normalized_cell_entropy": float(
            np.mean([row["normalized_cell_entropy"] for row in broadness_rows])
        ),
        "mean_maximum_cell_mass": float(
            np.mean([row["maximum_cell_mass"] for row in broadness_rows])
        ),
        "mean_effective_angular_sector_number": float(
            np.mean(
                [row["effective_angular_sector_number"] for row in broadness_rows]
            )
        ),
    }
    for key, value in metrics.items():
        checks[f"group_metric:{key}"] = (
            abs(float(value) - float(decision["metrics"][key])) < 1e-15
        )
    status = "pass" if all(checks.values()) else "fail"
    payload = {
        "schema": "KITE-SAT-AKB-direction-delay-mechanism-outcome-audit-v1",
        "status": status,
        "mechanism_decision_status": expected_status,
        "mechanism_implementation_lock_sha256": lock_hash,
        "mechanism_decision_sha256": sha256(DECISION),
        "checks": checks,
        "recomputed_decoding_exact_one_sided_p": decoding_p,
        "recomputed_reliability_exact_one_sided_p": reliability_p,
        "recomputed_metrics": metrics,
        "subject_artifact_sha256": hashes,
    }
    OUTPUT.write_text(json.dumps(payload, indent=2) + "\n", encoding="utf-8")
    observed = sha256(OUTPUT)
    OUTPUT.with_name(OUTPUT.name + ".sha256").write_text(
        f"{observed}  {OUTPUT.name}\n", encoding="ascii"
    )
    print(
        json.dumps(
            {
                "status": status,
                "mechanism_status": expected_status,
                "decoding_p": decoding_p,
                "reliability_p": reliability_p,
                "metrics": metrics,
                "sha256": observed,
            }
        )
    )
    if status != "pass":
        print(json.dumps({"failed": [key for key, value in checks.items() if not value]}))
        raise SystemExit(1)


if __name__ == "__main__":
    main()
```
