## Supplementary material for "Falsifiable substitution tests reveal task-structured neural evidence for auditory attention": Article_Analysis_Source_Code: audit_kite_sat_akb_patch_lock_v1.py

```
"""Preoutcome integrity audit for the AKB contiguous-patch experiment."""

from __future__ import annotations

import hashlib
import json
from pathlib import Path

import numpy as np

from kite_sat_patch_erasure_v1 import ANCHORS, PATCH_SIZE, fixed_patch_masks
from kite_st_hd_core_v1 import load_public_geometry


ROOT = Path(__file__).resolve().parents[1]
LOCK = ROOT / "docs" / "kite_sat_akb_patch_resilience_development_implementation_lock_v1.json"
LOCK_HASH = "c37f800c4ff7a0921f2651a66e7f02681c812aededda0ac12303da936b09765a"
OUTPUT = (
    ROOT
    / "results"
    / "kite_sat"
    / "akb_patch_resilience_development_v1"
    / "implementation_lock_audit_v1.json"
)
CONFIRMATION = (
    "S0", "S10", "S26", "S2", "S21", "S28", "S22",
    "S9", "S3", "S15", "S16", "S18", "S13", "S1",
)


def sha256(path: Path) -> str:
    digest = hashlib.sha256()
    with path.open("rb") as stream:
        for chunk in iter(lambda: stream.read(8 * 1024 * 1024), b""):
            digest.update(chunk)
    return digest.hexdigest()


def main() -> None:
    if OUTPUT.exists():
        raise RuntimeError("Refusing to overwrite patch implementation audit.")
    lock = json.loads(LOCK.read_text(encoding="utf-8"))
    checks = {"lock_hash": sha256(LOCK) == LOCK_HASH}
    for section in ("protocol",):
        record = lock[section]
        checks[f"artifact:{section}"] = sha256(ROOT / record["path"]) == record["sha256"]
    for name, record in lock["evidence_inputs"].items():
        checks[f"evidence:{name}"] = sha256(ROOT / record["path"]) == record["sha256"]
    for path_text, expected in lock["source_sha256"].items():
        checks[f"source:{path_text}"] = sha256(ROOT / path_text) == expected
    coordinate_path = ROOT / lock["evidence_inputs"]["public_coordinates"]["path"]
    geometry = load_public_geometry(coordinate_path)
    masks, members = fixed_patch_masks(geometry.labels, geometry.xy)
    checks["patch_anchor_order"] = list(ANCHORS) == lock["fixed_execution"]["patch_anchor_order"]
    checks["eight_patches"] = masks.shape == (8, 253)
    checks["each_patch_51"] = bool(np.all(np.sum(masks, axis=1) == PATCH_SIZE))
    checks["all_anchors_present"] = all(
        anchor in members[index] for index, anchor in enumerate(ANCHORS)
    )
    subject_root = OUTPUT.parent / "subjects"
    existing_subject_outputs = (
        sorted(path.name for path in subject_root.glob("*"))
        if subject_root.exists()
        else []
    )
    checks["zero_patch_subject_outputs"] = not existing_subject_outputs
    checks["zero_patch_group_output"] = not (OUTPUT.parent / "development_gate_v1.json").exists()
    raw_root = ROOT / "data" / "raw" / "kite_st_hd_zenodo4518754"
    confirmation_names = {f"{subject}.tar.gz" for subject in CONFIRMATION}
    found_confirmation = sorted(
        path.relative_to(ROOT).as_posix()
        for path in raw_root.rglob("*")
        if path.is_file() and path.name in confirmation_names
    )
    checks["no_confirmation_archive"] = not found_confirmation
    status = "pass" if all(checks.values()) else "fail"
    payload = {
        "schema": "KITE-SAT-AKB-patch-resilience-lock-audit-v1",
        "status": status,
        "lock_sha256": LOCK_HASH,
        "checks": checks,
        "patch_inventory": [
            {
                "anchor": anchor,
                "removed_count": int(np.sum(masks[index])),
                "removed_labels": list(members[index]),
            }
            for index, anchor in enumerate(ANCHORS)
        ],
        "existing_patch_subject_outputs": existing_subject_outputs,
        "found_confirmation_archives": found_confirmation,
    }
    OUTPUT.parent.mkdir(parents=True, exist_ok=True)
    OUTPUT.write_text(json.dumps(payload, indent=2) + "\n", encoding="utf-8")
    observed = sha256(OUTPUT)
    OUTPUT.with_name(OUTPUT.name + ".sha256").write_text(
        f"{observed}  {OUTPUT.name}\n", encoding="ascii"
    )
    print(json.dumps({"status": status, "sha256": observed, "checks": checks}))
    if status != "pass":
        raise SystemExit(1)


if __name__ == "__main__":
    main()
```
