## Supplementary material for "Falsifiable substitution tests reveal task-structured neural evidence for auditory attention": Article_Analysis_Source_Code: audit_kite_sat_akb_patch_outcome_v1.py

```
"""Independent outcome audit for the AKB contiguous-patch development gate."""

from __future__ import annotations

import hashlib
import itertools
import json
from pathlib import Path

import numpy as np


ROOT = Path(__file__).resolve().parents[1]
PROTOCOL = ROOT / "docs" / "kite_sat_akb_patch_resilience_development_protocol_v1.json"
PROTOCOL_HASH = "ff32e236996b761cc059e03f7f775aac84cd3322cab2d220cf44efff219d1cb4"
LOCK = ROOT / "docs" / "kite_sat_akb_patch_resilience_development_implementation_lock_v1.json"
LOCK_HASH = "c37f800c4ff7a0921f2651a66e7f02681c812aededda0ac12303da936b09765a"
GATE = (
    ROOT / "results" / "kite_sat" / "akb_patch_resilience_development_v1"
    / "development_gate_v1.json"
)
GATE_HASH = "524c922712ae5fb071fed5c7e9a86a124671d7bcc74297091367ddd797696cad"
OUTPUT = GATE.parent / "development_outcome_audit_v1.json"
SUBJECTS = (
    "S4", "S24", "S11", "S23", "S14", "S29",
    "S19", "S17", "S12", "S25", "S27", "S5",
)
ANCHORS = ("TP7H", "PO3", "PO4", "TP8H", "FT8H", "AF4", "AF3", "FT7H")
CONFIRMATION = (
    "S0", "S10", "S26", "S2", "S21", "S28", "S22",
    "S9", "S3", "S15", "S16", "S18", "S13", "S1",
)


def sha256(path: Path) -> str:
    digest = hashlib.sha256()
    with path.open("rb") as stream:
        for chunk in iter(lambda: stream.read(8 * 1024 * 1024), b""):
            digest.update(chunk)
    return digest.hexdigest()


def auc(left: list[float], right: list[float]) -> float:
    a = np.asarray(left, dtype=np.float64)
    b = np.asarray(right, dtype=np.float64)
    difference = a[:, None] - b[None, :]
    return float(np.mean(difference > 0) + 0.5 * np.mean(difference == 0))


def exact_upper(values: np.ndarray) -> float:
    observed = float(np.mean(values))
    exceed = sum(
        float(np.mean(values * np.asarray(signs))) >= observed - 1e-15
        for signs in itertools.product((-1.0, 1.0), repeat=len(values))
    )
    return exceed / (2 ** len(values))


def stable_seed(text: str) -> int:
    return int.from_bytes(hashlib.sha256(text.encode("utf-8")).digest()[:8], "little")


def main() -> None:
    if OUTPUT.exists():
        raise RuntimeError("Refusing to overwrite patch outcome audit.")
    gate = json.loads(GATE.read_text(encoding="utf-8"))
    lock = json.loads(LOCK.read_text(encoding="utf-8"))
    checks: dict[str, bool] = {
        "protocol_hash": sha256(PROTOCOL) == PROTOCOL_HASH,
        "implementation_lock_hash": sha256(LOCK) == LOCK_HASH,
        "gate_hash": sha256(GATE) == GATE_HASH,
        "gate_status": gate["status"] == "advance_patch_endpoint_to_confirmation",
        "gate_subject_order": gate["subject_order"] == list(SUBJECTS),
        "all_gate_checks_true": all(gate["gate_checks"].values()),
        "confirmation_download_false": gate["firewall"]["confirmation_archives_downloaded"]
        is False,
        "confirmation_read_false": gate["firewall"]["confirmation_subject_read"] is False,
        "paper_false": gate["firewall"]["paper_mutated"] is False,
    }
    for path_text, expected in lock["source_sha256"].items():
        checks[f"locked_source:{path_text}"] = sha256(ROOT / path_text) == expected
    records = []
    subject_hashes = {}
    for subject in SUBJECTS:
        path = GATE.parent / "subjects" / f"{subject}_patch_resilience_v1.json"
        record = json.loads(path.read_text(encoding="utf-8"))
        subject_hashes[subject] = sha256(path)
        checks[f"subject_hash:{subject}"] = (
            subject_hashes[subject] == gate["subject_artifact_sha256"][subject]
        )
        checks[f"subject_identity:{subject}"] = (
            record["status"] == "pass"
            and record["subject"] == subject
            and record["protocol_sha256"] == PROTOCOL_HASH
        )
        checks[f"subject_firewall:{subject}"] = (
            record["firewall"]["patch_variant_computed"] is False
            and record["firewall"]["confirmation_subject_read"] is False
        )
        checks[f"patch_order:{subject}"] = (
            [row["anchor"] for row in record["patches"]] == list(ANCHORS)
            and all(row["removed_count"] == 51 for row in record["patches"])
        )
        for estimator in ("global", "AKB"):
            recomputed_patch_means = []
            for patch_index, anchor in enumerate(ANCHORS):
                scores = record["condition_scores"][estimator]
                story1 = auc(scores["1L"][patch_index], scores["1R"][patch_index])
                story2 = auc(scores["2L"][patch_index], scores["2R"][patch_index])
                mean_auc = (story1 + story2) / 2
                saved = record["endpoints"][estimator]["patches"][patch_index]
                checks[f"AUC:{subject}:{estimator}:{anchor}"] = (
                    saved["anchor"] == anchor
                    and abs(story1 - saved["story1_auc"]) < 1e-15
                    and abs(story2 - saved["story2_auc"]) < 1e-15
                    and abs(mean_auc - saved["mean_auc"]) < 1e-15
                )
                recomputed_patch_means.append(mean_auc)
            checks[f"patch_average:{subject}:{estimator}"] = (
                abs(
                    float(np.mean(recomputed_patch_means))
                    - record["endpoints"][estimator]["patch_average_auc"]
                )
                < 1e-15
            )
        checks[f"gain:{subject}"] = (
            abs(
                record["endpoints"]["AKB"]["patch_average_auc"]
                - record["endpoints"]["global"]["patch_average_auc"]
                - record["patch_average_gain_auc"]
            )
            < 1e-15
        )
        records.append(record)
    global_auc = np.asarray(
        [record["endpoints"]["global"]["patch_average_auc"] for record in records]
    )
    akb_auc = np.asarray(
        [record["endpoints"]["AKB"]["patch_average_auc"] for record in records]
    )
    gain = akb_auc - global_auc
    patch_rows = []
    for patch_index, anchor in enumerate(ANCHORS):
        patch_global = np.asarray(
            [
                record["endpoints"]["global"]["patches"][patch_index]["mean_auc"]
                for record in records
            ]
        )
        patch_akb = np.asarray(
            [
                record["endpoints"]["AKB"]["patches"][patch_index]["mean_auc"]
                for record in records
            ]
        )
        patch_rows.append(
            {
                "anchor": anchor,
                "mean_global_auc": float(np.mean(patch_global)),
                "mean_AKB_auc": float(np.mean(patch_akb)),
                "mean_gain_auc": float(np.mean(patch_akb - patch_global)),
            }
        )
    rng = np.random.default_rng(
        stable_seed("KITE-SAT-AKB-patch-resilience-development-bootstrap-v1")
    )
    indices = rng.integers(0, len(gain), size=(20000, len(gain)))
    bootstrap = np.mean(gain[indices], axis=1)
    metrics = {
        "n": len(gain),
        "mean_patch_average_global_auc": float(np.mean(global_auc)),
        "mean_patch_average_AKB_auc": float(np.mean(akb_auc)),
        "mean_patch_average_gain_auc": float(np.mean(gain)),
        "median_patch_average_gain_auc": float(np.median(gain)),
        "gain_bootstrap_95_percent_ci": [
            float(np.quantile(bootstrap, 0.025)),
            float(np.quantile(bootstrap, 0.975)),
        ],
        "paired_gain_exact_one_sided_p": exact_upper(gain),
        "AKB_vs_chance_exact_one_sided_p": exact_upper(akb_auc - 0.5),
        "participants_positive_gain": int(np.sum(gain > 0)),
        "minimum_patch_specific_group_mean_AKB_auc": float(
            min(row["mean_AKB_auc"] for row in patch_rows)
        ),
        "minimum_patch_specific_group_mean_gain_auc": float(
            min(row["mean_gain_auc"] for row in patch_rows)
        ),
    }
    for key, value in metrics.items():
        saved = gate["metrics"][key]
        checks[f"group_metric:{key}"] = (
            bool(np.allclose(value, saved, atol=1e-15, rtol=0.0))
            if isinstance(value, list)
            else abs(float(value) - float(saved)) < 1e-15
        )
    checks["patch_group_metrics"] = all(
        saved["anchor"] == observed["anchor"]
        and abs(saved["mean_global_auc"] - observed["mean_global_auc"]) < 1e-15
        and abs(saved["mean_AKB_auc"] - observed["mean_AKB_auc"]) < 1e-15
        and abs(saved["mean_gain_auc"] - observed["mean_gain_auc"]) < 1e-15
        for saved, observed in zip(gate["patch_specific_group_metrics"], patch_rows)
    )
    raw_root = ROOT / "data" / "raw" / "kite_st_hd_zenodo4518754"
    confirmation_names = {f"{subject}.tar.gz" for subject in CONFIRMATION}
    found_confirmation = sorted(
        path.relative_to(ROOT).as_posix()
        for path in raw_root.rglob("*")
        if path.is_file() and path.name in confirmation_names
    )
    checks["no_confirmation_archive"] = not found_confirmation
    status = "pass" if all(checks.values()) else "fail"
    payload = {
        "schema": "KITE-SAT-AKB-patch-resilience-development-outcome-audit-v1",
        "status": status,
        "protocol_sha256": PROTOCOL_HASH,
        "implementation_lock_sha256": LOCK_HASH,
        "gate_sha256": GATE_HASH,
        "checks": checks,
        "recomputed_metrics": metrics,
        "recomputed_patch_specific_group_metrics": patch_rows,
        "subject_artifact_sha256": subject_hashes,
        "found_confirmation_archives": found_confirmation,
    }
    OUTPUT.write_text(json.dumps(payload, indent=2) + "\n", encoding="utf-8")
    observed = sha256(OUTPUT)
    OUTPUT.with_name(OUTPUT.name + ".sha256").write_text(
        f"{observed}  {OUTPUT.name}\n", encoding="ascii"
    )
    print(json.dumps({"status": status, "sha256": observed, "metrics": metrics}))
    if status != "pass":
        print(json.dumps({"failed": [key for key, value in checks.items() if not value]}))
        raise SystemExit(1)


if __name__ == "__main__":
    main()
```
