## Supplementary material for "Falsifiable substitution tests reveal task-structured neural evidence for auditory attention": Article_Analysis_Source_Code: audit_kite_sat_akb_stability_lock_v2.py

```
"""Preoutcome audit after the AKB boolean-token repair."""

from __future__ import annotations

import hashlib
import json
from pathlib import Path


ROOT = Path(__file__).resolve().parents[1]
LOCK = ROOT / "docs" / "kite_sat_akb_stability_development_implementation_lock_v2.json"
LOCK_HASH = "a8d5a68ca9b71ed42545ecfbe0dc4556673deea05f459d76735587ea5ec95c3c"
OUTPUT = ROOT / "results" / "kite_sat" / "akb_stability_development_v1" / "implementation_lock_audit_v2.json"
CONFIRMATION = ("S0", "S10", "S26", "S2", "S21", "S28", "S22", "S9", "S3", "S15", "S16", "S18", "S13", "S1")


def sha256(path: Path) -> str:
    digest = hashlib.sha256()
    with path.open("rb") as stream:
        for chunk in iter(lambda: stream.read(8 * 1024 * 1024), b""):
            digest.update(chunk)
    return digest.hexdigest()


def main() -> None:
    if OUTPUT.exists():
        raise RuntimeError("Refusing to overwrite AKB v2 lock audit.")
    lock = json.loads(LOCK.read_text(encoding="utf-8"))
    checks = {"lock_hash": sha256(LOCK) == LOCK_HASH}
    for key in ("supersedes", "incident", "protocol", "postengineering_screen"):
        record = lock[key]
        checks[f"artifact:{key}"] = sha256(ROOT / record["path"]) == record["sha256"]
    for path_text, expected in lock["sources"].items():
        checks[f"source:{path_text}"] = sha256(ROOT / path_text) == expected
    subject_root = OUTPUT.parent / "subjects"
    existing = (
        sorted(path.name for path in subject_root.glob("*"))
        if subject_root.exists()
        else []
    )
    checks["zero_subject_artifacts"] = not existing
    raw_root = ROOT / "data" / "raw" / "kite_st_hd_zenodo4518754"
    names = {f"{subject}.tar.gz" for subject in CONFIRMATION}
    found_confirmation = (
        sorted(
            path.relative_to(ROOT).as_posix()
            for path in raw_root.rglob("*")
            if path.is_file() and path.name in names
        )
        if raw_root.exists()
        else []
    )
    checks["no_confirmation_archive"] = not found_confirmation
    checks["repair_declares_no_science_change"] = (
        lock["repair_diff"]["scientific_change"] is False
    )
    status = "pass" if all(checks.values()) else "fail"
    payload = {
        "schema": "KITE-SAT-AKB-stability-lock-audit-v2",
        "status": status,
        "lock_sha256": LOCK_HASH,
        "checks": checks,
        "existing_subject_artifacts": existing,
        "found_confirmation_archives": found_confirmation,
    }
    OUTPUT.parent.mkdir(parents=True, exist_ok=True)
    OUTPUT.write_text(json.dumps(payload, indent=2) + "\n", encoding="utf-8")
    observed = sha256(OUTPUT)
    OUTPUT.with_name(OUTPUT.name + ".sha256").write_text(
        f"{observed}  {OUTPUT.name}\n", encoding="ascii"
    )
    print(json.dumps({"status": status, "sha256": observed, "checks": checks}))
    if status != "pass":
        raise SystemExit(1)


if __name__ == "__main__":
    main()
```
