## Supplementary material for "Falsifiable substitution tests reveal task-structured neural evidence for auditory attention": Article_Analysis_Source_Code: audit_kite_sat_akb_stability_outcome_v1.py

```
"""Independent outcome audit for the KITE-SAT AKB stability development gate.

This audit deliberately does not import either AKB production runner.  It
recomputes every participant AUC from saved block scores, both exact sign-flip
tests, the deterministic participant bootstrap, and all artifact hashes.
"""

from __future__ import annotations

import hashlib
import itertools
import json
from pathlib import Path

import numpy as np


ROOT = Path(__file__).resolve().parents[1]
PROTOCOL = ROOT / "docs" / "kite_sat_akb_stability_development_protocol_v1.json"
PROTOCOL_HASH = "12be0a01ed654cab61f54dffc6df84e34ac2bce5ddb17653ee81aea695a2b883"
LOCK = ROOT / "docs" / "kite_sat_akb_stability_development_implementation_lock_v2.json"
LOCK_HASH = "a8d5a68ca9b71ed42545ecfbe0dc4556673deea05f459d76735587ea5ec95c3c"
GATE = ROOT / "results" / "kite_sat" / "akb_stability_development_v1" / "development_gate_v1.json"
GATE_HASH = "6a29be774fb3d68016b0d4a6d59d6f627c7a8c297a67bf68696af8aa7a8b2e5f"
OUTPUT = GATE.parent / "development_outcome_audit_v1.json"
SUBJECTS = (
    "S4", "S24", "S11", "S23", "S14", "S29",
    "S19", "S17", "S12", "S25", "S27", "S5",
)
CONFIRMATION = (
    "S0", "S10", "S26", "S2", "S21", "S28", "S22",
    "S9", "S3", "S15", "S16", "S18", "S13", "S1",
)


def sha256(path: Path) -> str:
    digest = hashlib.sha256()
    with path.open("rb") as stream:
        for chunk in iter(lambda: stream.read(8 * 1024 * 1024), b""):
            digest.update(chunk)
    return digest.hexdigest()


def auc(left_scores: list[float], right_scores: list[float]) -> float:
    """Pairwise AUC: probability that a left trial outranks a right trial."""
    left = np.asarray(left_scores, dtype=np.float64)
    right = np.asarray(right_scores, dtype=np.float64)
    comparisons = left[:, None] - right[None, :]
    return float(np.mean(comparisons > 0) + 0.5 * np.mean(comparisons == 0))


def exact_upper(values: np.ndarray) -> float:
    observed = float(np.mean(values))
    exceed = 0
    for signs in itertools.product((-1.0, 1.0), repeat=len(values)):
        statistic = float(np.mean(values * np.asarray(signs, dtype=np.float64)))
        exceed += statistic >= observed - 1e-15
    return exceed / (2 ** len(values))


def stable_seed(text: str) -> int:
    return int.from_bytes(hashlib.sha256(text.encode("utf-8")).digest()[:8], "little")


def main() -> None:
    if OUTPUT.exists():
        raise RuntimeError("Refusing to overwrite AKB development outcome audit.")
    gate = json.loads(GATE.read_text(encoding="utf-8"))
    lock = json.loads(LOCK.read_text(encoding="utf-8"))
    checks: dict[str, bool] = {
        "protocol_hash": sha256(PROTOCOL) == PROTOCOL_HASH,
        "implementation_lock_hash": sha256(LOCK) == LOCK_HASH,
        "gate_hash": sha256(GATE) == GATE_HASH,
        "gate_status": gate["status"] == "advance_to_AKB_confirmation_lock",
        "gate_protocol": gate["protocol_sha256"] == PROTOCOL_HASH,
        "all_predeclared_subjects": gate["subject_order"] == list(SUBJECTS),
        "all_gate_checks_true": all(gate["gate_checks"].values()),
        "confirmation_read_false": gate["firewall"]["confirmation_subject_read"] is False,
        "confirmation_download_false": gate["firewall"]["confirmation_archives_downloaded"] is False,
        "paper_mutated_false": gate["firewall"]["paper_mutated"] is False,
    }
    for path_text, expected in lock["sources"].items():
        checks[f"locked_source:{path_text}"] = sha256(ROOT / path_text) == expected

    subject_root = GATE.parent / "subjects"
    global_values: list[float] = []
    akb_values: list[float] = []
    story1_values: list[float] = []
    story2_values: list[float] = []
    subject_hashes: dict[str, str] = {}
    baseline_hashes: dict[str, str] = {}
    model_hashes: dict[str, str] = {}
    recomputed_subjects: list[dict[str, float | str]] = []
    for subject in SUBJECTS:
        path = subject_root / f"{subject}_S16_softmax2_v1.json"
        record = json.loads(path.read_text(encoding="utf-8"))
        observed_hash = sha256(path)
        subject_hashes[subject] = observed_hash
        checks[f"subject_hash:{subject}"] = (
            observed_hash == gate["subject_artifact_sha256"][subject]
        )
        checks[f"subject_protocol:{subject}"] = record["protocol_sha256"] == PROTOCOL_HASH
        checks[f"subject_status:{subject}"] = (
            record["status"] == "pass" and record["subject"] == subject
        )
        checks[f"subject_firewall:{subject}"] = (
            record["firewall"]["confirmation_subject_read"] is False
        )
        baseline_path = ROOT / record["baseline_artifact"]["path"]
        model_path = ROOT / record["model_archive"]["path"]
        baseline_hashes[subject] = sha256(baseline_path)
        model_hashes[subject] = sha256(model_path)
        checks[f"baseline_hash:{subject}"] = (
            baseline_hashes[subject] == record["baseline_artifact"]["sha256"]
        )
        checks[f"model_hash:{subject}"] = (
            model_hashes[subject] == record["model_archive"]["sha256"]
        )
        baseline = json.loads(baseline_path.read_text(encoding="utf-8"))
        endpoints: dict[str, tuple[float, float, float]] = {}
        for estimator in ("global", "AKB"):
            scores = record["condition_scores"][estimator]
            story1 = auc(scores["1L"], scores["1R"])
            story2 = auc(scores["2L"], scores["2R"])
            mean = (story1 + story2) / 2.0
            endpoints[estimator] = (story1, story2, mean)
            saved = record["endpoints"][estimator]
            checks[f"AUC_recomputed:{subject}:{estimator}"] = (
                abs(story1 - saved["story1_auc"]) < 1e-15
                and abs(story2 - saved["story2_auc"]) < 1e-15
                and abs(mean - saved["mean_auc"]) < 1e-15
            )
        global_mean = endpoints["global"][2]
        akb_mean = endpoints["AKB"][2]
        checks[f"baseline_reproduced:{subject}"] = (
            abs(global_mean - baseline["mean_auc"]) < 1e-15
        )
        checks[f"gain_recomputed:{subject}"] = (
            abs((akb_mean - global_mean) - record["gain_auc"]) < 1e-15
        )
        global_values.append(global_mean)
        akb_values.append(akb_mean)
        story1_values.append(endpoints["AKB"][0])
        story2_values.append(endpoints["AKB"][1])
        recomputed_subjects.append(
            {
                "subject": subject,
                "global_auc": global_mean,
                "AKB_auc": akb_mean,
                "gain_auc": akb_mean - global_mean,
            }
        )

    global_array = np.asarray(global_values)
    akb_array = np.asarray(akb_values)
    gain = akb_array - global_array
    rng = np.random.default_rng(
        stable_seed("KITE-SAT-AKB-stability-development-bootstrap-v1")
    )
    indices = rng.integers(0, len(gain), size=(20000, len(gain)))
    bootstrap = np.mean(gain[indices], axis=1)
    recomputed = {
        "n": len(gain),
        "mean_global_auc": float(np.mean(global_array)),
        "mean_AKB_auc": float(np.mean(akb_array)),
        "mean_gain_auc": float(np.mean(gain)),
        "median_gain_auc": float(np.median(gain)),
        "gain_bootstrap_95_percent_ci": [
            float(np.quantile(bootstrap, 0.025)),
            float(np.quantile(bootstrap, 0.975)),
        ],
        "paired_gain_exact_one_sided_p": exact_upper(gain),
        "AKB_vs_chance_exact_one_sided_p": exact_upper(akb_array - 0.5),
        "participants_positive_gain": int(np.sum(gain > 0)),
        "participants_nonnegative_gain": int(np.sum(gain >= 0)),
        "story1_AKB_group_mean": float(np.mean(story1_values)),
        "story2_AKB_group_mean": float(np.mean(story2_values)),
    }
    for key, value in recomputed.items():
        saved = gate["metrics"][key]
        if isinstance(value, list):
            checks[f"group_metric:{key}"] = bool(
                np.allclose(value, saved, atol=1e-15, rtol=0.0)
            )
        else:
            checks[f"group_metric:{key}"] = abs(float(value) - float(saved)) < 1e-15

    raw_root = ROOT / "data" / "raw" / "kite_st_hd_zenodo4518754"
    confirmation_names = {f"{subject}.tar.gz" for subject in CONFIRMATION}
    found_confirmation = sorted(
        path.relative_to(ROOT).as_posix()
        for path in raw_root.rglob("*")
        if path.is_file() and path.name in confirmation_names
    )
    derived_root = ROOT / "data" / "derived" / "kite_st_hd_zenodo4518754"
    forbidden_derived_names = {
        f"{subject}_AAD_{condition}_preprocessed_v1.npz"
        for subject in CONFIRMATION
        for condition in ("1L", "1R", "2L", "2R")
    }
    found_confirmation_derived = sorted(
        path.relative_to(ROOT).as_posix()
        for path in derived_root.rglob("*.npz")
        if path.name in forbidden_derived_names
    )
    checks["no_confirmation_archive"] = not found_confirmation
    checks["no_confirmation_preprocessed_artifact"] = not found_confirmation_derived

    status = "pass" if all(checks.values()) else "fail"
    payload = {
        "schema": "KITE-SAT-AKB-stability-development-outcome-audit-v1",
        "status": status,
        "protocol_sha256": PROTOCOL_HASH,
        "implementation_lock_sha256": LOCK_HASH,
        "gate_sha256": GATE_HASH,
        "checks": checks,
        "recomputed_subjects": recomputed_subjects,
        "recomputed_metrics": recomputed,
        "subject_artifact_sha256": subject_hashes,
        "baseline_artifact_sha256": baseline_hashes,
        "model_archive_sha256": model_hashes,
        "found_confirmation_archives": found_confirmation,
        "found_confirmation_preprocessed_artifacts": found_confirmation_derived,
    }
    OUTPUT.write_text(json.dumps(payload, indent=2) + "\n", encoding="utf-8")
    observed = sha256(OUTPUT)
    OUTPUT.with_name(OUTPUT.name + ".sha256").write_text(
        f"{observed}  {OUTPUT.name}\n", encoding="ascii"
    )
    print(json.dumps({"status": status, "sha256": observed, "metrics": recomputed}))
    if status != "pass":
        failed = [key for key, value in checks.items() if not value]
        print(json.dumps({"failed_checks": failed}))
        raise SystemExit(1)


if __name__ == "__main__":
    main()
```
