## Supplementary material for "Falsifiable substitution tests reveal task-structured neural evidence for auditory attention": Article_Analysis_Source_Code: audit_kite_sat_formal_development_outcome_v1.py

```
"""Outcome integrity audit for the KITE-SAT formal development benchmark."""

from __future__ import annotations

import hashlib
import itertools
import json
from pathlib import Path

import numpy as np


ROOT = Path(__file__).resolve().parents[1]
PROTOCOL_HASH = "9d16d5499e281a582531b9d87a04e52da7228b3982fc220acb25add5d382c07e"
LOCK_HASH = "192a112abb1a5e81161b7c6ba85e8844d9ba5b03abbeb3050d51134eccaba67a"
GATE = ROOT / "results" / "kite_sat" / "formal_development_v1" / "development_gate_v1.json"
GATE_HASH = "c6e2bef9bae40e6fb1738b7712d7ba4b7e9f74d9eac0772405007505f44df207"
OUTPUT = ROOT / "results" / "kite_sat" / "formal_development_v1" / "development_outcome_audit_v1.json"
SUBJECTS = ("S4", "S24", "S11", "S23", "S14", "S29", "S19", "S17", "S12", "S25", "S27", "S5")
CONFIRMATION = ("S0", "S10", "S26", "S2", "S21", "S28", "S22", "S9", "S3", "S15", "S16", "S18", "S13", "S1")


def sha256(path: Path) -> str:
    digest = hashlib.sha256()
    with path.open("rb") as stream:
        for chunk in iter(lambda: stream.read(8 * 1024 * 1024), b""):
            digest.update(chunk)
    return digest.hexdigest()


def main() -> None:
    if OUTPUT.exists():
        raise RuntimeError("Refusing to overwrite formal development outcome audit.")
    gate = json.loads(GATE.read_text(encoding="utf-8"))
    checks = {
        "gate_hash": sha256(GATE) == GATE_HASH,
        "gate_status": gate["status"] == "advance_to_kakeya_method_development",
        "gate_protocol": gate["protocol_sha256"] == PROTOCOL_HASH,
        "all_predeclared_subjects": gate["subject_order"] == list(SUBJECTS),
        "all_gate_checks_true": all(gate["gate_checks"].values()),
        "confirmation_read_false": gate["firewall"]["confirmation_subject_read"] is False,
        "paper_mutated_false": gate["firewall"]["paper_mutated"] is False,
    }
    subject_root = GATE.parent / "subjects"
    endpoints = []
    artifact_hashes = {}
    model_hashes = {}
    for subject in SUBJECTS:
        path = subject_root / f"{subject}_speech_decoder_v1.json"
        record = json.loads(path.read_text(encoding="utf-8"))
        artifact_hashes[subject] = sha256(path)
        model_path = ROOT / record["model_archive"]["path"]
        model_hashes[subject] = sha256(model_path)
        checks[f"subject_hash:{subject}"] = (
            artifact_hashes[subject] == gate["subject_artifact_sha256"][subject]
        )
        checks[f"model_hash:{subject}"] = (
            model_hashes[subject] == record["model_archive"]["sha256"]
        )
        endpoints.append(record["mean_auc"])
    values = np.asarray(endpoints, dtype=np.float64) - 0.5
    observed = float(np.mean(values))
    exceed = sum(
        float(np.mean(values * np.asarray(signs))) >= observed - 1e-15
        for signs in itertools.product((-1.0, 1.0), repeat=len(values))
    )
    recomputed_p = exceed / (2 ** len(values))
    checks["exact_p_recomputed"] = (
        abs(recomputed_p - gate["metrics"]["exact_one_sided_sign_flip_p"]) < 1e-15
    )
    raw_root = ROOT / "data" / "raw" / "kite_st_hd_zenodo4518754"
    confirmation_names = {f"{subject}.tar.gz" for subject in CONFIRMATION}
    found_confirmation = (
        sorted(
            path.relative_to(ROOT).as_posix()
            for path in raw_root.rglob("*")
            if path.is_file() and path.name in confirmation_names
        )
        if raw_root.exists()
        else []
    )
    checks["no_confirmation_archive"] = not found_confirmation
    status = "pass" if all(checks.values()) else "fail"
    payload = {
        "schema": "KITE-SAT-formal-development-outcome-audit-v1",
        "status": status,
        "protocol_sha256": PROTOCOL_HASH,
        "implementation_lock_sha256": LOCK_HASH,
        "gate_sha256": GATE_HASH,
        "checks": checks,
        "recomputed_exact_p": recomputed_p,
        "subject_artifact_sha256": artifact_hashes,
        "model_archive_sha256": model_hashes,
        "found_confirmation_archives": found_confirmation,
    }
    OUTPUT.write_text(json.dumps(payload, indent=2) + "\n", encoding="utf-8")
    output_hash = sha256(OUTPUT)
    OUTPUT.with_name(OUTPUT.name + ".sha256").write_text(
        f"{output_hash}  {OUTPUT.name}\n", encoding="ascii"
    )
    print(json.dumps({"status": status, "sha256": output_hash, "recomputed_p": recomputed_p}))
    if status != "pass":
        raise SystemExit(1)


if __name__ == "__main__":
    main()
```
