## Supplementary material for "Falsifiable substitution tests reveal task-structured neural evidence for auditory attention": Article_Analysis_Source_Code: audit_kite_sat_sensor_density_outcome_v1.py

```
"""Independent audit of the post-confirmation sensor-density analysis."""

from __future__ import annotations

import itertools
import json
from pathlib import Path

import numpy as np

from kite_sat_sensor_density_lock_utils_v1 import (
    ROOT,
    sha256,
    validate_sensor_density_lock,
)


SUBJECTS = (
    "S0", "S10", "S2", "S21", "S28", "S22",
    "S9", "S15", "S16", "S18", "S13", "S1",
)
BUDGETS = (32, 64, 128, 253)
ROOT_RESULT = ROOT / "results" / "kite_sat" / "sensor_density_mechanism_v1"
DECISION = ROOT_RESULT / "sensor_density_decision_v1.json"
OUTPUT = ROOT_RESULT / "sensor_density_outcome_audit_v1.json"


def exact(values: np.ndarray) -> float:
    observed = float(np.mean(values))
    exceed = 0
    for signs in itertools.product((-1.0, 1.0), repeat=len(values)):
        exceed += (
            float(np.mean(values * np.asarray(signs))) >= observed - 1e-15
        )
    return exceed / (2 ** len(values))


def slope(values: np.ndarray) -> float:
    x = np.log2(np.asarray(BUDGETS, dtype=np.float64) / 32.0)
    x -= np.mean(x)
    return float(np.sum(x * values) / np.sum(x * x))


def main() -> None:
    if OUTPUT.exists():
        raise RuntimeError("Refusing to overwrite sensor-density audit.")
    lock, lock_hash = validate_sensor_density_lock(Path(__file__))
    decision = json.loads(DECISION.read_text(encoding="utf-8"))
    sidecar = DECISION.with_name(DECISION.name + ".sha256").read_text(
        encoding="ascii"
    ).strip().split()
    checks = {
        "decision_sidecar": (
            len(sidecar) == 2
            and sidecar[0].lower() == sha256(DECISION)
            and sidecar[1] == DECISION.name
        ),
        "decision_lock": (
            decision["sensor_density_implementation_lock_sha256"] == lock_hash
        ),
        "subject_order": decision["subject_order"] == list(SUBJECTS),
        "postconfirmation_label": (
            decision["inferential_status"] == "post-confirmation_mechanistic"
        ),
    }
    records = []
    hashes = {}
    for subject in SUBJECTS:
        path = ROOT_RESULT / "subjects" / f"{subject}_sensor_density_v1.json"
        record = json.loads(path.read_text(encoding="utf-8"))
        hashes[subject] = sha256(path)
        checks[f"subject:{subject}"] = (
            record["status"] == "pass"
            and record["subject"] == subject
            and record["sensor_density_implementation_lock_sha256"] == lock_hash
            and hashes[subject] == decision["subject_artifact_sha256"][subject]
            and max(record["full_budget_primary_absolute_differences"].values())
            <= 1e-12
        )
        records.append(record)
    gains = np.asarray(
        [
            [row["AKB_minus_global_auc"] for row in record["budget_results"]]
            for record in records
        ]
    )
    topology = np.asarray(
        [
            [
                row["identity_minus_mean_topology_controls_auc"]
                for row in record["budget_results"]
            ]
            for record in records
        ]
    )
    global_auc = np.asarray(
        [
            [row["global_mean_auc"] for row in record["budget_results"]]
            for record in records
        ]
    )
    akb_auc = np.asarray(
        [
            [row["AKB_mean_auc"] for row in record["budget_results"]]
            for record in records
        ]
    )
    expected_budget = []
    for index, budget in enumerate(BUDGETS):
        expected_budget.append(
            {
                "budget": budget,
                "mean_global_auc": float(np.mean(global_auc[:, index])),
                "mean_AKB_auc": float(np.mean(akb_auc[:, index])),
                "mean_AKB_minus_global_auc": float(np.mean(gains[:, index])),
                "AKB_minus_global_exact_one_sided_p": exact(gains[:, index]),
                "participants_positive_gain": int(np.sum(gains[:, index] > 0)),
                "mean_identity_minus_controls_auc": float(
                    np.mean(topology[:, index])
                ),
                "identity_minus_controls_exact_one_sided_p": exact(
                    topology[:, index]
                ),
                "participants_positive_topology_advantage": int(
                    np.sum(topology[:, index] > 0)
                ),
            }
        )
    checks["budget_metrics"] = all(
        all(
            (
                saved["budget"] == observed["budget"],
                abs(saved["mean_global_auc"] - observed["mean_global_auc"]) < 1e-15,
                abs(saved["mean_AKB_auc"] - observed["mean_AKB_auc"]) < 1e-15,
                abs(
                    saved["mean_AKB_minus_global_auc"]
                    - observed["mean_AKB_minus_global_auc"]
                )
                < 1e-15,
                abs(
                    saved["AKB_minus_global_exact_one_sided_p"]
                    - observed["AKB_minus_global_exact_one_sided_p"]
                )
                < 1e-15,
                saved["participants_positive_gain"]
                == observed["participants_positive_gain"],
                abs(
                    saved["mean_identity_minus_controls_auc"]
                    - observed["mean_identity_minus_controls_auc"]
                )
                < 1e-15,
                abs(
                    saved["identity_minus_controls_exact_one_sided_p"]
                    - observed["identity_minus_controls_exact_one_sided_p"]
                )
                < 1e-15,
                saved["participants_positive_topology_advantage"]
                == observed["participants_positive_topology_advantage"],
            )
        )
        for saved, observed in zip(decision["budget_metrics"], expected_budget)
    )
    gain_slopes = np.asarray([slope(row) for row in gains])
    topology_slopes = np.asarray([slope(row) for row in topology])
    expected_predictions = {
        "AKB_gain_density_slope_gt_0": {
            "mean_slope_auc_per_doubling": float(np.mean(gain_slopes)),
            "exact_one_sided_p": exact(gain_slopes),
            "participants_positive": int(np.sum(gain_slopes > 0)),
        },
        "topology_advantage_density_slope_gt_0": {
            "mean_slope_auc_per_doubling": float(np.mean(topology_slopes)),
            "exact_one_sided_p": exact(topology_slopes),
            "participants_positive": int(np.sum(topology_slopes > 0)),
        },
        "group_mean_gain_monotone_nondecreasing": bool(
            np.all(np.diff(np.mean(gains, axis=0)) >= -1e-15)
        ),
        "group_mean_topology_monotone_nondecreasing": bool(
            np.all(np.diff(np.mean(topology, axis=0)) >= -1e-15)
        ),
    }
    checks["predictions"] = (
        decision["frozen_predictions"] == expected_predictions
    )
    status = "pass" if all(bool(value) for value in checks.values()) else "fail"
    payload = {
        "schema": "KITE-SAT-sensor-density-mechanism-outcome-audit-v1",
        "status": status,
        "sensor_density_implementation_lock_sha256": lock_hash,
        "sensor_density_decision_sha256": sha256(DECISION),
        "checks": checks,
        "recomputed_budget_metrics": expected_budget,
        "recomputed_predictions": expected_predictions,
        "subject_artifact_sha256": hashes,
    }
    OUTPUT.write_text(json.dumps(payload, indent=2) + "\n", encoding="utf-8")
    observed = sha256(OUTPUT)
    OUTPUT.with_name(OUTPUT.name + ".sha256").write_text(
        f"{observed}  {OUTPUT.name}\n", encoding="ascii"
    )
    print(json.dumps({"status": status, "sha256": observed}))
    if status != "pass":
        raise SystemExit(1)


if __name__ == "__main__":
    main()
```
