## Supplementary material for "Falsifiable substitution tests reveal task-structured neural evidence for auditory attention": Article_Analysis_Source_Code: audit_kul_confirmation.py

```
"""Outcome-free integrity audit of the unsealed KUL even-ID partition."""

from __future__ import annotations

import argparse
import hashlib
import json
from collections import Counter
from pathlib import Path

import numpy as np
from scipy.io import loadmat

from audit_kul_discovery import base_part, clean_stimulus_name


CONFIRMATION_SUBJECTS = (2, 4, 6, 8, 10, 12, 14, 16)
EXPECTED_MD5 = {
    2: "69df6e89c29f5efd44c92bbc51a98359",
    4: "dfe3268d21838ad7d773aa4558439a1e",
    6: "da7f4321266a5cf85067a04568e01f16",
    8: "bc926d599ce90f2e38d71b7a4bf72076",
    10: "fffc929b1f71a9feb0aabc85504cd990",
    12: "d76e8526d83cb7f15420dd670957e0e2",
    14: "7767b3cc8ee58f55abc4f30fd0bf2214",
    16: "8b244536d002ab35b72f5ed6664946f0",
}
EXPECTED_LOCK_SHA256 = (
    "0e152a85fc86620d325a2414c683049ede0ccac0a29ab536bd73c899885886fc"
)


def digest(path: Path, algorithm: str) -> str:
    value = hashlib.new(algorithm)
    with path.open("rb") as handle:
        while block := handle.read(8 * 1024 * 1024):
            value.update(block)
    return value.hexdigest()


def main() -> None:
    parser = argparse.ArgumentParser()
    parser.add_argument(
        "--data-root",
        type=Path,
        default=Path("data/raw/kul_aad_confirmation"),
    )
    parser.add_argument(
        "--stimulus-root",
        type=Path,
        default=Path("data/raw/kul_aad/stimuli_extracted/stimuli"),
    )
    parser.add_argument(
        "--lock-file",
        type=Path,
        default=Path("docs/kul_candidate_lock.json"),
    )
    parser.add_argument(
        "--output",
        type=Path,
        default=Path(
            "results/kul_aad/confirmation_integrity_audit.json"
        ),
    )
    args = parser.parse_args()
    lock_hash = digest(args.lock_file, "sha256")
    if lock_hash != EXPECTED_LOCK_SHA256:
        raise RuntimeError(f"KUL candidate lock mismatch: {lock_hash}")
    observed_files = {
        int(path.stem[1:]) for path in args.data_root.glob("S*.mat")
    }
    if observed_files != set(CONFIRMATION_SUBJECTS):
        raise RuntimeError(
            f"Unexpected confirmation participants: {observed_files}"
        )

    hashes = {}
    rows = []
    global_valid = True
    for subject in CONFIRMATION_SUBJECTS:
        path = args.data_root / f"S{subject}.mat"
        observed_md5 = digest(path, "md5")
        hashes[f"S{subject}.mat"] = {
            "expected_md5": EXPECTED_MD5[subject],
            "observed_md5": observed_md5,
            "matches": observed_md5 == EXPECTED_MD5[subject],
            "bytes": path.stat().st_size,
        }
        global_valid &= observed_md5 == EXPECTED_MD5[subject]
        trials = loadmat(
            path, simplify_cells=True, variable_names=["trials"]
        )["trials"]
        if len(trials) != 20:
            raise RuntimeError(f"S{subject}: expected 20 trials")
        for index, trial in enumerate(trials, start=1):
            eeg = np.asarray(trial["RawData"]["EegData"])
            stimuli = [
                clean_stimulus_name(value)
                for value in np.atleast_1d(trial["stimuli"])
            ]
            attended_index = (
                0 if str(trial["attended_ear"]).upper() == "L" else 1
            )
            valid = bool(
                int(trial["TrialID"]) == index
                and str(trial["subject"]) == f"S{subject}"
                and int(trial["FileHeader"]["SampleRate"]) == 128
                and eeg.ndim == 2
                and eeg.shape[1] == 64
                and np.isfinite(eeg).all()
                and all(
                    (args.stimulus_root / name).is_file()
                    for name in stimuli
                )
            )
            global_valid &= valid
            rows.append(
                {
                    "subject": subject,
                    "trial": index,
                    "base_part_identity_group": base_part(trial),
                    "repetition": bool(trial["repetition"]),
                    "condition": str(trial["condition"]),
                    "attended_ear": str(trial["attended_ear"]),
                    "attended_audio": stimuli[attended_index],
                    "unattended_audio": stimuli[1 - attended_index],
                    "eeg_samples": int(eeg.shape[0]),
                    "valid": valid,
                }
            )
        del trials
    group_counts = Counter(
        (row["subject"], row["base_part_identity_group"]) for row in rows
    )
    grouping_valid = all(value == 5 for value in group_counts.values())
    global_valid &= grouping_valid
    report = {
        "dataset": "KUL auditory attention dataset",
        "partition": "even-ID one-shot confirmation",
        "candidate_lock_sha256": lock_hash,
        "participants": list(CONFIRMATION_SUBJECTS),
        "discovery_participants_loaded_by_this_script": [],
        "file_hashes": hashes,
        "summary": {
            "participant_count": len(CONFIRMATION_SUBJECTS),
            "trial_count": len(rows),
            "all_hashes_match": all(
                value["matches"] for value in hashes.values()
            ),
            "all_trials_valid": all(row["valid"] for row in rows),
            "five_trials_per_identity_group": grouping_valid,
            "condition_counts": dict(
                Counter(row["condition"] for row in rows)
            ),
            "attention_ear_counts": dict(
                Counter(row["attended_ear"] for row in rows)
            ),
            "global_integrity_pass": bool(global_valid),
        },
        "trials": rows,
    }
    args.output.parent.mkdir(parents=True, exist_ok=True)
    args.output.write_text(json.dumps(report, indent=2), encoding="utf-8")
    print(json.dumps(report["summary"], indent=2))
    if not global_valid:
        raise RuntimeError("KUL confirmation integrity audit failed")


if __name__ == "__main__":
    main()
```
