## Supplementary material for "Falsifiable substitution tests reveal task-structured neural evidence for auditory attention": Article_Analysis_Source_Code: audit_kul_discovery.py

```
"""Outcome-free integrity audit for the sealed KUL discovery partition.

Only odd-numbered participants are permitted.  The audit verifies the official
file hashes, MATLAB trial structure, exact audio-file mapping, EEG/audio
lengths, and the four base-part identity groups used for leakage-safe
cross-validation.  It deliberately computes no decoder or attention outcome.
"""

from __future__ import annotations

import argparse
import hashlib
import json
import re
from collections import Counter
from pathlib import Path

import numpy as np
from scipy.io import loadmat, wavfile


DISCOVERY_SUBJECTS = (1, 3, 5, 7, 9, 11, 13, 15)
EXPECTED_MD5 = {
    "S1.mat": "81a4ea96dc63f6acffb16405e071f27e",
    "S3.mat": "2fb8067dc26a5420377be480fdb80dc5",
    "S5.mat": "025953727a5b07797263210c34589700",
    "S7.mat": "0ae742f02b154897fd44a133e0aff30c",
    "S9.mat": "f12716937c3d53bbdcedb1364018b2f7",
    "S11.mat": "cbe8aaf6e7075f07746e644f5adefedb",
    "S13.mat": "9c277eceb8070f58c8c4655918eb3beb",
    "S15.mat": "d2d70041bd6317416e1c5266426e08d4",
    "stimuli.zip": "bfc4309b35387dfb2552de288c7c0b13",
    "README.txt.txt": "646f4117f725a8ce9e3ac27a6036f864",
    "preprocess_data.m": "51e257fa5a15850776e985847932d172",
}


def md5(path: Path) -> str:
    digest = hashlib.md5()
    with path.open("rb") as handle:
        while block := handle.read(8 * 1024 * 1024):
            digest.update(block)
    return digest.hexdigest()


def base_part(trial: dict[str, object]) -> int:
    part = int(trial["part"])
    return (part - 1) % 4 + 1


def clean_stimulus_name(value: object) -> str:
    name = str(value)
    return re.sub(r"_(?:hrtf|dry)\.wav$", "_dry.wav", name)


def main() -> None:
    parser = argparse.ArgumentParser()
    parser.add_argument(
        "--data-root", type=Path, default=Path("data/raw/kul_aad")
    )
    parser.add_argument(
        "--output",
        type=Path,
        default=Path("results/kul_aad/discovery_integrity_audit.json"),
    )
    args = parser.parse_args()

    observed_subject_files = {
        int(match.group(1))
        for path in args.data_root.glob("S*.mat")
        if (match := re.fullmatch(r"S(\d+)\.mat", path.name))
    }
    if observed_subject_files != set(DISCOVERY_SUBJECTS):
        raise RuntimeError(
            "KUL firewall violation or incomplete discovery partition: "
            f"observed={sorted(observed_subject_files)}"
        )

    hashes = {}
    for name, expected in EXPECTED_MD5.items():
        path = args.data_root / name
        actual = md5(path)
        hashes[name] = {
            "expected_md5": expected,
            "observed_md5": actual,
            "matches": actual == expected,
            "bytes": path.stat().st_size,
        }

    stimulus_root = args.data_root / "stimuli_extracted" / "stimuli"
    audio_metadata = {}
    for path in sorted(stimulus_root.glob("*_dry.wav")):
        sample_rate, audio = wavfile.read(path, mmap=True)
        audio_metadata[path.name] = {
            "sample_rate_hz": int(sample_rate),
            "samples": int(audio.shape[0]),
            "duration_seconds": float(audio.shape[0] / sample_rate),
            "channels": 1 if audio.ndim == 1 else int(audio.shape[1]),
            "dtype": str(audio.dtype),
        }

    rows = []
    global_valid = all(value["matches"] for value in hashes.values())
    for subject in DISCOVERY_SUBJECTS:
        path = args.data_root / f"S{subject}.mat"
        trials = loadmat(
            path, simplify_cells=True, variable_names=["trials"]
        )["trials"]
        if len(trials) != 20:
            raise RuntimeError(f"S{subject}: expected 20 trials")
        for index, trial in enumerate(trials, start=1):
            eeg = np.asarray(trial["RawData"]["EegData"])
            sample_rate = int(trial["FileHeader"]["SampleRate"])
            original_stimuli = [
                str(value) for value in np.atleast_1d(trial["stimuli"])
            ]
            clean_stimuli = [
                clean_stimulus_name(value) for value in original_stimuli
            ]
            attended_index = 0 if str(trial["attended_ear"]).upper() == "L" else 1
            unattended_index = 1 - attended_index
            audio_lengths = [
                audio_metadata[name]["samples"]
                / audio_metadata[name]["sample_rate_hz"]
                for name in clean_stimuli
            ]
            duration = eeg.shape[0] / sample_rate
            group = base_part(trial)
            filename_parts = [
                int(re.search(r"part(\d+)", name).group(1))
                for name in clean_stimuli
            ]
            row_valid = bool(
                int(trial["TrialID"]) == index
                and str(trial["subject"]) == f"S{subject}"
                and sample_rate == 128
                and eeg.ndim == 2
                and eeg.shape[1] == 64
                and np.isfinite(eeg).all()
                and len(set(filename_parts)) == 1
                and filename_parts[0] == group
                and all(name in audio_metadata for name in clean_stimuli)
                and all(value >= duration for value in audio_lengths)
            )
            global_valid &= row_valid
            rows.append(
                {
                    "subject": subject,
                    "trial": index,
                    "experiment": int(trial["experiment"]),
                    "part": int(trial["part"]),
                    "base_part_identity_group": group,
                    "repetition": bool(trial["repetition"]),
                    "condition": str(trial["condition"]),
                    "attended_ear": str(trial["attended_ear"]),
                    "attended_track": int(trial["attended_track"]),
                    "original_left_stimulus": original_stimuli[0],
                    "original_right_stimulus": original_stimuli[1],
                    "clean_left_stimulus": clean_stimuli[0],
                    "clean_right_stimulus": clean_stimuli[1],
                    "attended_clean_stimulus": clean_stimuli[attended_index],
                    "unattended_clean_stimulus": clean_stimuli[
                        unattended_index
                    ],
                    "eeg_samples": int(eeg.shape[0]),
                    "eeg_channels": int(eeg.shape[1]),
                    "sample_rate_hz": sample_rate,
                    "duration_seconds": duration,
                    "all_eeg_values_finite": bool(np.isfinite(eeg).all()),
                    "valid": row_valid,
                }
            )
        del trials

    group_counts = Counter(
        (row["subject"], row["base_part_identity_group"]) for row in rows
    )
    expected_grouping = all(value == 5 for value in group_counts.values())
    global_valid &= expected_grouping
    report = {
        "dataset": "KUL auditory attention dataset",
        "zenodo_record": "https://zenodo.org/records/4004271",
        "partition": "odd-participant discovery only",
        "participants": list(DISCOVERY_SUBJECTS),
        "confirmation_participants_loaded": [],
        "file_hashes": hashes,
        "audio_files": audio_metadata,
        "summary": {
            "participant_count": len(DISCOVERY_SUBJECTS),
            "trial_count": len(rows),
            "base_part_identity_groups_per_participant": 4,
            "trials_per_participant_identity_group": sorted(
                set(group_counts.values())
            ),
            "condition_counts": dict(
                sorted(Counter(row["condition"] for row in rows).items())
            ),
            "attention_ear_counts": dict(
                sorted(Counter(row["attended_ear"] for row in rows).items())
            ),
            "all_hashes_match": all(
                value["matches"] for value in hashes.values()
            ),
            "all_trials_valid": all(row["valid"] for row in rows),
            "identity_grouping_is_5_trials_per_group": expected_grouping,
            "global_integrity_pass": bool(global_valid),
        },
        "identity_firewall": {
            "group_definition": "((trial.part - 1) mod 4) + 1",
            "rule": (
                "Full-length trials, two-minute excerpts, presentation "
                "conditions, attention assignments, and all repeated "
                "presentations sharing a base part remain in one fold."
            ),
        },
        "trials": rows,
    }
    args.output.parent.mkdir(parents=True, exist_ok=True)
    args.output.write_text(json.dumps(report, indent=2), encoding="utf-8")
    print(json.dumps(report["summary"], indent=2))
    if not global_valid:
        raise RuntimeError("KUL discovery integrity audit failed")


if __name__ == "__main__":
    main()
```
