## Supplementary material for "Falsifiable substitution tests reveal task-structured neural evidence for auditory attention": Article_Analysis_Source_Code: audit_wz_hic_continuous_listener_transport_artifacts_v1.py

```
#!/usr/bin/env python3
"""Independent aggregation and hash audit of continuous-listener transport v1."""

from __future__ import annotations

import csv
import hashlib
import itertools
import json
from collections import defaultdict
from pathlib import Path

import numpy as np


ROOT = Path(__file__).resolve().parents[1]
RESULT_ROOT = (
    ROOT / "results" / "wz_hic_cross_dataset"
    / "continuous_listener_transport_v1"
)
DECISION = RESULT_ROOT / "continuous_listener_transport_decision_v1.json"
PARTICIPANTS = RESULT_ROOT / "participant_endpoints_v1.csv"
DIRECTIONS = RESULT_ROOT / "direction_endpoints_v1.csv"
OUTPUT = RESULT_ROOT / "independent_artifact_audit_v1.json"
DATASETS = (
    "KITE_ST_confirmation_complete_case_v2",
    "KUL_64ch",
    "Ear_SAAD_29ch",
)
ENDPOINTS = (
    "F1_fingerprint_listener_rank_advantage",
    "F2_fingerprint_physical_mapping_rank_advantage",
    "P1_predictive_listener_rank_advantage",
    "P2_predictive_physical_mapping_rank_advantage",
)
BOOTSTRAP_DRAWS = 100_000
BOOTSTRAP_SEED = 2026073001
RANDOMIZATION_DRAWS = 1_000_000
RANDOMIZATION_SEED = 2026073002


def sha256(path: Path) -> str:
    digest = hashlib.sha256()
    with path.open("rb") as handle:
        for block in iter(lambda: handle.read(8 * 1024 * 1024), b""):
            digest.update(block)
    return digest.hexdigest()


def sidecar_ok(path: Path) -> bool:
    fields = path.with_name(path.name + ".sha256").read_text(
        encoding="ascii"
    ).strip().split()
    return len(fields) == 2 and fields[0] == sha256(path) and fields[1] == path.name


def read_csv(path: Path) -> list[dict]:
    with path.open("r", encoding="utf-8", newline="") as handle:
        return list(csv.DictReader(handle))


def exact_sign_flip(values: np.ndarray) -> float:
    values = np.asarray(values, dtype=np.float64)
    observed = float(np.mean(values))
    null = []
    for signs in itertools.product((-1.0, 1.0), repeat=len(values)):
        null.append(float(np.mean(values * np.asarray(signs))))
    return float(np.mean(np.asarray(null) >= observed - 1e-15))


def dataset_summary(values: np.ndarray, seed: int) -> dict:
    values = np.asarray(values, dtype=np.float64)
    rng = np.random.default_rng(seed)
    samples = values[
        rng.integers(0, len(values), size=(BOOTSTRAP_DRAWS, len(values)))
    ]
    boot = np.mean(samples, axis=1)
    return {
        "n": len(values),
        "mean": float(np.mean(values)),
        "bootstrap_95": [
            float(np.quantile(boot, 0.025)),
            float(np.quantile(boot, 0.975)),
        ],
        "exact_one_sided_signflip_p": exact_sign_flip(values),
        "positive_participants": int(np.sum(values > 0)),
        "zero_participants": int(np.sum(values == 0)),
    }


def cross_summary(
    arrays: list[np.ndarray], bootstrap_seed: int, randomization_seed: int
) -> dict:
    observed = float(np.mean([np.mean(values) for values in arrays]))
    rng = np.random.default_rng(bootstrap_seed)
    boot = []
    for values in arrays:
        indices = rng.integers(
            0, len(values), size=(BOOTSTRAP_DRAWS, len(values))
        )
        boot.append(np.mean(values[indices], axis=1))
    bootstrap = np.mean(np.row_stack(boot), axis=0)
    rng = np.random.default_rng(randomization_seed)
    extreme = 0
    remaining = RANDOMIZATION_DRAWS
    while remaining:
        current = min(10_000, remaining)
        null = np.zeros(current, dtype=np.float64)
        for values in arrays:
            signs = rng.integers(
                0, 2, size=(current, len(values)), dtype=np.int8
            )
            signs = signs.astype(np.float64) * 2.0 - 1.0
            null += (signs @ values) / (len(arrays) * len(values))
        extreme += int(np.sum(null >= observed - 1e-15))
        remaining -= current
    return {
        "dataset_count": len(arrays),
        "participant_count": int(sum(len(values) for values in arrays)),
        "equal_dataset_mean": observed,
        "stratified_bootstrap_95": [
            float(np.quantile(bootstrap, 0.025)),
            float(np.quantile(bootstrap, 0.975)),
        ],
        "participant_signflip_randomization_p": float(
            (1 + extreme) / (1 + RANDOMIZATION_DRAWS)
        ),
        "bootstrap_draws": BOOTSTRAP_DRAWS,
        "randomization_draws": RANDOMIZATION_DRAWS,
    }


def maximum_error(left, right) -> float:
    if isinstance(left, dict):
        return max(
            [maximum_error(left[key], right[key]) for key in left] or [0.0]
        )
    if isinstance(left, list):
        return max(
            [maximum_error(a, b) for a, b in zip(left, right)] or [0.0]
        )
    if isinstance(left, (int, float)) and not isinstance(left, bool):
        return abs(float(left) - float(right))
    return 0.0 if left == right else float("inf")


def main() -> None:
    if OUTPUT.exists():
        raise RuntimeError("Refusing to overwrite independent artifact audit.")
    decision = json.loads(DECISION.read_text(encoding="utf-8"))
    participant_rows = read_csv(PARTICIPANTS)
    direction_rows = read_csv(DIRECTIONS)

    by_person = defaultdict(list)
    for row in direction_rows:
        by_person[(row["dataset"], row["subject"])].append(row)
    endpoint_sources = {
        ENDPOINTS[0]: "fingerprint_listener_rank_probability",
        ENDPOINTS[1]: "fingerprint_physical_mapping_rank_probability",
        ENDPOINTS[2]: "predictive_listener_rank_probability",
        ENDPOINTS[3]: "predictive_physical_mapping_rank_probability",
    }
    participant_errors = []
    for row in participant_rows:
        directions = by_person[(row["dataset"], row["subject"])]
        if len(directions) != 2:
            participant_errors.append(float("inf"))
            continue
        for endpoint, source in endpoint_sources.items():
            expected = np.mean([float(value[source]) for value in directions]) - 0.5
            participant_errors.append(abs(float(row[endpoint]) - expected))

    summaries = {}
    crosses = {}
    arrays_by_endpoint = {}
    for endpoint_index, endpoint in enumerate(ENDPOINTS):
        summaries[endpoint] = {}
        arrays = []
        for dataset_index, dataset in enumerate(DATASETS):
            values = np.asarray(
                [
                    float(row[endpoint])
                    for row in participant_rows
                    if row["dataset"] == dataset
                ]
            )
            arrays.append(values)
            summaries[endpoint][dataset] = dataset_summary(
                values,
                BOOTSTRAP_SEED + 100 * endpoint_index + dataset_index,
            )
        cross = cross_summary(
            arrays,
            BOOTSTRAP_SEED + 10_000 * endpoint_index,
            RANDOMIZATION_SEED + 10_000 * endpoint_index,
        )
        cross["positive_dataset_means"] = int(
            sum(summaries[endpoint][dataset]["mean"] > 0 for dataset in DATASETS)
        )
        cross["datasets_with_bootstrap_upper_below_zero"] = [
            dataset
            for dataset in DATASETS
            if summaries[endpoint][dataset]["bootstrap_95"][1] < 0
        ]
        crosses[endpoint] = cross
        arrays_by_endpoint[endpoint] = arrays

    def gate(endpoint: str) -> dict:
        row = crosses[endpoint]
        checks = {
            "equal_dataset_mean_gt_zero": row["equal_dataset_mean"] > 0,
            "bootstrap_lower_gt_zero": row["stratified_bootstrap_95"][0] > 0,
            "randomization_p_le_0_05": (
                row["participant_signflip_randomization_p"] <= 0.05
            ),
            "at_least_two_positive_dataset_means": (
                row["positive_dataset_means"] >= 2
            ),
            "no_dataset_bootstrap_upper_below_zero": not row[
                "datasets_with_bootstrap_upper_below_zero"
            ],
        }
        return {"pass": all(checks.values()), "checks": checks}

    scientific_gates = {
        "C1_fingerprint_listener": gate(ENDPOINTS[0]),
        "C2_fingerprint_mapping": gate(ENDPOINTS[1]),
        "C3_predictive_listener": gate(ENDPOINTS[2]),
        "C4_predictive_mapping": gate(ENDPOINTS[3]),
    }
    frozen_input_errors = []
    for record in decision["frozen_inputs"]:
        frozen_input_errors.append(
            0.0
            if sha256(ROOT / record["path"]) == record["sha256"]
            else float("inf")
        )
    declared_hash_errors = [
        0.0
        if sha256(ROOT / decision[name]["path"]) == decision[name]["sha256"]
        else float("inf")
        for name in ("protocol", "implementation_lock", "runner")
    ]
    checks = {
        "result_sidecar": sidecar_ok(DECISION),
        "participant_sidecar": sidecar_ok(PARTICIPANTS),
        "direction_sidecar": sidecar_ok(DIRECTIONS),
        "participant_rows_43": len(participant_rows) == 43,
        "direction_rows_86": len(direction_rows) == 86,
        "two_directions_each": (
            len(by_person) == 43
            and all(len(rows) == 2 for rows in by_person.values())
        ),
        "participant_reconstruction_max_abs_error": max(participant_errors),
        "dataset_summary_max_abs_error": maximum_error(
            summaries, decision["dataset_summaries"]
        ),
        "cross_summary_max_abs_error": maximum_error(
            crosses, decision["cross_dataset_summaries"]
        ),
        "scientific_gates_exact": all(
            decision["gates"][name] == value
            for name, value in scientific_gates.items()
        ),
        "all_frozen_inputs_match": max(frozen_input_errors or [0.0]) == 0.0,
        "declared_protocol_lock_runner_match": (
            max(declared_hash_errors) == 0.0
        ),
        "conjunction_exact": (
            decision["conjunction_pass"]
            == (
                decision["gates"]["C0_integrity"]["pass"]
                and all(value["pass"] for value in scientific_gates.values())
            )
        ),
    }
    tolerance = 1e-15
    passed = (
        all(
            value is True
            for key, value in checks.items()
            if not key.endswith("_error")
            and "max_abs_error" not in key
        )
        and checks["participant_reconstruction_max_abs_error"] <= tolerance
        and checks["dataset_summary_max_abs_error"] <= tolerance
        and checks["cross_summary_max_abs_error"] <= tolerance
    )
    output = {
        "schema": "WZ-HIC-continuous-listener-transport-independent-artifact-audit-v1",
        "status": "pass" if passed else "fail",
        "auditor": {
            "path": Path(__file__).relative_to(ROOT).as_posix(),
            "sha256": sha256(Path(__file__)),
            "does_not_import_project_runner": True,
        },
        "source_result": {
            "path": DECISION.relative_to(ROOT).as_posix(),
            "sha256": sha256(DECISION),
        },
        "checks": checks,
        "recomputed_scientific_gates": scientific_gates,
        "tolerance": tolerance,
    }
    with OUTPUT.open("x", encoding="utf-8") as handle:
        json.dump(output, handle, ensure_ascii=False, indent=2, allow_nan=False)
        handle.write("\n")
    sidecar = OUTPUT.with_name(OUTPUT.name + ".sha256")
    with sidecar.open("x", encoding="ascii") as handle:
        handle.write(f"{sha256(OUTPUT)}  {OUTPUT.name}\n")
    print(json.dumps(output, indent=2), flush=True)


if __name__ == "__main__":
    main()
```
