## Supplementary material for "Falsifiable substitution tests reveal task-structured neural evidence for auditory attention": Article_Analysis_Source_Code: audit_wz_hic_continuous_listener_transport_source_v1.py

```
#!/usr/bin/env python3
"""Independent source-level recomputation of continuous listener transport.

This auditor does not import either continuous-listener runner.  It rebuilds
the two source profiles and the cross-partition target contributions for all
43 participants from the frozen public-data derivatives, then independently
reconstructs the participant endpoints stored by the primary implementation.
"""

from __future__ import annotations

import csv
import hashlib
import itertools
import json
from pathlib import Path

import numpy as np
from scipy.io import loadmat
from scipy.signal import resample_poly
from scipy.stats import rankdata

from kite_kul_akb_v1 import biosemi64_geometry, fit_ridge, make_windows
from kite_sat_adaptive_broad_v1 import family_weights, reliability
from kite_sat_backward_v1 import block_design
from kite_sat_direction_delay_controls_v1 import TRANSFORMS, transform_vector
from kite_sat_geometry_v1 import cell_delta_contributions, partition_ids
from kite_st_hd_core_v1 import load_public_geometry
from run_kite_sat_ear_saad_external_subject_v1 import (
    BLOCKS_PER_TRIAL,
    N_CHANNELS,
    N_FEATURES,
    SAMPLES_PER_BLOCK,
    USABLE,
    block_design as ear_design,
    cell_contributions as ear_contributions,
    fit_from_blocks,
    sufficient_statistics,
)


ROOT = Path(__file__).resolve().parents[1]
RESULT_ROOT = (
    ROOT / "results" / "wz_hic_cross_dataset"
    / "continuous_listener_transport_v1"
)
PRIMARY = RESULT_ROOT / "participant_endpoints_v1.csv"
PRIMARY_DIRECTIONS = RESULT_ROOT / "direction_endpoints_v1.csv"
DECISION = RESULT_ROOT / "continuous_listener_transport_decision_v1.json"
OUTPUT = RESULT_ROOT / "independent_source_audit_v1.json"

KITE_SUBJECTS = (
    "S0", "S10", "S2", "S21", "S28", "S22",
    "S9", "S15", "S16", "S18", "S13", "S1",
)
KUL_SUBJECTS = tuple(f"S{number}" for number in range(1, 17))
EAR_SUBJECTS = tuple(f"S{number}" for number in range(1, 16))
KUL_SPLIT = {"A": (1, 4), "B": (2, 3)}


def sha256(path: Path) -> str:
    digest = hashlib.sha256()
    with path.open("rb") as handle:
        for chunk in iter(lambda: handle.read(8 * 1024 * 1024), b""):
            digest.update(chunk)
    return digest.hexdigest()


def rows(path: Path) -> list[dict]:
    with path.open("r", encoding="utf-8", newline="") as handle:
        return list(csv.DictReader(handle))


def corr(left: np.ndarray, right: np.ndarray) -> float:
    a = np.asarray(left, dtype=np.float64)
    b = np.asarray(right, dtype=np.float64)
    a -= np.mean(a)
    b -= np.mean(b)
    denominator = np.sqrt(np.sum(a * a) * np.sum(b * b))
    if denominator <= 1e-12:
        raise RuntimeError("Degenerate profile in source auditor.")
    return float(np.sum(a * b) / denominator)


def auc(scores: np.ndarray, labels: np.ndarray) -> float:
    scores = np.asarray(scores, dtype=np.float64)
    labels = np.asarray(labels, dtype=np.int64)
    positive = scores[labels == 1]
    negative = scores[labels == 0]
    joined = np.r_[positive, negative]
    ranks = rankdata(joined, method="average")
    u = np.sum(ranks[: len(positive)]) - len(positive) * (len(positive) + 1) / 2
    return float(u / (len(positive) * len(negative)))


def archive_model(saved: np.lib.npyio.NpzFile, prefix: str) -> dict:
    return {
        key: np.asarray(saved[f"{prefix}_{key}"], dtype=np.float64)
        for key in ("x_mean", "x_scale", "weights")
    }


def load_kite() -> list[dict]:
    audio_path = (
        ROOT / "data" / "derived" / "kite_st_hd_zenodo4518754"
        / "kite_sat_audio_features_v1.npz"
    )
    preprocessed_root = (
        ROOT / "data" / "derived" / "kite_st_hd_zenodo4518754"
        / "confirmation_preprocessed_v1"
    )
    result_root = (
        ROOT / "results" / "kite_sat" / "akb_confirmation_complete_case_v2"
        / "subjects"
    )
    coordinate_path = (
        ROOT / "downloads" / "candidate_audit" / "zenodo4518754_support"
        / "misc" / "misc" / "eeg255ch_locs.csv"
    )
    physical = {
        "1L": ("part1_track1", "part1_track2"),
        "1R": ("part1_track1", "part1_track2"),
        "2L": ("part2_track2", "part2_track1"),
        "2R": ("part2_track2", "part2_track1"),
    }
    with np.load(audio_path, allow_pickle=False) as saved:
        audio = {key: np.asarray(saved[key]) for key in saved.files}
    cell_ids = partition_ids(load_public_geometry(coordinate_path).xy)["S16"]
    people = []
    for subject in KITE_SUBJECTS:
        result_path = result_root / f"{subject}_AKB_confirmation_v1.json"
        result = json.loads(result_path.read_text(encoding="utf-8"))
        manifest_path = ROOT / result["upstream_manifest"]["path"]
        if sha256(manifest_path) != result["upstream_manifest"]["sha256"]:
            raise RuntimeError("KITE manifest hash mismatch.")
        model_path = ROOT / result["model_archive"]["path"]
        if sha256(model_path) != result["model_archive"]["sha256"]:
            raise RuntimeError("KITE model hash mismatch.")
        condition_rows = {}
        for condition in ("1L", "1R", "2L", "2R"):
            path = (
                preprocessed_root / subject
                / f"{subject}_AAD_{condition}_preprocessed_v1.npz"
            )
            if sha256(path) != result["preprocessed_sha256"][condition]:
                raise RuntimeError("KITE preprocessed hash mismatch.")
            with np.load(path, allow_pickle=False) as saved:
                eeg = np.asarray(saved["eeg"])
            offset = int(round(float(result["offset_seconds"][condition]) * 100))
            left_key, right_key = physical[condition]
            blocks = []
            for block_index in range(12):
                first = offset + 3000 * block_index
                stop = first + 3000
                design, left = block_design(
                    eeg[3000 * block_index : 3000 * (block_index + 1)],
                    audio[left_key][first:stop],
                )
                right = resample_poly(
                    audio[right_key][first:stop], 1, 5, axis=0
                )[:595, 0]
                blocks.append(
                    {
                        "design": design,
                        "left": left[:, 0],
                        "right": right,
                        "label": int(condition.endswith("L")),
                    }
                )
            condition_rows[condition] = blocks
        profiles = {
            "A": np.asarray(
                result["fold_reliabilities"]["train_story_1"],
                dtype=np.float64,
            ),
            "B": np.asarray(
                result["fold_reliabilities"]["train_story_2"],
                dtype=np.float64,
            ),
        }
        tests = {}
        with np.load(model_path, allow_pickle=False) as saved:
            for source, target, conditions, prefix in (
                ("A", "B", ("2L", "2R"), "train_story_1"),
                ("B", "A", ("1L", "1R"), "train_story_2"),
            ):
                model = archive_model(saved, prefix)
                deltas, labels = [], []
                for condition in conditions:
                    for block in condition_rows[condition]:
                        delta, _ = cell_delta_contributions(
                            block["design"],
                            model,
                            block["left"],
                            block["right"],
                            cell_ids,
                        )
                        deltas.append(delta)
                        labels.append(block["label"])
                tests[f"{source}_to_{target}"] = {
                    "source": source,
                    "target": target,
                    "delta": np.asarray(deltas),
                    "labels": np.asarray(labels),
                }
        people.append({"subject": subject, "profiles": profiles, "tests": tests})
        print(f"audit KITE {subject}", flush=True)
    return people


def kul_half_profile(windows: list, parts: tuple[int, int], ids: np.ndarray) -> np.ndarray:
    held_contributions = []
    for held_part in parts:
        train = [
            window
            for window in windows
            if window.part in parts and window.part != held_part
        ]
        model = fit_ridge(train)
        for window in windows:
            if window.part != held_part:
                continue
            delta, _ = cell_delta_contributions(
                window.design, model, window.left, window.right, ids
            )
            sign = 1.0 if window.attended_ear == "L" else -1.0
            held_contributions.append(sign * delta)
    return reliability(np.asarray(held_contributions))


def load_kul() -> list[dict]:
    odd_root = ROOT / "data" / "derived" / "kul_discovery_32hz"
    even_root = ROOT / "data" / "derived" / "kul_confirmation_32hz"
    audio_path = odd_root / "audio_powerlaw_subbands.npz"
    with np.load(audio_path, allow_pickle=False) as saved:
        audio = {
            key.removeprefix("audio::"): np.asarray(saved[key])
            for key in saved.files
            if key.startswith("audio::")
        }
    _, _, ids = biosemi64_geometry()
    people = []
    for subject_text in KUL_SUBJECTS:
        number = int(subject_text[1:])
        path = (odd_root if number % 2 else even_root) / f"S{number}.npz"
        with np.load(path, allow_pickle=False) as saved:
            metadata = json.loads(str(saved["metadata_json"]))
            eeg = [np.asarray(saved[f"eeg_{index:02d}"]) for index in range(1, 21)]
        windows = make_windows(eeg, metadata, audio)
        profiles = {
            half: kul_half_profile(windows, parts, ids)
            for half, parts in KUL_SPLIT.items()
        }
        tests = {}
        for source, target in (("A", "B"), ("B", "A")):
            model = fit_ridge(
                [window for window in windows if window.part in KUL_SPLIT[source]]
            )
            delta, labels = [], []
            for window in windows:
                if window.part not in KUL_SPLIT[target]:
                    continue
                contribution, _ = cell_delta_contributions(
                    window.design, model, window.left, window.right, ids
                )
                delta.append(contribution)
                labels.append(int(window.attended_ear == "L"))
            tests[f"{source}_to_{target}"] = {
                "source": source,
                "target": target,
                "delta": np.asarray(delta),
                "labels": np.asarray(labels),
            }
        people.append(
            {"subject": subject_text, "profiles": profiles, "tests": tests}
        )
        print(f"audit KUL {subject_text}", flush=True)
    return people


def balanced_split(attended: np.ndarray, video: np.ndarray) -> dict:
    candidates = []
    for first in itertools.combinations(range(6), 3):
        if 0 not in first:
            continue
        second = tuple(index for index in range(6) if index not in first)
        if not all(
            len(np.unique(values[list(indices)])) == 2
            for values in (attended, video)
            for indices in (first, second)
        ):
            continue
        imbalance = (
            abs(
                np.sum(attended[list(first)] == 1)
                - np.sum(attended[list(second)] == 1)
            )
            + abs(
                np.sum(video[list(first)] == 1)
                - np.sum(video[list(second)] == 1)
            )
        )
        candidates.append((int(imbalance), first, second))
    _, first, second = min(candidates)
    return {"A": first, "B": second}


def ear_half_profile(records: list[dict], trials: tuple[int, ...], ids: np.ndarray) -> np.ndarray:
    contributions = []
    for held_trial in trials:
        train_indices = [
            index
            for index, record in enumerate(records)
            if record["trial"] in trials and record["trial"] != held_trial
        ]
        model = fit_from_blocks(records, train_indices)
        for record in records:
            if record["trial"] != held_trial:
                continue
            delta, _ = ear_contributions(
                record["design"],
                model,
                record["speaker1"],
                record["speaker2"],
                ids,
            )
            contributions.append(
                (1.0 if record["label"] == 1 else -1.0) * delta
            )
    return reliability(np.asarray(contributions))


def load_ear() -> list[dict]:
    data_root = ROOT / "data" / "raw" / "ear_saad" / "preprocessedData"
    geometry_path = ROOT / "docs" / "kite_sat_ear_saad_scalp_geometry_v1.json"
    geometry = json.loads(geometry_path.read_text(encoding="utf-8"))
    sectors = np.asarray(geometry["angular_sector_id_0_to_7"], dtype=np.int64)
    lag_index = np.repeat(np.arange(6), N_CHANNELS)
    ids = np.tile(sectors, 6) + 8 * (lag_index >= 3)
    if ids.shape != (N_FEATURES,):
        raise RuntimeError("Ear geometry mismatch.")
    people = []
    for subject_text in EAR_SUBJECTS:
        number = int(subject_text[1:])
        data = loadmat(
            data_root / f"dataSubject{number}.mat", simplify_cells=True
        )
        eeg_trials = [
            np.nan_to_num(np.asarray(value, dtype=np.float64))
            for value in np.asarray(data["eegTrials"], dtype=object).ravel()
        ]
        envelope_trials = [
            np.asarray(value, dtype=np.float64)
            for value in np.asarray(data["envelopes"], dtype=object).ravel()
        ]
        attended = np.asarray(data["attSpeaker"]).ravel().astype(np.int64)
        video = np.asarray(data["videoCondition"]).ravel().astype(np.int64)
        split = balanced_split(attended, video)
        records = []
        for trial in range(6):
            for block in range(BLOCKS_PER_TRIAL):
                start = block * SAMPLES_PER_BLOCK
                stop = start + SAMPLES_PER_BLOCK
                design = ear_design(
                    eeg_trials[trial][start:stop, :N_CHANNELS]
                )
                envelopes = envelope_trials[trial][start:stop, :][:USABLE]
                target = envelopes[:, attended[trial] - 1]
                records.append(
                    {
                        "trial": trial,
                        "block": block,
                        "design": design,
                        "speaker1": envelopes[:, 0],
                        "speaker2": envelopes[:, 1],
                        "label": int(attended[trial] == 1),
                        "stats": sufficient_statistics(design, target),
                    }
                )
        profiles = {
            half: ear_half_profile(records, trials, ids)
            for half, trials in split.items()
        }
        tests = {}
        for source, target in (("A", "B"), ("B", "A")):
            train_indices = [
                index
                for index, record in enumerate(records)
                if record["trial"] in split[source]
            ]
            model = fit_from_blocks(records, train_indices)
            delta, labels = [], []
            for record in records:
                if record["trial"] not in split[target]:
                    continue
                contribution, _ = ear_contributions(
                    record["design"],
                    model,
                    record["speaker1"],
                    record["speaker2"],
                    ids,
                )
                delta.append(contribution)
                labels.append(record["label"])
            tests[f"{source}_to_{target}"] = {
                "source": source,
                "target": target,
                "delta": np.asarray(delta),
                "labels": np.asarray(labels),
            }
        people.append(
            {"subject": subject_text, "profiles": profiles, "tests": tests}
        )
        print(f"audit Ear-SAAD {subject_text}", flush=True)
    return people


def direction_metrics(index: int, people: list[dict], direction: str) -> dict:
    receiver = people[index]
    test = receiver["tests"][direction]
    source = test["source"]
    target = test["target"]
    source_profile = receiver["profiles"][source]
    target_profile = receiver["profiles"][target]
    donors = np.asarray(
        [
            person["profiles"][source]
            for donor_index, person in enumerate(people)
            if donor_index != index
        ]
    )
    same = corr(source_profile, target_profile)
    donor_corr = np.asarray([corr(profile, target_profile) for profile in donors])
    wrong_corr = np.asarray(
        [
            corr(
                transform_vector(source_profile, angular_shift, delay_swap),
                target_profile,
            )
            for angular_shift, delay_swap in TRANSFORMS
            if (angular_shift, delay_swap) != (0, 0)
        ]
    )
    fingerprint_listener = np.mean(
        (same > donor_corr) + 0.5 * (same == donor_corr)
    )
    fingerprint_mapping = np.mean(
        (same > wrong_corr) + 0.5 * (same == wrong_corr)
    )
    own_weight = family_weights(source_profile, "softmax2")
    donor_weights = np.asarray(
        [family_weights(profile, "softmax2") for profile in donors]
    )
    wrong_weights = np.asarray(
        [
            transform_vector(own_weight, angular_shift, delay_swap)
            for angular_shift, delay_swap in TRANSFORMS
            if (angular_shift, delay_swap) != (0, 0)
        ]
    )
    delta = test["delta"]
    labels = test["labels"]
    sign = np.where(labels == 1, 1.0, -1.0)
    own = sign * (delta @ own_weight)
    donor = sign[:, None] * (delta @ donor_weights.T)
    wrong = sign[:, None] * (delta @ wrong_weights.T)
    donor_scale = np.sqrt(np.mean(np.column_stack((own, donor)) ** 2, axis=1))
    wrong_scale = np.sqrt(np.mean(np.column_stack((own, wrong)) ** 2, axis=1))
    donor_scale = np.maximum(donor_scale, 1e-12)
    wrong_scale = np.maximum(wrong_scale, 1e-12)
    return {
        "same_profile_correlation": same,
        "mean_donor_profile_correlation": float(np.mean(donor_corr)),
        "mean_wrong_mapping_correlation": float(np.mean(wrong_corr)),
        "fingerprint_listener_rank_probability": float(fingerprint_listener),
        "fingerprint_physical_mapping_rank_probability": float(
            fingerprint_mapping
        ),
        "predictive_listener_rank_probability": float(
            np.mean((own[:, None] > donor) + 0.5 * (own[:, None] == donor))
        ),
        "predictive_physical_mapping_rank_probability": float(
            np.mean((own[:, None] > wrong) + 0.5 * (own[:, None] == wrong))
        ),
        "normalized_own_minus_donor_score": float(
            np.mean((own - np.mean(donor, axis=1)) / donor_scale)
        ),
        "normalized_physical_minus_wrong_score": float(
            np.mean((own - np.mean(wrong, axis=1)) / wrong_scale)
        ),
        "global_auc": auc(np.sum(delta, axis=1), labels),
        "own_profile_auc": auc(delta @ own_weight, labels),
    }


def main() -> None:
    if OUTPUT.exists():
        raise RuntimeError("Refusing to overwrite independent source audit.")
    primary_people = {
        (row["dataset"], row["subject"]): row for row in rows(PRIMARY)
    }
    primary_directions = {
        (row["dataset"], row["subject"], row["direction"]): row
        for row in rows(PRIMARY_DIRECTIONS)
    }
    maximum_participant_error = 0.0
    maximum_direction_error = 0.0
    reconstructed = 0
    direction_reconstructed = 0
    dataset_details = {}
    for dataset, loader in (
        ("KITE_ST_confirmation_complete_case_v2", load_kite),
        ("KUL_64ch", load_kul),
        ("Ear_SAAD_29ch", load_ear),
    ):
        people = loader()
        dataset_participant_error = 0.0
        dataset_direction_error = 0.0
        for index, person in enumerate(people):
            direction_rows = {
                direction: direction_metrics(index, people, direction)
                for direction in ("A_to_B", "B_to_A")
            }
            for direction, values in direction_rows.items():
                primary = primary_directions[
                    (dataset, person["subject"], direction)
                ]
                for key, value in values.items():
                    error = abs(float(primary[key]) - float(value))
                    dataset_direction_error = max(dataset_direction_error, error)
                direction_reconstructed += 1
            endpoint = {
                "F1_fingerprint_listener_rank_advantage": (
                    np.mean(
                        [
                            value["fingerprint_listener_rank_probability"]
                            for value in direction_rows.values()
                        ]
                    )
                    - 0.5
                ),
                "F2_fingerprint_physical_mapping_rank_advantage": (
                    np.mean(
                        [
                            value[
                                "fingerprint_physical_mapping_rank_probability"
                            ]
                            for value in direction_rows.values()
                        ]
                    )
                    - 0.5
                ),
                "P1_predictive_listener_rank_advantage": (
                    np.mean(
                        [
                            value["predictive_listener_rank_probability"]
                            for value in direction_rows.values()
                        ]
                    )
                    - 0.5
                ),
                "P2_predictive_physical_mapping_rank_advantage": (
                    np.mean(
                        [
                            value[
                                "predictive_physical_mapping_rank_probability"
                            ]
                            for value in direction_rows.values()
                        ]
                    )
                    - 0.5
                ),
                "same_minus_mean_donor_correlation": np.mean(
                    [
                        value["same_profile_correlation"]
                        - value["mean_donor_profile_correlation"]
                        for value in direction_rows.values()
                    ]
                ),
                "physical_minus_mean_wrong_correlation": np.mean(
                    [
                        value["same_profile_correlation"]
                        - value["mean_wrong_mapping_correlation"]
                        for value in direction_rows.values()
                    ]
                ),
                "normalized_own_minus_donor_score": np.mean(
                    [
                        value["normalized_own_minus_donor_score"]
                        for value in direction_rows.values()
                    ]
                ),
                "normalized_physical_minus_wrong_score": np.mean(
                    [
                        value["normalized_physical_minus_wrong_score"]
                        for value in direction_rows.values()
                    ]
                ),
                "global_auc": np.mean(
                    [value["global_auc"] for value in direction_rows.values()]
                ),
                "own_profile_auc": np.mean(
                    [
                        value["own_profile_auc"]
                        for value in direction_rows.values()
                    ]
                ),
            }
            primary = primary_people[(dataset, person["subject"])]
            for key, value in endpoint.items():
                error = abs(float(primary[key]) - float(value))
                dataset_participant_error = max(
                    dataset_participant_error, error
                )
            reconstructed += 1
        maximum_participant_error = max(
            maximum_participant_error, dataset_participant_error
        )
        maximum_direction_error = max(
            maximum_direction_error, dataset_direction_error
        )
        dataset_details[dataset] = {
            "participants": len(people),
            "maximum_participant_endpoint_absolute_error": (
                dataset_participant_error
            ),
            "maximum_direction_endpoint_absolute_error": (
                dataset_direction_error
            ),
        }
    tolerance = 1e-12
    passed = (
        reconstructed == 43
        and direction_reconstructed == 86
        and maximum_participant_error <= tolerance
        and maximum_direction_error <= tolerance
    )
    output = {
        "schema": "WZ-HIC-continuous-listener-transport-independent-source-audit-v1",
        "status": "pass" if passed else "fail",
        "auditor": {
            "path": Path(__file__).relative_to(ROOT).as_posix(),
            "sha256": sha256(Path(__file__)),
            "imports_continuous_listener_runner": False,
            "recomputes_profiles_and_target_contributions_from_source": True,
        },
        "source_decision": {
            "path": DECISION.relative_to(ROOT).as_posix(),
            "sha256": sha256(DECISION),
        },
        "participant_count": reconstructed,
        "direction_count": direction_reconstructed,
        "maximum_participant_endpoint_absolute_error": maximum_participant_error,
        "maximum_direction_endpoint_absolute_error": maximum_direction_error,
        "dataset_details": dataset_details,
        "tolerance": tolerance,
    }
    with OUTPUT.open("x", encoding="utf-8") as handle:
        json.dump(output, handle, ensure_ascii=False, indent=2, allow_nan=False)
        handle.write("\n")
    with OUTPUT.with_name(OUTPUT.name + ".sha256").open(
        "x", encoding="ascii"
    ) as handle:
        handle.write(f"{sha256(OUTPUT)}  {OUTPUT.name}\n")
    print(json.dumps(output, indent=2), flush=True)


if __name__ == "__main__":
    main()
```
