## Supplementary material for "Falsifiable substitution tests reveal task-structured neural evidence for auditory attention": Article_Analysis_Source_Code: audit_wz_hic_cross_dataset_command_identity_residual_v1.py

```
#!/usr/bin/env python3
"""Independent audit of cross-dataset command-identity residual control."""

from __future__ import annotations

import hashlib
import itertools
import json
from pathlib import Path

import numpy as np

from audit_wz_hic_asme30_crossphase_individual_geometry_v1 import (
    independent_lda_scores,
)


ROOT = Path(__file__).resolve().parents[1]
FEATURE_ASME = (
    ROOT / "data" / "derived" / "asme30_wz_hic_dataset_native_features_v2.npz"
)
FEATURE_KOJIMA = (
    ROOT / "data" / "derived" / "kojima2024b_wz_hic_confirmation_features_v1.npz"
)
PROTOCOL = (
    ROOT
    / "docs"
    / "wz_hic_cross_dataset_command_identity_residual_protocol_v1.json"
)
LOCK = (
    ROOT
    / "docs"
    / "wz_hic_cross_dataset_command_identity_residual_implementation_lock_v1.json"
)
REPRODUCIBILITY_AMENDMENT = (
    ROOT
    / "docs"
    / "wz_hic_cross_dataset_command_identity_residual_reproducibility_amendment_v1.json"
)
RESULT_DIR = (
    ROOT
    / "results"
    / "wz_hic_cross_dataset"
    / "command_identity_residual_v1"
)
RESULT = RESULT_DIR / "residual_decision_v1.json"
AUDIT = RESULT_DIR / "independent_source_audit_v1.json"
INDEPENDENT_LDA_SOURCE = (
    ROOT
    / "analysis"
    / "audit_wz_hic_asme30_crossphase_individual_geometry_v1.py"
)

ASME_PHYSICAL = (
    (17, 23, 5, 18, 20, 25, 21, 9, 15, 16),
    (1, 19, 4, 6, 7, 8, 10, 11, 12, 29),
    (26, 24, 3, 22, 2, 14, 13, 27, 28, 30),
)
ASME_WRONG = (
    (
        tuple(range(1, 31, 3)),
        tuple(range(2, 31, 3)),
        tuple(range(3, 31, 3)),
    ),
    (
        tuple(range(1, 11)),
        tuple(range(11, 21)),
        tuple(range(21, 31)),
    ),
)
KOJIMA_PHYSICAL = ((1, 2), (3, 4))
KOJIMA_WRONG = (((1, 3), (2, 4)), ((1, 4), (2, 3)))
KOJIMA_SUBJECTS = tuple("BDFHJLN")
BOOTSTRAP_SEED = 20260730
BOOTSTRAP_N = 20_000
TOLERANCE = 5e-7


def sha256(path: Path) -> str:
    digest = hashlib.sha256()
    with path.open("rb") as handle:
        for block in iter(lambda: handle.read(1024 * 1024), b""):
            digest.update(block)
    return digest.hexdigest()


def semantic_json_sha256(path: Path) -> str:
    """Hash result content after removing only rerun-time metadata."""

    def clean(value):
        if isinstance(value, dict):
            return {
                key: clean(item)
                for key, item in value.items()
                if key != "created_utc"
            }
        if isinstance(value, list):
            return [clean(item) for item in value]
        return value

    payload = json.loads(path.read_text(encoding="utf-8"))
    canonical = json.dumps(
        clean(payload), sort_keys=True, separators=(",", ":"), allow_nan=True
    ).encode("utf-8")
    return hashlib.sha256(canonical).hexdigest()


def make_labels(
    command: np.ndarray, target: np.ndarray, groups, n_commands: int
) -> np.ndarray:
    lookup = np.full(n_commands + 1, -1, dtype=int)
    for index, group in enumerate(groups):
        lookup[list(group)] = index
    result = np.full(len(command), "U", dtype="U1")
    target_mask = command == target
    result[target_mask] = "T"
    result[(lookup[command] == lookup[target]) & ~target_mask] = "S"
    return result


def residualize(
    train_x: np.ndarray,
    test_x: np.ndarray,
    train_command: np.ndarray,
    train_target: np.ndarray,
    test_command: np.ndarray,
    n_commands: int,
) -> tuple[np.ndarray, np.ndarray]:
    train_result = np.asarray(train_x, dtype=float).copy()
    test_result = np.asarray(test_x, dtype=float).copy()
    non_target = train_command != train_target
    for value in range(1, n_commands + 1):
        baseline = (train_command == value) & non_target
        command_mean = np.mean(train_x[baseline], axis=0)
        train_result[train_command == value] -= command_mean
        test_result[test_command == value] -= command_mean
    return train_result, test_result


def selections(
    command: np.ndarray, trial: np.ndarray, n_commands: int
) -> list[np.ndarray]:
    output = []
    for trial_id in sorted(int(value) for value in np.unique(trial)):
        ids = np.flatnonzero(trial == trial_id)
        chunks = []
        for value in range(1, n_commands + 1):
            value_ids = ids[command[ids] == value]
            if value_ids.size < 3:
                chunks = []
                break
            chunks.append(value_ids[:3])
        if len(chunks) == n_commands:
            output.append(np.concatenate(chunks))
    return output


def effects(
    score: np.ndarray,
    command: np.ndarray,
    target: np.ndarray,
    selected_trials: list[np.ndarray],
    groups,
    n_commands: int,
) -> list[float]:
    output = []
    for selected in selected_trials:
        labels = make_labels(
            command[selected], target[selected], groups, n_commands
        )
        output.append(
            float(score[selected][labels == "S"].mean())
            - float(score[selected][labels == "U"].mean())
        )
    return output


def cosine(left: np.ndarray, right: np.ndarray) -> float:
    return float(
        np.vdot(left, right)
        / (np.linalg.norm(left) * np.linalg.norm(right))
    )


def direction(
    train_x: np.ndarray,
    train_labels: np.ndarray,
    test_x: np.ndarray,
    command: np.ndarray,
    target: np.ndarray,
    selected_trials: list[np.ndarray],
    groups,
    n_commands: int,
) -> tuple[np.ndarray, np.ndarray, float]:
    outer = np.isin(train_labels, ("S", "U"))
    x = np.asarray(train_x[outer], dtype=float)
    labels = train_labels[outer]
    center = x.mean(axis=0)
    scale = x.std(axis=0, ddof=1)
    scale[~np.isfinite(scale) | (scale <= 1e-12)] = 1.0
    z_train = (x - center) / scale
    train_direction = z_train[labels == "S"].mean(axis=0) - z_train[
        labels == "U"
    ].mean(axis=0)
    z_test = (np.asarray(test_x, dtype=float) - center) / scale
    test_trial = []
    for selected in selected_trials:
        test_labels = make_labels(
            command[selected], target[selected], groups, n_commands
        )
        test_trial.append(
            z_test[selected][test_labels == "S"].mean(axis=0)
            - z_test[selected][test_labels == "U"].mean(axis=0)
        )
    test_direction = np.stack(test_trial).mean(axis=0)
    return (
        train_direction,
        test_direction,
        cosine(train_direction, test_direction),
    )


def mapping(
    train_x: np.ndarray,
    test_x: np.ndarray,
    train_command: np.ndarray,
    train_target: np.ndarray,
    test_command: np.ndarray,
    test_target: np.ndarray,
    selected_trials: list[np.ndarray],
    groups,
    n_commands: int,
) -> dict:
    labels = make_labels(
        train_command, train_target, groups, n_commands
    )
    score, _ = independent_lda_scores(train_x, labels, test_x)
    trial_values = effects(
        score,
        test_command,
        test_target,
        selected_trials,
        groups,
        n_commands,
    )
    train_direction, test_direction, alignment = direction(
        train_x,
        labels,
        test_x,
        test_command,
        test_target,
        selected_trials,
        groups,
        n_commands,
    )
    return {
        "trial_values": trial_values,
        "effect": float(np.mean(trial_values)),
        "train_direction": train_direction,
        "test_direction": test_direction,
        "alignment": alignment,
    }


def asme_subject(arrays: dict, subject: int) -> dict:
    train = (
        (arrays["subject"] == subject)
        & (arrays["task"] == "offline")
        & arrays["accepted"].astype(bool)
    )
    test = (
        (arrays["subject"] == subject)
        & (arrays["task"] == "online")
        & arrays["accepted"].astype(bool)
    )
    train_x = arrays["eeg_bins"][train, :, 1:].reshape(
        int(np.sum(train)), -1
    )
    test_x = arrays["eeg_bins"][test, :, 1:].reshape(
        int(np.sum(test)), -1
    )
    train_command = arrays["command"][train].astype(int)
    train_target = arrays["target"][train].astype(int)
    test_command = arrays["command"][test].astype(int)
    test_target = arrays["target"][test].astype(int)
    train_x, test_x = residualize(
        train_x,
        test_x,
        train_command,
        train_target,
        test_command,
        30,
    )
    test_selections = selections(
        test_command, arrays["trial"][test].astype(int), 30
    )
    physical = mapping(
        train_x,
        test_x,
        train_command,
        train_target,
        test_command,
        test_target,
        test_selections,
        ASME_PHYSICAL,
        30,
    )
    wrong = [
        mapping(
            train_x,
            test_x,
            train_command,
            train_target,
            test_command,
            test_target,
            test_selections,
            groups,
            30,
        )
        for groups in ASME_WRONG
    ]
    return {"subject": subject, "physical": physical, "wrong": wrong}


def kojima_subject(arrays: dict, subject: str) -> dict:
    base = (
        (arrays["subject"] == subject)
        & (arrays["task"] == "2stream")
        & arrays["accepted"].astype(bool)
    )
    runs = sorted(int(value) for value in np.unique(arrays["run"][base]))
    folds = []
    for heldout_run in runs:
        train = base & (arrays["run"] != heldout_run)
        test = base & (arrays["run"] == heldout_run)
        train_x = arrays["eeg_bins"][train, :, 1:].reshape(
            int(np.sum(train)), -1
        )
        test_x = arrays["eeg_bins"][test, :, 1:].reshape(
            int(np.sum(test)), -1
        )
        train_command = arrays["command"][train].astype(int)
        train_target = arrays["target"][train].astype(int)
        test_command = arrays["command"][test].astype(int)
        test_target = arrays["target"][test].astype(int)
        train_x, test_x = residualize(
            train_x,
            test_x,
            train_command,
            train_target,
            test_command,
            4,
        )
        test_selections = selections(
            test_command, arrays["trial"][test].astype(int), 4
        )
        folds.append(
            {
                "run": heldout_run,
                "physical": mapping(
                    train_x,
                    test_x,
                    train_command,
                    train_target,
                    test_command,
                    test_target,
                    test_selections,
                    KOJIMA_PHYSICAL,
                    4,
                ),
                "wrong": [
                    mapping(
                        train_x,
                        test_x,
                        train_command,
                        train_target,
                        test_command,
                        test_target,
                        test_selections,
                        groups,
                        4,
                    )
                    for groups in KOJIMA_WRONG
                ],
            }
        )
    return {
        "subject": subject,
        "runs": runs,
        "folds": folds,
        "physical_effect": float(
            np.mean(
                [
                    value
                    for fold in folds
                    for value in fold["physical"]["trial_values"]
                ]
            )
        ),
        "wrong_effects": [
            float(
                np.mean(
                    [
                        value
                        for fold in folds
                        for value in fold["wrong"][mapping_index][
                            "trial_values"
                        ]
                    ]
                )
            )
            for mapping_index in range(2)
        ],
        "physical_alignment": float(
            np.mean([fold["physical"]["alignment"] for fold in folds])
        ),
        "wrong_alignments": [
            float(
                np.mean(
                    [
                        fold["wrong"][mapping_index]["alignment"]
                        for fold in folds
                    ]
                )
            )
            for mapping_index in range(2)
        ],
    }


def bootstrap_ci(values: np.ndarray) -> list[float]:
    rng = np.random.default_rng(BOOTSTRAP_SEED)
    ids = rng.integers(0, len(values), size=(BOOTSTRAP_N, len(values)))
    return [
        float(value)
        for value in np.quantile(values[ids].mean(axis=1), [0.025, 0.975])
    ]


def exact_p(values: np.ndarray) -> float:
    observed = float(values.mean())
    permutations = [
        float(np.mean(values * np.asarray(signs)))
        for signs in itertools.product((-1.0, 1.0), repeat=len(values))
    ]
    return float(np.mean(np.asarray(permutations) >= observed - 1e-15))


def summarize(values: np.ndarray) -> dict:
    return {
        "mean": float(values.mean()),
        "bootstrap_95_CI_mean": bootstrap_ci(values),
        "n_positive": int(np.sum(values > 0)),
        "n_total": int(len(values)),
        "one_sided_exact_signflip_p": exact_p(values),
        "participant_values": [float(value) for value in values],
    }


def max_error(left, right) -> float:
    return float(
        np.max(
            np.abs(
                np.asarray(left, dtype=float)
                - np.asarray(right, dtype=float)
            )
        )
    )


def main() -> None:
    if AUDIT.exists():
        raise FileExistsError("Immutable audit already exists.")
    reported = json.loads(RESULT.read_text(encoding="utf-8"))
    lock = json.loads(LOCK.read_text(encoding="utf-8"))
    with np.load(FEATURE_ASME, allow_pickle=True) as source:
        asme_arrays = {key: source[key] for key in source.files}
    with np.load(FEATURE_KOJIMA, allow_pickle=True) as source:
        kojima_arrays = {key: source[key] for key in source.files}
    asme_records = [
        asme_subject(asme_arrays, subject) for subject in range(1, 11)
    ]
    asme_rows = []
    for participant_index, record in enumerate(asme_records):
        physical = record["physical"]
        wrong_1, wrong_2 = record["wrong"]
        between = float(
            np.mean(
                [
                    cosine(
                        physical["test_direction"],
                        asme_records[donor]["physical"][
                            "train_direction"
                        ],
                    )
                    for donor in range(len(asme_records))
                    if donor != participant_index
                ]
            )
        )
        asme_rows.append(
            {
                "physical": physical["effect"],
                "binding": physical["effect"]
                - max(wrong_1["effect"], wrong_2["effect"]),
                "identity": physical["alignment"] - between,
                "mapping_alignment": physical["alignment"]
                - max(wrong_1["alignment"], wrong_2["alignment"]),
            }
        )

    kojima_records = [
        kojima_subject(kojima_arrays, subject)
        for subject in KOJIMA_SUBJECTS
    ]
    kojima_rows = []
    for participant_index, record in enumerate(kojima_records):
        between_folds = []
        for fold_index, run in enumerate(record["runs"]):
            test_direction = record["folds"][fold_index]["physical"][
                "test_direction"
            ]
            between_folds.append(
                float(
                    np.mean(
                        [
                            cosine(
                                test_direction,
                                donor["folds"][
                                    donor["runs"].index(run)
                                ]["physical"]["train_direction"],
                            )
                            for donor_index, donor in enumerate(kojima_records)
                            if donor_index != participant_index
                        ]
                    )
                )
            )
        between = float(np.mean(between_folds))
        kojima_rows.append(
            {
                "physical": record["physical_effect"],
                "binding": record["physical_effect"]
                - max(record["wrong_effects"]),
                "identity": record["physical_alignment"] - between,
                "mapping_alignment": record["physical_alignment"]
                - max(record["wrong_alignments"]),
            }
        )
    recomputed_summaries = {
        "ASME30": {
            key: summarize(np.asarray([row[key] for row in asme_rows]))
            for key in asme_rows[0]
        },
        "Kojima2024B": {
            key: summarize(np.asarray([row[key] for row in kojima_rows]))
            for key in kojima_rows[0]
        },
    }
    summary_error = max(
        max_error(
            recomputed_summaries[dataset][metric][key],
            reported["summaries"][dataset][metric][key],
        )
        for dataset in recomputed_summaries
        for metric in recomputed_summaries[dataset]
        for key in (
            "mean",
            "bootstrap_95_CI_mean",
            "n_positive",
            "one_sided_exact_signflip_p",
            "participant_values",
        )
    )
    row_error = max(
        abs(
            float(row[metric])
            - float(reported_row[reported_key])
        )
        for dataset, rows, reported_rows in (
            (
                "ASME30",
                asme_rows,
                reported["participant_results"]["ASME30"],
            ),
            (
                "Kojima2024B",
                kojima_rows,
                reported["participant_results"]["Kojima2024B"],
            ),
        )
        for row, reported_row in zip(rows, reported_rows)
        for metric, reported_key in (
            ("physical", "physical_effect"),
            ("binding", "binding_specificity"),
            ("identity", "identity_advantage"),
            ("mapping_alignment", "mapping_alignment_specificity"),
        )
    )
    amendment = json.loads(
        REPRODUCIBILITY_AMENDMENT.read_text(encoding="utf-8")
    )
    if amendment["base_lock_sha256"] != sha256(LOCK):
        raise RuntimeError("Reproducibility amendment base-lock hash mismatch.")
    semantic_inputs = amendment["semantic_json_sha256"]
    frozen_hashes = {}
    for name, item in lock["frozen_files"].items():
        if name in semantic_inputs:
            expected = semantic_inputs[name]
            observed = semantic_json_sha256(ROOT / item["path"])
            mode = "semantic_json_excluding_created_utc"
        else:
            expected = item["sha256"]
            observed = sha256(ROOT / item["path"])
            mode = "byte_sha256"
        frozen_hashes[name] = {
            "mode": mode,
            "expected": expected,
            "observed": observed,
            "match": observed == expected,
        }
    checks = {
        "all_lock_hashes_match": all(
            item["match"] for item in frozen_hashes.values()
        ),
        "participant_values_match": row_error <= TOLERANCE,
        "summaries_match": summary_error <= TOLERANCE,
        "all_reported_gates_pass": all(reported["gates"].values()),
        "reported_conjunction_matches": all(reported["gates"].values())
        == reported["command_identity_explanation_disfavored"],
    }
    audit_pass = all(checks.values())
    output = {
        "audit": "WZ-HIC-cross-dataset-command-identity-residual-v1-independent-source-audit",
        "status": "PASS" if audit_pass else "FAIL",
        "audit_pass": audit_pass,
        "checks": checks,
        "absolute_tolerance": TOLERANCE,
        "numerical_errors": {
            "participant_max_abs": row_error,
            "summary_max_abs": summary_error,
        },
        "recomputed_summaries": recomputed_summaries,
        "frozen_hash_checks": frozen_hashes,
        "source_hashes": {
            "result": sha256(RESULT),
            "independent_lda_source": sha256(INDEPENDENT_LDA_SOURCE),
            "auditor": sha256(Path(__file__)),
            "reproducibility_amendment": sha256(REPRODUCIBILITY_AMENDMENT),
        },
        "interpretation": (
            "Physical, binding, identity and mapping-alignment effects in "
            "both datasets were independently recomputed using the separate "
            "source-audit Ledoit-Wolf implementation."
        ),
    }
    AUDIT.write_text(
        json.dumps(output, ensure_ascii=False, indent=2) + "\n",
        encoding="utf-8",
    )
    print(json.dumps(output, ensure_ascii=False, indent=2))
    if not audit_pass:
        raise SystemExit(1)


if __name__ == "__main__":
    main()
```
