## Supplementary material for "Falsifiable substitution tests reveal task-structured neural evidence for auditory attention": Article_Analysis_Source_Code: audit_wz_hic_individual_geometry_synthesis_v1.py

```
#!/usr/bin/env python3
"""Independent aggregation audit for individual-geometry synthesis v1."""

from __future__ import annotations

import csv
import hashlib
import json
from pathlib import Path


ROOT = Path(__file__).resolve().parents[1]
SYNTHESIS = (
    ROOT
    / "results"
    / "wz_hic_cross_dataset"
    / "individual_geometry_synthesis_v1"
    / "synthesis_v1.json"
)
CSV = SYNTHESIS.with_name("participant_metrics_v1.csv")
AUDIT = SYNTHESIS.with_name("independent_aggregation_audit_v1.json")


def sha256(path: Path) -> str:
    digest = hashlib.sha256()
    with path.open("rb") as handle:
        for block in iter(lambda: handle.read(1024 * 1024), b""):
            digest.update(block)
    return digest.hexdigest()


def main() -> None:
    if AUDIT.exists():
        raise FileExistsError("Immutable aggregation audit already exists.")
    synthesis = json.loads(SYNTHESIS.read_text(encoding="utf-8"))
    with CSV.open("r", encoding="utf-8-sig", newline="") as handle:
        rows = list(csv.DictReader(handle))
    source_checks = {}
    for relative, expected in synthesis["source_hashes"].items():
        observed = sha256(ROOT / relative)
        source_checks[relative] = {
            "expected": expected,
            "observed": observed,
            "match": observed == expected,
        }
    metrics = (
        "physical_effect",
        "binding_specificity",
        "nonocular_specificity",
        "identity_advantage",
        "mapping_alignment_specificity",
        "residual_physical_effect",
        "residual_binding_specificity",
    )
    counts = {
        metric: {
            "positive": sum(float(row[metric]) > 0 for row in rows),
            "total": len(rows),
        }
        for metric in metrics
    }
    dataset_counts = {
        dataset: sum(row["dataset"] == dataset for row in rows)
        for dataset in ("ASME30", "Kojima2024B")
    }
    checks = {
        "all_source_hashes_match": all(
            item["match"] for item in source_checks.values()
        ),
        "csv_has_17_unique_participants": len(rows) == 17
        and len(
            {(row["dataset"], row["participant"]) for row in rows}
        )
        == 17,
        "dataset_counts_match": dataset_counts
        == {"ASME30": 10, "Kojima2024B": 7},
        "positive_counts_match": counts
        == synthesis["participant_positive_counts_descriptive"],
        "all_source_conjunctions_pass": all(
            synthesis["source_conjunctions"].values()
        ),
        "shared_template_boundary_is_negative": synthesis[
            "shared_template_boundary"
        ]["shared_template_supported"]
        is False,
    }
    audit_pass = all(checks.values())
    output = {
        "audit": "WZ-HIC-individual-geometry-synthesis-v1-independent-aggregation-audit",
        "status": "PASS" if audit_pass else "FAIL",
        "audit_pass": audit_pass,
        "checks": checks,
        "dataset_counts": dataset_counts,
        "recomputed_positive_counts": counts,
        "source_hash_checks": source_checks,
        "source_hashes": {
            "synthesis": sha256(SYNTHESIS),
            "participant_csv": sha256(CSV),
            "auditor": sha256(Path(__file__)),
        },
    }
    AUDIT.write_text(
        json.dumps(output, ensure_ascii=False, indent=2) + "\n",
        encoding="utf-8",
    )
    print(json.dumps(output, ensure_ascii=False, indent=2))
    if not audit_pass:
        raise SystemExit(1)


if __name__ == "__main__":
    main()
```
