## Supplementary material for "Falsifiable substitution tests reveal task-structured neural evidence for auditory attention": Article_Analysis_Source_Code: audit_wz_hic_kojima2024b_crossrun_individual_geometry_v1.py

```
#!/usr/bin/env python3
"""Independent source audit of Kojima2024B cross-run individual geometry."""

from __future__ import annotations

import csv
import hashlib
import itertools
import json
import math
from pathlib import Path

import numpy as np
from sklearn.covariance import ledoit_wolf


ROOT = Path(__file__).resolve().parents[1]
FEATURES = (
    ROOT / "data" / "derived" / "kojima2024b_wz_hic_confirmation_features_v1.npz"
)
PROTOCOL = (
    ROOT
    / "docs"
    / "wz_hic_kojima2024b_crossrun_individual_geometry_protocol_v1.json"
)
LOCK = (
    ROOT
    / "docs"
    / "wz_hic_kojima2024b_crossrun_individual_geometry_implementation_lock_v1.json"
)
RESULT_DIR = (
    ROOT
    / "results"
    / "wz_hic_kojima2024b"
    / "crossrun_individual_geometry_v1"
)
RESULT = RESULT_DIR / "crossrun_decision_v1.json"
PARTICIPANT_CSV = RESULT_DIR / "participant_geometry_v1.csv"
TIME_CSV = RESULT_DIR / "time_resolved_v1.csv"
AUDIT = RESULT_DIR / "independent_source_audit_v1.json"

SUBJECTS = tuple("BDFHJLN")
PHYSICAL = ((1, 2), (3, 4))
WRONG = (((1, 3), (2, 4)), ((1, 4), (2, 3)))
HORIZON = 3
BOOTSTRAP_N = 20_000
BOOTSTRAP_SEED = 20260730
TOLERANCE = 5e-7


def sha256(path: Path) -> str:
    digest = hashlib.sha256()
    with path.open("rb") as handle:
        for block in iter(lambda: handle.read(1024 * 1024), b""):
            digest.update(block)
    return digest.hexdigest()


def labels(command: np.ndarray, target: np.ndarray, groups) -> np.ndarray:
    lookup = np.full(5, -1, dtype=int)
    flattened = []
    for index, group in enumerate(groups):
        lookup[list(group)] = index
        flattened.extend(group)
    if sorted(flattened) != [1, 2, 3, 4]:
        raise RuntimeError("Invalid pairing.")
    result = np.full(command.size, "U", dtype="U1")
    target_mask = command == target
    result[target_mask] = "T"
    result[(lookup[command] == lookup[target]) & ~target_mask] = "S"
    return result


def select_trials(
    command: np.ndarray, trial: np.ndarray
) -> list[np.ndarray]:
    selected_trials = []
    for trial_id in sorted(int(value) for value in np.unique(trial)):
        ids = np.flatnonzero(trial == trial_id)
        chunks = []
        for value in range(1, 5):
            value_ids = ids[command[ids] == value]
            if value_ids.size < HORIZON:
                chunks = []
                break
            chunks.append(value_ids[:HORIZON])
        if len(chunks) == 4:
            selected_trials.append(np.concatenate(chunks))
    return selected_trials


def independent_scores(
    train_x: np.ndarray,
    train_labels: np.ndarray,
    test_x: np.ndarray,
) -> tuple[np.ndarray, dict]:
    outer = np.isin(train_labels, ("S", "U"))
    x = np.asarray(train_x[outer], dtype=float)
    y = (train_labels[outer] == "S").astype(int)
    center = x.mean(axis=0)
    scale = x.std(axis=0, ddof=1)
    scale[~np.isfinite(scale) | (scale < 1e-12)] = 1.0
    z = (x - center) / scale
    mean_u = z[y == 0].mean(axis=0)
    mean_s = z[y == 1].mean(axis=0)
    residual = np.vstack(
        (z[y == 0] - mean_u, z[y == 1] - mean_s)
    )
    covariance, _ = ledoit_wolf(residual, assume_centered=True)
    delta = mean_s - mean_u
    try:
        direction = np.linalg.solve(covariance, delta)
    except np.linalg.LinAlgError:
        direction = np.linalg.pinv(covariance, hermitian=True) @ delta
    intercept = float(-0.5 * (mean_s + mean_u) @ direction)

    def decision(values: np.ndarray) -> np.ndarray:
        return (
            (np.asarray(values, dtype=float) - center) / scale
        ) @ direction + intercept

    train_score = decision(x)
    score_mean = float(train_score.mean())
    score_sd = float(train_score.std(ddof=1))
    return (decision(test_x) - score_mean) / score_sd, {
        "n_train_S": int(np.sum(train_labels == "S")),
        "n_train_U": int(np.sum(train_labels == "U")),
        "n_train_T_excluded": int(np.sum(train_labels == "T")),
        "training_score_mean": score_mean,
        "training_score_sd": score_sd,
    }


def score_trial_values(
    score: np.ndarray,
    command: np.ndarray,
    target: np.ndarray,
    selections: list[np.ndarray],
    groups,
) -> list[float]:
    values = []
    for selected in selections:
        test_labels = labels(
            command[selected], target[selected], groups
        )
        counts = tuple(
            int(np.sum(test_labels == value))
            for value in ("T", "S", "U")
        )
        if counts != (3, 3, 6):
            raise RuntimeError(f"Target-free counts failed: {counts}")
        values.append(
            float(score[selected][test_labels == "S"].mean())
            - float(score[selected][test_labels == "U"].mean())
        )
    return values


def cosine(left: np.ndarray, right: np.ndarray) -> float:
    return float(
        np.vdot(left, right)
        / (np.linalg.norm(left) * np.linalg.norm(right))
    )


def direction_pair(
    train_x: np.ndarray,
    train_labels: np.ndarray,
    test_x: np.ndarray,
    command: np.ndarray,
    target: np.ndarray,
    selections: list[np.ndarray],
    groups,
) -> tuple[np.ndarray, np.ndarray, float]:
    outer = np.isin(train_labels, ("S", "U"))
    x = np.asarray(train_x[outer], dtype=float)
    outer_labels = train_labels[outer]
    center = x.mean(axis=0)
    scale = x.std(axis=0, ddof=1)
    scale[~np.isfinite(scale) | (scale <= 1e-12)] = 1.0
    z_train = (x - center) / scale
    train_direction = z_train[outer_labels == "S"].mean(
        axis=0
    ) - z_train[outer_labels == "U"].mean(axis=0)
    z_test = (np.asarray(test_x, dtype=float) - center) / scale
    test_trial_directions = []
    for selected in selections:
        test_labels = labels(
            command[selected], target[selected], groups
        )
        test_trial_directions.append(
            z_test[selected][test_labels == "S"].mean(axis=0)
            - z_test[selected][test_labels == "U"].mean(axis=0)
        )
    test_direction = np.stack(test_trial_directions).mean(axis=0)
    return (
        train_direction,
        test_direction,
        cosine(train_direction, test_direction),
    )


def audit_mapping_fold(
    train_bins: np.ndarray,
    test_bins: np.ndarray,
    train_command: np.ndarray,
    train_target: np.ndarray,
    test_command: np.ndarray,
    test_target: np.ndarray,
    selections: list[np.ndarray],
    groups,
) -> dict:
    train_full = train_bins.reshape(train_bins.shape[0], -1)
    test_full = test_bins.reshape(test_bins.shape[0], -1)
    train_labels = labels(train_command, train_target, groups)
    score, fit = independent_scores(
        train_full, train_labels, test_full
    )
    full_values = score_trial_values(
        score, test_command, test_target, selections, groups
    )
    train_direction, test_direction, alignment = direction_pair(
        train_full,
        train_labels,
        test_full,
        test_command,
        test_target,
        selections,
        groups,
    )
    time_values = []
    for bin_index in range(10):
        time_score, _ = independent_scores(
            train_bins[:, :, bin_index],
            train_labels,
            test_bins[:, :, bin_index],
        )
        time_values.append(
            score_trial_values(
                time_score,
                test_command,
                test_target,
                selections,
                groups,
            )
        )
    return {
        "full": full_values,
        "time": time_values,
        "train_direction": train_direction,
        "test_direction": test_direction,
        "alignment": alignment,
        "fit": fit,
    }


def audit_subject(arrays: dict, subject: str) -> dict:
    base = (
        (arrays["subject"] == subject)
        & (arrays["task"] == "2stream")
        & arrays["accepted"].astype(bool)
    )
    runs = sorted(int(value) for value in np.unique(arrays["run"][base]))
    folds = []
    for heldout_run in runs:
        train = base & (arrays["run"] != heldout_run)
        test = base & (arrays["run"] == heldout_run)
        train_bins = arrays["eeg_bins"][train, :, 1:]
        test_bins = arrays["eeg_bins"][test, :, 1:]
        train_command = arrays["command"][train].astype(int)
        train_target = arrays["target"][train].astype(int)
        test_command = arrays["command"][test].astype(int)
        test_target = arrays["target"][test].astype(int)
        selections = select_trials(
            test_command, arrays["trial"][test].astype(int)
        )
        physical = audit_mapping_fold(
            train_bins,
            test_bins,
            train_command,
            train_target,
            test_command,
            test_target,
            selections,
            PHYSICAL,
        )
        wrong = [
            audit_mapping_fold(
                train_bins,
                test_bins,
                train_command,
                train_target,
                test_command,
                test_target,
                selections,
                groups,
            )
            for groups in WRONG
        ]
        train_eog = arrays["eog_bins"][train, :, 1:].reshape(
            int(np.sum(train)), -1
        )
        test_eog = arrays["eog_bins"][test, :, 1:].reshape(
            int(np.sum(test)), -1
        )
        physical_labels = labels(
            train_command, train_target, PHYSICAL
        )
        eog_score, eog_fit = independent_scores(
            train_eog, physical_labels, test_eog
        )
        eog = score_trial_values(
            eog_score,
            test_command,
            test_target,
            selections,
            PHYSICAL,
        )
        folds.append(
            {
                "heldout_run": heldout_run,
                "n_trials": len(selections),
                "physical": physical,
                "wrong": wrong,
                "eog": eog,
                "eog_fit": eog_fit,
            }
        )

    def all_full(kind: str, wrong_index: int | None = None):
        return np.asarray(
            [
                value
                for fold in folds
                for value in (
                    fold[kind]["full"]
                    if wrong_index is None
                    else fold[kind][wrong_index]["full"]
                )
            ]
        )

    physical_values = all_full("physical")
    wrong_values = [all_full("wrong", index) for index in range(2)]
    eog_values = np.asarray(
        [value for fold in folds for value in fold["eog"]]
    )

    def time_mean(kind: str, wrong_index: int | None = None):
        return np.asarray(
            [
                np.mean(
                    [
                        value
                        for fold in folds
                        for value in (
                            fold[kind]["time"][bin_index]
                            if wrong_index is None
                            else fold[kind][wrong_index]["time"][bin_index]
                        )
                    ]
                )
                for bin_index in range(10)
            ]
        )

    return {
        "subject": subject,
        "runs": runs,
        "folds": folds,
        "n_trials": len(physical_values),
        "physical_effect": float(physical_values.mean()),
        "wrong_effects": [float(values.mean()) for values in wrong_values],
        "eog_effect": float(eog_values.mean()),
        "physical_time": time_mean("physical"),
        "wrong_time": [
            time_mean("wrong", index) for index in range(2)
        ],
        "physical_alignment": float(
            np.mean([fold["physical"]["alignment"] for fold in folds])
        ),
        "wrong_alignments": [
            float(
                np.mean(
                    [
                        fold["wrong"][index]["alignment"]
                        for fold in folds
                    ]
                )
            )
            for index in range(2)
        ],
    }


def bootstrap_ci(values: np.ndarray) -> list[float]:
    rng = np.random.default_rng(BOOTSTRAP_SEED)
    ids = rng.integers(0, len(values), size=(BOOTSTRAP_N, len(values)))
    return [
        float(value)
        for value in np.quantile(values[ids].mean(axis=1), [0.025, 0.975])
    ]


def exact_p(values: np.ndarray) -> float:
    observed = float(values.mean())
    permuted = [
        float(np.mean(values * np.asarray(signs)))
        for signs in itertools.product((-1.0, 1.0), repeat=len(values))
    ]
    return float(np.mean(np.asarray(permuted) >= observed - 1e-15))


def summarize(values: np.ndarray) -> dict:
    return {
        "mean": float(values.mean()),
        "bootstrap_95_CI_mean": bootstrap_ci(values),
        "n_positive": int(np.sum(values > 0)),
        "n_total": int(len(values)),
        "one_sided_exact_signflip_p": exact_p(values),
        "participant_values": [float(value) for value in values],
    }


def t_stat(values: np.ndarray) -> np.ndarray:
    denominator = values.std(axis=0, ddof=1) / math.sqrt(len(values))
    return np.divide(
        values.mean(axis=0),
        denominator,
        out=np.zeros(values.shape[1]),
        where=denominator > 1e-12,
    )


def exact_max_t(values: np.ndarray):
    observed = t_stat(values)
    maxima = []
    for signs in itertools.product((-1.0, 1.0), repeat=len(values)):
        maxima.append(
            float(
                np.max(
                    np.abs(
                        t_stat(values * np.asarray(signs)[:, None])
                    )
                )
            )
        )
    maxima = np.asarray(maxima)
    p = (
        maxima[:, None] >= np.abs(observed)[None, :] - 1e-15
    ).mean(axis=0)
    return observed, p, maxima


def max_error(left, right) -> float:
    return float(
        np.max(
            np.abs(
                np.asarray(left, dtype=float)
                - np.asarray(right, dtype=float)
            )
        )
    )


def main() -> None:
    if AUDIT.exists():
        raise FileExistsError("Immutable audit already exists.")
    reported = json.loads(RESULT.read_text(encoding="utf-8"))
    protocol = json.loads(PROTOCOL.read_text(encoding="utf-8"))
    lock = json.loads(LOCK.read_text(encoding="utf-8"))
    with np.load(FEATURES, allow_pickle=True) as source:
        arrays = {key: source[key] for key in source.files}
    records = [audit_subject(arrays, subject) for subject in SUBJECTS]
    rows = []
    time_specificity = []
    for participant_index, record in enumerate(records):
        fold_between = []
        for fold_index, heldout_run in enumerate(record["runs"]):
            test_direction = record["folds"][fold_index]["physical"][
                "test_direction"
            ]
            donor_cosines = []
            for donor_index, donor in enumerate(records):
                if donor_index == participant_index:
                    continue
                donor_fold = donor["runs"].index(heldout_run)
                donor_cosines.append(
                    cosine(
                        test_direction,
                        donor["folds"][donor_fold]["physical"][
                            "train_direction"
                        ],
                    )
                )
            fold_between.append(float(np.mean(donor_cosines)))
        between = float(np.mean(fold_between))
        best_wrong_effect = max(record["wrong_effects"])
        best_wrong_alignment = max(record["wrong_alignments"])
        rows.append(
            {
                "subject": record["subject"],
                "n_runs": len(record["runs"]),
                "n_eligible_trials": record["n_trials"],
                "physical_full_effect": record["physical_effect"],
                "wrong_1_full_effect": record["wrong_effects"][0],
                "wrong_2_full_effect": record["wrong_effects"][1],
                "best_wrong_full_effect": best_wrong_effect,
                "binding_specificity": record["physical_effect"]
                - best_wrong_effect,
                "physical_eog_effect": record["eog_effect"],
                "nonocular_specificity": record["physical_effect"]
                - record["eog_effect"],
                "physical_within_alignment": record["physical_alignment"],
                "wrong_1_within_alignment": record["wrong_alignments"][0],
                "wrong_2_within_alignment": record["wrong_alignments"][1],
                "best_wrong_within_alignment": best_wrong_alignment,
                "mapping_alignment_specificity": record[
                    "physical_alignment"
                ]
                - best_wrong_alignment,
                "between_person_substitution": between,
                "identity_advantage": record["physical_alignment"] - between,
            }
        )
        time_specificity.append(
            record["physical_time"]
            - np.maximum(record["wrong_time"][0], record["wrong_time"][1])
        )
    time_specificity = np.stack(time_specificity)
    time_t, time_p, time_null = exact_max_t(time_specificity)
    summaries = {
        "physical_crossrun": summarize(
            np.asarray([row["physical_full_effect"] for row in rows])
        ),
        "binding_specificity": summarize(
            np.asarray([row["binding_specificity"] for row in rows])
        ),
        "nonocular_specificity": summarize(
            np.asarray([row["nonocular_specificity"] for row in rows])
        ),
        "identity_advantage": summarize(
            np.asarray([row["identity_advantage"] for row in rows])
        ),
        "mapping_alignment_specificity": summarize(
            np.asarray(
                [row["mapping_alignment_specificity"] for row in rows]
            )
        ),
        "physical_within_alignment": summarize(
            np.asarray(
                [row["physical_within_alignment"] for row in rows]
            )
        ),
        "between_person_substitution": summarize(
            np.asarray(
                [row["between_person_substitution"] for row in rows]
            )
        ),
    }

    def gate(name: str, required: int) -> bool:
        item = summaries[name]
        return (
            item["mean"] > 0
            and item["bootstrap_95_CI_mean"][0] > 0
            and item["n_positive"] >= required
            and item["one_sided_exact_signflip_p"] <= 0.05
        )

    gates = {
        "K0_coverage": all(
            len(record["runs"]) == 6 and record["n_trials"] >= 23
            for record in records
        ),
        "K1_physical_crossrun": gate("physical_crossrun", 7),
        "K2_binding_specificity": gate("binding_specificity", 6),
        "K3_nonocular_specificity": gate("nonocular_specificity", 6),
        "K4_identity_advantage": gate("identity_advantage", 6),
        "K5_mapping_alignment_specificity": gate(
            "mapping_alignment_specificity", 6
        ),
        "K6_time_localization": bool(
            np.any(
                (time_specificity.mean(axis=0) > 0)
                & (time_p <= 0.05)
            )
        ),
    }
    numeric_keys = [
        key
        for key in rows[0]
        if key not in ("subject", "n_runs", "n_eligible_trials")
    ]
    participant_error = max(
        abs(float(row[key]) - float(reported_row[key]))
        for row, reported_row in zip(
            rows, reported["participant_geometry"]
        )
        for key in numeric_keys
    )
    summary_error = max(
        max_error(
            summaries[name][key], reported["summaries"][name][key]
        )
        for name in summaries
        for key in (
            "mean",
            "bootstrap_95_CI_mean",
            "one_sided_exact_signflip_p",
            "participant_values",
        )
    )
    reported_time = reported["time_resolved_binding_specificity"]
    time_errors = {
        "mean": max_error(
            time_specificity.mean(axis=0),
            [
                row["mean_physical_minus_best_wrong"]
                for row in reported_time
            ],
        ),
        "t": max_error(
            time_t, [row["t_value"] for row in reported_time]
        ),
        "p": max_error(
            time_p, [row["max_t_fwer_p"] for row in reported_time]
        ),
        "participant": max_error(
            time_specificity.T,
            [row["participant_values"] for row in reported_time],
        ),
        "null_95": abs(
            float(np.quantile(time_null, 0.95))
            - float(reported["time_max_t"]["null_95_percentile"])
        ),
    }
    with PARTICIPANT_CSV.open(
        "r", encoding="utf-8-sig", newline=""
    ) as handle:
        participant_csv = list(csv.DictReader(handle))
    csv_error = max(
        abs(float(row[key]) - float(csv_row[key]))
        for row, csv_row in zip(rows, participant_csv)
        for key in numeric_keys
    )
    frozen_hashes = {}
    for name, item in lock["frozen_files"].items():
        observed = sha256(ROOT / item["path"])
        frozen_hashes[name] = {
            "expected": item["sha256"],
            "observed": observed,
            "match": observed == item["sha256"],
        }
    errors = {
        "participant_json_max_abs": participant_error,
        "participant_csv_max_abs": csv_error,
        "summary_max_abs": summary_error,
        **{f"time_{key}_max_abs": value for key, value in time_errors.items()},
    }
    checks = {
        "all_lock_hashes_match": all(
            item["match"] for item in frozen_hashes.values()
        ),
        "feature_hash_matches_protocol": sha256(FEATURES)
        == protocol["input"]["sha256"],
        "participant_values_match": participant_error <= TOLERANCE,
        "participant_csv_matches": csv_error <= TOLERANCE,
        "summaries_match": summary_error <= TOLERANCE,
        "time_results_match": max(time_errors.values()) <= TOLERANCE,
        "gates_match": gates == reported["gates"],
        "conjunction_matches": all(gates.values())
        == reported["cross_dataset_geometry_supported"],
        "target_events_excluded": all(
            fold["physical"]["fit"]["n_train_T_excluded"] > 0
            and all(
                wrong["fit"]["n_train_T_excluded"] > 0
                for wrong in fold["wrong"]
            )
            and fold["eog_fit"]["n_train_T_excluded"] > 0
            for record in records
            for fold in record["folds"]
        ),
        "coverage_matches": all(
            len(record["runs"]) == 6 and record["n_trials"] >= 23
            for record in records
        ),
        "csv_row_counts_match": len(participant_csv) == 7,
    }
    audit_pass = all(checks.values())
    output = {
        "audit": "WZ-HIC-Kojima2024B-crossrun-individual-geometry-v1-independent-source-audit",
        "status": "PASS" if audit_pass else "FAIL",
        "audit_pass": audit_pass,
        "absolute_tolerance": TOLERANCE,
        "checks": checks,
        "numerical_errors": errors,
        "recomputed": {
            "gates": gates,
            "summaries": summaries,
            "time_mean": [
                float(value) for value in time_specificity.mean(axis=0)
            ],
            "time_t": [float(value) for value in time_t],
            "time_max_t_fwer_p": [float(value) for value in time_p],
        },
        "frozen_hash_checks": frozen_hashes,
        "source_hashes": {
            "result": sha256(RESULT),
            "participant_csv": sha256(PARTICIPANT_CSV),
            "time_csv": sha256(TIME_CSV),
            "auditor": sha256(Path(__file__)),
        },
        "interpretation": (
            "The complete Kojima2024B conjunction was independently "
            "recomputed without importing the runner or LDA core."
        ),
    }
    AUDIT.write_text(
        json.dumps(output, ensure_ascii=False, indent=2) + "\n",
        encoding="utf-8",
    )
    print(json.dumps(output, ensure_ascii=False, indent=2))
    if not audit_pass:
        raise SystemExit(1)


if __name__ == "__main__":
    main()
```
