## Supplementary material for "Falsifiable substitution tests reveal task-structured neural evidence for auditory attention": Article_Analysis_Source_Code: audit_wz_hic_target_free_mechanism_v1.py

```
#!/usr/bin/env python3
"""Independently audit the saved target-free WZ-HIC decision."""

from __future__ import annotations

import hashlib
import itertools
import json
from datetime import datetime, timezone
from pathlib import Path

import numpy as np


ROOT = Path(__file__).resolve().parents[1]
RESULT = (
    ROOT
    / "results"
    / "wz_hic_cross_dataset"
    / "target_free_mechanism_v1"
    / "target_free_decision_v1.json"
)
AUDIT = RESULT.with_name("independent_outcome_audit_v1.json")
PROTOCOL = ROOT / "docs" / "wz_hic_target_free_mechanism_protocol_v1.json"
LOCK = (
    ROOT
    / "docs"
    / "wz_hic_target_free_mechanism_implementation_lock_v1c.json"
)
KOJIMA_FEATURES = (
    ROOT
    / "data"
    / "derived"
    / "kojima2024b_wz_hic_confirmation_features_v1.npz"
)
ASME_FEATURES = (
    ROOT / "data" / "derived" / "asme30_wz_hic_dataset_native_features_v2.npz"
)
KOJIMA_PRIMARY = (
    ROOT
    / "results"
    / "wz_hic_kojima2024b"
    / "confirmation_v1"
    / "confirmation_decision_v1.json"
)
ASME_PRIMARY = (
    ROOT
    / "results"
    / "wz_hic_asme30"
    / "dataset_native_v2"
    / "dataset_native_decision_v2.json"
)
BOOTSTRAP_SEED = 20260730
BOOTSTRAP_DRAWS = 20_000
TOL = 1e-12


def sha256(path: Path) -> str:
    h = hashlib.sha256()
    with path.open("rb") as f:
        for block in iter(lambda: f.read(1024 * 1024), b""):
            h.update(block)
    return h.hexdigest()


def close(a, b, tol=TOL) -> bool:
    if np.isnan(a) and np.isnan(b):
        return True
    return bool(abs(float(a) - float(b)) <= tol)


def exact_signflip_mean_p(values, null=0.0) -> float:
    x = np.asarray(values, dtype=float) - float(null)
    x = x[np.isfinite(x)]
    observed = float(np.mean(x))
    count = 0
    for signs in itertools.product((-1.0, 1.0), repeat=int(x.size)):
        statistic = float(np.mean(x * np.asarray(signs)))
        if statistic >= observed - 1e-15:
            count += 1
    return float(count / (2 ** int(x.size)))


def independent_summary(values, null=0.0) -> dict:
    x = np.asarray(values, dtype=float)
    x = x[np.isfinite(x)]
    rng = np.random.default_rng(BOOTSTRAP_SEED)
    index = rng.integers(0, x.size, size=(BOOTSTRAP_DRAWS, x.size))
    means = np.mean(x[index], axis=1)
    return {
        "mean": float(np.mean(x)),
        "bootstrap_95_CI_mean": [
            float(np.quantile(means, 0.025)),
            float(np.quantile(means, 0.975)),
        ],
        "n_positive": int(np.sum(x > null)),
        "n_total": int(x.size),
        "one_sided_exact_signflip_p": exact_signflip_mean_p(x, null),
        "participant_values": [float(value) for value in x],
    }


def compare_summary(saved: dict, observed: dict, prefix: str, failures: list):
    for field in (
        "mean",
        "n_positive",
        "n_total",
        "one_sided_exact_signflip_p",
    ):
        if not close(saved[field], observed[field]):
            failures.append(
                f"{prefix}.{field}: {saved[field]} != {observed[field]}"
            )
    if not np.allclose(
        saved["bootstrap_95_CI_mean"],
        observed["bootstrap_95_CI_mean"],
        atol=TOL,
        rtol=0,
    ):
        failures.append(f"{prefix}.bootstrap_95_CI_mean mismatch")
    if not np.allclose(
        saved["participant_values"],
        observed["participant_values"],
        atol=TOL,
        rtol=0,
        equal_nan=True,
    ):
        failures.append(f"{prefix}.participant_values mismatch")


def main() -> None:
    if AUDIT.exists():
        raise FileExistsError(AUDIT)
    failures = []
    d = json.loads(RESULT.read_text(encoding="utf-8"))
    expected_provenance = {
        "protocol_sha256": sha256(PROTOCOL),
        "implementation_lock_sha256": sha256(LOCK),
        "kojima_features_sha256": sha256(KOJIMA_FEATURES),
        "asme_features_sha256": sha256(ASME_FEATURES),
        "kojima_primary_result_sha256": sha256(KOJIMA_PRIMARY),
        "asme_primary_result_sha256": sha256(ASME_PRIMARY),
    }
    if d["provenance"] != expected_provenance:
        failures.append("Saved provenance does not match current inputs.")

    expected_ids = {
        "Kojima2024B": list("BDFHJLN"),
        "ASME30": ["1", "2", "3", "4", "5", "7", "8", "9", "10"],
    }
    if d["evaluable_participants"] != expected_ids:
        failures.append("Evaluable participant identities mismatch.")

    recomputed = {}
    count_audit = {}
    for dataset, ids in expected_ids.items():
        participants = d["participants"][dataset]
        absolute = []
        mapping = []
        nonocular = []
        free_minus_inclusive = []
        count_audit[dataset] = {}
        for subject in ids:
            item = participants[subject]
            free = item["target_free_EEG"]["certificate_AUC"]
            eog = item["target_free_EOG"]["certificate_AUC"]
            if dataset == "Kojima2024B":
                wrong_values = [
                    value["certificate_AUC"]
                    for value in item[
                        "target_free_wrong_pairings"
                    ].values()
                ]
                expected_free_counts = (3, 6)
                expected_inclusive_counts = (6, 6)
            else:
                wrong_values = [
                    value["certificate_AUC"]
                    for value in item[
                        "target_free_wrong_mappings"
                    ].values()
                ]
                expected_free_counts = (27, 30)
                expected_inclusive_counts = (30, 30)
            best_wrong = max(wrong_values)
            independently_computed = {
                "EEG_minus_EOG": free - eog,
                "EEG_minus_best_wrong": free - best_wrong,
                "target_free_minus_inclusive": (
                    free
                    - item["target_inclusive_EEG"]["certificate_AUC"]
                ),
            }
            for field, value in independently_computed.items():
                if not close(item["contrasts"][field], value):
                    failures.append(
                        f"{dataset}.{subject}.{field} mismatch"
                    )
            free_counts = (
                item["target_free_EEG"]["minimum_parent_event_count"],
                item["target_free_EEG"]["maximum_parent_event_count"],
            )
            inclusive_counts = (
                item["target_inclusive_EEG"][
                    "minimum_parent_event_count"
                ],
                item["target_inclusive_EEG"][
                    "maximum_parent_event_count"
                ],
            )
            if free_counts != expected_free_counts:
                failures.append(
                    f"{dataset}.{subject} target-free event counts "
                    f"{free_counts} != {expected_free_counts}"
                )
            if inclusive_counts != expected_inclusive_counts:
                failures.append(
                    f"{dataset}.{subject} inclusive event counts "
                    f"{inclusive_counts} != {expected_inclusive_counts}"
                )
            count_audit[dataset][subject] = {
                "target_free": list(free_counts),
                "target_inclusive": list(inclusive_counts),
            }
            absolute.append(free)
            mapping.append(independently_computed["EEG_minus_best_wrong"])
            nonocular.append(independently_computed["EEG_minus_EOG"])
            free_minus_inclusive.append(
                independently_computed["target_free_minus_inclusive"]
            )

        recomputed[dataset] = {
            "target_free_absolute": independent_summary(absolute, 0.5),
            "mapping_specificity": independent_summary(mapping, 0.0),
            "nonocular": independent_summary(nonocular, 0.0),
            "target_free_minus_inclusive": independent_summary(
                free_minus_inclusive, 0.0
            ),
        }
        for name, value in recomputed[dataset].items():
            compare_summary(
                d["summaries"][dataset][name],
                value,
                f"{dataset}.{name}",
                failures,
            )

    ks = recomputed["Kojima2024B"]
    ass = recomputed["ASME30"]
    gates = {
        "T1_Kojima_target_free_absolute": (
            ks["target_free_absolute"]["mean"] >= 0.75
            and ks["target_free_absolute"]["n_positive"] >= 6
            and ks["target_free_absolute"]["n_total"] == 7
            and ks["target_free_absolute"]["one_sided_exact_signflip_p"]
            <= 0.05
        ),
        "T2_Kojima_mapping_specificity": (
            ks["mapping_specificity"]["mean"] >= 0.15
            and ks["mapping_specificity"]["n_positive"] >= 6
            and ks["mapping_specificity"]["one_sided_exact_signflip_p"]
            <= 0.05
        ),
        "T3_Kojima_nonocular": (
            ks["nonocular"]["mean"] >= 0.05
            and ks["nonocular"]["n_positive"] >= 6
            and ks["nonocular"]["one_sided_exact_signflip_p"] <= 0.05
        ),
        "T4_ASME_target_free_absolute": (
            ass["target_free_absolute"]["mean"] >= 0.80
            and ass["target_free_absolute"]["n_positive"] >= 8
            and ass["target_free_absolute"]["n_total"] == 9
            and ass["target_free_absolute"]["one_sided_exact_signflip_p"]
            <= 0.05
        ),
        "T5_ASME_mapping_specificity": (
            ass["mapping_specificity"]["mean"] >= 0.15
            and ass["mapping_specificity"]["n_positive"] >= 8
            and ass["mapping_specificity"]["one_sided_exact_signflip_p"]
            <= 0.05
        ),
        "T6_ASME_nonocular": (
            ass["nonocular"]["mean"] >= 0.10
            and ass["nonocular"]["n_positive"] >= 8
            and ass["nonocular"]["one_sided_exact_signflip_p"] <= 0.05
        ),
        "implementation_equivalence": (
            d["implementation_equivalence"][
                "maximum_absolute_AUC_error"
            ]
            <= 1e-10
        ),
    }
    if d["gates"] != gates:
        failures.append("Saved gates do not match independent gates.")
    if not all(gates.values()):
        failures.append("Not all independently recomputed gates pass.")
    if d["status"] != "target_free_cross_dataset_mechanism_supported":
        failures.append("Saved status is inconsistent with passing gates.")

    audit = {
        "route": "WZ-HIC-target-free-mechanism-independent-audit-v1",
        "created_utc": datetime.now(timezone.utc).isoformat(),
        "status": "PASS" if not failures else "FAIL",
        "failures": failures,
        "audited_result_sha256": sha256(RESULT),
        "provenance": expected_provenance,
        "independently_recomputed_summaries": recomputed,
        "independently_recomputed_gates": gates,
        "event_count_audit": count_audit,
        "scope": (
            "Independent arithmetic, gate, provenance, event-count and "
            "archived-equivalence audit of the saved participant metrics. "
            "The locked signal-level runner and synthetic exclusion test "
            "remain the signal-level validation."
        ),
    }
    AUDIT.write_text(json.dumps(audit, indent=2) + "\n", encoding="utf-8")
    print(
        json.dumps(
            {
                "status": audit["status"],
                "n_failures": len(failures),
                "audit": str(AUDIT),
                "sha256": sha256(AUDIT),
            },
            indent=2,
        )
    )
    if failures:
        raise SystemExit(1)


if __name__ == "__main__":
    main()
```
