## Supplementary material for "Falsifiable substitution tests reveal task-structured neural evidence for auditory attention": Article_Analysis_Source_Code: audit_wz_hic_wearable_calibration_artifacts_v2.py

```
#!/usr/bin/env python3
"""Audit cross-implementation artifact consistency after the write collision."""

from __future__ import annotations

import csv
import hashlib
import json
import math
from datetime import datetime, timezone
from pathlib import Path


ROOT = Path(__file__).resolve().parents[1]
OUTDIR = (
    ROOT
    / "results"
    / "wz_hic_cross_dataset"
    / "wearable_calibration_v1"
)
RESULT = OUTDIR / "wearable_calibration_decision_v1.json"
PARTICIPANT_CSV = OUTDIR / "participant_metrics_v1.csv"
MONTAGE_CSV = OUTDIR / "montage_curve_v1.csv"
CALIBRATION_CSV = OUTDIR / "calibration_curve_v1.csv"
LOCK_V1 = (
    ROOT
    / "docs"
    / "wz_hic_cross_dataset_wearable_calibration_implementation_lock_v1.json"
)
AUDIT = OUTDIR / "final_v1_artifact_audit_v2.json"
TOLERANCE = 5e-12


def sha256(path: Path) -> str:
    digest = hashlib.sha256()
    with path.open("rb") as handle:
        for block in iter(lambda: handle.read(1024 * 1024), b""):
            digest.update(block)
    return digest.hexdigest()


def read_csv(path: Path) -> list[dict]:
    with path.open("r", newline="", encoding="utf-8-sig") as handle:
        return list(csv.DictReader(handle))


def error(left: float, right: float) -> float:
    return abs(float(left) - float(right))


def scalar_support(item: dict, required: int) -> bool:
    return (
        item["mean"] > 0
        and item["bootstrap_95_CI_mean"][0] > 0
        and item["n_positive"] >= required
        and item["one_sided_exact_signflip_p"] <= 0.05
    )


def main() -> None:
    if AUDIT.exists():
        raise FileExistsError("Immutable artifact audit exists.")
    result = json.loads(RESULT.read_text(encoding="utf-8"))
    participant_rows = read_csv(PARTICIPANT_CSV)
    montage_rows = read_csv(MONTAGE_CSV)
    calibration_rows = read_csv(CALIBRATION_CSV)

    participants = {
        "ASME30": [str(value) for value in range(1, 11)],
        "Kojima2024B": list("BDFHJLN"),
    }
    participant_metrics = (
        "physical_effect",
        "binding_specificity",
        "nonocular_specificity",
        "identity_advantage",
        "mapping_alignment_specificity",
        "physical_alignment",
        "between_person_substitution",
    )
    participant_errors = []
    for dataset, participant_order in participants.items():
        for montage_n in (8, 16, 32, 64):
            rows = [
                row
                for row in participant_rows
                if row["dataset"] == dataset
                and int(row["montage_channels"]) == montage_n
            ]
            observed_order = [row["participant"] for row in rows]
            if observed_order != participant_order:
                raise RuntimeError(
                    f"Participant order mismatch: {dataset}/{montage_n}"
                )
            for metric in participant_metrics:
                expected = result["montage_summaries"][dataset][
                    str(montage_n)
                ][metric]["participant_values"]
                participant_errors.extend(
                    error(row[metric], value)
                    for row, value in zip(rows, expected)
                )

    montage_errors = []
    for row in montage_rows:
        item = result["montage_summaries"][row["dataset"]][
            str(int(row["montage_channels"]))
        ][row["metric"]]
        montage_errors.extend(
            [
                error(row["mean"], item["mean"]),
                error(row["ci_low"], item["bootstrap_95_CI_mean"][0]),
                error(row["ci_high"], item["bootstrap_95_CI_mean"][1]),
                error(row["n_positive"], item["n_positive"]),
                error(row["n_total"], item["n_total"]),
                error(
                    row["p_one_sided_exact"],
                    item["one_sided_exact_signflip_p"],
                ),
            ]
        )

    calibration_errors = []
    for row in calibration_rows:
        item = result["calibration_summaries"][row["dataset"]][
            str(int(row["training_runs"]))
        ][row["metric"]]
        calibration_errors.extend(
            [
                error(row["mean"], item["mean"]),
                error(row["ci_low"], item["bootstrap_95_CI_mean"][0]),
                error(row["ci_high"], item["bootstrap_95_CI_mean"][1]),
                error(row["n_positive"], item["n_positive"]),
                error(row["n_total"], item["n_total"]),
                error(
                    row["p_one_sided_exact"],
                    item["one_sided_exact_signflip_p"],
                ),
            ]
        )

    required = {"ASME30": 8, "Kojima2024B": 6}
    summaries = result["montage_summaries"]
    calibration = result["calibration_summaries"]

    def both_montage(metric: str) -> bool:
        return all(
            scalar_support(summaries[d]["8"][metric], required[d])
            for d in required
        )

    def both_calibration(metric: str) -> bool:
        return all(
            scalar_support(calibration[d]["3"][metric], required[d])
            for d in required
        )

    recomputed_gates = {
        "W0_integrity": all(result["integrity"].values()),
        "W1_eight_channel_physical": both_montage("physical_effect"),
        "W2_eight_channel_binding": both_montage(
            "binding_specificity"
        ),
        "W3_eight_channel_nonocular": both_montage(
            "nonocular_specificity"
        ),
        "W4_eight_channel_identity": both_montage(
            "identity_advantage"
        ),
        "W5_eight_channel_retention": all(
            math.isfinite(value) and value >= 0.50
            for value in result["eight_channel_binding_retention"].values()
        )
        and all(
            summaries[d]["64"]["binding_specificity"]["mean"] > 0
            for d in required
        ),
        "W6_half_calibration_physical": both_calibration(
            "physical_effect"
        ),
        "W7_half_calibration_binding": both_calibration(
            "binding_specificity"
        ),
    }
    checks = {
        "participant_csv_matches_v2_json": (
            max(participant_errors, default=0.0) <= TOLERANCE
        ),
        "montage_csv_matches_v2_json": (
            max(montage_errors, default=0.0) <= TOLERANCE
        ),
        "calibration_csv_matches_v2_json": (
            max(calibration_errors, default=0.0) <= TOLERANCE
        ),
        "gates_recompute_exactly": recomputed_gates == result["gates"],
        "v1_lock_hash_matches_provenance": (
            sha256(LOCK_V1)
            == result["provenance"]["implementation_lock_sha256"]
        ),
        "v1_runner_hash_matches_provenance": (
            json.loads(LOCK_V1.read_text(encoding="utf-8"))["runner_sha256"]
            == result["provenance"]["runner_sha256"]
        ),
    }
    output = {
        "audit_id": "WZ-HIC-wearable-calibration-final-v1-artifact-audit-v2",
        "created_utc": datetime.now(timezone.utc).isoformat(),
        "status": "PASS" if all(checks.values()) else "FAIL",
        "scope": (
            "Artifact-level consistency of the final files after the write "
            "collision ended. The final decision JSON identifies the locked "
            "v1 runner and lock. This audit does not independently recompute "
            "EEG models."
        ),
        "checks": checks,
        "maximum_absolute_errors": {
            "participant_csv_vs_v2_json": max(
                participant_errors, default=0.0
            ),
            "montage_csv_vs_v2_json": max(montage_errors, default=0.0),
            "calibration_csv_vs_v2_json": max(
                calibration_errors, default=0.0
            ),
        },
        "recomputed_gates": recomputed_gates,
        "source_hashes": {
            "decision_json": sha256(RESULT),
            "participant_csv": sha256(PARTICIPANT_CSV),
            "montage_csv": sha256(MONTAGE_CSV),
            "calibration_csv": sha256(CALIBRATION_CSV),
            "implementation_lock_v1": sha256(LOCK_V1),
        },
        "interpretation": (
            "The final CSVs and the final v1 decision JSON agree exactly if "
            "this audit passes. The earlier failed v1 audit is retained as "
            "the collision diagnostic. A source-level audit is still "
            "required."
        ),
    }
    OUTDIR.mkdir(parents=True, exist_ok=True)
    AUDIT.write_text(
        json.dumps(output, ensure_ascii=False, indent=2) + "\n",
        encoding="utf-8",
    )
    print(json.dumps(output, ensure_ascii=False, indent=2))
    if output["status"] != "PASS":
        raise SystemExit(1)


if __name__ == "__main__":
    main()
```
