## Supplementary material for "Falsifiable substitution tests reveal task-structured neural evidence for auditory attention": Article_Analysis_Source_Code: audit_wz_hic_wearable_calibration_source_v1.py

```
#!/usr/bin/env python3
"""Independent source-level audit of primary wearable/calibration outcomes."""

from __future__ import annotations

import hashlib
import itertools
import json
import math
from datetime import datetime, timezone
from pathlib import Path

import numpy as np
from joblib import Parallel, delayed
from sklearn.covariance import ledoit_wolf
from threadpoolctl import threadpool_limits


ROOT = Path(__file__).resolve().parents[1]
PROTOCOL = (
    ROOT
    / "docs"
    / "wz_hic_wearable_calibration_source_audit_protocol_v1.json"
)
ASME_FEATURES = (
    ROOT
    / "data"
    / "derived"
    / "asme30_wz_hic_dataset_native_features_v2.npz"
)
KOJIMA_FEATURES = (
    ROOT
    / "data"
    / "derived"
    / "kojima2024b_wz_hic_confirmation_features_v1.npz"
)
DECISION = (
    ROOT
    / "results"
    / "wz_hic_cross_dataset"
    / "wearable_calibration_v1"
    / "wearable_calibration_decision_v1.json"
)
AUDIT = (
    ROOT
    / "results"
    / "wz_hic_cross_dataset"
    / "wearable_calibration_v1"
    / "independent_source_audit_v1.json"
)
ASME_SHA256 = (
    "e767e08dce8e46d1589203e76289a5acf64284166efc54325b945108e9559017"
)
KOJIMA_SHA256 = (
    "a3874f1b8dfb8af1d9329f95e4a6da9a0cb86935d1b17f294e7601c092cc32c9"
)
DECISION_SEMANTIC_SHA256 = (
    "5100658437336f44246c08f149e2bc87b82fd1ca91460fe47e75f4b6616e3390"
)
REPRODUCIBILITY_AMENDMENT = (
    ROOT
    / "docs"
    / "wz_hic_wearable_source_audit_reproducibility_amendment_v1.json"
)
TOLERANCE = 5e-6
HORIZON = 3
BOOTSTRAP_N = 20_000
BOOTSTRAP_SEED = 20260730
N_JOBS = 8
MONTAGE = ("Fz", "FCz", "C3", "Cz", "C4", "CPz", "Pz", "Oz")
ASME_SUBJECTS = tuple(range(1, 11))
KOJIMA_SUBJECTS = tuple("BDFHJLN")
ASME_PHYSICAL = (
    (17, 23, 5, 18, 20, 25, 21, 9, 15, 16),
    (1, 19, 4, 6, 7, 8, 10, 11, 12, 29),
    (26, 24, 3, 22, 2, 14, 13, 27, 28, 30),
)
ASME_WRONG = (
    (
        tuple(range(1, 31, 3)),
        tuple(range(2, 31, 3)),
        tuple(range(3, 31, 3)),
    ),
    (
        tuple(range(1, 11)),
        tuple(range(11, 21)),
        tuple(range(21, 31)),
    ),
)
KOJIMA_PHYSICAL = ((1, 2), (3, 4))
KOJIMA_WRONG = (((1, 3), (2, 4)), ((1, 4), (2, 3)))


def sha256(path: Path) -> str:
    digest = hashlib.sha256()
    with path.open("rb") as handle:
        for block in iter(lambda: handle.read(1024 * 1024), b""):
            digest.update(block)
    return digest.hexdigest()


def semantic_json_sha256(path: Path) -> str:
    """Hash all decision content except the new-run timestamp."""

    def clean(value):
        if isinstance(value, dict):
            return {
                key: clean(item)
                for key, item in value.items()
                if key != "created_utc"
            }
        if isinstance(value, list):
            return [clean(item) for item in value]
        return value

    payload = json.loads(path.read_text(encoding="utf-8"))
    canonical = json.dumps(
        clean(payload), sort_keys=True, separators=(",", ":"), allow_nan=True
    ).encode("utf-8")
    return hashlib.sha256(canonical).hexdigest()


def labels_for(
    command: np.ndarray,
    target: np.ndarray,
    groups,
    n_commands: int,
) -> np.ndarray:
    lookup = np.full(n_commands + 1, -1, dtype=int)
    members = []
    for group_id, group in enumerate(groups):
        lookup[list(group)] = group_id
        members.extend(int(value) for value in group)
    if sorted(members) != list(range(1, n_commands + 1)):
        raise RuntimeError("Mapping is not a partition.")
    output = np.full(len(command), "U", dtype="U1")
    is_target = command == target
    output[is_target] = "T"
    output[(lookup[command] == lookup[target]) & ~is_target] = "S"
    return output


def residualize(
    train_x: np.ndarray,
    test_x: np.ndarray,
    train_command: np.ndarray,
    train_target: np.ndarray,
    test_command: np.ndarray,
    n_commands: int,
) -> tuple[np.ndarray, np.ndarray]:
    train_out = np.asarray(train_x, dtype=float).copy()
    test_out = np.asarray(test_x, dtype=float).copy()
    non_target = train_command != train_target
    for value in range(1, n_commands + 1):
        ids = (train_command == value) & non_target
        if not np.any(ids):
            raise RuntimeError(f"No command baseline: {value}")
        center = np.mean(train_x[ids], axis=0)
        train_out[train_command == value] -= center
        test_out[test_command == value] -= center
    return train_out, test_out


def complete_trials(
    command: np.ndarray, trial: np.ndarray, n_commands: int
) -> list[np.ndarray]:
    output = []
    for trial_id in sorted(int(value) for value in np.unique(trial)):
        trial_ids = np.flatnonzero(trial == trial_id)
        chunks = []
        for command_value in range(1, n_commands + 1):
            ids = trial_ids[command[trial_ids] == command_value]
            if len(ids) < HORIZON:
                chunks = []
                break
            chunks.append(ids[:HORIZON])
        if len(chunks) == n_commands:
            output.append(np.concatenate(chunks))
    return output


def fit_lda_scores(
    train_x: np.ndarray,
    train_labels: np.ndarray,
    test_x: np.ndarray,
) -> np.ndarray:
    keep = np.isin(train_labels, ("S", "U"))
    x = np.asarray(train_x[keep], dtype=float)
    y = (train_labels[keep] == "S").astype(int)
    if set(np.unique(y)) != {0, 1}:
        raise RuntimeError("Binary labels missing.")
    mean = np.mean(x, axis=0)
    scale = np.std(x, axis=0, ddof=1)
    scale[~np.isfinite(scale) | (scale < 1e-12)] = 1.0
    z = (x - mean) / scale
    mean_0 = np.mean(z[y == 0], axis=0)
    mean_1 = np.mean(z[y == 1], axis=0)
    centered = np.vstack(
        (z[y == 0] - mean_0, z[y == 1] - mean_1)
    )
    covariance, _ = ledoit_wolf(centered, assume_centered=True)
    delta = mean_1 - mean_0
    try:
        direction = np.linalg.solve(covariance, delta)
    except np.linalg.LinAlgError:
        direction = np.linalg.pinv(covariance, hermitian=True) @ delta
    intercept = float(-0.5 * (mean_1 + mean_0) @ direction)

    def decision(values: np.ndarray) -> np.ndarray:
        return ((np.asarray(values, dtype=float) - mean) / scale) @ (
            direction
        ) + intercept

    training_score = decision(x)
    training_mean = float(np.mean(training_score))
    training_sd = float(np.std(training_score, ddof=1))
    if not np.isfinite(training_sd) or training_sd <= 1e-12:
        raise RuntimeError("Degenerate training score.")
    return (decision(test_x) - training_mean) / training_sd


def cosine(left: np.ndarray, right: np.ndarray) -> float:
    denominator = float(np.linalg.norm(left) * np.linalg.norm(right))
    if denominator <= 1e-12:
        raise RuntimeError("Degenerate direction.")
    return float(np.vdot(left, right) / denominator)


def heldout_trial_effects(
    score: np.ndarray,
    command: np.ndarray,
    target: np.ndarray,
    selections: list[np.ndarray],
    groups,
    n_commands: int,
) -> list[float]:
    expected_s = (len(groups[0]) - 1) * HORIZON
    expected_u = (n_commands - len(groups[0])) * HORIZON
    output = []
    for selected in selections:
        labels = labels_for(
            command[selected], target[selected], groups, n_commands
        )
        counts = tuple(
            int(np.sum(labels == value)) for value in ("T", "S", "U")
        )
        if counts != (HORIZON, expected_s, expected_u):
            raise RuntimeError(f"Unexpected T/S/U: {counts}")
        output.append(
            float(np.mean(score[selected][labels == "S"]))
            - float(np.mean(score[selected][labels == "U"]))
        )
    return output


def direction_pair(
    train_x: np.ndarray,
    train_labels: np.ndarray,
    test_x: np.ndarray,
    test_command: np.ndarray,
    test_target: np.ndarray,
    selections: list[np.ndarray],
    groups,
    n_commands: int,
) -> tuple[np.ndarray, np.ndarray, float]:
    keep = np.isin(train_labels, ("S", "U"))
    x = np.asarray(train_x[keep], dtype=float)
    labels = train_labels[keep]
    mean = np.mean(x, axis=0)
    scale = np.std(x, axis=0, ddof=1)
    scale[~np.isfinite(scale) | (scale <= 1e-12)] = 1.0
    z_train = (x - mean) / scale
    training = np.mean(z_train[labels == "S"], axis=0) - np.mean(
        z_train[labels == "U"], axis=0
    )
    z_test = (np.asarray(test_x, dtype=float) - mean) / scale
    heldout_trials = []
    for selected in selections:
        test_labels = labels_for(
            test_command[selected],
            test_target[selected],
            groups,
            n_commands,
        )
        heldout_trials.append(
            np.mean(z_test[selected][test_labels == "S"], axis=0)
            - np.mean(z_test[selected][test_labels == "U"], axis=0)
        )
    heldout = np.mean(np.stack(heldout_trials), axis=0)
    return training, heldout, cosine(training, heldout)


def mapping(
    train_x: np.ndarray,
    test_x: np.ndarray,
    train_command: np.ndarray,
    train_target: np.ndarray,
    test_command: np.ndarray,
    test_target: np.ndarray,
    selections: list[np.ndarray],
    groups,
    n_commands: int,
) -> dict:
    train_labels = labels_for(
        train_command, train_target, groups, n_commands
    )
    score = fit_lda_scores(train_x, train_labels, test_x)
    trial_effects = heldout_trial_effects(
        score,
        test_command,
        test_target,
        selections,
        groups,
        n_commands,
    )
    training, heldout, alignment = direction_pair(
        train_x,
        train_labels,
        test_x,
        test_command,
        test_target,
        selections,
        groups,
        n_commands,
    )
    return {
        "trial_effects": trial_effects,
        "effect": float(np.mean(trial_effects)),
        "training_direction": training,
        "heldout_direction": heldout,
        "alignment": alignment,
    }


def feature_matrix(
    arrays: dict, mask: np.ndarray, *, eog: bool = False
) -> np.ndarray:
    if eog:
        bins = arrays["eog_bins"][mask, :, 1:]
    else:
        names = [str(value) for value in arrays["channel_names"]]
        indices = np.asarray([names.index(name) for name in MONTAGE])
        bins = arrays["eeg_bins"][mask][:, indices, 1:]
    return bins.reshape(bins.shape[0], -1)


def analyze_partition(
    arrays: dict,
    train: np.ndarray,
    test: np.ndarray,
    physical_groups,
    wrong_groups,
    n_commands: int,
    *,
    include_eog: bool,
) -> dict:
    train_command = arrays["command"][train].astype(int)
    train_target = arrays["target"][train].astype(int)
    test_command = arrays["command"][test].astype(int)
    test_target = arrays["target"][test].astype(int)
    selections = complete_trials(
        test_command, arrays["trial"][test].astype(int), n_commands
    )
    train_x, test_x = residualize(
        feature_matrix(arrays, train),
        feature_matrix(arrays, test),
        train_command,
        train_target,
        test_command,
        n_commands,
    )
    physical = mapping(
        train_x,
        test_x,
        train_command,
        train_target,
        test_command,
        test_target,
        selections,
        physical_groups,
        n_commands,
    )
    wrong = [
        mapping(
            train_x,
            test_x,
            train_command,
            train_target,
            test_command,
            test_target,
            selections,
            groups,
            n_commands,
        )
        for groups in wrong_groups
    ]
    eog_effect = None
    if include_eog:
        train_eog, test_eog = residualize(
            feature_matrix(arrays, train, eog=True),
            feature_matrix(arrays, test, eog=True),
            train_command,
            train_target,
            test_command,
            n_commands,
        )
        eog_effect = mapping(
            train_eog,
            test_eog,
            train_command,
            train_target,
            test_command,
            test_target,
            selections,
            physical_groups,
            n_commands,
        )["effect"]
    return {
        "n_test_trials": len(selections),
        "physical": physical,
        "wrong": wrong,
        "eog_effect": eog_effect,
    }


def collapse(partition: dict) -> dict:
    wrong_effects = [item["effect"] for item in partition["wrong"]]
    wrong_alignments = [item["alignment"] for item in partition["wrong"]]
    return {
        "physical_effect": partition["physical"]["effect"],
        "binding_specificity": (
            partition["physical"]["effect"] - max(wrong_effects)
        ),
        "nonocular_specificity": (
            None
            if partition["eog_effect"] is None
            else partition["physical"]["effect"] - partition["eog_effect"]
        ),
        "physical_alignment": partition["physical"]["alignment"],
        "mapping_alignment_specificity": (
            partition["physical"]["alignment"] - max(wrong_alignments)
        ),
    }


def full_asme(arrays: dict) -> list[dict]:
    accepted = arrays["accepted"].astype(bool)

    def one(subject: int) -> dict:
        train = (
            (arrays["subject"] == subject)
            & (arrays["task"] == "offline")
            & accepted
        )
        test = (
            (arrays["subject"] == subject)
            & (arrays["task"] == "online")
            & accepted
        )
        partition = analyze_partition(
            arrays,
            train,
            test,
            ASME_PHYSICAL,
            ASME_WRONG,
            30,
            include_eog=True,
        )
        return {
            "participant": str(subject),
            "partition": partition,
            **collapse(partition),
        }

    with threadpool_limits(limits=1):
        records = Parallel(n_jobs=N_JOBS, prefer="threads")(
            delayed(one)(subject) for subject in ASME_SUBJECTS
        )
    for index, record in enumerate(records):
        heldout = record["partition"]["physical"]["heldout_direction"]
        between = np.mean(
            [
                cosine(
                    heldout,
                    donor["partition"]["physical"]["training_direction"],
                )
                for donor_index, donor in enumerate(records)
                if donor_index != index
            ]
        )
        record["identity_advantage"] = (
            record["physical_alignment"] - between
        )
    return records


def full_kojima(arrays: dict) -> list[dict]:
    accepted = arrays["accepted"].astype(bool)

    def one(subject: str) -> dict:
        base = (
            (arrays["subject"] == subject)
            & (arrays["task"] == "2stream")
            & accepted
        )
        runs = sorted(int(value) for value in np.unique(arrays["run"][base]))
        folds = []
        for heldout in runs:
            partition = analyze_partition(
                arrays,
                base & (arrays["run"] != heldout),
                base & (arrays["run"] == heldout),
                KOJIMA_PHYSICAL,
                KOJIMA_WRONG,
                4,
                include_eog=True,
            )
            folds.append(partition)
        physical_values = [
            value
            for fold in folds
            for value in fold["physical"]["trial_effects"]
        ]
        wrong_values = [
            [
                value
                for fold in folds
                for value in fold["wrong"][wrong_index]["trial_effects"]
            ]
            for wrong_index in range(2)
        ]
        eog_values = [
            fold["eog_effect"]
            for fold in folds
            for _ in fold["physical"]["trial_effects"]
        ]
        physical_effect = float(np.mean(physical_values))
        wrong_effects = [float(np.mean(values)) for values in wrong_values]
        physical_alignment = float(
            np.mean([fold["physical"]["alignment"] for fold in folds])
        )
        wrong_alignments = [
            float(
                np.mean(
                    [
                        fold["wrong"][wrong_index]["alignment"]
                        for fold in folds
                    ]
                )
            )
            for wrong_index in range(2)
        ]
        return {
            "participant": subject,
            "folds": folds,
            "physical_effect": physical_effect,
            "binding_specificity": physical_effect - max(wrong_effects),
            "nonocular_specificity": physical_effect - float(
                np.mean(eog_values)
            ),
            "physical_alignment": physical_alignment,
            "mapping_alignment_specificity": (
                physical_alignment - max(wrong_alignments)
            ),
        }

    with threadpool_limits(limits=1):
        records = Parallel(n_jobs=N_JOBS, prefer="threads")(
            delayed(one)(subject) for subject in KOJIMA_SUBJECTS
        )
    for subject_index, record in enumerate(records):
        between = []
        for fold_index, fold in enumerate(record["folds"]):
            heldout = fold["physical"]["heldout_direction"]
            between.append(
                np.mean(
                    [
                        cosine(
                            heldout,
                            donor["folds"][fold_index]["physical"][
                                "training_direction"
                            ],
                        )
                        for donor_index, donor in enumerate(records)
                        if donor_index != subject_index
                    ]
                )
            )
        record["identity_advantage"] = (
            record["physical_alignment"] - float(np.mean(between))
        )
    return records


def calibration_asme(arrays: dict) -> list[dict]:
    accepted = arrays["accepted"].astype(bool)

    def one(subject: int) -> dict:
        offline = (
            (arrays["subject"] == subject)
            & (arrays["task"] == "offline")
            & accepted
        )
        test = (
            (arrays["subject"] == subject)
            & (arrays["task"] == "online")
            & accepted
        )
        runs = sorted(int(value) for value in np.unique(arrays["run"][offline]))
        cells = []
        for subset in itertools.combinations(runs, 3):
            partition = analyze_partition(
                arrays,
                offline & np.isin(arrays["run"], subset),
                test,
                ASME_PHYSICAL,
                ASME_WRONG,
                30,
                include_eog=False,
            )
            cells.append(collapse(partition))
        if len(cells) != 20:
            raise RuntimeError("ASME C(6,3) enumeration failed.")
        return {
            "participant": str(subject),
            "physical_effect": float(
                np.mean([cell["physical_effect"] for cell in cells])
            ),
            "binding_specificity": float(
                np.mean([cell["binding_specificity"] for cell in cells])
            ),
        }

    with threadpool_limits(limits=1):
        return Parallel(n_jobs=N_JOBS, prefer="threads")(
            delayed(one)(subject) for subject in ASME_SUBJECTS
        )


def calibration_kojima(arrays: dict) -> list[dict]:
    accepted = arrays["accepted"].astype(bool)

    def one(subject: str) -> dict:
        base = (
            (arrays["subject"] == subject)
            & (arrays["task"] == "2stream")
            & accepted
        )
        runs = sorted(int(value) for value in np.unique(arrays["run"][base]))
        cells = []
        for heldout in runs:
            available = [run for run in runs if run != heldout]
            test = base & (arrays["run"] == heldout)
            for subset in itertools.combinations(available, 3):
                partition = analyze_partition(
                    arrays,
                    base & np.isin(arrays["run"], subset),
                    test,
                    KOJIMA_PHYSICAL,
                    KOJIMA_WRONG,
                    4,
                    include_eog=False,
                )
                cells.append(collapse(partition))
        if len(cells) != 60:
            raise RuntimeError("Kojima 6*C(5,3) enumeration failed.")
        return {
            "participant": subject,
            "physical_effect": float(
                np.mean([cell["physical_effect"] for cell in cells])
            ),
            "binding_specificity": float(
                np.mean([cell["binding_specificity"] for cell in cells])
            ),
        }

    with threadpool_limits(limits=1):
        return Parallel(n_jobs=N_JOBS, prefer="threads")(
            delayed(one)(subject) for subject in KOJIMA_SUBJECTS
        )


def bootstrap_ci(values: np.ndarray) -> list[float]:
    rng = np.random.default_rng(BOOTSTRAP_SEED)
    ids = rng.integers(0, len(values), size=(BOOTSTRAP_N, len(values)))
    return [
        float(value)
        for value in np.quantile(values[ids].mean(axis=1), [0.025, 0.975])
    ]


def exact_p(values: np.ndarray) -> float:
    observed = float(np.mean(values))
    permuted = [
        float(np.mean(values * np.asarray(signs)))
        for signs in itertools.product((-1.0, 1.0), repeat=len(values))
    ]
    return float(np.mean(np.asarray(permuted) >= observed - 1e-15))


def summarize(records: list[dict], metric: str) -> dict:
    values = np.asarray([record[metric] for record in records], dtype=float)
    return {
        "mean": float(np.mean(values)),
        "bootstrap_95_CI_mean": bootstrap_ci(values),
        "n_positive": int(np.sum(values > 0)),
        "n_total": len(values),
        "one_sided_exact_signflip_p": exact_p(values),
        "participant_values": [float(value) for value in values],
    }


def supported(item: dict, required: int) -> bool:
    return (
        item["mean"] > 0
        and item["bootstrap_95_CI_mean"][0] > 0
        and item["n_positive"] >= required
        and item["one_sided_exact_signflip_p"] <= 0.05
    )


def numeric_errors(observed: dict, expected: dict) -> list[float]:
    return [
        abs(observed["mean"] - expected["mean"]),
        abs(
            observed["bootstrap_95_CI_mean"][0]
            - expected["bootstrap_95_CI_mean"][0]
        ),
        abs(
            observed["bootstrap_95_CI_mean"][1]
            - expected["bootstrap_95_CI_mean"][1]
        ),
        abs(observed["n_positive"] - expected["n_positive"]),
        abs(observed["n_total"] - expected["n_total"]),
        abs(
            observed["one_sided_exact_signflip_p"]
            - expected["one_sided_exact_signflip_p"]
        ),
        *[
            abs(left - right)
            for left, right in zip(
                observed["participant_values"],
                expected["participant_values"],
            )
        ],
    ]


def main() -> None:
    if AUDIT.exists():
        raise FileExistsError("Immutable source audit exists.")
    amendment = json.loads(
        REPRODUCIBILITY_AMENDMENT.read_text(encoding="utf-8")
    )
    if (
        amendment["decision_semantic_sha256"]
        != DECISION_SEMANTIC_SHA256
        or sha256(ROOT / amendment["source_result_generator"]["path"])
        != amendment["source_result_generator"]["sha256"]
    ):
        raise RuntimeError("Wearable reproducibility amendment mismatch.")
    observed_hashes = {
        "asme": sha256(ASME_FEATURES),
        "kojima": sha256(KOJIMA_FEATURES),
        "decision_semantic": semantic_json_sha256(DECISION),
    }
    expected_hashes = {
        "asme": ASME_SHA256,
        "kojima": KOJIMA_SHA256,
        "decision_semantic": DECISION_SEMANTIC_SHA256,
    }
    if observed_hashes != expected_hashes:
        raise RuntimeError(
            f"Frozen source mismatch: {observed_hashes} != {expected_hashes}"
        )
    expected = json.loads(DECISION.read_text(encoding="utf-8"))
    with np.load(ASME_FEATURES, allow_pickle=True) as source:
        asme = {key: source[key] for key in source.files}
    with np.load(KOJIMA_FEATURES, allow_pickle=True) as source:
        kojima = {key: source[key] for key in source.files}

    full = {
        "ASME30": full_asme(asme),
        "Kojima2024B": full_kojima(kojima),
    }
    calibration = {
        "ASME30": calibration_asme(asme),
        "Kojima2024B": calibration_kojima(kojima),
    }
    full_metrics = (
        "physical_effect",
        "binding_specificity",
        "nonocular_specificity",
        "identity_advantage",
        "mapping_alignment_specificity",
    )
    calibration_metrics = ("physical_effect", "binding_specificity")
    summaries = {
        dataset: {
            metric: summarize(records, metric)
            for metric in full_metrics
        }
        for dataset, records in full.items()
    }
    calibration_summaries = {
        dataset: {
            metric: summarize(records, metric)
            for metric in calibration_metrics
        }
        for dataset, records in calibration.items()
    }
    errors = []
    for dataset in ("ASME30", "Kojima2024B"):
        for metric in full_metrics:
            errors.extend(
                numeric_errors(
                    summaries[dataset][metric],
                    expected["montage_summaries"][dataset]["8"][metric],
                )
            )
        for metric in calibration_metrics:
            errors.extend(
                numeric_errors(
                    calibration_summaries[dataset][metric],
                    expected["calibration_summaries"][dataset]["3"][metric],
                )
            )
    required = {"ASME30": 8, "Kojima2024B": 6}

    def both_full(metric: str) -> bool:
        return all(
            supported(summaries[d][metric], required[d]) for d in required
        )

    def both_calibration(metric: str) -> bool:
        return all(
            supported(calibration_summaries[d][metric], required[d])
            for d in required
        )

    recomputed_gates = {
        "W1_eight_channel_physical": both_full("physical_effect"),
        "W2_eight_channel_binding": both_full("binding_specificity"),
        "W3_eight_channel_nonocular": both_full("nonocular_specificity"),
        "W4_eight_channel_identity": both_full("identity_advantage"),
        "W6_half_calibration_physical": both_calibration(
            "physical_effect"
        ),
        "W7_half_calibration_binding": both_calibration(
            "binding_specificity"
        ),
    }
    expected_gates = {
        key: expected["gates"][key] for key in recomputed_gates
    }
    max_error = max(errors, default=0.0)
    checks = {
        "frozen_hashes": observed_hashes == expected_hashes,
        "participant_and_summary_values": max_error <= TOLERANCE,
        "gates_exact": recomputed_gates == expected_gates,
        "asme_full_participants": len(full["ASME30"]) == 10,
        "kojima_full_participants": len(full["Kojima2024B"]) == 7,
        "asme_three_run_participants": len(calibration["ASME30"]) == 10,
        "kojima_three_run_participants": (
            len(calibration["Kojima2024B"]) == 7
        ),
    }
    output = {
        "audit_id": "WZ-HIC-wearable-calibration-independent-source-audit-v1",
        "created_utc": datetime.now(timezone.utc).isoformat(),
        "status": "PASS" if all(checks.values()) else "FAIL",
        "checks": checks,
        "maximum_absolute_error": max_error,
        "tolerance": TOLERANCE,
        "recomputed_gates": recomputed_gates,
        "expected_gates": expected_gates,
        "eight_channel_summaries": summaries,
        "three_run_summaries": calibration_summaries,
        "provenance": {
            "audit_protocol_sha256": sha256(PROTOCOL),
            "auditor_sha256": sha256(Path(__file__)),
            "asme_features_sha256": observed_hashes["asme"],
            "kojima_features_sha256": observed_hashes["kojima"],
            "decision_sha256": sha256(DECISION),
            "decision_semantic_sha256": observed_hashes[
                "decision_semantic"
            ],
            "reproducibility_amendment_sha256": sha256(
                REPRODUCIBILITY_AMENDMENT
            ),
        },
        "independence_statement": (
            "The auditor imports no project runner or core. It independently "
            "implements labels, command residuals, complete trials, "
            "Ledoit-Wolf LDA, held-out scores, neural directions, cross-person "
            "substitution, all fixed three-run subsets, bootstrap, sign flips, "
            "and gates."
        ),
    }
    AUDIT.write_text(
        json.dumps(output, ensure_ascii=False, indent=2) + "\n",
        encoding="utf-8",
    )
    print(
        json.dumps(
            {
                "status": output["status"],
                "checks": output["checks"],
                "maximum_absolute_error": max_error,
                "recomputed_gates": recomputed_gates,
                "audit_sha256": sha256(AUDIT),
            },
            ensure_ascii=False,
            indent=2,
        )
    )
    if output["status"] != "PASS":
        raise SystemExit(1)


if __name__ == "__main__":
    main()
```
