## Supplementary material for "Falsifiable substitution tests reveal task-structured neural evidence for auditory attention": Article_Analysis_Source_Code: build_article_claim_ledger_v1.py

```
"""Create a machine-readable claim ledger for the unified article draft."""

from __future__ import annotations

import hashlib
import json
from datetime import datetime, timezone
from pathlib import Path
from typing import Any


ROOT = Path(__file__).resolve().parents[1]
OUT = ROOT / "manuscript_archive_v1"
JSON_OUT = OUT / "claim_ledger_v1.json"
MD_OUT = OUT / "claim_ledger_v1.md"


def sha256(path: Path) -> str:
    digest = hashlib.sha256()
    with path.open("rb") as stream:
        for chunk in iter(lambda: stream.read(1024 * 1024), b""):
            digest.update(chunk)
    return digest.hexdigest()


def read_json(relative: str) -> Any:
    return json.loads((ROOT / relative).read_text(encoding="utf-8"))


def get_path(data: Any, path: str) -> Any:
    value = data
    for token in path.split("."):
        if token.isdigit() and isinstance(value, list):
            value = value[int(token)]
        else:
            value = value[token]
    return value


def main() -> None:
    OUT.mkdir(parents=True, exist_ok=True)
    source_cache: dict[str, Any] = {}

    def add(
        claim_id: str,
        section: str,
        wording: str,
        source: str,
        json_path: str,
        standing: str,
        guardrail: str,
    ) -> dict[str, Any]:
        if source not in source_cache:
            source_cache[source] = read_json(source)
        value = get_path(source_cache[source], json_path)
        return {
            "claim_id": claim_id,
            "section": section,
            "wording": wording,
            "source": source,
            "source_sha256": sha256(ROOT / source),
            "json_path": json_path,
            "value": value,
            "standing": standing,
            "guardrail": guardrail,
        }

    projection = "results/multimodal_projection_breadth_synthesis/synthesis.json"
    kite = "results/kite_sat/article_evidence_synthesis_v1/evidence_synthesis_v1.json"
    target = (
        "results/wz_hic_cross_dataset/target_free_mechanism_v1/"
        "target_free_decision_v1.json"
    )
    asme_primary = (
        "results/wz_hic_asme30/dataset_native_v2/"
        "dataset_native_decision_v2.json"
    )
    continuous = (
        "results/wz_hic_cross_dataset/continuous_listener_transport_v1/"
        "continuous_listener_transport_decision_v1.json"
    )
    shared = (
        "results/wz_hic_cross_dataset/continuous_shared_template_v1/"
        "continuous_shared_template_decision_v1.json"
    )
    individual = (
        "results/wz_hic_cross_dataset/individual_geometry_synthesis_v1/"
        "synthesis_v1.json"
    )
    residual = (
        "results/wz_hic_cross_dataset/command_identity_residual_v1/"
        "residual_decision_v1.json"
    )
    wearable = (
        "results/wz_hic_cross_dataset/wearable_calibration_v1/"
        "wearable_calibration_decision_v1.json"
    )

    claims = [
        add(
            "PB-N",
            "Results 1",
            "Three locked projection-breadth tests included 41 participants.",
            projection,
            "independent_locked_synthesis.participants_total",
            "secondary locked-study synthesis",
            "Do not include the eight discovery participants.",
        ),
        add(
            "PB-POS",
            "Results 1",
            "Thirty-two of 41 participants improved.",
            projection,
            "independent_locked_synthesis.participants_improved_total",
            "secondary locked-study synthesis",
            "Participant count, not independent-study count.",
        ),
        add(
            "PB-GAIN",
            "Results 1",
            "The study-equal accuracy gain was 0.020117.",
            projection,
            "independent_locked_synthesis.study_equal_mean_gain",
            "secondary synthesis",
            "A modest accuracy gain; not a theorem or mechanism estimate.",
        ),
        add(
            "PB-CI",
            "Results 1",
            "Projection-breadth gain 95% interval.",
            projection,
            "independent_locked_synthesis.study_equal_stratified_bootstrap_95",
            "secondary synthesis",
            "Stratified participant bootstrap.",
        ),
        add(
            "PB-CONJ",
            "Results 1",
            "Replicability-conjunction P was 0.011719.",
            projection,
            "independent_locked_synthesis.replicability_conjunction_p",
            "conservative conjunction",
            "Maximum of the three locked one-sided P values.",
        ),
        add(
            "KITE-CONF",
            "Results 1",
            "Participant-disjoint KITE-ST bank gain was 0.054398 AUC.",
            kite,
            "cohort_endpoints.2.AKB_minus_comparator",
            "within-site independent-participant confirmation",
            "Not an independent-site confirmation.",
        ),
        add(
            "KITE-TOPO",
            "Results 1",
            "Physical topology exceeded 15 capacity-matched mappings by 0.077836 AUC.",
            kite,
            "capacity_matched_topology_mechanism.mean_identity_minus_15_controls_auc",
            "post-primary frozen mechanism test",
            "Reliability-profile stability subclaim failed and must not be implied.",
        ),
        add(
            "CONT-N",
            "Results 2",
            "Continuous listener transport included 43 participants.",
            continuous,
            "participant_count",
            "frozen cross-dataset experiment",
            "Three previously available public datasets, not pristine unseen datasets.",
        ),
        add(
            "CONT-DIRECTIONS",
            "Results 2",
            "There were 86 A-to-B or B-to-A transport directions.",
            continuous,
            "direction_count",
            "frozen cross-dataset experiment",
            "Two directions per participant; not 86 independent participants.",
        ),
        add(
            "F1",
            "Results 2",
            "Static own-listener profile rank advantage was 0.167623.",
            continuous,
            "cross_dataset_summaries.F1_fingerprint_listener_rank_advantage.equal_dataset_mean",
            "frozen primary endpoint",
            "KITE-ST dataset-specific effect is null; use 'cross-dataset' only.",
        ),
        add(
            "F2",
            "Results 2",
            "Static physical-mapping profile rank advantage was 0.258611.",
            continuous,
            "cross_dataset_summaries.F2_fingerprint_physical_mapping_rank_advantage.equal_dataset_mean",
            "frozen primary endpoint",
            "KITE-ST dataset-specific effect is uncertain.",
        ),
        add(
            "P1",
            "Results 2",
            "Held-speech own-listener versus donor rank advantage was 0.096895.",
            continuous,
            "cross_dataset_summaries.P1_predictive_listener_rank_advantage.equal_dataset_mean",
            "frozen primary endpoint",
            "Rank advantage, not classification-accuracy gain.",
        ),
        add(
            "P1-CI",
            "Results 2",
            "Held-speech listener advantage 95% interval.",
            continuous,
            "cross_dataset_summaries.P1_predictive_listener_rank_advantage.stratified_bootstrap_95",
            "frozen primary endpoint",
            "Three datasets weighted equally.",
        ),
        add(
            "P2",
            "Results 2",
            "Held-speech physical versus wrong-mapping rank advantage was 0.137109.",
            continuous,
            "cross_dataset_summaries.P2_predictive_physical_mapping_rank_advantage.equal_dataset_mean",
            "frozen primary endpoint",
            "Rank advantage, not classification-accuracy gain.",
        ),
        add(
            "P2-CI",
            "Results 2",
            "Held-speech mapping advantage 95% interval.",
            continuous,
            "cross_dataset_summaries.P2_predictive_physical_mapping_rank_advantage.stratified_bootstrap_95",
            "frozen primary endpoint",
            "Three datasets weighted equally.",
        ),
        add(
            "G1",
            "Results 2",
            "Own listener exceeded leave-one-listener mean-profile template by 0.122523.",
            shared,
            "cross_dataset_summaries.G1_own_vs_mean_profile_rank_advantage.equal_dataset_mean",
            "frozen strong comparator",
            "Supports evidence ranking, not a universal AUC gain.",
        ),
        add(
            "G2",
            "Results 2",
            "Own listener exceeded leave-one-listener mean-weight template by 0.121947.",
            shared,
            "cross_dataset_summaries.G2_own_vs_mean_weight_rank_advantage.equal_dataset_mean",
            "frozen strong comparator",
            "Supports evidence ranking, not a universal AUC gain.",
        ),
        add(
            "TF-KOJ",
            "Results 3",
            "Kojima target-free parent-error AUC was 0.967772.",
            target,
            "summaries.Kojima2024B.target_free_absolute.mean",
            "frozen post-primary mechanism audit",
            "Oracle target removal; not deployable online.",
        ),
        add(
            "TF-ASME",
            "Results 3",
            "ASME30 target-free parent-error AUC was 0.965432.",
            target,
            "summaries.ASME30.target_free_absolute.mean",
            "frozen post-primary mechanism audit",
            "Oracle target removal; not deployable online.",
        ),
        add(
            "MAP-KOJ",
            "Results 3",
            "Kojima true-binding specificity was 0.399618 AUC.",
            target,
            "summaries.Kojima2024B.mapping_specificity.mean",
            "frozen mechanism gate",
            "Physical grouping compared with matched wrong pairings.",
        ),
        add(
            "MAP-ASME",
            "Results 3",
            "ASME30 true-binding specificity was 0.240547 AUC.",
            target,
            "summaries.ASME30.mapping_specificity.mean",
            "frozen mechanism gate",
            "Physical grouping compared with matched wrong pairings.",
        ),
        add(
            "EOG-KOJ",
            "Results 3",
            "Kojima EEG-minus-EOG target-free AUC was 0.322057.",
            target,
            "summaries.Kojima2024B.nonocular.mean",
            "frozen mechanism gate",
            "EOG control does not exclude every possible artifact.",
        ),
        add(
            "EOG-ASME",
            "Results 3",
            "ASME30 EEG-minus-EOG target-free AUC was 0.317937.",
            target,
            "summaries.ASME30.nonocular.mean",
            "frozen mechanism gate",
            "EOG control does not exclude every possible artifact.",
        ),
        add(
            "EVENT-ASME",
            "Results 4",
            "ASME cross-phase physical effect was 0.935083.",
            individual,
            "dataset_summaries.ASME30.geometry.physical_crossphase.mean",
            "dataset-specific frozen transport",
            "Standardized effect, not AUC.",
        ),
        add(
            "EVENT-KOJ",
            "Results 4",
            "Kojima cross-run physical effect was 1.014133.",
            individual,
            "dataset_summaries.Kojima2024B.geometry.physical_crossrun.mean",
            "dataset-specific frozen transport",
            "Standardized effect, not AUC.",
        ),
        add(
            "EVENT-POS",
            "Results 4",
            "Physical, binding, and nonocular effects were positive in 17/17 participants.",
            individual,
            "participant_positive_counts_descriptive",
            "descriptive cross-study count",
            "Do not perform pooled 17-person inference.",
        ),
        add(
            "RESID-STATUS",
            "Results 4",
            "Training-only command residual passed all nine frozen gates.",
            residual,
            "command_identity_explanation_disfavored",
            "frozen post-primary control",
            "Residualization uses only outer-training nontarget events.",
        ),
        add(
            "WEAR-CRITERIA",
            "Results 5",
            "Seven of eight wearable/calibration criteria were satisfied.",
            wearable,
            "gates",
            "frozen robustness boundary",
            "W4 eight-channel listener-specific consistency failed.",
        ),
        add(
            "WEAR-RET-ASME",
            "Results 5",
            "ASME eight-channel binding retention was 0.590735.",
            wearable,
            "eight_channel_binding_retention.ASME30",
            "frozen robustness analysis",
            "Ratio relative to 64-channel binding effect.",
        ),
        add(
            "WEAR-RET-KOJ",
            "Results 5",
            "Kojima eight-channel binding retention was 0.768500.",
            wearable,
            "eight_channel_binding_retention.Kojima2024B",
            "frozen robustness analysis",
            "Ratio relative to 64-channel binding effect.",
        ),
        add(
            "ASME-EVAL6-N",
            "Methods 3",
            "ASME30 participant 6 contributed 15 flat-decoder test trials.",
            asme_primary,
            "participants.6.metrics.EEG_physical.n_trials",
            "outcome-defined AUC evaluability; reported transparently",
            "Trial count alone does not determine AUC evaluability.",
        ),
        add(
            "ASME-EVAL6-COR",
            "Methods 3",
            "All 15 flat predictions for ASME30 participant 6 were parent-correct.",
            asme_primary,
            "participants.6.metrics.EEG_physical.n_parent_correct",
            "outcome-defined AUC evaluability; reported transparently",
            "This perfect parent-level count does not imply perfect child-command decoding.",
        ),
        add(
            "ASME-EVAL6-ERR",
            "Methods 3",
            "ASME30 participant 6 had zero parent-error trials, so parent-error AUC was undefined.",
            asme_primary,
            "participants.6.metrics.EEG_physical.n_parent_error",
            "outcome-defined AUC evaluability; reported transparently",
            "Undefined AUC is not missing data; the exploratory all-participant mapping contrast is reported separately in SI Appendix.",
        ),
    ]

    output = {
        "schema": "article-unified-claim-ledger-v1",
        "created_utc": datetime.now(timezone.utc).isoformat(),
        "purpose": (
            "Bind every quantitative main-text claim to an authoritative frozen "
            "artifact and an explicit interpretation boundary."
        ),
        "claims": claims,
    }
    JSON_OUT.write_text(
        json.dumps(output, ensure_ascii=False, indent=2) + "\n",
        encoding="utf-8",
    )

    lines = [
        "# Unified article claim ledger v1",
        "",
        "Each row binds a manuscript claim to a frozen source artifact. "
        "The guardrail is part of the claim and must travel with it.",
        "",
        "| ID | Section | Value | Source path | Standing | Guardrail |",
        "|---|---|---:|---|---|---|",
    ]
    for claim in claims:
        value = json.dumps(claim["value"], ensure_ascii=False)
        wording = claim["wording"].replace("|", "\\|")
        guardrail = claim["guardrail"].replace("|", "\\|")
        lines.append(
            f"| {claim['claim_id']} | {claim['section']} | `{value}` | "
            f"`{claim['source']}::{claim['json_path']}` | "
            f"{claim['standing']} — {wording} | {guardrail} |"
        )
    lines.extend(
        [
            "",
            "## Global wording prohibitions",
            "",
            "- Do not call rank advantage an accuracy gain.",
            "- Do not call target-free scoring an online-deployable certificate.",
            "- Do not call local hashing a public preregistration.",
            "- Do not claim that the Kakeya theorem mathematically implies an EEG result.",
            "- Do not pool the 17 event participants inferentially across datasets.",
            "- Do not claim a universal AUC benefit of listener-specific weighting.",
            "- Do not conceal the KITE-ST static-profile null or the ASME eight-channel W4 failure.",
            "",
        ]
    )
    MD_OUT.write_text("\n".join(lines), encoding="utf-8")
    print(f"Saved {JSON_OUT}")
    print(f"Saved {MD_OUT}")


if __name__ == "__main__":
    main()
```
