## Supplementary material for "Falsifiable substitution tests reveal task-structured neural evidence for auditory attention": Article_Analysis_Source_Code: build_article_figures_v2.py

```
"""Build the six figures for the polished article v2 manuscript.

The script reads only final result artifacts. It does not refit a model or
recompute a scientific endpoint. Every numerical panel can therefore be
traced to a JSON/CSV artifact included in the public replication package.
"""

from __future__ import annotations

import csv
import json
import math
from pathlib import Path

import matplotlib.pyplot as plt
import numpy as np
from matplotlib.patches import FancyArrowPatch, FancyBboxPatch, Rectangle


ROOT = Path(__file__).resolve().parents[1]
OUT = ROOT / "article_outputs" / "figures"
OUT.mkdir(parents=True, exist_ok=True)

BLUE = "#176B87"
ORANGE = "#C65D36"
GREEN = "#3B8C6E"
PURPLE = "#6B3FA0"
GRAY = "#5B6470"
LIGHT = "#EEF3F6"
DATASET_COLORS = {
    "KITE_ST_confirmation_complete_case_v2": "#4C78A8",
    "KUL_64ch": "#E07B39",
    "Ear_SAAD_29ch": "#4C956C",
}
DATASET_LABELS = {
    "KITE_ST_confirmation_complete_case_v2": "KITE-ST\n253 ch",
    "KUL_64ch": "KUL\n64 ch",
    "Ear_SAAD_29ch": "Ear-SAAD\n29 ch",
}


def load_json(relative: str) -> dict:
    return json.loads((ROOT / relative).read_text(encoding="utf-8"))


def load_csv(relative: str) -> list[dict[str, str]]:
    with (ROOT / relative).open("r", encoding="utf-8-sig", newline="") as stream:
        return list(csv.DictReader(stream))


def style() -> None:
    plt.rcParams.update(
        {
            "font.family": "DejaVu Sans",
            "font.size": 10,
            "axes.spines.top": False,
            "axes.spines.right": False,
            "axes.linewidth": 0.8,
            "axes.titleweight": "bold",
            "axes.titlesize": 11,
            "axes.labelsize": 10,
            "xtick.labelsize": 9,
            "ytick.labelsize": 9,
            "legend.fontsize": 8.5,
            "figure.dpi": 160,
            "savefig.dpi": 300,
            "pdf.fonttype": 42,
            "ps.fonttype": 42,
        }
    )


def save(fig: plt.Figure, stem: str) -> None:
    validate_rounded_box_text(fig)
    fig.savefig(OUT / f"{stem}.png", bbox_inches="tight", facecolor="white")
    fig.savefig(OUT / f"{stem}.pdf", bbox_inches="tight", facecolor="white")
    fig.savefig(OUT / f"{stem}.svg", bbox_inches="tight", facecolor="white")
    plt.close(fig)


def panel_label(ax: plt.Axes, label: str) -> None:
    ax.text(
        -0.11,
        1.04,
        label,
        transform=ax.transAxes,
        fontsize=14,
        weight="bold",
        va="top",
    )


def jitter(n: int, width: float = 0.07) -> np.ndarray:
    if n <= 1:
        return np.zeros(n)
    return np.linspace(-width, width, n)


def mean_ci(ax: plt.Axes, x: float, mean: float, ci: list[float], color: str) -> None:
    ax.errorbar(
        [x],
        [mean],
        yerr=[[mean - ci[0]], [ci[1] - mean]],
        fmt="o",
        ms=7,
        color=color,
        ecolor=color,
        elinewidth=1.8,
        capsize=3,
        zorder=5,
    )


def rounded_box(
    ax: plt.Axes,
    xy: tuple[float, float],
    width: float,
    height: float,
    text: str,
    face: str,
    edge: str = "none",
    text_color: str = "#17202A",
    size: float = 9.5,
    weight: str = "normal",
) -> tuple[FancyBboxPatch, plt.Text]:
    box = FancyBboxPatch(
        xy,
        width,
        height,
        boxstyle="round,pad=0.012,rounding_size=0.018",
        facecolor=face,
        edgecolor=edge,
        linewidth=1.0,
    )
    ax.add_patch(box)
    label = ax.text(
        xy[0] + width / 2,
        xy[1] + height / 2,
        text,
        ha="center",
        va="center",
        color=text_color,
        fontsize=size,
        weight=weight,
    )
    # Keep a direct pairing so the final renderer can prove that every label
    # remains within its own background box. Matplotlib text does not wrap to
    # patch width automatically.
    label._paired_rounded_box = box
    return box, label


def validate_rounded_box_text(fig: plt.Figure) -> None:
    """Fail figure generation when a rounded-box label crosses its frame."""
    fig.canvas.draw()
    renderer = fig.canvas.get_renderer()
    failures = []
    for axis_index, ax in enumerate(fig.axes):
        for label in ax.texts:
            box = getattr(label, "_paired_rounded_box", None)
            if box is None:
                continue
            text_extent = label.get_window_extent(renderer=renderer)
            box_extent = box.get_window_extent(renderer=renderer)
            inset = 1.0
            if (
                text_extent.x0 < box_extent.x0 + inset
                or text_extent.x1 > box_extent.x1 - inset
                or text_extent.y0 < box_extent.y0 + inset
                or text_extent.y1 > box_extent.y1 - inset
            ):
                failures.append(
                    {
                        "axis": axis_index,
                        "text": label.get_text().replace("\n", " / "),
                        "text_extent": tuple(round(value, 1) for value in text_extent.extents),
                        "box_extent": tuple(round(value, 1) for value in box_extent.extents),
                    }
                )
    if failures:
        raise RuntimeError(f"Rounded-box text overflow: {failures}")


def arrow(
    ax: plt.Axes,
    start: tuple[float, float],
    end: tuple[float, float],
    color: str = GRAY,
    style: str = "-|>",
    linestyle: str = "-",
    mutation_scale: float = 12,
    linewidth: float = 1.4,
) -> None:
    ax.add_patch(
        FancyArrowPatch(
            start,
            end,
            arrowstyle=style,
            mutation_scale=mutation_scale,
            linewidth=linewidth,
            color=color,
            linestyle=linestyle,
        )
    )


def figure1() -> None:
    fig, axes = plt.subplots(2, 2, figsize=(12.5, 8.2))
    fig.suptitle(
        "From a Kakeya proof strategy to falsifiable attribution of auditory-attention evidence",
        fontsize=15,
        weight="bold",
        y=0.99,
    )

    ax = axes[0, 0]
    ax.set_xlim(0, 1)
    ax.set_ylim(0, 1)
    ax.axis("off")
    panel_label(ax, "A")
    ax.set_title(
        "Proof strategy:\nresolve a family, not a single direction",
        loc="left",
        fontsize=10.5,
        pad=4,
    )
    stage_x = (0.01, 0.28, 0.55, 0.82)
    stage_labels = ("Many\ndirections", "Overlap\nmultiplicity", "Convex-set\ncontrol", "Multiscale\nrefinement")
    stage_faces = ("#DDEAF4", "#E8E0F3", "#E0F0E8", "#F7E7D6")
    stage_sizes = (10.2, 9.2, 9.0, 9.2)
    for x, label, face, label_size in zip(stage_x, stage_labels, stage_faces, stage_sizes):
        rounded_box(ax, (x, 0.68), 0.17, 0.17, label, face, size=label_size, weight="bold")
    for x0, x1 in ((0.19, 0.27), (0.46, 0.54), (0.73, 0.81)):
        arrow(ax, (x0, 0.765), (x1, 0.765), mutation_scale=9, linewidth=1.25)
    rounded_box(
        ax,
        (0.14, 0.23),
        0.72,
        0.21,
        "Concentrated or “sticky” configurations\nrequire finer structure across scales",
        "#F4F5F7",
        edge="#A8B0B8",
        size=10.5,
    )
    arrow(ax, (0.635, 0.66), (0.635, 0.46), mutation_scale=10)
    ax.text(
        0.5,
        0.08,
        "Wang–Zahl and Guth–Wang–Zahl concern Euclidean tube families.",
        ha="center",
        fontsize=9,
        color=GRAY,
    )

    ax = axes[0, 1]
    ax.set_xlim(0, 1)
    ax.set_ylim(0, 1)
    ax.axis("off")
    panel_label(ax, "B")
    ax.set_title(
        "Translation boundary:\na testing strategy, not a neural theorem",
        loc="left",
        fontsize=10.5,
        pad=4,
    )
    rounded_box(
        ax,
        (0.02, 0.59),
        0.38,
        0.22,
        "Mathematical theorem\nEuclidean tube families",
        "#E8E0F3",
        size=10.5,
    )
    rounded_box(
        ax,
        (0.60, 0.59),
        0.38,
        0.22,
        "Auditory estimands\nheld-out EEG evidence",
        "#DDEAF4",
        size=10.5,
    )
    ax.text(
        0.50,
        0.70,
        "×",
        ha="center",
        va="center",
        color="#B3261E",
        fontsize=38,
        fontfamily="DejaVu Sans",
    )
    ax.text(0.50, 0.49, "Not a theorem-to-EEG deduction", ha="center", color="#B3261E", weight="bold")
    rounded_box(
        ax,
        (0.08, 0.10),
        0.84,
        0.30,
        "Transferred test logic:\nfamily > strongest member\nphysical mapping > arbitrary grouping\nheld-out transport + matched substitutions",
        "#F7E7D6",
        size=10.2,
    )
    arrow(ax, (0.21, 0.57), (0.34, 0.42), linestyle="--", mutation_scale=9, linewidth=1.2)
    arrow(ax, (0.79, 0.57), (0.66, 0.42), linestyle="--", mutation_scale=9, linewidth=1.2)

    ax = axes[1, 0]
    ax.set_xlim(0, 1)
    ax.set_ylim(0, 1)
    ax.axis("off")
    panel_label(ax, "C")
    ax.set_title(
        "Two auditory systems,\none falsifiable estimand",
        loc="left",
        fontsize=10.5,
        pad=4,
    )
    rounded_box(
        ax,
        (0.03, 0.59),
        0.44,
        0.25,
        "Continuous speech\ndirection × delay cells\ndisjoint source → target",
        "#DDEAF4",
        size=9.3,
        weight="bold",
    )
    rounded_box(
        ax,
        (0.53, 0.59),
        0.44,
        0.25,
        "Event-based auditory BCI\nparent stream ×\nchild command\nphase/run transport",
        "#E0F0E8",
        size=8.8,
        weight="bold",
    )
    rounded_box(
        ax,
        (0.12, 0.17),
        0.76,
        0.23,
        "Common question:\ndoes the proposed family retain\nfunction in held-out data?",
        "#F4F5F7",
        edge="#A8B0B8",
        size=10.5,
    )
    arrow(ax, (0.25, 0.57), (0.40, 0.42), mutation_scale=10)
    arrow(ax, (0.75, 0.57), (0.60, 0.42), mutation_scale=10)

    ax = axes[1, 1]
    ax.set_xlim(0, 1)
    ax.set_ylim(0, 1)
    ax.axis("off")
    panel_label(ax, "D")
    ax.set_title(
        "Matched substitutions make\nthe interpretation falsifiable",
        loc="left",
        fontsize=10.5,
        pad=4,
    )
    labels = [
        ("Leading direction", "Direction family", "#DDEAF4"),
        ("Remapped coordinates", "Physical mapping", "#E8E0F3"),
        ("Other / pooled listener", "Matched listener", "#E0F0E8"),
        ("Target / command /\nEOG-only substitutions", "EEG evidence exceeding\nan EOG-only comparator", "#F7E7D6"),
    ]
    y = 0.78
    for row_index, (left, right, face) in enumerate(labels):
        label_size = 8.0 if row_index == 3 else 8.8
        rounded_box(ax, (0.015, y - 0.07), 0.41, 0.14, left, "#F4F5F7", edge="#B8BEC5", size=label_size)
        ax.text(0.50, y, "vs", ha="center", va="center", weight="bold", color=GRAY)
        rounded_box(ax, (0.575, y - 0.07), 0.41, 0.14, right, face, size=label_size, weight="bold")
        y -= 0.20
    ax.text(
        0.5,
        0.04,
        "The proposed structure must transport; matched alternatives must fail.",
        ha="center",
        fontsize=8.8,
        color=GRAY,
    )

    fig.subplots_adjust(hspace=0.28, wspace=0.16, top=0.91)
    save(fig, "figure1_framework")


def figure2() -> None:
    projection_files = [
        (
            "KUL even",
            "results/kul_aad/locked_confirmation/participants.csv",
            "equal_k4_gain_vs_dominant_k1",
            "#4C78A8",
        ),
        (
            "DTU",
            "results/dtu_aad/locked_external/participants.csv",
            "equal_k4_gain_vs_dominant_k1",
            "#E07B39",
        ),
        (
            "Ear-SAAD",
            "results/ear_saad/locked_multimodal_external/primary_participants.csv",
            "gain_equal_minus_k1",
            "#4C956C",
        ),
    ]
    confirm = load_json(
        "results/kite_sat/akb_confirmation_complete_case_v2/confirmation_decision_v1.json"
    )
    mechanism = load_json(
        "results/kite_sat/akb_confirmation_complete_case_direction_delay_v2/mechanism_decision_v1.json"
    )
    synthesis = load_json("results/kite_sat/article_evidence_synthesis_v1/evidence_synthesis_v1.json")

    fig, axes = plt.subplots(2, 2, figsize=(12.2, 8.2))
    fig.suptitle(
        "Continuous-speech information extends beyond the leading projection and depends on direction–delay structure",
        fontsize=15,
        weight="bold",
    )
    ax = axes[0, 0]
    panel_label(ax, "A")
    ax.set_title("Four CCA margins versus the leading margin")
    for i, (label, path, column, color) in enumerate(projection_files):
        values = np.array([float(row[column]) for row in load_csv(path)])
        ax.scatter(i + jitter(len(values)), values, s=28, color=color, alpha=0.65)
        ax.plot([i - 0.16, i + 0.16], [values.mean(), values.mean()], color="black", lw=2.5)
    ax.axhline(0, color="#777", lw=1, ls="--")
    ax.set_xticks(range(3), [x[0] for x in projection_files])
    ax.set_ylabel("Accuracy gain")
    ax.text(
        0.02,
        0.95,
        "32/41 positive\nstudy-equal +0.0201\n95% CI 0.0125–0.0279",
        transform=ax.transAxes,
        va="top",
        fontsize=9,
    )

    ax = axes[0, 1]
    panel_label(ax, "B")
    ax.set_title("Participant-disjoint KITE-ST confirmation")
    rows = confirm["participant_results"]
    baseline = np.array([row["intact_global_auc"] for row in rows])
    bank = np.array([row["intact_AKB_auc"] for row in rows])
    for i in range(len(rows)):
        ax.plot([0, 1], [baseline[i], bank[i]], color="#BEC4C9", lw=1)
    ax.scatter(np.zeros(len(rows)), baseline, color=GRAY, s=28, zorder=3)
    ax.scatter(np.ones(len(rows)), bank, color=BLUE, s=28, zorder=3)
    ax.set_xticks([0, 1], ["Global model", "8 × 2 bank"])
    ax.set_ylabel("Held-story AUC")
    ax.text(
        0.02,
        0.95,
        "Gain +0.0544\n95% CI 0.0188–0.0987\n10/12 positive; P=0.00391",
        transform=ax.transAxes,
        va="top",
        fontsize=9,
    )

    ax = axes[1, 0]
    panel_label(ax, "C")
    ax.set_title("Physical topology versus 15 capacity-matched mappings")
    values = np.array(
        [row["intact_identity_advantage_auc"] for row in mechanism["participant_results"]]
    )
    ax.scatter(jitter(len(values), 0.05), values, color=PURPLE, s=34, alpha=0.75)
    ax.plot([-0.12, 0.12], [values.mean(), values.mean()], color="black", lw=2.8)
    ax.axhline(0, color="#777", lw=1, ls="--")
    ax.set_xlim(-0.35, 0.35)
    ax.set_xticks([0], ["Physical − mean wrong"])
    ax.set_ylabel("AUC advantage")
    ax.text(
        0.04,
        0.94,
        "Mean +0.0778\n12/12 positive; P=0.000244\nsame cells, parameters, and weight histogram",
        transform=ax.transAxes,
        va="top",
        fontsize=9,
    )

    ax = axes[1, 1]
    panel_label(ax, "D")
    ax.set_title("Independent retraining across sensor density")
    budgets = synthesis["sensor_density_boundary"]["budgets"]
    x = [row["budget"] for row in budgets]
    gain = [row["mean_AKB_minus_global_auc"] for row in budgets]
    topo = [row["mean_identity_minus_controls_auc"] for row in budgets]
    ax.plot(x, gain, "-o", color=BLUE, lw=2, label="Bank − global")
    ax.plot(x, topo, "-s", color=ORANGE, lw=2, label="Physical − remapped")
    ax.axhline(0, color="#777", lw=1, ls="--")
    ax.set_xscale("log", base=2)
    ax.set_xticks(x, [str(v) for v in x])
    ax.set_xlabel("EEG sensors")
    ax.set_ylabel("AUC advantage")
    ax.legend(
        frameon=True,
        facecolor="white",
        edgecolor="none",
        framealpha=1.0,
        loc="lower right",
        bbox_to_anchor=(0.98, 0.04),
        fontsize=7.5,
    )
    ax.text(
        0.03,
        0.95,
        "All four densities positive;\nno significant gain-by-density slope",
        transform=ax.transAxes,
        va="top",
        fontsize=9,
        bbox={"facecolor": "white", "edgecolor": "none", "alpha": 1.0, "pad": 1.8},
        zorder=10,
    )

    fig.tight_layout(rect=(0, 0, 1, 0.94))
    save(fig, "figure2_continuous_structure")


def endpoint_panel(
    ax: plt.Axes,
    rows: list[dict[str, str]],
    summary: dict,
    field: str,
    title: str,
    letter: str,
) -> None:
    panel_label(ax, letter)
    ax.set_title(title)
    order = list(DATASET_COLORS)
    for i, dataset in enumerate(order):
        values = np.array(
            [float(row[field]) for row in rows if row["dataset"] == dataset]
        )
        color = DATASET_COLORS[dataset]
        ax.scatter(i + jitter(len(values)), values, color=color, alpha=0.58, s=24)
        stats = summary[field][dataset]
        mean_ci(ax, i, stats["mean"], stats["bootstrap_95"], color)
    cross = summary["_cross"][field]
    mean_ci(
        ax,
        3.55,
        cross["equal_dataset_mean"],
        cross["stratified_bootstrap_95"],
        PURPLE,
    )
    ax.axhline(0, color="#777", lw=1, ls="--")
    ax.set_xticks(
        [0, 1, 2, 3.55],
        [DATASET_LABELS[x] for x in order] + ["Study-\nequal"],
    )
    p_value = cross["participant_signflip_randomization_p"]
    p_label = "P<10⁻⁶" if p_value < 1e-6 else (f"P={p_value:.6f}" if p_value < 0.001 else f"P={p_value:.3g}")
    ax.text(
        0.98,
        0.05,
        p_label,
        transform=ax.transAxes,
        ha="right",
        color=GRAY,
        fontsize=8.5,
    )


def figure3() -> None:
    transport = load_json(
        "results/wz_hic_cross_dataset/continuous_listener_transport_v1/"
        "continuous_listener_transport_decision_v1.json"
    )
    templates = load_json(
        "results/wz_hic_cross_dataset/continuous_shared_template_v1/"
        "continuous_shared_template_decision_v1.json"
    )
    rows = load_csv(
        "results/wz_hic_cross_dataset/continuous_listener_transport_v1/"
        "participant_endpoints_v1.csv"
    )
    template_rows = load_csv(
        "results/wz_hic_cross_dataset/continuous_shared_template_v1/"
        "participant_endpoints_v1.csv"
    )
    transport_summary = dict(transport["dataset_summaries"])
    transport_summary["_cross"] = transport["cross_dataset_summaries"]
    template_summary = dict(templates["dataset_summaries"])
    template_summary["_cross"] = templates["cross_dataset_summaries"]

    panels = [
        (
            rows,
            transport_summary,
            "F1_fingerprint_listener_rank_advantage",
            "Profile: own listener vs donors",
            "A",
        ),
        (
            rows,
            transport_summary,
            "F2_fingerprint_physical_mapping_rank_advantage",
            "Profile: physical vs wrong mappings",
            "B",
        ),
        (
            rows,
            transport_summary,
            "P1_predictive_listener_rank_advantage",
            "Held speech: own listener vs donors",
            "C",
        ),
        (
            rows,
            transport_summary,
            "P2_predictive_physical_mapping_rank_advantage",
            "Held speech: physical vs wrong mappings",
            "D",
        ),
        (
            template_rows,
            template_summary,
            "G1_own_vs_mean_profile_rank_advantage",
            "Held speech: own vs mean-profile template",
            "E",
        ),
        (
            template_rows,
            template_summary,
            "G2_own_vs_mean_weight_rank_advantage",
            "Held speech: own vs mean-weight template",
            "F",
        ),
    ]

    fig, axes = plt.subplots(3, 2, figsize=(12.3, 11.1), sharey=True)
    fig.suptitle(
        "Structure-dependent held-speech evidence across nonoverlapping continuous speech",
        fontsize=15,
        weight="bold",
    )
    for ax, args in zip(axes.flat, panels):
        endpoint_panel(ax, *args)
    for ax in axes[:, 0]:
        ax.set_ylabel("Rank advantage over chance")
    axes[0, 0].set_ylim(-0.54, 0.54)
    axes[0, 0].text(
        0.02,
        0.05,
        "KITE-ST static profile recurrence is null",
        transform=axes[0, 0].transAxes,
        fontsize=8.5,
        color=GRAY,
    )
    axes[2, 0].text(
        0.02,
        0.05,
        "Receiver excluded from pooled template",
        transform=axes[2, 0].transAxes,
        fontsize=8.5,
        color=GRAY,
    )
    fig.text(
        0.5,
        0.012,
        "Rank advantage is the probability of winning a matched comparison minus 0.5; it is not an accuracy gain.",
        ha="center",
        fontsize=9,
        color=GRAY,
    )
    fig.tight_layout(rect=(0, 0.035, 1, 0.955))
    save(fig, "figure3_continuous_transport")


def figure4() -> None:
    decision = load_json(
        "results/wz_hic_cross_dataset/target_free_mechanism_v1/target_free_decision_v1.json"
    )
    rows = []
    for dataset, participants in decision["evaluable_participants"].items():
        for participant in participants:
            item = decision["participants"][dataset][participant]
            rows.append(
                {
                    "dataset": dataset,
                    "participant": participant,
                    "target_free_eeg_auc": item["target_free_EEG"]["certificate_AUC"],
                    "target_free_minus_best_wrong_auc": item["contrasts"]["EEG_minus_best_wrong"],
                }
            )
    fig, axes = plt.subplots(2, 2, figsize=(12.2, 8.5))
    fig.suptitle(
        "Non-target sibling events identify parent-stream errors hidden by a flat auditory BCI",
        fontsize=15,
        weight="bold",
    )

    ax = axes[0, 0]
    panel_label(ax, "A")
    ax.set_title("Flat child decision and separate parent score")
    ax.set_xlim(0, 1)
    ax.set_ylim(0, 1)
    ax.axis("off")
    rounded_box(ax, (0.02, 0.68), 0.40, 0.18, "Flat decoder\nselects child command ĉ", "#DDEAF4", size=11, weight="bold")
    rounded_box(ax, (0.59, 0.68), 0.36, 0.18, "Parent score\nevaluates stream of ĉ", "#E0F0E8", size=11, weight="bold")
    arrow(ax, (0.42, 0.77), (0.58, 0.77))
    rounded_box(ax, (0.05, 0.24), 0.36, 0.20, "Error type 1\nwrong parent stream", "#F7DAD4", size=11)
    rounded_box(ax, (0.59, 0.24), 0.36, 0.20, "Error type 2\nright parent, wrong child", "#F7E7D6", size=11)
    arrow(ax, (0.23, 0.66), (0.23, 0.46))
    arrow(ax, (0.77, 0.66), (0.77, 0.46))
    ax.text(
        0.5,
        0.07,
        "Target-event-excluded check removes every known-target event using oracle labels;\nit is mechanism-oriented, not causal identification or an online score.",
        ha="center",
        fontsize=9,
        color=GRAY,
    )

    datasets = ["Kojima2024B", "ASME30"]
    colors = [BLUE, ORANGE]
    labels = ["Kojima\n(n=7)", "ASME30\n(n=9)"]

    ax = axes[0, 1]
    panel_label(ax, "B")
    ax.set_title("Target-event-excluded parent-error AUC")
    for i, (dataset, color) in enumerate(zip(datasets, colors)):
        values = np.array(
            [float(row["target_free_eeg_auc"]) for row in rows if row["dataset"] == dataset]
        )
        ax.scatter(i + jitter(len(values)), values, color=color, alpha=0.65, s=28)
        stats = decision["summaries"][dataset]["target_free_absolute"]
        mean_ci(ax, i, stats["mean"], stats["bootstrap_95_CI_mean"], color)
    ax.axhline(0.5, color="#777", ls="--", lw=1)
    ax.set_xticks([0, 1], labels)
    ax.set_ylim(0.45, 1.04)
    ax.set_ylabel("AUC")

    ax = axes[1, 0]
    panel_label(ax, "C")
    ax.set_title("Physical parent binding versus matched wrong bindings")
    for i, (dataset, color) in enumerate(zip(datasets, colors)):
        values = np.array(
            [
                float(row["target_free_minus_best_wrong_auc"])
                for row in rows
                if row["dataset"] == dataset
            ]
        )
        ax.scatter(i + jitter(len(values)), values, color=color, alpha=0.65, s=28)
        stats = decision["summaries"][dataset]["mapping_specificity"]
        mean_ci(ax, i, stats["mean"], stats["bootstrap_95_CI_mean"], color)
    ax.axhline(0, color="#777", ls="--", lw=1)
    ax.set_xticks([0, 1], labels)
    ax.set_ylabel("AUC difference")

    ax = axes[1, 1]
    panel_label(ax, "D")
    ax.set_title("EEG–EOG contrast; target exclusion changes little")
    width = 0.28
    for i, (dataset, color) in enumerate(zip(datasets, colors)):
        eeg_eog = decision["summaries"][dataset]["nonocular"]
        removal = decision["summaries"][dataset]["target_free_minus_inclusive"]
        for j, (stats, hatch) in enumerate(((eeg_eog, ""), (removal, "//"))):
            x = i + (j - 0.5) * width
            ax.bar(
                x,
                stats["mean"],
                width=width * 0.85,
                color=color if j == 0 else "white",
                edgecolor=color,
                hatch=hatch,
                linewidth=1.4,
            )
            ax.errorbar(
                x,
                stats["mean"],
                yerr=[
                    [stats["mean"] - stats["bootstrap_95_CI_mean"][0]],
                    [stats["bootstrap_95_CI_mean"][1] - stats["mean"]],
                ],
                color="black",
                capsize=3,
                lw=1,
            )
    ax.axhline(0, color="#777", ls="--", lw=1)
    ax.set_xticks([0, 1], labels)
    ax.set_ylabel("AUC difference")
    ax.legend(
        [
            Rectangle((0, 0), 1, 1, facecolor=GRAY),
            Rectangle((0, 0), 1, 1, facecolor="white", edgecolor=GRAY, hatch="//"),
        ],
        ["EEG − EOG", "Excluded − inclusive"],
        frameon=False,
        loc="upper right",
    )

    fig.tight_layout(rect=(0, 0, 1, 0.94))
    save(fig, "figure4_target_free_parent")


def figure5() -> None:
    data = load_json(
        "results/wz_hic_cross_dataset/individual_geometry_synthesis_v1/synthesis_v1.json"
    )
    rows = data["participant_rows"]
    colors = {
        "ASME30": BLUE,
        "Kojima2024B": ORANGE,
        "wrong": "#D98C35",
        "eog": "#8A8A8A",
    }
    display_name = {"ASME30": "ASME30", "Kojima2024B": "Kojima"}
    datasets = ("ASME30", "Kojima2024B")

    def bootstrap_ci(values: np.ndarray) -> tuple[np.ndarray, np.ndarray]:
        rng = np.random.default_rng(20260730)
        ids = rng.integers(0, values.shape[0], size=(20_000, values.shape[0]))
        draws = values[ids].mean(axis=1)
        return np.quantile(draws, 0.025, axis=0), np.quantile(draws, 0.975, axis=0)

    with plt.rc_context({"font.size": 8.5, "axes.titlesize": 10.2, "axes.labelsize": 8.8}):
        fig, axes = plt.subplots(2, 2, figsize=(7.25, 6.3))
        ax_a, ax_b, ax_c, ax_d = axes.flat

        # A: held-out effect and capacity-matched controls by dataset.
        metric_specs = [
            ("physical_effect", "Physical EEG", None),
            ("best_wrong_effect", "Best wrong", colors["wrong"]),
            ("eog_effect", "EOG", colors["eog"]),
        ]
        base_positions = {"ASME30": 0.0, "Kojima2024B": 4.2}
        for dataset in datasets:
            dataset_rows = [row for row in rows if row["dataset"] == dataset]
            positions = base_positions[dataset] + np.arange(3)
            for row in dataset_rows:
                ax_a.plot(
                    positions,
                    [float(row[key]) for key, _, _ in metric_specs],
                    color="#D0D0D0",
                    linewidth=0.65,
                    zorder=1,
                )
            for position, (key, _, fixed_color) in zip(positions, metric_specs):
                values = np.asarray([float(row[key]) for row in dataset_rows])
                color = fixed_color or colors[dataset]
                ax_a.scatter(
                    np.full(len(values), position), values, s=23, color=color,
                    edgecolor="white", linewidth=0.5, zorder=3,
                )
                ax_a.plot(
                    [position - 0.16, position + 0.16],
                    [values.mean(), values.mean()], color="black", linewidth=2, zorder=4,
                )
            ax_a.text(
                positions.mean(), -0.21, display_name[dataset], ha="center", va="center",
                color=colors[dataset], fontweight="bold", transform=ax_a.transData,
            )
        ax_a.axhline(0, color="#555555", linestyle="--", linewidth=0.7)
        ax_a.set_xticks(
            [0, 1, 2, 4.2, 5.2, 6.2],
            [label for _, label, _ in metric_specs] * 2,
            rotation=22,
        )
        ax_a.set_ylabel("Held-out S−U effect (training SD)")
        ax_a.set_ylim(-0.28, 1.55)
        ax_a.set_title("Held-out physical parent binding", loc="left")
        ax_a.text(
            0.02, 0.97, "Physical and binding effects: 17/17 positive",
            transform=ax_a.transAxes, ha="left", va="top", fontsize=7.5,
        )

        # B: listener identity and physical mapping specificity.
        jitter_offsets = {"ASME30": -0.09, "Kojima2024B": 0.09}
        for dataset in datasets:
            dataset_rows = [row for row in rows if row["dataset"] == dataset]
            for x_value, metric in ((0, "identity_advantage"), (1, "mapping_alignment_specificity")):
                values = np.asarray([float(row[metric]) for row in dataset_rows])
                xpos = x_value + jitter_offsets[dataset]
                ax_b.scatter(
                    np.full(len(values), xpos), values, s=28, color=colors[dataset],
                    edgecolor="white", linewidth=0.5,
                    label=display_name[dataset] if x_value == 0 else None,
                )
                ax_b.plot(
                    [xpos - 0.08, xpos + 0.08], [values.mean(), values.mean()],
                    color="black", linewidth=1.7,
                )
        ax_b.axhline(0, color="#555555", linestyle="--", linewidth=0.7)
        ax_b.set_xticks(
            [0, 1], ["Same-person −\nother-person", "Physical −\nbest-wrong alignment"]
        )
        ax_b.set_ylabel("Direction-cosine contrast")
        ax_b.set_title("Participant and binding identity", loc="left")
        ax_b.legend(frameon=False, fontsize=7.5, loc="upper right")
        ax_b.text(
            0.02, 0.04, "Identity: 16/17 positive\nMapping alignment: 14/17 positive",
            transform=ax_b.transAxes, ha="left", va="bottom", fontsize=7.4,
        )

        # C: command residualization leaves participant-level effects intact.
        markers = {"ASME30": "o", "Kojima2024B": "s"}
        for dataset in datasets:
            dataset_rows = [row for row in rows if row["dataset"] == dataset]
            raw_physical = np.asarray([float(row["physical_effect"]) for row in dataset_rows])
            residual_physical = np.asarray(
                [float(row["residual_physical_effect"]) for row in dataset_rows]
            )
            raw_binding = np.asarray([float(row["binding_specificity"]) for row in dataset_rows])
            residual_binding = np.asarray(
                [float(row["residual_binding_specificity"]) for row in dataset_rows]
            )
            ax_c.scatter(
                raw_physical, residual_physical, s=31, marker=markers[dataset],
                color=colors[dataset], edgecolor="white", linewidth=0.5,
                label=f"{display_name[dataset]}: physical",
            )
            ax_c.scatter(
                raw_binding, residual_binding, s=31, marker=markers[dataset],
                facecolor="white", edgecolor=colors[dataset], linewidth=1.1,
                label=f"{display_name[dataset]}: binding",
            )
        all_values = [
            float(row[key])
            for row in rows
            for key in (
                "physical_effect", "residual_physical_effect",
                "binding_specificity", "residual_binding_specificity",
            )
        ]
        lower, upper = min(all_values) - 0.05, max(all_values) + 0.05
        ax_c.plot([lower, upper], [lower, upper], color="#555555", linestyle="--", linewidth=0.8)
        ax_c.set_xlim(lower, upper)
        ax_c.set_ylim(lower, upper)
        ax_c.set_aspect("equal", adjustable="box", anchor="W")
        ax_c.set_xlabel("Original target-event-excluded effect")
        ax_c.set_ylabel("After training-only command residualization", labelpad=6)
        ax_c.set_title("Command-identity residual control", loc="left")
        ax_c.legend(frameon=False, fontsize=6.8, loc="lower right")

        # D: fixed-bin time profiles and within-dataset maximum-t decisions.
        centers = np.arange(50, 1000, 100)
        for dataset in datasets:
            time_rows = data["dataset_summaries"][dataset]["time_resolved"]
            values = np.asarray([row["participant_values"] for row in time_rows], dtype=float).T
            mean = values.mean(axis=0)
            low, high = bootstrap_ci(values)
            ax_d.fill_between(centers, low, high, color=colors[dataset], alpha=0.14, linewidth=0)
            ax_d.plot(
                centers, mean, color=colors[dataset], marker="o", markersize=3.3,
                linewidth=1.8, label=display_name[dataset],
            )
            for center, row, y_value in zip(centers, time_rows, mean):
                if float(row["max_t_fwer_p"]) <= 0.05:
                    ax_d.text(
                        center, y_value + (0.055 if dataset == "ASME30" else -0.055),
                        "●", color=colors[dataset], ha="center", va="center", fontsize=5.7,
                    )
        ax_d.axhline(0, color="#555555", linestyle="--", linewidth=0.7)
        ax_d.set_xlim(0, 1000)
        ax_d.set_xticks(np.arange(0, 1001, 200))
        ax_d.set_xlabel("Time after auditory event (ms)")
        ax_d.set_ylabel("Physical − best-wrong held-out effect")
        ax_d.set_title("Fixed-bin binding specificity", loc="left")
        ax_d.legend(frameon=False, fontsize=7.5, loc="upper right")
        ax_d.text(
            0.02, 0.04, "Dots: within-dataset exact max-T FWER p ≤ 0.05",
            transform=ax_d.transAxes, ha="left", va="bottom", fontsize=7.1,
        )

        fig.suptitle(
            "Task-defined event directions transport within listeners across phases or runs",
            fontsize=11, fontweight="bold", y=0.995,
        )
        fig.text(
            0.01, 0.006,
            "Known target-command events excluded with oracle labels; models evaluated on held-out phases or runs. "
            "No cross-dataset model training; inference remains dataset specific.",
            fontsize=6.7,
        )
        fig.tight_layout(rect=(0.025, 0.04, 1, 0.965), h_pad=2.25, w_pad=1.7)

        # Equal aspect previously centered panel C within its grid cell. Anchor it
        # to the west and place panel letters in figure coordinates so A/C share
        # the same left edge and C cannot overlap its vertical-axis title.
        fig.canvas.draw()
        ax_c.set_anchor("W")
        fig.canvas.draw()
        if abs(ax_a.get_position().x0 - ax_c.get_position().x0) > 0.002:
            raise RuntimeError("Figure 5 panels A and C are not left-aligned")
        for label, axis in zip(("A", "B", "C", "D"), axes.flat):
            box = axis.get_position()
            x_offset = 0.095 if label == "C" else 0.055
            y_offset = 0.032 if label == "D" else 0.012
            fig.text(
                box.x0 - x_offset, box.y1 + y_offset, label,
                fontsize=12, fontweight="bold", va="bottom", ha="left",
            )
    save(fig, "figure5_event_transport")


def figure6() -> None:
    wearable = load_json(
        "results/wz_hic_cross_dataset/wearable_calibration_v1/"
        "wearable_calibration_decision_v1.json"
    )
    fig, axes = plt.subplots(2, 2, figsize=(12.2, 8.6))
    fig.suptitle(
        "Physical binding survives wearable reduction, but listener specificity is less robust",
        fontsize=14.5,
        weight="bold",
    )
    dataset_names = ["ASME30", "Kojima2024B"]
    colors = [BLUE, ORANGE]

    ax = axes[0, 0]
    panel_label(ax, "A")
    ax.set_title("Physical binding across electrode budgets")
    for dataset, color in zip(dataset_names, colors):
        summaries = wearable["montage_summaries"][dataset]
        x = sorted(int(key) for key in summaries)
        y = [summaries[str(key)]["binding_specificity"]["mean"] for key in x]
        lo = [summaries[str(key)]["binding_specificity"]["bootstrap_95_CI_mean"][0] for key in x]
        hi = [summaries[str(key)]["binding_specificity"]["bootstrap_95_CI_mean"][1] for key in x]
        ax.errorbar(x, y, yerr=[np.array(y) - lo, np.array(hi) - y], marker="o", color=color, capsize=3, label=dataset.replace("Kojima2024B", "Kojima"))
    ax.axhline(0, color="#777", ls="--", lw=1)
    ax.set_xscale("log", base=2)
    ax.set_xticks([8, 16, 32, 64], ["8", "16", "32", "64"])
    ax.set_xlabel("EEG channels")
    ax.set_ylabel("Physical − best wrong (z)")
    ax.legend(frameon=False)

    ax = axes[0, 1]
    panel_label(ax, "B")
    ax.set_title("Listener-specific transport across electrode budgets")
    for dataset, color in zip(dataset_names, colors):
        summaries = wearable["montage_summaries"][dataset]
        x = sorted(int(key) for key in summaries)
        y = [summaries[str(key)]["identity_advantage"]["mean"] for key in x]
        lo = [summaries[str(key)]["identity_advantage"]["bootstrap_95_CI_mean"][0] for key in x]
        hi = [summaries[str(key)]["identity_advantage"]["bootstrap_95_CI_mean"][1] for key in x]
        ax.errorbar(x, y, yerr=[np.array(y) - lo, np.array(hi) - y], marker="o", color=color, capsize=3, label=dataset.replace("Kojima2024B", "Kojima"))
    ax.axhline(0, color="#777", ls="--", lw=1)
    ax.set_xscale("log", base=2)
    ax.set_xticks([8, 16, 32, 64], ["8", "16", "32", "64"])
    ax.set_xlabel("EEG channels")
    ax.set_ylabel("Own − other listener cosine")
    ax.text(
        0.04,
        0.12,
        "ASME30, 8 ch: mean CI > 0,\nbut 7/10 positive < specified 8/10 rule",
        transform=ax.transAxes,
        fontsize=8.5,
        color=GRAY,
    )

    ax = axes[1, 0]
    panel_label(ax, "C")
    ax.set_title("Eight-channel binding retention")
    retention = [
        wearable["eight_channel_binding_retention"][dataset] for dataset in dataset_names
    ]
    x = np.arange(2)
    ax.bar(x, retention, 0.52, color=colors, alpha=0.9)
    ax.axhline(0.50, color="#777", ls=":", lw=1.2)
    ax.axhline(0, color="#777", ls="--", lw=1)
    ax.set_xticks(x, [name.replace("Kojima2024B", "Kojima") for name in dataset_names])
    ax.set_ylabel("8/64-channel retention ratio")
    ax.text(0.03, 0.52, "specified minimum", transform=ax.get_yaxis_transform(), fontsize=8, color=GRAY, va="bottom")

    ax = axes[1, 1]
    panel_label(ax, "D")
    ax.set_title("Three-run binding specificity")
    three_run = [
        wearable["calibration_summaries"][dataset]["3"]["binding_specificity"]
        for dataset in dataset_names
    ]
    for i, (stats, color) in enumerate(zip(three_run, colors)):
        value = stats["mean"]
        ci = stats["bootstrap_95_CI_mean"]
        ax.bar(i, value, width=0.52, color=color, alpha=0.85)
        ax.errorbar(i, value, yerr=[[value - ci[0]], [ci[1] - value]], color="black", capsize=3)
    ax.axhline(0, color="#777", ls="--", lw=1)
    ax.set_xticks(range(2), [name.replace("Kojima2024B", "Kojima") for name in dataset_names])
    ax.set_ylabel("Physical − best wrong (z)")

    fig.tight_layout(rect=(0, 0, 1, 0.94))
    save(fig, "figure6_boundaries")


def main() -> None:
    style()
    figure1()
    figure2()
    figure3()
    figure4()
    figure5()
    figure6()
    print(f"Saved six figures to {OUT}")


if __name__ == "__main__":
    main()
```
