## Supplementary material for "Falsifiable substitution tests reveal task-structured neural evidence for auditory attention": Article_Analysis_Source_Code: build_article_kite_summary_v1.py

```
#!/usr/bin/env python3
"""Build the article-only KITE summary from audited reported-result inputs.

This deliberately excludes AASD, ESAA, and other exploratory branches that are
outside the six-dataset article scope.  It is a deterministic reporting step;
it does not refit an EEG model.
"""

from __future__ import annotations

import hashlib
import json
from pathlib import Path


ROOT = Path(__file__).resolve().parents[1]
CONFIRM = (
    ROOT / "results" / "kite_sat" / "akb_confirmation_complete_case_v2"
    / "confirmation_decision_v1.json"
)
CONFIRM_AUDIT = CONFIRM.with_name("confirmation_outcome_audit_v1.json")
MECHANISM = (
    ROOT / "results" / "kite_sat"
    / "akb_confirmation_complete_case_direction_delay_v2"
    / "mechanism_decision_v1.json"
)
MECHANISM_AUDIT = MECHANISM.with_name("mechanism_outcome_audit_precision_v2.json")
DENSITY = (
    ROOT / "results" / "kite_sat" / "sensor_density_mechanism_v1"
    / "sensor_density_decision_v1.json"
)
DENSITY_AUDIT = DENSITY.with_name("sensor_density_outcome_audit_v1.json")
OUTPUT = (
    ROOT / "results" / "kite_sat" / "article_evidence_synthesis_v1"
    / "evidence_synthesis_v1.json"
)


def sha256(path: Path) -> str:
    digest = hashlib.sha256()
    with path.open("rb") as stream:
        for chunk in iter(lambda: stream.read(1024 * 1024), b""):
            digest.update(chunk)
    return digest.hexdigest()


def load(path: Path) -> dict:
    return json.loads(path.read_text(encoding="utf-8"))


def source(path: Path) -> dict[str, str]:
    return {
        "path": path.relative_to(ROOT).as_posix(),
        "sha256": sha256(path),
    }


def main() -> None:
    if OUTPUT.exists():
        raise RuntimeError("Refusing to overwrite the article-only KITE summary.")
    confirm, mechanism, density = load(CONFIRM), load(MECHANISM), load(DENSITY)
    confirm_audit = load(CONFIRM_AUDIT)
    mechanism_audit = load(MECHANISM_AUDIT)
    density_audit = load(DENSITY_AUDIT)
    if (
        confirm_audit.get("status") != "pass"
        or mechanism_audit.get("status") != "pass"
        or density_audit.get("status") != "pass"
    ):
        raise RuntimeError("A required KITE result audit did not pass.")
    cm = confirm["metrics"]
    mm = mechanism["metrics"]
    budgets = density["budget_metrics"]
    payload = {
        "schema": "article-only-KITE-evidence-synthesis-v1",
        "status": "pass",
        "scope": (
            "Reported KITE-ST confirmation, capacity-matched topology, and "
            "sensor-density results only; exploratory external branches excluded."
        ),
        "cohort_endpoints": [
            {"cohort": "reserved-development-baseline", "reported": False},
            {"cohort": "reserved-development-AKB", "reported": False},
            {
                "cohort": "KITE-ST high-density participant-disjoint complete cases",
                "standing": "independent-participant confirmation within one public acquisition site",
                "endpoint": "intact held-story AUC",
                "n": cm["n"],
                "comparator_performance": cm["mean_intact_global_auc"],
                "AKB_performance": cm["mean_intact_AKB_auc"],
                "AKB_minus_comparator": cm["mean_intact_gain_auc"],
                "bootstrap_95": cm["intact_gain_bootstrap_95_percent_ci"],
                "exact_one_sided_p": confirm["fixed_sequence_decision"]["test_1_intact_gain"]["exact_one_sided_p"],
                "participants_positive_gain": cm["participants_positive_intact_gain"],
                "status": confirm["status"],
            },
        ],
        "capacity_matched_topology_mechanism": {
            "standing": "post-primary test against the mean of 15 prespecified remappings",
            "mean_identity_minus_15_controls_auc": mm["mean_intact_identity_advantage_auc"],
            "median_advantage_auc": mm["median_intact_identity_advantage_auc"],
            "participants_positive": mm["participants_positive_intact_identity_advantage"],
            "exact_one_sided_p": mechanism["fixed_sequence"]["decoding_identity_advantage"]["exact_one_sided_p"],
            "angular_only_advantage_auc": mm["mean_angular_only_advantage_auc"],
            "delay_swapped_advantage_auc": mm["mean_delay_swapped_advantage_auc"],
            "patch_identity_advantage_auc": mm["mean_patch_identity_advantage_auc"],
            "reliability_stability_subclaim": {
                "mean_advantage": mm["mean_reliability_identity_advantage"],
                "participants_positive": mm["participants_positive_reliability_identity_advantage"],
                "exact_one_sided_p": mechanism["fixed_sequence"]["reliability_identity_advantage"]["raw_exact_one_sided_p"],
                "supported": mechanism["fixed_sequence"]["reliability_identity_advantage"]["rejected_with_mechanistic_standing"],
            },
            "broadness_descriptive": {
                "effective_cells_of_16": mm["mean_effective_cell_number"],
                "normalized_entropy": mm["mean_normalized_cell_entropy"],
                "maximum_cell_mass": mm["mean_maximum_cell_mass"],
                "effective_angular_sectors_of_8": mm["mean_effective_angular_sector_number"],
            },
        },
        "sensor_density_boundary": {
            "standing": "post-confirmation mechanism-oriented analysis",
            "budgets": budgets,
            "claim_boundary": density["claim_boundary"],
        },
        "sources": {
            "confirmation": source(CONFIRM),
            "confirmation_audit": source(CONFIRM_AUDIT),
            "mechanism": source(MECHANISM),
            "mechanism_audit": source(MECHANISM_AUDIT),
            "sensor_density": source(DENSITY),
            "sensor_density_audit": source(DENSITY_AUDIT),
        },
    }
    OUTPUT.parent.mkdir(parents=True, exist_ok=True)
    OUTPUT.write_text(
        json.dumps(payload, indent=2, allow_nan=False) + "\n", encoding="utf-8"
    )
    observed = sha256(OUTPUT)
    OUTPUT.with_name(OUTPUT.name + ".sha256").write_text(
        f"{observed}  {OUTPUT.name}\n", encoding="ascii"
    )
    print(json.dumps({"status": "pass", "sha256": observed}))


if __name__ == "__main__":
    main()
```
