## Supplementary material for "Falsifiable substitution tests reveal task-structured neural evidence for auditory attention": Article_Analysis_Source_Code: build_asme30_wz_hic_dataset_native_features_v2.py

```
#!/usr/bin/env python3
"""Build ASME30 features using the source article's online preprocessing."""

from __future__ import annotations

import argparse
import hashlib
import json
import platform
import sys
from datetime import datetime, timezone
from pathlib import Path

import mne
import numpy as np
import scipy
from scipy.signal import butter, sosfilt

from build_asme30_wz_hic_features_v1 import read_channels, read_unique_events
from wz_hic_asme30_core_v1 import BIN_EDGES_S, labels_for_mapping


ROOT = Path(__file__).resolve().parents[1]
STRUCTURE = ROOT / "data" / "raw" / "asme30_structure"
SIGNAL = ROOT / "data" / "raw" / "asme30_signal"
OUTPUT = (
    ROOT / "data" / "derived" / "asme30_wz_hic_dataset_native_features_v2.npz"
)
AUDIT = (
    ROOT
    / "data"
    / "inventory"
    / "asme30_wz_hic_dataset_native_feature_audit_v2.json"
)
PROTOCOL = ROOT / "docs" / "wz_hic_asme30_dataset_native_protocol_v2.json"
LOCK = ROOT / "docs" / "wz_hic_asme30_dataset_native_implementation_lock_v2.json"
SIGNAL_MANIFEST = ROOT / "data" / "inventory" / "asme30_signal_manifest_v1.json"


def sha256(path: Path) -> str:
    h = hashlib.sha256()
    with path.open("rb") as f:
        for block in iter(lambda: f.read(1024 * 1024), b""):
            h.update(block)
    return h.hexdigest()


def filter_native(edf_path: Path, channel_rows) -> tuple[np.ndarray, np.ndarray]:
    raw = mne.io.read_raw_edf(
        edf_path, preload=True, infer_types=False, verbose="ERROR"
    )
    expected = [row["name"] for row in channel_rows]
    if raw.ch_names != expected or float(raw.info["sfreq"]) != 1000.0:
        raise RuntimeError(f"EDF metadata mismatch: {edf_path}")
    data = (raw.get_data() * 1e6).astype(np.float32)
    del raw
    eeg_idx = [i for i, row in enumerate(channel_rows) if row["type"] == "EEG"]
    eog_idx = [i for i, row in enumerate(channel_rows) if row["type"] == "EOG"]
    eeg, eog = data[eeg_idx], data[eog_idx]
    del data
    # The article describes a second-order 1-40 Hz Butterworth applied in the
    # forward direction to the complete offline/online stream.
    sos = butter(2, [1.0, 40.0], btype="bandpass", fs=1000.0, output="sos")
    return (
        sosfilt(sos, eeg, axis=1).astype(np.float32),
        sosfilt(sos, eog, axis=1).astype(np.float32),
    )


def process_run(subject: int, task: str, run: int) -> dict:
    stem = f"sub-{subject}_ses-1_task-{task}_run-{run}"
    structure_dir = STRUCTURE / f"sub-{subject}" / "ses-1" / "eeg"
    signal_dir = SIGNAL / f"sub-{subject}" / "ses-1" / "eeg"
    events = read_unique_events(structure_dir / f"{stem}_events.tsv")
    channels = read_channels(structure_dir / f"{stem}_channels.tsv")
    eeg, eog = filter_native(signal_dir / f"{stem}_eeg.edf", channels)
    bin_bounds = [
        (
            int(round((lo + 0.1) * 1000)),
            int(round((hi + 0.1) * 1000)),
        )
        for lo, hi in zip(BIN_EDGES_S[:-1], BIN_EDGES_S[1:])
    ]
    out = {
        key: []
        for key in (
            "eeg_bins",
            "eog_bins",
            "prestim_raw",
            "subject",
            "task",
            "run",
            "trial",
            "target",
            "command",
            "label",
            "sample",
            "accepted",
        )
    }
    target, trial_order = None, 0
    for event in events:
        code = int(float(event["trial_type"]))
        if 201 <= code <= 230:
            target, trial_order = code - 200, trial_order + 1
            continue
        if not (1 <= code <= 130):
            continue
        command = code % 100
        if not (1 <= command <= 30):
            continue
        if target is None or (code >= 101) != (command == target):
            raise RuntimeError(f"Event/target mismatch: {stem}")
        onset = int(float(event["sample"]))
        start, stop = onset - 100, onset + 1000
        if start < 0 or stop > eeg.shape[1]:
            raise RuntimeError(f"Epoch out of range: {stem}")
        epoch_eeg, epoch_eog = eeg[:, start:stop], eog[:, start:stop]
        eeg_bins = np.stack(
            [epoch_eeg[:, lo:hi].mean(axis=1) for lo, hi in bin_bounds], axis=1
        )
        eog_bins = np.stack(
            [epoch_eog[:, lo:hi].mean(axis=1) for lo, hi in bin_bounds], axis=1
        )
        out["eeg_bins"].append(eeg_bins.astype(np.float32))
        out["eog_bins"].append(eog_bins.astype(np.float32))
        out["prestim_raw"].append(eeg_bins[:, 0].astype(np.float32))
        out["subject"].append(subject)
        out["task"].append(task)
        out["run"].append(run)
        out["trial"].append((1000 if task == "online" else 0) + run * 10 + trial_order)
        out["target"].append(target)
        out["command"].append(command)
        out["sample"].append(onset)
        out["accepted"].append(True)
    if trial_order != 5 or len(out["command"]) != 2250:
        raise RuntimeError(f"Unexpected run structure: {stem}")
    out["label"] = labels_for_mapping(
        np.asarray(out["command"]), np.asarray(out["target"])
    ).tolist()
    return out


def main() -> None:
    parser = argparse.ArgumentParser()
    parser.add_argument("--overwrite", action="store_true")
    args = parser.parse_args()
    if OUTPUT.exists() and not args.overwrite:
        raise FileExistsError(OUTPUT)
    if not LOCK.exists():
        raise FileNotFoundError("Dataset-native implementation is not locked.")
    manifest = json.loads(SIGNAL_MANIFEST.read_text(encoding="utf-8"))
    if manifest["status"] != "PASS" or manifest["n_edf_files"] != 90:
        raise RuntimeError("Signal manifest incomplete.")

    keys = (
        "eeg_bins",
        "eog_bins",
        "prestim_raw",
        "subject",
        "task",
        "run",
        "trial",
        "target",
        "command",
        "label",
        "sample",
        "accepted",
    )
    merged = {key: [] for key in keys}
    run_summaries = []
    for subject in range(1, 11):
        for task, n_runs in (("offline", 6), ("online", 3)):
            for run in range(1, n_runs + 1):
                item = process_run(subject, task, run)
                for key in keys:
                    merged[key].extend(item[key])
                run_summaries.append(
                    {
                        "subject": subject,
                        "task": task,
                        "run": run,
                        "n_events": 2250,
                        "n_accepted": 2250,
                    }
                )
                print(f"sub-{subject} {task} run-{run}: 2250/2250", flush=True)
    arrays = {
        "eeg_bins": np.stack(merged["eeg_bins"]).astype(np.float32),
        "eog_bins": np.stack(merged["eog_bins"]).astype(np.float32),
        "prestim_raw": np.stack(merged["prestim_raw"]).astype(np.float32),
        "subject": np.asarray(merged["subject"], dtype=np.int8),
        "task": np.asarray(merged["task"], dtype="U7"),
        "run": np.asarray(merged["run"], dtype=np.int8),
        "trial": np.asarray(merged["trial"], dtype=np.int16),
        "target": np.asarray(merged["target"], dtype=np.int8),
        "command": np.asarray(merged["command"], dtype=np.int8),
        "label": np.asarray(merged["label"], dtype="U1"),
        "sample": np.asarray(merged["sample"], dtype=np.int32),
        "accepted": np.asarray(merged["accepted"], dtype=bool),
        "bin_edges_s": BIN_EDGES_S,
        "channel_names": np.asarray(
            [
                row["name"]
                for row in read_channels(
                    STRUCTURE
                    / "sub-1"
                    / "ses-1"
                    / "eeg"
                    / "sub-1_ses-1_task-offline_run-1_channels.tsv"
                )
                if row["type"] == "EEG"
            ],
            dtype="U8",
        ),
        "eog_names": np.asarray(["vEOG", "hEOG"], dtype="U8"),
    }
    OUTPUT.parent.mkdir(parents=True, exist_ok=True)
    np.savez_compressed(OUTPUT, **arrays)
    audit = {
        "status": "PASS",
        "created_utc": datetime.now(timezone.utc).isoformat(),
        "protocol_sha256": sha256(PROTOCOL),
        "implementation_lock_sha256": sha256(LOCK),
        "signal_manifest_sha256": sha256(SIGNAL_MANIFEST),
        "output": OUTPUT.relative_to(ROOT).as_posix(),
        "output_sha256": sha256(OUTPUT),
        "shape_eeg_bins": list(arrays["eeg_bins"].shape),
        "shape_eog_bins": list(arrays["eog_bins"].shape),
        "n_events_total": int(arrays["accepted"].size),
        "n_events_accepted": int(np.sum(arrays["accepted"])),
        "n_events_rejected": 0,
        "run_summaries": run_summaries,
        "software": {
            "python": sys.version,
            "platform": platform.platform(),
            "numpy": np.__version__,
            "scipy": scipy.__version__,
            "mne": mne.__version__,
        },
        "epistemic_label": (
            "Dataset-native feature robustness after input QC; no decoder outcome "
            "was computed in this builder."
        ),
    }
    AUDIT.parent.mkdir(parents=True, exist_ok=True)
    AUDIT.write_text(json.dumps(audit, indent=2) + "\n", encoding="utf-8")
    print(f"features={OUTPUT} sha256={audit['output_sha256']}")
    print(f"audit={AUDIT} sha256={sha256(AUDIT)}")


if __name__ == "__main__":
    main()
```
