## Supplementary material for "Falsifiable substitution tests reveal task-structured neural evidence for auditory attention": Article_Analysis_Source_Code: build_asme30_wz_hic_features_v1.py

```
#!/usr/bin/env python3
"""Build fixed ASME30 event features for the external WZ-HIC-v1 test."""

from __future__ import annotations

import argparse
import csv
import hashlib
import json
import platform
import re
import sys
from datetime import datetime, timezone
from pathlib import Path

import mne
import numpy as np
import scipy
from scipy.signal import butter, resample_poly, sosfiltfilt

from wz_hic_asme30_core_v1 import BIN_EDGES_S, labels_for_mapping


ROOT = Path(__file__).resolve().parents[1]
STRUCTURE = ROOT / "data" / "raw" / "asme30_structure"
SIGNAL = ROOT / "data" / "raw" / "asme30_signal"
OUTPUT = ROOT / "data" / "derived" / "asme30_wz_hic_features_v1.npz"
AUDIT = ROOT / "data" / "inventory" / "asme30_wz_hic_feature_audit_v1.json"
PROTOCOL = ROOT / "docs" / "wz_hic_asme30_external_protocol_v1.json"
LOCK = ROOT / "docs" / "wz_hic_asme30_external_implementation_lock_v1.json"
SIGNAL_MANIFEST = ROOT / "data" / "inventory" / "asme30_signal_manifest_v1.json"


def sha256(path: Path) -> str:
    h = hashlib.sha256()
    with path.open("rb") as f:
        for block in iter(lambda: f.read(1024 * 1024), b""):
            h.update(block)
    return h.hexdigest()


def read_channels(path: Path) -> list[dict[str, str]]:
    with path.open("r", encoding="utf-8-sig", newline="") as f:
        return list(csv.DictReader(f, delimiter="\t"))


def read_unique_events(path: Path) -> list[dict[str, str]]:
    with path.open("r", encoding="utf-8-sig", newline="") as f:
        rows = list(csv.DictReader(f, delimiter="\t"))
    keys = ("onset", "duration", "trial_type", "value", "sample")
    out = []
    previous = None
    duplicates = 0
    for row in rows:
        key = tuple(row[k] for k in keys)
        if key == previous:
            duplicates += 1
            continue
        out.append(row)
        previous = key
    if len(rows) != 2 * len(out) or duplicates != len(out):
        raise RuntimeError(f"Unexpected event duplication in {path}")
    return out


def filter_resample(
    edf_path: Path, channel_rows: list[dict[str, str]]
) -> tuple[np.ndarray, np.ndarray]:
    raw = mne.io.read_raw_edf(
        edf_path, preload=True, infer_types=False, verbose="ERROR"
    )
    expected = [row["name"] for row in channel_rows]
    if raw.ch_names != expected:
        raise RuntimeError(
            f"EDF/sidecar channel mismatch in {edf_path}: "
            f"{raw.ch_names!r} != {expected!r}"
        )
    if float(raw.info["sfreq"]) != 1000.0:
        raise RuntimeError(f"Unexpected sampling frequency: {raw.info['sfreq']}")
    data = (raw.get_data() * 1e6).astype(np.float32)  # V -> microvolts
    del raw
    eeg_idx = [i for i, row in enumerate(channel_rows) if row["type"] == "EEG"]
    eog_idx = [i for i, row in enumerate(channel_rows) if row["type"] == "EOG"]
    if len(eeg_idx) != 64 or len(eog_idx) != 2:
        raise RuntimeError(f"Unexpected channel types in {edf_path}")
    eeg = data[eeg_idx]
    eog = data[eog_idx]
    del data
    eeg -= eeg.mean(axis=0, keepdims=True)
    sos = butter(4, [0.1, 8.0], btype="bandpass", fs=1000.0, output="sos")
    eeg = sosfiltfilt(sos, eeg, axis=1).astype(np.float32)
    eog = sosfiltfilt(sos, eog, axis=1).astype(np.float32)
    eeg = resample_poly(eeg, 1, 4, axis=1).astype(np.float32)
    eog = resample_poly(eog, 1, 4, axis=1).astype(np.float32)
    return eeg, eog


def process_run(subject: int, task: str, run: int) -> dict:
    stem = f"sub-{subject}_ses-1_task-{task}_run-{run}"
    structure_dir = STRUCTURE / f"sub-{subject}" / "ses-1" / "eeg"
    signal_dir = SIGNAL / f"sub-{subject}" / "ses-1" / "eeg"
    events_path = structure_dir / f"{stem}_events.tsv"
    channels_path = structure_dir / f"{stem}_channels.tsv"
    edf_path = signal_dir / f"{stem}_eeg.edf"
    if not edf_path.exists():
        raise FileNotFoundError(edf_path)
    events = read_unique_events(events_path)
    channel_rows = read_channels(channels_path)
    eeg, eog = filter_resample(edf_path, channel_rows)
    fs = 250
    start_offset, stop_offset = -25, 250
    baseline_stop = 25
    bin_bounds = [
        (
            int(round((lo + 0.1) * fs)),
            int(round((hi + 0.1) * fs)),
        )
        for lo, hi in zip(BIN_EDGES_S[:-1], BIN_EDGES_S[1:])
    ]

    out = {
        key: []
        for key in (
            "eeg_bins",
            "eog_bins",
            "prestim_raw",
            "subject",
            "task",
            "run",
            "trial",
            "target",
            "command",
            "label",
            "sample",
            "accepted",
        )
    }
    target = None
    trial_order = 0
    for event in events:
        code = int(float(event["trial_type"]))
        if 201 <= code <= 230:
            target = code - 200
            trial_order += 1
            continue
        if not (1 <= code <= 130):
            continue
        if target is None:
            raise RuntimeError(f"Stimulus before target marker in {events_path}")
        command = code % 100
        if not (1 <= command <= 30):
            continue
        marker_is_target = code >= 101
        if marker_is_target != (command == target):
            raise RuntimeError(f"Target encoding mismatch in {events_path}")
        sample_1000 = int(float(event["sample"]))
        onset = int(round(sample_1000 / 4))
        start, stop = onset + start_offset, onset + stop_offset
        if start < 0 or stop > eeg.shape[1]:
            raise RuntimeError(f"Epoch out of range in {edf_path}: {sample_1000}")
        epoch_eeg_raw = eeg[:, start:stop].copy()
        epoch_eog_raw = eog[:, start:stop].copy()
        prestim_raw = epoch_eeg_raw[:, :baseline_stop].mean(axis=1)
        epoch_eeg = (
            epoch_eeg_raw
            - epoch_eeg_raw[:, :baseline_stop].mean(axis=1, keepdims=True)
        )
        epoch_eog = (
            epoch_eog_raw
            - epoch_eog_raw[:, :baseline_stop].mean(axis=1, keepdims=True)
        )
        accepted = bool(np.max(np.ptp(epoch_eeg, axis=1)) <= 200.0)
        eeg_bins = np.stack(
            [epoch_eeg[:, lo:hi].mean(axis=1) for lo, hi in bin_bounds], axis=1
        )
        eog_bins = np.stack(
            [epoch_eog[:, lo:hi].mean(axis=1) for lo, hi in bin_bounds], axis=1
        )
        out["eeg_bins"].append(eeg_bins.astype(np.float32))
        out["eog_bins"].append(eog_bins.astype(np.float32))
        out["prestim_raw"].append(prestim_raw.astype(np.float32))
        out["subject"].append(subject)
        out["task"].append(task)
        out["run"].append(run)
        out["trial"].append((1000 if task == "online" else 0) + run * 10 + trial_order)
        out["target"].append(target)
        out["command"].append(command)
        out["sample"].append(sample_1000)
        out["accepted"].append(accepted)
    if trial_order != 5 or len(out["command"]) != 2250:
        raise RuntimeError(
            f"Unexpected run structure {subject} {task} {run}: "
            f"{trial_order} trials, {len(out['command'])} stimuli"
        )
    out["label"] = labels_for_mapping(
        np.asarray(out["command"]), np.asarray(out["target"])
    ).tolist()
    del eeg, eog
    return out


def main() -> None:
    parser = argparse.ArgumentParser()
    parser.add_argument("--overwrite", action="store_true")
    args = parser.parse_args()
    if OUTPUT.exists() and not args.overwrite:
        raise FileExistsError(f"Use --overwrite explicitly: {OUTPUT}")
    if not LOCK.exists():
        raise FileNotFoundError("External implementation must be locked first.")
    manifest = json.loads(SIGNAL_MANIFEST.read_text(encoding="utf-8"))
    if manifest["status"] != "PASS" or manifest["n_edf_files"] != 90:
        raise RuntimeError("Signal manifest is incomplete.")

    merged = {
        key: []
        for key in (
            "eeg_bins",
            "eog_bins",
            "prestim_raw",
            "subject",
            "task",
            "run",
            "trial",
            "target",
            "command",
            "label",
            "sample",
            "accepted",
        )
    }
    run_summaries = []
    for subject in range(1, 11):
        for task, n_runs in (("offline", 6), ("online", 3)):
            for run in range(1, n_runs + 1):
                item = process_run(subject, task, run)
                for key in merged:
                    merged[key].extend(item[key])
                run_summaries.append(
                    {
                        "subject": subject,
                        "task": task,
                        "run": run,
                        "n_events": len(item["accepted"]),
                        "n_accepted": int(sum(item["accepted"])),
                    }
                )
                print(
                    f"sub-{subject} {task} run-{run}: "
                    f"{sum(item['accepted'])}/2250 accepted",
                    flush=True,
                )
    arrays = {
        "eeg_bins": np.stack(merged["eeg_bins"]).astype(np.float32),
        "eog_bins": np.stack(merged["eog_bins"]).astype(np.float32),
        "prestim_raw": np.stack(merged["prestim_raw"]).astype(np.float32),
        "subject": np.asarray(merged["subject"], dtype=np.int8),
        "task": np.asarray(merged["task"], dtype="U7"),
        "run": np.asarray(merged["run"], dtype=np.int8),
        "trial": np.asarray(merged["trial"], dtype=np.int16),
        "target": np.asarray(merged["target"], dtype=np.int8),
        "command": np.asarray(merged["command"], dtype=np.int8),
        "label": np.asarray(merged["label"], dtype="U1"),
        "sample": np.asarray(merged["sample"], dtype=np.int32),
        "accepted": np.asarray(merged["accepted"], dtype=bool),
        "bin_edges_s": BIN_EDGES_S,
        "channel_names": np.asarray(
            [
                row["name"]
                for row in read_channels(
                    STRUCTURE
                    / "sub-1"
                    / "ses-1"
                    / "eeg"
                    / "sub-1_ses-1_task-offline_run-1_channels.tsv"
                )
                if row["type"] == "EEG"
            ],
            dtype="U8",
        ),
        "eog_names": np.asarray(["vEOG", "hEOG"], dtype="U8"),
    }
    OUTPUT.parent.mkdir(parents=True, exist_ok=True)
    np.savez_compressed(OUTPUT, **arrays)
    accepted = arrays["accepted"]
    audit = {
        "status": "PASS",
        "created_utc": datetime.now(timezone.utc).isoformat(),
        "protocol_sha256": sha256(PROTOCOL),
        "implementation_lock_sha256": sha256(LOCK),
        "signal_manifest_sha256": sha256(SIGNAL_MANIFEST),
        "output": OUTPUT.relative_to(ROOT).as_posix(),
        "output_sha256": sha256(OUTPUT),
        "shape_eeg_bins": list(arrays["eeg_bins"].shape),
        "shape_eog_bins": list(arrays["eog_bins"].shape),
        "shape_prestim_raw": list(arrays["prestim_raw"].shape),
        "n_events_total": int(accepted.size),
        "n_events_accepted": int(np.sum(accepted)),
        "n_events_rejected": int(np.sum(~accepted)),
        "run_summaries": run_summaries,
        "software": {
            "python": sys.version,
            "platform": platform.platform(),
            "numpy": np.__version__,
            "scipy": scipy.__version__,
            "mne": mne.__version__,
        },
        "outcome_access": (
            "Feature extraction used only frozen event labels and fixed signal "
            "operations; it did not fit a decoder or compute online accuracy."
        ),
    }
    AUDIT.parent.mkdir(parents=True, exist_ok=True)
    AUDIT.write_text(json.dumps(audit, indent=2) + "\n", encoding="utf-8")
    print(f"features={OUTPUT} sha256={audit['output_sha256']}")
    print(f"audit={AUDIT} sha256={sha256(AUDIT)}")


if __name__ == "__main__":
    main()
```
