## Supplementary material for "Falsifiable substitution tests reveal task-structured neural evidence for auditory attention": Article_Analysis_Source_Code: build_kojima2024b_wz_hic_confirmation_features_v1.py

```
#!/usr/bin/env python
"""Build the sealed-participant features for WZ-HIC-v1 confirmation."""

from __future__ import annotations

import argparse
import hashlib
import json
import re
from datetime import datetime, timezone
from pathlib import Path

import numpy as np

from build_kojima2024b_wz_hif_discovery_features_v1 import (
    BIN_EDGES,
    parse_header,
    process_run,
)


ROOT = Path(__file__).resolve().parents[1]
DATA_ROOT = ROOT / "data" / "raw" / "kojima2024b_structure_all"
OUTPUT = (
    ROOT / "data" / "derived" / "kojima2024b_wz_hic_confirmation_features_v1.npz"
)
AUDIT = (
    ROOT
    / "data"
    / "inventory"
    / "kojima2024b_wz_hic_confirmation_feature_audit_v1.json"
)
PROTOCOL = ROOT / "docs" / "wz_hic_kojima2024b_confirmation_protocol_v1.json"
CONFIRMATION = tuple("BDFHJLN")
TASKS = ("2stream", "4stream")


def sha256(path: Path) -> str:
    h = hashlib.sha256()
    with path.open("rb") as f:
        for block in iter(lambda: f.read(1024 * 1024), b""):
            h.update(block)
    return h.hexdigest()


def main() -> None:
    parser = argparse.ArgumentParser()
    parser.add_argument("--overwrite", action="store_true")
    args = parser.parse_args()
    if OUTPUT.exists() and not args.overwrite:
        raise FileExistsError(OUTPUT)
    if list(DATA_ROOT.glob("sub-*/nasatlx/*")):
        raise RuntimeError("NASA-TLX must remain sealed.")
    merged = {
        k: []
        for k in (
            "eeg_bins",
            "eog_bins",
            "subject",
            "task",
            "run",
            "trial",
            "target",
            "command",
            "label",
            "accepted",
        )
    }
    source_files, run_summaries = [], []
    final_header = None
    for subject in CONFIRMATION:
        eeg_dir = DATA_ROOT / f"sub-{subject}" / "eeg"
        for task in TASKS:
            vhdrs = sorted(
                eeg_dir.glob(f"sub-{subject}_task-{task}_run-*_eeg.vhdr"),
                key=lambda p: int(re.search(r"_run-(\d+)_", p.name).group(1)),
            )
            if len(vhdrs) != 6:
                raise RuntimeError(f"{subject} {task}: {len(vhdrs)} headers.")
            for vhdr in vhdrs:
                item = process_run(subject, task, vhdr)
                for key in merged:
                    merged[key].extend(item[key])
                final_header = parse_header(vhdr)
                data_path = final_header["data_path"]
                source_files.append(
                    {
                        "path": str(data_path.relative_to(ROOT)).replace("\\", "/"),
                        "bytes": data_path.stat().st_size,
                        "sha256": sha256(data_path),
                    }
                )
                run_summaries.append(
                    {
                        "subject": subject,
                        "task": task,
                        "run": int(item["run"][0]),
                        "n_events": len(item["accepted"]),
                        "n_accepted": int(sum(item["accepted"])),
                    }
                )
                print(
                    subject,
                    task,
                    item["run"][0],
                    sum(item["accepted"]),
                    "/",
                    len(item["accepted"]),
                    flush=True,
                )
    arrays = {
        "eeg_bins": np.stack(merged["eeg_bins"]).astype(np.float32),
        "eog_bins": np.stack(merged["eog_bins"]).astype(np.float32),
        "subject": np.asarray(merged["subject"], dtype="U1"),
        "task": np.asarray(merged["task"], dtype="U8"),
        "run": np.asarray(merged["run"], dtype=np.int16),
        "trial": np.asarray(merged["trial"], dtype=np.int16),
        "target": np.asarray(merged["target"], dtype=np.int8),
        "command": np.asarray(merged["command"], dtype=np.int8),
        "label": np.asarray(merged["label"], dtype="U1"),
        "accepted": np.asarray(merged["accepted"], dtype=bool),
        "bin_edges_s": BIN_EDGES,
        "channel_names": np.asarray(
            [c["name"] for c in final_header["channels"][:64]], dtype="U8"
        ),
        "eog_names": np.asarray(["vEOG", "hEOG"], dtype="U8"),
    }
    OUTPUT.parent.mkdir(parents=True, exist_ok=True)
    np.savez_compressed(OUTPUT, **arrays)
    audit = {
        "status": "PASS",
        "created_utc": datetime.now(timezone.utc).isoformat(),
        "protocol_sha256": sha256(PROTOCOL),
        "output": str(OUTPUT.relative_to(ROOT)).replace("\\", "/"),
        "output_sha256": sha256(OUTPUT),
        "shape_eeg_bins": list(arrays["eeg_bins"].shape),
        "shape_eog_bins": list(arrays["eog_bins"].shape),
        "n_events_total": int(arrays["accepted"].size),
        "n_events_accepted": int(arrays["accepted"].sum()),
        "n_events_rejected": int((~arrays["accepted"]).sum()),
        "source_files": source_files,
        "run_summaries": run_summaries,
        "NASA_TLX_value_files_present": len(
            list(DATA_ROOT.glob("sub-*/nasatlx/*"))
        ),
        "signal_split": "confirmation_only",
    }
    AUDIT.parent.mkdir(parents=True, exist_ok=True)
    AUDIT.write_text(
        json.dumps(audit, ensure_ascii=False, indent=2) + "\n", encoding="utf-8"
    )
    print(OUTPUT, sha256(OUTPUT))
    print(AUDIT, sha256(AUDIT))


if __name__ == "__main__":
    main()
```
