## Supplementary material for "Falsifiable substitution tests reveal task-structured neural evidence for auditory attention": Article_Analysis_Source_Code: build_kojima2024b_wz_hif_discovery_features_v1.py

```
#!/usr/bin/env python
"""Build outcome-blind event features for the WZ-HIF-v1 discovery split.

The script reads only the EEG binaries for A,C,E,G,I,K,M,O and never reads the
NASA-TLX folders. Each run is processed independently and the output contains
fixed 100-ms EEG/EOG bin means plus event metadata, not analysis outcomes.
"""

from __future__ import annotations

import argparse
import hashlib
import json
import re
from datetime import datetime, timezone
from pathlib import Path

import numpy as np
from scipy.signal import butter, resample_poly, sosfiltfilt


ROOT = Path(__file__).resolve().parents[1]
DATA_ROOT = ROOT / "data" / "raw" / "kojima2024b_structure_all"
OUTPUT = ROOT / "data" / "derived" / "kojima2024b_wz_hif_discovery_features_v1.npz"
AUDIT = (
    ROOT
    / "data"
    / "inventory"
    / "kojima2024b_wz_hif_discovery_feature_audit_v1.json"
)
PROTOCOL = ROOT / "docs" / "wz_hif_kojima2024b_discovery_protocol_v1.json"
DISCOVERY = tuple("ACEGIKMO")
CONFIRMATION = tuple("BDFHJLN")
TASKS = ("2stream", "4stream")
BIN_EDGES = np.round(np.arange(-0.1, 1.0001, 0.1), 10)


def sha256(path: Path) -> str:
    h = hashlib.sha256()
    with path.open("rb") as f:
        for block in iter(lambda: f.read(1024 * 1024), b""):
            h.update(block)
    return h.hexdigest()


def parse_header(path: Path) -> dict:
    text = path.read_text(encoding="utf-8", errors="replace")
    get = lambda pattern: re.search(pattern, text, flags=re.MULTILINE).group(1).strip()
    n_channels = int(get(r"^NumberOfChannels=(.+)$"))
    interval_us = float(get(r"^SamplingInterval=(.+)$"))
    data_file = get(r"^DataFile=(.+)$")
    marker_file = get(r"^MarkerFile=(.+)$")
    fmt = get(r"^BinaryFormat=(.+)$")
    channels = []
    for line in text.splitlines():
        match = re.match(r"^Ch(\d+)=(.*)$", line)
        if not match:
            continue
        fields = match.group(2).split(",")
        channels.append(
            {
                "index": int(match.group(1)),
                "name": fields[0],
                "resolution": float(fields[2]),
            }
        )
    channels.sort(key=lambda x: x["index"])
    if fmt != "INT_16" or n_channels != 66 or len(channels) != 66:
        raise RuntimeError(f"Unexpected BrainVision header: {path}")
    return {
        "n_channels": n_channels,
        "sampling_hz": 1e6 / interval_us,
        "data_path": path.parent / data_file,
        "marker_path": path.parent / marker_file,
        "channels": channels,
    }


def parse_markers(path: Path) -> list[tuple[int, str]]:
    events = []
    for line in path.read_text(encoding="utf-8", errors="replace").splitlines():
        if not re.match(r"^Mk\d+=", line):
            continue
        fields = line.split("=", 1)[1].split(",")
        if len(fields) < 3 or fields[0] != "Stimulus":
            continue
        desc = fields[1].replace(" ", "")
        if not desc.startswith("S"):
            continue
        events.append((int(fields[2]) - 1, desc))
    events.sort()
    return events


def label_2stream(command: int, target: int) -> str:
    if command == target:
        return "T"
    target_parent = 1 if target in (1, 2) else 2
    command_parent = 1 if command in (1, 2) else 2
    return "S" if command_parent == target_parent else "U"


def filter_resample_run(header: dict) -> tuple[np.ndarray, np.ndarray]:
    raw_path = header["data_path"]
    if not raw_path.exists():
        raise FileNotFoundError(raw_path)
    resolutions = np.asarray(
        [c["resolution"] for c in header["channels"]], dtype=np.float32
    )
    binary = np.memmap(raw_path, dtype="<i2", mode="r")
    if binary.size % 66:
        raise RuntimeError(f"Binary length is not divisible by 66: {raw_path}")
    data = np.asarray(binary.reshape(-1, 66), dtype=np.float32)
    data *= resolutions[None, :]  # microvolts
    eeg = data[:, :64]
    eeg -= eeg.mean(axis=1, keepdims=True)
    eog = data[:, 64:66]

    fs = float(header["sampling_hz"])
    if fs != 1000.0:
        raise RuntimeError(f"Expected 1000 Hz, found {fs}.")
    sos = butter(4, [0.1, 8.0], btype="bandpass", fs=fs, output="sos")
    eeg = sosfiltfilt(sos, eeg, axis=0).astype(np.float32)
    eog = sosfiltfilt(sos, eog, axis=0).astype(np.float32)
    eeg_250 = resample_poly(eeg, 1, 4, axis=0).astype(np.float32)
    eog_250 = resample_poly(eog, 1, 4, axis=0).astype(np.float32)
    return eeg_250, eog_250


def process_run(subject: str, task: str, vhdr: Path) -> dict[str, list]:
    header = parse_header(vhdr)
    events = parse_markers(header["marker_path"])
    eeg, eog = filter_resample_run(header)
    fs = 250
    start_offset = int(round(-0.1 * fs))
    stop_offset = int(round(1.0 * fs))
    baseline_stop = int(round(0.1 * fs))

    out = {
        "eeg_bins": [],
        "eog_bins": [],
        "subject": [],
        "task": [],
        "run": [],
        "trial": [],
        "target": [],
        "command": [],
        "label": [],
        "accepted": [],
    }
    run_match = re.search(r"_run-(\d+)_", vhdr.name)
    run = int(run_match.group(1))
    target = None
    trial_order = 0
    for position_1000, description in events:
        code = int(description[1:])
        if 201 <= code <= 204:
            target = code - 200
            trial_order += 1
            continue
        if not (101 <= code <= 104 or 111 <= code <= 114):
            continue
        if target is None:
            raise RuntimeError(f"Deviant before trial marker in {vhdr}.")
        command = code - 110 if code >= 111 else code - 100
        onset = int(round(position_1000 / 4))
        start, stop = onset + start_offset, onset + stop_offset
        if start < 0 or stop > eeg.shape[0]:
            continue
        epoch_eeg = eeg[start:stop].T.copy()
        epoch_eog = eog[start:stop].T.copy()
        epoch_eeg -= epoch_eeg[:, :baseline_stop].mean(axis=1, keepdims=True)
        epoch_eog -= epoch_eog[:, :baseline_stop].mean(axis=1, keepdims=True)
        accepted = bool(np.ptp(epoch_eeg, axis=1).max() <= 200.0)
        eeg_bins = np.stack(
            [
                epoch_eeg[:, int(round((edge + 0.1) * fs)) : int(
                    round((BIN_EDGES[i + 1] + 0.1) * fs)
                )].mean(axis=1)
                for i, edge in enumerate(BIN_EDGES[:-1])
            ],
            axis=1,
        )
        eog_bins = np.stack(
            [
                epoch_eog[:, int(round((edge + 0.1) * fs)) : int(
                    round((BIN_EDGES[i + 1] + 0.1) * fs)
                )].mean(axis=1)
                for i, edge in enumerate(BIN_EDGES[:-1])
            ],
            axis=1,
        )
        label = (
            label_2stream(command, target)
            if task == "2stream"
            else ("T" if command == target else "U")
        )
        out["eeg_bins"].append(eeg_bins.astype(np.float32))
        out["eog_bins"].append(eog_bins.astype(np.float32))
        out["subject"].append(subject)
        out["task"].append(task)
        out["run"].append(run)
        out["trial"].append(run * 10 + trial_order)
        out["target"].append(target)
        out["command"].append(command)
        out["label"].append(label)
        out["accepted"].append(accepted)

    if trial_order != 4 or len(out["target"]) != 240:
        raise RuntimeError(
            f"Unexpected structure {subject} {task} run {run}: "
            f"{trial_order} trials, {len(out['target'])} deviants."
        )
    return out


def main() -> None:
    parser = argparse.ArgumentParser()
    parser.add_argument("--overwrite", action="store_true")
    args = parser.parse_args()
    if OUTPUT.exists() and not args.overwrite:
        raise FileExistsError(f"Use --overwrite explicitly: {OUTPUT}")
    if not PROTOCOL.exists():
        raise FileNotFoundError(PROTOCOL)

    confirmation_binaries = []
    for subject in CONFIRMATION:
        confirmation_binaries.extend(
            (DATA_ROOT / f"sub-{subject}" / "eeg").glob("*.eeg")
        )
    if confirmation_binaries:
        raise RuntimeError(
            f"Confirmation seal broken: {len(confirmation_binaries)} binaries exist."
        )
    nasa_files = list(DATA_ROOT.glob("sub-*/nasatlx/*"))
    if nasa_files:
        raise RuntimeError(f"NASA seal broken: {len(nasa_files)} value files exist.")

    merged: dict[str, list] = {
        k: []
        for k in (
            "eeg_bins",
            "eog_bins",
            "subject",
            "task",
            "run",
            "trial",
            "target",
            "command",
            "label",
            "accepted",
        )
    }
    source_files = []
    run_summaries = []
    for subject in DISCOVERY:
        eeg_dir = DATA_ROOT / f"sub-{subject}" / "eeg"
        for task in TASKS:
            vhdrs = sorted(
                eeg_dir.glob(f"sub-{subject}_task-{task}_run-*_eeg.vhdr"),
                key=lambda p: int(re.search(r"_run-(\d+)_", p.name).group(1)),
            )
            if len(vhdrs) != 6:
                raise RuntimeError(f"{subject} {task}: expected 6 headers, got {len(vhdrs)}")
            for vhdr in vhdrs:
                item = process_run(subject, task, vhdr)
                for key in merged:
                    merged[key].extend(item[key])
                data_path = parse_header(vhdr)["data_path"]
                source_files.append(
                    {
                        "path": str(data_path.relative_to(ROOT)).replace("\\", "/"),
                        "bytes": data_path.stat().st_size,
                        "sha256": sha256(data_path),
                    }
                )
                run_summaries.append(
                    {
                        "subject": subject,
                        "task": task,
                        "run": int(item["run"][0]),
                        "n_events": len(item["accepted"]),
                        "n_accepted": int(sum(item["accepted"])),
                    }
                )
                print(
                    subject,
                    task,
                    item["run"][0],
                    sum(item["accepted"]),
                    "/",
                    len(item["accepted"]),
                    flush=True,
                )

    arrays = {
        "eeg_bins": np.stack(merged["eeg_bins"]).astype(np.float32),
        "eog_bins": np.stack(merged["eog_bins"]).astype(np.float32),
        "subject": np.asarray(merged["subject"], dtype="U1"),
        "task": np.asarray(merged["task"], dtype="U8"),
        "run": np.asarray(merged["run"], dtype=np.int16),
        "trial": np.asarray(merged["trial"], dtype=np.int16),
        "target": np.asarray(merged["target"], dtype=np.int8),
        "command": np.asarray(merged["command"], dtype=np.int8),
        "label": np.asarray(merged["label"], dtype="U1"),
        "accepted": np.asarray(merged["accepted"], dtype=bool),
        "bin_edges_s": BIN_EDGES,
        "channel_names": np.asarray(
            [c["name"] for c in parse_header(vhdr)["channels"][:64]], dtype="U8"
        ),
        "eog_names": np.asarray(["vEOG", "hEOG"], dtype="U8"),
    }
    OUTPUT.parent.mkdir(parents=True, exist_ok=True)
    np.savez_compressed(OUTPUT, **arrays)

    accepted = arrays["accepted"]
    audit = {
        "status": "PASS",
        "created_utc": datetime.now(timezone.utc).isoformat(),
        "protocol_sha256": sha256(PROTOCOL),
        "output": str(OUTPUT.relative_to(ROOT)).replace("\\", "/"),
        "output_sha256": sha256(OUTPUT),
        "shape_eeg_bins": list(arrays["eeg_bins"].shape),
        "shape_eog_bins": list(arrays["eog_bins"].shape),
        "n_events_total": int(accepted.size),
        "n_events_accepted": int(accepted.sum()),
        "n_events_rejected": int((~accepted).sum()),
        "source_files": source_files,
        "run_summaries": run_summaries,
        "confirmation_binaries_present": 0,
        "nasa_tlx_value_files_present": 0,
        "signal_split": "discovery_only",
    }
    AUDIT.parent.mkdir(parents=True, exist_ok=True)
    AUDIT.write_text(
        json.dumps(audit, ensure_ascii=False, indent=2) + "\n", encoding="utf-8"
    )
    print(OUTPUT, sha256(OUTPUT))
    print(AUDIT, sha256(AUDIT))


if __name__ == "__main__":
    main()
```
