## Supplementary material for "Falsifiable substitution tests reveal task-structured neural evidence for auditory attention": Article_Analysis_Source_Code: build_wz_hic_individual_geometry_synthesis_v1.py

```
#!/usr/bin/env python3
"""Build a descriptive cross-dataset synthesis of individual WZ-HIC geometry."""

from __future__ import annotations

import csv
import hashlib
import json
from datetime import datetime, timezone
from pathlib import Path


ROOT = Path(__file__).resolve().parents[1]
ASME = (
    ROOT
    / "results"
    / "wz_hic_asme30"
    / "crossphase_individual_geometry_v1"
    / "crossphase_decision_v1.json"
)
ASME_AUDIT = (
    ROOT
    / "results"
    / "wz_hic_asme30"
    / "crossphase_individual_geometry_v1"
    / "independent_source_audit_v1.json"
)
KOJIMA = (
    ROOT
    / "results"
    / "wz_hic_kojima2024b"
    / "crossrun_individual_geometry_v1"
    / "crossrun_decision_v1.json"
)
KOJIMA_AUDIT = (
    ROOT
    / "results"
    / "wz_hic_kojima2024b"
    / "crossrun_individual_geometry_v1"
    / "independent_source_audit_v1.json"
)
RESIDUAL = (
    ROOT
    / "results"
    / "wz_hic_cross_dataset"
    / "command_identity_residual_v1"
    / "residual_decision_v1.json"
)
RESIDUAL_AUDIT = (
    ROOT
    / "results"
    / "wz_hic_cross_dataset"
    / "command_identity_residual_v1"
    / "independent_source_audit_v1.json"
)
SHARED_TEMPLATE = (
    ROOT
    / "results"
    / "wz_hic_asme30"
    / "target_free_spatiotemporal_v1"
    / "spatiotemporal_decision_v1.json"
)
SHARED_TEMPLATE_AUDIT = (
    ROOT
    / "results"
    / "wz_hic_asme30"
    / "target_free_spatiotemporal_v1"
    / "independent_audit_v2.json"
)
OUTDIR = (
    ROOT
    / "results"
    / "wz_hic_cross_dataset"
    / "individual_geometry_synthesis_v1"
)
OUTPUT = OUTDIR / "synthesis_v1.json"
CSV = OUTDIR / "participant_metrics_v1.csv"


def sha256(path: Path) -> str:
    digest = hashlib.sha256()
    with path.open("rb") as handle:
        for block in iter(lambda: handle.read(1024 * 1024), b""):
            digest.update(block)
    return digest.hexdigest()


def main() -> None:
    if OUTPUT.exists() or CSV.exists():
        raise FileExistsError("Immutable synthesis v1 already exists.")
    asme = json.loads(ASME.read_text(encoding="utf-8"))
    kojima = json.loads(KOJIMA.read_text(encoding="utf-8"))
    residual = json.loads(RESIDUAL.read_text(encoding="utf-8"))
    shared = json.loads(SHARED_TEMPLATE.read_text(encoding="utf-8"))
    audits = {
        "ASME30": json.loads(ASME_AUDIT.read_text(encoding="utf-8")),
        "Kojima2024B": json.loads(
            KOJIMA_AUDIT.read_text(encoding="utf-8")
        ),
        "command_residual": json.loads(
            RESIDUAL_AUDIT.read_text(encoding="utf-8")
        ),
        "shared_template": json.loads(
            SHARED_TEMPLATE_AUDIT.read_text(encoding="utf-8")
        ),
    }
    if not all(item["audit_pass"] for item in audits.values()):
        raise RuntimeError("A source audit did not pass.")

    rows = []
    residual_lookup = {
        dataset: {
            str(row["subject"]): row
            for row in residual["participant_results"][dataset]
        }
        for dataset in ("ASME30", "Kojima2024B")
    }
    for dataset, source, split_label in (
        ("ASME30", asme, "offline-to-later-online"),
        ("Kojima2024B", kojima, "leave-one-run-out"),
    ):
        for row in source["participant_geometry"]:
            residual_row = residual_lookup[dataset][str(row["subject"])]
            rows.append(
                {
                    "dataset": dataset,
                    "participant": str(row["subject"]),
                    "split": split_label,
                    "physical_effect": row["physical_full_effect"],
                    "best_wrong_effect": row["best_wrong_full_effect"],
                    "binding_specificity": row["binding_specificity"],
                    "eog_effect": row["physical_eog_effect"],
                    "nonocular_specificity": row["nonocular_specificity"],
                    "within_person_alignment": row[
                        "physical_within_alignment"
                    ],
                    "between_person_substitution": row[
                        "between_person_substitution"
                    ],
                    "identity_advantage": row["identity_advantage"],
                    "mapping_alignment_specificity": row[
                        "mapping_alignment_specificity"
                    ],
                    "residual_physical_effect": residual_row[
                        "physical_effect"
                    ],
                    "residual_binding_specificity": residual_row[
                        "binding_specificity"
                    ],
                    "physical_retention_ratio": residual_row[
                        "residual_to_raw_physical_ratio"
                    ],
                    "binding_retention_ratio": residual_row[
                        "residual_to_raw_binding_ratio"
                    ],
                }
            )

    positive_counts = {
        metric: {
            "positive": sum(float(row[metric]) > 0 for row in rows),
            "total": len(rows),
        }
        for metric in (
            "physical_effect",
            "binding_specificity",
            "nonocular_specificity",
            "identity_advantage",
            "mapping_alignment_specificity",
            "residual_physical_effect",
            "residual_binding_specificity",
        )
    }
    output = {
        "synthesis": "WZ-HIC-cross-dataset-individual-geometry-v1",
        "created_utc": datetime.now(timezone.utc).isoformat(),
        "analysis_class": (
            "Post-outcome descriptive source synthesis. Dataset-specific "
            "locked inference is retained; no pooled four-point regression "
            "or exchangeability claim is made."
        ),
        "status": "cross_dataset_individual_geometry_evidence_chain_complete",
        "source_conjunctions": {
            "ASME30_all_geometry_gates": all(asme["gates"].values()),
            "Kojima2024B_all_geometry_gates": all(
                kojima["gates"].values()
            ),
            "command_identity_all_gates": all(
                residual["gates"].values()
            ),
            "all_independent_audits_pass": all(
                item["audit_pass"] for item in audits.values()
            ),
        },
        "dataset_summaries": {
            "ASME30": {
                "n": 10,
                "split": "offline-to-later-online",
                "geometry": asme["summaries"],
                "command_residual": residual["summaries"]["ASME30"],
                "time_resolved": asme[
                    "time_resolved_binding_specificity"
                ],
            },
            "Kojima2024B": {
                "n": 7,
                "split": "leave-one-run-out",
                "geometry": kojima["summaries"],
                "command_residual": residual["summaries"][
                    "Kojima2024B"
                ],
                "time_resolved": kojima[
                    "time_resolved_binding_specificity"
                ],
            },
        },
        "participant_positive_counts_descriptive": positive_counts,
        "shared_template_boundary": {
            "dataset": "ASME30 artifact-rejected v1",
            "shared_template_supported": shared[
                "mechanism_localization_supported"
            ],
            "gates": shared["gates"],
            "mapping_specificity_mean": shared[
                "mapping_specificity"
            ]["mean"],
            "mapping_specificity_CI": shared[
                "mapping_specificity"
            ]["bootstrap_95_CI_mean"],
            "mapping_specificity_p": shared[
                "mapping_specificity"
            ]["one_sided_exact_signflip_p"],
            "max_t_fwer_min": shared["spatiotemporal"][
                "strongest_absolute_t_cell"
            ]["max_t_fwer_p"],
            "interpretation": (
                "A single cross-participant scalp-time template failed under "
                "a different, artifact-rejected preprocessing route. It is "
                "a boundary result, not a matched statistical comparator to "
                "the dataset-native individualized analysis."
            ),
        },
        "claim": (
            "Across two public hierarchical auditory BCI datasets, all 17 "
            "participants showed positive target-free physical-direction "
            "transport and positive physical-binding specificity; 16/17 "
            "showed an identity advantage. The effects persisted after "
            "training-only command-identity residualization. The supported "
            "object is therefore an individually transported task-native "
            "binding direction, not a mandatory group-shared scalp template."
        ),
        "participant_rows": rows,
        "source_hashes": {
            path.relative_to(ROOT).as_posix(): sha256(path)
            for path in (
                ASME,
                ASME_AUDIT,
                KOJIMA,
                KOJIMA_AUDIT,
                RESIDUAL,
                RESIDUAL_AUDIT,
                SHARED_TEMPLATE,
                SHARED_TEMPLATE_AUDIT,
            )
        },
        "interpretation_boundary": [
            "The 17 participants are described across studies; inferential p values remain dataset specific.",
            "Command residualization disfavors stable command identity but cannot remove every acoustic or task confound.",
            "The shared-template failure and individualized success use different ASME preprocessing and are not a direct paired estimator comparison.",
            "No cortical-source, clinical-benefit, direct-Kakeya-theorem or deployable-oracle claim is made."
        ],
    }
    OUTDIR.mkdir(parents=True, exist_ok=True)
    OUTPUT.write_text(
        json.dumps(output, ensure_ascii=False, indent=2) + "\n",
        encoding="utf-8",
    )
    with CSV.open("x", newline="", encoding="utf-8-sig") as handle:
        writer = csv.DictWriter(handle, fieldnames=list(rows[0]))
        writer.writeheader()
        writer.writerows(rows)
    print(
        json.dumps(
            {
                "output": OUTPUT.relative_to(ROOT).as_posix(),
                "output_sha256": sha256(OUTPUT),
                "csv_sha256": sha256(CSV),
                "status": output["status"],
                "positive_counts": positive_counts,
            },
            ensure_ascii=False,
            indent=2,
        )
    )


if __name__ == "__main__":
    main()
```
