## Supplementary material for "Falsifiable substitution tests reveal task-structured neural evidence for auditory attention": Article_Analysis_Source_Code: discover_kul_wang_cca.py

```
"""Leakage-safe KUL discovery screen of Wang-inspired CCA aggregations.

Each outer fold holds out one of four complete base-part identity groups.
Full presentations, two-minute excerpts, HRTF/dry versions, both attention
assignments, and all repetitions of that part are therefore absent from
training.  Models are participant-specific and all normalization, PCA, CCA,
and projections are fit using the other three identity groups only.

The structural screen compares:
  * conventional dominant-projection CCA;
  * breadth across mutually orthogonal canonical projections;
  * median and lower-quartile projection evidence (non-concentration);
  * separate early/late neural-lag views and their two-ends consensus;
  * agreement between a full window and its two dyadic half-windows.

These are theorem-motivated hypotheses, not deductions from the Kakeya result.
Only odd-ID discovery participants are accepted by this script.
"""

from __future__ import annotations

import argparse
import csv
import json
from collections import defaultdict
from pathlib import Path

import numpy as np
from scipy.linalg import eigh, svd

from audit_kul_discovery import DISCOVERY_SUBJECTS


FS = 32
N_PCS = 16
MAX_COMPONENTS = 8
LAG_SAMPLES = 16
AUDIO_LAG_SAMPLES = 16
EARLY_LAGS = 9
REGULARIZATION = 0.1
TRIM_SECONDS = 5
WINDOW_SECONDS = (1, 2, 5, 10)


def standardize_columns(value: np.ndarray) -> np.ndarray:
    centered = value - value.mean(axis=0, keepdims=True)
    scale = centered.std(axis=0, keepdims=True)
    scale[scale < 1e-8] = 1.0
    return centered / scale


def lag_matrix(value: np.ndarray) -> np.ndarray:
    n = value.shape[0] - LAG_SAMPLES
    return np.concatenate(
        [value[lag : lag + n] for lag in range(LAG_SAMPLES + 1)],
        axis=1,
    )


def audio_lag_matrix(value: np.ndarray) -> np.ndarray:
    n = value.shape[0] - LAG_SAMPLES
    return np.concatenate(
        [
            value[lag : lag + n]
            for lag in range(AUDIO_LAG_SAMPLES + 1)
        ],
        axis=1,
    )


def pca_basis(eeg_trials: list[np.ndarray]) -> tuple[np.ndarray, np.ndarray]:
    covariance = np.zeros((eeg_trials[0].shape[1],) * 2)
    count = 0
    trim = TRIM_SECONDS * FS
    for value in eeg_trials:
        normalized = standardize_columns(value)
        segment = normalized[trim:-trim]
        covariance += segment.T @ segment
        count += segment.shape[0]
    covariance /= count
    eigenvalues, eigenvectors = eigh(covariance)
    order = np.argsort(eigenvalues)[::-1][:N_PCS]
    scale = np.sqrt(np.maximum(eigenvalues[order], 1e-8))
    return eigenvectors[:, order], scale


def inverse_square_root(value: np.ndarray) -> np.ndarray:
    scale = np.trace(value) / value.shape[0]
    eigenvalues, eigenvectors = eigh(
        value + REGULARIZATION * scale * np.eye(value.shape[0])
    )
    eigenvalues = np.maximum(eigenvalues, 1e-10)
    return (eigenvectors / np.sqrt(eigenvalues)) @ eigenvectors.T


def fit_cca(
    cxx: np.ndarray, cyy: np.ndarray, cxy: np.ndarray
) -> tuple[np.ndarray, np.ndarray, np.ndarray]:
    wx = inverse_square_root(cxx)
    wy = inverse_square_root(cyy)
    left, canonical, right_t = svd(
        wx @ cxy @ wy, full_matrices=False, check_finite=False
    )
    return wx @ left, wy @ right_t.T, canonical


def correlation_columns(first: np.ndarray, second: np.ndarray) -> np.ndarray:
    a = first - first.mean(axis=0, keepdims=True)
    b = second - second.mean(axis=0, keepdims=True)
    denominator = np.linalg.norm(a, axis=0) * np.linalg.norm(b, axis=0)
    return np.divide(
        np.sum(a * b, axis=0),
        denominator,
        out=np.zeros(a.shape[1]),
        where=denominator > 1e-12,
    )


def load_subject(
    root: Path, subject: int, audio: dict[str, np.ndarray]
) -> tuple[list[np.ndarray], list[dict[str, object]], list[np.ndarray]]:
    archive = np.load(root / f"S{subject}.npz")
    metadata = json.loads(str(archive["metadata_json"]))
    eeg = [archive[f"eeg_{index:02d}"] for index in range(1, 21)]
    attended = [
        audio[str(row["attended_audio"])][: int(row["samples"])]
        for row in metadata
    ]
    return eeg, metadata, attended


def sufficient_statistics(
    eeg: list[np.ndarray],
    attended: list[np.ndarray],
    training_indices: list[int],
    basis: np.ndarray,
    pca_scale: np.ndarray,
) -> dict[str, tuple[np.ndarray, np.ndarray, np.ndarray]]:
    full_dim = N_PCS * (LAG_SAMPLES + 1)
    early_dim = N_PCS * EARLY_LAGS
    late_dim = full_dim - early_dim
    audio_dim = attended[training_indices[0]].shape[1] * (
        AUDIO_LAG_SAMPLES + 1
    )
    cxx = {
        "full": np.zeros((full_dim, full_dim)),
        "early": np.zeros((early_dim, early_dim)),
        "late": np.zeros((late_dim, late_dim)),
    }
    cxy = {
        "full": np.zeros((full_dim, audio_dim)),
        "early": np.zeros((early_dim, audio_dim)),
        "late": np.zeros((late_dim, audio_dim)),
    }
    cyy = np.zeros((audio_dim, audio_dim))
    count = 0
    trim = TRIM_SECONDS * FS
    for index in training_indices:
        projected = (
            standardize_columns(eeg[index]) @ basis
        ) / pca_scale
        full = lag_matrix(projected)
        views = {
            "full": full,
            "early": full[:, :early_dim],
            "late": full[:, early_dim:],
        }
        y = audio_lag_matrix(standardize_columns(attended[index]))
        usable = min(full.shape[0], y.shape[0])
        selection = slice(trim, usable - trim)
        ys = y[selection]
        cyy += ys.T @ ys
        for name, x in views.items():
            xs = x[selection]
            cxx[name] += xs.T @ xs
            cxy[name] += xs.T @ ys
        count += ys.shape[0]
    output = {}
    for name in ("full", "early", "late"):
        output[name] = (
            cxx[name] / count,
            cyy / count,
            cxy[name] / count,
        )
    return output


def weighted_mean(value: np.ndarray, weights: np.ndarray) -> float:
    weights = weights / weights.sum()
    return float(np.dot(value, weights))


def candidate_scores(value: np.ndarray, prefix: str) -> dict[str, float]:
    positive = np.maximum(value, 0)
    negative = np.maximum(-value, 0)
    ordered = np.sort(value)
    return {
        f"{prefix}_mean": float(np.mean(value)),
        f"{prefix}_median": float(np.median(value)),
        f"{prefix}_q25": float(np.quantile(value, 0.25)),
        f"{prefix}_trim_max": float(np.mean(ordered[:-1])),
        f"{prefix}_lcb25": float(np.mean(value) - 0.25 * np.std(value)),
        f"{prefix}_lcb50": float(np.mean(value) - 0.50 * np.std(value)),
        f"{prefix}_positive_l2": float(np.sqrt(np.mean(positive**2))),
        f"{prefix}_signed_square": float(
            np.sqrt(np.mean(positive**2))
            - np.sqrt(np.mean(negative**2))
        ),
    }


def fixed_method_margins(
    full_attended: np.ndarray,
    full_unattended: np.ndarray,
    early_attended: np.ndarray,
    early_unattended: np.ndarray,
    late_attended: np.ndarray,
    late_unattended: np.ndarray,
    full_canonical: np.ndarray,
    nested: dict[str, object],
) -> dict[str, float]:
    full = full_attended - full_unattended
    early = early_attended - early_unattended
    late = late_attended - late_unattended
    canonical4 = full_canonical[:4]
    canonical8 = full_canonical[:8]
    capped8 = np.minimum(canonical8, np.median(canonical8))
    early1, late1 = float(early[0]), float(late[0])
    combined4 = np.concatenate([early[:4], late[:4]])
    methods = {
        "full_k1": float(full[0]),
        "full_equal_k2": float(np.mean(full[:2])),
        "full_equal_k4": float(np.mean(full[:4])),
        "full_equal_k8": float(np.mean(full[:8])),
        "full_canonical_k4": weighted_mean(full[:4], canonical4),
        "full_canonical_k8": weighted_mean(full[:8], canonical8),
        "full_capped_k8": weighted_mean(full[:8], capped8),
        "full_median_k4": float(np.median(full[:4])),
        "full_median_k8": float(np.median(full[:8])),
        "full_q25_k4": float(np.quantile(full[:4], 0.25)),
        "full_q25_k8": float(np.quantile(full[:8], 0.25)),
        "early_k1": early1,
        "late_k1": late1,
        "two_ends_k1_mean": (early1 + late1) / 2,
        "two_ends_k1_min": min(early1, late1),
        "two_ends_k1_penalty25": (
            (early1 + late1) / 2 - 0.25 * abs(early1 - late1)
        ),
        "two_ends_k4_mean": float(np.mean(combined4)),
        "two_ends_k4_median": float(np.median(combined4)),
        "two_ends_k4_q25": float(np.quantile(combined4, 0.25)),
    }
    for component in range(MAX_COMPONENTS):
        methods[f"full_component_{component + 1}"] = float(full[component])
    for components in (4, 8):
        attended_scores = candidate_scores(
            full_attended[:components], f"candidate_k{components}"
        )
        unattended_scores = candidate_scores(
            full_unattended[:components], f"candidate_k{components}"
        )
        for name in attended_scores:
            methods[name] = attended_scores[name] - unattended_scores[name]
    two_ends_attended = np.concatenate(
        [early_attended[:4], late_attended[:4]]
    )
    two_ends_unattended = np.concatenate(
        [early_unattended[:4], late_unattended[:4]]
    )
    attended_scores = candidate_scores(
        two_ends_attended, "candidate_two_ends_k4"
    )
    unattended_scores = candidate_scores(
        two_ends_unattended, "candidate_two_ends_k4"
    )
    for name in attended_scores:
        methods[name] = attended_scores[name] - unattended_scores[name]
    selected_k = int(nested["selected_k"])
    reliability = np.asarray(nested["reliability_weights"], dtype=float)
    shrunk = np.asarray(nested["shrunk_weights"], dtype=float)
    methods["nested_selected_equal"] = float(
        np.mean(full[:selected_k])
    )
    methods["nested_selected_canonical"] = weighted_mean(
        full[:selected_k], full_canonical[:selected_k]
    )
    methods["nested_reliability_capped"] = float(
        np.dot(full[:selected_k], reliability[:selected_k])
    )
    methods["nested_reliability_shrunk"] = float(
        np.dot(full[:selected_k], shrunk[:selected_k])
    )
    return methods


def window_component_correlations(
    x_projection: np.ndarray,
    attended_projection: np.ndarray,
    unattended_projection: np.ndarray,
    start: int,
    stop: int,
    components: int,
) -> tuple[np.ndarray, np.ndarray]:
    x = x_projection[start:stop, :components]
    return (
        correlation_columns(
            x, attended_projection[start:stop, :components]
        ),
        correlation_columns(
            x, unattended_projection[start:stop, :components]
        ),
    )


def evaluate_trial(
    eeg: np.ndarray,
    attended_audio: np.ndarray,
    unattended_audio: np.ndarray,
    basis: np.ndarray,
    pca_scale: np.ndarray,
    models: dict[str, tuple[np.ndarray, np.ndarray, np.ndarray]],
    nested: dict[str, object],
) -> dict[int, dict[str, list[float]]]:
    projected = (standardize_columns(eeg) @ basis) / pca_scale
    full_x = lag_matrix(projected)
    early_dim = N_PCS * EARLY_LAGS
    x_views = {
        "full": full_x,
        "early": full_x[:, :early_dim],
        "late": full_x[:, early_dim:],
    }
    attended_y = audio_lag_matrix(standardize_columns(attended_audio))
    unattended_y = audio_lag_matrix(standardize_columns(unattended_audio))
    usable = min(
        full_x.shape[0], attended_y.shape[0], unattended_y.shape[0]
    )
    projections = {}
    for name, (a, b, canonical) in models.items():
        projections[name] = (
            x_views[name][:usable] @ a[:, :MAX_COMPONENTS],
            attended_y[:usable] @ b[:, :MAX_COMPONENTS],
            unattended_y[:usable] @ b[:, :MAX_COMPONENTS],
            canonical,
        )

    result: dict[int, dict[str, list[float]]] = {}
    trim = TRIM_SECONDS * FS
    for seconds in WINDOW_SECONDS:
        window = seconds * FS
        values: dict[str, list[float]] = defaultdict(list)
        for start in range(trim, usable - trim - window + 1, window):
            stop = start + window
            correlations = {}
            for name in ("full", "early", "late"):
                x, ya, yu, _ = projections[name]
                correlations[name] = window_component_correlations(
                    x, ya, yu, start, stop, MAX_COMPONENTS
                )
            methods = fixed_method_margins(
                correlations["full"][0],
                correlations["full"][1],
                correlations["early"][0],
                correlations["early"][1],
                correlations["late"][0],
                correlations["late"][1],
                projections["full"][3],
                nested,
            )
            if seconds >= 2:
                midpoint = start + window // 2
                half_margins = []
                for half_start, half_stop in (
                    (start, midpoint),
                    (midpoint, stop),
                ):
                    x, ya, yu, _ = projections["full"]
                    half_attended, half_unattended = (
                        window_component_correlations(
                            x,
                            ya,
                            yu,
                            half_start,
                            half_stop,
                            4,
                        )
                    )
                    half_margins.append(
                        np.mean(half_attended - half_unattended)
                    )
                full_scale = methods["full_equal_k4"]
                half_scale = float(np.mean(half_margins))
                methods["dyadic_k4_mean"] = (
                    full_scale + half_scale
                ) / 2
                methods["dyadic_k4_min"] = min(full_scale, half_scale)
            for method, margin in methods.items():
                values[method].append(float(margin))
        result[seconds] = values
    return result


def inner_trial_component_margins(
    eeg: np.ndarray,
    attended_audio: np.ndarray,
    unattended_audio: np.ndarray,
    basis: np.ndarray,
    pca_scale: np.ndarray,
    model: tuple[np.ndarray, np.ndarray, np.ndarray],
) -> np.ndarray:
    projected = (standardize_columns(eeg) @ basis) / pca_scale
    x = lag_matrix(projected)
    attended_y = audio_lag_matrix(standardize_columns(attended_audio))
    unattended_y = audio_lag_matrix(standardize_columns(unattended_audio))
    usable = min(x.shape[0], attended_y.shape[0], unattended_y.shape[0])
    a, b, _ = model
    xp = x[:usable] @ a[:, :MAX_COMPONENTS]
    ya = attended_y[:usable] @ b[:, :MAX_COMPONENTS]
    yu = unattended_y[:usable] @ b[:, :MAX_COMPONENTS]
    trim = TRIM_SECONDS * FS
    window = 5 * FS
    values = []
    for start in range(trim, usable - trim - window + 1, window):
        attended_r, unattended_r = window_component_correlations(
            xp, ya, yu, start, start + window, MAX_COMPONENTS
        )
        values.append(attended_r - unattended_r)
    return np.asarray(values)


def nested_direction_rule(
    eeg: list[np.ndarray],
    metadata: list[dict[str, object]],
    attended: list[np.ndarray],
    audio: dict[str, np.ndarray],
    outer_held_out_part: int,
) -> tuple[dict[str, object], dict[str, object]]:
    available_parts = [
        part for part in (1, 2, 3, 4) if part != outer_held_out_part
    ]
    trial_values = []
    for validation_part in available_parts:
        training_indices = [
            index
            for index, row in enumerate(metadata)
            if int(row["base_part_identity_group"])
            not in (outer_held_out_part, validation_part)
        ]
        validation_indices = [
            index
            for index, row in enumerate(metadata)
            if int(row["base_part_identity_group"]) == validation_part
        ]
        basis, pca_scale = pca_basis(
            [eeg[index] for index in training_indices]
        )
        statistics = sufficient_statistics(
            eeg, attended, training_indices, basis, pca_scale
        )
        model = fit_cca(*statistics["full"])
        for index in validation_indices:
            row = metadata[index]
            unattended_audio = audio[str(row["unattended_audio"])][
                : int(row["samples"])
            ]
            trial_values.append(
                inner_trial_component_margins(
                    eeg[index],
                    attended[index],
                    unattended_audio,
                    basis,
                    pca_scale,
                    model,
                )
            )
    component_accuracy = np.mean(
        [np.mean(value > 0, axis=0) for value in trial_values], axis=0
    )
    candidate_k = (1, 2, 4, 8)
    ensemble_accuracy = {
        components: float(
            np.mean(
                [
                    np.mean(np.mean(value[:, :components], axis=1) > 0)
                    for value in trial_values
                ]
            )
        )
        for components in candidate_k
    }
    selected_k = min(
        candidate_k,
        key=lambda components: (
            -ensemble_accuracy[components],
            components,
        ),
    )
    excess = np.maximum(component_accuracy[:selected_k] - 0.5, 0)
    if not np.any(excess > 0):
        reliability = np.ones(selected_k) / selected_k
    else:
        positive = excess[excess > 0]
        capped = np.minimum(excess, np.median(positive))
        reliability = capped / capped.sum()
    shrunk = 0.5 * reliability + 0.5 / selected_k
    padded_reliability = np.zeros(MAX_COMPONENTS)
    padded_shrunk = np.zeros(MAX_COMPONENTS)
    padded_reliability[:selected_k] = reliability
    padded_shrunk[:selected_k] = shrunk
    rule = {
        "selected_k": selected_k,
        "reliability_weights": padded_reliability.tolist(),
        "shrunk_weights": padded_shrunk.tolist(),
    }
    audit = {
        "outer_held_out_part": outer_held_out_part,
        "inner_validation_parts": available_parts,
        "component_accuracy": component_accuracy.tolist(),
        "ensemble_accuracy": {
            str(key): value for key, value in ensemble_accuracy.items()
        },
        **rule,
    }
    return rule, audit


def summarize(
    trial_rows: list[dict[str, object]]
) -> tuple[list[dict[str, object]], dict[str, object]]:
    grouped = defaultdict(list)
    for row in trial_rows:
        grouped[
            (
                int(row["subject"]),
                str(row["method"]),
                int(row["window_seconds"]),
            )
        ].append(row)
    subject_rows = []
    for (subject, method, seconds), rows in sorted(grouped.items()):
        subject_rows.append(
            {
                "subject": subject,
                "method": method,
                "window_seconds": seconds,
                "accuracy": float(
                    np.mean([float(row["accuracy"]) for row in rows])
                ),
                "mean_margin": float(
                    np.mean([float(row["mean_margin"]) for row in rows])
                ),
                "trials": len(rows),
            }
        )
    methods = defaultdict(list)
    for row in subject_rows:
        methods[
            (str(row["method"]), int(row["window_seconds"]))
        ].append(row)
    payload = {}
    for (method, seconds), rows in sorted(methods.items()):
        accuracies = np.asarray([float(row["accuracy"]) for row in rows])
        baseline = {
            int(row["subject"]): float(row["accuracy"])
            for row in subject_rows
            if row["method"] == "full_k1"
            and int(row["window_seconds"]) == seconds
        }
        gains = np.asarray(
            [
                float(row["accuracy"]) - baseline[int(row["subject"])]
                for row in rows
            ]
        )
        payload[f"{method}|w={seconds}"] = {
            "method": method,
            "window_seconds": seconds,
            "participant_mean_accuracy": float(accuracies.mean()),
            "participant_standard_error": float(
                accuracies.std(ddof=1) / np.sqrt(accuracies.size)
            ),
            "participant_mean_gain_vs_full_k1": float(gains.mean()),
            "participants_improved_vs_full_k1": int(np.sum(gains > 0)),
            "participants_tied_vs_full_k1": int(np.sum(gains == 0)),
            "participant_accuracies": accuracies.tolist(),
            "participant_gains_vs_full_k1": gains.tolist(),
        }
    return subject_rows, payload


def write_csv(path: Path, rows: list[dict[str, object]]) -> None:
    with path.open("w", encoding="utf-8", newline="") as handle:
        writer = csv.DictWriter(handle, fieldnames=list(rows[0]))
        writer.writeheader()
        writer.writerows(rows)


def main() -> None:
    global AUDIO_LAG_SAMPLES
    parser = argparse.ArgumentParser()
    parser.add_argument(
        "--input-dir",
        type=Path,
        default=Path("data/derived/kul_discovery_32hz"),
    )
    parser.add_argument(
        "--output-dir",
        type=Path,
        default=Path("results/kul_aad/discovery_wang_cca"),
    )
    parser.add_argument(
        "--audio-lag-samples",
        type=int,
        choices=(0, 4, 8, 16),
        default=16,
    )
    parser.add_argument("--skip-nested", action="store_true")
    parser.add_argument(
        "--full-only",
        action="store_true",
        help="Exclude all experiment-3 repeated two-minute trials.",
    )
    args = parser.parse_args()
    AUDIO_LAG_SAMPLES = args.audio_lag_samples
    manifest = json.loads(
        (args.input_dir / "manifest.json").read_text(encoding="utf-8")
    )
    observed = tuple(int(value) for value in manifest["participants"])
    if observed != DISCOVERY_SUBJECTS:
        raise RuntimeError(
            f"KUL confirmation firewall violation: observed={observed}"
        )
    if manifest["confirmation_participants_loaded"]:
        raise RuntimeError("Confirmation participants present in manifest")
    audio_archive = np.load(
        args.input_dir / str(manifest["audio_cache"])
    )
    audio = {
        name.removeprefix("audio::"): audio_archive[name]
        for name in audio_archive.files
        if name.startswith("audio::")
    }

    trial_rows = []
    canonical_rows = []
    nested_rows = []
    for subject in DISCOVERY_SUBJECTS:
        eeg, metadata, attended = load_subject(
            args.input_dir, subject, audio
        )
        for held_out_part in (1, 2, 3, 4):
            if args.skip_nested:
                nested_rule = {
                    "selected_k": 4,
                    "reliability_weights": [
                        0.25,
                        0.25,
                        0.25,
                        0.25,
                        0,
                        0,
                        0,
                        0,
                    ],
                    "shrunk_weights": [
                        0.25,
                        0.25,
                        0.25,
                        0.25,
                        0,
                        0,
                        0,
                        0,
                    ],
                }
                nested_audit = {
                    "outer_held_out_part": held_out_part,
                    "status": "skipped_fixed_four_direction_placeholder",
                    **nested_rule,
                }
            else:
                nested_rule, nested_audit = nested_direction_rule(
                    eeg,
                    metadata,
                    attended,
                    audio,
                    held_out_part,
                )
            nested_rows.append(
                {
                    "subject": subject,
                    **nested_audit,
                }
            )
            training_indices = [
                index
                for index, row in enumerate(metadata)
                if int(row["base_part_identity_group"]) != held_out_part
                and (not args.full_only or not bool(row["repetition"]))
            ]
            testing_indices = [
                index
                for index, row in enumerate(metadata)
                if int(row["base_part_identity_group"]) == held_out_part
                and (not args.full_only or not bool(row["repetition"]))
            ]
            expected = (6, 2) if args.full_only else (15, 5)
            if (len(training_indices), len(testing_indices)) != expected:
                raise RuntimeError("Identity-fold cardinality failure")
            basis, pca_scale = pca_basis(
                [eeg[index] for index in training_indices]
            )
            statistics = sufficient_statistics(
                eeg,
                attended,
                training_indices,
                basis,
                pca_scale,
            )
            models = {
                name: fit_cca(*values)
                for name, values in statistics.items()
            }
            for name, (_, _, canonical) in models.items():
                canonical_rows.append(
                    {
                        "subject": subject,
                        "held_out_part": held_out_part,
                        "view": name,
                        **{
                            f"canonical_{index + 1}": float(canonical[index])
                            for index in range(MAX_COMPONENTS)
                        },
                    }
                )
            for index in testing_indices:
                row = metadata[index]
                unattended_audio = audio[str(row["unattended_audio"])][
                    : int(row["samples"])
                ]
                evaluated = evaluate_trial(
                    eeg[index],
                    attended[index],
                    unattended_audio,
                    basis,
                    pca_scale,
                    models,
                    nested_rule,
                )
                for seconds, method_values in evaluated.items():
                    for method, margins in method_values.items():
                        values = np.asarray(margins)
                        trial_rows.append(
                            {
                                "subject": subject,
                                "held_out_part": held_out_part,
                                "trial": int(row["trial"]),
                                "condition": str(row["condition"]),
                                "repetition": int(bool(row["repetition"])),
                                "attended_ear": str(row["attended_ear"]),
                                "method": method,
                                "window_seconds": seconds,
                                "windows": int(values.size),
                                "accuracy": float(np.mean(values > 0)),
                                "mean_margin": float(values.mean()),
                            }
                        )
            print(
                f"S{subject:02d}, held-out identity part "
                f"{held_out_part}: complete",
                flush=True,
            )
        del eeg, attended

    subject_rows, model_summary = summarize(trial_rows)
    args.output_dir.mkdir(parents=True, exist_ok=True)
    write_csv(args.output_dir / "trial_results.csv", trial_rows)
    write_csv(args.output_dir / "subject_results.csv", subject_rows)
    write_csv(args.output_dir / "canonical_spectra.csv", canonical_rows)
    (args.output_dir / "nested_direction_selection.json").write_text(
        json.dumps(nested_rows, indent=2), encoding="utf-8"
    )
    summary = {
        "design": {
            "partition": "odd-participant discovery only",
            "participants": list(DISCOVERY_SUBJECTS),
            "cross_validation": (
                "within-participant leave-one-base-part-identity-out"
            ),
            "training_trials_per_fold": 15,
            "testing_trials_per_fold": 5,
            "normalization_pca_cca_fit_on_training_only": True,
            "participant_is_inferential_unit": True,
            "trial_equal_weighting_within_participant": True,
            "repeated_trials_included": not args.full_only,
            "sample_rate_hz": FS,
            "pca_components": N_PCS,
            "lag_range_ms": [0, 500],
            "audio_lag_range_ms": [
                0,
                1000 * AUDIO_LAG_SAMPLES / FS,
            ],
            "regularization": REGULARIZATION,
            "window_seconds": list(WINDOW_SECONDS),
        },
        "models": model_summary,
    }
    (args.output_dir / "summary.json").write_text(
        json.dumps(summary, indent=2), encoding="utf-8"
    )
    ordered = sorted(
        model_summary.values(),
        key=lambda value: value["participant_mean_accuracy"],
        reverse=True,
    )
    for result in ordered[:30]:
        print(
            f"{result['method']:28s} w={result['window_seconds']:2d} "
            f"acc={result['participant_mean_accuracy']:.4f} "
            f"gain={result['participant_mean_gain_vs_full_k1']:+.4f} "
            f"improved={result['participants_improved_vs_full_k1']}/8"
        )


if __name__ == "__main__":
    main()
```
