## Supplementary material for "Falsifiable substitution tests reveal task-structured neural evidence for auditory attention": Article_Analysis_Source_Code: download_asme30_signal_v1.py

```
#!/usr/bin/env python3
"""Download only the 90 offline/online EDFs required by the frozen protocol."""

from __future__ import annotations

import concurrent.futures
import hashlib
import json
import os
from pathlib import Path

import requests


ROOT = Path(__file__).resolve().parents[1]
OUT = ROOT / "data" / "raw" / "asme30_signal"
MANIFEST = ROOT / "data" / "inventory" / "asme30_signal_manifest_v1.json"
PROTOCOL = ROOT / "docs" / "wz_hic_asme30_external_protocol_v1.json"
LOCK = ROOT / "docs" / "wz_hic_asme30_external_implementation_lock_v1.json"
API = "https://dataverse.harvard.edu/api"
DOI = "doi:10.7910/DVN/TYRCWL"


def digest(path: Path, algorithm: str) -> str:
    h = hashlib.new(algorithm)
    with path.open("rb") as f:
        for block in iter(lambda: f.read(4 * 1024 * 1024), b""):
            h.update(block)
    return h.hexdigest()


def main() -> None:
    if not PROTOCOL.exists() or not LOCK.exists():
        raise FileNotFoundError("Freeze protocol and implementation before EDF access.")
    response = requests.get(
        f"{API}/datasets/:persistentId/",
        params={"persistentId": DOI},
        timeout=(30, 120),
    )
    response.raise_for_status()
    version = response.json()["data"]["latestVersion"]
    selected = [
        item
        for item in version["files"]
        if item["label"].endswith("_eeg.edf")
        and (
            "task-offline" in item["label"]
            or "task-online" in item["label"]
        )
    ]
    selected.sort(key=lambda x: (x.get("directoryLabel", ""), x["label"]))
    if len(selected) != 90:
        raise RuntimeError(f"Expected 90 EDFs, found {len(selected)}")
    if any("derivatives" in item.get("directoryLabel", "") for item in selected):
        raise RuntimeError("Derivative entered signal selection.")

    def download(item: dict) -> dict:
        data_file = item["dataFile"]
        target = OUT / item.get("directoryLabel", "") / item["label"]
        target.parent.mkdir(parents=True, exist_ok=True)
        expected_size = int(data_file["filesize"])
        expected_md5 = data_file["checksum"]["value"].lower()
        if target.exists() and target.stat().st_size == expected_size:
            observed_md5 = digest(target, "md5")
            if observed_md5 == expected_md5:
                return {
                    "directory": item.get("directoryLabel", ""),
                    "label": item["label"],
                    "dataverse_file_id": data_file["id"],
                    "bytes": expected_size,
                    "md5": observed_md5,
                    "sha256": digest(target, "sha256"),
                    "path": target.relative_to(ROOT).as_posix(),
                    "download_status": "reused_verified",
                }
        partial = target.with_suffix(target.suffix + ".partial")
        if partial.exists():
            partial.unlink()
        with requests.get(
            f"{API}/access/datafile/{data_file['id']}",
            stream=True,
            timeout=(30, 900),
        ) as r:
            r.raise_for_status()
            with partial.open("wb") as f:
                for chunk in r.iter_content(chunk_size=4 * 1024 * 1024):
                    if chunk:
                        f.write(chunk)
                f.flush()
                os.fsync(f.fileno())
        if partial.stat().st_size != expected_size:
            raise RuntimeError(
                f"Size mismatch {item['label']}: "
                f"{partial.stat().st_size} != {expected_size}"
            )
        observed_md5 = digest(partial, "md5")
        if observed_md5 != expected_md5:
            raise RuntimeError(
                f"MD5 mismatch {item['label']}: {observed_md5} != {expected_md5}"
            )
        partial.replace(target)
        record = {
            "directory": item.get("directoryLabel", ""),
            "label": item["label"],
            "dataverse_file_id": data_file["id"],
            "bytes": expected_size,
            "md5": observed_md5,
            "sha256": digest(target, "sha256"),
            "path": target.relative_to(ROOT).as_posix(),
            "download_status": "downloaded_verified",
        }
        print(f"{item['label']} {expected_size}", flush=True)
        return record

    records = []
    with concurrent.futures.ThreadPoolExecutor(max_workers=3) as pool:
        for record in pool.map(download, selected):
            records.append(record)
    records.sort(key=lambda x: x["path"])
    manifest = {
        "status": "PASS",
        "dataset": "ASME-speller 30-class",
        "persistent_id": DOI,
        "version": f"{version['versionNumber']}.{version['versionMinorNumber']}",
        "license": version["license"],
        "scope": (
            "Only six offline and three online EEG EDF files for each of ten "
            "participants; no derivative classifier outputs."
        ),
        "n_edf_files": len(records),
        "total_bytes": sum(item["bytes"] for item in records),
        "all_md5_verified": all(item["md5"] for item in records),
        "derivative_files_downloaded": 0,
        "protocol_sha256": digest(PROTOCOL, "sha256"),
        "implementation_lock_sha256": digest(LOCK, "sha256"),
        "files": records,
    }
    MANIFEST.parent.mkdir(parents=True, exist_ok=True)
    MANIFEST.write_text(json.dumps(manifest, indent=2) + "\n", encoding="utf-8")
    print(f"manifest={MANIFEST}")
    print(f"sha256={digest(MANIFEST, 'sha256')}")
    print(f"files={len(records)} bytes={manifest['total_bytes']}")


if __name__ == "__main__":
    main()
```
