## Supplementary material for "Falsifiable substitution tests reveal task-structured neural evidence for auditory attention": Article_Analysis_Source_Code: download_asme30_structure_metadata_v1.py

```
"""Download outcome-blind BIDS structure for the 2026 30-class ASME-speller."""

from __future__ import annotations

import concurrent.futures
import hashlib
import json
from pathlib import Path

import requests


ROOT = Path(__file__).resolve().parents[1]
OUT = ROOT / "data" / "raw" / "asme30_structure"
MANIFEST = ROOT / "data" / "inventory" / "asme30_structure_manifest_v1.json"
API = "https://dataverse.harvard.edu/api"
DOI = "doi:10.7910/DVN/TYRCWL"


def digest(path: Path, algorithm: str) -> str:
    h = hashlib.new(algorithm)
    h.update(path.read_bytes())
    return h.hexdigest()


def main() -> None:
    j = requests.get(
        f"{API}/datasets/:persistentId/",
        params={"persistentId": DOI},
        timeout=60,
    ).json()["data"]["latestVersion"]
    selected = []
    for item in j["files"]:
        label = item["label"]
        original = item["dataFile"].get("originalFileName", "")
        if label in {"README.md", "dataset_description.json", "participants.json"}:
            selected.append(item)
            continue
        task = "task-offline" in label or "task-online" in label
        structure = (
            label.endswith("_events.json")
            or label.endswith("_eeg.json")
            or original.endswith("_events.tsv")
            or original.endswith("_channels.tsv")
        )
        if task and structure:
            selected.append(item)
    if any(x["label"].endswith(".edf") for x in selected):
        raise RuntimeError("Signal file entered structure selection.")
    if len(selected) != 3 + 10 * 9 * 4:
        raise RuntimeError(f"Unexpected structure file count: {len(selected)}")

    def download(item: dict) -> dict:
        d = item["dataFile"]
        original = d.get("originalFileName")
        label = original or item["label"]
        target = OUT / item.get("directoryLabel", "") / label
        target.parent.mkdir(parents=True, exist_ok=True)
        params = {"format": "original"} if original else {}
        r = requests.get(
            f"{API}/access/datafile/{d['id']}", params=params, timeout=60
        )
        r.raise_for_status()
        target.write_bytes(r.content)
        md5 = digest(target, "md5")
        official = d["checksum"]["value"].lower()
        # Dataverse tabular checksum can refer to the preserved original or the
        # normalized tabular representation depending on ingest. Size+local hash
        # are always recorded; a mismatch is explicit rather than silently hidden.
        return {
            "directory": item.get("directoryLabel", ""),
            "label": item["label"],
            "original_file_name": original,
            "dataverse_file_id": d["id"],
            "official_bytes": d["filesize"],
            "downloaded_bytes": target.stat().st_size,
            "official_md5": official,
            "local_md5": md5,
            "official_md5_match": md5 == official,
            "local_sha256": digest(target, "sha256"),
            "path": str(target.relative_to(ROOT)).replace("\\", "/"),
        }

    records = []
    with concurrent.futures.ThreadPoolExecutor(max_workers=10) as pool:
        for record in pool.map(download, selected):
            records.append(record)
    records.sort(key=lambda x: x["path"])
    manifest = {
        "status": "PASS",
        "dataset": "ASME-speller 30-class",
        "persistent_id": DOI,
        "version": f"{j['versionNumber']}.{j['versionMinorNumber']}",
        "license": j["license"],
        "scope": (
            "README, dataset/participant JSON, and offline/online BIDS "
            "events/channels/EEG sidecars only; no EDF, stimuli, derivatives, or "
            "online classification result values"
        ),
        "n_files": len(records),
        "total_downloaded_bytes": sum(x["downloaded_bytes"] for x in records),
        "EDF_files_downloaded": 0,
        "derivative_result_files_downloaded": 0,
        "files": records,
    }
    MANIFEST.parent.mkdir(parents=True, exist_ok=True)
    MANIFEST.write_text(
        json.dumps(manifest, ensure_ascii=False, indent=2) + "\n", encoding="utf-8"
    )
    print(MANIFEST)
    print(digest(MANIFEST, "sha256"))
    print(len(records), manifest["total_downloaded_bytes"])


if __name__ == "__main__":
    main()
```
