## Supplementary material for "Falsifiable substitution tests reveal task-structured neural evidence for auditory attention": Article_Analysis_Source_Code: download_kite_st_hd_confirmation_v1.py

```
"""Download exactly the 14 frozen KITE-ST-HD confirmation archives."""

from __future__ import annotations

import argparse
import hashlib
import json
import os
import time
import urllib.request
from pathlib import Path

from kite_sat_confirmation_lock_utils_v1 import ROOT, validate_confirmation_lock


SUBJECTS = (
    "S0", "S10", "S26", "S2", "S21", "S28", "S22",
    "S9", "S3", "S15", "S16", "S18", "S13", "S1",
)
FILES = {
    "S0": (1595382432, "878639e9403ae50bf49e586e6fd2e247"),
    "S10": (1537620418, "992e460a5a5681fd295f2f54cd45bb4f"),
    "S26": (1570883148, "f8c1eef8e4cd744694799de7d9dbae43"),
    "S2": (1594184702, "9f8632f0308472a22ddcff38a5d29ea5"),
    "S21": (1549811083, "5af735a0c6de6b8ba38f9668e17a87f0"),
    "S28": (1636182867, "4e583685ed7b7c8d76d0bba9a8f2c51c"),
    "S22": (1661550873, "c24ac9b4e43746260dccb50638b22648"),
    "S9": (1584883076, "2456d15a55f4fbfe83665312027f8ae0"),
    "S3": (1613014619, "80da47754e91f581b987c9a152e1d824"),
    "S15": (1682151858, "c067faa8327a27d099f75f0d31b46168"),
    "S16": (1635646725, "a23f11c39e942ae2c99e579cd4c2d5bc"),
    "S18": (1660394720, "9f639b5e11989224736f5b0fc817638a"),
    "S13": (1618698966, "616a5b5a455f34e5e1d7347f8e65bb8a"),
    "S1": (1586358973, "4142b7c3f2e6191e00aaf23490133886"),
}
OUTPUT_ROOT = (
    ROOT / "data" / "raw" / "kite_st_hd_zenodo4518754" / "confirmation_archives"
)
METADATA_ROOT = (
    ROOT / "data" / "metadata" / "kite_st_hd_confirmation_downloads_v1"
)
SUMMARY = (
    ROOT / "results" / "kite_st_hd" / "confirmation_v1"
    / "confirmation_acquisition_summary_v1.json"
)
BASE = "https://zenodo.org/api/records/4518754/files"


def file_hash(path: Path, algorithm: str) -> str:
    digest = hashlib.new(algorithm)
    with path.open("rb") as stream:
        for chunk in iter(lambda: stream.read(8 * 1024 * 1024), b""):
            digest.update(chunk)
    return digest.hexdigest()


def retrieve(subject: str, lock_hash: str) -> dict:
    expected_bytes, expected_md5 = FILES[subject]
    OUTPUT_ROOT.mkdir(parents=True, exist_ok=True)
    METADATA_ROOT.mkdir(parents=True, exist_ok=True)
    final = OUTPUT_ROOT / f"{subject}.tar.gz"
    partial = final.with_name(final.name + ".part")
    if final.exists():
        if final.stat().st_size != expected_bytes:
            raise RuntimeError(f"Existing confirmation archive size mismatch: {final}")
        if file_hash(final, "md5") != expected_md5:
            raise RuntimeError(f"Existing confirmation archive MD5 mismatch: {final}")
        status = "existing_archive_verified"
    else:
        offset = partial.stat().st_size if partial.exists() else 0
        if offset > expected_bytes:
            raise RuntimeError(f"Partial archive exceeds expected size: {partial}")
        request = urllib.request.Request(
            f"{BASE}/{subject}.tar.gz/content",
            headers={
                "User-Agent": "KITE-SAT-independent-confirmation/1.0",
                "Range": f"bytes={offset}-",
            },
        )
        started = time.time()
        with urllib.request.urlopen(request, timeout=120) as response:
            code = getattr(response, "status", response.getcode())
            if offset and code != 206:
                offset = 0
                mode = "wb"
            else:
                mode = "ab" if offset else "wb"
            with partial.open(mode) as stream:
                while True:
                    chunk = response.read(8 * 1024 * 1024)
                    if not chunk:
                        break
                    stream.write(chunk)
                    offset += len(chunk)
                    if offset % (256 * 1024 * 1024) < len(chunk):
                        print(
                            json.dumps(
                                {
                                    "event": "confirmation_download_progress",
                                    "subject": subject,
                                    "bytes": offset,
                                    "expected": expected_bytes,
                                }
                            ),
                            flush=True,
                        )
        if partial.stat().st_size != expected_bytes:
            raise RuntimeError(f"Downloaded confirmation size mismatch: {partial}")
        if file_hash(partial, "md5") != expected_md5:
            raise RuntimeError(f"Downloaded confirmation MD5 mismatch: {partial}")
        os.replace(partial, final)
        status = "downloaded_and_verified"
        print(
            json.dumps(
                {
                    "event": "confirmation_download_complete",
                    "subject": subject,
                    "seconds": time.time() - started,
                }
            ),
            flush=True,
        )
    record = {
        "schema": "KITE-ST-HD-confirmation-acquisition-v1",
        "status": status,
        "subject": subject,
        "source_record": "https://zenodo.org/records/4518754",
        "source_url": f"{BASE}/{subject}.tar.gz/content",
        "path": final.relative_to(ROOT).as_posix(),
        "bytes": expected_bytes,
        "md5": expected_md5,
        "sha256": file_hash(final, "sha256"),
        "confirmation_implementation_lock_sha256": lock_hash,
        "archive_opened_by_acquisition": False,
        "eeg_numerical_sample_read_by_acquisition": False,
    }
    record_path = METADATA_ROOT / f"{subject}_archive_acquisition_v1.json"
    if record_path.exists():
        previous = json.loads(record_path.read_text(encoding="utf-8"))
        immutable = {key: value for key, value in record.items() if key != "status"}
        if any(previous.get(key) != value for key, value in immutable.items()):
            raise RuntimeError(f"Existing confirmation acquisition record mismatch: {subject}")
    else:
        record_path.write_text(json.dumps(record, indent=2) + "\n", encoding="utf-8")
        record_hash = file_hash(record_path, "sha256")
        record_path.with_name(record_path.name + ".sha256").write_text(
            f"{record_hash}  {record_path.name}\n", encoding="ascii"
        )
    return record


def main() -> None:
    parser = argparse.ArgumentParser()
    parser.add_argument("--subject", choices=SUBJECTS)
    parser.add_argument("--all", action="store_true")
    args = parser.parse_args()
    if bool(args.subject) == bool(args.all):
        raise RuntimeError("Choose exactly one of --subject or --all.")
    lock, lock_hash = validate_confirmation_lock(Path(__file__))
    if tuple(lock["confirmation_subject_order"]) != SUBJECTS:
        raise RuntimeError("Confirmation subjects differ from implementation lock.")
    selected = SUBJECTS if args.all else (args.subject,)
    records = [retrieve(subject, lock_hash) for subject in selected]
    if args.all:
        if SUMMARY.exists():
            raise RuntimeError("Refusing to overwrite confirmation acquisition summary.")
        payload = {
            "schema": "KITE-ST-HD-confirmation-acquisition-summary-v1",
            "status": "pass",
            "confirmation_implementation_lock_sha256": lock_hash,
            "subjects": list(SUBJECTS),
            "archive_count": len(records),
            "total_bytes": int(sum(record["bytes"] for record in records)),
            "records": records,
        }
        SUMMARY.parent.mkdir(parents=True, exist_ok=True)
        SUMMARY.write_text(json.dumps(payload, indent=2) + "\n", encoding="utf-8")
        observed = file_hash(SUMMARY, "sha256")
        SUMMARY.with_name(SUMMARY.name + ".sha256").write_text(
            f"{observed}  {SUMMARY.name}\n", encoding="ascii"
        )
    print(json.dumps({"status": "pass", "subjects": list(selected)}))


if __name__ == "__main__":
    main()
```
