## Supplementary material for "Falsifiable substitution tests reveal task-structured neural evidence for auditory attention": Article_Analysis_Source_Code: download_kite_st_hd_development_v1.py

```
"""Resumable firewall-limited acquisition of the 12 development archives."""

from __future__ import annotations

import argparse
import hashlib
import json
import os
import time
import urllib.request
from pathlib import Path

from kite_st_hd_lock_utils_v1 import ROOT, validate_implementation_lock


SUBJECTS = (
    "S7", "S4", "S24", "S11", "S23", "S14",
    "S29", "S19", "S17", "S12", "S25", "S27",
)
FILES = {
    "S7": (1531972990, "dd59db3a63c2da91cb4d83dbf618e81a"),
    "S4": (1610163270, "4c6b7b7662fed95e7cf19eef15ceb727"),
    "S24": (1562760875, "bffd98033aec15c18a6a7604760df920"),
    "S11": (1534385366, "a04cf071d7cfa2e0c4be664ac349efd6"),
    "S23": (1655100979, "c69dcb5801759fa2ae5b903e8243faf1"),
    "S14": (1615117902, "189ac6e6feec4880e38673df18ba316e"),
    "S29": (1586569020, "05f3e910867faff6fa615e7e96aee4bd"),
    "S19": (1560570367, "3d7ab1442dbab1573837a8c125c771a2"),
    "S17": (1604413128, "2da02728804ade257ec564b327368bdb"),
    "S12": (1591853456, "f2d9451ee1cd490992bcd4dbfe424356"),
    "S25": (1552337050, "aaad26facc4e5f68d00da9041d9ce728"),
    "S27": (1618113174, "59a7b2a5282da7e8f960cd6884dc1ac3"),
}
OUTPUT_ROOT = (
    ROOT / "data" / "raw" / "kite_st_hd_zenodo4518754" / "development_archives"
)
METADATA_ROOT = ROOT / "data" / "metadata" / "kite_st_hd_development_downloads"
SUMMARY = (
    ROOT
    / "results"
    / "kite_st_hd"
    / "development"
    / "development_acquisition_summary_v1.json"
)
BASE = "https://zenodo.org/api/records/4518754/files"


def file_hash(path: Path, algorithm: str) -> str:
    digest = hashlib.new(algorithm)
    with path.open("rb") as stream:
        for chunk in iter(lambda: stream.read(8 * 1024 * 1024), b""):
            digest.update(chunk)
    return digest.hexdigest()


def retrieve(subject: str, lock_hash: str) -> dict:
    expected_bytes, expected_md5 = FILES[subject]
    OUTPUT_ROOT.mkdir(parents=True, exist_ok=True)
    METADATA_ROOT.mkdir(parents=True, exist_ok=True)
    final = OUTPUT_ROOT / f"{subject}.tar.gz"
    partial = final.with_name(final.name + ".part")
    if final.exists():
        if final.stat().st_size != expected_bytes:
            raise RuntimeError(f"Existing archive size mismatch: {final}")
        if file_hash(final, "md5") != expected_md5:
            raise RuntimeError(f"Existing archive MD5 mismatch: {final}")
        status = "already_present_and_verified"
    else:
        offset = partial.stat().st_size if partial.exists() else 0
        if offset > expected_bytes:
            raise RuntimeError(f"Partial file exceeds expected size: {partial}")
        request = urllib.request.Request(
            f"{BASE}/{subject}.tar.gz/content",
            headers={
                "User-Agent": "KITE-ST-HD-development-acquisition/1.0",
                "Range": f"bytes={offset}-",
            },
        )
        started = time.time()
        with urllib.request.urlopen(request, timeout=120) as response:
            code = getattr(response, "status", response.getcode())
            if offset and code != 206:
                offset = 0
                mode = "wb"
            else:
                mode = "ab" if offset else "wb"
            with partial.open(mode) as stream:
                while True:
                    chunk = response.read(8 * 1024 * 1024)
                    if not chunk:
                        break
                    stream.write(chunk)
                    offset += len(chunk)
                    if offset % (256 * 1024 * 1024) < len(chunk):
                        print(
                            json.dumps(
                                {
                                    "event": "download_progress",
                                    "subject": subject,
                                    "bytes": offset,
                                    "expected": expected_bytes,
                                }
                            ),
                            flush=True,
                        )
        if partial.stat().st_size != expected_bytes:
            raise RuntimeError(f"Downloaded archive size mismatch: {partial}")
        if file_hash(partial, "md5") != expected_md5:
            raise RuntimeError(f"Downloaded archive MD5 mismatch: {partial}")
        os.replace(partial, final)
        status = "downloaded_and_verified"
        print(
            json.dumps(
                {
                    "event": "download_complete",
                    "subject": subject,
                    "seconds": time.time() - started,
                }
            ),
            flush=True,
        )
    observed_sha256 = file_hash(final, "sha256")
    record = {
        "schema": "KITE-ST-HD-development-acquisition-v1",
        "status": status,
        "subject": subject,
        "source_record": "https://zenodo.org/records/4518754",
        "source_url": f"{BASE}/{subject}.tar.gz/content",
        "path": final.relative_to(ROOT).as_posix(),
        "bytes": expected_bytes,
        "md5": expected_md5,
        "sha256": observed_sha256,
        "implementation_lock_sha256": lock_hash,
        "archive_opened": False,
        "eeg_numerical_sample_read": False,
        "confirmation_file_acquired": False,
    }
    record_path = METADATA_ROOT / f"{subject}_archive_acquisition_v1.json"
    if record_path.exists():
        previous = json.loads(record_path.read_text(encoding="utf-8"))
        immutable = {
            key: value
            for key, value in record.items()
            if key != "status"
        }
        if any(previous.get(key) != value for key, value in immutable.items()):
            raise RuntimeError(f"Existing acquisition record mismatch: {subject}")
    else:
        record_path.write_text(json.dumps(record, indent=2) + "\n", encoding="utf-8")
        record_hash = file_hash(record_path, "sha256")
        record_path.with_name(record_path.name + ".sha256").write_text(
            f"{record_hash}  {record_path.name}\n",
            encoding="ascii",
        )
    return record


def main() -> None:
    parser = argparse.ArgumentParser()
    parser.add_argument("--subject", choices=SUBJECTS)
    parser.add_argument("--all", action="store_true")
    args = parser.parse_args()
    if bool(args.subject) == bool(args.all):
        raise RuntimeError("Choose exactly one of --subject or --all.")
    lock, lock_hash = validate_implementation_lock(Path(__file__))
    if tuple(lock["development_subjects"]) != SUBJECTS:
        raise RuntimeError("Development subject order differs from lock.")
    selected = SUBJECTS if args.all else (args.subject,)
    records = []
    for subject in selected:
        records.append(retrieve(subject, lock_hash))
    if args.all:
        if SUMMARY.exists():
            raise RuntimeError("Refusing to overwrite development acquisition summary.")
        payload = {
            "schema": "KITE-ST-HD-development-acquisition-summary-v1",
            "status": "pass",
            "implementation_lock_sha256": lock_hash,
            "subjects": list(SUBJECTS),
            "archive_count": len(records),
            "total_bytes": int(sum(item["bytes"] for item in records)),
            "records": records,
            "confirmation_file_acquired": False,
        }
        SUMMARY.parent.mkdir(parents=True, exist_ok=True)
        SUMMARY.write_text(json.dumps(payload, indent=2) + "\n", encoding="utf-8")
        observed = file_hash(SUMMARY, "sha256")
        SUMMARY.with_name(SUMMARY.name + ".sha256").write_text(
            f"{observed}  {SUMMARY.name}\n",
            encoding="ascii",
        )
    print(json.dumps({"status": "pass", "subjects": list(selected)}))


if __name__ == "__main__":
    main()
```
