## Supplementary material for "Falsifiable substitution tests reveal task-structured neural evidence for auditory attention": Article_Analysis_Source_Code: download_kojima2024b_structure_metadata_v1.py

```
"""Download outcome-blind structure metadata for Kojima2024B.

Only README, BrainVision headers, and marker files for ASME-2stream and
ASME-4stream are acquired. EEG binary files and NASA-TLX values are
deliberately excluded.
"""

from __future__ import annotations

import concurrent.futures
import hashlib
import json
from pathlib import Path

import requests


ROOT = Path(__file__).resolve().parents[1]
DOI = "doi:10.7910/DVN/1UJDV6"
API = "https://dataverse.harvard.edu/api"
OUT = ROOT / "data" / "raw" / "kojima2024b_structure_all"
MANIFEST = ROOT / "data" / "inventory" / "kojima2024b_structure_manifest_v1.json"
SUBJECTS = tuple("ABCDEFGHIJKLMNO")


def file_hash(path: Path, algorithm: str) -> str:
    digest = hashlib.new(algorithm)
    with path.open("rb") as handle:
        while block := handle.read(1024 * 1024):
            digest.update(block)
    return digest.hexdigest()


def main() -> None:
    response = requests.get(
        f"{API}/datasets/:persistentId/",
        params={"persistentId": DOI},
        timeout=60,
    )
    response.raise_for_status()
    latest = response.json()["data"]["latestVersion"]
    if (latest["versionNumber"], latest["versionMinorNumber"]) != (1, 0):
        raise RuntimeError("Unexpected Dataverse version")

    selected = []
    for item in latest["files"]:
        label = item["label"]
        directory = item.get("directoryLabel", "")
        if label == "README.md":
            selected.append(item)
            continue
        is_task = "task-2stream" in label or "task-4stream" in label
        is_structure = label.endswith(".vhdr") or label.endswith(".vmrk")
        if directory.endswith("/eeg") and is_task and is_structure:
            selected.append(item)
    expected = 1 + len(SUBJECTS) * 12 * 2
    if len(selected) != expected:
        raise RuntimeError(f"Expected {expected} structure files, got {len(selected)}")
    if any(
        item["label"].endswith(".eeg")
        or "nasatlx" in item.get("directoryLabel", "").lower()
        for item in selected
    ):
        raise RuntimeError("Outcome-blind selection includes forbidden data")

    def download_one(item: dict) -> dict:
        data_file = item["dataFile"]
        relative = (
            Path(item.get("directoryLabel", "")) / item["label"]
            if item.get("directoryLabel")
            else Path(item["label"])
        )
        target = OUT / relative
        target.parent.mkdir(parents=True, exist_ok=True)
        expected_md5 = data_file["checksum"]["value"].lower()
        expected_size = int(data_file["filesize"])
        if not target.exists():
            download = requests.get(
                f"{API}/access/datafile/{data_file['id']}", timeout=60
            )
            download.raise_for_status()
            target.write_bytes(download.content)
        if target.stat().st_size != expected_size:
            raise RuntimeError(f"Size mismatch: {target}")
        observed_md5 = file_hash(target, "md5")
        if observed_md5 != expected_md5:
            raise RuntimeError(f"MD5 mismatch: {target}")
        return {
            "directory": item.get("directoryLabel", ""),
            "label": item["label"],
            "dataverse_file_id": data_file["id"],
            "bytes": expected_size,
            "official_md5": expected_md5,
            "local_md5": observed_md5,
            "local_sha256": file_hash(target, "sha256"),
            "local_path": str(target.relative_to(ROOT)).replace("\\", "/"),
        }

    records = []
    with concurrent.futures.ThreadPoolExecutor(max_workers=10) as pool:
        futures = [pool.submit(download_one, item) for item in selected]
        for index, future in enumerate(
            concurrent.futures.as_completed(futures), start=1
        ):
            records.append(future.result())
            if index % 50 == 0:
                print(f"VERIFIED {index}/{len(selected)} structure files")

    records.sort(key=lambda record: (record["directory"], record["label"]))
    summary = {
        "dataset": "Kojima2024B four-class ASME BCI",
        "persistent_id": DOI,
        "dataset_version": "1.0",
        "release_time": latest["releaseTime"],
        "scope": (
            "Outcome-blind structure only: README plus 2stream/4stream vhdr "
            "and vmrk; no EEG binary and no NASA-TLX values"
        ),
        "subjects": list(SUBJECTS),
        "n_files": len(records),
        "total_bytes": sum(record["bytes"] for record in records),
        "forbidden_EEG_files_downloaded": 0,
        "forbidden_NASA_TLX_files_downloaded": 0,
        "files": records,
    }
    MANIFEST.parent.mkdir(parents=True, exist_ok=True)
    MANIFEST.write_text(
        json.dumps(summary, indent=2, ensure_ascii=False) + "\n",
        encoding="utf-8",
    )
    print(f"MANIFEST {MANIFEST}")
    print(f"SHA256 {file_hash(MANIFEST, 'sha256')}")
    print(f"TOTAL {summary['n_files']} files {summary['total_bytes']:,} bytes")


if __name__ == "__main__":
    main()
```
