## Supplementary material for "Falsifiable substitution tests reveal task-structured neural evidence for auditory attention": Article_Analysis_Source_Code: download_kojima2024b_wz_hic_confirmation_eeg_v1.py

```
"""Download and verify only the WZ-HIC-v1 confirmation EEG binaries."""

from __future__ import annotations

import concurrent.futures
import hashlib
import json
import shutil
import threading
from datetime import datetime, timezone
from pathlib import Path

import requests


ROOT = Path(__file__).resolve().parents[1]
DATA_ROOT = ROOT / "data" / "raw" / "kojima2024b_structure_all"
DOI = "doi:10.7910/DVN/1UJDV6"
API = "https://dataverse.harvard.edu/api"
PROTOCOL = ROOT / "docs" / "wz_hic_kojima2024b_confirmation_protocol_v1.json"
MANIFEST = (
    ROOT
    / "data"
    / "inventory"
    / "kojima2024b_wz_hic_confirmation_eeg_manifest_v1.json"
)
SUBJECTS = tuple("BDFHJLN")
print_lock = threading.Lock()


def file_hash(path: Path, algorithm: str) -> str:
    h = hashlib.new(algorithm)
    with path.open("rb") as f:
        for block in iter(lambda: f.read(4 * 1024 * 1024), b""):
            h.update(block)
    return h.hexdigest()


def main() -> None:
    if not PROTOCOL.exists():
        raise FileNotFoundError(PROTOCOL)
    if list(DATA_ROOT.glob("sub-*/nasatlx/*")):
        raise RuntimeError("NASA-TLX must remain sealed.")
    response = requests.get(
        f"{API}/datasets/:persistentId/",
        params={"persistentId": DOI},
        timeout=60,
    )
    response.raise_for_status()
    latest = response.json()["data"]["latestVersion"]
    if (latest["versionNumber"], latest["versionMinorNumber"]) != (1, 0):
        raise RuntimeError("Unexpected Dataverse version.")
    selected = []
    for item in latest["files"]:
        directory = item.get("directoryLabel", "")
        label = item["label"]
        if (
            any(directory == f"sub-{s}/eeg" for s in SUBJECTS)
            and label.endswith(".eeg")
            and ("task-2stream" in label or "task-4stream" in label)
        ):
            selected.append(item)
    expected = len(SUBJECTS) * 2 * 6
    if len(selected) != expected:
        raise RuntimeError(f"Expected {expected}, got {len(selected)}.")
    total = sum(int(item["dataFile"]["filesize"]) for item in selected)
    free = shutil.disk_usage(ROOT).free
    if free < total + 5 * 1024**3:
        raise RuntimeError(f"Insufficient safety margin: free={free}, total={total}")
    print(f"SELECTED {len(selected)} files {total:,} bytes", flush=True)

    def download_one(item: dict) -> dict:
        d = item["dataFile"]
        target = DATA_ROOT / item["directoryLabel"] / item["label"]
        target.parent.mkdir(parents=True, exist_ok=True)
        part = target.with_suffix(".eeg.part")
        size = int(d["filesize"])
        md5 = d["checksum"]["value"].lower()
        if target.exists() and (
            target.stat().st_size != size or file_hash(target, "md5") != md5
        ):
            raise RuntimeError(f"Existing mismatch: {target}")
        if not target.exists():
            start = part.stat().st_size if part.exists() else 0
            headers = {"Range": f"bytes={start}-"} if start else {}
            with requests.get(
                f"{API}/access/datafile/{d['id']}",
                headers=headers,
                stream=True,
                timeout=(30, 120),
            ) as r:
                r.raise_for_status()
                resumed = start > 0 and r.status_code == 206
                mode = "ab" if resumed else "wb"
                with part.open(mode) as f:
                    for block in r.iter_content(4 * 1024 * 1024):
                        if block:
                            f.write(block)
            if part.stat().st_size != size:
                raise RuntimeError(f"Size mismatch: {part}")
            part.replace(target)
        local_md5 = file_hash(target, "md5")
        if local_md5 != md5:
            raise RuntimeError(f"MD5 mismatch: {target}")
        record = {
            "directory": item["directoryLabel"],
            "label": item["label"],
            "dataverse_file_id": d["id"],
            "bytes": size,
            "official_md5": md5,
            "local_md5": local_md5,
            "local_sha256": file_hash(target, "sha256"),
            "local_path": str(target.relative_to(ROOT)).replace("\\", "/"),
        }
        with print_lock:
            print(f"VERIFIED {record['local_path']}", flush=True)
        return record

    records = []
    with concurrent.futures.ThreadPoolExecutor(max_workers=4) as pool:
        futures = [pool.submit(download_one, item) for item in selected]
        for future in concurrent.futures.as_completed(futures):
            records.append(future.result())
    records.sort(key=lambda x: (x["directory"], x["label"]))
    manifest = {
        "status": "PASS",
        "created_utc": datetime.now(timezone.utc).isoformat(),
        "persistent_id": DOI,
        "dataset_version": "1.0",
        "protocol_sha256": file_hash(PROTOCOL, "sha256"),
        "scope": "WZ-HIC-v1 confirmation 2stream and 4stream EEG only",
        "participants": list(SUBJECTS),
        "n_files": len(records),
        "total_bytes": sum(x["bytes"] for x in records),
        "NASA_TLX_value_files_present": len(
            list(DATA_ROOT.glob("sub-*/nasatlx/*"))
        ),
        "files": records,
    }
    if manifest["NASA_TLX_value_files_present"]:
        raise RuntimeError("NASA-TLX appeared during acquisition.")
    MANIFEST.write_text(
        json.dumps(manifest, ensure_ascii=False, indent=2) + "\n", encoding="utf-8"
    )
    print(MANIFEST)
    print(file_hash(MANIFEST, "sha256"))


if __name__ == "__main__":
    main()
```
