## Supplementary material for "Falsifiable substitution tests reveal task-structured neural evidence for auditory attention": Article_Analysis_Source_Code: extract_dtu_aad.py

```
"""Build the locked 32-Hz compact DTU external-evaluation dataset."""

from __future__ import annotations

import argparse
import hashlib
import json
from pathlib import Path

import numpy as np
from scipy.io import loadmat
from scipy.signal import butter, sosfiltfilt

from audit_dtu_aad import SUBJECTS, story_identity, trial_audio_paths
from extract_kul_discovery import (
    FS_MODEL,
    center_frequencies,
    process_audio,
    save_npz_atomic,
)


EXPECTED_LOCK_SHA256 = (
    "f87d8bf4ac291d2c1f2bb57138f0175335dc58a9624a205fbf5c5777dcf89440"
)


def sha256(path: Path) -> str:
    digest = hashlib.sha256()
    with path.open("rb") as handle:
        while block := handle.read(1024 * 1024):
            digest.update(block)
    return digest.hexdigest()


def extract_audio_cache(
    audio_root: Path, destination: Path
) -> dict[str, np.ndarray]:
    centers = center_frequencies()
    arrays: dict[str, np.ndarray] = {
        "center_frequencies_hz": centers.astype(np.float64),
        "metadata_json": np.asarray(
            json.dumps(
                {
                    "front_end": (
                        "same real FIR gammatone power-law subbands as "
                        "locked KUL candidate"
                    ),
                    "model_sample_rate_hz": FS_MODEL,
                    "center_frequencies_hz": centers.tolist(),
                }
            )
        ),
    }
    for path in sorted(audio_root.glob("*.wav")):
        print(f"DTU audio envelope: {path.name}", flush=True)
        arrays[f"audio::{path.name}"] = process_audio(path, centers)
    save_npz_atomic(destination, arrays)
    return {
        name.removeprefix("audio::"): value
        for name, value in arrays.items()
        if name.startswith("audio::")
    }


def process_eeg(value: np.ndarray) -> np.ndarray:
    eeg = np.asarray(value[:, :64], dtype=np.float64)
    bandpass = butter(
        4, (1.0, 9.0), btype="bandpass", fs=64, output="sos"
    )
    eeg = sosfiltfilt(bandpass, eeg, axis=0)
    eeg = eeg[::2]
    if eeg.shape != (1600, 64) or not np.isfinite(eeg).all():
        raise RuntimeError(f"Invalid processed DTU EEG shape {eeg.shape}")
    return eeg.astype(np.float32)


def main() -> None:
    parser = argparse.ArgumentParser()
    parser.add_argument(
        "--data-root", type=Path, default=Path("data/raw/dtu_aad")
    )
    parser.add_argument(
        "--output-dir",
        type=Path,
        default=Path("data/derived/dtu_aad_32hz"),
    )
    parser.add_argument(
        "--lock-file",
        type=Path,
        default=Path("docs/dtu_external_lock.json"),
    )
    parser.add_argument("--rebuild-audio", action="store_true")
    args = parser.parse_args()
    lock_hash = sha256(args.lock_file)
    if lock_hash != EXPECTED_LOCK_SHA256:
        raise RuntimeError(
            f"DTU lock hash mismatch: observed={lock_hash}"
        )

    args.output_dir.mkdir(parents=True, exist_ok=True)
    audio_cache_path = args.output_dir / "audio_powerlaw_subbands.npz"
    audio_root = args.data_root / "audio_extracted"
    if args.rebuild_audio or not audio_cache_path.exists():
        audio = extract_audio_cache(audio_root, audio_cache_path)
    else:
        archive = np.load(audio_cache_path)
        audio = {
            name.removeprefix("audio::"): archive[name]
            for name in archive.files
            if name.startswith("audio::")
        }

    manifest = {
        "dataset": "DTU auditory attention dataset",
        "participants": list(SUBJECTS),
        "sample_rate_hz": FS_MODEL,
        "scalp_channels": 64,
        "excluded_channels": ["EXG1", "EXG2"],
        "passband_hz": [1, 9],
        "audio_cache": audio_cache_path.name,
        "lock_file": str(args.lock_file),
        "lock_sha256": lock_hash,
        "participant_files": [],
    }
    preprocessed_root = args.data_root / "preprocessed_extracted"
    for subject in SUBJECTS:
        source = preprocessed_root / f"S{subject}_data_preproc.mat"
        data = loadmat(
            source,
            squeeze_me=True,
            struct_as_record=False,
            variable_names=["data"],
        )["data"]
        eeg_trials = list(np.atleast_1d(data.eeg))
        events = list(np.atleast_1d(data.event.eeg))
        paths = trial_audio_paths(data)
        arrays: dict[str, np.ndarray] = {}
        metadata = []
        for index, (attended_name, unattended_name) in enumerate(
            paths, start=1
        ):
            eeg = process_eeg(eeg_trials[index - 1])
            usable = min(
                eeg.shape[0],
                audio[attended_name].shape[0],
                audio[unattended_name].shape[0],
            )
            if usable != 1600:
                raise RuntimeError(
                    f"S{subject} T{index}: unexpected length {usable}"
                )
            attended_story = story_identity(attended_name)
            unattended_story = story_identity(unattended_name)
            arrays[f"eeg_{index:02d}"] = eeg[:usable]
            metadata.append(
                {
                    "subject": subject,
                    "trial": index,
                    "attended_audio": attended_name,
                    "unattended_audio": unattended_name,
                    "attended_story_identity": attended_story,
                    "unattended_story_identity": unattended_story,
                    "unordered_story_pair_group": "|".join(
                        sorted([attended_story, unattended_story])
                    ),
                    "attention_code": int(events[index - 1].value),
                    "samples": usable,
                }
            )
        arrays["metadata_json"] = np.asarray(json.dumps(metadata))
        destination = args.output_dir / f"S{subject}.npz"
        save_npz_atomic(destination, arrays)
        manifest["participant_files"].append(
            {"subject": subject, "file": destination.name}
        )
        print(f"DTU S{subject:02d}: 60 trials complete", flush=True)
        del data, arrays

    (args.output_dir / "manifest.json").write_text(
        json.dumps(manifest, indent=2), encoding="utf-8"
    )
    print(json.dumps(manifest, indent=2))


if __name__ == "__main__":
    main()
```
