## Supplementary material for "Falsifiable substitution tests reveal task-structured neural evidence for auditory attention": Article_Analysis_Source_Code: extract_kul_confirmation.py

```
"""Extract the hash-locked KUL even-ID confirmation partition at 32 Hz."""

from __future__ import annotations

import argparse
import hashlib
import json
from pathlib import Path

import numpy as np
from scipy.io import loadmat

from audit_kul_confirmation import (
    CONFIRMATION_SUBJECTS,
    EXPECTED_LOCK_SHA256,
)
from audit_kul_discovery import base_part, clean_stimulus_name
from extract_kul_discovery import process_eeg, save_npz_atomic


def sha256(path: Path) -> str:
    digest = hashlib.sha256()
    with path.open("rb") as handle:
        while block := handle.read(1024 * 1024):
            digest.update(block)
    return digest.hexdigest()


def main() -> None:
    parser = argparse.ArgumentParser()
    parser.add_argument(
        "--data-root",
        type=Path,
        default=Path("data/raw/kul_aad_confirmation"),
    )
    parser.add_argument(
        "--audio-cache",
        type=Path,
        default=Path(
            "data/derived/kul_discovery_32hz/audio_powerlaw_subbands.npz"
        ),
    )
    parser.add_argument(
        "--output-dir",
        type=Path,
        default=Path("data/derived/kul_confirmation_32hz"),
    )
    parser.add_argument(
        "--lock-file",
        type=Path,
        default=Path("docs/kul_candidate_lock.json"),
    )
    args = parser.parse_args()
    lock_hash = sha256(args.lock_file)
    if lock_hash != EXPECTED_LOCK_SHA256:
        raise RuntimeError(f"KUL candidate lock mismatch: {lock_hash}")
    observed = {
        int(path.stem[1:]) for path in args.data_root.glob("S*.mat")
    }
    if observed != set(CONFIRMATION_SUBJECTS):
        raise RuntimeError(
            f"Unexpected confirmation participant set: {observed}"
        )
    audio_archive = np.load(args.audio_cache)
    audio = {
        name.removeprefix("audio::"): audio_archive[name]
        for name in audio_archive.files
        if name.startswith("audio::")
    }
    if (
        len(audio) != 16
        or audio_archive["center_frequencies_hz"].size != 15
    ):
        raise RuntimeError("Locked KUL audio cache structure mismatch")

    args.output_dir.mkdir(parents=True, exist_ok=True)
    manifest = {
        "dataset": "KUL auditory attention dataset",
        "partition": "even-ID one-shot confirmation",
        "participants": list(CONFIRMATION_SUBJECTS),
        "discovery_participants_loaded": [],
        "sample_rate_hz": 32,
        "reference": "Cz, released channel 48",
        "passband_hz": [1, 9],
        "audio_cache": str(args.audio_cache),
        "candidate_lock": str(args.lock_file),
        "candidate_lock_sha256": lock_hash,
        "participant_files": [],
    }
    for subject in CONFIRMATION_SUBJECTS:
        path = args.data_root / f"S{subject}.mat"
        trials = loadmat(
            path, simplify_cells=True, variable_names=["trials"]
        )["trials"]
        arrays: dict[str, np.ndarray] = {}
        metadata = []
        for index, trial in enumerate(trials, start=1):
            eeg = process_eeg(trial["RawData"]["EegData"])
            original = [
                str(value) for value in np.atleast_1d(trial["stimuli"])
            ]
            clean = [clean_stimulus_name(value) for value in original]
            attended_index = (
                0 if str(trial["attended_ear"]).upper() == "L" else 1
            )
            usable = min(
                eeg.shape[0],
                audio[clean[0]].shape[0],
                audio[clean[1]].shape[0],
            )
            arrays[f"eeg_{index:02d}"] = eeg[:usable]
            metadata.append(
                {
                    "subject": subject,
                    "trial": index,
                    "experiment": int(trial["experiment"]),
                    "part": int(trial["part"]),
                    "base_part_identity_group": base_part(trial),
                    "repetition": bool(trial["repetition"]),
                    "condition": str(trial["condition"]),
                    "attended_ear": str(trial["attended_ear"]),
                    "attended_track": int(trial["attended_track"]),
                    "left_audio": clean[0],
                    "right_audio": clean[1],
                    "attended_audio": clean[attended_index],
                    "unattended_audio": clean[1 - attended_index],
                    "samples": usable,
                }
            )
        arrays["metadata_json"] = np.asarray(json.dumps(metadata))
        destination = args.output_dir / f"S{subject}.npz"
        save_npz_atomic(destination, arrays)
        manifest["participant_files"].append(
            {"subject": subject, "file": destination.name}
        )
        print(f"KUL confirmation S{subject:02d}: complete", flush=True)
        del trials, arrays
    (args.output_dir / "manifest.json").write_text(
        json.dumps(manifest, indent=2), encoding="utf-8"
    )
    print(json.dumps(manifest, indent=2))


if __name__ == "__main__":
    main()
```
