## Supplementary material for "Falsifiable substitution tests reveal task-structured neural evidence for auditory attention": Article_Analysis_Source_Code: extract_kul_discovery.py

```
"""Create the compact, odd-participant KUL discovery dataset at 32 Hz.

The acoustic front end follows the public KUL example script: audio is
resampled to 8 kHz, passed through 1.5-ERB-spaced real gammatone filters from
150 Hz to just below 4 kHz, compressed with a 0.6 power law, resampled to
128 Hz, band-pass filtered at 1--9 Hz, and downsampled to 32 Hz.  EEG is
Cz-referenced (channel 48 in the released 1-based indexing), filtered by the
same 1--9 Hz passband, and downsampled from 128 to 32 Hz.

Only odd-numbered discovery participants can be read.  Each participant is
stored in a separate compressed NPZ containing variable-length trial arrays;
the acoustic cache is stored once and referred to by exact clean WAV name.
"""

from __future__ import annotations

import argparse
import json
import re
from pathlib import Path

import numpy as np
from scipy.io import loadmat, wavfile
from scipy.signal import butter, oaconvolve, resample_poly, sosfiltfilt

from audit_kul_discovery import (
    DISCOVERY_SUBJECTS,
    base_part,
    clean_stimulus_name,
)


FS_AUDIO_INTERMEDIATE = 8000
FS_INTERMEDIATE = 128
FS_MODEL = 32
POWER = 0.6
LOW_HZ = 1.0
HIGH_HZ = 9.0
ERB_SPACING = 1.5
GAMMATONE_ORDER = 4
GAMMATONE_BW_MULTIPLIER = 1.5
GAMMATONE_DURATION_SECONDS = 0.080


def hz_to_erb_rate(frequency: np.ndarray | float) -> np.ndarray:
    return 21.4 * np.log10(1.0 + 4.37e-3 * np.asarray(frequency))


def erb_rate_to_hz(rate: np.ndarray | float) -> np.ndarray:
    return (10.0 ** (np.asarray(rate) / 21.4) - 1.0) / 4.37e-3


def center_frequencies() -> np.ndarray:
    low_rate = float(hz_to_erb_rate(150.0))
    high_rate = float(hz_to_erb_rate(4000.0))
    auditory_range = high_rate - low_rate
    intervals = int(np.floor(auditory_range / ERB_SPACING))
    remainder = auditory_range - intervals * ERB_SPACING
    rates = (
        low_rate
        + np.arange(intervals + 1) * ERB_SPACING
        + remainder / 2
    )
    return erb_rate_to_hz(rates)


def erb_bandwidth(frequency: float) -> float:
    return 24.7 * (4.37e-3 * frequency + 1.0)


def real_gammatone_fir(frequency: float) -> np.ndarray:
    length = round(GAMMATONE_DURATION_SECONDS * FS_AUDIO_INTERMEDIATE)
    time = np.arange(length, dtype=np.float64) / FS_AUDIO_INTERMEDIATE
    beta = (
        1.019
        * erb_bandwidth(frequency)
        * GAMMATONE_BW_MULTIPLIER
    )
    impulse = (
        time ** (GAMMATONE_ORDER - 1)
        * np.cos(2 * np.pi * frequency * time)
        * np.exp(-2 * np.pi * beta * time)
    )
    response = np.sum(
        impulse
        * np.exp(
            -2j
            * np.pi
            * frequency
            * np.arange(length)
            / FS_AUDIO_INTERMEDIATE
        )
    )
    impulse /= max(abs(response), 1e-12)
    return impulse


def process_audio(path: Path, centers: np.ndarray) -> np.ndarray:
    sample_rate, raw = wavfile.read(path)
    audio = np.asarray(raw, dtype=np.float64)
    if np.issubdtype(raw.dtype, np.integer):
        audio /= max(abs(np.iinfo(raw.dtype).min), np.iinfo(raw.dtype).max)
    if audio.ndim != 1:
        audio = audio.mean(axis=1)
    audio = resample_poly(audio, FS_AUDIO_INTERMEDIATE, sample_rate)
    bandpass = butter(
        4,
        (LOW_HZ, HIGH_HZ),
        btype="bandpass",
        fs=FS_INTERMEDIATE,
        output="sos",
    )
    output = np.empty(
        (
            int(np.ceil(audio.size * FS_INTERMEDIATE / FS_AUDIO_INTERMEDIATE))
            // (FS_INTERMEDIATE // FS_MODEL),
            centers.size,
        ),
        dtype=np.float32,
    )
    for index, center in enumerate(centers):
        impulse = real_gammatone_fir(float(center))
        filtered = oaconvolve(audio, impulse, mode="full")[: audio.size]
        envelope = np.abs(filtered) ** POWER
        envelope = resample_poly(
            envelope, FS_INTERMEDIATE, FS_AUDIO_INTERMEDIATE
        )
        envelope = sosfiltfilt(bandpass, envelope)
        envelope = envelope[:: FS_INTERMEDIATE // FS_MODEL]
        if envelope.size < output.shape[0]:
            output = output[: envelope.size]
        output[:, index] = envelope[: output.shape[0]]
    if not np.isfinite(output).all():
        raise RuntimeError(f"Non-finite audio envelope: {path}")
    return output


def save_npz_atomic(path: Path, arrays: dict[str, np.ndarray]) -> None:
    temporary = path.with_suffix(path.suffix + ".tmp")
    with temporary.open("wb") as handle:
        np.savez_compressed(handle, **arrays)
    temporary.replace(path)


def extract_audio_cache(
    stimulus_root: Path, destination: Path
) -> dict[str, np.ndarray]:
    centers = center_frequencies()
    arrays: dict[str, np.ndarray] = {
        "center_frequencies_hz": centers.astype(np.float64),
        "metadata_json": np.asarray(
            json.dumps(
                {
                    "source": "KUL public preprocess_data.m",
                    "front_end": "real FIR gammatone power-law subbands",
                    "audio_intermediate_sample_rate_hz": FS_AUDIO_INTERMEDIATE,
                    "intermediate_sample_rate_hz": FS_INTERMEDIATE,
                    "model_sample_rate_hz": FS_MODEL,
                    "passband_hz": [LOW_HZ, HIGH_HZ],
                    "power": POWER,
                    "erb_spacing": ERB_SPACING,
                    "gammatone_order": GAMMATONE_ORDER,
                    "gammatone_bandwidth_multiplier": (
                        GAMMATONE_BW_MULTIPLIER
                    ),
                    "center_frequencies_hz": centers.tolist(),
                }
            )
        ),
    }
    for path in sorted(stimulus_root.glob("*_dry.wav")):
        print(f"Audio envelope: {path.name}", flush=True)
        arrays[f"audio::{path.name}"] = process_audio(path, centers)
    save_npz_atomic(destination, arrays)
    return {
        name.removeprefix("audio::"): value
        for name, value in arrays.items()
        if name.startswith("audio::")
    }


def load_audio_cache(path: Path) -> dict[str, np.ndarray]:
    archive = np.load(path)
    return {
        name.removeprefix("audio::"): archive[name]
        for name in archive.files
        if name.startswith("audio::")
    }


def process_eeg(value: np.ndarray) -> np.ndarray:
    eeg = np.asarray(value, dtype=np.float64)
    eeg = eeg - eeg[:, [47]]
    bandpass = butter(
        4,
        (LOW_HZ, HIGH_HZ),
        btype="bandpass",
        fs=FS_INTERMEDIATE,
        output="sos",
    )
    eeg = sosfiltfilt(bandpass, eeg, axis=0)
    eeg = eeg[:: FS_INTERMEDIATE // FS_MODEL]
    if not np.isfinite(eeg).all():
        raise RuntimeError("Non-finite processed EEG")
    return eeg.astype(np.float32)


def main() -> None:
    parser = argparse.ArgumentParser()
    parser.add_argument(
        "--data-root", type=Path, default=Path("data/raw/kul_aad")
    )
    parser.add_argument(
        "--output-dir",
        type=Path,
        default=Path("data/derived/kul_discovery_32hz"),
    )
    parser.add_argument("--rebuild-audio", action="store_true")
    args = parser.parse_args()

    observed = {
        int(match.group(1))
        for path in args.data_root.glob("S*.mat")
        if (match := re.fullmatch(r"S(\d+)\.mat", path.name))
    }
    if observed != set(DISCOVERY_SUBJECTS):
        raise RuntimeError(
            "KUL confirmation firewall violation or incomplete discovery "
            f"partition: observed={sorted(observed)}"
        )

    args.output_dir.mkdir(parents=True, exist_ok=True)
    audio_cache_path = args.output_dir / "audio_powerlaw_subbands.npz"
    stimulus_root = args.data_root / "stimuli_extracted" / "stimuli"
    if args.rebuild_audio or not audio_cache_path.exists():
        audio = extract_audio_cache(stimulus_root, audio_cache_path)
    else:
        audio = load_audio_cache(audio_cache_path)

    manifest = {
        "partition": "odd-participant discovery only",
        "participants": list(DISCOVERY_SUBJECTS),
        "confirmation_participants_loaded": [],
        "sample_rate_hz": FS_MODEL,
        "reference": "Cz, released channel 48",
        "passband_hz": [LOW_HZ, HIGH_HZ],
        "identity_group": "((part - 1) mod 4) + 1",
        "audio_cache": audio_cache_path.name,
        "participant_files": [],
    }
    for subject in DISCOVERY_SUBJECTS:
        source = args.data_root / f"S{subject}.mat"
        trials = loadmat(
            source, simplify_cells=True, variable_names=["trials"]
        )["trials"]
        arrays: dict[str, np.ndarray] = {}
        metadata = []
        for index, trial in enumerate(trials, start=1):
            eeg = process_eeg(trial["RawData"]["EegData"])
            original = [
                str(value) for value in np.atleast_1d(trial["stimuli"])
            ]
            clean = [clean_stimulus_name(value) for value in original]
            attended_index = (
                0 if str(trial["attended_ear"]).upper() == "L" else 1
            )
            max_audio_samples = min(audio[name].shape[0] for name in clean)
            usable = min(eeg.shape[0], max_audio_samples)
            arrays[f"eeg_{index:02d}"] = eeg[:usable]
            metadata.append(
                {
                    "subject": subject,
                    "trial": index,
                    "experiment": int(trial["experiment"]),
                    "part": int(trial["part"]),
                    "base_part_identity_group": base_part(trial),
                    "repetition": bool(trial["repetition"]),
                    "condition": str(trial["condition"]),
                    "attended_ear": str(trial["attended_ear"]),
                    "attended_track": int(trial["attended_track"]),
                    "left_audio": clean[0],
                    "right_audio": clean[1],
                    "attended_audio": clean[attended_index],
                    "unattended_audio": clean[1 - attended_index],
                    "samples": usable,
                }
            )
            print(
                f"S{subject:02d} T{index:02d}: "
                f"{usable / FS_MODEL:.1f} s",
                flush=True,
            )
        arrays["metadata_json"] = np.asarray(json.dumps(metadata))
        destination = args.output_dir / f"S{subject}.npz"
        save_npz_atomic(destination, arrays)
        manifest["participant_files"].append(
            {"subject": subject, "file": destination.name}
        )
        del trials, arrays

    (args.output_dir / "manifest.json").write_text(
        json.dumps(manifest, indent=2), encoding="utf-8"
    )
    print(json.dumps(manifest, indent=2))


if __name__ == "__main__":
    main()
```
