## Supplementary material for "Falsifiable substitution tests reveal task-structured neural evidence for auditory attention": Article_Analysis_Source_Code: inspect_kite_st_hd_engineering_v1.py

```
"""Value-blind Curry parsing and signal-QC inspection for KITE-ST-HD.

Only the three protocol-declared engineering participants are accepted.  Each
recording is inspected independently.  The code contains no L/R subtraction,
no true-label decoder, no edge selection and no geometry endpoint.
"""

from __future__ import annotations

import argparse
import hashlib
import json
import re
from datetime import datetime
from pathlib import Path

import numpy as np


ROOT = Path(__file__).resolve().parents[1]
PROTOCOL = ROOT / "docs" / "kite_st_hd_preoutcome_protocol_v1.json"
PROTOCOL_HASH = "f1096ab5ed46a54e26a95c2f19f8e2619c5a8feac768c7fb77d1e85d7f09feb8"
AMENDMENT = ROOT / "docs" / "kite_st_hd_preoutcome_amendment_01.json"
AMENDMENT_HASH = "d53e96bc121cc1ec200f44ebad32bfbd8e295a28e1007c2b5373b18806719ded"
WORK = ROOT / "data" / "raw" / "kite_st_hd_zenodo4518754" / "engineering_work"
ACQUISITION = ROOT / "data" / "metadata" / "kite_st_hd_engineering_downloads"
OUTPUT_ROOT = ROOT / "results" / "kite_st_hd" / "engineering"
WHITELIST = ("S20", "S6", "S8")
CONDITIONS = ("1L", "1R", "2L", "2R")


def file_hash(path: Path, algorithm: str = "sha256") -> str:
    digest = hashlib.new(algorithm)
    with path.open("rb") as stream:
        for chunk in iter(lambda: stream.read(8 * 1024 * 1024), b""):
            digest.update(chunk)
    return digest.hexdigest()


def lock_check() -> dict:
    if file_hash(PROTOCOL) != PROTOCOL_HASH:
        raise RuntimeError("Protocol hash mismatch.")
    if file_hash(AMENDMENT) != AMENDMENT_HASH:
        raise RuntimeError("Coordinate amendment hash mismatch.")
    protocol = json.loads(PROTOCOL.read_text(encoding="utf-8"))
    if tuple(protocol["sample_split"]["engineering_only"]) != WHITELIST:
        raise RuntimeError("Engineering whitelist mismatch.")
    return protocol


def scalar(text: str, key: str) -> str:
    match = re.search(rf"^\s*{re.escape(key)}\s*=\s*(.*?)\s*$", text, re.MULTILINE)
    if match is None:
        raise RuntimeError(f"Missing Curry DAP field {key}.")
    return match.group(1)


def block_lines(text: str, name: str) -> list[str]:
    pattern = (
        rf"^{re.escape(name)} START_LIST[^\n]*\n"
        rf"(.*?)"
        rf"^{re.escape(name)} END_LIST"
    )
    match = re.search(pattern, text, re.MULTILINE | re.DOTALL)
    if match is None:
        raise RuntimeError(f"Missing Curry RS3 list {name}.")
    return [line.strip() for line in match.group(1).splitlines() if line.strip()]


def parse_header(base: Path) -> dict:
    dap = base.with_suffix(".dap").read_text(encoding="utf-8", errors="strict")
    rs3 = base.with_suffix(".rs3").read_text(encoding="utf-8", errors="strict")
    scalp = block_lines(rs3, "LABELS")
    others = block_lines(rs3, "LABELS_OTHERS")
    labels = scalp + others
    fields = {
        key: int(scalar(dap, key))
        for key in (
            "NumSamples",
            "NumTrials",
            "NumChannels",
            "StartYear",
            "StartMonth",
            "StartDay",
            "StartHour",
            "StartMin",
            "StartSec",
            "StartMillisec",
        )
    }
    fields["SampleFreqHz"] = float(scalar(dap, "SampleFreqHz"))
    fields["DataFormat"] = scalar(dap, "DataFormat")
    fields["DataByteOrder"] = scalar(dap, "DataByteOrder")
    fields["DataSampOrder"] = scalar(dap, "DataSampOrder")
    fields["scalp_labels"] = scalp
    fields["other_labels"] = others
    fields["labels"] = labels
    fields["start_iso"] = datetime(
        fields["StartYear"],
        fields["StartMonth"],
        fields["StartDay"],
        fields["StartHour"],
        fields["StartMin"],
        fields["StartSec"],
        fields["StartMillisec"] * 1000,
    ).isoformat()
    if len(scalp) != 253 or others != ["XtraL", "XtraR", "Trigger"]:
        raise RuntimeError(
            f"Unexpected channel groups for {base.name}: {len(scalp)}, {others}"
        )
    if len(labels) != fields["NumChannels"] or fields["NumChannels"] != 256:
        raise RuntimeError("Curry channel count does not match label lists.")
    if fields["DataFormat"] != "FLOAT":
        raise RuntimeError("Only FLOAT Curry data are accepted.")
    if fields["DataByteOrder"] != "INTEL" or fields["DataSampOrder"] != "SAMP":
        raise RuntimeError("Unexpected Curry byte/sample order.")
    return fields


def fixed_qc_indices(start: int, stop: int, fs: int) -> np.ndarray:
    """Twenty non-overlapping one-second slices, fixed without labels."""
    width = fs
    starts = np.linspace(start, stop - width, 20, dtype=np.int64)
    return np.concatenate(
        [np.arange(value, value + width, dtype=np.int64) for value in starts]
    )


def inspect_recording(base: Path) -> dict:
    header = parse_header(base)
    n_samples = header["NumSamples"]
    n_channels = header["NumChannels"]
    sampling = int(header["SampleFreqHz"])
    dat = base.with_suffix(".dat")
    expected_bytes = n_samples * n_channels * 4
    if dat.stat().st_size != expected_bytes:
        raise RuntimeError(f"Curry DAT size mismatch: {dat}")
    data = np.memmap(
        dat,
        dtype="<f4",
        mode="r",
        shape=(n_samples, n_channels),
        order="C",
    )
    trigger = np.asarray(data[:, -1])
    finite_trigger = np.isfinite(trigger)
    if not finite_trigger.all():
        raise RuntimeError(f"Nonfinite trigger values in {dat}.")
    baseline = float(np.median(trigger[: 5 * sampling]))
    active = trigger > baseline
    rising = np.flatnonzero(active & ~np.r_[False, active[:-1]])
    falling = np.flatnonzero(~active & np.r_[False, active[:-1]])
    if len(rising) < 2:
        raise RuntimeError(f"Insufficient trigger rises in {dat}.")
    official_start = int(rising[0])
    official_stop = int(rising[-1])
    official_seconds = (official_stop - official_start) / sampling
    central_samples = 360 * sampling
    center = (official_start + official_stop) // 2
    central_start = center - central_samples // 2
    central_stop = central_start + central_samples
    if central_start < 0 or central_stop > n_samples:
        raise RuntimeError(f"Central 360-second crop is out of bounds: {dat}.")

    indices = fixed_qc_indices(central_start, central_stop, sampling)
    scalp_raw = np.asarray(data[indices, :253], dtype=np.float64)
    auxiliary = np.asarray(data[indices, 253:255], dtype=np.float64)
    finite_fraction = float(np.isfinite(scalp_raw).mean())
    if finite_fraction != 1.0:
        raise RuntimeError(f"Nonfinite scalp values in fixed QC slices: {dat}.")
    scalp_centered = scalp_raw - np.median(scalp_raw, axis=0, keepdims=True)
    scalp_car = scalp_centered - np.mean(scalp_centered, axis=1, keepdims=True)
    medians = np.median(scalp_car, axis=0)
    robust_scale = 1.4826 * np.median(
        np.abs(scalp_car - medians[None, :]),
        axis=0,
    )
    rms = np.sqrt(np.mean(scalp_car * scalp_car, axis=0))
    aux_centered = auxiliary - np.median(auxiliary, axis=0, keepdims=True)
    aux_scale = 1.4826 * np.median(np.abs(aux_centered), axis=0)

    mne_check: dict
    try:
        import mne

        raw = mne.io.read_raw_curry(str(base), preload=False, verbose="ERROR")
        direct = np.asarray(data[:5, :3], dtype=np.float64).T * 1e-6
        observed = raw.get_data(picks=[0, 1, 2], start=0, stop=5)
        mne_check = {
            "status": "pass",
            "n_times": int(raw.n_times),
            "channels": len(raw.ch_names),
            "sampling_hz": float(raw.info["sfreq"]),
            "last_labels": raw.ch_names[-3:],
            "maximum_absolute_first_sample_difference_volts": float(
                np.max(np.abs(direct - observed))
            ),
        }
        if (
            raw.n_times != n_samples
            or len(raw.ch_names) != n_channels
            or raw.ch_names[-3:] != ["XtraL", "XtraR", "Trigger"]
            or mne_check["maximum_absolute_first_sample_difference_volts"] > 1e-12
        ):
            raise RuntimeError(f"MNE/direct Curry check failed: {base}")
    except Exception as error:
        raise RuntimeError(f"MNE/direct Curry cross-reader failure: {base}") from error

    del data
    return {
        "recording": base.name,
        "header": {
            key: header[key]
            for key in (
                "NumSamples",
                "NumTrials",
                "NumChannels",
                "SampleFreqHz",
                "DataFormat",
                "DataByteOrder",
                "DataSampOrder",
                "start_iso",
            )
        },
        "file_bytes": dat.stat().st_size,
        "groups": {
            "scalp_channels": 253,
            "auxiliary_labels": header["other_labels"][:2],
            "trigger_label": header["other_labels"][2],
        },
        "trigger": {
            "baseline": baseline,
            "unique_values": [float(value) for value in np.unique(trigger)],
            "rising_edges": len(rising),
            "falling_edges": len(falling),
            "first_rise_sample": official_start,
            "last_rise_sample": official_stop,
            "median_inter_rise_samples": float(np.median(np.diff(rising))),
            "minimum_inter_rise_samples": int(np.min(np.diff(rising))),
            "maximum_inter_rise_samples": int(np.max(np.diff(rising))),
            "official_first_to_last_seconds": official_seconds,
        },
        "fixed_central_crop": {
            "start_sample": central_start,
            "stop_sample_exclusive": central_stop,
            "seconds": 360,
            "blocks_30_seconds": 12,
        },
        "value_blind_qc": {
            "fixed_one_second_slices": 20,
            "scalp_finite_fraction": finite_fraction,
            "car_robust_scale_summary_raw_units": {
                "minimum": float(np.min(robust_scale)),
                "q25": float(np.quantile(robust_scale, 0.25)),
                "median": float(np.median(robust_scale)),
                "q75": float(np.quantile(robust_scale, 0.75)),
                "maximum": float(np.max(robust_scale)),
            },
            "car_rms_summary_raw_units": {
                "minimum": float(np.min(rms)),
                "median": float(np.median(rms)),
                "maximum": float(np.max(rms)),
            },
            "auxiliary_robust_scale_raw_units": [
                float(value) for value in aux_scale
            ],
            "per_scalp_channel_robust_scale_raw_units": [
                float(value) for value in robust_scale
            ],
        },
        "mne_direct_cross_reader": mne_check,
    }


def main() -> None:
    parser = argparse.ArgumentParser()
    parser.add_argument("subject", choices=WHITELIST)
    args = parser.parse_args()
    protocol = lock_check()
    subject = args.subject
    acquisition_path = ACQUISITION / f"{subject}_archive_acquisition_v1.json"
    acquisition = json.loads(acquisition_path.read_text(encoding="utf-8"))
    if acquisition["subject"] != subject or not acquisition["status"].endswith("verified"):
        raise RuntimeError("Engineering acquisition record is invalid.")
    base_root = WORK / subject
    if not base_root.is_dir():
        raise FileNotFoundError(base_root)
    OUTPUT_ROOT.mkdir(parents=True, exist_ok=True)
    output = OUTPUT_ROOT / f"{subject}_value_blind_inspection_v1.json"
    if output.exists():
        raise RuntimeError(f"Refusing to overwrite engineering inspection: {output}")

    records = [
        inspect_recording(base_root / f"{subject}_AAD_{condition}")
        for condition in CONDITIONS
    ]
    by_start = sorted(records, key=lambda record: record["header"]["start_iso"])
    payload = {
        "schema": "KITE-ST-HD-value-blind-engineering-inspection-v1",
        "status": "pass",
        "subject": subject,
        "scope": "Independent per-recording parsing, trigger timing, fixed-slice signal QC and cross-reader equivalence.",
        "protocol_sha256": PROTOCOL_HASH,
        "amendment_sha256": AMENDMENT_HASH,
        "archive_acquisition_record": acquisition_path.relative_to(ROOT).as_posix(),
        "archive_sha256": acquisition["sha256"],
        "acquisition_order": [
            record["recording"] for record in by_start
        ],
        "records": records,
        "firewall": {
            "engineering_whitelist": list(WHITELIST),
            "true_L_minus_R_contrast_computed": False,
            "true_label_decoder_computed": False,
            "attention_edges_selected": False,
            "geometry_endpoint_computed": False,
            "development_or_confirmation_file_read": False,
            "paper_mutated": False,
        },
        "next_use": "Pool only label-blind QC summaries across all three engineering subjects, then freeze preprocessing thresholds and the implementation.",
    }
    output.write_text(json.dumps(payload, indent=2) + "\n", encoding="utf-8")
    digest = file_hash(output)
    output.with_name(output.name + ".sha256").write_text(
        f"{digest}  {output.name}\n",
        encoding="ascii",
    )
    print(json.dumps({"status": "pass", "output": str(output), "sha256": digest}))


if __name__ == "__main__":
    main()
```
