## Supplementary material for "Falsifiable substitution tests reveal task-structured neural evidence for auditory attention": Article_Analysis_Source_Code: kite_kul_akb_v1.py

```
"""Frozen KUL translation of the adaptive broad direction-delay decoder."""

from __future__ import annotations

from dataclasses import dataclass

import mne
import numpy as np
from scipy.linalg import cho_factor, cho_solve


FS = 32
TRIM = 5 * FS
WINDOW = 30 * FS
LAGS = np.asarray((0, 2, 3, 5, 6, 8), dtype=np.int64)
N_CHANNELS = 64


@dataclass(frozen=True)
class Window:
    trial: int
    part: int
    window_index: int
    attended_ear: str
    condition: str
    repetition: bool
    design: np.ndarray
    left: np.ndarray
    right: np.ndarray


def biosemi64_geometry() -> tuple[tuple[str, ...], np.ndarray, np.ndarray]:
    montage = mne.channels.make_standard_montage("biosemi64")
    labels = tuple(montage.ch_names)
    if len(labels) != N_CHANNELS:
        raise RuntimeError("MNE biosemi64 montage does not have 64 channels.")
    positions = montage.get_positions()["ch_pos"]
    xyz = np.row_stack([positions[label] for label in labels])
    xy = xyz[:, :2]
    angle = np.arctan2(xy[:, 1], xy[:, 0])
    sector = (
        np.floor((angle + np.pi) / (2 * np.pi) * 8).astype(np.int64) % 8
    )
    ids = np.tile(sector, len(LAGS))
    ids += np.repeat((0, 0, 0, 8, 8, 8), N_CHANNELS)
    if ids.shape != (384,) or np.any(np.bincount(ids, minlength=16) == 0):
        raise RuntimeError("Degenerate KUL direction-delay partition.")
    return labels, xy, ids


def make_design(eeg: np.ndarray) -> np.ndarray:
    value = np.asarray(eeg, dtype=np.float64)
    if value.shape != (WINDOW, N_CHANNELS):
        raise ValueError("KUL window must be 960 x 64.")
    common = WINDOW - int(LAGS[-1])
    design = np.column_stack(
        [value[int(lag) : int(lag) + common] for lag in LAGS]
    )
    if design.shape != (952, 384) or not np.isfinite(design).all():
        raise RuntimeError("Invalid KUL lag design.")
    return design.astype(np.float32)


def make_windows(
    eeg_trials: list[np.ndarray],
    metadata: list[dict],
    audio: dict[str, np.ndarray],
) -> list[Window]:
    if len(eeg_trials) != 20 or len(metadata) != 20:
        raise ValueError("KUL expects 20 trials per participant.")
    windows = []
    for eeg, row in zip(eeg_trials, metadata):
        samples = min(int(row["samples"]), len(eeg))
        left_full = np.mean(audio[str(row["left_audio"])], axis=1)
        right_full = np.mean(audio[str(row["right_audio"])], axis=1)
        samples = min(samples, len(left_full), len(right_full))
        usable = samples - 2 * TRIM
        count = max(0, usable // WINDOW)
        for window_index in range(count):
            first = TRIM + window_index * WINDOW
            stop = first + WINDOW
            design = make_design(eeg[first:stop])
            common = len(design)
            windows.append(
                Window(
                    trial=int(row["trial"]),
                    part=int(row["base_part_identity_group"]),
                    window_index=window_index,
                    attended_ear=str(row["attended_ear"]),
                    condition=str(row["condition"]),
                    repetition=bool(row["repetition"]),
                    design=design,
                    left=np.asarray(left_full[first : first + common], dtype=np.float64),
                    right=np.asarray(right_full[first : first + common], dtype=np.float64),
                )
            )
    if len(windows) != 132:
        raise RuntimeError(f"Frozen KUL window count differs: {len(windows)}")
    if {window.part for window in windows} != {1, 2, 3, 4}:
        raise RuntimeError("KUL base-part groups are incomplete.")
    return windows


def fit_ridge(windows: list[Window]) -> dict[str, np.ndarray | float]:
    if len(windows) < 20:
        raise ValueError("Too few KUL training windows.")
    dimension = len(LAGS) * N_CHANNELS
    count = sum(len(window.design) for window in windows)
    sum_x = np.zeros(dimension, dtype=np.float64)
    sum_x2 = np.zeros(dimension, dtype=np.float64)
    sum_y = 0.0
    for window in windows:
        x = np.asarray(window.design, dtype=np.float64)
        y = window.left if window.attended_ear == "L" else window.right
        sum_x += np.sum(x, axis=0)
        sum_x2 += np.sum(x * x, axis=0)
        sum_y += float(np.sum(y))
    x_mean = sum_x / count
    variance = np.maximum(sum_x2 / count - x_mean * x_mean, 0.0)
    x_scale = np.sqrt(variance)
    x_scale = np.where(x_scale > 1e-12, x_scale, 1.0)
    y_mean = sum_y / count
    gram = np.zeros((dimension, dimension), dtype=np.float64)
    cross = np.zeros(dimension, dtype=np.float64)
    for window in windows:
        x = (np.asarray(window.design, dtype=np.float64) - x_mean) / x_scale
        y = (
            window.left if window.attended_ear == "L" else window.right
        ) - y_mean
        gram += x.T @ x
        cross += x.T @ y
    gram_energy = float(np.mean(np.diag(gram)))
    alpha = gram_energy
    gram.flat[:: dimension + 1] += alpha
    factor = cho_factor(gram, lower=True, check_finite=False)
    weights = cho_solve(factor, cross, check_finite=False)
    if not np.isfinite(weights).all():
        raise RuntimeError("Nonfinite KUL ridge model.")
    return {
        "x_mean": x_mean,
        "x_scale": x_scale,
        "y_mean": y_mean,
        "weights": weights,
        "gram_energy": gram_energy,
        "alpha": alpha,
    }
```
