## Supplementary material for "Falsifiable substitution tests reveal task-structured neural evidence for auditory attention": Article_Analysis_Source_Code: kite_sat_adaptive_broad_v1.py

```
"""Adaptive cross-fitted broad weights for KITE-SAT."""

from __future__ import annotations

import numpy as np


PARTITIONS = ("P8", "P16", "S16")
WEIGHT_FAMILIES = ("positive", "signed", "softmax2", "top_half")
CANDIDATE_ORDER = tuple(
    f"{partition}_{family}"
    for partition in PARTITIONS
    for family in WEIGHT_FAMILIES
) + ("multiscale_positive",)


def reliability(aligned_contributions: np.ndarray) -> np.ndarray:
    values = np.asarray(aligned_contributions, dtype=np.float64)
    if values.ndim != 2 or len(values) < 4:
        raise ValueError("AKB reliability expects held blocks x cells.")
    rms = np.sqrt(np.mean(values * values, axis=0))
    output = np.mean(values, axis=0) / np.maximum(rms, 1e-12)
    if np.any(output < -1.0 - 1e-12) or np.any(output > 1.0 + 1e-12):
        raise RuntimeError("AKB reliability escaped its theoretical bounds.")
    return np.clip(output, -1.0, 1.0)


def family_weights(reliability_values: np.ndarray, family: str) -> np.ndarray:
    values = np.asarray(reliability_values, dtype=np.float64)
    if family == "positive":
        weights = np.maximum(values, 0.0)
    elif family == "signed":
        weights = values.copy()
    elif family == "softmax2":
        shifted = 2.0 * values
        weights = np.exp(shifted - np.max(shifted))
    elif family == "top_half":
        threshold = float(np.median(values))
        weights = ((values >= threshold) & (values > 0.0)).astype(np.float64)
    else:
        raise ValueError(f"Unknown AKB weight family: {family}")
    normalizer = float(np.sum(np.abs(weights)))
    if normalizer <= 1e-12:
        weights = np.ones_like(values)
        normalizer = float(len(weights))
    return weights / normalizer * len(weights)


def adaptive_score(
    delta_contributions: np.ndarray,
    reliability_values: np.ndarray,
    family: str,
) -> float:
    delta = np.asarray(delta_contributions, dtype=np.float64)
    weights = family_weights(reliability_values, family)
    if delta.shape != weights.shape:
        raise ValueError("AKB contribution/reliability shape mismatch.")
    return float(np.sum(delta * weights))
```
