## Supplementary material for "Falsifiable substitution tests reveal task-structured neural evidence for auditory attention": Article_Analysis_Source_Code: kite_sat_backward_v1.py

```
"""Cross-story backward stimulus reconstruction for KITE-SAT engineering."""

from __future__ import annotations

import numpy as np
from scipy.signal import resample_poly


DECODER_HZ = 20
LAGS_SAMPLES = np.arange(6, dtype=np.int64)
RIDGE_RATIOS = (0.001, 0.01, 0.1, 1.0)


def block_design(eeg_100hz: np.ndarray, audio_100hz: np.ndarray) -> tuple[np.ndarray, np.ndarray]:
    """Return causal neural-lag design and aligned envelope/onset targets."""
    eeg = np.asarray(eeg_100hz, dtype=np.float64)
    audio = np.asarray(audio_100hz, dtype=np.float64)
    if eeg.shape != (3000, 253) or audio.shape != (3000, 2):
        raise ValueError("KITE-SAT backward decoder expects one 30-s matched block.")
    eeg20 = resample_poly(eeg, 1, 5, axis=0)
    audio20 = resample_poly(audio, 1, 5, axis=0)
    common = len(eeg20) - int(LAGS_SAMPLES[-1])
    design = np.column_stack(
        [eeg20[int(lag) : int(lag) + common] for lag in LAGS_SAMPLES]
    )
    target = audio20[:common]
    if design.shape != (595, 1518) or target.shape != (595, 2):
        raise RuntimeError("Unexpected KITE-SAT backward-decoder block shape.")
    if not np.isfinite(design).all() or not np.isfinite(target).all():
        raise RuntimeError("Nonfinite KITE-SAT backward-decoder block.")
    return design, target


def fit_ridge_family(
    training_designs: list[np.ndarray],
    training_targets: list[np.ndarray],
) -> dict[float, dict[str, np.ndarray]]:
    """Fit the fixed ridge family with one eigendecomposition."""
    x = np.vstack(training_designs).astype(np.float64, copy=False)
    y = np.vstack(training_targets).astype(np.float64, copy=False)
    x_mean = np.mean(x, axis=0)
    x_scale = np.std(x, axis=0)
    x_scale = np.where(x_scale > 1e-12, x_scale, 1.0)
    y_mean = np.mean(y, axis=0)
    standardized = (x - x_mean) / x_scale
    centered_target = y - y_mean
    gram = standardized.T @ standardized
    cross = standardized.T @ centered_target
    eigenvalues, eigenvectors = np.linalg.eigh(gram)
    projected = eigenvectors.T @ cross
    energy = float(np.mean(np.diag(gram)))
    models: dict[float, dict[str, np.ndarray]] = {}
    for ratio in RIDGE_RATIOS:
        weights = eigenvectors @ (projected / (eigenvalues[:, None] + ratio * energy))
        models[ratio] = {
            "x_mean": x_mean,
            "x_scale": x_scale,
            "y_mean": y_mean,
            "weights": weights,
        }
    return models


def predict(design: np.ndarray, model: dict[str, np.ndarray]) -> np.ndarray:
    standardized = (np.asarray(design, dtype=np.float64) - model["x_mean"]) / model[
        "x_scale"
    ]
    return standardized @ model["weights"] + model["y_mean"]


def pearson(first: np.ndarray, second: np.ndarray) -> float:
    a = np.asarray(first, dtype=np.float64)
    b = np.asarray(second, dtype=np.float64)
    a = a - np.mean(a)
    b = b - np.mean(b)
    denominator = np.sqrt(np.sum(a * a) * np.sum(b * b))
    return float(np.sum(a * b) / max(float(denominator), 1e-12))
```
