## Supplementary material for "Falsifiable substitution tests reveal task-structured neural evidence for auditory attention": Article_Analysis_Source_Code: kite_sat_confirmation_complete_case_lock_utils_v2.py

```
"""Hash validation for the post-QC complete-case AKB confirmation v2."""

from __future__ import annotations

import hashlib
import json
from pathlib import Path


ROOT = Path(__file__).resolve().parents[1]
PROTOCOL = ROOT / "docs" / "kite_sat_akb_confirmation_complete_case_protocol_v2.json"
PROTOCOL_SHA256 = "64d3ef6866567f1da0a91c32fb095e44d788b4283de4ddc9a54e9341ec589807"
LOCK = (
    ROOT / "docs"
    / "kite_sat_akb_confirmation_complete_case_implementation_lock_v2.json"
)


def sha256(path: Path) -> str:
    digest = hashlib.sha256()
    with path.open("rb") as stream:
        for chunk in iter(lambda: stream.read(8 * 1024 * 1024), b""):
            digest.update(chunk)
    return digest.hexdigest()


def parse_sidecar(path: Path) -> tuple[str, str]:
    fields = path.read_text(encoding="ascii").strip().split()
    if len(fields) != 2:
        raise RuntimeError(f"Invalid SHA256 sidecar: {path}")
    return fields[0].lower(), fields[1]


def validate_complete_case_lock(caller: Path) -> tuple[dict, str]:
    if sha256(PROTOCOL) != PROTOCOL_SHA256:
        raise RuntimeError("Complete-case confirmation protocol hash mismatch.")
    sidecar_hash, sidecar_name = parse_sidecar(
        LOCK.with_name(LOCK.name + ".sha256")
    )
    observed_lock_hash = sha256(LOCK)
    if sidecar_name != LOCK.name or sidecar_hash != observed_lock_hash:
        raise RuntimeError("Complete-case implementation lock sidecar mismatch.")
    payload = json.loads(LOCK.read_text(encoding="utf-8"))
    if (
        payload["status"]
        != "locked_after_label_blind_QC_and_before_any_confirmation_attention_score"
        or payload["protocol_sha256"] != PROTOCOL_SHA256
    ):
        raise RuntimeError("Complete-case implementation lock status mismatch.")
    relative = caller.resolve().relative_to(ROOT.resolve()).as_posix()
    expected = payload["source_sha256"].get(relative)
    if expected is None or sha256(caller) != expected:
        raise RuntimeError(f"Caller is not complete-case locked: {relative}")
    for path_text, expected_hash in payload["source_sha256"].items():
        if sha256(ROOT / path_text) != expected_hash:
            raise RuntimeError(f"Complete-case source mismatch: {path_text}")
    for name, record in payload["frozen_inputs"].items():
        if sha256(ROOT / record["path"]) != record["sha256"]:
            raise RuntimeError(f"Complete-case frozen input mismatch: {name}")
    return payload, observed_lock_hash
```
