## Supplementary material for "Falsifiable substitution tests reveal task-structured neural evidence for auditory attention": Article_Analysis_Source_Code: kite_sat_confirmation_lock_utils_v1.py

```
"""Dynamic source/hash validation for the frozen KITE-SAT confirmation."""

from __future__ import annotations

import hashlib
import json
from pathlib import Path


ROOT = Path(__file__).resolve().parents[1]
PROTOCOL = ROOT / "docs" / "kite_sat_akb_confirmation_protocol_v1.json"
PROTOCOL_HASH = "36f8a83a3a6b80b7e2123706707a7dea6af43fc6cc41448b3e6483efb5b46847"
LOCK = ROOT / "docs" / "kite_sat_akb_confirmation_implementation_lock_v1.json"


def sha256(path: Path) -> str:
    digest = hashlib.sha256()
    with path.open("rb") as stream:
        for chunk in iter(lambda: stream.read(8 * 1024 * 1024), b""):
            digest.update(chunk)
    return digest.hexdigest()


def parse_sidecar(path: Path) -> tuple[str, str]:
    fields = path.read_text(encoding="ascii").strip().split()
    if len(fields) != 2:
        raise RuntimeError(f"Invalid SHA256 sidecar: {path}")
    return fields[0].lower(), fields[1]


def validate_confirmation_lock(caller: Path) -> tuple[dict, str]:
    if sha256(PROTOCOL) != PROTOCOL_HASH:
        raise RuntimeError("KITE-SAT confirmation protocol hash mismatch.")
    sidecar_hash, sidecar_name = parse_sidecar(LOCK.with_name(LOCK.name + ".sha256"))
    observed_lock_hash = sha256(LOCK)
    if sidecar_name != LOCK.name or sidecar_hash != observed_lock_hash:
        raise RuntimeError("KITE-SAT confirmation implementation-lock sidecar mismatch.")
    payload = json.loads(LOCK.read_text(encoding="utf-8"))
    if payload.get("status") != "locked_before_any_confirmation_archive_download_or_numeric_read":
        raise RuntimeError("KITE-SAT confirmation lock has the wrong status.")
    if payload.get("protocol_sha256") != PROTOCOL_HASH:
        raise RuntimeError("KITE-SAT confirmation lock/protocol mismatch.")
    relative = caller.resolve().relative_to(ROOT.resolve()).as_posix()
    expected = payload["source_sha256"].get(relative)
    if expected is None or sha256(caller) != expected:
        raise RuntimeError(f"Caller is not bound by confirmation lock: {relative}")
    for path_text, expected_hash in payload["source_sha256"].items():
        if sha256(ROOT / path_text) != expected_hash:
            raise RuntimeError(f"Confirmation locked source mismatch: {path_text}")
    for name, record in payload["frozen_inputs"].items():
        if sha256(ROOT / record["path"]) != record["sha256"]:
            raise RuntimeError(f"Confirmation frozen input mismatch: {name}")
    return payload, observed_lock_hash
```
