## Supplementary material for "Falsifiable substitution tests reveal task-structured neural evidence for auditory attention": Article_Analysis_Source_Code: kite_sat_direction_delay_controls_v1.py

```
"""Fixed capacity-matched direction-delay transformations for AKB."""

from __future__ import annotations

import numpy as np


TRANSFORMS = tuple((angular_shift, delay_swap) for delay_swap in range(2) for angular_shift in range(8))


def transform_vector(
    values: np.ndarray,
    angular_shift: int,
    delay_swap: int,
) -> np.ndarray:
    """Assign source ((s-k) mod 8, b xor d) to output test cell (s,b)."""
    array = np.asarray(values, dtype=np.float64)
    if array.shape != (16,):
        raise ValueError("Direction-delay transformation expects 16 values.")
    if angular_shift not in range(8) or delay_swap not in (0, 1):
        raise ValueError("Invalid direction-delay transformation.")
    output = np.empty(16, dtype=np.float64)
    for delay_bin in range(2):
        for sector in range(8):
            source_sector = (sector - angular_shift) % 8
            source_delay = delay_bin ^ delay_swap
            output[sector + 8 * delay_bin] = array[
                source_sector + 8 * source_delay
            ]
    return output


def transformed_matrix(values: np.ndarray) -> np.ndarray:
    """Return identity first, then all 15 fixed nonidentity controls."""
    return np.row_stack(
        [transform_vector(values, shift, swap) for shift, swap in TRANSFORMS]
    )
```
