## Supplementary material for "Falsifiable substitution tests reveal task-structured neural evidence for auditory attention": Article_Analysis_Source_Code: kite_sat_geometry_locked_v1.py

```
"""Locked P16-drop2 broad transform for KITE-SAT method development."""

from __future__ import annotations

import numpy as np

from kite_sat_geometry_v1 import cell_delta_contributions


def p16_drop2_score(
    design: np.ndarray,
    model: dict[str, np.ndarray | float],
    left: np.ndarray,
    right: np.ndarray,
    p16_ids: np.ndarray,
) -> tuple[float, float, np.ndarray]:
    delta, global_margin = cell_delta_contributions(
        design, model, left, right, p16_ids
    )
    if len(delta) != 16:
        raise RuntimeError("Locked KITE-SAT P16 partition does not have 16 cells.")
    order = np.argsort(-np.abs(delta), kind="stable")
    retained = delta.copy()
    retained[order[:2]] = 0.0
    return float(np.sum(retained)), global_margin, order[:2].astype(np.int32)
```
