## Supplementary material for "Falsifiable substitution tests reveal task-structured neural evidence for auditory attention": Article_Analysis_Source_Code: kite_sat_geometry_v1.py

```
"""Polynomial-partition broad contributions on the KITE-SAT scalp-latency field."""

from __future__ import annotations

import numpy as np

from kite_sat_backward_v1 import pearson


CANDIDATE_ORDER = (
    "P8_drop1",
    "P8_clip75",
    "P16_drop2",
    "P16_clip75",
    "S16_drop2",
    "S16_clip75",
    "multiscale_clip75",
)


def partition_ids(xy: np.ndarray) -> dict[str, np.ndarray]:
    coordinates = np.asarray(xy, dtype=np.float64)
    if coordinates.shape != (253, 2):
        raise ValueError("KITE-SAT geometry expects 253 public coordinates.")
    x = np.tile(coordinates[:, 0], 6)
    y = np.tile(coordinates[:, 1], 6)
    lag_index = np.repeat(np.arange(6), 253)
    late = lag_index >= 3
    radius2_electrode = np.sum(coordinates * coordinates, axis=1)
    radius2 = np.tile(radius2_electrode, 6)
    median_radius2 = float(np.median(radius2_electrode))
    p8 = (x >= 0).astype(np.int32) + 2 * (y >= 0) + 4 * late
    p16 = p8 + 8 * (radius2 >= median_radius2)
    angle = np.arctan2(y, x)
    sector = (
        np.floor((angle + np.pi) / (2 * np.pi) * 8).astype(np.int32) % 8
    )
    s16 = sector + 8 * late
    output = {"P8": p8, "P16": p16, "S16": s16}
    expected = {"P8": 8, "P16": 16, "S16": 16}
    for name, ids in output.items():
        counts = np.bincount(ids, minlength=expected[name])
        if len(ids) != 1518 or len(counts) != expected[name] or np.any(counts == 0):
            raise RuntimeError(f"Degenerate KITE-SAT partition: {name}")
    return output


def cell_delta_contributions(
    design: np.ndarray,
    model: dict[str, np.ndarray | float],
    left: np.ndarray,
    right: np.ndarray,
    ids: np.ndarray,
) -> tuple[np.ndarray, float]:
    x = (
        np.asarray(design, dtype=np.float64) - np.asarray(model["x_mean"])
    ) / np.asarray(model["x_scale"])
    weighted = x * np.asarray(model["weights"])[None, :]
    global_reconstruction = np.sum(weighted, axis=1)
    global_centered = global_reconstruction - np.mean(global_reconstruction)
    left_centered = np.asarray(left, dtype=np.float64) - np.mean(left)
    right_centered = np.asarray(right, dtype=np.float64) - np.mean(right)
    denom_left = np.sqrt(
        np.sum(global_centered * global_centered)
        * np.sum(left_centered * left_centered)
    )
    denom_right = np.sqrt(
        np.sum(global_centered * global_centered)
        * np.sum(right_centered * right_centered)
    )
    cells = int(np.max(ids)) + 1
    delta = np.empty(cells, dtype=np.float64)
    for cell in range(cells):
        cell_reconstruction = np.sum(weighted[:, ids == cell], axis=1)
        cell_reconstruction -= np.mean(cell_reconstruction)
        contribution_left = float(
            np.sum(cell_reconstruction * left_centered) / max(denom_left, 1e-12)
        )
        contribution_right = float(
            np.sum(cell_reconstruction * right_centered) / max(denom_right, 1e-12)
        )
        delta[cell] = contribution_left - contribution_right
    global_margin = pearson(global_reconstruction, left) - pearson(
        global_reconstruction, right
    )
    if abs(float(np.sum(delta)) - global_margin) > 1e-10:
        raise RuntimeError("Cell contributions do not reproduce the global margin.")
    return delta, global_margin


def _drop(values: np.ndarray, count: int) -> float:
    transformed = np.asarray(values, dtype=np.float64).copy()
    order = np.argsort(-np.abs(transformed), kind="stable")
    transformed[order[:count]] = 0.0
    return float(np.sum(transformed))


def _clip75(values: np.ndarray) -> float:
    array = np.asarray(values, dtype=np.float64)
    cap = float(np.quantile(np.abs(array), 0.75))
    return float(np.sum(np.clip(array, -cap, cap)))


def geometry_scores(
    design: np.ndarray,
    model: dict[str, np.ndarray | float],
    left: np.ndarray,
    right: np.ndarray,
    partitions: dict[str, np.ndarray],
) -> dict[str, float]:
    p8, global_margin = cell_delta_contributions(
        design, model, left, right, partitions["P8"]
    )
    p16, check_p16 = cell_delta_contributions(
        design, model, left, right, partitions["P16"]
    )
    s16, check_s16 = cell_delta_contributions(
        design, model, left, right, partitions["S16"]
    )
    if (
        abs(global_margin - check_p16) > 1e-12
        or abs(global_margin - check_s16) > 1e-12
    ):
        raise RuntimeError("Partition global margins disagree.")
    scores = {
        "global": global_margin,
        "P8_drop1": _drop(p8, 1),
        "P8_clip75": _clip75(p8),
        "P16_drop2": _drop(p16, 2),
        "P16_clip75": _clip75(p16),
        "S16_drop2": _drop(s16, 2),
        "S16_clip75": _clip75(s16),
    }
    scores["multiscale_clip75"] = float(
        np.mean(
            [
                scores["P8_clip75"],
                scores["P16_clip75"],
                scores["S16_clip75"],
            ]
        )
    )
    return scores
```
