## Supplementary material for "Falsifiable substitution tests reveal task-structured neural evidence for auditory attention": Article_Analysis_Source_Code: kite_sat_locked_decoder_v1.py

```
"""Frozen single-ridge speech-envelope decoder for KITE-SAT development."""

from __future__ import annotations

import numpy as np
from scipy.linalg import cho_factor, cho_solve


RIDGE_RATIO = 1.0


def fit_locked_ridge(
    training_designs: list[np.ndarray],
    training_targets: list[np.ndarray],
) -> dict[str, np.ndarray | float]:
    x = np.vstack(training_designs).astype(np.float64, copy=False)
    y = np.concatenate(training_targets).astype(np.float64, copy=False)
    if x.ndim != 2 or x.shape[1] != 1518 or x.shape[0] != len(y):
        raise ValueError("Unexpected locked KITE-SAT training shape.")
    x_mean = np.mean(x, axis=0)
    x_scale = np.std(x, axis=0)
    x_scale = np.where(x_scale > 1e-12, x_scale, 1.0)
    y_mean = float(np.mean(y))
    standardized = (x - x_mean) / x_scale
    centered = y - y_mean
    gram = standardized.T @ standardized
    cross = standardized.T @ centered
    gram_energy = float(np.mean(np.diag(gram)))
    alpha = RIDGE_RATIO * gram_energy
    regularized = gram
    regularized.flat[:: regularized.shape[0] + 1] += alpha
    factor = cho_factor(regularized, lower=True, check_finite=False)
    weights = cho_solve(factor, cross, check_finite=False)
    if not np.isfinite(weights).all():
        raise RuntimeError("Locked KITE-SAT ridge fit is nonfinite.")
    return {
        "x_mean": x_mean,
        "x_scale": x_scale,
        "y_mean": y_mean,
        "weights": weights,
        "gram_energy": gram_energy,
        "alpha": alpha,
    }


def predict_locked(
    design: np.ndarray,
    model: dict[str, np.ndarray | float],
) -> np.ndarray:
    standardized = (
        np.asarray(design, dtype=np.float64) - np.asarray(model["x_mean"])
    ) / np.asarray(model["x_scale"])
    return standardized @ np.asarray(model["weights"]) + float(model["y_mean"])
```
