## Supplementary material for "Falsifiable substitution tests reveal task-structured neural evidence for auditory attention": Article_Analysis_Source_Code: kite_sat_mechanism_lock_utils_v1.py

```
"""Dynamic validation for the frozen AKB direction-delay mechanism analysis."""

from __future__ import annotations

import hashlib
import json
from pathlib import Path


ROOT = Path(__file__).resolve().parents[1]
PROTOCOL = ROOT / "docs" / "kite_sat_akb_direction_delay_mechanism_protocol_v1.json"
PROTOCOL_HASH = "0bf4cfed0219bc0dd14ae8d7810f7bf9b07afbe550ac8ccfafbf7125418e4f5c"
LOCK = ROOT / "docs" / "kite_sat_akb_direction_delay_mechanism_implementation_lock_v1.json"


def sha256(path: Path) -> str:
    digest = hashlib.sha256()
    with path.open("rb") as stream:
        for chunk in iter(lambda: stream.read(8 * 1024 * 1024), b""):
            digest.update(chunk)
    return digest.hexdigest()


def validate_mechanism_lock(caller: Path) -> tuple[dict, str]:
    if sha256(PROTOCOL) != PROTOCOL_HASH:
        raise RuntimeError("AKB mechanism protocol hash mismatch.")
    sidecar = LOCK.with_name(LOCK.name + ".sha256")
    fields = sidecar.read_text(encoding="ascii").strip().split()
    observed = sha256(LOCK)
    if len(fields) != 2 or fields[0].lower() != observed or fields[1] != LOCK.name:
        raise RuntimeError("AKB mechanism lock sidecar mismatch.")
    lock = json.loads(LOCK.read_text(encoding="utf-8"))
    if lock.get("status") != "locked_before_any_confirmation_or_control_score":
        raise RuntimeError("AKB mechanism lock status mismatch.")
    if lock.get("protocol_sha256") != PROTOCOL_HASH:
        raise RuntimeError("AKB mechanism lock/protocol mismatch.")
    relative = caller.resolve().relative_to(ROOT.resolve()).as_posix()
    if lock["source_sha256"].get(relative) != sha256(caller):
        raise RuntimeError(f"Caller not bound by AKB mechanism lock: {relative}")
    for path_text, expected in lock["source_sha256"].items():
        if sha256(ROOT / path_text) != expected:
            raise RuntimeError(f"AKB mechanism source mismatch: {path_text}")
    for name, record in lock["frozen_inputs"].items():
        if sha256(ROOT / record["path"]) != record["sha256"]:
            raise RuntimeError(f"AKB mechanism frozen input mismatch: {name}")
    return lock, observed
```
