## Supplementary material for "Falsifiable substitution tests reveal task-structured neural evidence for auditory attention": Article_Analysis_Source_Code: kite_sat_patch_erasure_v1.py

```
"""Outcome-agnostic contiguous sensor-erasure primitives for KITE-SAT."""

from __future__ import annotations

import numpy as np


ANCHORS = ("TP7H", "PO3", "PO4", "TP8H", "FT8H", "AF4", "AF3", "FT7H")
PATCH_SIZE = 51


def fixed_patch_masks(
    labels: tuple[str, ...],
    coordinates: np.ndarray,
) -> tuple[np.ndarray, tuple[tuple[str, ...], ...]]:
    """Return the eight locked nearest-neighbor scalp patches."""
    xy = np.asarray(coordinates, dtype=np.float64)
    if len(labels) != 253 or xy.shape != (253, 2) or len(set(labels)) != 253:
        raise ValueError("Expected 253 unique public-coordinate electrodes.")
    masks = np.zeros((len(ANCHORS), 253), dtype=bool)
    members: list[tuple[str, ...]] = []
    for patch_index, anchor in enumerate(ANCHORS):
        if anchor not in labels:
            raise RuntimeError(f"Locked patch anchor is absent: {anchor}")
        anchor_index = labels.index(anchor)
        distance2 = np.sum((xy - xy[anchor_index]) ** 2, axis=1)
        # lexsort makes public coordinate-file order the exact tie breaker.
        order = np.lexsort((np.arange(len(labels)), distance2))
        selected = order[:PATCH_SIZE]
        masks[patch_index, selected] = True
        members.append(tuple(labels[index] for index in selected))
    if not np.all(np.sum(masks, axis=1) == PATCH_SIZE):
        raise RuntimeError("A fixed KITE-SAT patch has the wrong size.")
    if not all(masks[index, labels.index(anchor)] for index, anchor in enumerate(ANCHORS)):
        raise RuntimeError("A fixed patch does not contain its anchor.")
    return masks, tuple(members)


def erased_cell_delta_contributions(
    design: np.ndarray,
    model: dict[str, np.ndarray | float],
    left: np.ndarray,
    right: np.ndarray,
    ids: np.ndarray,
    removed_sensors: np.ndarray,
) -> tuple[np.ndarray, np.ndarray]:
    """Recompute S16 cell contributions after each locked sensor patch.

    Missing features are set to their training-model means, hence to zero in
    standardized coordinates.  Outputs are patch x cell deltas and patch
    global margins.
    """
    x = np.asarray(design, dtype=np.float64)
    labels = np.asarray(ids, dtype=np.int64)
    removed = np.asarray(removed_sensors, dtype=bool)
    if x.shape != (595, 1518) or labels.shape != (1518,):
        raise ValueError("Unexpected KITE-SAT design or cell-ID shape.")
    if removed.ndim != 2 or removed.shape[1] != 253:
        raise ValueError("Sensor-erasure masks must be patch x 253.")
    if np.min(labels) != 0 or np.max(labels) != 15:
        raise ValueError("Patch erasure expects the frozen 16-cell S16 partition.")
    standardized = (
        x - np.asarray(model["x_mean"], dtype=np.float64)
    ) / np.asarray(model["x_scale"], dtype=np.float64)
    weighted = standardized * np.asarray(model["weights"], dtype=np.float64)[None, :]
    by_lag_sensor = weighted.reshape(595, 6, 253)
    by_delay_sensor = np.stack(
        (
            np.sum(by_lag_sensor[:, :3, :], axis=1),
            np.sum(by_lag_sensor[:, 3:, :], axis=1),
        ),
        axis=1,
    )
    sector = labels[:253]
    if not np.array_equal(labels, np.tile(sector, 6) + np.repeat((0, 0, 0, 8, 8, 8), 253)):
        raise RuntimeError("S16 feature ordering is inconsistent with six lag blocks.")
    keep = (~removed).astype(np.float64)
    n_patches = len(removed)
    cell_reconstruction = np.zeros((n_patches, 595, 16), dtype=np.float64)
    for delay_bin in range(2):
        for angle_sector in range(8):
            sensor_mask = sector == angle_sector
            cell = angle_sector + 8 * delay_bin
            cell_reconstruction[:, :, cell] = (
                by_delay_sensor[:, delay_bin, sensor_mask]
                @ keep[:, sensor_mask].T
            ).T
    global_reconstruction = np.sum(cell_reconstruction, axis=2)
    global_centered = global_reconstruction - np.mean(
        global_reconstruction, axis=1, keepdims=True
    )
    cell_centered = cell_reconstruction - np.mean(
        cell_reconstruction, axis=1, keepdims=True
    )
    left_centered = np.asarray(left, dtype=np.float64) - np.mean(left)
    right_centered = np.asarray(right, dtype=np.float64) - np.mean(right)
    global_energy = np.sum(global_centered * global_centered, axis=1)
    denominator_left = np.sqrt(
        global_energy * np.sum(left_centered * left_centered)
    )
    denominator_right = np.sqrt(
        global_energy * np.sum(right_centered * right_centered)
    )
    contribution_left = np.einsum(
        "pnc,n->pc", cell_centered, left_centered
    ) / np.maximum(denominator_left[:, None], 1e-12)
    contribution_right = np.einsum(
        "pnc,n->pc", cell_centered, right_centered
    ) / np.maximum(denominator_right[:, None], 1e-12)
    delta = contribution_left - contribution_right
    global_margin = (
        np.einsum("pn,n->p", global_centered, left_centered)
        / np.maximum(denominator_left, 1e-12)
        - np.einsum("pn,n->p", global_centered, right_centered)
        / np.maximum(denominator_right, 1e-12)
    )
    if not np.allclose(np.sum(delta, axis=1), global_margin, atol=1e-10, rtol=0.0):
        raise RuntimeError("Erased S16 cells do not reproduce the global margins.")
    if not np.isfinite(delta).all() or not np.isfinite(global_margin).all():
        raise RuntimeError("Nonfinite erased KITE-SAT score.")
    return delta, global_margin
```
