## Supplementary material for "Falsifiable substitution tests reveal task-structured neural evidence for auditory attention": Article_Analysis_Source_Code: kite_sat_sensor_density_lock_utils_v1.py

```
"""Hash validation for the post-confirmation sensor-density mechanism."""

from __future__ import annotations

import hashlib
import json
from pathlib import Path


ROOT = Path(__file__).resolve().parents[1]
PROTOCOL = ROOT / "docs" / "kite_sat_sensor_density_mechanism_protocol_v1.json"
PROTOCOL_SHA256 = "db704901422b45686819e294afdd60ac4ffafbc9b7c75a2f871ab1049a740f86"
LOCK = ROOT / "docs" / "kite_sat_sensor_density_implementation_lock_v1.json"


def sha256(path: Path) -> str:
    digest = hashlib.sha256()
    with path.open("rb") as stream:
        for chunk in iter(lambda: stream.read(8 * 1024 * 1024), b""):
            digest.update(chunk)
    return digest.hexdigest()


def validate_sensor_density_lock(caller: Path) -> tuple[dict, str]:
    if sha256(PROTOCOL) != PROTOCOL_SHA256:
        raise RuntimeError("Sensor-density protocol hash mismatch.")
    fields = LOCK.with_name(LOCK.name + ".sha256").read_text(
        encoding="ascii"
    ).strip().split()
    observed_lock = sha256(LOCK)
    if len(fields) != 2 or fields[0].lower() != observed_lock or fields[1] != LOCK.name:
        raise RuntimeError("Sensor-density implementation lock sidecar mismatch.")
    payload = json.loads(LOCK.read_text(encoding="utf-8"))
    if (
        payload["status"]
        != "locked_after_primary_confirmation_and_before_any_sensor_budget_decoder"
        or payload["protocol_sha256"] != PROTOCOL_SHA256
    ):
        raise RuntimeError("Sensor-density implementation lock status mismatch.")
    relative = caller.resolve().relative_to(ROOT.resolve()).as_posix()
    expected = payload["source_sha256"].get(relative)
    if expected is None or sha256(caller) != expected:
        raise RuntimeError(f"Caller is not sensor-density locked: {relative}")
    for path_text, expected_hash in payload["source_sha256"].items():
        if sha256(ROOT / path_text) != expected_hash:
            raise RuntimeError(f"Sensor-density source mismatch: {path_text}")
    for name, record in payload["frozen_inputs"].items():
        if sha256(ROOT / record["path"]) != record["sha256"]:
            raise RuntimeError(f"Sensor-density frozen input mismatch: {name}")
    return payload, observed_lock
```
