## Supplementary material for "Falsifiable substitution tests reveal task-structured neural evidence for auditory attention": Article_Analysis_Source_Code: kite_st_hd_core_v1.py

```
"""Core signal and scalp-time geometry primitives for KITE-ST-HD-v1.

The functions are outcome-agnostic and are exercised on synthetic constructs
before any development EEG is available.
"""

from __future__ import annotations

import csv
import hashlib
import math
from dataclasses import dataclass
from pathlib import Path

import numpy as np
from scipy.signal import get_window


LAGS_AT_100HZ = np.asarray([2, 4, 6, 8, 10, 12], dtype=np.int64)
PHASE_FREQUENCIES_HZ = np.arange(4.0, 15.0, 1.0)


def stable_seed(*parts: object) -> int:
    text = "|".join(str(part) for part in parts)
    return int.from_bytes(hashlib.sha256(text.encode("utf-8")).digest()[:8], "little")


def normalize_label(label: str) -> str:
    return label.strip().upper()


@dataclass(frozen=True)
class ScalpGeometry:
    labels: tuple[str, ...]
    xy: np.ndarray
    primary: np.ndarray
    frontal: np.ndarray


def load_public_geometry(path: Path) -> ScalpGeometry:
    labels: list[str] = []
    xy: list[tuple[float, float]] = []
    with path.open("r", encoding="utf-8-sig", newline="") as stream:
        for row in csv.DictReader(stream):
            theta = float(row["Theta"])
            phi = float(row["Phi"])
            if not math.isfinite(theta) or not math.isfinite(phi):
                continue
            theta_rad = math.radians(theta)
            phi_rad = math.radians(phi)
            labels.append(normalize_label(row["Channel Name"]))
            xy.append(
                (
                    theta_rad * math.cos(phi_rad) / math.pi,
                    theta_rad * math.sin(phi_rad) / math.pi,
                )
            )
    coordinates = np.asarray(xy, dtype=np.float64)
    if len(labels) != 253 or coordinates.shape != (253, 2):
        raise RuntimeError("Expected 253 finite public scalp coordinates.")
    if len(set(labels)) != 253:
        raise RuntimeError("Public coordinate labels are not unique.")
    frontal = np.flatnonzero(coordinates[:, 1] > 0.30)
    primary = np.flatnonzero(coordinates[:, 1] <= 0.30)
    if len(frontal) != 31 or len(primary) != 222:
        raise RuntimeError("Locked frontal/primary partition does not reproduce.")
    checks = {label: coordinates[labels.index(label)] for label in ("CZ", "C3", "C4", "FZ", "OZ")}
    if (
        not np.allclose(checks["CZ"], 0.0)
        or not checks["C3"][0] < 0 < checks["C4"][0]
        or not checks["FZ"][1] > 0 > checks["OZ"][1]
    ):
        raise RuntimeError("Public coordinate orientation checks failed.")
    return ScalpGeometry(tuple(labels), coordinates, primary, frontal)


def all_unordered_edges(n_nodes: int) -> np.ndarray:
    return np.column_stack(np.triu_indices(n_nodes, k=1)).astype(np.int32)


def standardize_samples(block: np.ndarray) -> np.ndarray:
    x = np.asarray(block, dtype=np.float64)
    if x.ndim != 2 or len(x) < 20:
        raise ValueError("A signal block must be samples x channels.")
    x = x - np.mean(x, axis=0, keepdims=True)
    scale = np.sqrt(np.mean(x * x, axis=0))
    scale = np.where(scale > 1e-12, scale, 1.0)
    return x / scale


def lag_antisymmetric(
    block: np.ndarray,
    lags: np.ndarray = LAGS_AT_100HZ,
) -> tuple[np.ndarray, np.ndarray]:
    """Return average and per-lag directed antisymmetric association."""
    x = standardize_samples(block)
    lags = np.asarray(lags, dtype=np.int64)
    maximum = int(np.max(lags))
    base = x[: len(x) - maximum]
    per_lag = []
    for lag in lags:
        future = x[int(lag) : int(lag) + len(base)]
        cross = base.T @ future / len(base)
        per_lag.append(0.5 * (cross - cross.T))
    stacked = np.asarray(per_lag)
    return np.mean(stacked, axis=0), stacked


def _windowed_fft(
    block: np.ndarray,
    sampling_hz: float,
    segment_seconds: float = 2.0,
    step_seconds: float = 1.0,
) -> tuple[np.ndarray, np.ndarray]:
    x = standardize_samples(block)
    width = int(round(segment_seconds * sampling_hz))
    step = int(round(step_seconds * sampling_hz))
    if width < 8 or step < 1 or len(x) < width:
        raise ValueError("Block is too short for phase-slope estimation.")
    starts = np.arange(0, len(x) - width + 1, step, dtype=np.int64)
    window = get_window("hann", width, fftbins=True).astype(np.float64)
    spectra = []
    for start in starts:
        spectra.append(
            np.fft.rfft(x[start : start + width] * window[:, None], axis=0)
        )
    frequencies = np.fft.rfftfreq(width, d=1.0 / sampling_hz)
    return np.asarray(spectra), frequencies


def phase_slope_transport(
    block: np.ndarray,
    sampling_hz: float = 100.0,
    frequencies_hz: np.ndarray = PHASE_FREQUENCIES_HZ,
) -> np.ndarray:
    """Fixed-band phase-slope index matrix; output is antisymmetric."""
    spectra, frequencies = _windowed_fft(block, sampling_hz)
    bins = np.asarray(
        [int(np.argmin(np.abs(frequencies - value))) for value in frequencies_hz],
        dtype=np.int64,
    )
    if len(np.unique(bins)) != len(bins):
        raise RuntimeError("Requested phase-slope frequencies do not map uniquely.")
    coherencies = []
    for index in bins:
        z = spectra[:, index, :]
        spectrum = z.conj().T @ z / len(z)
        power = np.maximum(np.real(np.diag(spectrum)), 1e-12)
        denominator = np.sqrt(power[:, None] * power[None, :])
        coherencies.append(spectrum / denominator)
    coherencies = np.asarray(coherencies)
    psi = np.zeros(coherencies.shape[1:], dtype=np.float64)
    for first, second in zip(coherencies[:-1], coherencies[1:]):
        psi += np.imag(np.conj(first) * second)
    return 0.5 * (psi - psi.T)


def upper_edge_values(matrix: np.ndarray, edges: np.ndarray) -> np.ndarray:
    values = np.asarray(matrix)
    return values[edges[:, 0], edges[:, 1]]


def paired_bounded_effect(left: np.ndarray, right: np.ndarray) -> np.ndarray:
    difference = np.asarray(left, dtype=np.float64) - np.asarray(right, dtype=np.float64)
    mean = np.mean(difference, axis=0)
    rms = np.sqrt(np.mean(difference * difference, axis=0))
    return mean / (rms + 1e-12)


def cross_story_edge_score(part1: np.ndarray, part2: np.ndarray) -> np.ndarray:
    first = np.asarray(part1, dtype=np.float64)
    second = np.asarray(part2, dtype=np.float64)
    if first.shape != second.shape or first.ndim != 2:
        raise ValueError("Cross-story arrays must share participant x edge shape.")
    n = len(first)
    if n < 2:
        raise ValueError("At least two participants are required.")
    return (
        first.sum(axis=0) * second.sum(axis=0)
        + second.sum(axis=0) * first.sum(axis=0)
        - 2.0 * np.sum(first * second, axis=0)
    ) / (2.0 * n * (n - 1))


def select_edges(score: np.ndarray, count: int) -> np.ndarray:
    score = np.asarray(score, dtype=np.float64)
    if not 0 < count <= len(score):
        raise ValueError("Invalid selected edge count.")
    order = np.lexsort((np.arange(len(score)), -score))
    return order[:count].astype(np.int32)


def all_sign_patterns(n_participants: int) -> np.ndarray:
    integers = np.arange(1 << n_participants, dtype=np.uint32)
    bits = (integers[:, None] >> np.arange(n_participants)) & 1
    return np.where(bits == 0, 1.0, -1.0)


def exact_upper_p(null: np.ndarray, observed: float) -> float:
    values = np.asarray(null, dtype=np.float64)
    return float(np.mean(values >= observed - 1e-15))


def candidate_signflip_max_statistics(
    candidate_splits: dict[str, tuple[np.ndarray, np.ndarray]],
    selected_count: int,
) -> tuple[dict[str, float], dict[str, np.ndarray], np.ndarray]:
    names = tuple(candidate_splits)
    n = len(next(iter(candidate_splits.values()))[0])
    patterns = all_sign_patterns(n)
    observed: dict[str, float] = {}
    nulls: dict[str, np.ndarray] = {}
    for name in names:
        part1, part2 = candidate_splits[name]
        local = []
        for signs in patterns:
            score = cross_story_edge_score(
                part1 * signs[:, None],
                part2 * signs[:, None],
            )
            selected = select_edges(score, selected_count)
            local.append(float(np.mean(score[selected])))
        nulls[name] = np.asarray(local)
        observed[name] = float(local[0])
    maximum = np.max(np.column_stack([nulls[name] for name in names]), axis=1)
    return observed, nulls, maximum


def tube_segments(
    xy: np.ndarray,
    edges: np.ndarray,
    signs: np.ndarray,
    lag_seconds: np.ndarray | float,
    time_scale_seconds: float,
) -> tuple[np.ndarray, np.ndarray, np.ndarray]:
    """Create oriented finite segments in R2_scalp x R_time."""
    coordinates = np.asarray(xy, dtype=np.float64)
    edge_array = np.asarray(edges, dtype=np.int64)
    sign_array = np.where(np.asarray(signs) >= 0, 1, -1)
    lag = np.broadcast_to(np.asarray(lag_seconds, dtype=np.float64), len(edge_array))
    if time_scale_seconds <= 0 or np.any(lag <= 0):
        raise ValueError("Positive lag and time scale are required.")
    source = np.where(sign_array[:, None] > 0, edge_array[:, :1], edge_array[:, 1:])
    target = np.where(sign_array[:, None] > 0, edge_array[:, 1:], edge_array[:, :1])
    start = np.column_stack(
        [
            coordinates[source[:, 0], 0],
            coordinates[source[:, 0], 1],
            np.zeros(len(edge_array)),
        ]
    )
    stop = np.column_stack(
        [
            coordinates[target[:, 0], 0],
            coordinates[target[:, 0], 1],
            lag / time_scale_seconds,
        ]
    )
    vector = stop - start
    length = np.linalg.norm(vector, axis=1)
    direction = vector / np.maximum(length[:, None], 1e-12)
    return start, stop, direction


def direction_diagnostics(
    direction: np.ndarray,
    edges: np.ndarray,
    n_nodes: int,
    seed: int,
    triple_samples: int = 20000,
) -> dict[str, float]:
    directions = np.asarray(direction, dtype=np.float64)
    moment = directions.T @ directions / len(directions)
    eigenvalues = np.linalg.eigvalsh(moment)
    eigenvalues = np.maximum(eigenvalues, 0.0)
    probabilities = eigenvalues / np.sum(eigenvalues)
    effective_rank = float(np.exp(-np.sum(probabilities * np.log(probabilities + 1e-15))))
    rng = np.random.default_rng(seed)
    triples = rng.integers(0, len(directions), size=(triple_samples, 3))
    determinants = np.abs(
        np.einsum(
            "ij,ij->i",
            directions[triples[:, 0]],
            np.cross(directions[triples[:, 1]], directions[triples[:, 2]]),
        )
    )
    degree = np.bincount(np.asarray(edges).ravel(), minlength=n_nodes).astype(float)
    degree_probability = degree / np.sum(degree)
    return {
        "direction_minimum_eigenvalue": float(eigenvalues[0]),
        "direction_effective_rank": effective_rank,
        "trilinear_determinant_mean": float(np.mean(determinants)),
        "trilinear_determinant_q75": float(np.quantile(determinants, 0.75)),
        "sensor_degree_hhi": float(np.sum(degree_probability * degree_probability)),
        "maximum_sensor_degree_fraction": float(np.max(degree) / len(edges)),
    }


def point_segment_squared_distance(
    points: np.ndarray,
    start: np.ndarray,
    stop: np.ndarray,
) -> np.ndarray:
    vector = stop - start
    denominator = float(np.dot(vector, vector))
    if denominator <= 1e-15:
        return np.sum((points - start) ** 2, axis=1)
    projection = np.clip(((points - start) @ vector) / denominator, 0.0, 1.0)
    closest = start + projection[:, None] * vector
    return np.sum((points - closest) ** 2, axis=1)


def fixed_integration_points(count: int, seed: int) -> np.ndarray:
    rng = np.random.default_rng(seed)
    return np.column_stack(
        [
            rng.uniform(-0.68, 0.68, size=count),
            rng.uniform(-0.68, 0.55, size=count),
            rng.uniform(-0.10, 1.10, size=count),
        ]
    )


def tube_occupation_diagnostics(
    start: np.ndarray,
    stop: np.ndarray,
    radius: float,
    integration_points: np.ndarray,
) -> dict[str, float]:
    if radius <= 0:
        raise ValueError("Tube radius must be positive.")
    points = np.asarray(integration_points, dtype=np.float64)
    multiplicity = np.zeros(len(points), dtype=np.int32)
    for first, second in zip(start, stop):
        multiplicity += (
            point_segment_squared_distance(points, first, second) <= radius * radius
        )
    occupied = multiplicity > 0
    if not np.any(occupied):
        raise RuntimeError("No integration point is occupied by the tube family.")
    occupied_multiplicity = multiplicity[occupied]
    return {
        "union_fraction": float(np.mean(occupied)),
        "occupied_multiplicity_median": float(np.median(occupied_multiplicity)),
        "occupied_multiplicity_q90": float(np.quantile(occupied_multiplicity, 0.90)),
        "occupied_multiplicity_maximum": int(np.max(occupied_multiplicity)),
        "mean_multiplicity_all_points": float(np.mean(multiplicity)),
    }


def covering_slope(
    start: np.ndarray,
    stop: np.ndarray,
    scales: tuple[float, ...] = (0.20, 0.10, 0.05),
    points_per_segment: int = 41,
) -> dict[str, object]:
    sampled = []
    fractions = np.linspace(0.0, 1.0, points_per_segment)
    for first, second in zip(start, stop):
        sampled.append(first + fractions[:, None] * (second - first))
    points = np.row_stack(sampled)
    counts = []
    for scale in scales:
        cells = np.floor(points / scale).astype(np.int64)
        counts.append(len(np.unique(cells, axis=0)))
    x = np.log(1.0 / np.asarray(scales))
    y = np.log(np.asarray(counts, dtype=np.float64))
    slope = float(np.polyfit(x, y, 1)[0])
    return {
        "scales": list(scales),
        "occupied_cover_counts": counts,
        "log_covering_slope": slope,
    }
```
