## Supplementary material for "Falsifiable substitution tests reveal task-structured neural evidence for auditory attention": Article_Analysis_Source_Code: kite_st_hd_features_v1.py

```
"""Frozen feature contract for KITE-ST-HD development and confirmation.

All functions operate on one recording at a time.  No condition label,
participant split, edge selection or group outcome enters this module.
"""

from __future__ import annotations

from dataclasses import dataclass

import numpy as np
from scipy.signal import get_window

from kite_st_hd_core_v1 import (
    LAGS_AT_100HZ,
    PHASE_FREQUENCIES_HZ,
    all_unordered_edges,
    lag_antisymmetric,
    standardize_samples,
    upper_edge_values,
)


SAMPLING_HZ = 100
BLOCK_SECONDS = 30
BLOCK_SAMPLES = SAMPLING_HZ * BLOCK_SECONDS
BLOCK_COUNT = 12
CLIP_STANDARD_DEVIATIONS = 8.0
FRONTAL_PC_COUNT = 5


@dataclass(frozen=True)
class FeatureGeometry:
    primary_indices: np.ndarray
    frontal_indices: np.ndarray
    primary_xy: np.ndarray
    frontal_xy: np.ndarray
    primary_edges: np.ndarray
    frontal_edges: np.ndarray
    posterior_left_indices: np.ndarray
    posterior_right_indices: np.ndarray


def feature_geometry(public_geometry) -> FeatureGeometry:
    primary = np.asarray(public_geometry.primary, dtype=np.int32)
    frontal = np.asarray(public_geometry.frontal, dtype=np.int32)
    xy = np.asarray(public_geometry.xy, dtype=np.float64)
    posterior_left = np.flatnonzero((xy[:, 1] < -0.15) & (xy[:, 0] < -0.10))
    posterior_right = np.flatnonzero((xy[:, 1] < -0.15) & (xy[:, 0] > 0.10))
    if len(primary) != 222 or len(frontal) != 31:
        raise RuntimeError("Frozen primary/frontal geometry does not reproduce.")
    if len(posterior_left) < 10 or len(posterior_right) < 10:
        raise RuntimeError("Frozen posterior alpha regions are unexpectedly small.")
    return FeatureGeometry(
        primary_indices=primary,
        frontal_indices=frontal,
        primary_xy=xy[primary],
        frontal_xy=xy[frontal],
        primary_edges=all_unordered_edges(len(primary)),
        frontal_edges=all_unordered_edges(len(frontal)),
        posterior_left_indices=posterior_left.astype(np.int32),
        posterior_right_indices=posterior_right.astype(np.int32),
    )


def robust_block(block: np.ndarray) -> np.ndarray:
    standardized = standardize_samples(block)
    clipped = np.clip(
        standardized,
        -CLIP_STANDARD_DEVIATIONS,
        CLIP_STANDARD_DEVIATIONS,
    )
    return standardize_samples(clipped)


def regress_frontal_pcs(primary: np.ndarray, frontal: np.ndarray) -> np.ndarray:
    target = robust_block(primary)
    nuisance = robust_block(frontal)
    u, singular, _ = np.linalg.svd(nuisance, full_matrices=False)
    if len(singular) < FRONTAL_PC_COUNT or singular[FRONTAL_PC_COUNT - 1] <= 1e-10:
        raise RuntimeError("Frontal nuisance PCs are degenerate.")
    scores = u[:, :FRONTAL_PC_COUNT]
    residual = target - scores @ (scores.T @ target)
    return robust_block(residual)


def _windowed_coherencies(block: np.ndarray) -> tuple[np.ndarray, float]:
    x = robust_block(block)
    width = 2 * SAMPLING_HZ
    step = SAMPLING_HZ
    starts = np.arange(0, len(x) - width + 1, step, dtype=np.int64)
    if len(starts) != 29:
        raise RuntimeError("Frozen phase-slope window count does not reproduce.")
    window = get_window("hann", width, fftbins=True).astype(np.float64)
    spectra = np.asarray(
        [
            np.fft.rfft(x[start : start + width] * window[:, None], axis=0)
            for start in starts
        ]
    )
    frequencies = np.fft.rfftfreq(width, d=1.0 / SAMPLING_HZ)
    bins = np.asarray(
        [
            int(np.argmin(np.abs(frequencies - value)))
            for value in PHASE_FREQUENCIES_HZ
        ],
        dtype=np.int32,
    )
    if len(np.unique(bins)) != len(PHASE_FREQUENCIES_HZ):
        raise RuntimeError("Frozen phase-slope bins are not unique.")
    coherencies = []
    for index in bins:
        z = spectra[:, index, :]
        spectrum = z.conj().T @ z / len(z)
        power = np.maximum(np.real(np.diag(spectrum)), 1e-12)
        coherencies.append(spectrum / np.sqrt(power[:, None] * power[None, :]))
    spacing = float(frequencies[bins[1]] - frequencies[bins[0]])
    if not np.allclose(np.diff(frequencies[bins]), spacing) or spacing != 1.0:
        raise RuntimeError("Frozen phase-frequency spacing does not reproduce.")
    return np.asarray(coherencies), spacing


def phase_slope_and_carrier_delay(
    block: np.ndarray,
) -> tuple[np.ndarray, np.ndarray]:
    """Return antisymmetric PSI and a label-blind absolute carrier delay.

    The carrier delay is the absolute circular-mean group delay of adjacent
    coherencies, truncated to the candidate's predeclared 20--120 ms support.
    It is used only as the tube's time coordinate and never as a decoder value.
    """
    coherencies, spacing = _windowed_coherencies(block)
    increments = np.conj(coherencies[:-1]) * coherencies[1:]
    psi = np.sum(np.imag(increments), axis=0)
    psi = 0.5 * (psi - psi.T)
    weights = np.abs(increments)
    unit = increments / np.maximum(weights, 1e-12)
    mean_unit = np.sum(weights * unit, axis=0) / np.maximum(
        np.sum(weights, axis=0), 1e-12
    )
    signed_delay = -np.angle(mean_unit) / (2.0 * np.pi * spacing)
    carrier = np.clip(np.abs(signed_delay), 0.020, 0.120)
    np.fill_diagonal(carrier, 0.020)
    return psi, carrier


def lag_features(
    block: np.ndarray,
    edges: np.ndarray,
) -> tuple[np.ndarray, np.ndarray]:
    average, per_lag = lag_antisymmetric(robust_block(block))
    return (
        upper_edge_values(average, edges),
        np.asarray(
            [upper_edge_values(matrix, edges) for matrix in per_lag],
            dtype=np.float64,
        ),
    )


def phase_features(
    block: np.ndarray,
    edges: np.ndarray,
) -> tuple[np.ndarray, np.ndarray]:
    matrix, delay = phase_slope_and_carrier_delay(block)
    return upper_edge_values(matrix, edges), upper_edge_values(delay, edges)


def symmetric_zero_lag(block: np.ndarray, edges: np.ndarray) -> np.ndarray:
    x = robust_block(block)
    covariance = x.T @ x / len(x)
    return upper_edge_values(covariance, edges)


def posterior_alpha_lateralization(
    block_all_scalp: np.ndarray,
    left_indices: np.ndarray,
    right_indices: np.ndarray,
) -> float:
    x = robust_block(block_all_scalp)
    window = get_window("hann", len(x), fftbins=True).astype(np.float64)
    spectrum = np.fft.rfft(x * window[:, None], axis=0)
    frequencies = np.fft.rfftfreq(len(x), d=1.0 / SAMPLING_HZ)
    bins = (frequencies >= 8.0) & (frequencies <= 13.0)
    power = np.mean(np.abs(spectrum[bins]) ** 2, axis=0)
    log_power = np.log(np.maximum(power, 1e-12))
    return float(np.mean(log_power[right_indices]) - np.mean(log_power[left_indices]))


def extract_recording_features(
    eeg: np.ndarray,
    geometry: FeatureGeometry,
) -> dict[str, np.ndarray]:
    values = np.asarray(eeg, dtype=np.float64)
    if values.shape != (BLOCK_COUNT * BLOCK_SAMPLES, 253):
        raise ValueError("Expected one 360-second, 253-channel recording.")
    output: dict[str, list] = {
        "A_primary": [],
        "A_primary_per_lag": [],
        "B_primary": [],
        "B_primary_carrier_delay_seconds": [],
        "A_frontal_pc_residual": [],
        "B_frontal_pc_residual": [],
        "A_frontal_only": [],
        "B_frontal_only": [],
        "A_time_reversed": [],
        "B_time_reversed": [],
        "zero_lag_symmetric_primary": [],
        "posterior_alpha_right_minus_left": [],
    }
    for block_index in range(BLOCK_COUNT):
        whole = values[
            block_index * BLOCK_SAMPLES : (block_index + 1) * BLOCK_SAMPLES
        ]
        primary = whole[:, geometry.primary_indices]
        frontal = whole[:, geometry.frontal_indices]
        a_primary, a_per_lag = lag_features(primary, geometry.primary_edges)
        b_primary, b_delay = phase_features(primary, geometry.primary_edges)
        residual = regress_frontal_pcs(primary, frontal)
        a_residual, _ = lag_features(residual, geometry.primary_edges)
        b_residual, _ = phase_features(residual, geometry.primary_edges)
        a_frontal, _ = lag_features(frontal, geometry.frontal_edges)
        b_frontal, _ = phase_features(frontal, geometry.frontal_edges)
        a_reversed, _ = lag_features(primary[::-1], geometry.primary_edges)
        b_reversed, _ = phase_features(primary[::-1], geometry.primary_edges)
        output["A_primary"].append(a_primary)
        output["A_primary_per_lag"].append(a_per_lag)
        output["B_primary"].append(b_primary)
        output["B_primary_carrier_delay_seconds"].append(b_delay)
        output["A_frontal_pc_residual"].append(a_residual)
        output["B_frontal_pc_residual"].append(b_residual)
        output["A_frontal_only"].append(a_frontal)
        output["B_frontal_only"].append(b_frontal)
        output["A_time_reversed"].append(a_reversed)
        output["B_time_reversed"].append(b_reversed)
        output["zero_lag_symmetric_primary"].append(
            symmetric_zero_lag(primary, geometry.primary_edges)
        )
        output["posterior_alpha_right_minus_left"].append(
            posterior_alpha_lateralization(
                whole,
                geometry.posterior_left_indices,
                geometry.posterior_right_indices,
            )
        )
    arrays = {
        name: np.asarray(items, dtype=np.float32)
        for name, items in output.items()
    }
    expected = {
        "A_primary": (12, 24531),
        "A_primary_per_lag": (12, 6, 24531),
        "B_primary": (12, 24531),
        "B_primary_carrier_delay_seconds": (12, 24531),
        "A_frontal_pc_residual": (12, 24531),
        "B_frontal_pc_residual": (12, 24531),
        "A_frontal_only": (12, 465),
        "B_frontal_only": (12, 465),
        "A_time_reversed": (12, 24531),
        "B_time_reversed": (12, 24531),
        "zero_lag_symmetric_primary": (12, 24531),
        "posterior_alpha_right_minus_left": (12,),
    }
    for name, shape in expected.items():
        if arrays[name].shape != shape or not np.isfinite(arrays[name]).all():
            raise RuntimeError(f"Feature contract failed for {name}.")
    if np.any(arrays["B_primary_carrier_delay_seconds"] < 0.020 - 1e-7):
        raise RuntimeError("Phase carrier delay is below its frozen support.")
    if np.any(arrays["B_primary_carrier_delay_seconds"] > 0.120 + 1e-7):
        raise RuntimeError("Phase carrier delay is above its frozen support.")
    return arrays


def pooled_carrier_lags(
    condition_features: list[dict[str, np.ndarray]],
) -> tuple[np.ndarray, np.ndarray]:
    """Pool all four recordings without labels to bind one lag per edge."""
    if len(condition_features) != 4:
        raise ValueError("Carrier-lag pooling requires all four recordings.")
    per_lag = np.concatenate(
        [item["A_primary_per_lag"] for item in condition_features], axis=0
    )
    strength = np.mean(per_lag.astype(np.float64) ** 2, axis=0)
    selected_lag = np.argmax(strength, axis=0)
    lag_seconds = LAGS_AT_100HZ[selected_lag].astype(np.float64) / SAMPLING_HZ
    phase_delays = np.concatenate(
        [
            item["B_primary_carrier_delay_seconds"]
            for item in condition_features
        ],
        axis=0,
    )
    phase_lag_seconds = np.median(phase_delays.astype(np.float64), axis=0)
    if lag_seconds.shape != (24531,) or phase_lag_seconds.shape != (24531,):
        raise RuntimeError("Pooled carrier-lag contract failed.")
    return lag_seconds.astype(np.float32), phase_lag_seconds.astype(np.float32)
```
