## Supplementary material for "Falsifiable substitution tests reveal task-structured neural evidence for auditory attention": Article_Analysis_Source_Code: kite_st_hd_geometry_v1.py

```
"""Frozen finite-tube geometry and matched structural nulls for KITE-ST-HD."""

from __future__ import annotations

from collections import defaultdict

import numpy as np

from kite_st_hd_core_v1 import (
    covering_slope,
    direction_diagnostics,
    fixed_integration_points,
    stable_seed,
    tube_occupation_diagnostics,
    tube_segments,
)


TIME_SCALE_SECONDS = 0.120
RADII = (0.025, 0.035, 0.050)
PRIMARY_RADIUS = 0.035
INTEGRATION_POINT_COUNT = 12000
NESTED_INTEGRATION_POINT_COUNT = 2000
MATCHED_FAMILY_COUNT = 1000
SELECTED_EDGE_COUNT = 200
TRIPLE_SAMPLES = 5000
COVERING_SCALES = (0.20, 0.10, 0.05)
SEGMENT_SHADE_POINTS = 21
CONVEX_BALL_RADIUS = 0.25
CONVEX_SLAB_WIDTH = 0.20


def spatial_length_deciles(xy: np.ndarray, edges: np.ndarray) -> tuple[np.ndarray, np.ndarray]:
    coordinates = np.asarray(xy, dtype=np.float64)
    pairs = np.asarray(edges, dtype=np.int64)
    lengths = np.linalg.norm(coordinates[pairs[:, 1]] - coordinates[pairs[:, 0]], axis=1)
    boundaries = np.quantile(lengths, np.arange(1, 10) / 10.0)
    bins = np.digitize(lengths, boundaries, right=True).astype(np.int8)
    return lengths, bins


def carrier_lag_bins(lag_seconds: np.ndarray) -> np.ndarray:
    lag = np.asarray(lag_seconds, dtype=np.float64)
    if np.any(lag < 0.020 - 1e-9) or np.any(lag > 0.120 + 1e-9):
        raise ValueError("Carrier lags are outside 20--120 ms.")
    return np.digitize(lag, [0.04, 0.06, 0.08, 0.10], right=True).astype(np.int8)


def _shade_points(start: np.ndarray, stop: np.ndarray) -> np.ndarray:
    fraction = np.linspace(0.0, 1.0, SEGMENT_SHADE_POINTS)
    return np.row_stack(
        [first + fraction[:, None] * (second - first) for first, second in zip(start, stop)]
    )


def convex_concentration(start: np.ndarray, stop: np.ndarray) -> dict[str, float]:
    points = _shade_points(start, stop)
    x_centers = (-0.40, 0.0, 0.40)
    y_centers = (-0.40, 0.0, 0.40)
    z_centers = (0.0, 0.5, 1.0)
    centers = np.asarray(
        [(x, y, z) for x in x_centers for y in y_centers for z in z_centers],
        dtype=np.float64,
    )
    squared = np.sum((points[:, None, :] - centers[None, :, :]) ** 2, axis=2)
    ball_fraction = np.mean(squared <= CONVEX_BALL_RADIUS**2, axis=0)
    slab_centers = (
        np.asarray([-0.50, -0.25, 0.0, 0.25, 0.50]),
        np.asarray([-0.50, -0.25, 0.0, 0.25, 0.50]),
        np.asarray([0.0, 0.25, 0.50, 0.75, 1.0]),
    )
    slab_fractions = []
    for axis, centers_axis in enumerate(slab_centers):
        slab_fractions.extend(
            np.mean(
                np.abs(points[:, axis, None] - centers_axis[None, :])
                <= CONVEX_SLAB_WIDTH / 2.0,
                axis=0,
            )
        )
    return {
        "maximum_fixed_ball_fraction": float(np.max(ball_fraction)),
        "maximum_axis_aligned_slab_fraction": float(np.max(slab_fractions)),
    }


def family_geometry(
    xy: np.ndarray,
    all_edges: np.ndarray,
    selected_indices: np.ndarray,
    signs: np.ndarray,
    carrier_lag_seconds: np.ndarray,
    integration_points: np.ndarray,
    seed: int,
) -> dict:
    selected = np.asarray(selected_indices, dtype=np.int64)
    pairs = np.asarray(all_edges, dtype=np.int64)[selected]
    lag = np.asarray(carrier_lag_seconds, dtype=np.float64)[selected]
    start, stop, direction = tube_segments(
        xy,
        pairs,
        signs,
        lag,
        TIME_SCALE_SECONDS,
    )
    occupation = {
        str(radius): tube_occupation_diagnostics(
            start,
            stop,
            radius,
            integration_points,
        )
        for radius in RADII
    }
    diagnostics = direction_diagnostics(
        direction,
        pairs,
        n_nodes=len(xy),
        seed=seed,
        triple_samples=TRIPLE_SAMPLES,
    )
    diagnostics.update(
        covering_slope(
            start,
            stop,
            scales=COVERING_SCALES,
            points_per_segment=41,
        )
    )
    diagnostics.update(convex_concentration(start, stop))
    return {
        "selected_edge_indices": selected,
        "selected_edge_pairs": pairs,
        "signs": np.asarray(signs, dtype=np.int8),
        "carrier_lag_seconds": lag,
        "occupation": occupation,
        "diagnostics": diagnostics,
    }


def family_union_fraction(
    xy: np.ndarray,
    all_edges: np.ndarray,
    selected_indices: np.ndarray,
    signs: np.ndarray,
    carrier_lag_seconds: np.ndarray,
    integration_points: np.ndarray,
    radius: float = PRIMARY_RADIUS,
) -> float:
    selected = np.asarray(selected_indices, dtype=np.int64)
    pairs = np.asarray(all_edges, dtype=np.int64)[selected]
    lag = np.asarray(carrier_lag_seconds, dtype=np.float64)[selected]
    start, stop, _ = tube_segments(
        xy,
        pairs,
        signs,
        lag,
        TIME_SCALE_SECONDS,
    )
    return tube_occupation_diagnostics(
        start,
        stop,
        radius,
        integration_points,
    )["union_fraction"]


def matched_structural_family_indices(
    all_edges: np.ndarray,
    xy: np.ndarray,
    carrier_lag_seconds: np.ndarray,
    selected_indices: np.ndarray,
    signs: np.ndarray,
    rng: np.random.Generator,
) -> tuple[np.ndarray, np.ndarray, dict]:
    _, length_bin = spatial_length_deciles(xy, all_edges)
    lag_bin = carrier_lag_bins(carrier_lag_seconds)
    selected = np.asarray(selected_indices, dtype=np.int64)
    selected_strata: dict[tuple[int, int], list[int]] = defaultdict(list)
    pool_strata: dict[tuple[int, int], list[int]] = defaultdict(list)
    for index, key in enumerate(zip(length_bin, lag_bin)):
        pool_strata[(int(key[0]), int(key[1]))].append(index)
    for position, index in enumerate(selected):
        key = (int(length_bin[index]), int(lag_bin[index]))
        selected_strata[key].append(position)
    sampled_by_position = np.empty(len(selected), dtype=np.int64)
    for key, positions in selected_strata.items():
        pool = np.asarray(pool_strata[key], dtype=np.int64)
        if len(pool) < len(positions):
            raise RuntimeError("A matched structural stratum is too small.")
        sampled = rng.choice(pool, size=len(positions), replace=False)
        sampled_by_position[np.asarray(positions)] = sampled
    sampled_signs = np.asarray(signs, dtype=np.int8)[rng.permutation(len(signs))]
    audit = {
        "selected_length_bin_counts": np.bincount(
            length_bin[selected], minlength=10
        ).tolist(),
        "sampled_length_bin_counts": np.bincount(
            length_bin[sampled_by_position], minlength=10
        ).tolist(),
        "selected_lag_bin_counts": np.bincount(
            lag_bin[selected], minlength=5
        ).tolist(),
        "sampled_lag_bin_counts": np.bincount(
            lag_bin[sampled_by_position], minlength=5
        ).tolist(),
        "selected_positive_signs": int(np.sum(np.asarray(signs) > 0)),
        "sampled_positive_signs": int(np.sum(sampled_signs > 0)),
    }
    return sampled_by_position, sampled_signs, audit


def matched_geometry_distribution(
    xy: np.ndarray,
    all_edges: np.ndarray,
    carrier_lag_seconds: np.ndarray,
    selected_indices: np.ndarray,
    signs: np.ndarray,
    seed_label: str,
    family_count: int = MATCHED_FAMILY_COUNT,
    integration_point_count: int = INTEGRATION_POINT_COUNT,
) -> dict:
    points = fixed_integration_points(
        integration_point_count,
        stable_seed(seed_label, "integration"),
    )
    observed = family_geometry(
        xy,
        all_edges,
        selected_indices,
        signs,
        carrier_lag_seconds,
        points,
        stable_seed(seed_label, "observed"),
    )
    rng = np.random.default_rng(stable_seed(seed_label, "matched-families"))
    distributions = {
        "union_0.025": [],
        "union_0.035": [],
        "union_0.05": [],
        "direction_minimum_eigenvalue": [],
        "direction_effective_rank": [],
        "trilinear_determinant_mean": [],
        "trilinear_determinant_q75": [],
        "sensor_degree_hhi": [],
        "maximum_sensor_degree_fraction": [],
        "log_covering_slope": [],
        "maximum_fixed_ball_fraction": [],
        "maximum_axis_aligned_slab_fraction": [],
    }
    first_audit = None
    for family_index in range(family_count):
        sampled, sampled_signs, audit = matched_structural_family_indices(
            all_edges,
            xy,
            carrier_lag_seconds,
            selected_indices,
            signs,
            rng,
        )
        if first_audit is None:
            first_audit = audit
        geometry = family_geometry(
            xy,
            all_edges,
            sampled,
            sampled_signs,
            carrier_lag_seconds,
            points,
            stable_seed(seed_label, "matched", family_index),
        )
        for radius in RADII:
            distributions[f"union_{radius}"].append(
                geometry["occupation"][str(radius)]["union_fraction"]
            )
        for name in (
            "direction_minimum_eigenvalue",
            "direction_effective_rank",
            "trilinear_determinant_mean",
            "trilinear_determinant_q75",
            "sensor_degree_hhi",
            "maximum_sensor_degree_fraction",
            "log_covering_slope",
            "maximum_fixed_ball_fraction",
            "maximum_axis_aligned_slab_fraction",
        ):
            distributions[name].append(geometry["diagnostics"][name])
    arrays = {
        name: np.asarray(values, dtype=np.float64)
        for name, values in distributions.items()
    }
    observed_flat = {
        f"union_{radius}": observed["occupation"][str(radius)]["union_fraction"]
        for radius in RADII
    }
    observed_flat.update(
        {
            name: observed["diagnostics"][name]
            for name in (
                "direction_minimum_eigenvalue",
                "direction_effective_rank",
                "trilinear_determinant_mean",
                "trilinear_determinant_q75",
                "sensor_degree_hhi",
                "maximum_sensor_degree_fraction",
                "log_covering_slope",
                "maximum_fixed_ball_fraction",
                "maximum_axis_aligned_slab_fraction",
            )
        }
    )
    comparison = {}
    for name, value in observed_flat.items():
        null = arrays[name]
        standard = float(np.std(null, ddof=1))
        comparison[name] = {
            "observed": float(value),
            "matched_mean": float(np.mean(null)),
            "matched_median": float(np.median(null)),
            "matched_standard_deviation": standard,
            "z": float((value - np.mean(null)) / standard) if standard > 0 else 0.0,
            "upper_tail_p": float((1 + np.sum(null >= value)) / (len(null) + 1)),
            "lower_tail_p": float((1 + np.sum(null <= value)) / (len(null) + 1)),
        }
    return {
        "observed": observed,
        "comparison": comparison,
        "matched_distributions": arrays,
        "first_family_matching_audit": first_audit,
        "family_count": family_count,
        "integration_point_count": integration_point_count,
    }
```
