## Supplementary material for "Falsifiable substitution tests reveal task-structured neural evidence for auditory attention": Article_Analysis_Source_Code: kite_st_hd_lock_utils_v2.py

```
"""Dynamic hash-bound validation of the KITE-ST-HD-v2 implementation lock."""

from __future__ import annotations

import hashlib
import json
from pathlib import Path


ROOT = Path(__file__).resolve().parents[1]
LOCK = ROOT / "docs" / "kite_st_hd_implementation_lock_v2.json"
PROTOCOL = ROOT / "docs" / "kite_st_hd_preoutcome_protocol_v2.json"
PROTOCOL_HASH = "67ca142eb44a391b555612f4d11f77077b8063916ab1be75c9b3f19f912579f0"


def sha256(path: Path) -> str:
    digest = hashlib.sha256()
    with path.open("rb") as stream:
        for chunk in iter(lambda: stream.read(8 * 1024 * 1024), b""):
            digest.update(chunk)
    return digest.hexdigest()


def parse_sidecar(path: Path) -> tuple[str, str]:
    fields = path.read_text(encoding="ascii").strip().split()
    if len(fields) != 2:
        raise RuntimeError(f"Invalid SHA256 sidecar: {path}")
    return fields[0].lower(), fields[1]


def confirmation_numeric_files() -> list[str]:
    confirmation = {
        "S0", "S10", "S26", "S2", "S21", "S28", "S22",
        "S9", "S3", "S15", "S16", "S18", "S13", "S1",
    }
    roots = (
        ROOT / "data" / "raw" / "kite_st_hd_zenodo4518754",
        ROOT / "data" / "derived" / "kite_st_hd_zenodo4518754",
    )
    suffixes = (".tar.gz", ".dat", ".npy", ".npz", ".mat", ".fif", ".edf")
    found = []
    for base in roots:
        if not base.exists():
            continue
        for path in base.rglob("*"):
            if not path.is_file() or not path.name.lower().endswith(suffixes):
                continue
            if any(
                part == subject
                or part.startswith(subject + "_AAD_")
                or part == subject + ".tar.gz"
                for subject in confirmation
                for part in path.parts
            ):
                found.append(path.relative_to(ROOT).as_posix())
    return sorted(set(found))


def validate_implementation_lock(caller: Path) -> tuple[dict, str]:
    if sha256(PROTOCOL) != PROTOCOL_HASH:
        raise RuntimeError("KITE-ST-HD-v2 protocol hash mismatch.")
    sidecar_hash, sidecar_name = parse_sidecar(
        LOCK.with_name(LOCK.name + ".sha256")
    )
    observed_lock_hash = sha256(LOCK)
    if sidecar_name != LOCK.name or sidecar_hash != observed_lock_hash:
        raise RuntimeError("V2 implementation-lock sidecar mismatch.")
    payload = json.loads(LOCK.read_text(encoding="utf-8"))
    if payload.get("status") != "locked_before_any_v2_development_archive_open_or_numeric_read":
        raise RuntimeError("V2 implementation lock does not have the required status.")
    if payload.get("v2_protocol_sha256") != PROTOCOL_HASH:
        raise RuntimeError("V2 implementation lock parent mismatch.")
    relative = caller.resolve().relative_to(ROOT.resolve()).as_posix()
    expected = payload["locked_source_hashes"].get(relative)
    if expected is None or sha256(caller) != expected:
        raise RuntimeError(f"Caller is not hash-bound by v2 lock: {relative}")
    for path_text, expected_hash in payload["locked_source_hashes"].items():
        if sha256(ROOT / path_text) != expected_hash:
            raise RuntimeError(f"V2 locked source mismatch: {path_text}")
    hits = confirmation_numeric_files()
    if hits:
        raise RuntimeError(f"V2 confirmation firewall violation: {hits}")
    return payload, observed_lock_hash
```
