## Supplementary material for "Falsifiable substitution tests reveal task-structured neural evidence for auditory attention": Article_Analysis_Source_Code: kite_st_hd_statistics_v1.py

```
"""Frozen attention-effect, exact-inference and decoding statistics."""

from __future__ import annotations

import numpy as np
from scipy.stats import rankdata

from kite_st_hd_core_v1 import (
    all_sign_patterns,
    cross_story_edge_score,
    exact_upper_p,
    paired_bounded_effect,
    select_edges,
)


CONDITIONS = ("1L", "1R", "2L", "2R")


def participant_story_effects(features: np.ndarray) -> tuple[np.ndarray, np.ndarray]:
    """Return bounded part-1 and part-2 L-minus-R effects."""
    values = np.asarray(features, dtype=np.float64)
    if values.ndim != 4 or values.shape[1:3] != (4, 12):
        raise ValueError("Expected participant x condition x block x feature.")
    first = np.asarray(
        [paired_bounded_effect(subject[0], subject[1]) for subject in values]
    )
    second = np.asarray(
        [paired_bounded_effect(subject[2], subject[3]) for subject in values]
    )
    return first, second


def story_order_contrasts(acquisition_orders: np.ndarray) -> tuple[np.ndarray, np.ndarray]:
    orders = np.asarray(acquisition_orders, dtype=np.float64)
    if orders.ndim != 2 or orders.shape[1] != 4:
        raise ValueError("Expected participant x four acquisition ranks.")
    if any(len(np.unique(row)) != 4 for row in orders):
        raise ValueError("Each participant must have four unique acquisition ranks.")
    return orders[:, 0] - orders[:, 1], orders[:, 2] - orders[:, 3]


def residualize_order(effect: np.ndarray, contrast: np.ndarray) -> np.ndarray:
    """Remove the centered acquisition-order contrast while preserving means."""
    values = np.asarray(effect, dtype=np.float64)
    q = np.asarray(contrast, dtype=np.float64)
    if values.ndim != 2 or q.shape != (len(values),):
        raise ValueError("Order residualization dimensions do not match.")
    centered = q - np.mean(q)
    denominator = float(centered @ centered)
    if denominator <= 1e-12:
        return values.copy()
    beta = centered @ values / denominator
    residual = values - centered[:, None] * beta[None, :]
    if not np.allclose(np.mean(residual, axis=0), np.mean(values, axis=0), atol=1e-12):
        raise RuntimeError("Order residualization did not preserve group means.")
    return residual


def observed_candidate_fit(
    candidate_effects: dict[str, tuple[np.ndarray, np.ndarray]],
    order_contrasts: tuple[np.ndarray, np.ndarray],
    selected_count: int,
) -> dict[str, dict]:
    q1, q2 = order_contrasts
    fitted: dict[str, dict] = {}
    for name, (first, second) in candidate_effects.items():
        r1 = residualize_order(first, q1)
        r2 = residualize_order(second, q2)
        score = cross_story_edge_score(r1, r2)
        selected = select_edges(score, selected_count)
        signed_mean = np.mean(r1[:, selected] + r2[:, selected], axis=0) / 2.0
        signs = np.where(signed_mean >= 0.0, 1, -1).astype(np.int8)
        combined_weight = signed_mean.astype(np.float64)
        norm = float(np.linalg.norm(combined_weight))
        if norm <= 1e-12:
            combined_weight = signs.astype(np.float64) / np.sqrt(len(signs))
        else:
            combined_weight /= norm
        fitted[name] = {
            "part1_residualized": r1,
            "part2_residualized": r2,
            "edge_score": score,
            "selected_edge_indices": selected,
            "selected_edge_signs": signs,
            "selected_statistic": float(np.mean(score[selected])),
            "combined_template_weight": combined_weight,
        }
    return fitted


def exact_candidate_familywise(
    candidate_effects: dict[str, tuple[np.ndarray, np.ndarray]],
    order_contrasts: tuple[np.ndarray, np.ndarray],
    selected_count: int,
) -> dict:
    names = tuple(candidate_effects)
    n = len(next(iter(candidate_effects.values()))[0])
    patterns = all_sign_patterns(n)
    q1, q2 = (np.asarray(value, dtype=np.float64) for value in order_contrasts)
    nulls = {name: np.empty(len(patterns), dtype=np.float64) for name in names}
    for pattern_index, signs in enumerate(patterns):
        for name in names:
            first, second = candidate_effects[name]
            r1 = residualize_order(first * signs[:, None], q1 * signs)
            r2 = residualize_order(second * signs[:, None], q2 * signs)
            score = cross_story_edge_score(r1, r2)
            selected = select_edges(score, selected_count)
            nulls[name][pattern_index] = float(np.mean(score[selected]))
    maximum = np.max(np.column_stack([nulls[name] for name in names]), axis=1)
    observed = {name: float(nulls[name][0]) for name in names}
    familywise_p = {
        name: exact_upper_p(maximum, observed[name]) for name in names
    }
    winner = max(names, key=lambda name: (observed[name], -names.index(name)))
    return {
        "observed": observed,
        "familywise_p": familywise_p,
        "winner": winner,
        "winner_observed": observed[winner],
        "winner_familywise_p": familywise_p[winner],
        "pattern_count": len(patterns),
        "nulls": nulls,
        "maximum_null": maximum,
    }


def binary_auc(left_scores: np.ndarray, right_scores: np.ndarray) -> float:
    left = np.asarray(left_scores, dtype=np.float64)
    right = np.asarray(right_scores, dtype=np.float64)
    joined = np.r_[left, right]
    ranks = rankdata(joined, method="average")
    n_left = len(left)
    n_right = len(right)
    if n_left == 0 or n_right == 0:
        raise ValueError("AUC requires both classes.")
    u = np.sum(ranks[:n_left]) - n_left * (n_left + 1) / 2.0
    return float(u / (n_left * n_right))


def label_blind_standardize(left: np.ndarray, right: np.ndarray) -> tuple[np.ndarray, np.ndarray]:
    joined = np.row_stack([left, right]).astype(np.float64)
    joined -= np.mean(joined, axis=0, keepdims=True)
    scale = np.sqrt(np.mean(joined * joined, axis=0))
    scale = np.where(scale > 1e-12, scale, 1.0)
    joined /= scale[None, :]
    return joined[: len(left)], joined[len(left) :]


def score_story(
    left: np.ndarray,
    right: np.ndarray,
    weights: np.ndarray,
) -> float:
    standardized_left, standardized_right = label_blind_standardize(left, right)
    return binary_auc(standardized_left @ weights, standardized_right @ weights)


def leave_one_participant_out_cross_story_auc(
    features: np.ndarray,
    acquisition_orders: np.ndarray,
    selected_count: int,
) -> dict:
    values = np.asarray(features, dtype=np.float64)
    first, second = participant_story_effects(values)
    q1, q2 = story_order_contrasts(acquisition_orders)
    aucs = []
    directional = []
    fold_edge_indices = []
    n = len(values)
    for held_out in range(n):
        training = np.arange(n) != held_out
        r1 = residualize_order(first[training], q1[training])
        r2 = residualize_order(second[training], q2[training])
        edge_score = cross_story_edge_score(r1, r2)
        selected = select_edges(edge_score, selected_count)
        weight1 = np.mean(r1[:, selected], axis=0)
        weight2 = np.mean(r2[:, selected], axis=0)
        norm1 = float(np.linalg.norm(weight1))
        norm2 = float(np.linalg.norm(weight2))
        if norm1 <= 1e-12 or norm2 <= 1e-12:
            auc12 = auc21 = 0.5
        else:
            weight1 /= norm1
            weight2 /= norm2
            auc12 = score_story(
                values[held_out, 2][:, selected],
                values[held_out, 3][:, selected],
                weight1,
            )
            auc21 = score_story(
                values[held_out, 0][:, selected],
                values[held_out, 1][:, selected],
                weight2,
            )
        directional.append([auc12, auc21])
        aucs.append((auc12 + auc21) / 2.0)
        fold_edge_indices.append(selected)
    aucs_array = np.asarray(aucs)
    centered = aucs_array - 0.5
    patterns = all_sign_patterns(n)
    null = np.mean(patterns * centered[None, :], axis=1)
    observed = float(np.mean(centered))
    return {
        "participant_auc": aucs_array,
        "directional_auc": np.asarray(directional),
        "mean_auc": float(np.mean(aucs_array)),
        "participants_above_chance": int(np.sum(aucs_array > 0.5)),
        "exact_one_sided_p": exact_upper_p(null, observed),
        "pattern_count": len(patterns),
        "fold_selected_edge_indices": fold_edge_indices,
    }


def frozen_template_auc(
    features: np.ndarray,
    selected: np.ndarray,
    weights: np.ndarray,
) -> dict:
    values = np.asarray(features, dtype=np.float64)
    selected = np.asarray(selected, dtype=np.int64)
    weight = np.asarray(weights, dtype=np.float64)
    if weight.shape != (len(selected),):
        raise ValueError("Frozen-template weight dimensions do not match.")
    participant_aucs = []
    directional = []
    for subject in values:
        first = score_story(subject[0][:, selected], subject[1][:, selected], weight)
        second = score_story(subject[2][:, selected], subject[3][:, selected], weight)
        directional.append([first, second])
        participant_aucs.append((first + second) / 2.0)
    participant_aucs = np.asarray(participant_aucs)
    centered = participant_aucs - 0.5
    patterns = all_sign_patterns(len(values))
    null = np.mean(patterns * centered[None, :], axis=1)
    observed = float(np.mean(centered))
    return {
        "participant_auc": participant_aucs,
        "directional_auc": np.asarray(directional),
        "mean_auc": float(np.mean(participant_aucs)),
        "participants_above_chance": int(np.sum(participant_aucs > 0.5)),
        "exact_one_sided_p": exact_upper_p(null, observed),
        "pattern_count": len(patterns),
    }


def fixed_local_erasure_patches(xy: np.ndarray, sensors_per_patch: int = 25) -> list[dict]:
    coordinates = np.asarray(xy, dtype=np.float64)
    if coordinates.shape != (222, 2):
        raise ValueError("Local-erasure atlas expects the 222 primary sensors.")
    anchors = []
    for radius, angles in (
        (0.45, np.arange(0, 2 * np.pi, np.pi / 4.0)),
        (0.20, np.arange(0, 2 * np.pi, np.pi / 2.0)),
    ):
        anchors.extend(
            [(radius * np.cos(angle), radius * np.sin(angle)) for angle in angles]
        )
    patches = []
    for index, anchor in enumerate(np.asarray(anchors)):
        distance = np.linalg.norm(coordinates - anchor[None, :], axis=1)
        sensors = np.argsort(distance, kind="stable")[:sensors_per_patch]
        patches.append(
            {
                "patch_id": f"patch_{index + 1:02d}",
                "anchor_xy": anchor,
                "sensor_indices": sensors.astype(np.int32),
            }
        )
    return patches


def local_erasure_frozen_template_auc(
    features: np.ndarray,
    all_edges: np.ndarray,
    selected: np.ndarray,
    weights: np.ndarray,
    xy: np.ndarray,
) -> dict:
    edges = np.asarray(all_edges, dtype=np.int64)
    selected = np.asarray(selected, dtype=np.int64)
    selected_pairs = edges[selected]
    weight = np.asarray(weights, dtype=np.float64)
    patches = fixed_local_erasure_patches(xy)
    results = []
    for patch in patches:
        sensors = patch["sensor_indices"]
        retain = ~(
            np.isin(selected_pairs[:, 0], sensors)
            | np.isin(selected_pairs[:, 1], sensors)
        )
        if np.sum(retain) < 10:
            raise RuntimeError("A local-erasure patch retained fewer than 10 edges.")
        retained_weight = weight[retain]
        retained_weight /= np.linalg.norm(retained_weight)
        auc = frozen_template_auc(
            features,
            selected[retain],
            retained_weight,
        )
        results.append(
            {
                "patch_id": patch["patch_id"],
                "anchor_xy": np.asarray(patch["anchor_xy"], dtype=float),
                "sensor_indices": sensors,
                "retained_edge_count": int(np.sum(retain)),
                "mean_auc": auc["mean_auc"],
                "participant_auc": auc["participant_auc"],
                "exact_one_sided_p": auc["exact_one_sided_p"],
            }
        )
    worst = min(results, key=lambda item: item["mean_auc"])
    return {
        "patch_count": len(results),
        "sensors_per_patch": 25,
        "results": results,
        "worst_patch_id": worst["patch_id"],
        "worst_patch_mean_auc": worst["mean_auc"],
    }
```
