## Supplementary material for "Falsifiable substitution tests reveal task-structured neural evidence for auditory attention": Article_Analysis_Source_Code: kite_st_hd_subject_io_v1.py

```
"""Outcome-blind subject-level preprocessing and feature I/O for KITE-ST-HD."""

from __future__ import annotations

import hashlib
import json
from pathlib import Path

import numpy as np

import preprocess_kite_st_hd_engineering_v1 as preprocessing
from kite_st_hd_core_v1 import load_public_geometry
from kite_st_hd_features_v1 import (
    extract_recording_features,
    feature_geometry,
    pooled_carrier_lags,
)


ROOT = Path(__file__).resolve().parents[1]
CONDITIONS = ("1L", "1R", "2L", "2R")
COORDINATES = (
    ROOT
    / "downloads"
    / "candidate_audit"
    / "zenodo4518754_support"
    / "misc"
    / "misc"
    / "eeg255ch_locs.csv"
)


def sha256(path: Path) -> str:
    digest = hashlib.sha256()
    with path.open("rb") as stream:
        for chunk in iter(lambda: stream.read(8 * 1024 * 1024), b""):
            digest.update(chunk)
    return digest.hexdigest()


def preprocess_subject(
    subject: str,
    work_root: Path,
    output_dir: Path,
) -> list[dict]:
    if output_dir.exists():
        raise RuntimeError(f"Refusing existing subject preprocessing: {output_dir}")
    if not (work_root / subject).is_dir():
        raise RuntimeError(f"Subject work directory is absent: {work_root / subject}")
    output_dir.mkdir(parents=True)
    geometry = load_public_geometry(COORDINATES)
    original_work = preprocessing.WORK
    try:
        preprocessing.WORK = work_root
        records = [
            preprocessing.process_recording(subject, condition, output_dir, geometry)
            for condition in CONDITIONS
        ]
    finally:
        preprocessing.WORK = original_work
    return records


def build_feature_archive(
    subject: str,
    preprocessing_records: list[dict],
    output_path: Path,
) -> dict:
    if output_path.exists():
        raise RuntimeError(f"Refusing existing feature archive: {output_path}")
    if [item["recording"] for item in preprocessing_records] != [
        f"{subject}_AAD_{condition}" for condition in CONDITIONS
    ]:
        raise RuntimeError("Preprocessing record order does not match frozen conditions.")
    public_geometry = load_public_geometry(COORDINATES)
    geometry = feature_geometry(public_geometry)
    condition_features: list[dict[str, np.ndarray]] = []
    input_hashes: dict[str, str] = {}
    for record in preprocessing_records:
        input_path = ROOT / record["output"]["path"]
        observed = sha256(input_path)
        if observed != record["output"]["sha256"]:
            raise RuntimeError(f"Preprocessed hash mismatch: {input_path}")
        with np.load(input_path, allow_pickle=False) as saved:
            features = extract_recording_features(saved["eeg"], geometry)
        condition_features.append(features)
        input_hashes[record["recording"]] = observed
    carrier_a, carrier_b = pooled_carrier_lags(condition_features)
    stacked = {
        name: np.stack([item[name] for item in condition_features], axis=0)
        for name in condition_features[0]
    }
    output_path.parent.mkdir(parents=True, exist_ok=True)
    np.savez_compressed(
        output_path,
        **stacked,
        A_pooled_carrier_lag_seconds=carrier_a,
        B_pooled_carrier_lag_seconds=carrier_b,
        condition_labels=np.asarray(CONDITIONS),
        primary_global_indices=geometry.primary_indices,
        frontal_global_indices=geometry.frontal_indices,
        primary_xy=geometry.primary_xy.astype(np.float32),
        primary_edges=geometry.primary_edges,
        frontal_edges=geometry.frontal_edges,
        posterior_left_global_indices=geometry.posterior_left_indices,
        posterior_right_global_indices=geometry.posterior_right_indices,
    )
    output_hash = sha256(output_path)
    output_path.with_name(output_path.name + ".sha256").write_text(
        f"{output_hash}  {output_path.name}\n",
        encoding="ascii",
    )
    with np.load(output_path, allow_pickle=False) as saved:
        shapes = {name: list(saved[name].shape) for name in saved.files}
        finite = {
            name: bool(np.isfinite(saved[name]).all())
            for name in saved.files
            if saved[name].dtype.kind in "fci"
        }
    if not all(finite.values()):
        raise RuntimeError("Feature archive contains nonfinite numerical values.")
    return {
        "subject": subject,
        "path": output_path.relative_to(ROOT).as_posix(),
        "bytes": output_path.stat().st_size,
        "sha256": output_hash,
        "preprocessed_input_sha256": input_hashes,
        "array_shapes": shapes,
        "all_numeric_arrays_finite": all(finite.values()),
        "condition_contrast_computed": False,
        "true_label_decoder_computed": False,
        "edge_selection_computed": False,
        "empirical_geometry_endpoint_computed": False,
    }


def write_subject_manifest(
    path: Path,
    payload: dict,
) -> str:
    if path.exists():
        raise RuntimeError(f"Refusing existing subject manifest: {path}")
    path.parent.mkdir(parents=True, exist_ok=True)
    path.write_text(json.dumps(payload, indent=2) + "\n", encoding="utf-8")
    observed = sha256(path)
    path.with_name(path.name + ".sha256").write_text(
        f"{observed}  {path.name}\n",
        encoding="ascii",
    )
    return observed
```
