## Supplementary material for "Falsifiable substitution tests reveal task-structured neural evidence for auditory attention": Article_Analysis_Source_Code: kite_st_hd_subject_io_v2.py

```
"""KITE-ST-HD-v2 subject I/O after the disclosed S7 label-blind QC stop.

The only preprocessing change from v1 is a frozen maximum of 15 robust-scale
bad channels per 253-channel recording.  Thresholds, interpolation and every
signal/feature operation are unchanged.
"""

from __future__ import annotations

from pathlib import Path

import numpy as np

import preprocess_kite_st_hd_engineering_v1 as preprocessing
from inspect_kite_st_hd_engineering_v1 import fixed_qc_indices
from kite_st_hd_core_v1 import load_public_geometry
from kite_st_hd_subject_io_v1 import (
    CONDITIONS,
    COORDINATES,
    ROOT,
    build_feature_archive,
    sha256,
    write_subject_manifest,
)


MAXIMUM_BAD_CHANNELS_V2 = 15


def bad_channels_v2(
    data: np.memmap,
    central_start: int,
    central_stop: int,
) -> tuple[np.ndarray, np.ndarray, np.ndarray]:
    indices = fixed_qc_indices(
        central_start,
        central_stop,
        preprocessing.RAW_SAMPLING_HZ,
    )
    raw = np.asarray(data[indices, :253], dtype=np.float64)
    medians = np.median(raw, axis=0)
    centered = raw - medians[None, :]
    car = centered - np.mean(centered, axis=1, keepdims=True)
    scale = preprocessing.robust_scale(car)
    ratio = scale / np.median(scale)
    bad = np.flatnonzero(
        ~np.isfinite(ratio)
        | (ratio < preprocessing.LOW_RATIO)
        | (ratio > preprocessing.HIGH_RATIO)
    )
    if len(bad) > MAXIMUM_BAD_CHANNELS_V2:
        raise RuntimeError(
            f"More than {MAXIMUM_BAD_CHANNELS_V2} bad scalp channels: {bad.tolist()}"
        )
    return bad.astype(np.int32), medians, ratio


def preprocess_subject_v2(
    subject: str,
    work_root: Path,
    output_dir: Path,
) -> list[dict]:
    if output_dir.exists():
        raise RuntimeError(f"Refusing existing v2 preprocessing: {output_dir}")
    if not (work_root / subject).is_dir():
        raise RuntimeError(f"Subject work directory is absent: {work_root / subject}")
    output_dir.mkdir(parents=True)
    geometry = load_public_geometry(COORDINATES)
    original_work = preprocessing.WORK
    original_bad_channels = preprocessing.bad_channels
    try:
        preprocessing.WORK = work_root
        preprocessing.bad_channels = bad_channels_v2
        records = [
            preprocessing.process_recording(subject, condition, output_dir, geometry)
            for condition in CONDITIONS
        ]
    finally:
        preprocessing.WORK = original_work
        preprocessing.bad_channels = original_bad_channels
    return records
```
