## Supplementary material for "Falsifiable substitution tests reveal task-structured neural evidence for auditory attention": Article_Analysis_Source_Code: prepare_kite_st_hd_confirmation_complete_case_ready_v2.py

```
"""Create the compatibility-ready record for the frozen 12-subject v2 analysis."""

from __future__ import annotations

import json
from pathlib import Path

from kite_sat_confirmation_complete_case_lock_utils_v2 import (
    ROOT,
    sha256,
    validate_complete_case_lock,
)


DISPOSITION = (
    ROOT / "results" / "kite_st_hd" / "confirmation_complete_case_v2"
    / "preprocessing_disposition_v2.json"
)
OUTPUT = DISPOSITION.parent / "confirmation_preprocessing_ready_v2.json"


def main() -> None:
    if OUTPUT.exists():
        raise RuntimeError("Refusing to overwrite complete-case ready record.")
    lock, lock_hash = validate_complete_case_lock(Path(__file__))
    disposition = json.loads(DISPOSITION.read_text(encoding="utf-8"))
    subjects = lock["confirmation_subject_order"]
    if (
        disposition["status"] != "pass"
        or disposition["pass_subject_order"] != subjects
        or disposition["pass_count"] != len(subjects)
        or disposition["technical_failure_count"] != 2
        or disposition["attention_outcome_computed"]
        or disposition["AKB_score_computed"]
        or disposition["original_v1_decision_remains"] != "technical_failure"
    ):
        raise RuntimeError("Complete-case preprocessing disposition mismatch.")
    ready_subjects = []
    for expected, record in zip(subjects, disposition["pass_records"]):
        if expected != record["subject"]:
            raise RuntimeError("Complete-case pass order mismatch.")
        manifest_path = ROOT / record["manifest_path"]
        feature_path = ROOT / record["feature_path"]
        if (
            sha256(manifest_path) != record["manifest_sha256"]
            or sha256(feature_path) != record["feature_sha256"]
            or len(record["preprocessed_sha256"]) != 4
        ):
            raise RuntimeError(f"Complete-case hash mismatch: {expected}")
        ready_subjects.append(
            {
                "subject": expected,
                "manifest_path": record["manifest_path"],
                "manifest_sha256": record["manifest_sha256"],
                "maximum_bad_channel_count": record["maximum_bad_channel_count"],
                "preprocessed_sha256": record["preprocessed_sha256"],
                "feature_archive_sha256": record["feature_sha256"],
            }
        )
    payload = {
        "schema": "KITE-ST-HD-confirmation-preprocessing-ready-v2-complete-case",
        "status": "pass",
        "confirmation_implementation_lock_sha256": lock_hash,
        "subject_order": subjects,
        "subject_count": len(subjects),
        "subjects": ready_subjects,
        "all_four_recordings_present": True,
        "technical_failure_subjects_excluded_before_outcome": ["S26", "S3"],
        "replacement_subjects": [],
        "attention_outcome_computed": False,
        "AKB_score_computed": False,
        "paper_mutated": False,
    }
    OUTPUT.write_text(json.dumps(payload, indent=2) + "\n", encoding="utf-8")
    observed = sha256(OUTPUT)
    OUTPUT.with_name(OUTPUT.name + ".sha256").write_text(
        f"{observed}  {OUTPUT.name}\n", encoding="ascii"
    )
    print(json.dumps({"status": "pass", "subjects": len(subjects), "sha256": observed}))


if __name__ == "__main__":
    main()
```
