## Supplementary material for "Falsifiable substitution tests reveal task-structured neural evidence for auditory attention": Article_Analysis_Source_Code: preprocess_kite_st_hd_engineering_v1.py

```
"""Frozen value-blind preprocessing for the three KITE-ST-HD engineering subjects."""

from __future__ import annotations

import argparse
import hashlib
import json
from pathlib import Path

import numpy as np
from scipy.signal import butter, resample_poly, sosfiltfilt

from inspect_kite_st_hd_engineering_v1 import (
    CONDITIONS,
    fixed_qc_indices,
    parse_header,
)
from kite_st_hd_core_v1 import load_public_geometry, normalize_label


ROOT = Path(__file__).resolve().parents[1]
WHITELIST = ("S20", "S6", "S8")
WORK = ROOT / "data" / "raw" / "kite_st_hd_zenodo4518754" / "engineering_work"
OUTPUT_ROOT = (
    ROOT
    / "data"
    / "derived"
    / "kite_st_hd_zenodo4518754"
    / "engineering_preprocessed_v1"
)
RESULT_ROOT = ROOT / "results" / "kite_st_hd" / "engineering"
SUMMARY = RESULT_ROOT / "engineering_summary_v1.json"
SUMMARY_HASH = "005d39fea2e606c00fe82004b85365ec843006eef410fe2bfcaf06e53f5456ce"
COORDINATES = (
    ROOT
    / "downloads"
    / "candidate_audit"
    / "zenodo4518754_support"
    / "misc"
    / "misc"
    / "eeg255ch_locs.csv"
)
LOW_RATIO = 0.15
HIGH_RATIO = 8.0
RAW_SAMPLING_HZ = 1000
OUTPUT_SAMPLING_HZ = 100
PADDED_SECONDS = 366
TRIM_SECONDS = 3
FINAL_SECONDS = 360
NEIGHBORS = 6
CURRY_TO_COORDINATE_LABEL = {
    # The released Curry headers consistently use PPO8H at row 48 whereas the
    # released coordinate CSV uses PPO8 at the same unique row.
    "PPO8H": "PPO8",
}


def sha256(path: Path) -> str:
    digest = hashlib.sha256()
    with path.open("rb") as stream:
        for chunk in iter(lambda: stream.read(8 * 1024 * 1024), b""):
            digest.update(chunk)
    return digest.hexdigest()


def longest_trigger_train(data: np.memmap, sampling: int) -> np.ndarray:
    trigger = np.asarray(data[:, -1])
    baseline = float(np.median(trigger[: 5 * sampling]))
    active = trigger > baseline
    rising = np.flatnonzero(active & ~np.r_[False, active[:-1]])
    gaps = np.diff(rising)
    bad = np.flatnonzero((gaps < 0.95 * sampling) | (gaps > 1.05 * sampling))
    boundaries = np.r_[0, bad + 1, len(rising)]
    groups = [
        rising[first:stop]
        for first, stop in zip(boundaries[:-1], boundaries[1:])
    ]
    train = max(groups, key=lambda values: (len(values), -int(values[0])))
    if len(train) < 390:
        raise RuntimeError("No valid long trigger train.")
    if np.any((np.diff(train) < 0.95 * sampling) | (np.diff(train) > 1.05 * sampling)):
        raise RuntimeError("Selected trigger train is not continuous at one Hz.")
    return train


def robust_scale(values: np.ndarray) -> np.ndarray:
    median = np.median(values, axis=0)
    return 1.4826 * np.median(np.abs(values - median[None, :]), axis=0)


def bad_channels(
    data: np.memmap,
    central_start: int,
    central_stop: int,
) -> tuple[np.ndarray, np.ndarray, np.ndarray]:
    indices = fixed_qc_indices(central_start, central_stop, RAW_SAMPLING_HZ)
    raw = np.asarray(data[indices, :253], dtype=np.float64)
    medians = np.median(raw, axis=0)
    centered = raw - medians[None, :]
    car = centered - np.mean(centered, axis=1, keepdims=True)
    scale = robust_scale(car)
    ratio = scale / np.median(scale)
    bad = np.flatnonzero(
        ~np.isfinite(ratio) | (ratio < LOW_RATIO) | (ratio > HIGH_RATIO)
    )
    if len(bad) > 5:
        raise RuntimeError(f"More than five bad scalp channels: {bad.tolist()}")
    return bad.astype(np.int32), medians, ratio


def reference_signal(
    data: np.memmap,
    start: int,
    stop: int,
    medians: np.ndarray,
    good: np.ndarray,
    chunk: int = 32,
) -> np.ndarray:
    reference = np.zeros(stop - start, dtype=np.float64)
    for first in range(0, len(good), chunk):
        indices = good[first : first + chunk]
        values = np.asarray(data[start:stop, :253][:, indices], dtype=np.float64)
        values -= medians[indices][None, :]
        reference += np.sum(values, axis=1)
    reference /= len(good)
    return reference


def filter_resample_scalp(
    data: np.memmap,
    start: int,
    stop: int,
    medians: np.ndarray,
    reference: np.ndarray,
    chunk: int = 24,
) -> np.ndarray:
    sos = butter(
        4,
        [1.0, 15.0],
        btype="bandpass",
        fs=RAW_SAMPLING_HZ,
        output="sos",
    )
    padded_output = PADDED_SECONDS * OUTPUT_SAMPLING_HZ
    output = np.empty((padded_output, 253), dtype=np.float32)
    for first in range(0, 253, chunk):
        indices = np.arange(first, min(first + chunk, 253))
        values = np.asarray(data[start:stop, :253][:, indices], dtype=np.float64)
        values -= medians[indices][None, :]
        values -= reference[:, None]
        filtered = sosfiltfilt(sos, values, axis=0)
        downsampled = resample_poly(filtered, 1, 10, axis=0)
        if len(downsampled) != padded_output:
            raise RuntimeError("Unexpected resampled scalp length.")
        output[:, indices] = downsampled.astype(np.float32)
    trim = TRIM_SECONDS * OUTPUT_SAMPLING_HZ
    return output[trim:-trim]


def filter_resample_auxiliary(
    data: np.memmap,
    start: int,
    stop: int,
) -> np.ndarray:
    sos = butter(
        4,
        [1.0, 15.0],
        btype="bandpass",
        fs=RAW_SAMPLING_HZ,
        output="sos",
    )
    values = np.asarray(data[start:stop, 253:255], dtype=np.float64)
    values -= np.median(values, axis=0, keepdims=True)
    filtered = sosfiltfilt(sos, values, axis=0)
    downsampled = resample_poly(filtered, 1, 10, axis=0)
    trim = TRIM_SECONDS * OUTPUT_SAMPLING_HZ
    return downsampled[trim:-trim].astype(np.float32)


def regress_auxiliary(scalp: np.ndarray, auxiliary: np.ndarray) -> np.ndarray:
    aux = np.asarray(auxiliary, dtype=np.float64)
    derivative = np.vstack([np.zeros((1, 2)), np.diff(aux, axis=0)])
    design = np.column_stack([aux, derivative])
    design -= np.mean(design, axis=0, keepdims=True)
    scale = np.sqrt(np.mean(design * design, axis=0))
    if np.any(scale <= 1e-12) or not np.isfinite(scale).all():
        raise RuntimeError("Auxiliary nuisance design is degenerate.")
    design /= scale[None, :]
    values = np.asarray(scalp, dtype=np.float64)
    beta, _, _, _ = np.linalg.lstsq(design, values, rcond=None)
    residual = values - design @ beta
    return residual.astype(np.float32)


def interpolate_bad(
    scalp: np.ndarray,
    bad: np.ndarray,
    xy: np.ndarray,
) -> tuple[np.ndarray, dict[str, list]]:
    values = np.asarray(scalp, dtype=np.float32).copy()
    good = np.setdiff1d(np.arange(253), bad)
    details: dict[str, list] = {}
    for target in bad:
        distance = np.linalg.norm(xy[good] - xy[int(target)], axis=1)
        order = np.argsort(distance, kind="stable")[:NEIGHBORS]
        neighbors = good[order]
        weights = 1.0 / np.maximum(distance[order], 1e-8)
        weights /= np.sum(weights)
        values[:, int(target)] = values[:, neighbors] @ weights.astype(np.float32)
        details[str(int(target))] = [
            {
                "neighbor": int(neighbor),
                "weight": float(weight),
            }
            for neighbor, weight in zip(neighbors, weights)
        ]
    return values, details


def process_recording(
    subject: str,
    condition: str,
    output_dir: Path,
    geometry,
) -> dict:
    base = WORK / subject / f"{subject}_AAD_{condition}"
    header = parse_header(base)
    if int(header["SampleFreqHz"]) != RAW_SAMPLING_HZ:
        raise RuntimeError("Unexpected raw sampling frequency.")
    raw_labels = tuple(normalize_label(label) for label in header["scalp_labels"])
    labels = tuple(CURRY_TO_COORDINATE_LABEL.get(label, label) for label in raw_labels)
    if labels != geometry.labels:
        raise RuntimeError("Curry scalp labels do not match public geometry.")
    data = np.memmap(
        base.with_suffix(".dat"),
        dtype="<f4",
        mode="r",
        shape=(header["NumSamples"], header["NumChannels"]),
        order="C",
    )
    train = longest_trigger_train(data, RAW_SAMPLING_HZ)
    center = (int(train[0]) + int(train[-1])) // 2
    padded_samples = PADDED_SECONDS * RAW_SAMPLING_HZ
    padded_start = center - padded_samples // 2
    padded_stop = padded_start + padded_samples
    central_start = center - FINAL_SECONDS * RAW_SAMPLING_HZ // 2
    central_stop = central_start + FINAL_SECONDS * RAW_SAMPLING_HZ
    if padded_start < 0 or padded_stop > len(data):
        raise RuntimeError("Padded crop is outside recording bounds.")
    bad, medians, scale_ratio = bad_channels(data, central_start, central_stop)
    good = np.setdiff1d(np.arange(253), bad)
    reference = reference_signal(data, padded_start, padded_stop, medians, good)
    scalp = filter_resample_scalp(
        data,
        padded_start,
        padded_stop,
        medians,
        reference,
    )
    auxiliary = filter_resample_auxiliary(data, padded_start, padded_stop)
    if scalp.shape != (36000, 253) or auxiliary.shape != (36000, 2):
        raise RuntimeError("Unexpected frozen preprocessing output shape.")
    residual = regress_auxiliary(scalp, auxiliary)
    interpolated, interpolation = interpolate_bad(residual, bad, geometry.xy)
    if not np.isfinite(interpolated).all():
        raise RuntimeError("Preprocessed scalp data contain nonfinite values.")
    output = output_dir / f"{subject}_AAD_{condition}_preprocessed_v1.npz"
    np.savez_compressed(
        output,
        eeg=interpolated.astype(np.float32),
        xtra=auxiliary.astype(np.float32),
        sampling_hz=np.asarray(OUTPUT_SAMPLING_HZ, dtype=np.int32),
        channel_names=np.asarray(labels),
        bad_channel_indices=bad,
        trigger_train_first=np.asarray(int(train[0]), dtype=np.int64),
        trigger_train_last=np.asarray(int(train[-1]), dtype=np.int64),
        padded_start=np.asarray(padded_start, dtype=np.int64),
        padded_stop=np.asarray(padded_stop, dtype=np.int64),
    )
    output_hash = sha256(output)
    output.with_name(output.name + ".sha256").write_text(
        f"{output_hash}  {output.name}\n",
        encoding="ascii",
    )
    result = {
        "recording": base.name,
        "input_dat_bytes": base.with_suffix(".dat").stat().st_size,
        "start_iso": header["start_iso"],
        "channel_label_aliases": {
            raw: mapped
            for raw, mapped in zip(raw_labels, labels)
            if raw != mapped
        },
        "trigger_train": {
            "rising_edges": len(train),
            "first_sample": int(train[0]),
            "last_sample": int(train[-1]),
            "duration_seconds": float((train[-1] - train[0]) / RAW_SAMPLING_HZ),
        },
        "crop": {
            "padded_start": padded_start,
            "padded_stop": padded_stop,
            "padded_seconds": PADDED_SECONDS,
            "trim_seconds_each_end": TRIM_SECONDS,
            "output_seconds": FINAL_SECONDS,
        },
        "bad_channels": {
            "count": len(bad),
            "indices": [int(value) for value in bad],
            "labels": [labels[int(value)] for value in bad],
            "minimum_scale_ratio": float(np.min(scale_ratio)),
            "maximum_scale_ratio": float(np.max(scale_ratio)),
            "interpolation": interpolation,
        },
        "nuisance": {
            "labels": ["XtraL", "XtraR"],
            "columns": 4,
        },
        "output": {
            "path": output.relative_to(ROOT).as_posix(),
            "bytes": output.stat().st_size,
            "sha256": output_hash,
            "eeg_shape": list(interpolated.shape),
            "eeg_dtype": str(interpolated.dtype),
            "xtra_shape": list(auxiliary.shape),
            "finite": True,
            "median_channel_rms": float(
                np.median(np.sqrt(np.mean(interpolated.astype(np.float64) ** 2, axis=0)))
            ),
        },
    }
    del data
    return result


def main() -> None:
    parser = argparse.ArgumentParser()
    parser.add_argument("subject", choices=WHITELIST)
    args = parser.parse_args()
    if sha256(SUMMARY) != SUMMARY_HASH:
        raise RuntimeError("Engineering summary hash mismatch.")
    summary = json.loads(SUMMARY.read_text(encoding="utf-8"))
    choices = summary["fixed_preprocessing_choices"]
    if (
        choices["bad_channel_scale_ratio_low"] != LOW_RATIO
        or choices["bad_channel_scale_ratio_high"] != HIGH_RATIO
        or choices["padded_crop_seconds"] != PADDED_SECONDS
        or choices["filter_padding_trim_seconds_each_end"] != TRIM_SECONDS
    ):
        raise RuntimeError("Preprocessing constants differ from engineering summary.")
    geometry = load_public_geometry(COORDINATES)
    subject = args.subject
    output_dir = OUTPUT_ROOT / subject
    result_path = RESULT_ROOT / f"{subject}_preprocessing_engineering_v1.json"
    if result_path.exists() or (output_dir.exists() and any(output_dir.iterdir())):
        raise RuntimeError(
            f"Refusing to overwrite preprocessing output: {output_dir}/{result_path}"
        )
    output_dir.mkdir(parents=True, exist_ok=True)
    records = [
        process_recording(subject, condition, output_dir, geometry)
        for condition in CONDITIONS
    ]
    payload = {
        "schema": "KITE-ST-HD-engineering-preprocessing-v1",
        "status": "pass",
        "subject": subject,
        "engineering_summary_sha256": SUMMARY_HASH,
        "records": records,
        "acquisition_order": [
            record["recording"]
            for record in sorted(records, key=lambda item: item["start_iso"])
        ],
        "firewall": {
            "true_L_minus_R_contrast_computed": False,
            "true_label_decoder_computed": False,
            "attention_edges_selected": False,
            "empirical_geometry_endpoint_computed": False,
            "development_or_confirmation_file_read": False,
            "paper_mutated": False,
        },
    }
    result_path.write_text(json.dumps(payload, indent=2) + "\n", encoding="utf-8")
    result_hash = sha256(result_path)
    result_path.with_name(result_path.name + ".sha256").write_text(
        f"{result_hash}  {result_path.name}\n",
        encoding="ascii",
    )
    print(
        json.dumps(
            {
                "status": "pass",
                "subject": subject,
                "output": str(result_path),
                "sha256": result_hash,
            }
        )
    )


if __name__ == "__main__":
    main()
```
