## Supplementary material for "Falsifiable substitution tests reveal task-structured neural evidence for auditory attention": Article_Analysis_Source_Code: run_kite_sat_akb_confirmation_complete_case_subject_v2.py

```
"""Run the byte-identical AKB v1 subject engine on the frozen v2 complete cases."""

from __future__ import annotations

from pathlib import Path

import run_kite_sat_akb_confirmation_subject_v1 as engine
from kite_sat_confirmation_complete_case_lock_utils_v2 import (
    ROOT,
    validate_complete_case_lock,
)


SUBJECTS = (
    "S0", "S10", "S2", "S21", "S28", "S22",
    "S9", "S15", "S16", "S18", "S13", "S1",
)
PRIMARY_ROOT = (
    ROOT / "results" / "kite_sat" / "akb_confirmation_complete_case_v2"
)
READY = (
    ROOT / "results" / "kite_st_hd" / "confirmation_complete_case_v2"
    / "confirmation_preprocessing_ready_v2.json"
)
MODEL_ROOT = (
    ROOT / "data" / "derived" / "kite_st_hd_zenodo4518754"
    / "kite_sat_confirmation_complete_case_models_v2"
)


def validate_adapter(_: Path):
    return validate_complete_case_lock(Path(__file__))


def main() -> None:
    engine.SUBJECTS = SUBJECTS
    engine.READY = READY
    engine.OUTPUT_ROOT = PRIMARY_ROOT / "subjects"
    engine.MODEL_ROOT = MODEL_ROOT
    engine.validate_confirmation_lock = validate_adapter
    engine.main()


if __name__ == "__main__":
    main()
```
