## Supplementary material for "Falsifiable substitution tests reveal task-structured neural evidence for auditory attention": Article_Analysis_Source_Code: run_kite_sat_akb_confirmation_group_v1.py

```
"""Frozen fixed-sequence group decision for the KITE-SAT AKB confirmation."""

from __future__ import annotations

import hashlib
import itertools
import json
from pathlib import Path

import numpy as np

from download_kite_st_hd_confirmation_v1 import SUBJECTS
from kite_sat_confirmation_lock_utils_v1 import ROOT, sha256, validate_confirmation_lock
from kite_sat_patch_erasure_v1 import ANCHORS


SUBJECT_ROOT = ROOT / "results" / "kite_sat" / "akb_confirmation_v1" / "subjects"
OUTPUT = (
    ROOT / "results" / "kite_sat" / "akb_confirmation_v1"
    / "confirmation_decision_v1.json"
)


def exact_one_sided(values: np.ndarray) -> float:
    observed = float(np.mean(values))
    exceed = sum(
        float(np.mean(values * np.asarray(signs))) >= observed - 1e-15
        for signs in itertools.product((-1.0, 1.0), repeat=len(values))
    )
    return exceed / (2 ** len(values))


def seed(text: str) -> int:
    return int.from_bytes(hashlib.sha256(text.encode("utf-8")).digest()[:8], "little")


def bootstrap_ci(values: np.ndarray, endpoint: str) -> list[float]:
    rng = np.random.default_rng(
        seed(f"KITE-SAT-AKB-confirmation-bootstrap-v1|{endpoint}")
    )
    indices = rng.integers(0, len(values), size=(100000, len(values)))
    means = np.mean(values[indices], axis=1)
    return [
        float(np.quantile(means, 0.025)),
        float(np.quantile(means, 0.975)),
    ]


def main() -> None:
    if OUTPUT.exists():
        raise RuntimeError("Refusing to overwrite AKB confirmation decision.")
    lock, lock_hash = validate_confirmation_lock(Path(__file__))
    if tuple(lock["confirmation_subject_order"]) != SUBJECTS:
        raise RuntimeError("Confirmation subjects differ from lock.")
    records, hashes, model_hashes = [], {}, {}
    ready_hashes = set()
    for subject in SUBJECTS:
        path = SUBJECT_ROOT / f"{subject}_AKB_confirmation_v1.json"
        record = json.loads(path.read_text(encoding="utf-8"))
        if (
            record["schema"] != "KITE-SAT-AKB-independent-confirmation-subject-v1"
            or record["status"] != "pass"
            or record["subject"] != subject
            or record["confirmation_implementation_lock_sha256"] != lock_hash
            or record["firewall"]["alternative_AKB_candidate_computed"]
            or record["firewall"]["alternative_patch_family_computed"]
            or record["firewall"]["paper_mutated"]
        ):
            raise RuntimeError(f"Invalid AKB confirmation artifact: {subject}")
        if [row["anchor"] for row in record["patches"]] != list(ANCHORS):
            raise RuntimeError(f"Patch order mismatch: {subject}")
        hashes[subject] = sha256(path)
        model_path = ROOT / record["model_archive"]["path"]
        model_hashes[subject] = sha256(model_path)
        if model_hashes[subject] != record["model_archive"]["sha256"]:
            raise RuntimeError(f"Confirmation model hash mismatch: {subject}")
        ready_hashes.add(record["preprocessing_ready"]["sha256"])
        records.append(record)
    if len(ready_hashes) != 1:
        raise RuntimeError("Confirmation subjects used different preprocessing-ready records.")
    intact_global = np.asarray(
        [record["intact_endpoints"]["global"]["mean_auc"] for record in records]
    )
    intact_akb = np.asarray(
        [record["intact_endpoints"]["AKB"]["mean_auc"] for record in records]
    )
    patch_global = np.asarray(
        [record["patch_endpoints"]["global"]["patch_average_auc"] for record in records]
    )
    patch_akb = np.asarray(
        [record["patch_endpoints"]["AKB"]["patch_average_auc"] for record in records]
    )
    intact_gain = intact_akb - intact_global
    patch_gain = patch_akb - patch_global
    intact_p = exact_one_sided(intact_gain)
    patch_raw_p = exact_one_sided(patch_gain)
    test1_rejected = intact_p <= 0.05
    test2_eligible = test1_rejected
    test2_rejected = test2_eligible and patch_raw_p <= 0.05
    if test1_rejected and test2_rejected:
        status = "intact_and_patch_confirmed"
    elif test1_rejected:
        status = "intact_only_confirmed"
    else:
        status = "not_confirmed"
    patch_metrics = []
    for patch_index, anchor in enumerate(ANCHORS):
        local_global = np.asarray(
            [
                record["patch_endpoints"]["global"]["patches"][patch_index]["mean_auc"]
                for record in records
            ]
        )
        local_akb = np.asarray(
            [
                record["patch_endpoints"]["AKB"]["patches"][patch_index]["mean_auc"]
                for record in records
            ]
        )
        patch_metrics.append(
            {
                "anchor": anchor,
                "mean_global_auc": float(np.mean(local_global)),
                "mean_AKB_auc": float(np.mean(local_akb)),
                "mean_gain_auc": float(np.mean(local_akb - local_global)),
            }
        )
    metrics = {
        "n": len(records),
        "mean_intact_global_auc": float(np.mean(intact_global)),
        "mean_intact_AKB_auc": float(np.mean(intact_akb)),
        "mean_intact_gain_auc": float(np.mean(intact_gain)),
        "median_intact_gain_auc": float(np.median(intact_gain)),
        "intact_gain_bootstrap_95_percent_ci": bootstrap_ci(
            intact_gain, "intact_gain"
        ),
        "intact_AKB_vs_chance_exact_one_sided_p": exact_one_sided(
            intact_akb - 0.5
        ),
        "participants_positive_intact_gain": int(np.sum(intact_gain > 0)),
        "story1_intact_AKB_group_mean": float(
            np.mean(
                [
                    record["intact_endpoints"]["AKB"]["story1_auc"]
                    for record in records
                ]
            )
        ),
        "story2_intact_AKB_group_mean": float(
            np.mean(
                [
                    record["intact_endpoints"]["AKB"]["story2_auc"]
                    for record in records
                ]
            )
        ),
        "mean_patch_average_global_auc": float(np.mean(patch_global)),
        "mean_patch_average_AKB_auc": float(np.mean(patch_akb)),
        "mean_patch_average_gain_auc": float(np.mean(patch_gain)),
        "median_patch_average_gain_auc": float(np.median(patch_gain)),
        "patch_gain_bootstrap_95_percent_ci": bootstrap_ci(
            patch_gain, "patch_gain"
        ),
        "patch_AKB_vs_chance_exact_one_sided_p": exact_one_sided(
            patch_akb - 0.5
        ),
        "participants_positive_patch_gain": int(np.sum(patch_gain > 0)),
        "minimum_patch_specific_group_mean_AKB_auc": float(
            min(row["mean_AKB_auc"] for row in patch_metrics)
        ),
        "minimum_patch_specific_group_mean_gain_auc": float(
            min(row["mean_gain_auc"] for row in patch_metrics)
        ),
    }
    payload = {
        "schema": "KITE-SAT-AKB-independent-confirmation-decision-v1",
        "status": status,
        "confirmation_implementation_lock_sha256": lock_hash,
        "preprocessing_ready_sha256": next(iter(ready_hashes)),
        "subject_order": list(SUBJECTS),
        "subject_artifact_sha256": hashes,
        "model_archive_sha256": model_hashes,
        "fixed_sequence_decision": {
            "familywise_one_sided_alpha": 0.05,
            "test_1_intact_gain": {
                "exact_one_sided_p": intact_p,
                "rejected": test1_rejected,
            },
            "test_2_patch_gain": {
                "eligible_after_test_1": test2_eligible,
                "raw_exact_one_sided_p": patch_raw_p,
                "rejected_with_confirmatory_standing": test2_rejected,
            },
        },
        "participant_results": [
            {
                "subject": record["subject"],
                "intact_global_auc": record["intact_endpoints"]["global"]["mean_auc"],
                "intact_AKB_auc": record["intact_endpoints"]["AKB"]["mean_auc"],
                "intact_gain_auc": record["intact_gain_auc"],
                "patch_global_auc": record["patch_endpoints"]["global"][
                    "patch_average_auc"
                ],
                "patch_AKB_auc": record["patch_endpoints"]["AKB"][
                    "patch_average_auc"
                ],
                "patch_gain_auc": record["patch_average_gain_auc"],
            }
            for record in records
        ],
        "patch_specific_group_metrics": patch_metrics,
        "metrics": metrics,
        "firewall": {
            "alternative_candidate_selected": False,
            "alternative_patch_selected": False,
            "paper_mutated": False,
        },
    }
    OUTPUT.parent.mkdir(parents=True, exist_ok=True)
    OUTPUT.write_text(json.dumps(payload, indent=2) + "\n", encoding="utf-8")
    observed = sha256(OUTPUT)
    OUTPUT.with_name(OUTPUT.name + ".sha256").write_text(
        f"{observed}  {OUTPUT.name}\n", encoding="ascii"
    )
    print(
        json.dumps(
            {
                "status": status,
                "fixed_sequence": payload["fixed_sequence_decision"],
                "metrics": metrics,
                "sha256": observed,
            }
        )
    )


if __name__ == "__main__":
    main()
```
