## Supplementary material for "Falsifiable substitution tests reveal task-structured neural evidence for auditory attention": Article_Analysis_Source_Code: run_kite_sat_akb_confirmation_subject_v1.py

```
"""Run the single frozen AKB method for one independent confirmation subject."""

from __future__ import annotations

import argparse
import json
from pathlib import Path

import numpy as np
from scipy.signal import resample_poly

from download_kite_st_hd_confirmation_v1 import SUBJECTS
from kite_sat_adaptive_broad_v1 import adaptive_score, reliability
from kite_sat_backward_v1 import block_design
from kite_sat_confirmation_lock_utils_v1 import ROOT, sha256, validate_confirmation_lock
from kite_sat_geometry_v1 import cell_delta_contributions, partition_ids
from kite_sat_locked_decoder_v1 import fit_locked_ridge
from kite_sat_patch_erasure_v1 import ANCHORS, erased_cell_delta_contributions, fixed_patch_masks
from kite_st_hd_core_v1 import load_public_geometry
from kite_st_hd_statistics_v1 import binary_auc


CONDITIONS = ("1L", "1R", "2L", "2R")
PREPROCESSED_ROOT = (
    ROOT / "data" / "derived" / "kite_st_hd_zenodo4518754"
    / "confirmation_preprocessed_v1"
)
MANIFEST_ROOT = ROOT / "results" / "kite_st_hd" / "confirmation_v1" / "subjects"
READY = (
    ROOT / "results" / "kite_st_hd" / "confirmation_v1"
    / "confirmation_preprocessing_ready_v1.json"
)
AUDIO_CACHE = (
    ROOT / "data" / "derived" / "kite_st_hd_zenodo4518754"
    / "kite_sat_audio_features_v1.npz"
)
COORDINATES = (
    ROOT / "downloads" / "candidate_audit" / "zenodo4518754_support"
    / "misc" / "misc" / "eeg255ch_locs.csv"
)
OUTPUT_ROOT = ROOT / "results" / "kite_sat" / "akb_confirmation_v1" / "subjects"
MODEL_ROOT = (
    ROOT / "data" / "derived" / "kite_st_hd_zenodo4518754"
    / "kite_sat_confirmation_models_v1"
)
PHYSICAL = {
    "1L": ("part1_track1", "part1_track2"),
    "1R": ("part1_track1", "part1_track2"),
    "2L": ("part2_track2", "part2_track1"),
    "2R": ("part2_track2", "part2_track1"),
}


def main() -> None:
    parser = argparse.ArgumentParser()
    parser.add_argument("subject", choices=SUBJECTS)
    args = parser.parse_args()
    subject = args.subject
    output = OUTPUT_ROOT / f"{subject}_AKB_confirmation_v1.json"
    model_output = MODEL_ROOT / f"{subject}_AKB_confirmation_models_v1.npz"
    if output.exists() or model_output.exists():
        raise RuntimeError(f"Refusing to overwrite AKB confirmation: {subject}")
    lock, lock_hash = validate_confirmation_lock(Path(__file__))
    if tuple(lock["confirmation_subject_order"]) != SUBJECTS:
        raise RuntimeError("Confirmation subject list differs from lock.")
    ready = json.loads(READY.read_text(encoding="utf-8"))
    if (
        ready["status"] != "pass"
        or ready["subject_order"] != list(SUBJECTS)
        or ready["subject_count"] != len(SUBJECTS)
        or not ready["all_four_recordings_present"]
        or ready["confirmation_implementation_lock_sha256"] != lock_hash
        or ready["attention_outcome_computed"]
        or ready["AKB_score_computed"]
    ):
        raise RuntimeError("All-subject confirmation preprocessing is not ready.")
    ready_subject = next(
        item for item in ready["subjects"] if item["subject"] == subject
    )
    manifest_path = MANIFEST_ROOT / f"{subject}_production_manifest_v1.json"
    manifest = json.loads(manifest_path.read_text(encoding="utf-8"))
    if (
        sha256(manifest_path) != ready_subject["manifest_sha256"]
        or manifest["status"] != "pass"
        or manifest["subject"] != subject
    ):
        raise RuntimeError("Confirmation preprocessing manifest mismatch.")
    preprocessing = {
        item["recording"].rsplit("_", 1)[-1]: item
        for item in manifest["preprocessing_records"]
    }
    with np.load(AUDIO_CACHE, allow_pickle=False) as saved:
        full_audio = {name: saved[name] for name in saved.files}
    geometry = load_public_geometry(COORDINATES)
    s16 = partition_ids(geometry.xy)["S16"]
    patches, patch_members = fixed_patch_masks(geometry.labels, geometry.xy)
    designs: dict[str, list[np.ndarray]] = {}
    audio_blocks: dict[str, list[tuple[np.ndarray, np.ndarray]]] = {}
    attended: dict[str, list[np.ndarray]] = {}
    input_hashes, offsets = {}, {}
    for condition in CONDITIONS:
        path = (
            PREPROCESSED_ROOT / subject
            / f"{subject}_AAD_{condition}_preprocessed_v1.npz"
        )
        input_hashes[condition] = sha256(path)
        if input_hashes[condition] != preprocessing[condition]["output"]["sha256"]:
            raise RuntimeError(f"Confirmation input hash mismatch: {subject} {condition}")
        with np.load(path, allow_pickle=False) as saved:
            eeg = saved["eeg"]
        record = preprocessing[condition]
        offset_seconds = (
            record["crop"]["padded_start"]
            + 3000
            - record["trigger_train"]["first_sample"]
        ) / 1000.0
        offsets[condition] = offset_seconds
        offset = int(round(offset_seconds * 100))
        left_name, right_name = PHYSICAL[condition]
        designs[condition], audio_blocks[condition], attended[condition] = [], [], []
        for block_index in range(12):
            first = offset + block_index * 3000
            stop = first + 3000
            left100 = full_audio[left_name][first:stop]
            right100 = full_audio[right_name][first:stop]
            design, left_features = block_design(
                eeg[block_index * 3000 : (block_index + 1) * 3000], left100
            )
            right_features = resample_poly(right100, 1, 5, axis=0)[:595]
            left, right = left_features[:, 0], right_features[:, 0]
            designs[condition].append(design)
            audio_blocks[condition].append((left, right))
            attended[condition].append(left if condition.endswith("L") else right)

    intact_scores = {
        estimator: {condition: [] for condition in CONDITIONS}
        for estimator in ("global", "AKB")
    }
    patch_scores = {
        estimator: {
            condition: [[] for _ in ANCHORS]
            for condition in CONDITIONS
        }
        for estimator in ("global", "AKB")
    }
    fold_reliabilities, model_arrays = {}, {}
    for train_story, test_story in (("1", "2"), ("2", "1")):
        train_conditions = (f"{train_story}L", f"{train_story}R")
        held_aligned = []
        for fit_parity in (0, 1):
            held_parity = 1 - fit_parity
            calibration_model = fit_locked_ridge(
                [
                    designs[condition][index]
                    for condition in train_conditions
                    for index in range(12)
                    if index % 2 == fit_parity
                ],
                [
                    attended[condition][index]
                    for condition in train_conditions
                    for index in range(12)
                    if index % 2 == fit_parity
                ],
            )
            for condition in train_conditions:
                alignment = 1.0 if condition.endswith("L") else -1.0
                for index in range(12):
                    if index % 2 != held_parity:
                        continue
                    left, right = audio_blocks[condition][index]
                    delta, _ = cell_delta_contributions(
                        designs[condition][index],
                        calibration_model,
                        left,
                        right,
                        s16,
                    )
                    held_aligned.append(alignment * delta)
        fold_reliability = reliability(np.asarray(held_aligned))
        fold_reliabilities[f"train_story_{train_story}"] = [
            float(value) for value in fold_reliability
        ]
        full_model = fit_locked_ridge(
            [
                designs[condition][index]
                for condition in train_conditions
                for index in range(12)
            ],
            [
                attended[condition][index]
                for condition in train_conditions
                for index in range(12)
            ],
        )
        for name in ("x_mean", "x_scale", "weights"):
            model_arrays[f"train_story_{train_story}_{name}"] = np.asarray(
                full_model[name]
            )
        for name in ("y_mean", "alpha", "gram_energy"):
            model_arrays[f"train_story_{train_story}_{name}"] = np.asarray(
                full_model[name]
            )
        for condition in (f"{test_story}L", f"{test_story}R"):
            for design, (left, right) in zip(
                designs[condition], audio_blocks[condition]
            ):
                delta, global_margin = cell_delta_contributions(
                    design, full_model, left, right, s16
                )
                intact_scores["global"][condition].append(global_margin)
                intact_scores["AKB"][condition].append(
                    adaptive_score(delta, fold_reliability, "softmax2")
                )
                erased_delta, erased_global = erased_cell_delta_contributions(
                    design, full_model, left, right, s16, patches
                )
                for patch_index in range(len(ANCHORS)):
                    patch_scores["global"][condition][patch_index].append(
                        float(erased_global[patch_index])
                    )
                    patch_scores["AKB"][condition][patch_index].append(
                        adaptive_score(
                            erased_delta[patch_index],
                            fold_reliability,
                            "softmax2",
                        )
                    )
    intact_endpoints = {}
    for estimator in ("global", "AKB"):
        story1 = binary_auc(
            intact_scores[estimator]["1L"], intact_scores[estimator]["1R"]
        )
        story2 = binary_auc(
            intact_scores[estimator]["2L"], intact_scores[estimator]["2R"]
        )
        intact_endpoints[estimator] = {
            "story1_auc": story1,
            "story2_auc": story2,
            "mean_auc": (story1 + story2) / 2,
        }
    patch_endpoints = {}
    for estimator in ("global", "AKB"):
        rows = []
        for patch_index, anchor in enumerate(ANCHORS):
            story1 = binary_auc(
                patch_scores[estimator]["1L"][patch_index],
                patch_scores[estimator]["1R"][patch_index],
            )
            story2 = binary_auc(
                patch_scores[estimator]["2L"][patch_index],
                patch_scores[estimator]["2R"][patch_index],
            )
            rows.append(
                {
                    "anchor": anchor,
                    "story1_auc": story1,
                    "story2_auc": story2,
                    "mean_auc": (story1 + story2) / 2,
                }
            )
        patch_endpoints[estimator] = {
            "patches": rows,
            "patch_average_auc": float(np.mean([row["mean_auc"] for row in rows])),
        }
    MODEL_ROOT.mkdir(parents=True, exist_ok=True)
    np.savez_compressed(model_output, **model_arrays)
    model_hash = sha256(model_output)
    model_output.with_name(model_output.name + ".sha256").write_text(
        f"{model_hash}  {model_output.name}\n", encoding="ascii"
    )
    payload = {
        "schema": "KITE-SAT-AKB-independent-confirmation-subject-v1",
        "status": "pass",
        "subject": subject,
        "confirmation_implementation_lock_sha256": lock_hash,
        "preprocessing_ready": {
            "path": READY.relative_to(ROOT).as_posix(),
            "sha256": sha256(READY),
        },
        "upstream_manifest": {
            "path": manifest_path.relative_to(ROOT).as_posix(),
            "sha256": sha256(manifest_path),
        },
        "audio_cache_sha256": sha256(AUDIO_CACHE),
        "preprocessed_sha256": input_hashes,
        "offset_seconds": offsets,
        "patches": [
            {
                "anchor": anchor,
                "removed_count": int(np.sum(patches[index])),
                "removed_labels": list(patch_members[index]),
            }
            for index, anchor in enumerate(ANCHORS)
        ],
        "intact_endpoints": intact_endpoints,
        "intact_gain_auc": (
            intact_endpoints["AKB"]["mean_auc"]
            - intact_endpoints["global"]["mean_auc"]
        ),
        "patch_endpoints": patch_endpoints,
        "patch_average_gain_auc": (
            patch_endpoints["AKB"]["patch_average_auc"]
            - patch_endpoints["global"]["patch_average_auc"]
        ),
        "intact_condition_scores": intact_scores,
        "patch_condition_scores": patch_scores,
        "fold_reliabilities": fold_reliabilities,
        "model_archive": {
            "path": model_output.relative_to(ROOT).as_posix(),
            "bytes": model_output.stat().st_size,
            "sha256": model_hash,
        },
        "firewall": {
            "alternative_AKB_candidate_computed": False,
            "alternative_patch_family_computed": False,
            "paper_mutated": False,
        },
    }
    OUTPUT_ROOT.mkdir(parents=True, exist_ok=True)
    output.write_text(json.dumps(payload, indent=2) + "\n", encoding="utf-8")
    observed = sha256(output)
    output.with_name(output.name + ".sha256").write_text(
        f"{observed}  {output.name}\n", encoding="ascii"
    )
    print(
        json.dumps(
            {
                "subject": subject,
                "intact_global_auc": intact_endpoints["global"]["mean_auc"],
                "intact_AKB_auc": intact_endpoints["AKB"]["mean_auc"],
                "intact_gain": payload["intact_gain_auc"],
                "patch_global_auc": patch_endpoints["global"]["patch_average_auc"],
                "patch_AKB_auc": patch_endpoints["AKB"]["patch_average_auc"],
                "patch_gain": payload["patch_average_gain_auc"],
                "sha256": observed,
            }
        )
    )


if __name__ == "__main__":
    main()
```
