## Supplementary material for "Falsifiable substitution tests reveal task-structured neural evidence for auditory attention": Article_Analysis_Source_Code: run_kite_sat_akb_mechanism_group_v1.py

```
"""Frozen group decision for AKB direction-delay mechanism controls."""

from __future__ import annotations

import itertools
import json
from pathlib import Path

import numpy as np

from download_kite_st_hd_confirmation_v1 import SUBJECTS
from kite_sat_mechanism_lock_utils_v1 import ROOT, sha256, validate_mechanism_lock


SUBJECT_ROOT = (
    ROOT / "results" / "kite_sat" / "akb_direction_delay_mechanism_v1"
    / "subjects"
)
OUTPUT = SUBJECT_ROOT.parent / "mechanism_decision_v1.json"
PRIMARY_DECISION = (
    ROOT / "results" / "kite_sat" / "akb_confirmation_v1"
    / "confirmation_decision_v1.json"
)


def exact_one_sided(values: np.ndarray) -> float:
    observed = float(np.mean(values))
    exceed = sum(
        float(np.mean(values * np.asarray(signs))) >= observed - 1e-15
        for signs in itertools.product((-1.0, 1.0), repeat=len(values))
    )
    return exceed / (2 ** len(values))


def main() -> None:
    if OUTPUT.exists():
        raise RuntimeError("Refusing to overwrite AKB mechanism decision.")
    lock, lock_hash = validate_mechanism_lock(Path(__file__))
    if tuple(lock["confirmation_subject_order"]) != SUBJECTS:
        raise RuntimeError("Mechanism subject order differs from lock.")
    primary = json.loads(PRIMARY_DECISION.read_text(encoding="utf-8"))
    primary_confirmed = primary["status"] in (
        "intact_and_patch_confirmed",
        "intact_only_confirmed",
    )
    records, hashes = [], {}
    for subject in SUBJECTS:
        path = SUBJECT_ROOT / f"{subject}_direction_delay_mechanism_v1.json"
        record = json.loads(path.read_text(encoding="utf-8"))
        if (
            record["status"] != "pass"
            or record["subject"] != subject
            or record["mechanism_implementation_lock_sha256"] != lock_hash
            or record["firewall"]["control_selected_by_outcome"]
            or record["firewall"]["primary_method_changed"]
            or record["firewall"]["paper_mutated"]
        ):
            raise RuntimeError(f"Invalid AKB mechanism artifact: {subject}")
        hashes[subject] = sha256(path)
        records.append(record)
    decoding = np.asarray(
        [record["intact_identity_advantage"] for record in records]
    )
    reliability = np.asarray(
        [record["reliability_identity_advantage"] for record in records]
    )
    patch = np.asarray(
        [record["patch_identity_advantage"] for record in records]
    )
    decoding_p = exact_one_sided(decoding)
    decoding_rejected = decoding_p <= 0.05
    reliability_eligible = decoding_rejected
    reliability_p = exact_one_sided(reliability)
    reliability_rejected = reliability_eligible and reliability_p <= 0.05
    mechanism_supported = primary_confirmed and decoding_rejected
    if mechanism_supported and reliability_rejected:
        status = "full_direction_delay_stability_supported"
    elif mechanism_supported:
        status = "direction_delay_decoding_mechanism_supported"
    else:
        status = "mechanism_not_supported"
    broadness_rows = [
        record["broadness"][story]
        for record in records
        for story in ("train_story_1", "train_story_2")
    ]
    metrics = {
        "n": len(records),
        "mean_intact_identity_advantage_auc": float(np.mean(decoding)),
        "median_intact_identity_advantage_auc": float(np.median(decoding)),
        "participants_positive_intact_identity_advantage": int(np.sum(decoding > 0)),
        "mean_angular_only_advantage_auc": float(
            np.mean(
                [record["intact_angular_only_advantage"] for record in records]
            )
        ),
        "mean_delay_swapped_advantage_auc": float(
            np.mean(
                [record["intact_delay_swapped_advantage"] for record in records]
            )
        ),
        "mean_patch_identity_advantage_auc": float(np.mean(patch)),
        "mean_reliability_identity_advantage": float(np.mean(reliability)),
        "participants_positive_reliability_identity_advantage": int(
            np.sum(reliability > 0)
        ),
        "mean_effective_cell_number": float(
            np.mean([row["effective_cell_number"] for row in broadness_rows])
        ),
        "mean_normalized_cell_entropy": float(
            np.mean([row["normalized_cell_entropy"] for row in broadness_rows])
        ),
        "mean_maximum_cell_mass": float(
            np.mean([row["maximum_cell_mass"] for row in broadness_rows])
        ),
        "mean_effective_angular_sector_number": float(
            np.mean(
                [
                    row["effective_angular_sector_number"]
                    for row in broadness_rows
                ]
            )
        ),
    }
    payload = {
        "schema": "KITE-SAT-AKB-direction-delay-mechanism-decision-v1",
        "status": status,
        "mechanism_implementation_lock_sha256": lock_hash,
        "primary_confirmation_status": primary["status"],
        "primary_confirmation_sha256": sha256(PRIMARY_DECISION),
        "subject_order": list(SUBJECTS),
        "subject_artifact_sha256": hashes,
        "fixed_sequence": {
            "primary_intact_AKB_confirmed": primary_confirmed,
            "decoding_identity_advantage": {
                "exact_one_sided_p": decoding_p,
                "rejected": decoding_rejected,
                "mechanistic_standing": mechanism_supported,
            },
            "reliability_identity_advantage": {
                "eligible_after_decoding_test": reliability_eligible,
                "raw_exact_one_sided_p": reliability_p,
                "rejected_with_mechanistic_standing": reliability_rejected,
            },
        },
        "participant_results": [
            {
                "subject": record["subject"],
                "intact_identity_advantage_auc": record[
                    "intact_identity_advantage"
                ],
                "angular_only_advantage_auc": record[
                    "intact_angular_only_advantage"
                ],
                "delay_swapped_advantage_auc": record[
                    "intact_delay_swapped_advantage"
                ],
                "patch_identity_advantage_auc": record[
                    "patch_identity_advantage"
                ],
                "reliability_identity_advantage": record[
                    "reliability_identity_advantage"
                ],
            }
            for record in records
        ],
        "metrics": metrics,
        "firewall": {
            "control_selected_by_outcome": False,
            "primary_decision_changed": False,
            "paper_mutated": False,
        },
    }
    OUTPUT.write_text(json.dumps(payload, indent=2) + "\n", encoding="utf-8")
    observed = sha256(OUTPUT)
    OUTPUT.with_name(OUTPUT.name + ".sha256").write_text(
        f"{observed}  {OUTPUT.name}\n", encoding="ascii"
    )
    print(
        json.dumps(
            {
                "status": status,
                "fixed_sequence": payload["fixed_sequence"],
                "metrics": metrics,
                "sha256": observed,
            }
        )
    )


if __name__ == "__main__":
    main()
```
