## Supplementary material for "Falsifiable substitution tests reveal task-structured neural evidence for auditory attention": Article_Analysis_Source_Code: run_kite_sat_akb_mechanism_subject_v1.py

```
"""Capacity-matched direction-delay controls for one confirmation subject."""

from __future__ import annotations

import argparse
import json
from pathlib import Path

import numpy as np
from scipy.signal import resample_poly

from download_kite_st_hd_confirmation_v1 import SUBJECTS
from kite_sat_adaptive_broad_v1 import family_weights
from kite_sat_backward_v1 import block_design
from kite_sat_direction_delay_controls_v1 import TRANSFORMS, transformed_matrix
from kite_sat_geometry_v1 import cell_delta_contributions, partition_ids
from kite_sat_mechanism_lock_utils_v1 import ROOT, sha256, validate_mechanism_lock
from kite_sat_patch_erasure_v1 import ANCHORS, erased_cell_delta_contributions, fixed_patch_masks
from kite_st_hd_core_v1 import load_public_geometry
from kite_st_hd_statistics_v1 import binary_auc


CONDITIONS = ("1L", "1R", "2L", "2R")
PREPROCESSED_ROOT = (
    ROOT / "data" / "derived" / "kite_st_hd_zenodo4518754"
    / "confirmation_preprocessed_v1"
)
MANIFEST_ROOT = ROOT / "results" / "kite_st_hd" / "confirmation_v1" / "subjects"
PRIMARY_ROOT = ROOT / "results" / "kite_sat" / "akb_confirmation_v1"
MODEL_ROOT = (
    ROOT / "data" / "derived" / "kite_st_hd_zenodo4518754"
    / "kite_sat_confirmation_models_v1"
)
AUDIO_CACHE = (
    ROOT / "data" / "derived" / "kite_st_hd_zenodo4518754"
    / "kite_sat_audio_features_v1.npz"
)
COORDINATES = (
    ROOT / "downloads" / "candidate_audit" / "zenodo4518754_support"
    / "misc" / "misc" / "eeg255ch_locs.csv"
)
OUTPUT_ROOT = (
    ROOT / "results" / "kite_sat" / "akb_direction_delay_mechanism_v1"
    / "subjects"
)
PHYSICAL = {
    "1L": ("part1_track1", "part1_track2"),
    "1R": ("part1_track1", "part1_track2"),
    "2L": ("part2_track2", "part2_track1"),
    "2R": ("part2_track2", "part2_track1"),
}


def centered_correlation(first: np.ndarray, second: np.ndarray) -> float:
    a = np.asarray(first, dtype=np.float64) - np.mean(first)
    b = np.asarray(second, dtype=np.float64) - np.mean(second)
    return float(
        np.sum(a * b)
        / max(float(np.sqrt(np.sum(a * a) * np.sum(b * b))), 1e-12)
    )


def broadness(reliability: np.ndarray) -> dict[str, float]:
    weights = family_weights(reliability, "softmax2")
    probability = weights / np.sum(weights)
    entropy = float(-np.sum(probability * np.log(probability + 1e-300)))
    angular = probability[:8] + probability[8:]
    angular_entropy = float(-np.sum(angular * np.log(angular + 1e-300)))
    return {
        "effective_cell_number": float(np.exp(entropy)),
        "normalized_cell_entropy": float(entropy / np.log(16)),
        "maximum_cell_mass": float(np.max(probability)),
        "effective_angular_sector_number": float(np.exp(angular_entropy)),
    }


def main() -> None:
    parser = argparse.ArgumentParser()
    parser.add_argument("subject", choices=SUBJECTS)
    args = parser.parse_args()
    subject = args.subject
    output = OUTPUT_ROOT / f"{subject}_direction_delay_mechanism_v1.json"
    if output.exists():
        raise RuntimeError(f"Refusing to overwrite mechanism result: {subject}")
    lock, mechanism_lock_hash = validate_mechanism_lock(Path(__file__))
    if tuple(lock["confirmation_subject_order"]) != SUBJECTS:
        raise RuntimeError("Mechanism subject order differs from lock.")
    decision_path = PRIMARY_ROOT / "confirmation_decision_v1.json"
    audit_path = PRIMARY_ROOT / "confirmation_outcome_audit_v1.json"
    decision = json.loads(decision_path.read_text(encoding="utf-8"))
    audit = json.loads(audit_path.read_text(encoding="utf-8"))
    if (
        audit["status"] != "pass"
        or audit["confirmation_decision_sha256"] != sha256(decision_path)
        or decision["subject_order"] != list(SUBJECTS)
    ):
        raise RuntimeError("Audited primary confirmation decision is unavailable.")
    primary_path = PRIMARY_ROOT / "subjects" / f"{subject}_AKB_confirmation_v1.json"
    primary = json.loads(primary_path.read_text(encoding="utf-8"))
    if sha256(primary_path) != decision["subject_artifact_sha256"][subject]:
        raise RuntimeError("Primary confirmation subject hash mismatch.")
    manifest_path = MANIFEST_ROOT / f"{subject}_production_manifest_v1.json"
    manifest = json.loads(manifest_path.read_text(encoding="utf-8"))
    preprocessing = {
        item["recording"].rsplit("_", 1)[-1]: item
        for item in manifest["preprocessing_records"]
    }
    model_path = MODEL_ROOT / f"{subject}_AKB_confirmation_models_v1.npz"
    if sha256(model_path) != primary["model_archive"]["sha256"]:
        raise RuntimeError("Primary confirmation model hash mismatch.")
    with np.load(model_path, allow_pickle=False) as saved:
        models = {name: saved[name] for name in saved.files}
    with np.load(AUDIO_CACHE, allow_pickle=False) as saved:
        full_audio = {name: saved[name] for name in saved.files}
    geometry = load_public_geometry(COORDINATES)
    s16 = partition_ids(geometry.xy)["S16"]
    patches, _ = fixed_patch_masks(geometry.labels, geometry.xy)
    designs: dict[str, list[np.ndarray]] = {}
    audio_blocks: dict[str, list[tuple[np.ndarray, np.ndarray]]] = {}
    for condition in CONDITIONS:
        path = (
            PREPROCESSED_ROOT / subject
            / f"{subject}_AAD_{condition}_preprocessed_v1.npz"
        )
        if sha256(path) != primary["preprocessed_sha256"][condition]:
            raise RuntimeError(f"Mechanism input hash mismatch: {subject} {condition}")
        with np.load(path, allow_pickle=False) as saved:
            eeg = saved["eeg"]
        record = preprocessing[condition]
        offset = int(
            round(
                (
                    record["crop"]["padded_start"]
                    + 3000
                    - record["trigger_train"]["first_sample"]
                )
                / 10.0
            )
        )
        left_name, right_name = PHYSICAL[condition]
        designs[condition], audio_blocks[condition] = [], []
        for block_index in range(12):
            first = offset + block_index * 3000
            stop = first + 3000
            left100 = full_audio[left_name][first:stop]
            right100 = full_audio[right_name][first:stop]
            design, left_features = block_design(
                eeg[block_index * 3000 : (block_index + 1) * 3000], left100
            )
            right_features = resample_poly(right100, 1, 5, axis=0)[:595]
            designs[condition].append(design)
            audio_blocks[condition].append(
                (left_features[:, 0], right_features[:, 0])
            )
    intact_scores = {
        condition: [[] for _ in TRANSFORMS] for condition in CONDITIONS
    }
    patch_scores = {
        condition: [
            [[] for _ in TRANSFORMS] for _ in ANCHORS
        ]
        for condition in CONDITIONS
    }
    reliability_vectors = {}
    for train_story, test_story in (("1", "2"), ("2", "1")):
        reliability = np.asarray(
            primary["fold_reliabilities"][f"train_story_{train_story}"],
            dtype=np.float64,
        )
        reliability_vectors[f"train_story_{train_story}"] = reliability
        weight_matrix = transformed_matrix(
            family_weights(reliability, "softmax2")
        )
        model = {
            "x_mean": models[f"train_story_{train_story}_x_mean"],
            "x_scale": models[f"train_story_{train_story}_x_scale"],
            "weights": models[f"train_story_{train_story}_weights"],
            "y_mean": float(models[f"train_story_{train_story}_y_mean"]),
        }
        for condition in (f"{test_story}L", f"{test_story}R"):
            for design, (left, right) in zip(
                designs[condition], audio_blocks[condition]
            ):
                delta, _ = cell_delta_contributions(
                    design, model, left, right, s16
                )
                transform_scores = weight_matrix @ delta
                for transform_index, score in enumerate(transform_scores):
                    intact_scores[condition][transform_index].append(float(score))
                erased_delta, _ = erased_cell_delta_contributions(
                    design, model, left, right, s16, patches
                )
                erased_scores = erased_delta @ weight_matrix.T
                for patch_index in range(len(ANCHORS)):
                    for transform_index in range(len(TRANSFORMS)):
                        patch_scores[condition][patch_index][
                            transform_index
                        ].append(float(erased_scores[patch_index, transform_index]))
    transform_results = []
    for transform_index, (shift, swap) in enumerate(TRANSFORMS):
        story1 = binary_auc(
            intact_scores["1L"][transform_index],
            intact_scores["1R"][transform_index],
        )
        story2 = binary_auc(
            intact_scores["2L"][transform_index],
            intact_scores["2R"][transform_index],
        )
        patch_aucs = []
        for patch_index in range(len(ANCHORS)):
            patch_story1 = binary_auc(
                patch_scores["1L"][patch_index][transform_index],
                patch_scores["1R"][patch_index][transform_index],
            )
            patch_story2 = binary_auc(
                patch_scores["2L"][patch_index][transform_index],
                patch_scores["2R"][patch_index][transform_index],
            )
            patch_aucs.append((patch_story1 + patch_story2) / 2)
        transform_results.append(
            {
                "angular_shift": shift,
                "delay_swap": bool(swap),
                "story1_auc": story1,
                "story2_auc": story2,
                "intact_mean_auc": (story1 + story2) / 2,
                "patch_average_auc": float(np.mean(patch_aucs)),
            }
        )
    identity = transform_results[0]
    if (
        abs(identity["intact_mean_auc"] - primary["intact_endpoints"]["AKB"]["mean_auc"])
        > 1e-12
        or abs(
            identity["patch_average_auc"]
            - primary["patch_endpoints"]["AKB"]["patch_average_auc"]
        )
        > 1e-12
    ):
        raise RuntimeError("Mechanism identity did not reproduce primary AKB.")
    intact_controls = np.asarray(
        [row["intact_mean_auc"] for row in transform_results[1:]]
    )
    patch_controls = np.asarray(
        [row["patch_average_auc"] for row in transform_results[1:]]
    )
    r1 = reliability_vectors["train_story_1"]
    r2 = reliability_vectors["train_story_2"]
    transformed_r2 = transformed_matrix(r2)
    reliability_correlations = np.asarray(
        [centered_correlation(r1, row) for row in transformed_r2]
    )
    payload = {
        "schema": "KITE-SAT-AKB-direction-delay-mechanism-subject-v1",
        "status": "pass",
        "subject": subject,
        "mechanism_implementation_lock_sha256": mechanism_lock_hash,
        "primary_confirmation": {
            "decision_status": decision["status"],
            "decision_sha256": sha256(decision_path),
            "outcome_audit_sha256": sha256(audit_path),
            "subject_artifact_sha256": sha256(primary_path),
            "model_archive_sha256": sha256(model_path),
        },
        "transform_order": [
            {"angular_shift": shift, "delay_swap": bool(swap)}
            for shift, swap in TRANSFORMS
        ],
        "intact_condition_scores": intact_scores,
        "patch_condition_scores": patch_scores,
        "transform_results": transform_results,
        "intact_identity_advantage": float(
            identity["intact_mean_auc"] - np.mean(intact_controls)
        ),
        "intact_angular_only_advantage": float(
            identity["intact_mean_auc"]
            - np.mean(
                [row["intact_mean_auc"] for row in transform_results[1:8]]
            )
        ),
        "intact_delay_swapped_advantage": float(
            identity["intact_mean_auc"]
            - np.mean(
                [row["intact_mean_auc"] for row in transform_results[8:]]
            )
        ),
        "patch_identity_advantage": float(
            identity["patch_average_auc"] - np.mean(patch_controls)
        ),
        "reliability_correlations": [
            float(value) for value in reliability_correlations
        ],
        "reliability_vectors": {
            key: [float(value) for value in vector]
            for key, vector in reliability_vectors.items()
        },
        "reliability_identity_advantage": float(
            reliability_correlations[0] - np.mean(reliability_correlations[1:])
        ),
        "broadness": {
            "train_story_1": broadness(r1),
            "train_story_2": broadness(r2),
        },
        "firewall": {
            "control_selected_by_outcome": False,
            "primary_method_changed": False,
            "paper_mutated": False,
        },
    }
    OUTPUT_ROOT.mkdir(parents=True, exist_ok=True)
    output.write_text(json.dumps(payload, indent=2) + "\n", encoding="utf-8")
    observed = sha256(output)
    output.with_name(output.name + ".sha256").write_text(
        f"{observed}  {output.name}\n", encoding="ascii"
    )
    print(
        json.dumps(
            {
                "subject": subject,
                "intact_identity_advantage": payload["intact_identity_advantage"],
                "reliability_identity_advantage": payload[
                    "reliability_identity_advantage"
                ],
                "patch_identity_advantage": payload["patch_identity_advantage"],
                "sha256": observed,
            }
        )
    )


if __name__ == "__main__":
    main()
```
