## Supplementary material for "Falsifiable substitution tests reveal task-structured neural evidence for auditory attention": Article_Analysis_Source_Code: run_kite_sat_akb_patch_group_v1.py

```
"""Group gate for the locked KITE-SAT AKB contiguous-patch experiment."""

from __future__ import annotations

import hashlib
import itertools
import json
from pathlib import Path

import numpy as np


ROOT = Path(__file__).resolve().parents[1]
PROTOCOL = ROOT / "docs" / "kite_sat_akb_patch_resilience_development_protocol_v1.json"
PROTOCOL_HASH = "ff32e236996b761cc059e03f7f775aac84cd3322cab2d220cf44efff219d1cb4"
SUBJECTS = (
    "S4", "S24", "S11", "S23", "S14", "S29",
    "S19", "S17", "S12", "S25", "S27", "S5",
)
ANCHORS = ("TP7H", "PO3", "PO4", "TP8H", "FT8H", "AF4", "AF3", "FT7H")
SUBJECT_ROOT = (
    ROOT / "results" / "kite_sat" / "akb_patch_resilience_development_v1" / "subjects"
)
OUTPUT = SUBJECT_ROOT.parent / "development_gate_v1.json"


def sha256(path: Path) -> str:
    digest = hashlib.sha256()
    with path.open("rb") as stream:
        for chunk in iter(lambda: stream.read(8 * 1024 * 1024), b""):
            digest.update(chunk)
    return digest.hexdigest()


def exact_one_sided(values: np.ndarray) -> float:
    observed = float(np.mean(values))
    exceed = sum(
        float(np.mean(values * np.asarray(signs))) >= observed - 1e-15
        for signs in itertools.product((-1.0, 1.0), repeat=len(values))
    )
    return exceed / (2 ** len(values))


def stable_seed(text: str) -> int:
    return int.from_bytes(hashlib.sha256(text.encode("utf-8")).digest()[:8], "little")


def main() -> None:
    if OUTPUT.exists():
        raise RuntimeError("Refusing to overwrite patch-resilience development gate.")
    if sha256(PROTOCOL) != PROTOCOL_HASH:
        raise RuntimeError("Patch-resilience protocol hash mismatch.")
    records, hashes = [], {}
    for subject in SUBJECTS:
        path = SUBJECT_ROOT / f"{subject}_patch_resilience_v1.json"
        record = json.loads(path.read_text(encoding="utf-8"))
        if (
            record["status"] != "pass"
            or record["subject"] != subject
            or record["protocol_sha256"] != PROTOCOL_HASH
            or record["firewall"]["patch_variant_computed"]
            or record["firewall"]["confirmation_subject_read"]
        ):
            raise RuntimeError(f"Invalid patch artifact for {subject}.")
        for estimator in ("global", "AKB"):
            if [row["anchor"] for row in record["endpoints"][estimator]["patches"]] != list(ANCHORS):
                raise RuntimeError(f"Patch order mismatch for {subject}.")
        hashes[subject] = sha256(path)
        records.append(record)
    global_auc = np.asarray(
        [record["endpoints"]["global"]["patch_average_auc"] for record in records]
    )
    akb_auc = np.asarray(
        [record["endpoints"]["AKB"]["patch_average_auc"] for record in records]
    )
    gain = akb_auc - global_auc
    patch_metrics = []
    for patch_index, anchor in enumerate(ANCHORS):
        patch_global = np.asarray(
            [
                record["endpoints"]["global"]["patches"][patch_index]["mean_auc"]
                for record in records
            ]
        )
        patch_akb = np.asarray(
            [
                record["endpoints"]["AKB"]["patches"][patch_index]["mean_auc"]
                for record in records
            ]
        )
        patch_metrics.append(
            {
                "anchor": anchor,
                "mean_global_auc": float(np.mean(patch_global)),
                "mean_AKB_auc": float(np.mean(patch_akb)),
                "mean_gain_auc": float(np.mean(patch_akb - patch_global)),
            }
        )
    rng = np.random.default_rng(
        stable_seed("KITE-SAT-AKB-patch-resilience-development-bootstrap-v1")
    )
    indices = rng.integers(0, len(gain), size=(20000, len(gain)))
    boot = np.mean(gain[indices], axis=1)
    metrics = {
        "n": len(gain),
        "mean_patch_average_global_auc": float(np.mean(global_auc)),
        "mean_patch_average_AKB_auc": float(np.mean(akb_auc)),
        "mean_patch_average_gain_auc": float(np.mean(gain)),
        "median_patch_average_gain_auc": float(np.median(gain)),
        "gain_bootstrap_95_percent_ci": [
            float(np.quantile(boot, 0.025)),
            float(np.quantile(boot, 0.975)),
        ],
        "paired_gain_exact_one_sided_p": exact_one_sided(gain),
        "AKB_vs_chance_exact_one_sided_p": exact_one_sided(akb_auc - 0.5),
        "participants_positive_gain": int(np.sum(gain > 0)),
        "minimum_patch_specific_group_mean_AKB_auc": float(
            min(row["mean_AKB_auc"] for row in patch_metrics)
        ),
        "minimum_patch_specific_group_mean_gain_auc": float(
            min(row["mean_gain_auc"] for row in patch_metrics)
        ),
    }
    checks = {
        "mean_patch_average_AKB_auc_at_least_0.59": (
            metrics["mean_patch_average_AKB_auc"] >= 0.59
        ),
        "mean_patch_average_gain_at_least_0.01": (
            metrics["mean_patch_average_gain_auc"] >= 0.01
        ),
        "paired_gain_p_at_most_0.10": (
            metrics["paired_gain_exact_one_sided_p"] <= 0.10
        ),
        "positive_gain_at_least_7": metrics["participants_positive_gain"] >= 7,
        "minimum_patch_AKB_at_least_0.55": (
            metrics["minimum_patch_specific_group_mean_AKB_auc"] >= 0.55
        ),
    }
    advance = all(checks.values())
    payload = {
        "schema": "KITE-SAT-AKB-patch-resilience-development-gate-v1",
        "status": (
            "advance_patch_endpoint_to_confirmation"
            if advance
            else "patch_resilience_development_failed"
        ),
        "protocol_sha256": PROTOCOL_HASH,
        "subject_order": list(SUBJECTS),
        "subject_artifact_sha256": hashes,
        "subject_results": [
            {
                "subject": record["subject"],
                "global_patch_average_auc": record["endpoints"]["global"][
                    "patch_average_auc"
                ],
                "AKB_patch_average_auc": record["endpoints"]["AKB"][
                    "patch_average_auc"
                ],
                "gain_auc": record["patch_average_gain_auc"],
            }
            for record in records
        ],
        "patch_specific_group_metrics": patch_metrics,
        "metrics": metrics,
        "gate_checks": checks,
        "advance": advance,
        "firewall": {
            "confirmation_archives_downloaded": False,
            "confirmation_subject_read": False,
            "paper_mutated": False,
        },
    }
    OUTPUT.parent.mkdir(parents=True, exist_ok=True)
    OUTPUT.write_text(json.dumps(payload, indent=2) + "\n", encoding="utf-8")
    observed = sha256(OUTPUT)
    OUTPUT.with_name(OUTPUT.name + ".sha256").write_text(
        f"{observed}  {OUTPUT.name}\n", encoding="ascii"
    )
    print(
        json.dumps(
            {
                "status": payload["status"],
                "metrics": metrics,
                "checks": checks,
                "sha256": observed,
            }
        )
    )


if __name__ == "__main__":
    main()
```
