## Supplementary material for "Falsifiable substitution tests reveal task-structured neural evidence for auditory attention": Article_Analysis_Source_Code: run_kite_sat_akb_patch_subject_v1.py

```
"""Locked contiguous-patch resilience result for one AKB development subject."""

from __future__ import annotations

import argparse
import hashlib
import json
from pathlib import Path

import numpy as np
from scipy.signal import resample_poly

from kite_sat_adaptive_broad_v1 import adaptive_score
from kite_sat_backward_v1 import block_design
from kite_sat_geometry_v1 import partition_ids
from kite_sat_patch_erasure_v1 import ANCHORS, fixed_patch_masks, erased_cell_delta_contributions
from kite_st_hd_core_v1 import load_public_geometry
from kite_st_hd_statistics_v1 import binary_auc


ROOT = Path(__file__).resolve().parents[1]
PROTOCOL = ROOT / "docs" / "kite_sat_akb_patch_resilience_development_protocol_v1.json"
PROTOCOL_HASH = "ff32e236996b761cc059e03f7f775aac84cd3322cab2d220cf44efff219d1cb4"
SUBJECTS = (
    "S4", "S24", "S11", "S23", "S14", "S29",
    "S19", "S17", "S12", "S25", "S27", "S5",
)
CONDITIONS = ("1L", "1R", "2L", "2R")
PREPROCESSED_ROOT = (
    ROOT / "data" / "derived" / "kite_st_hd_zenodo4518754" / "development_preprocessed_v2"
)
MANIFEST_ROOT = ROOT / "results" / "kite_st_hd" / "development_v2" / "subjects"
MODEL_ROOT = (
    ROOT
    / "data"
    / "derived"
    / "kite_st_hd_zenodo4518754"
    / "kite_sat_formal_development_models_v1"
)
INTACT_ROOT = ROOT / "results" / "kite_sat" / "akb_stability_development_v1" / "subjects"
INTACT_GATE = (
    ROOT / "results" / "kite_sat" / "akb_stability_development_v1" / "development_gate_v1.json"
)
INTACT_GATE_HASH = "6a29be774fb3d68016b0d4a6d59d6f627c7a8c297a67bf68696af8aa7a8b2e5f"
AUDIO_CACHE = (
    ROOT
    / "data"
    / "derived"
    / "kite_st_hd_zenodo4518754"
    / "kite_sat_audio_features_v1.npz"
)
COORDINATES = (
    ROOT
    / "downloads"
    / "candidate_audit"
    / "zenodo4518754_support"
    / "misc"
    / "misc"
    / "eeg255ch_locs.csv"
)
OUTPUT_ROOT = (
    ROOT / "results" / "kite_sat" / "akb_patch_resilience_development_v1" / "subjects"
)
PHYSICAL = {
    "1L": ("part1_track1", "part1_track2"),
    "1R": ("part1_track1", "part1_track2"),
    "2L": ("part2_track2", "part2_track1"),
    "2R": ("part2_track2", "part2_track1"),
}


def sha256(path: Path) -> str:
    digest = hashlib.sha256()
    with path.open("rb") as stream:
        for chunk in iter(lambda: stream.read(8 * 1024 * 1024), b""):
            digest.update(chunk)
    return digest.hexdigest()


def main() -> None:
    parser = argparse.ArgumentParser()
    parser.add_argument("subject", choices=SUBJECTS)
    args = parser.parse_args()
    subject = args.subject
    output = OUTPUT_ROOT / f"{subject}_patch_resilience_v1.json"
    if output.exists():
        raise RuntimeError(f"Refusing to overwrite patch result for {subject}.")
    if sha256(PROTOCOL) != PROTOCOL_HASH:
        raise RuntimeError("Patch-resilience protocol hash mismatch.")
    if sha256(INTACT_GATE) != INTACT_GATE_HASH:
        raise RuntimeError("Intact AKB development gate hash mismatch.")
    intact_path = INTACT_ROOT / f"{subject}_S16_softmax2_v1.json"
    intact = json.loads(intact_path.read_text(encoding="utf-8"))
    intact_gate = json.loads(INTACT_GATE.read_text(encoding="utf-8"))
    if (
        sha256(intact_path) != intact_gate["subject_artifact_sha256"][subject]
        or intact["status"] != "pass"
        or intact["firewall"]["confirmation_subject_read"]
    ):
        raise RuntimeError("Invalid frozen intact AKB artifact.")
    manifest_path = MANIFEST_ROOT / f"{subject}_production_manifest_v1.json"
    manifest = json.loads(manifest_path.read_text(encoding="utf-8"))
    records = {
        item["recording"].rsplit("_", 1)[-1]: item
        for item in manifest["preprocessing_records"]
    }
    model_path = MODEL_ROOT / f"{subject}_speech_decoder_models_v1.npz"
    with np.load(model_path, allow_pickle=False) as saved:
        saved_models = {name: saved[name] for name in saved.files}
    with np.load(AUDIO_CACHE, allow_pickle=False) as saved:
        full_audio = {name: saved[name] for name in saved.files}
    geometry = load_public_geometry(COORDINATES)
    s16 = partition_ids(geometry.xy)["S16"]
    patches, patch_members = fixed_patch_masks(geometry.labels, geometry.xy)
    designs: dict[str, list[np.ndarray]] = {}
    audio_blocks: dict[str, list[tuple[np.ndarray, np.ndarray]]] = {}
    for condition in CONDITIONS:
        with np.load(
            PREPROCESSED_ROOT
            / subject
            / f"{subject}_AAD_{condition}_preprocessed_v1.npz",
            allow_pickle=False,
        ) as saved:
            eeg = saved["eeg"]
        record = records[condition]
        offset = int(
            round(
                (
                    record["crop"]["padded_start"]
                    + 3000
                    - record["trigger_train"]["first_sample"]
                )
                / 10.0
            )
        )
        left_name, right_name = PHYSICAL[condition]
        designs[condition], audio_blocks[condition] = [], []
        for block_index in range(12):
            first = offset + block_index * 3000
            stop = first + 3000
            left100 = full_audio[left_name][first:stop]
            right100 = full_audio[right_name][first:stop]
            design, left_features = block_design(
                eeg[block_index * 3000 : (block_index + 1) * 3000], left100
            )
            right_features = resample_poly(right100, 1, 5, axis=0)[:595]
            designs[condition].append(design)
            audio_blocks[condition].append(
                (left_features[:, 0], right_features[:, 0])
            )
    condition_scores = {
        estimator: {
            condition: [[] for _ in ANCHORS]
            for condition in CONDITIONS
        }
        for estimator in ("global", "AKB")
    }
    for train_story, test_story in (("1", "2"), ("2", "1")):
        reliability = np.asarray(
            intact["fold_reliabilities"][f"train_story_{train_story}"],
            dtype=np.float64,
        )
        full_model = {
            "x_mean": saved_models[f"train_story_{train_story}_x_mean"],
            "x_scale": saved_models[f"train_story_{train_story}_x_scale"],
            "weights": saved_models[f"train_story_{train_story}_weights"],
            "y_mean": float(saved_models[f"train_story_{train_story}_y_mean"]),
        }
        for condition in (f"{test_story}L", f"{test_story}R"):
            for design, (left, right) in zip(
                designs[condition], audio_blocks[condition]
            ):
                delta, global_margin = erased_cell_delta_contributions(
                    design, full_model, left, right, s16, patches
                )
                for patch_index in range(len(ANCHORS)):
                    condition_scores["global"][condition][patch_index].append(
                        float(global_margin[patch_index])
                    )
                    condition_scores["AKB"][condition][patch_index].append(
                        adaptive_score(
                            delta[patch_index], reliability, "softmax2"
                        )
                    )
    endpoints: dict[str, dict] = {}
    for estimator in ("global", "AKB"):
        patch_rows = []
        for patch_index, anchor in enumerate(ANCHORS):
            story1 = binary_auc(
                condition_scores[estimator]["1L"][patch_index],
                condition_scores[estimator]["1R"][patch_index],
            )
            story2 = binary_auc(
                condition_scores[estimator]["2L"][patch_index],
                condition_scores[estimator]["2R"][patch_index],
            )
            patch_rows.append(
                {
                    "anchor": anchor,
                    "story1_auc": story1,
                    "story2_auc": story2,
                    "mean_auc": (story1 + story2) / 2.0,
                }
            )
        endpoints[estimator] = {
            "patches": patch_rows,
            "patch_average_auc": float(
                np.mean([row["mean_auc"] for row in patch_rows])
            ),
        }
    payload = {
        "schema": "KITE-SAT-AKB-patch-resilience-development-subject-v1",
        "status": "pass",
        "subject": subject,
        "protocol_sha256": PROTOCOL_HASH,
        "intact_AKB_artifact": {
            "path": intact_path.relative_to(ROOT).as_posix(),
            "sha256": sha256(intact_path),
        },
        "model_archive": {
            "path": model_path.relative_to(ROOT).as_posix(),
            "sha256": sha256(model_path),
        },
        "patches": [
            {
                "anchor": anchor,
                "removed_count": int(np.sum(patches[index])),
                "removed_labels": list(patch_members[index]),
            }
            for index, anchor in enumerate(ANCHORS)
        ],
        "endpoints": endpoints,
        "patch_average_gain_auc": (
            endpoints["AKB"]["patch_average_auc"]
            - endpoints["global"]["patch_average_auc"]
        ),
        "condition_scores": condition_scores,
        "firewall": {
            "patch_variant_computed": False,
            "confirmation_subject_read": False,
            "paper_mutated": False,
        },
    }
    OUTPUT_ROOT.mkdir(parents=True, exist_ok=True)
    output.write_text(json.dumps(payload, indent=2) + "\n", encoding="utf-8")
    observed = sha256(output)
    output.with_name(output.name + ".sha256").write_text(
        f"{observed}  {output.name}\n", encoding="ascii"
    )
    print(
        json.dumps(
            {
                "subject": subject,
                "global_patch_average_auc": endpoints["global"]["patch_average_auc"],
                "AKB_patch_average_auc": endpoints["AKB"]["patch_average_auc"],
                "gain": payload["patch_average_gain_auc"],
                "sha256": observed,
            }
        )
    )


if __name__ == "__main__":
    main()
```
