## Supplementary material for "Falsifiable substitution tests reveal task-structured neural evidence for auditory attention": Article_Analysis_Source_Code: run_kite_sat_akb_stability_subject_v1.py

```
"""Locked S16-softmax2 AKB result for one KITE-SAT development subject."""

from __future__ import annotations

import argparse
import hashlib
import json
from pathlib import Path

import numpy as np
from scipy.signal import resample_poly

from kite_sat_adaptive_broad_v1 import adaptive_score, reliability
from kite_sat_backward_v1 import block_design
from kite_sat_geometry_v1 import cell_delta_contributions, partition_ids
from kite_sat_locked_decoder_v1 import fit_locked_ridge
from kite_st_hd_core_v1 import load_public_geometry
from kite_st_hd_statistics_v1 import binary_auc


ROOT = Path(__file__).resolve().parents[1]
PROTOCOL = ROOT / "docs" / "kite_sat_akb_stability_development_protocol_v1.json"
PROTOCOL_HASH = "12be0a01ed654cab61f54dffc6df84e34ac2bce5ddb17653ee81aea695a2b883"
SUBJECTS = ("S4", "S24", "S11", "S23", "S14", "S29", "S19", "S17", "S12", "S25", "S27", "S5")
CONDITIONS = ("1L", "1R", "2L", "2R")
PREPROCESSED_ROOT = ROOT / "data" / "derived" / "kite_st_hd_zenodo4518754" / "development_preprocessed_v2"
MANIFEST_ROOT = ROOT / "results" / "kite_st_hd" / "development_v2" / "subjects"
MODEL_ROOT = ROOT / "data" / "derived" / "kite_st_hd_zenodo4518754" / "kite_sat_formal_development_models_v1"
BASELINE_ROOT = ROOT / "results" / "kite_sat" / "formal_development_v1" / "subjects"
AUDIO_CACHE = ROOT / "data" / "derived" / "kite_st_hd_zenodo4518754" / "kite_sat_audio_features_v1.npz"
COORDINATES = ROOT / "downloads" / "candidate_audit" / "zenodo4518754_support" / "misc" / "misc" / "eeg255ch_locs.csv"
OUTPUT_ROOT = ROOT / "results" / "kite_sat" / "akb_stability_development_v1" / "subjects"
PHYSICAL = {
    "1L": ("part1_track1", "part1_track2"),
    "1R": ("part1_track1", "part1_track2"),
    "2L": ("part2_track2", "part2_track1"),
    "2R": ("part2_track2", "part2_track1"),
}


def sha256(path: Path) -> str:
    digest = hashlib.sha256()
    with path.open("rb") as stream:
        for chunk in iter(lambda: stream.read(8 * 1024 * 1024), b""):
            digest.update(chunk)
    return digest.hexdigest()


def main() -> None:
    parser = argparse.ArgumentParser()
    parser.add_argument("subject", choices=SUBJECTS)
    args = parser.parse_args()
    subject = args.subject
    output = OUTPUT_ROOT / f"{subject}_S16_softmax2_v1.json"
    if output.exists():
        raise RuntimeError(f"Refusing to overwrite AKB result for {subject}.")
    if sha256(PROTOCOL) != PROTOCOL_HASH:
        raise RuntimeError("AKB stability protocol hash mismatch.")
    baseline_path = BASELINE_ROOT / f"{subject}_speech_decoder_v1.json"
    baseline = json.loads(baseline_path.read_text(encoding="utf-8"))
    manifest_path = MANIFEST_ROOT / f"{subject}_production_manifest_v1.json"
    manifest = json.loads(manifest_path.read_text(encoding="utf-8"))
    records = {
        item["recording"].rsplit("_", 1)[-1]: item
        for item in manifest["preprocessing_records"]
    }
    model_path = MODEL_ROOT / f"{subject}_speech_decoder_models_v1.npz"
    with np.load(model_path, allow_pickle=False) as saved:
        saved_models = {name: saved[name] for name in saved.files}
    with np.load(AUDIO_CACHE, allow_pickle=False) as saved:
        full_audio = {name: saved[name] for name in saved.files}
    s16 = partition_ids(load_public_geometry(COORDINATES).xy)["S16"]
    designs: dict[str, list[np.ndarray]] = {}
    audio_blocks: dict[str, list[tuple[np.ndarray, np.ndarray]]] = {}
    attended: dict[str, list[np.ndarray]] = {}
    for condition in CONDITIONS:
        with np.load(
            PREPROCESSED_ROOT / subject / f"{subject}_AAD_{condition}_preprocessed_v1.npz",
            allow_pickle=False,
        ) as saved:
            eeg = saved["eeg"]
        record = records[condition]
        offset = int(
            round(
                (
                    record["crop"]["padded_start"]
                    + 3000
                    - record["trigger_train"]["first_sample"]
                )
                / 10.0
            )
        )
        left_name, right_name = PHYSICAL[condition]
        designs[condition], audio_blocks[condition], attended[condition] = [], [], []
        for block_index in range(12):
            first = offset + block_index * 3000
            stop = first + 3000
            left100 = full_audio[left_name][first:stop]
            right100 = full_audio[right_name][first:stop]
            design, left_features = block_design(
                eeg[block_index * 3000 : (block_index + 1) * 3000], left100
            )
            right_features = resample_poly(right100, 1, 5, axis=0)[:595]
            left, right = left_features[:, 0], right_features[:, 0]
            designs[condition].append(design)
            audio_blocks[condition].append((left, right))
            attended[condition].append(left if condition.endswith("L") else right)
    condition_scores = {
        "global": {condition: [] for condition in CONDITIONS},
        "AKB": {condition: [] for condition in CONDITIONS},
    }
    fold_reliabilities = {}
    for train_story, test_story in (("1", "2"), ("2", "1")):
        train_conditions = (f"{train_story}L", f"{train_story}R")
        test_conditions = (f"{test_story}L", f"{test_story}R")
        held_aligned = []
        for fit_parity in (0, 1):
            held_parity = 1 - fit_parity
            calibration_model = fit_locked_ridge(
                [
                    designs[condition][index]
                    for condition in train_conditions
                    for index in range(12)
                    if index % 2 == fit_parity
                ],
                [
                    attended[condition][index]
                    for condition in train_conditions
                    for index in range(12)
                    if index % 2 == fit_parity
                ],
            )
            for condition in train_conditions:
                alignment = 1.0 if condition.endswith("L") else -1.0
                for index in range(12):
                    if index % 2 != held_parity:
                        continue
                    left, right = audio_blocks[condition][index]
                    delta, _ = cell_delta_contributions(
                        designs[condition][index],
                        calibration_model,
                        left,
                        right,
                        s16,
                    )
                    held_aligned.append(alignment * delta)
        fold_reliability = reliability(np.asarray(held_aligned))
        fold_reliabilities[f"train_story_{train_story}"] = [
            float(value) for value in fold_reliability
        ]
        full_model = {
            "x_mean": saved_models[f"train_story_{train_story}_x_mean"],
            "x_scale": saved_models[f"train_story_{train_story}_x_scale"],
            "weights": saved_models[f"train_story_{train_story}_weights"],
            "y_mean": float(saved_models[f"train_story_{train_story}_y_mean"]),
        }
        for condition in test_conditions:
            for design, (left, right) in zip(
                designs[condition], audio_blocks[condition]
            ):
                delta, global_margin = cell_delta_contributions(
                    design, full_model, left, right, s16
                )
                condition_scores["global"][condition].append(global_margin)
                condition_scores["AKB"][condition].append(
                    adaptive_score(delta, fold_reliability, "softmax2")
                )
    endpoints = {}
    for name in ("global", "AKB"):
        story1 = binary_auc(
            condition_scores[name]["1L"], condition_scores[name]["1R"]
        )
        story2 = binary_auc(
            condition_scores[name]["2L"], condition_scores[name]["2R"]
        )
        endpoints[name] = {
            "story1_auc": story1,
            "story2_auc": story2,
            "mean_auc": (story1 + story2) / 2.0,
        }
    reproduction_error = abs(endpoints["global"]["mean_auc"] - baseline["mean_auc"])
    if reproduction_error > 1e-12:
        raise RuntimeError(f"AKB baseline reproduction failed: {reproduction_error}")
    payload = {
        "schema": "KITE-SAT-AKB-stability-development-subject-v1",
        "status": "pass",
        "subject": subject,
        "protocol_sha256": PROTOCOL_HASH,
        "baseline_artifact": {
            "path": baseline_path.relative_to(ROOT).as_posix(),
            "sha256": sha256(baseline_path),
        },
        "model_archive": {
            "path": model_path.relative_to(ROOT).as_posix(),
            "sha256": sha256(model_path),
        },
        "endpoints": endpoints,
        "gain_auc": endpoints["AKB"]["mean_auc"] - endpoints["global"]["mean_auc"],
        "baseline_reproduction_error": reproduction_error,
        "condition_scores": condition_scores,
        "fold_reliabilities": fold_reliabilities,
        "firewall": {
            "other_AKB_candidate_computed": False,
            "confirmation_subject_read": False,
            "paper_mutated": False,
        },
    }
    OUTPUT_ROOT.mkdir(parents=True, exist_ok=True)
    output.write_text(json.dumps(payload, indent=2) + "\n", encoding="utf-8")
    observed = sha256(output)
    output.with_name(output.name + ".sha256").write_text(
        f"{observed}  {output.name}\n", encoding="ascii"
    )
    print(
        json.dumps(
            {
                "subject": subject,
                "global_auc": endpoints["global"]["mean_auc"],
                "AKB_auc": endpoints["AKB"]["mean_auc"],
                "gain": payload["gain_auc"],
                "sha256": observed,
            }
        )
    )


if __name__ == "__main__":
    main()
```
