## Supplementary material for "Falsifiable substitution tests reveal task-structured neural evidence for auditory attention": Article_Analysis_Source_Code: run_kite_sat_development_subject_v1.py

```
"""Produce one frozen KITE-SAT speech-anchored development result."""

from __future__ import annotations

import argparse
import hashlib
import json
from pathlib import Path

import numpy as np
from scipy.signal import resample_poly

from kite_sat_backward_v1 import block_design, pearson
from kite_sat_locked_decoder_v1 import fit_locked_ridge, predict_locked
from kite_st_hd_statistics_v1 import binary_auc


ROOT = Path(__file__).resolve().parents[1]
PROTOCOL = ROOT / "docs" / "kite_sat_formal_development_protocol_v1.json"
PROTOCOL_HASH = "9d16d5499e281a582531b9d87a04e52da7228b3982fc220acb25add5d382c07e"
PREPROCESSED_ROOT = ROOT / "data" / "derived" / "kite_st_hd_zenodo4518754" / "development_preprocessed_v2"
MANIFEST_ROOT = ROOT / "results" / "kite_st_hd" / "development_v2" / "subjects"
AUDIO_CACHE = ROOT / "data" / "derived" / "kite_st_hd_zenodo4518754" / "kite_sat_audio_features_v1.npz"
OUTPUT_ROOT = ROOT / "results" / "kite_sat" / "formal_development_v1" / "subjects"
MODEL_ROOT = ROOT / "data" / "derived" / "kite_st_hd_zenodo4518754" / "kite_sat_formal_development_models_v1"
SUBJECTS = ("S4", "S24", "S11", "S23", "S14", "S29", "S19", "S17", "S12", "S25", "S27", "S5")
CONDITIONS = ("1L", "1R", "2L", "2R")
PHYSICAL = {
    "1L": ("part1_track1", "part1_track2"),
    "1R": ("part1_track1", "part1_track2"),
    "2L": ("part2_track2", "part2_track1"),
    "2R": ("part2_track2", "part2_track1"),
}


def sha256(path: Path) -> str:
    digest = hashlib.sha256()
    with path.open("rb") as stream:
        for chunk in iter(lambda: stream.read(8 * 1024 * 1024), b""):
            digest.update(chunk)
    return digest.hexdigest()


def main() -> None:
    parser = argparse.ArgumentParser()
    parser.add_argument("subject", choices=SUBJECTS)
    args = parser.parse_args()
    subject = args.subject
    output = OUTPUT_ROOT / f"{subject}_speech_decoder_v1.json"
    model_output = MODEL_ROOT / f"{subject}_speech_decoder_models_v1.npz"
    if output.exists() or model_output.exists():
        raise RuntimeError(f"Refusing to overwrite KITE-SAT output for {subject}.")
    if sha256(PROTOCOL) != PROTOCOL_HASH:
        raise RuntimeError("KITE-SAT formal protocol hash mismatch.")
    manifest_path = MANIFEST_ROOT / f"{subject}_production_manifest_v1.json"
    manifest = json.loads(manifest_path.read_text(encoding="utf-8"))
    if manifest["status"] != "pass" or manifest["subject"] != subject:
        raise RuntimeError("Invalid upstream development manifest.")
    records = {
        item["recording"].rsplit("_", 1)[-1]: item
        for item in manifest["preprocessing_records"]
    }
    with np.load(AUDIO_CACHE, allow_pickle=False) as saved:
        full_audio = {name: saved[name] for name in saved.files}
    designs: dict[str, list[np.ndarray]] = {}
    physical_audio: dict[str, list[tuple[np.ndarray, np.ndarray]]] = {}
    attended: dict[str, list[np.ndarray]] = {}
    offsets: dict[str, float] = {}
    input_hashes: dict[str, str] = {}
    for condition in CONDITIONS:
        path = PREPROCESSED_ROOT / subject / f"{subject}_AAD_{condition}_preprocessed_v1.npz"
        input_hashes[condition] = sha256(path)
        with np.load(path, allow_pickle=False) as saved:
            eeg = saved["eeg"]
        record = records[condition]
        offset_seconds = (
            record["crop"]["padded_start"] + 3000 - record["trigger_train"]["first_sample"]
        ) / 1000.0
        offsets[condition] = offset_seconds
        offset = int(round(offset_seconds * 100))
        left_name, right_name = PHYSICAL[condition]
        designs[condition] = []
        physical_audio[condition] = []
        attended[condition] = []
        for block_index in range(12):
            first = offset + block_index * 3000
            stop = first + 3000
            left = full_audio[left_name][first:stop]
            right = full_audio[right_name][first:stop]
            design, left_features = block_design(
                eeg[block_index * 3000 : (block_index + 1) * 3000], left
            )
            right_features = resample_poly(right, 1, 5, axis=0)[:595]
            designs[condition].append(design)
            physical_audio[condition].append((left_features[:, 0], right_features[:, 0]))
            attended[condition].append(
                left_features[:, 0] if condition.endswith("L") else right_features[:, 0]
            )
    story_results: dict[str, object] = {}
    model_arrays: dict[str, np.ndarray] = {}
    for train_story, test_story in (("1", "2"), ("2", "1")):
        training_conditions = (f"{train_story}L", f"{train_story}R")
        testing_conditions = (f"{test_story}L", f"{test_story}R")
        training_designs = [
            design for condition in training_conditions for design in designs[condition]
        ]
        training_targets = [
            target for condition in training_conditions for target in attended[condition]
        ]
        model = fit_locked_ridge(training_designs, training_targets)
        for name in ("x_mean", "x_scale", "weights"):
            model_arrays[f"train_story_{train_story}_{name}"] = np.asarray(model[name])
        model_arrays[f"train_story_{train_story}_y_mean"] = np.asarray(model["y_mean"])
        model_arrays[f"train_story_{train_story}_alpha"] = np.asarray(model["alpha"])
        condition_scores: dict[str, list[float]] = {}
        condition_correlations: dict[str, list[dict[str, float]]] = {}
        for condition in testing_conditions:
            condition_scores[condition] = []
            condition_correlations[condition] = []
            for design, (left, right) in zip(designs[condition], physical_audio[condition]):
                reconstruction = predict_locked(design, model)
                corr_left = pearson(reconstruction, left)
                corr_right = pearson(reconstruction, right)
                condition_scores[condition].append(corr_left - corr_right)
                condition_correlations[condition].append(
                    {"left": corr_left, "right": corr_right}
                )
        auc = binary_auc(
            condition_scores[f"{test_story}L"], condition_scores[f"{test_story}R"]
        )
        story_results[test_story] = {
            "trained_story": train_story,
            "tested_story": test_story,
            "auc": auc,
            "left_minus_right_scores": condition_scores,
            "correlations": condition_correlations,
            "alpha": float(model["alpha"]),
            "gram_energy": float(model["gram_energy"]),
        }
    story_aucs = [story_results[story]["auc"] for story in ("1", "2")]
    MODEL_ROOT.mkdir(parents=True, exist_ok=True)
    np.savez_compressed(model_output, **model_arrays)
    model_hash = sha256(model_output)
    model_output.with_name(model_output.name + ".sha256").write_text(
        f"{model_hash}  {model_output.name}\n", encoding="ascii"
    )
    payload = {
        "schema": "KITE-SAT-formal-development-subject-v1",
        "status": "pass",
        "subject": subject,
        "protocol_sha256": PROTOCOL_HASH,
        "upstream_manifest": {
            "path": manifest_path.relative_to(ROOT).as_posix(),
            "sha256": sha256(manifest_path),
        },
        "audio_cache_sha256": sha256(AUDIO_CACHE),
        "preprocessed_sha256": input_hashes,
        "offset_seconds": offsets,
        "story_results": story_results,
        "mean_auc": float(np.mean(story_aucs)),
        "minimum_story_auc": float(np.min(story_aucs)),
        "model_archive": {
            "path": model_output.relative_to(ROOT).as_posix(),
            "bytes": model_output.stat().st_size,
            "sha256": model_hash,
        },
        "firewall": {
            "confirmation_subject_read": False,
            "paper_mutated": False,
        },
    }
    OUTPUT_ROOT.mkdir(parents=True, exist_ok=True)
    output.write_text(json.dumps(payload, indent=2) + "\n", encoding="utf-8")
    observed = sha256(output)
    output.with_name(output.name + ".sha256").write_text(
        f"{observed}  {output.name}\n", encoding="ascii"
    )
    print(
        json.dumps(
            {
                "subject": subject,
                "mean_auc": payload["mean_auc"],
                "story1_auc": story_results["1"]["auc"],
                "story2_auc": story_results["2"]["auc"],
                "sha256": observed,
            }
        )
    )


if __name__ == "__main__":
    main()
```
