## Supplementary material for "Falsifiable substitution tests reveal task-structured neural evidence for auditory attention": Article_Analysis_Source_Code: run_kite_sat_ear_saad_external_subject_v1.py

```
#!/usr/bin/env python3
"""Run the frozen KITE-SAT AKB on one Ear-SAAD scalp-EEG participant."""

from __future__ import annotations

import argparse
import hashlib
import json
from pathlib import Path

import numpy as np
from scipy.io import loadmat
from scipy.linalg import cho_factor, cho_solve
from scipy.stats import rankdata

from kite_sat_adaptive_broad_v1 import family_weights, reliability
from kite_sat_direction_delay_controls_v1 import TRANSFORMS, transformed_matrix


ROOT = Path(__file__).resolve().parents[1]
PROTOCOL = ROOT / "docs" / "kite_sat_ear_saad_external_protocol_v1.json"
IMPLEMENTATION_LOCK = (
    ROOT / "docs" / "kite_sat_ear_saad_external_implementation_lock_v2.json"
)
GEOMETRY = ROOT / "docs" / "kite_sat_ear_saad_scalp_geometry_v1.json"
DATA_ROOT = ROOT / "data" / "raw" / "ear_saad" / "preprocessedData"
OUTPUT_ROOT = ROOT / "results" / "kite_sat" / "ear_saad_external_v1" / "subjects"
MODEL_ROOT = (
    ROOT / "data" / "derived" / "ear_saad" / "kite_sat_external_models_v1"
)
PROTOCOL_SHA256 = "6ed5c37fe2869072646e632dff6a807c4a91c162c1ab3048fc52bc235fb9d906"
SUBJECTS = tuple(range(1, 16))
LAGS = np.arange(6, dtype=np.int64)
N_CHANNELS = 29
N_FEATURES = N_CHANNELS * len(LAGS)
SAMPLES_PER_BLOCK = 600
USABLE = SAMPLES_PER_BLOCK - int(LAGS[-1])
BLOCKS_PER_TRIAL = 20
TEMPORAL_SHIFT = 1200


def sha256(path: Path) -> str:
    digest = hashlib.sha256()
    with path.open("rb") as handle:
        for block in iter(lambda: handle.read(1024 * 1024), b""):
            digest.update(block)
    return digest.hexdigest()


def validate_lock() -> tuple[dict, str]:
    if sha256(PROTOCOL) != PROTOCOL_SHA256:
        raise RuntimeError("Ear-SAAD protocol hash mismatch.")
    if not IMPLEMENTATION_LOCK.exists():
        raise RuntimeError("Ear-SAAD implementation lock is unavailable.")
    lock = json.loads(IMPLEMENTATION_LOCK.read_text(encoding="utf-8"))
    lock_hash = sha256(IMPLEMENTATION_LOCK)
    if (
        lock["status"] != "frozen_before_any_KITE-SAT_Ear-SAAD_outcome"
        or lock["protocol_sha256"] != PROTOCOL_SHA256
        or lock["source_sha256"][Path(__file__).name] != sha256(Path(__file__))
    ):
        raise RuntimeError("Ear-SAAD implementation lock mismatch.")
    return lock, lock_hash


def binary_auc(positive: np.ndarray, negative: np.ndarray) -> float:
    left = np.asarray(positive, dtype=np.float64)
    right = np.asarray(negative, dtype=np.float64)
    joined = np.r_[left, right]
    ranks = rankdata(joined, method="average")
    u = np.sum(ranks[: len(left)]) - len(left) * (len(left) + 1) / 2.0
    return float(u / (len(left) * len(right)))


def balanced_accuracy(scores: np.ndarray, labels: np.ndarray) -> float:
    scores = np.asarray(scores, dtype=np.float64)
    labels = np.asarray(labels, dtype=np.int64)
    positive = labels == 1
    negative = labels == 0
    if not np.any(positive) or not np.any(negative):
        return float("nan")
    tpr = np.mean((scores[positive] > 0) + 0.5 * (scores[positive] == 0))
    tnr = np.mean((scores[negative] < 0) + 0.5 * (scores[negative] == 0))
    return float((tpr + tnr) / 2.0)


def block_design(eeg: np.ndarray) -> np.ndarray:
    array = np.asarray(eeg, dtype=np.float64)
    if array.shape != (SAMPLES_PER_BLOCK, N_CHANNELS):
        raise ValueError(f"Unexpected Ear-SAAD scalp block: {array.shape}")
    design = np.column_stack(
        [array[int(lag) : int(lag) + USABLE] for lag in LAGS]
    )
    if design.shape != (USABLE, N_FEATURES) or not np.isfinite(design).all():
        raise RuntimeError("Invalid Ear-SAAD design.")
    return design


def sufficient_statistics(design: np.ndarray, target: np.ndarray) -> dict[str, np.ndarray | float | int]:
    x = np.asarray(design, dtype=np.float64)
    y = np.asarray(target, dtype=np.float64)
    if x.shape != (USABLE, N_FEATURES) or y.shape != (USABLE,):
        raise ValueError("Unexpected block sufficient-statistic shape.")
    return {
        "n": len(x),
        "sum_x": np.sum(x, axis=0),
        "sum_x2": np.sum(x * x, axis=0),
        "sum_y": float(np.sum(y)),
        "xtx": x.T @ x,
        "xty": x.T @ y,
    }


def fit_from_blocks(records: list[dict], indices: list[int]) -> dict[str, np.ndarray | float]:
    if not indices:
        raise ValueError("No training blocks.")
    n = int(sum(int(records[index]["stats"]["n"]) for index in indices))
    sum_x = np.sum([records[index]["stats"]["sum_x"] for index in indices], axis=0)
    sum_x2 = np.sum([records[index]["stats"]["sum_x2"] for index in indices], axis=0)
    sum_y = float(sum(float(records[index]["stats"]["sum_y"]) for index in indices))
    xtx = np.sum([records[index]["stats"]["xtx"] for index in indices], axis=0)
    xty = np.sum([records[index]["stats"]["xty"] for index in indices], axis=0)
    x_mean = sum_x / n
    variance = np.maximum(sum_x2 / n - x_mean * x_mean, 0.0)
    x_scale = np.sqrt(variance)
    x_scale = np.where(x_scale > 1e-12, x_scale, 1.0)
    y_mean = sum_y / n
    centered_gram = xtx - n * np.outer(x_mean, x_mean)
    gram = centered_gram / np.outer(x_scale, x_scale)
    cross = (xty - x_mean * sum_y) / x_scale
    gram = 0.5 * (gram + gram.T)
    gram_energy = float(np.mean(np.diag(gram)))
    alpha = gram_energy
    regularized = gram.copy()
    regularized.flat[:: len(regularized) + 1] += alpha
    factor = cho_factor(regularized, lower=True, check_finite=False)
    weights = cho_solve(factor, cross, check_finite=False)
    if not np.isfinite(weights).all():
        raise RuntimeError("Nonfinite Ear-SAAD ridge weights.")
    return {
        "x_mean": x_mean,
        "x_scale": x_scale,
        "y_mean": float(y_mean),
        "weights": weights,
        "gram_energy": gram_energy,
        "alpha": alpha,
    }


def cell_contributions(
    design: np.ndarray,
    model: dict[str, np.ndarray | float],
    speaker1: np.ndarray,
    speaker2: np.ndarray,
    ids: np.ndarray,
) -> tuple[np.ndarray, float]:
    x = (
        np.asarray(design, dtype=np.float64) - np.asarray(model["x_mean"])
    ) / np.asarray(model["x_scale"])
    weighted = x * np.asarray(model["weights"])[None, :]
    reconstruction = np.sum(weighted, axis=1)
    centered_reconstruction = reconstruction - np.mean(reconstruction)
    first = np.asarray(speaker1, dtype=np.float64) - np.mean(speaker1)
    second = np.asarray(speaker2, dtype=np.float64) - np.mean(speaker2)
    denominator_first = np.sqrt(
        np.sum(centered_reconstruction**2) * np.sum(first**2)
    )
    denominator_second = np.sqrt(
        np.sum(centered_reconstruction**2) * np.sum(second**2)
    )
    delta = np.empty(16, dtype=np.float64)
    for cell in range(16):
        component = np.sum(weighted[:, ids == cell], axis=1)
        component -= np.mean(component)
        delta[cell] = (
            np.sum(component * first) / max(float(denominator_first), 1e-12)
            - np.sum(component * second) / max(float(denominator_second), 1e-12)
        )
    first_correlation = np.sum(centered_reconstruction * first) / max(
        float(denominator_first), 1e-12
    )
    second_correlation = np.sum(centered_reconstruction * second) / max(
        float(denominator_second), 1e-12
    )
    margin = float(first_correlation - second_correlation)
    if abs(float(np.sum(delta)) - margin) > 1e-10:
        raise RuntimeError("Ear-SAAD cell contributions are not additive.")
    return delta, margin


def broadness(reliability_values: np.ndarray) -> dict[str, float]:
    weights = family_weights(reliability_values, "softmax2")
    probability = weights / np.sum(weights)
    entropy = float(-np.sum(probability * np.log(probability + 1e-300)))
    angular = probability[:8] + probability[8:]
    angular_entropy = float(-np.sum(angular * np.log(angular + 1e-300)))
    return {
        "effective_cell_number": float(np.exp(entropy)),
        "normalized_cell_entropy": float(entropy / np.log(16)),
        "maximum_cell_mass": float(np.max(probability)),
        "effective_angular_sector_number": float(np.exp(angular_entropy)),
    }


def main() -> None:
    parser = argparse.ArgumentParser()
    parser.add_argument("subject", type=int, choices=SUBJECTS)
    args = parser.parse_args()
    subject = args.subject
    output = OUTPUT_ROOT / f"S{subject}_KITE_SAT_Ear_SAAD_v1.json"
    model_output = MODEL_ROOT / f"S{subject}_KITE_SAT_Ear_SAAD_models_v1.npz"
    if output.exists() or model_output.exists():
        raise RuntimeError(f"Refusing to overwrite Ear-SAAD S{subject}.")
    lock, lock_hash = validate_lock()
    protocol = json.loads(PROTOCOL.read_text(encoding="utf-8"))
    subject_protocol = next(
        row for row in protocol["dataset"]["participants"] if row["subject"] == subject
    )
    input_path = ROOT / subject_protocol["path"]
    if sha256(input_path) != subject_protocol["sha256"]:
        raise RuntimeError("Ear-SAAD participant input hash mismatch.")
    geometry = json.loads(GEOMETRY.read_text(encoding="utf-8"))
    if sha256(GEOMETRY) != protocol["frozen_geometry"]["sha256"]:
        raise RuntimeError("Ear-SAAD geometry hash mismatch.")
    sectors = np.asarray(geometry["angular_sector_id_0_to_7"], dtype=np.int64)
    lag_index = np.repeat(np.arange(6), N_CHANNELS)
    cell_ids = np.tile(sectors, 6) + 8 * (lag_index >= 3)
    if (
        cell_ids.shape != (N_FEATURES,)
        or np.any(np.bincount(cell_ids, minlength=16) == 0)
    ):
        raise RuntimeError("Degenerate Ear-SAAD direction-delay cells.")

    data = loadmat(input_path, simplify_cells=True)
    channels = [str(value) for value in np.asarray(data["channels"]).ravel()]
    if channels[:N_CHANNELS] != geometry["labels_in_data_order"]:
        raise RuntimeError("Ear-SAAD scalp channel order differs from geometry.")
    eeg_trials = [
        np.nan_to_num(
            np.asarray(value, dtype=np.float64),
            nan=0.0,
            posinf=0.0,
            neginf=0.0,
        )
        for value in np.asarray(data["eegTrials"], dtype=object).ravel()
    ]
    envelope_trials = [
        np.asarray(value, dtype=np.float64)
        for value in np.asarray(data["envelopes"], dtype=object).ravel()
    ]
    attended_speaker = np.asarray(data["attSpeaker"]).ravel().astype(np.int64)
    attended_ear = np.asarray(data["attendedEar"]).ravel().astype(np.int64)
    video = np.asarray(data["videoCondition"]).ravel().astype(np.int64)

    records: list[dict] = []
    for trial in range(6):
        shifted = np.roll(envelope_trials[trial], -TEMPORAL_SHIFT, axis=0)
        for block in range(BLOCKS_PER_TRIAL):
            start = block * SAMPLES_PER_BLOCK
            stop = start + SAMPLES_PER_BLOCK
            design = block_design(eeg_trials[trial][start:stop, :N_CHANNELS])
            envelopes = envelope_trials[trial][start:stop, :][:USABLE]
            shifted_envelopes = shifted[start:stop, :][:USABLE]
            target = envelopes[:, int(attended_speaker[trial]) - 1]
            records.append(
                {
                    "trial": trial,
                    "block": block,
                    "design": design,
                    "speaker1": envelopes[:, 0],
                    "speaker2": envelopes[:, 1],
                    "shifted_speaker1": shifted_envelopes[:, 0],
                    "shifted_speaker2": shifted_envelopes[:, 1],
                    "label": 1 if attended_speaker[trial] == 1 else 0,
                    "attended_ear": int(attended_ear[trial]),
                    "video": int(video[trial]),
                    "stats": sufficient_statistics(design, target),
                }
            )

    held_rows = []
    model_arrays: dict[str, np.ndarray] = {}
    fold_rows = []
    for held_trial in range(6):
        training = [
            index for index, row in enumerate(records)
            if row["trial"] != held_trial
        ]
        held = [
            index for index, row in enumerate(records)
            if row["trial"] == held_trial
        ]
        held_aligned = []
        for fit_parity in (0, 1):
            fit_indices = [
                index for index in training
                if records[index]["block"] % 2 == fit_parity
            ]
            calibration_indices = [
                index for index in training
                if records[index]["block"] % 2 != fit_parity
            ]
            calibration_model = fit_from_blocks(records, fit_indices)
            for index in calibration_indices:
                row = records[index]
                delta, _ = cell_contributions(
                    row["design"],
                    calibration_model,
                    row["speaker1"],
                    row["speaker2"],
                    cell_ids,
                )
                alignment = 1.0 if row["label"] == 1 else -1.0
                held_aligned.append(alignment * delta)
        if len(held_aligned) != 100:
            raise RuntimeError("Ear-SAAD calibration block count mismatch.")
        fold_reliability = reliability(np.asarray(held_aligned))
        identity_weights = family_weights(fold_reliability, "softmax2")
        weight_matrix = transformed_matrix(identity_weights)
        full_model = fit_from_blocks(records, training)
        prefix = f"fold_{held_trial + 1}"
        for name in ("x_mean", "x_scale", "weights"):
            model_arrays[f"{prefix}_{name}"] = np.asarray(full_model[name])
        for name in ("y_mean", "alpha", "gram_energy"):
            model_arrays[f"{prefix}_{name}"] = np.asarray(full_model[name])
        model_arrays[f"{prefix}_reliability"] = fold_reliability
        for index in held:
            row = records[index]
            delta, global_margin = cell_contributions(
                row["design"],
                full_model,
                row["speaker1"],
                row["speaker2"],
                cell_ids,
            )
            transform_scores = weight_matrix @ delta
            null_delta, null_global = cell_contributions(
                row["design"],
                full_model,
                row["shifted_speaker1"],
                row["shifted_speaker2"],
                cell_ids,
            )
            null_identity = float(weight_matrix[0] @ null_delta)
            held_rows.append(
                {
                    "trial": int(row["trial"] + 1),
                    "block": int(row["block"] + 1),
                    "label_speaker1": int(row["label"]),
                    "attended_ear_code": int(row["attended_ear"]),
                    "video_code": int(row["video"]),
                    "global_margin": global_margin,
                    "transform_scores": [float(value) for value in transform_scores],
                    "temporal_null_global_margin": null_global,
                    "temporal_null_AKB_margin": null_identity,
                }
            )
        fold_rows.append(
            {
                "held_trial": held_trial + 1,
                "training_trials": [value + 1 for value in range(6) if value != held_trial],
                "training_blocks": len(training),
                "held_blocks": len(held),
                "calibration_held_contributions": len(held_aligned),
                "reliability": [float(value) for value in fold_reliability],
                "broadness": broadness(fold_reliability),
            }
        )

    labels = np.asarray([row["label_speaker1"] for row in held_rows], dtype=np.int64)
    global_scores = np.asarray([row["global_margin"] for row in held_rows])
    transform_scores = np.asarray([row["transform_scores"] for row in held_rows])
    null_global = np.asarray(
        [row["temporal_null_global_margin"] for row in held_rows]
    )
    null_akb = np.asarray(
        [row["temporal_null_AKB_margin"] for row in held_rows]
    )
    positive = labels == 1
    negative = labels == 0
    global_auc = binary_auc(global_scores[positive], global_scores[negative])
    transform_aucs = np.asarray(
        [
            binary_auc(transform_scores[positive, index], transform_scores[negative, index])
            for index in range(16)
        ]
    )
    null_global_auc = binary_auc(null_global[positive], null_global[negative])
    null_akb_auc = binary_auc(null_akb[positive], null_akb[negative])
    identity_auc = float(transform_aucs[0])

    strata = {}
    for name, field, value in (
        ("video", "video_code", 1),
        ("fixation", "video_code", 0),
        ("attended_ear_code_1", "attended_ear_code", 1),
        ("attended_ear_code_2", "attended_ear_code", 2),
    ):
        selected = np.asarray([row[field] == value for row in held_rows])
        strata[name] = {
            "blocks": int(np.sum(selected)),
            "global_balanced_accuracy": balanced_accuracy(
                global_scores[selected], labels[selected]
            ),
            "AKB_balanced_accuracy": balanced_accuracy(
                transform_scores[selected, 0], labels[selected]
            ),
        }

    MODEL_ROOT.mkdir(parents=True, exist_ok=True)
    np.savez_compressed(model_output, **model_arrays)
    model_hash = sha256(model_output)
    Path(str(model_output) + ".sha256").write_text(
        f"{model_hash}  {model_output.name}\n", encoding="ascii"
    )
    payload = {
        "schema": "KITE-SAT-Ear-SAAD-external-subject-v1",
        "status": "pass",
        "subject": subject,
        "protocol_sha256": PROTOCOL_SHA256,
        "implementation_lock_sha256": lock_hash,
        "input": {
            "path": str(input_path.relative_to(ROOT)).replace("\\", "/"),
            "sha256": sha256(input_path),
        },
        "geometry_sha256": sha256(GEOMETRY),
        "cell_counts": np.bincount(cell_ids, minlength=16).tolist(),
        "outer_folds": fold_rows,
        "held_block_rows": held_rows,
        "endpoints": {
            "global_auc": global_auc,
            "AKB_identity_auc": identity_auc,
            "AKB_minus_global_auc": identity_auc - global_auc,
            "global_balanced_accuracy": balanced_accuracy(global_scores, labels),
            "AKB_balanced_accuracy": balanced_accuracy(
                transform_scores[:, 0], labels
            ),
            "transform_aucs": [float(value) for value in transform_aucs],
            "identity_minus_mean_15_controls_auc": float(
                transform_aucs[0] - np.mean(transform_aucs[1:])
            ),
            "temporal_null_global_auc": null_global_auc,
            "temporal_null_AKB_auc": null_akb_auc,
        },
        "strata": strata,
        "model_archive": {
            "path": str(model_output.relative_to(ROOT)).replace("\\", "/"),
            "bytes": model_output.stat().st_size,
            "sha256": model_hash,
        },
        "firewall": {
            "alternative_AKB_candidate_computed": False,
            "alternative_geometry_computed": False,
            "ear_modality_endpoint_computed": False,
            "paper_mutated": False,
        },
    }
    OUTPUT_ROOT.mkdir(parents=True, exist_ok=True)
    output.write_text(
        json.dumps(payload, indent=2, ensure_ascii=False) + "\n", encoding="utf-8"
    )
    observed = sha256(output)
    Path(str(output) + ".sha256").write_text(
        f"{observed}  {output.name}\n", encoding="ascii"
    )
    print(json.dumps({
        "subject": subject,
        "global_auc": global_auc,
        "AKB_auc": identity_auc,
        "gain_auc": identity_auc - global_auc,
        "identity_advantage_auc": payload["endpoints"][
            "identity_minus_mean_15_controls_auc"
        ],
        "sha256": observed,
    }))


if __name__ == "__main__":
    main()
```
