## Supplementary material for "Falsifiable substitution tests reveal task-structured neural evidence for auditory attention": Article_Analysis_Source_Code: run_kite_sat_formal_development_group_v1.py

```
"""Locked group gate for the KITE-SAT formal development benchmark."""

from __future__ import annotations

import hashlib
import itertools
import json
from pathlib import Path

import numpy as np

from kite_st_hd_core_v1 import stable_seed


ROOT = Path(__file__).resolve().parents[1]
PROTOCOL = ROOT / "docs" / "kite_sat_formal_development_protocol_v1.json"
PROTOCOL_HASH = "9d16d5499e281a582531b9d87a04e52da7228b3982fc220acb25add5d382c07e"
SUBJECT_ROOT = ROOT / "results" / "kite_sat" / "formal_development_v1" / "subjects"
OUTPUT = ROOT / "results" / "kite_sat" / "formal_development_v1" / "development_gate_v1.json"
SUBJECTS = ("S4", "S24", "S11", "S23", "S14", "S29", "S19", "S17", "S12", "S25", "S27", "S5")


def sha256(path: Path) -> str:
    digest = hashlib.sha256()
    with path.open("rb") as stream:
        for chunk in iter(lambda: stream.read(8 * 1024 * 1024), b""):
            digest.update(chunk)
    return digest.hexdigest()


def exact_sign_flip(values: np.ndarray) -> tuple[float, int]:
    observed = float(np.mean(values))
    exceed = 0
    total = 0
    for signs in itertools.product((-1.0, 1.0), repeat=len(values)):
        statistic = float(np.mean(values * np.asarray(signs)))
        exceed += statistic >= observed - 1e-15
        total += 1
    return exceed / total, total


def main() -> None:
    if OUTPUT.exists():
        raise RuntimeError("Refusing to overwrite KITE-SAT formal development gate.")
    if sha256(PROTOCOL) != PROTOCOL_HASH:
        raise RuntimeError("KITE-SAT formal protocol hash mismatch.")
    records = []
    subject_hashes = {}
    for subject in SUBJECTS:
        path = SUBJECT_ROOT / f"{subject}_speech_decoder_v1.json"
        payload = json.loads(path.read_text(encoding="utf-8"))
        if (
            payload["status"] != "pass"
            or payload["subject"] != subject
            or payload["protocol_sha256"] != PROTOCOL_HASH
            or payload["firewall"]["confirmation_subject_read"]
        ):
            raise RuntimeError(f"Invalid KITE-SAT subject artifact: {subject}")
        subject_hashes[subject] = sha256(path)
        records.append(payload)
    endpoints = np.asarray([item["mean_auc"] for item in records], dtype=np.float64)
    story1 = np.asarray(
        [item["story_results"]["1"]["auc"] for item in records], dtype=np.float64
    )
    story2 = np.asarray(
        [item["story_results"]["2"]["auc"] for item in records], dtype=np.float64
    )
    p_value, permutations = exact_sign_flip(endpoints - 0.5)
    rng = np.random.default_rng(
        stable_seed("KITE-SAT-formal-development-bootstrap-v1")
    )
    bootstrap = np.mean(
        endpoints[rng.integers(0, len(endpoints), size=(20000, len(endpoints)))],
        axis=1,
    )
    metrics = {
        "n": len(endpoints),
        "mean_auc": float(np.mean(endpoints)),
        "median_auc": float(np.median(endpoints)),
        "bootstrap_95_percent_ci": [
            float(np.quantile(bootstrap, 0.025)),
            float(np.quantile(bootstrap, 0.975)),
        ],
        "exact_one_sided_sign_flip_p": p_value,
        "exact_permutations": permutations,
        "story1_group_mean_auc": float(np.mean(story1)),
        "story2_group_mean_auc": float(np.mean(story2)),
        "participants_above_chance": int(np.sum(endpoints > 0.5)),
    }
    checks = {
        "p_at_most_0.025": p_value <= 0.025,
        "mean_auc_at_least_0.58": metrics["mean_auc"] >= 0.58,
        "story1_mean_at_least_0.55": metrics["story1_group_mean_auc"] >= 0.55,
        "story2_mean_at_least_0.55": metrics["story2_group_mean_auc"] >= 0.55,
        "participants_above_chance_at_least_8": metrics[
            "participants_above_chance"
        ]
        >= 8,
    }
    advance = all(checks.values())
    output = {
        "schema": "KITE-SAT-formal-development-gate-v1",
        "status": "advance_to_kakeya_method_development" if advance else "formal_development_failed",
        "protocol_sha256": PROTOCOL_HASH,
        "subject_order": list(SUBJECTS),
        "subject_artifact_sha256": subject_hashes,
        "subject_results": [
            {
                "subject": item["subject"],
                "mean_auc": item["mean_auc"],
                "story1_auc": item["story_results"]["1"]["auc"],
                "story2_auc": item["story_results"]["2"]["auc"],
                "minimum_story_auc": item["minimum_story_auc"],
            }
            for item in records
        ],
        "metrics": metrics,
        "gate_checks": checks,
        "advance": advance,
        "firewall": {
            "confirmation_archives_downloaded": False,
            "confirmation_subject_read": False,
            "paper_mutated": False,
        },
        "interpretation": (
            "Development benchmark only; passing authorizes method development, "
            "not a scientific claim or confirmation download."
        ),
    }
    OUTPUT.parent.mkdir(parents=True, exist_ok=True)
    OUTPUT.write_text(json.dumps(output, indent=2) + "\n", encoding="utf-8")
    observed = sha256(OUTPUT)
    OUTPUT.with_name(OUTPUT.name + ".sha256").write_text(
        f"{observed}  {OUTPUT.name}\n", encoding="ascii"
    )
    print(
        json.dumps(
            {
                "status": output["status"],
                "metrics": metrics,
                "checks": checks,
                "sha256": observed,
            }
        )
    )


if __name__ == "__main__":
    main()
```
