## Supplementary material for "Falsifiable substitution tests reveal task-structured neural evidence for auditory attention": Article_Analysis_Source_Code: run_kite_sat_geometry_development_subject_v1.py

```
"""Compute the locked P16-drop2 KITE-SAT result for one development subject."""

from __future__ import annotations

import argparse
import hashlib
import json
from pathlib import Path

import numpy as np
from scipy.signal import resample_poly

from kite_sat_backward_v1 import block_design
from kite_sat_geometry_locked_v1 import p16_drop2_score
from kite_sat_geometry_v1 import partition_ids
from kite_st_hd_core_v1 import load_public_geometry
from kite_st_hd_statistics_v1 import binary_auc


ROOT = Path(__file__).resolve().parents[1]
PROTOCOL = ROOT / "docs" / "kite_sat_geometry_development_protocol_v1.json"
PROTOCOL_HASH = "bb5c327b23fb898e2b6278a4595670cfc8e1e41ade75b24707544a8578e55200"
SUBJECTS = ("S4", "S24", "S11", "S23", "S14", "S29", "S19", "S17", "S12", "S25", "S27", "S5")
CONDITIONS = ("1L", "1R", "2L", "2R")
PREPROCESSED_ROOT = ROOT / "data" / "derived" / "kite_st_hd_zenodo4518754" / "development_preprocessed_v2"
MANIFEST_ROOT = ROOT / "results" / "kite_st_hd" / "development_v2" / "subjects"
MODEL_ROOT = ROOT / "data" / "derived" / "kite_st_hd_zenodo4518754" / "kite_sat_formal_development_models_v1"
BASELINE_ROOT = ROOT / "results" / "kite_sat" / "formal_development_v1" / "subjects"
AUDIO_CACHE = ROOT / "data" / "derived" / "kite_st_hd_zenodo4518754" / "kite_sat_audio_features_v1.npz"
COORDINATES = ROOT / "downloads" / "candidate_audit" / "zenodo4518754_support" / "misc" / "misc" / "eeg255ch_locs.csv"
OUTPUT_ROOT = ROOT / "results" / "kite_sat" / "geometry_development_v1" / "subjects"
PHYSICAL = {
    "1L": ("part1_track1", "part1_track2"),
    "1R": ("part1_track1", "part1_track2"),
    "2L": ("part2_track2", "part2_track1"),
    "2R": ("part2_track2", "part2_track1"),
}


def sha256(path: Path) -> str:
    digest = hashlib.sha256()
    with path.open("rb") as stream:
        for chunk in iter(lambda: stream.read(8 * 1024 * 1024), b""):
            digest.update(chunk)
    return digest.hexdigest()


def main() -> None:
    parser = argparse.ArgumentParser()
    parser.add_argument("subject", choices=SUBJECTS)
    args = parser.parse_args()
    subject = args.subject
    output = OUTPUT_ROOT / f"{subject}_P16_drop2_v1.json"
    if output.exists():
        raise RuntimeError(f"Refusing to overwrite geometry result for {subject}.")
    if sha256(PROTOCOL) != PROTOCOL_HASH:
        raise RuntimeError("Geometry development protocol hash mismatch.")
    baseline_path = BASELINE_ROOT / f"{subject}_speech_decoder_v1.json"
    baseline = json.loads(baseline_path.read_text(encoding="utf-8"))
    manifest_path = MANIFEST_ROOT / f"{subject}_production_manifest_v1.json"
    manifest = json.loads(manifest_path.read_text(encoding="utf-8"))
    records = {
        item["recording"].rsplit("_", 1)[-1]: item
        for item in manifest["preprocessing_records"]
    }
    model_path = MODEL_ROOT / f"{subject}_speech_decoder_models_v1.npz"
    with np.load(model_path, allow_pickle=False) as saved:
        saved_models = {name: saved[name] for name in saved.files}
    with np.load(AUDIO_CACHE, allow_pickle=False) as saved:
        full_audio = {name: saved[name] for name in saved.files}
    p16 = partition_ids(load_public_geometry(COORDINATES).xy)["P16"]
    condition_scores = {
        "geometry": {condition: [] for condition in CONDITIONS},
        "global": {condition: [] for condition in CONDITIONS},
    }
    dropped_cells: dict[str, list[list[int]]] = {condition: [] for condition in CONDITIONS}
    for condition in CONDITIONS:
        test_story = condition[0]
        train_story = "2" if test_story == "1" else "1"
        model = {
            "x_mean": saved_models[f"train_story_{train_story}_x_mean"],
            "x_scale": saved_models[f"train_story_{train_story}_x_scale"],
            "weights": saved_models[f"train_story_{train_story}_weights"],
            "y_mean": float(saved_models[f"train_story_{train_story}_y_mean"]),
        }
        with np.load(
            PREPROCESSED_ROOT / subject / f"{subject}_AAD_{condition}_preprocessed_v1.npz",
            allow_pickle=False,
        ) as saved:
            eeg = saved["eeg"]
        record = records[condition]
        offset = int(
            round(
                (
                    record["crop"]["padded_start"]
                    + 3000
                    - record["trigger_train"]["first_sample"]
                )
                / 10.0
            )
        )
        left_name, right_name = PHYSICAL[condition]
        for block_index in range(12):
            first = offset + block_index * 3000
            stop = first + 3000
            left100 = full_audio[left_name][first:stop]
            right100 = full_audio[right_name][first:stop]
            design, left_features = block_design(
                eeg[block_index * 3000 : (block_index + 1) * 3000], left100
            )
            right_features = resample_poly(right100, 1, 5, axis=0)[:595]
            geometry_score, global_score, dropped = p16_drop2_score(
                design,
                model,
                left_features[:, 0],
                right_features[:, 0],
                p16,
            )
            condition_scores["geometry"][condition].append(geometry_score)
            condition_scores["global"][condition].append(global_score)
            dropped_cells[condition].append([int(value) for value in dropped])
    endpoints = {}
    for name in ("global", "geometry"):
        story1 = binary_auc(
            condition_scores[name]["1L"], condition_scores[name]["1R"]
        )
        story2 = binary_auc(
            condition_scores[name]["2L"], condition_scores[name]["2R"]
        )
        endpoints[name] = {
            "story1_auc": story1,
            "story2_auc": story2,
            "mean_auc": (story1 + story2) / 2.0,
        }
    reproduction_error = abs(endpoints["global"]["mean_auc"] - baseline["mean_auc"])
    if reproduction_error > 1e-12:
        raise RuntimeError(
            f"Frozen baseline reproduction failed for {subject}: {reproduction_error}"
        )
    payload = {
        "schema": "KITE-SAT-geometry-development-subject-v1",
        "status": "pass",
        "subject": subject,
        "protocol_sha256": PROTOCOL_HASH,
        "baseline_artifact": {
            "path": baseline_path.relative_to(ROOT).as_posix(),
            "sha256": sha256(baseline_path),
        },
        "model_archive": {
            "path": model_path.relative_to(ROOT).as_posix(),
            "sha256": sha256(model_path),
        },
        "endpoints": endpoints,
        "gain_auc": endpoints["geometry"]["mean_auc"] - endpoints["global"]["mean_auc"],
        "baseline_reproduction_error": reproduction_error,
        "condition_scores": condition_scores,
        "dropped_P16_cells": dropped_cells,
        "firewall": {
            "discarded_geometry_candidate_computed": False,
            "confirmation_subject_read": False,
            "paper_mutated": False,
        },
    }
    OUTPUT_ROOT.mkdir(parents=True, exist_ok=True)
    output.write_text(json.dumps(payload, indent=2) + "\n", encoding="utf-8")
    observed = sha256(output)
    output.with_name(output.name + ".sha256").write_text(
        f"{observed}  {output.name}\n", encoding="ascii"
    )
    print(
        json.dumps(
            {
                "subject": subject,
                "global_auc": endpoints["global"]["mean_auc"],
                "geometry_auc": endpoints["geometry"]["mean_auc"],
                "gain": payload["gain_auc"],
                "sha256": observed,
            }
        )
    )


if __name__ == "__main__":
    main()
```
