## Supplementary material for "Falsifiable substitution tests reveal task-structured neural evidence for auditory attention": Article_Analysis_Source_Code: run_kite_sat_sensor_density_group_v1.py

```
"""Aggregate the frozen post-confirmation sensor-density mechanism analysis."""

from __future__ import annotations

import itertools
import json
from pathlib import Path

import numpy as np

from kite_sat_sensor_density_lock_utils_v1 import (
    ROOT,
    sha256,
    validate_sensor_density_lock,
)
from run_kite_sat_sensor_density_subject_v1 import BUDGETS, OUTPUT_ROOT, SUBJECTS


OUTPUT = OUTPUT_ROOT.parent / "sensor_density_decision_v1.json"


def exact_one_sided(values: np.ndarray) -> float:
    observed = float(np.mean(values))
    exceed = sum(
        float(np.mean(values * np.asarray(signs))) >= observed - 1e-15
        for signs in itertools.product((-1.0, 1.0), repeat=len(values))
    )
    return exceed / (2 ** len(values))


def slope(values: np.ndarray) -> float:
    x = np.log2(np.asarray(BUDGETS, dtype=np.float64) / 32.0)
    centered = x - np.mean(x)
    return float(np.sum(centered * values) / np.sum(centered * centered))


def main() -> None:
    if OUTPUT.exists():
        raise RuntimeError("Refusing to overwrite sensor-density group result.")
    lock, lock_hash = validate_sensor_density_lock(Path(__file__))
    if tuple(lock["subject_order"]) != SUBJECTS:
        raise RuntimeError("Sensor-density subject order differs from lock.")
    records = []
    hashes = {}
    for subject in SUBJECTS:
        path = OUTPUT_ROOT / f"{subject}_sensor_density_v1.json"
        record = json.loads(path.read_text(encoding="utf-8"))
        sidecar = path.with_name(path.name + ".sha256").read_text(
            encoding="ascii"
        ).strip().split()
        hashes[subject] = sha256(path)
        if (
            record["status"] != "pass"
            or record["subject"] != subject
            or record["sensor_density_implementation_lock_sha256"] != lock_hash
            or [row["budget"] for row in record["budget_results"]] != list(BUDGETS)
            or max(record["full_budget_primary_absolute_differences"].values())
            > 1e-12
            or len(sidecar) != 2
            or sidecar[0].lower() != hashes[subject]
            or sidecar[1] != path.name
            or record["firewall"]["sensor_subset_selected_by_outcome"]
            or record["firewall"]["ridge_or_AKB_parameter_changed"]
            or record["firewall"]["paper_mutated"]
        ):
            raise RuntimeError(f"Invalid sensor-density subject result: {subject}")
        records.append(record)
    gains = np.asarray(
        [
            [row["AKB_minus_global_auc"] for row in record["budget_results"]]
            for record in records
        ],
        dtype=np.float64,
    )
    topology = np.asarray(
        [
            [
                row["identity_minus_mean_topology_controls_auc"]
                for row in record["budget_results"]
            ]
            for record in records
        ],
        dtype=np.float64,
    )
    global_auc = np.asarray(
        [
            [row["global_mean_auc"] for row in record["budget_results"]]
            for record in records
        ]
    )
    akb_auc = np.asarray(
        [
            [row["AKB_mean_auc"] for row in record["budget_results"]]
            for record in records
        ]
    )
    gain_slopes = np.asarray([slope(row) for row in gains])
    topology_slopes = np.asarray([slope(row) for row in topology])
    mean_gains = np.mean(gains, axis=0)
    mean_topology = np.mean(topology, axis=0)
    budget_metrics = []
    for index, budget in enumerate(BUDGETS):
        budget_metrics.append(
            {
                "budget": budget,
                "mean_global_auc": float(np.mean(global_auc[:, index])),
                "mean_AKB_auc": float(np.mean(akb_auc[:, index])),
                "mean_AKB_minus_global_auc": float(mean_gains[index]),
                "AKB_minus_global_exact_one_sided_p": exact_one_sided(
                    gains[:, index]
                ),
                "participants_positive_gain": int(np.sum(gains[:, index] > 0)),
                "mean_identity_minus_controls_auc": float(mean_topology[index]),
                "identity_minus_controls_exact_one_sided_p": exact_one_sided(
                    topology[:, index]
                ),
                "participants_positive_topology_advantage": int(
                    np.sum(topology[:, index] > 0)
                ),
            }
        )
    predictions = {
        "AKB_gain_density_slope_gt_0": {
            "mean_slope_auc_per_doubling": float(np.mean(gain_slopes)),
            "exact_one_sided_p": exact_one_sided(gain_slopes),
            "participants_positive": int(np.sum(gain_slopes > 0)),
        },
        "topology_advantage_density_slope_gt_0": {
            "mean_slope_auc_per_doubling": float(np.mean(topology_slopes)),
            "exact_one_sided_p": exact_one_sided(topology_slopes),
            "participants_positive": int(np.sum(topology_slopes > 0)),
        },
        "group_mean_gain_monotone_nondecreasing": bool(
            np.all(np.diff(mean_gains) >= -1e-15)
        ),
        "group_mean_topology_monotone_nondecreasing": bool(
            np.all(np.diff(mean_topology) >= -1e-15)
        ),
    }
    payload = {
        "schema": "KITE-SAT-sensor-density-mechanism-decision-v1",
        "status": "pass",
        "inferential_status": "post-confirmation_mechanistic",
        "sensor_density_implementation_lock_sha256": lock_hash,
        "subject_order": list(SUBJECTS),
        "subject_artifact_sha256": hashes,
        "budget_metrics": budget_metrics,
        "frozen_predictions": predictions,
        "participant_slopes": [
            {
                "subject": subject,
                "AKB_gain_slope_per_doubling": float(gain_slopes[index]),
                "topology_advantage_slope_per_doubling": float(
                    topology_slopes[index]
                ),
            }
            for index, subject in enumerate(SUBJECTS)
        ],
        "participant_budget_gain_matrix": gains.tolist(),
        "participant_budget_topology_matrix": topology.tolist(),
        "claim_boundary": "Exploratory post-confirmation density mechanism; primary confirmatory p values are unchanged.",
        "paper_mutated": False,
    }
    OUTPUT.write_text(json.dumps(payload, indent=2) + "\n", encoding="utf-8")
    observed = sha256(OUTPUT)
    OUTPUT.with_name(OUTPUT.name + ".sha256").write_text(
        f"{observed}  {OUTPUT.name}\n", encoding="ascii"
    )
    print(
        json.dumps(
            {
                "status": "pass",
                "budget_metrics": budget_metrics,
                "predictions": predictions,
                "sha256": observed,
            }
        )
    )


if __name__ == "__main__":
    main()
```
