## Supplementary material for "Falsifiable substitution tests reveal task-structured neural evidence for auditory attention": Article_Analysis_Source_Code: run_kite_sat_sensor_density_subject_v1.py

```
"""Post-confirmation nested sensor-density mechanism analysis for one subject."""

from __future__ import annotations

import argparse
import json
from pathlib import Path

import numpy as np
from scipy.linalg import cho_factor, cho_solve
from scipy.signal import resample_poly

from kite_sat_adaptive_broad_v1 import adaptive_score, family_weights, reliability
from kite_sat_backward_v1 import block_design
from kite_sat_direction_delay_controls_v1 import TRANSFORMS, transformed_matrix
from kite_sat_geometry_v1 import cell_delta_contributions, partition_ids
from kite_sat_sensor_density_lock_utils_v1 import (
    ROOT,
    sha256,
    validate_sensor_density_lock,
)
from kite_st_hd_core_v1 import load_public_geometry
from kite_st_hd_statistics_v1 import binary_auc


SUBJECTS = (
    "S0", "S10", "S2", "S21", "S28", "S22",
    "S9", "S15", "S16", "S18", "S13", "S1",
)
BUDGETS = (32, 64, 128, 253)
CONDITIONS = ("1L", "1R", "2L", "2R")
PREPROCESSED_ROOT = (
    ROOT / "data" / "derived" / "kite_st_hd_zenodo4518754"
    / "confirmation_preprocessed_v1"
)
MANIFEST_ROOT = ROOT / "results" / "kite_st_hd" / "confirmation_v1" / "subjects"
PRIMARY_ROOT = (
    ROOT / "results" / "kite_sat" / "akb_confirmation_complete_case_v2"
)
AUDIO_CACHE = (
    ROOT / "data" / "derived" / "kite_st_hd_zenodo4518754"
    / "kite_sat_audio_features_v1.npz"
)
COORDINATES = (
    ROOT / "downloads" / "candidate_audit" / "zenodo4518754_support"
    / "misc" / "misc" / "eeg255ch_locs.csv"
)
SUBSETS = ROOT / "docs" / "kite_sat_nested_sensor_budgets_v1.json"
OUTPUT_ROOT = (
    ROOT / "results" / "kite_sat" / "sensor_density_mechanism_v1" / "subjects"
)
PHYSICAL = {
    "1L": ("part1_track1", "part1_track2"),
    "1R": ("part1_track1", "part1_track2"),
    "2L": ("part2_track2", "part2_track1"),
    "2R": ("part2_track2", "part2_track1"),
}


def fit_generic(
    training_designs: list[np.ndarray],
    training_targets: list[np.ndarray],
) -> dict[str, np.ndarray | float]:
    x = np.vstack(training_designs).astype(np.float64, copy=False)
    y = np.concatenate(training_targets).astype(np.float64, copy=False)
    if x.ndim != 2 or x.shape[0] != len(y):
        raise ValueError("Unexpected generic density-decoder training shape.")
    x_mean = np.mean(x, axis=0)
    x_scale = np.std(x, axis=0)
    x_scale = np.where(x_scale > 1e-12, x_scale, 1.0)
    y_mean = float(np.mean(y))
    standardized = (x - x_mean) / x_scale
    centered = y - y_mean
    gram = standardized.T @ standardized
    cross = standardized.T @ centered
    energy = float(np.mean(np.diag(gram)))
    alpha = energy
    gram.flat[:: gram.shape[0] + 1] += alpha
    factor = cho_factor(gram, lower=True, check_finite=False)
    weights = cho_solve(factor, cross, check_finite=False)
    if not np.isfinite(weights).all():
        raise RuntimeError("Nonfinite sensor-density decoder weights.")
    return {
        "x_mean": x_mean,
        "x_scale": x_scale,
        "y_mean": y_mean,
        "weights": weights,
        "gram_energy": energy,
        "alpha": alpha,
    }


def main() -> None:
    parser = argparse.ArgumentParser()
    parser.add_argument("subject", choices=SUBJECTS)
    args = parser.parse_args()
    subject = args.subject
    output = OUTPUT_ROOT / f"{subject}_sensor_density_v1.json"
    if output.exists():
        raise RuntimeError(f"Refusing to overwrite sensor-density result: {subject}")
    lock, lock_hash = validate_sensor_density_lock(Path(__file__))
    if tuple(lock["subject_order"]) != SUBJECTS:
        raise RuntimeError("Sensor-density subject order differs from lock.")
    decision_path = PRIMARY_ROOT / "confirmation_decision_v1.json"
    audit_path = PRIMARY_ROOT / "confirmation_outcome_audit_v1.json"
    decision = json.loads(decision_path.read_text(encoding="utf-8"))
    audit = json.loads(audit_path.read_text(encoding="utf-8"))
    if (
        decision["status"] != "intact_and_patch_confirmed"
        or audit["status"] != "pass"
        or audit["confirmation_decision_sha256"] != sha256(decision_path)
        or decision["subject_order"] != list(SUBJECTS)
    ):
        raise RuntimeError("Audited complete-case primary decision is unavailable.")
    primary_path = PRIMARY_ROOT / "subjects" / f"{subject}_AKB_confirmation_v1.json"
    primary = json.loads(primary_path.read_text(encoding="utf-8"))
    if sha256(primary_path) != decision["subject_artifact_sha256"][subject]:
        raise RuntimeError("Primary subject artifact mismatch.")
    subset_payload = json.loads(SUBSETS.read_text(encoding="utf-8"))
    if (
        subset_payload["status"] != "outcome_independent_geometry_only"
        or [row["budget"] for row in subset_payload["budgets"]] != list(BUDGETS)
    ):
        raise RuntimeError("Nested sensor subset artifact mismatch.")
    selected = {
        row["budget"]: np.asarray(row["sorted_indices"], dtype=np.int64)
        for row in subset_payload["budgets"]
    }
    manifest_path = MANIFEST_ROOT / f"{subject}_production_manifest_v1.json"
    manifest = json.loads(manifest_path.read_text(encoding="utf-8"))
    preprocessing = {
        item["recording"].rsplit("_", 1)[-1]: item
        for item in manifest["preprocessing_records"]
    }
    with np.load(AUDIO_CACHE, allow_pickle=False) as saved:
        full_audio = {name: saved[name] for name in saved.files}
    geometry = load_public_geometry(COORDINATES)
    full_s16 = partition_ids(geometry.xy)["S16"]
    designs: dict[str, list[np.ndarray]] = {}
    audio_blocks: dict[str, list[tuple[np.ndarray, np.ndarray]]] = {}
    attended: dict[str, list[np.ndarray]] = {}
    for condition in CONDITIONS:
        path = (
            PREPROCESSED_ROOT / subject
            / f"{subject}_AAD_{condition}_preprocessed_v1.npz"
        )
        if sha256(path) != primary["preprocessed_sha256"][condition]:
            raise RuntimeError(f"Sensor-density input mismatch: {subject} {condition}")
        with np.load(path, allow_pickle=False) as saved:
            eeg = saved["eeg"]
        record = preprocessing[condition]
        offset = int(
            round(
                (
                    record["crop"]["padded_start"]
                    + 3000
                    - record["trigger_train"]["first_sample"]
                )
                / 10.0
            )
        )
        left_name, right_name = PHYSICAL[condition]
        designs[condition], audio_blocks[condition], attended[condition] = [], [], []
        for block_index in range(12):
            first = offset + block_index * 3000
            stop = first + 3000
            left100 = full_audio[left_name][first:stop]
            right100 = full_audio[right_name][first:stop]
            design, left_features = block_design(
                eeg[block_index * 3000 : (block_index + 1) * 3000],
                left100,
            )
            right_features = resample_poly(right100, 1, 5, axis=0)[:595]
            left, right = left_features[:, 0], right_features[:, 0]
            designs[condition].append(design)
            audio_blocks[condition].append((left, right))
            attended[condition].append(
                left if condition.endswith("L") else right
            )

    budget_results = []
    for budget in BUDGETS:
        feature_indices = np.concatenate(
            [lag * 253 + selected[budget] for lag in range(6)]
        )
        ids = full_s16[feature_indices]
        if np.any(np.bincount(ids, minlength=16) == 0):
            raise RuntimeError(f"Empty S16 cell at budget {budget}.")
        budget_designs = {
            condition: [design[:, feature_indices] for design in designs[condition]]
            for condition in CONDITIONS
        }
        scores = {
            condition: {
                "global": [],
                "transforms": [[] for _ in TRANSFORMS],
            }
            for condition in CONDITIONS
        }
        fold_reliabilities = {}
        for train_story, test_story in (("1", "2"), ("2", "1")):
            train_conditions = (f"{train_story}L", f"{train_story}R")
            held_aligned = []
            for fit_parity in (0, 1):
                held_parity = 1 - fit_parity
                calibration = fit_generic(
                    [
                        budget_designs[condition][index]
                        for condition in train_conditions
                        for index in range(12)
                        if index % 2 == fit_parity
                    ],
                    [
                        attended[condition][index]
                        for condition in train_conditions
                        for index in range(12)
                        if index % 2 == fit_parity
                    ],
                )
                for condition in train_conditions:
                    alignment = 1.0 if condition.endswith("L") else -1.0
                    for index in range(12):
                        if index % 2 != held_parity:
                            continue
                        left, right = audio_blocks[condition][index]
                        delta, _ = cell_delta_contributions(
                            budget_designs[condition][index],
                            calibration,
                            left,
                            right,
                            ids,
                        )
                        held_aligned.append(alignment * delta)
            fold_reliability = reliability(np.asarray(held_aligned))
            fold_reliabilities[f"train_story_{train_story}"] = [
                float(value) for value in fold_reliability
            ]
            weight_matrix = transformed_matrix(
                family_weights(fold_reliability, "softmax2")
            )
            full_model = fit_generic(
                [
                    budget_designs[condition][index]
                    for condition in train_conditions
                    for index in range(12)
                ],
                [
                    attended[condition][index]
                    for condition in train_conditions
                    for index in range(12)
                ],
            )
            for condition in (f"{test_story}L", f"{test_story}R"):
                for design, (left, right) in zip(
                    budget_designs[condition], audio_blocks[condition]
                ):
                    delta, global_margin = cell_delta_contributions(
                        design, full_model, left, right, ids
                    )
                    scores[condition]["global"].append(global_margin)
                    transformed_scores = weight_matrix @ delta
                    for transform_index, value in enumerate(transformed_scores):
                        scores[condition]["transforms"][transform_index].append(
                            float(value)
                        )
        global_story1 = binary_auc(
            scores["1L"]["global"], scores["1R"]["global"]
        )
        global_story2 = binary_auc(
            scores["2L"]["global"], scores["2R"]["global"]
        )
        transform_aucs = []
        for transform_index, (shift, swap) in enumerate(TRANSFORMS):
            story1 = binary_auc(
                scores["1L"]["transforms"][transform_index],
                scores["1R"]["transforms"][transform_index],
            )
            story2 = binary_auc(
                scores["2L"]["transforms"][transform_index],
                scores["2R"]["transforms"][transform_index],
            )
            transform_aucs.append(
                {
                    "angular_shift": shift,
                    "delay_swap": bool(swap),
                    "story1_auc": story1,
                    "story2_auc": story2,
                    "mean_auc": (story1 + story2) / 2,
                }
            )
        global_mean = (global_story1 + global_story2) / 2
        identity = transform_aucs[0]["mean_auc"]
        control_mean = float(
            np.mean([row["mean_auc"] for row in transform_aucs[1:]])
        )
        budget_results.append(
            {
                "budget": budget,
                "selected_indices": [int(value) for value in selected[budget]],
                "feature_count": len(feature_indices),
                "cell_feature_counts": [
                    int(value)
                    for value in np.bincount(ids, minlength=16)
                ],
                "global_story1_auc": global_story1,
                "global_story2_auc": global_story2,
                "global_mean_auc": global_mean,
                "AKB_story1_auc": transform_aucs[0]["story1_auc"],
                "AKB_story2_auc": transform_aucs[0]["story2_auc"],
                "AKB_mean_auc": identity,
                "AKB_minus_global_auc": identity - global_mean,
                "identity_minus_mean_topology_controls_auc": (
                    identity - control_mean
                ),
                "transform_aucs": transform_aucs,
                "fold_reliabilities": fold_reliabilities,
            }
        )
    full = next(row for row in budget_results if row["budget"] == 253)
    equivalence = {
        "global": abs(
            full["global_mean_auc"]
            - primary["intact_endpoints"]["global"]["mean_auc"]
        ),
        "AKB": abs(
            full["AKB_mean_auc"]
            - primary["intact_endpoints"]["AKB"]["mean_auc"]
        ),
        "gain": abs(full["AKB_minus_global_auc"] - primary["intact_gain_auc"]),
    }
    if max(equivalence.values()) > 1e-12:
        raise RuntimeError("Full sensor budget does not reproduce primary endpoint.")
    payload = {
        "schema": "KITE-SAT-sensor-density-mechanism-subject-v1",
        "status": "pass",
        "subject": subject,
        "sensor_density_implementation_lock_sha256": lock_hash,
        "primary_subject_artifact_sha256": sha256(primary_path),
        "subset_artifact_sha256": sha256(SUBSETS),
        "budget_results": budget_results,
        "full_budget_primary_absolute_differences": equivalence,
        "firewall": {
            "sensor_subset_selected_by_outcome": False,
            "ridge_or_AKB_parameter_changed": False,
            "paper_mutated": False,
        },
    }
    OUTPUT_ROOT.mkdir(parents=True, exist_ok=True)
    output.write_text(json.dumps(payload, indent=2) + "\n", encoding="utf-8")
    observed = sha256(output)
    output.with_name(output.name + ".sha256").write_text(
        f"{observed}  {output.name}\n", encoding="ascii"
    )
    print(
        json.dumps(
            {
                "subject": subject,
                "gains": {
                    str(row["budget"]): row["AKB_minus_global_auc"]
                    for row in budget_results
                },
                "topology": {
                    str(row["budget"]): row[
                        "identity_minus_mean_topology_controls_auc"
                    ]
                    for row in budget_results
                },
                "sha256": observed,
            }
        )
    )


if __name__ == "__main__":
    main()
```
