## Supplementary material for "Falsifiable substitution tests reveal task-structured neural evidence for auditory attention": Article_Analysis_Source_Code: run_kite_st_hd_confirmation_preprocessing_s3_packaging_recovery_v1.py

```
"""Frozen recovery for the S3 archive's missing, unused Curry .ceo sidecar.

This script exists only because the public S3 archive omits
S3/S3_AAD_1L.ceo.  The scientific preprocessing reads .dap, .rs3 and .dat;
it never reads .ceo.  The exception is bound by the accompanying packaging
addendum and refuses every archive layout other than the exact observed one.
"""

from __future__ import annotations

import json
import shutil
import tarfile
from pathlib import Path, PurePosixPath

import run_kite_st_hd_confirmation_preprocessing_subject_v1 as frozen
from kite_sat_confirmation_lock_utils_v1 import ROOT, sha256, validate_confirmation_lock


SUBJECT = "S3"
MISSING_UNUSED_MEMBER = "S3/S3_AAD_1L.ceo"
ADDENDUM = ROOT / "docs" / "kite_sat_akb_confirmation_s3_packaging_addendum_v1.json"
STARTED_SHA256 = "d2d3b38965b9fd073ec25e0eb1ec90e785c78de96fdcbd6c3e71fad856f9cb91"
ARCHIVE_SHA256 = "00ac2ac5089fc7f14d6d0c28777891ef77f2fa82c6f7dc2b4e3be5345c3f4bdd"


def validate_addendum() -> tuple[dict, str]:
    sidecar = ADDENDUM.with_name(ADDENDUM.name + ".sha256")
    fields = sidecar.read_text(encoding="ascii").strip().split()
    observed = sha256(ADDENDUM)
    if len(fields) != 2 or fields[0].lower() != observed or fields[1] != ADDENDUM.name:
        raise RuntimeError("S3 packaging addendum sidecar mismatch.")
    payload = json.loads(ADDENDUM.read_text(encoding="utf-8"))
    relative = Path(__file__).resolve().relative_to(ROOT.resolve()).as_posix()
    if (
        payload["status"] != "frozen_before_any_confirmation_attention_or_AKB_score"
        or payload["subject"] != SUBJECT
        or payload["missing_member"] != MISSING_UNUSED_MEMBER
        or payload["recovery_source"]["path"] != relative
        or payload["recovery_source"]["sha256"] != sha256(Path(__file__))
    ):
        raise RuntimeError("S3 packaging addendum/source mismatch.")
    return payload, observed


def exact_extract(archive: Path) -> dict:
    work_subject = frozen.WORK_ROOT / SUBJECT
    if work_subject.exists():
        raise RuntimeError(f"Refusing existing S3 work directory: {work_subject}")
    expected = frozen.expected_member_names(SUBJECT)
    expected.remove(MISSING_UNUSED_MEMBER)
    frozen.WORK_ROOT.mkdir(parents=True, exist_ok=True)
    with tarfile.open(archive, mode="r:gz") as stream:
        members = stream.getmembers()
        observed = {member.name for member in members}
        if observed != expected:
            raise RuntimeError("S3 archive differs from the exact packaging exception.")
        for member in members:
            parsed = PurePosixPath(member.name)
            if parsed.is_absolute() or ".." in parsed.parts:
                raise RuntimeError(f"Unsafe S3 archive path: {member.name}")
            if member.issym() or member.islnk() or member.isdev():
                raise RuntimeError(f"Unsupported S3 archive member: {member.name}")
            if parsed.parts[0] != SUBJECT:
                raise RuntimeError(f"S3 member escapes root: {member.name}")
        stream.extractall(frozen.WORK_ROOT, members=members, filter="data")
    files = sorted(
        path.relative_to(frozen.WORK_ROOT).as_posix()
        for path in work_subject.rglob("*")
        if path.is_file()
    )
    if len(files) != 15:
        raise RuntimeError("S3 exact packaging exception did not yield 15 files.")
    required_used = {
        f"{SUBJECT}/{SUBJECT}_AAD_{condition}{extension}"
        for condition in frozen.CONDITIONS
        for extension in (".dap", ".dat", ".rs3")
    }
    if not required_used.issubset(observed):
        raise RuntimeError("S3 is missing a preprocessing-consumed member.")
    return {
        "member_count": len(members),
        "file_count": len(files),
        "files": files,
        "exact_missing_unused_member": MISSING_UNUSED_MEMBER,
        "all_preprocessing_consumed_members_present": True,
    }


def remove_work_subject() -> None:
    target = (frozen.WORK_ROOT / SUBJECT).resolve()
    if target.parent != frozen.WORK_ROOT.resolve() or target.name != SUBJECT:
        raise RuntimeError("Refusing unsafe S3 cleanup target.")
    if not target.is_dir():
        raise RuntimeError("S3 cleanup target is absent.")
    shutil.rmtree(target)
    if target.exists():
        raise RuntimeError("S3 cleanup did not complete.")


def main() -> None:
    original_script = Path(frozen.__file__).resolve()
    lock, lock_hash = validate_confirmation_lock(original_script)
    addendum, addendum_hash = validate_addendum()
    if tuple(lock["confirmation_subject_order"]) != frozen.SUBJECTS:
        raise RuntimeError("Confirmation subject list differs from original lock.")
    archive = frozen.ARCHIVE_ROOT / f"{SUBJECT}.tar.gz"
    acquisition_path = frozen.METADATA_ROOT / f"{SUBJECT}_archive_acquisition_v1.json"
    acquisition = json.loads(acquisition_path.read_text(encoding="utf-8"))
    expected_bytes, expected_md5 = frozen.FILES[SUBJECT]
    if (
        archive.stat().st_size != expected_bytes
        or acquisition["bytes"] != expected_bytes
        or acquisition["md5"] != expected_md5
        or acquisition["sha256"] != ARCHIVE_SHA256
        or sha256(archive) != ARCHIVE_SHA256
        or acquisition["confirmation_implementation_lock_sha256"] != lock_hash
    ):
        raise RuntimeError("S3 archive/acquisition contract mismatch.")
    started = frozen.RESULT_ROOT / f"{SUBJECT}_preprocessing_started_v1.json"
    final_manifest = frozen.RESULT_ROOT / f"{SUBJECT}_production_manifest_v1.json"
    output_dir = frozen.PREPROCESSED_ROOT / SUBJECT
    feature_path = frozen.FEATURE_ROOT / f"{SUBJECT}_features_v1.npz"
    if sha256(started) != STARTED_SHA256:
        raise RuntimeError("S3 original one-shot start record mismatch.")
    started_payload = json.loads(started.read_text(encoding="utf-8"))
    if (
        started_payload["subject"] != SUBJECT
        or started_payload["archive_sha256"] != ARCHIVE_SHA256
        or started_payload["confirmation_implementation_lock_sha256"] != lock_hash
    ):
        raise RuntimeError("S3 original one-shot start payload mismatch.")
    if any(
        path.exists()
        for path in (
            final_manifest,
            output_dir,
            feature_path,
            frozen.WORK_ROOT / SUBJECT,
        )
    ):
        raise RuntimeError("S3 recovery refuses any partial scientific output.")
    extraction = exact_extract(archive)
    preprocessing_records = frozen.preprocess_subject_v2(
        SUBJECT, frozen.WORK_ROOT, output_dir
    )
    feature_record = frozen.build_feature_archive(
        SUBJECT, preprocessing_records, feature_path
    )
    remove_work_subject()
    maximum_bad = max(
        record["bad_channels"]["count"] for record in preprocessing_records
    )
    payload = {
        "schema": "KITE-ST-HD-confirmation-subject-preprocessing-v1",
        "status": "pass",
        "subject": SUBJECT,
        "confirmation_implementation_lock_sha256": lock_hash,
        "archive": {
            "path": archive.relative_to(ROOT).as_posix(),
            "bytes": expected_bytes,
            "md5": expected_md5,
            "sha256": ARCHIVE_SHA256,
        },
        "extraction": extraction,
        "preprocessing_records": preprocessing_records,
        "maximum_bad_channel_count": maximum_bad,
        "maximum_bad_channel_rule": 15,
        "feature_archive": feature_record,
        "temporary_extracted_work_removed_after_verified_derivatives": True,
        "packaging_exception": {
            "kind": "missing_unused_Curry_ceo_sidecar",
            "missing_member": MISSING_UNUSED_MEMBER,
            "scientific_preprocessing_code_changed": False,
            "preprocessing_consumed_extensions": [".dap", ".dat", ".rs3"],
            "addendum_path": ADDENDUM.relative_to(ROOT).as_posix(),
            "addendum_sha256": addendum_hash,
            "recovery_source_sha256": addendum["recovery_source"]["sha256"],
            "original_failed_before_extraction_or_numeric_read": True,
        },
        "firewall": {
            "attention_contrast_computed": False,
            "AKB_score_computed": False,
            "paper_mutated": False,
        },
    }
    observed = frozen.write_subject_manifest(final_manifest, payload)
    print(
        json.dumps(
            {
                "status": "pass",
                "subject": SUBJECT,
                "maximum_bad_channels": maximum_bad,
                "manifest_sha256": observed,
                "packaging_addendum_sha256": addendum_hash,
            }
        )
    )


if __name__ == "__main__":
    main()
```
