## Supplementary material for "Falsifiable substitution tests reveal task-structured neural evidence for auditory attention": Article_Analysis_Source_Code: run_kite_st_hd_confirmation_qc_record_v1.py

```
#!/usr/bin/env python3
"""Execute and record the two prespecified KITE confirmation QC failures.

The frozen participant script is not modified.  This orchestration layer runs
it as a subprocess, verifies the expected label-blind failure, removes only
partial preprocessing products inside the workflow workspace, and writes the
machine-readable technical-failure record consumed by the complete-case gate.
"""

from __future__ import annotations

import argparse
import ast
import json
import re
import shutil
import subprocess
import sys
from pathlib import Path

import run_kite_st_hd_confirmation_preprocessing_subject_v1 as frozen
from kite_sat_confirmation_lock_utils_v1 import ROOT, sha256, validate_confirmation_lock


EXPECTED = {"S26": 16, "S3": 19}
ORIGINAL = ROOT / "analysis" / "run_kite_st_hd_confirmation_preprocessing_subject_v1.py"
RECOVERY = ROOT / "analysis" / "run_kite_st_hd_confirmation_preprocessing_s3_packaging_recovery_v1.py"
LOG_ROOT = ROOT / "workflow_failure_logs"


def execute(script: Path, subject: str | None, label: str) -> tuple[int, str, Path]:
    command = [sys.executable, str(script)]
    if subject is not None:
        command.append(subject)
    completed = subprocess.run(
        command,
        cwd=ROOT,
        stdout=subprocess.PIPE,
        stderr=subprocess.STDOUT,
        text=True,
        encoding="utf-8",
        errors="replace",
    )
    print(completed.stdout, end="", flush=True)
    LOG_ROOT.mkdir(parents=True, exist_ok=True)
    log = LOG_ROOT / f"{label}.log"
    log.write_text(completed.stdout, encoding="utf-8")
    return completed.returncode, completed.stdout, log


def safe_remove(path: Path, allowed_parent: Path) -> None:
    resolved = path.resolve()
    if resolved.parent != allowed_parent.resolve():
        raise RuntimeError(f"refusing unsafe technical-failure cleanup: {path}")
    if resolved.is_dir():
        shutil.rmtree(resolved)
    elif resolved.exists():
        resolved.unlink()


def parse_bad_channels(output: str, expected_count: int) -> list[int]:
    matches = re.findall(
        r"More than 15 bad scalp channels:\s*(\[[0-9, ]+\])", output
    )
    if len(matches) != 1:
        raise RuntimeError("expected one frozen bad-channel failure")
    values = ast.literal_eval(matches[0])
    if not isinstance(values, list) or len(values) != expected_count:
        raise RuntimeError(f"unexpected bad-channel count: {values}")
    return [int(value) for value in values]


def main() -> None:
    parser = argparse.ArgumentParser()
    parser.add_argument("subject", choices=tuple(EXPECTED))
    args = parser.parse_args()
    subject = args.subject
    output = frozen.RESULT_ROOT / f"{subject}_technical_failure_v1.json"
    if output.exists():
        raise RuntimeError(f"refusing to overwrite technical-failure record: {output}")

    original_code, original_text, original_log = execute(
        ORIGINAL, subject, f"KITE_confirmation_{subject}_original"
    )
    if original_code == 0:
        raise RuntimeError(f"{subject} unexpectedly passed frozen preprocessing")

    event_records: list[dict[str, object]] = []
    failure_text = original_text
    failure_log = original_log
    if subject == "S3":
        if "Confirmation archive members differ for S3" not in original_text:
            raise RuntimeError("S3 did not show the exact packaging exception")
        event_records.append(
            {
                "stage": "archive_member_validation",
                "outcome": "failed_before_extraction_or_numeric_read",
                "reason": "S3/S3_AAD_1L.ceo is absent from the public archive.",
                "log": original_log.relative_to(ROOT).as_posix(),
                "log_sha256": sha256(original_log),
            }
        )
        recovery_code, failure_text, failure_log = execute(
            RECOVERY, None, "KITE_confirmation_S3_packaging_recovery"
        )
        if recovery_code == 0:
            raise RuntimeError("S3 recovery unexpectedly passed frozen QC")

    bad = parse_bad_channels(failure_text, EXPECTED[subject])
    event_records.append(
        {
            "stage": "frozen_label_blind_signal_QC",
            "outcome": "technical_failure",
            "bad_channel_indices": bad,
            "bad_channel_count": len(bad),
            "maximum_allowed": 15,
            "log": failure_log.relative_to(ROOT).as_posix(),
            "log_sha256": sha256(failure_log),
        }
    )

    safe_remove(frozen.WORK_ROOT / subject, frozen.WORK_ROOT)
    safe_remove(frozen.PREPROCESSED_ROOT / subject, frozen.PREPROCESSED_ROOT)
    feature = frozen.FEATURE_ROOT / f"{subject}_features_v1.npz"
    if feature.exists():
        safe_remove(feature, frozen.FEATURE_ROOT)

    lock, lock_hash = validate_confirmation_lock(ORIGINAL)
    started = frozen.RESULT_ROOT / f"{subject}_preprocessing_started_v1.json"
    acquisition_path = frozen.METADATA_ROOT / f"{subject}_archive_acquisition_v1.json"
    acquisition = json.loads(acquisition_path.read_text(encoding="utf-8"))
    canonical = (
        ROOT / "frozen_inputs" / "KITE_ST_HD" / "confirmation_QC"
        / f"{subject}_technical_failure_v1.json"
    )
    canonical_payload = json.loads(canonical.read_text(encoding="utf-8"))
    common_provenance_mismatch = (
        canonical_payload["schema"]
        != "KITE-ST-HD-confirmation-technical-failure-v1"
        or canonical_payload["status"] != "technical_failure"
        or canonical_payload["subject"] != subject
        or canonical_payload["confirmation_protocol_sha256"]
        != lock["protocol_sha256"]
        or canonical_payload["confirmation_implementation_lock_sha256"]
        != lock_hash
        or canonical_payload["archive_sha256"] != acquisition["sha256"]
    )
    if common_provenance_mismatch:
        raise RuntimeError("canonical QC provenance does not match this rerun")
    if subject == "S26":
        if (
            canonical_payload.get("preprocessing_started_manifest_sha256")
            != sha256(started)
        ):
            raise RuntimeError("canonical S26 start-manifest hash mismatch")
        canonical_bad = canonical_payload["bad_channel_indices"]
    else:
        addendum = canonical_payload.get("packaging_addendum", {})
        addendum_path = ROOT / addendum.get("path", "__missing__")
        if (
            not addendum_path.is_file()
            or addendum.get("sha256") != sha256(addendum_path)
        ):
            raise RuntimeError("canonical S3 packaging-addendum mismatch")
        qc_events = [
            event for event in canonical_payload["events"]
            if "bad_channel_indices" in event
        ]
        if len(qc_events) != 1:
            raise RuntimeError("canonical S3 QC event is not unique")
        canonical_bad = qc_events[0]["bad_channel_indices"]
    if canonical_bad != bad:
        raise RuntimeError("recomputed bad-channel set differs from canonical QC")

    output.parent.mkdir(parents=True, exist_ok=True)
    output.write_bytes(canonical.read_bytes())
    observed = sha256(output)
    output.with_name(output.name + ".sha256").write_text(
        f"{observed}  {output.name}\n", encoding="ascii"
    )
    print(
        json.dumps(
            {
                "status": "technical_failure_recorded",
                "subject": subject,
                "bad_channel_count": len(bad),
                "record_sha256": observed,
            }
        )
    )


if __name__ == "__main__":
    main()
```
