## Supplementary material for "Falsifiable substitution tests reveal task-structured neural evidence for auditory attention": Article_Analysis_Source_Code: run_kite_st_hd_development_analysis_v1.py

```
"""One-shot, gate-driven KITE-ST-HD development analysis."""

from __future__ import annotations

import json
from pathlib import Path

import numpy as np

from kite_st_hd_core_v1 import (
    all_sign_patterns,
    cross_story_edge_score,
    exact_upper_p,
    fixed_integration_points,
    select_edges,
    stable_seed,
)
from kite_st_hd_geometry_v1 import (
    MATCHED_FAMILY_COUNT,
    NESTED_INTEGRATION_POINT_COUNT,
    PRIMARY_RADIUS,
    SELECTED_EDGE_COUNT,
    family_union_fraction,
    matched_geometry_distribution,
)
from kite_st_hd_lock_utils_v1 import ROOT, confirmation_numeric_files, sha256, validate_implementation_lock
from kite_st_hd_statistics_v1 import (
    exact_candidate_familywise,
    frozen_template_auc,
    leave_one_participant_out_cross_story_auc,
    local_erasure_frozen_template_auc,
    observed_candidate_fit,
    participant_story_effects,
    residualize_order,
    story_order_contrasts,
)


SUBJECTS = (
    "S7", "S4", "S24", "S11", "S23", "S14",
    "S29", "S19", "S17", "S12", "S25", "S27",
)
CONDITIONS = ("1L", "1R", "2L", "2R")
RESULT_ROOT = ROOT / "results" / "kite_st_hd" / "development"
SUBJECT_RESULT_ROOT = RESULT_ROOT / "subjects"
STARTED = RESULT_ROOT / "development_analysis_started_v1.json"
RESULT = RESULT_ROOT / "development_gate_v1.json"
NULLS = RESULT_ROOT / "development_null_distributions_v1.npz"
TEMPLATE = RESULT_ROOT / "development_template_v1.npz"


def jsonable(value):
    if isinstance(value, np.ndarray):
        return value.tolist()
    if isinstance(value, (np.integer,)):
        return int(value)
    if isinstance(value, (np.floating,)):
        return float(value)
    if isinstance(value, dict):
        return {str(key): jsonable(item) for key, item in value.items()}
    if isinstance(value, (list, tuple)):
        return [jsonable(item) for item in value]
    return value


def write_json_once(path: Path, payload: dict) -> str:
    if path.exists():
        raise RuntimeError(f"Refusing to overwrite: {path}")
    path.parent.mkdir(parents=True, exist_ok=True)
    path.write_text(json.dumps(jsonable(payload), indent=2) + "\n", encoding="utf-8")
    observed = sha256(path)
    path.with_name(path.name + ".sha256").write_text(
        f"{observed}  {path.name}\n",
        encoding="ascii",
    )
    return observed


def load_subject_contracts(lock_hash: str) -> tuple[list[dict], np.ndarray]:
    manifests = []
    orders = []
    for subject in SUBJECTS:
        path = SUBJECT_RESULT_ROOT / f"{subject}_production_manifest_v1.json"
        sidecar = path.with_name(path.name + ".sha256")
        fields = sidecar.read_text(encoding="ascii").strip().split()
        if len(fields) != 2 or fields[0] != sha256(path) or fields[1] != path.name:
            raise RuntimeError(f"Subject manifest sidecar mismatch: {subject}")
        payload = json.loads(path.read_text(encoding="utf-8"))
        if (
            payload.get("status") != "pass"
            or payload.get("subject") != subject
            or payload.get("implementation_lock_sha256") != lock_hash
            or any(payload["firewall"].values())
        ):
            raise RuntimeError(f"Subject production contract failed: {subject}")
        feature_path = ROOT / payload["feature_archive"]["path"]
        if sha256(feature_path) != payload["feature_archive"]["sha256"]:
            raise RuntimeError(f"Feature archive hash mismatch: {subject}")
        order_names = payload["acquisition_order"]
        if sorted(order_names) != sorted(
            f"{subject}_AAD_{condition}" for condition in CONDITIONS
        ):
            raise RuntimeError(f"Acquisition order is invalid: {subject}")
        rank = np.empty(4, dtype=np.int8)
        for acquisition_rank, name in enumerate(order_names):
            condition = name.rsplit("_", 1)[-1]
            rank[CONDITIONS.index(condition)] = acquisition_rank
        orders.append(rank)
        manifests.append(payload)
    return manifests, np.asarray(orders)


def load_array(manifests: list[dict], key: str) -> np.ndarray:
    arrays = []
    for payload in manifests:
        path = ROOT / payload["feature_archive"]["path"]
        with np.load(path, allow_pickle=False) as saved:
            arrays.append(saved[key])
    result = np.stack(arrays, axis=0)
    if not np.isfinite(result).all():
        raise RuntimeError(f"Nonfinite development feature: {key}")
    return result


def load_invariant(manifests: list[dict], key: str) -> np.ndarray:
    values = []
    for payload in manifests:
        path = ROOT / payload["feature_archive"]["path"]
        with np.load(path, allow_pickle=False) as saved:
            values.append(saved[key])
    first = values[0]
    if not all(np.array_equal(first, value) for value in values[1:]):
        raise RuntimeError(f"Subject invariant differs: {key}")
    return first


def fixed_selected_statistic(
    features: np.ndarray,
    acquisition_orders: np.ndarray,
    selected: np.ndarray,
) -> tuple[float, np.ndarray, np.ndarray]:
    first, second = participant_story_effects(features)
    q1, q2 = story_order_contrasts(acquisition_orders)
    r1 = residualize_order(first, q1)
    r2 = residualize_order(second, q2)
    score = cross_story_edge_score(r1, r2)
    return float(np.mean(score[selected])), r1, r2


def nested_geometry_signflip(
    first: np.ndarray,
    second: np.ndarray,
    order_contrasts: tuple[np.ndarray, np.ndarray],
    xy: np.ndarray,
    edges: np.ndarray,
    carrier: np.ndarray,
    observed_selected: np.ndarray,
    observed_signs: np.ndarray,
    seed_label: str,
) -> dict:
    points = fixed_integration_points(
        NESTED_INTEGRATION_POINT_COUNT,
        stable_seed(seed_label, "nested-integration"),
    )
    observed_union = family_union_fraction(
        xy,
        edges,
        observed_selected,
        observed_signs,
        carrier,
        points,
        PRIMARY_RADIUS,
    )
    patterns = all_sign_patterns(len(first))
    q1, q2 = order_contrasts
    null = np.empty(len(patterns), dtype=np.float64)
    for pattern_index, signs in enumerate(patterns):
        r1 = residualize_order(first * signs[:, None], q1 * signs)
        r2 = residualize_order(second * signs[:, None], q2 * signs)
        score = cross_story_edge_score(r1, r2)
        selected = select_edges(score, SELECTED_EDGE_COUNT)
        signed_mean = np.mean(r1[:, selected] + r2[:, selected], axis=0) / 2.0
        orientation = np.where(signed_mean >= 0.0, 1, -1)
        null[pattern_index] = family_union_fraction(
            xy,
            edges,
            selected,
            orientation,
            carrier,
            points,
            PRIMARY_RADIUS,
        )
    return {
        "observed_union": observed_union,
        "null": null,
        "exact_one_sided_p": exact_upper_p(null, observed_union),
        "pattern_count": len(patterns),
        "integration_point_count": NESTED_INTEGRATION_POINT_COUNT,
    }


def main() -> None:
    lock, lock_hash = validate_implementation_lock(Path(__file__))
    if tuple(lock["development_subjects"]) != SUBJECTS:
        raise RuntimeError("Development subjects differ from implementation lock.")
    if any(path.exists() for path in (STARTED, RESULT, NULLS, TEMPLATE)):
        raise RuntimeError("One-shot development analysis refusal.")
    manifests, acquisition_orders = load_subject_contracts(lock_hash)
    write_json_once(
        STARTED,
        {
            "schema": "KITE-ST-HD-development-analysis-start-v1",
            "implementation_lock_sha256": lock_hash,
            "subjects": list(SUBJECTS),
            "subject_manifest_sha256": {
                subject: sha256(
                    SUBJECT_RESULT_ROOT / f"{subject}_production_manifest_v1.json"
                )
                for subject in SUBJECTS
            },
            "one_shot_started": True,
        },
    )
    primary_a = load_array(manifests, "A_primary")
    primary_b = load_array(manifests, "B_primary")
    effects_a = participant_story_effects(primary_a)
    effects_b = participant_story_effects(primary_b)
    order_contrasts = story_order_contrasts(acquisition_orders)
    candidate_effects = {"A": effects_a, "B": effects_b}
    familywise = exact_candidate_familywise(
        candidate_effects,
        order_contrasts,
        SELECTED_EDGE_COUNT,
    )
    fits = observed_candidate_fit(
        candidate_effects,
        order_contrasts,
        SELECTED_EDGE_COUNT,
    )
    winner = familywise["winner"]
    d1 = (
        familywise["winner_observed"] > 0.0
        and familywise["winner_familywise_p"] <= 0.01
    )
    null_payload = {
        "candidate_A_exact_null": familywise["nulls"]["A"],
        "candidate_B_exact_null": familywise["nulls"]["B"],
        "candidate_maximum_exact_null": familywise["maximum_null"],
    }
    gates = {
        "D1_reproducibility": bool(d1),
        "D2_spatiotemporal_escape": False,
        "D3_transverse_multiscale": False,
        "D4_cross_story_function": False,
        "D5_confound_resistance": False,
        "D6_synthetic_and_integrity": False,
    }
    payload = {
        "schema": "KITE-ST-HD-development-gate-v1",
        "status": "development_failed_D1" if not d1 else "analysis_in_progress",
        "implementation_lock_sha256": lock_hash,
        "subjects": list(SUBJECTS),
        "acquisition_order_ranks_LR_by_condition": acquisition_orders,
        "order_contrasts": {
            "part1_L_minus_R": order_contrasts[0],
            "part2_L_minus_R": order_contrasts[1],
        },
        "candidate_familywise": {
            "observed": familywise["observed"],
            "familywise_p": familywise["familywise_p"],
            "winner": winner,
            "winner_observed": familywise["winner_observed"],
            "winner_familywise_p": familywise["winner_familywise_p"],
            "pattern_count": familywise["pattern_count"],
        },
        "gates": gates,
        "paper_mutated": False,
        "confirmation_acquired_or_read": False,
    }
    if d1:
        winner_primary = primary_a if winner == "A" else primary_b
        fit = fits[winner]
        selected = fit["selected_edge_indices"]
        selected_signs = fit["selected_edge_signs"]
        loo = leave_one_participant_out_cross_story_auc(
            winner_primary,
            acquisition_orders,
            SELECTED_EDGE_COUNT,
        )
        d4 = (
            loo["mean_auc"] >= 0.60
            and loo["exact_one_sided_p"] <= 0.05
            and loo["participants_above_chance"] >= 8
        )
        gates["D4_cross_story_function"] = bool(d4)
        edges = load_invariant(manifests, "primary_edges")
        xy = load_invariant(manifests, "primary_xy").astype(np.float64)
        carrier_key = (
            "A_pooled_carrier_lag_seconds"
            if winner == "A"
            else "B_pooled_carrier_lag_seconds"
        )
        carrier = np.median(load_array(manifests, carrier_key), axis=0)
        geometry = matched_geometry_distribution(
            xy,
            edges,
            carrier,
            selected,
            selected_signs,
            f"KITE-ST-HD-development-{winner}",
            family_count=MATCHED_FAMILY_COUNT,
        )
        nested = nested_geometry_signflip(
            candidate_effects[winner][0],
            candidate_effects[winner][1],
            order_contrasts,
            xy,
            edges,
            carrier,
            selected,
            selected_signs,
            f"KITE-ST-HD-development-{winner}",
        )
        for name, values in geometry["matched_distributions"].items():
            null_payload[f"geometry_matched_{name}"] = values
        null_payload["geometry_nested_signflip_union"] = nested["null"]
        comparison = geometry["comparison"]
        primary_comparison = comparison["union_0.035"]
        d2 = (
            primary_comparison["z"] >= 1.645
            and nested["exact_one_sided_p"] <= 0.05
        )
        d3 = (
            comparison["direction_minimum_eigenvalue"]["observed"]
            > comparison["direction_minimum_eigenvalue"]["matched_median"]
            and comparison["direction_effective_rank"]["observed"]
            > comparison["direction_effective_rank"]["matched_median"]
            and comparison["trilinear_determinant_mean"]["observed"]
            > comparison["trilinear_determinant_mean"]["matched_median"]
            and comparison["union_0.025"]["z"] > 0.0
            and comparison["union_0.05"]["z"] > 0.0
        )
        gates["D2_spatiotemporal_escape"] = bool(d2)
        gates["D3_transverse_multiscale"] = bool(d3)
        residual_key = f"{winner}_frontal_pc_residual"
        frontal_key = f"{winner}_frontal_only"
        reversed_key = f"{winner}_time_reversed"
        residual_features = load_array(manifests, residual_key)
        residual_stat, _, _ = fixed_selected_statistic(
            residual_features,
            acquisition_orders,
            selected,
        )
        frontal_features = load_array(manifests, frontal_key)
        frontal_effects = participant_story_effects(frontal_features)
        frontal_fit = observed_candidate_fit(
            {"frontal": frontal_effects},
            order_contrasts,
            SELECTED_EDGE_COUNT,
        )["frontal"]
        reversed_features = load_array(manifests, reversed_key)
        _, reverse_r1, reverse_r2 = fixed_selected_statistic(
            reversed_features,
            acquisition_orders,
            selected,
        )
        original_vector = np.mean(
            fit["part1_residualized"][:, selected]
            + fit["part2_residualized"][:, selected],
            axis=0,
        )
        reversed_vector = np.mean(
            reverse_r1[:, selected] + reverse_r2[:, selected],
            axis=0,
        )
        time_reversal_correlation = float(
            np.corrcoef(original_vector, reversed_vector)[0, 1]
        )
        zero_features = load_array(manifests, "zero_lag_symmetric_primary")
        zero_stat, _, _ = fixed_selected_statistic(
            zero_features,
            acquisition_orders,
            selected,
        )
        nested_points = fixed_integration_points(
            NESTED_INTEGRATION_POINT_COUNT,
            stable_seed("KITE-ST-HD-development-channel-order"),
        )
        rng = np.random.default_rng(
            stable_seed("KITE-ST-HD-development-channel-order-permutation")
        )
        permuted_xy = xy[rng.permutation(len(xy))]
        permuted_union = family_union_fraction(
            permuted_xy,
            edges,
            selected,
            selected_signs,
            carrier,
            nested_points,
            PRIMARY_RADIUS,
        )
        true_union_same_points = family_union_fraction(
            xy,
            edges,
            selected,
            selected_signs,
            carrier,
            nested_points,
            PRIMARY_RADIUS,
        )
        d5 = (
            residual_stat > 0.0
            and frontal_fit["selected_statistic"] <= fit["selected_statistic"]
            and time_reversal_correlation <= -0.80
            and zero_stat < fit["selected_statistic"]
            and abs(permuted_union - true_union_same_points) > 1e-6
        )
        gates["D5_confound_resistance"] = bool(d5)
        integrity = {}
        for path_text, expected_hash in lock["engineering_evidence_hashes"].items():
            path = ROOT / path_text
            integrity[path_text] = {
                "expected_sha256": expected_hash,
                "observed_sha256": sha256(path),
                "pass": sha256(path) == expected_hash,
            }
        d6 = (
            all(item["pass"] for item in integrity.values())
            and not confirmation_numeric_files()
        )
        gates["D6_synthetic_and_integrity"] = bool(d6)
        alpha = load_array(
            manifests,
            "posterior_alpha_right_minus_left",
        )[..., None]
        alpha_loo = leave_one_participant_out_cross_story_auc(
            alpha,
            acquisition_orders,
            selected_count=1,
        )
        full_template_auc = frozen_template_auc(
            winner_primary,
            selected,
            fit["combined_template_weight"],
        )
        local_erasure = local_erasure_frozen_template_auc(
            winner_primary,
            edges,
            selected,
            fit["combined_template_weight"],
            xy,
        )
        payload.update(
            {
                "status": (
                    "development_pass_all_gates"
                    if all(gates.values())
                    else "development_failed_conjunction"
                ),
                "winner": {
                    "candidate": winner,
                    "selected_edge_count": len(selected),
                    "selected_edge_indices": selected,
                    "selected_edge_pairs": edges[selected],
                    "selected_edge_signs": selected_signs,
                    "selected_statistic": fit["selected_statistic"],
                    "carrier_lag_seconds_selected": carrier[selected],
                },
                "geometry": {
                    "family_count": geometry["family_count"],
                    "integration_point_count": geometry["integration_point_count"],
                    "comparison": comparison,
                    "observed_occupation": geometry["observed"]["occupation"],
                    "observed_diagnostics": geometry["observed"]["diagnostics"],
                    "matching_audit": geometry["first_family_matching_audit"],
                    "nested_signflip": {
                        key: value
                        for key, value in nested.items()
                        if key != "null"
                    },
                },
                "cross_story_function": {
                    "leave_one_participant_out": {
                        key: value
                        for key, value in loo.items()
                        if key != "fold_selected_edge_indices"
                    },
                    "descriptive_full_development_frozen_template": full_template_auc,
                    "descriptive_local_erasure": local_erasure,
                    "ordinary_posterior_alpha_benchmark": {
                        key: value
                        for key, value in alpha_loo.items()
                        if key != "fold_selected_edge_indices"
                    },
                },
                "confound_controls": {
                    "frontal_pc_residual_fixed_selected_statistic": residual_stat,
                    "frontal_only_selected_statistic": frontal_fit["selected_statistic"],
                    "time_reversal_selected_vector_correlation": time_reversal_correlation,
                    "zero_lag_symmetric_fixed_selected_statistic": zero_stat,
                    "true_geometry_union_channel_order_audit": true_union_same_points,
                    "permuted_geometry_union_channel_order_audit": permuted_union,
                },
                "integrity": integrity,
            }
        )
        if all(gates.values()):
            np.savez_compressed(
                TEMPLATE,
                candidate=np.asarray(winner),
                selected_edge_indices=selected.astype(np.int32),
                selected_edge_pairs=edges[selected].astype(np.int32),
                selected_edge_signs=selected_signs.astype(np.int8),
                combined_template_weight=fit["combined_template_weight"].astype(
                    np.float64
                ),
                carrier_lag_seconds=carrier[selected].astype(np.float32),
                all_edge_carrier_lag_seconds=carrier.astype(np.float32),
                primary_xy=xy.astype(np.float32),
                primary_edges=edges.astype(np.int32),
                development_subjects=np.asarray(SUBJECTS),
                implementation_lock_sha256=np.asarray(lock_hash),
            )
            template_hash = sha256(TEMPLATE)
            TEMPLATE.with_name(TEMPLATE.name + ".sha256").write_text(
                f"{template_hash}  {TEMPLATE.name}\n",
                encoding="ascii",
            )
            payload["development_template"] = {
                "path": TEMPLATE.relative_to(ROOT).as_posix(),
                "bytes": TEMPLATE.stat().st_size,
                "sha256": template_hash,
            }
    np.savez_compressed(NULLS, **null_payload)
    null_hash = sha256(NULLS)
    NULLS.with_name(NULLS.name + ".sha256").write_text(
        f"{null_hash}  {NULLS.name}\n",
        encoding="ascii",
    )
    payload["null_distributions"] = {
        "path": NULLS.relative_to(ROOT).as_posix(),
        "bytes": NULLS.stat().st_size,
        "sha256": null_hash,
        "arrays": {name: list(np.asarray(value).shape) for name, value in null_payload.items()},
    }
    payload["confirmation_authorized"] = bool(all(gates.values()))
    result_hash = write_json_once(RESULT, payload)
    print(
        json.dumps(
            {
                "status": payload["status"],
                "gates": gates,
                "result": str(RESULT),
                "sha256": result_hash,
            }
        )
    )


if __name__ == "__main__":
    main()
```
