## Supplementary material for "Falsifiable substitution tests reveal task-structured neural evidence for auditory attention": Article_Analysis_Source_Code: run_kite_st_hd_development_analysis_v2.py

```
"""V2 cohort wrapper around the byte-frozen KITE-ST-HD analysis engine."""

from __future__ import annotations

from pathlib import Path

import run_kite_st_hd_development_analysis_v1 as engine
from kite_st_hd_lock_utils_v2 import (
    ROOT,
    confirmation_numeric_files,
    validate_implementation_lock,
)


SUBJECTS = (
    "S4", "S24", "S11", "S23", "S14", "S29",
    "S19", "S17", "S12", "S25", "S27", "S5",
)
RESULT_ROOT = ROOT / "results" / "kite_st_hd" / "development_v2"


def main() -> None:
    engine.SUBJECTS = SUBJECTS
    engine.RESULT_ROOT = RESULT_ROOT
    engine.SUBJECT_RESULT_ROOT = RESULT_ROOT / "subjects"
    engine.STARTED = RESULT_ROOT / "development_analysis_started_v2.json"
    engine.RESULT = RESULT_ROOT / "development_gate_v2.json"
    engine.NULLS = RESULT_ROOT / "development_null_distributions_v2.npz"
    engine.TEMPLATE = RESULT_ROOT / "development_template_v2.npz"
    engine.validate_implementation_lock = validate_implementation_lock
    engine.confirmation_numeric_files = confirmation_numeric_files
    engine.main()


if __name__ == "__main__":
    main()
```
