## Supplementary material for "Falsifiable substitution tests reveal task-structured neural evidence for auditory attention": Article_Analysis_Source_Code: run_kite_st_hd_development_subject_v2.py

```
"""One-shot KITE-ST-HD-v2 production for one development subject."""

from __future__ import annotations

import argparse
import json
import shutil
import tarfile
from pathlib import Path, PurePosixPath

from download_kite_st_hd_development_v2 import FILES, SUBJECTS
from kite_st_hd_lock_utils_v2 import ROOT, sha256, validate_implementation_lock
from kite_st_hd_subject_io_v1 import build_feature_archive, write_subject_manifest
from kite_st_hd_subject_io_v2 import preprocess_subject_v2


ARCHIVE_ROOT = (
    ROOT / "data" / "raw" / "kite_st_hd_zenodo4518754" / "development_archives"
)
WORK_ROOT = (
    ROOT / "data" / "raw" / "kite_st_hd_zenodo4518754" / "development_work_v2"
)
PREPROCESSED_ROOT = (
    ROOT
    / "data"
    / "derived"
    / "kite_st_hd_zenodo4518754"
    / "development_preprocessed_v2"
)
FEATURE_ROOT = (
    ROOT
    / "data"
    / "derived"
    / "kite_st_hd_zenodo4518754"
    / "development_features_v2"
)
RESULT_ROOT = ROOT / "results" / "kite_st_hd" / "development_v2" / "subjects"
METADATA_ROOT = ROOT / "data" / "metadata" / "kite_st_hd_development_downloads_v2"
CONDITIONS = ("1L", "1R", "2L", "2R")
EXTENSIONS = (".ceo", ".dap", ".dat", ".rs3")


def expected_member_names(subject: str) -> set[str]:
    names = {subject}
    names.update(
        f"{subject}/{subject}_AAD_{condition}{extension}"
        for condition in CONDITIONS
        for extension in EXTENSIONS
    )
    return names


def validate_and_extract(subject: str, archive: Path) -> dict:
    work_subject = WORK_ROOT / subject
    if work_subject.exists():
        raise RuntimeError(f"Refusing existing v2 work directory: {work_subject}")
    WORK_ROOT.mkdir(parents=True, exist_ok=True)
    with tarfile.open(archive, mode="r:gz") as stream:
        members = stream.getmembers()
        if {member.name for member in members} != expected_member_names(subject):
            raise RuntimeError(f"V2 archive members differ for {subject}.")
        for member in members:
            parsed = PurePosixPath(member.name)
            if parsed.is_absolute() or ".." in parsed.parts:
                raise RuntimeError(f"Unsafe v2 archive path: {member.name}")
            if member.issym() or member.islnk() or member.isdev():
                raise RuntimeError(f"Unsupported v2 archive member: {member.name}")
            if parsed.parts[0] != subject:
                raise RuntimeError(f"V2 archive member escapes root: {member.name}")
        stream.extractall(WORK_ROOT, members=members, filter="data")
    files = sorted(
        path.relative_to(WORK_ROOT).as_posix()
        for path in work_subject.rglob("*")
        if path.is_file()
    )
    if len(files) != 16:
        raise RuntimeError("Extracted v2 subject does not have 16 files.")
    return {"member_count": len(members), "file_count": len(files), "files": files}


def remove_work_subject(subject: str) -> None:
    target = (WORK_ROOT / subject).resolve()
    if target.parent != WORK_ROOT.resolve() or target.name != subject:
        raise RuntimeError("Refusing unsafe v2 cleanup target.")
    if not target.is_dir():
        raise RuntimeError("V2 cleanup target is absent.")
    shutil.rmtree(target)
    if target.exists():
        raise RuntimeError("V2 cleanup did not complete.")


def main() -> None:
    parser = argparse.ArgumentParser()
    parser.add_argument("subject", choices=SUBJECTS)
    args = parser.parse_args()
    subject = args.subject
    lock, lock_hash = validate_implementation_lock(Path(__file__))
    if tuple(lock["development_subjects"]) != SUBJECTS:
        raise RuntimeError("V2 subject list differs from lock.")
    archive = ARCHIVE_ROOT / f"{subject}.tar.gz"
    expected_bytes, expected_md5 = FILES[subject]
    acquisition_path = METADATA_ROOT / f"{subject}_archive_acquisition_v2.json"
    acquisition = json.loads(acquisition_path.read_text(encoding="utf-8"))
    if (
        archive.stat().st_size != expected_bytes
        or acquisition["bytes"] != expected_bytes
        or acquisition["md5"] != expected_md5
        or acquisition["sha256"] != sha256(archive)
        or acquisition["v2_implementation_lock_sha256"] != lock_hash
    ):
        raise RuntimeError("V2 archive/acquisition contract mismatch.")
    started = RESULT_ROOT / f"{subject}_production_started_v2.json"
    # The byte-frozen analysis engine reads this compatibility filename inside
    # the dedicated development_v2 root; the payload schema remains v2.
    final_manifest = RESULT_ROOT / f"{subject}_production_manifest_v1.json"
    output_dir = PREPROCESSED_ROOT / subject
    feature_path = FEATURE_ROOT / f"{subject}_features_v2.npz"
    if any(
        path.exists()
        for path in (
            started,
            final_manifest,
            output_dir,
            feature_path,
            WORK_ROOT / subject,
        )
    ):
        raise RuntimeError(f"V2 one-shot refusal for existing output: {subject}")
    write_subject_manifest(
        started,
        {
            "schema": "KITE-ST-HD-v2-development-production-start-v2",
            "subject": subject,
            "v2_implementation_lock_sha256": lock_hash,
            "archive_path": archive.relative_to(ROOT).as_posix(),
            "archive_sha256": acquisition["sha256"],
            "one_shot_started": True,
        },
    )
    extraction = validate_and_extract(subject, archive)
    preprocessing_records = preprocess_subject_v2(subject, WORK_ROOT, output_dir)
    feature_record = build_feature_archive(
        subject,
        preprocessing_records,
        feature_path,
    )
    remove_work_subject(subject)
    acquisition_order = [
        item["recording"]
        for item in sorted(preprocessing_records, key=lambda item: item["start_iso"])
    ]
    maximum_bad = max(item["bad_channels"]["count"] for item in preprocessing_records)
    payload = {
        "schema": "KITE-ST-HD-v2-development-subject-production-v2",
        "status": "pass",
        "subject": subject,
        "implementation_lock_sha256": lock_hash,
        "archive": {
            "path": archive.relative_to(ROOT).as_posix(),
            "bytes": expected_bytes,
            "md5": expected_md5,
            "sha256": acquisition["sha256"],
        },
        "extraction": extraction,
        "preprocessing_records": preprocessing_records,
        "maximum_bad_channel_count": maximum_bad,
        "maximum_bad_channel_rule": 15,
        "feature_archive": feature_record,
        "acquisition_order": acquisition_order,
        "temporary_extracted_work_removed_after_verified_derivatives": True,
        "firewall": {
            "condition_contrast_computed": False,
            "true_label_decoder_computed": False,
            "edge_selection_computed": False,
            "empirical_geometry_endpoint_computed": False,
            "confirmation_file_read": False,
            "paper_mutated": False,
        },
    }
    observed = write_subject_manifest(final_manifest, payload)
    print(
        json.dumps(
            {
                "status": "pass",
                "subject": subject,
                "manifest": str(final_manifest),
                "sha256": observed,
            }
        )
    )


if __name__ == "__main__":
    main()
```
