## Supplementary material for "Falsifiable substitution tests reveal task-structured neural evidence for auditory attention": Article_Analysis_Source_Code: run_wz_hic_asme30_crossphase_individual_geometry_v1.py

```
#!/usr/bin/env python3
"""Frozen ASME30 target-free cross-phase individualized geometry analysis."""

from __future__ import annotations

import csv
import hashlib
import itertools
import json
import math
from datetime import datetime, timezone
from pathlib import Path

import numpy as np

from wz_hic_asme30_core_v1 import (
    PHYSICAL_GROUPS,
    WRONG_GROUPS,
    labels_for_mapping,
)
from wz_hif_core_v1 import fit_shrinkage_lda


ROOT = Path(__file__).resolve().parents[1]
PROTOCOL = (
    ROOT
    / "docs"
    / "wz_hic_asme30_crossphase_individual_geometry_protocol_v1.json"
)
LOCK = (
    ROOT
    / "docs"
    / "wz_hic_asme30_crossphase_individual_geometry_implementation_lock_v1.json"
)
FEATURES = (
    ROOT
    / "data"
    / "derived"
    / "asme30_wz_hic_dataset_native_features_v2.npz"
)
OUTDIR = (
    ROOT
    / "results"
    / "wz_hic_asme30"
    / "crossphase_individual_geometry_v1"
)
RESULT = OUTDIR / "crossphase_decision_v1.json"
PARTICIPANT_CSV = OUTDIR / "participant_geometry_v1.csv"
TIME_CSV = OUTDIR / "time_resolved_v1.csv"

PROTOCOL_SHA256 = (
    "bf38a35a1fd6ba09accb702efde08eed9ba1596a0849d1c25bdd9bd835bed04d"
)
FEATURES_SHA256 = (
    "e767e08dce8e46d1589203e76289a5acf64284166efc54325b945108e9559017"
)
HORIZON = 3
BOOTSTRAP_N = 20_000
BOOTSTRAP_SEED = 20260730


def sha256(path: Path) -> str:
    digest = hashlib.sha256()
    with path.open("rb") as handle:
        for block in iter(lambda: handle.read(1024 * 1024), b""):
            digest.update(block)
    return digest.hexdigest()


def validate_frozen_inputs() -> None:
    observed = {
        "protocol": sha256(PROTOCOL),
        "features": sha256(FEATURES),
    }
    expected = {
        "protocol": PROTOCOL_SHA256,
        "features": FEATURES_SHA256,
    }
    if observed != expected:
        raise RuntimeError(f"Frozen input mismatch: {observed} != {expected}")
    if not LOCK.exists():
        raise FileNotFoundError("Implementation lock must exist before outcomes.")
    lock = json.loads(LOCK.read_text(encoding="utf-8"))
    if lock.get("synthetic_test_status") != "PASS":
        raise RuntimeError("Synthetic test was not locked as PASS.")
    if lock.get("scientific_outcomes_available_at_lock") is not False:
        raise RuntimeError("Implementation lock is not pre-outcome.")


def trial_horizon_indices(
    command: np.ndarray, trial: np.ndarray
) -> tuple[list[np.ndarray], list[int]]:
    selections = []
    incomplete = []
    for trial_id in sorted(int(value) for value in np.unique(trial)):
        ids = np.flatnonzero(trial == trial_id)
        chunks = []
        for value in range(1, 31):
            command_ids = ids[command[ids] == value]
            if command_ids.size < HORIZON:
                chunks = []
                break
            chunks.append(command_ids[:HORIZON])
        if len(chunks) == 30:
            selections.append(np.concatenate(chunks))
        else:
            incomplete.append(trial_id)
    return selections, incomplete


def fit_standardized_scores(
    train_x: np.ndarray,
    train_labels: np.ndarray,
    test_x: np.ndarray,
) -> tuple[np.ndarray, dict]:
    keep = np.isin(train_labels, ("S", "U"))
    y = (train_labels[keep] == "S").astype(int)
    model = fit_shrinkage_lda(train_x[keep], y)
    train_scores = model.decision_function(train_x[keep])
    mean = float(np.mean(train_scores))
    sd = float(np.std(train_scores, ddof=1))
    if not np.isfinite(sd) or sd <= 1e-12:
        raise RuntimeError("Degenerate training-score scale.")
    test_scores = (model.decision_function(test_x) - mean) / sd
    return test_scores, {
        "n_train_S": int(np.sum(train_labels == "S")),
        "n_train_U": int(np.sum(train_labels == "U")),
        "n_train_T_excluded": int(np.sum(train_labels == "T")),
        "training_score_mean": mean,
        "training_score_sd": sd,
    }


def trial_balanced_score_effect(
    scores: np.ndarray,
    command: np.ndarray,
    target: np.ndarray,
    selections: list[np.ndarray],
    groups,
) -> tuple[float, list[float]]:
    trial_values = []
    for selected in selections:
        target_values = np.unique(target[selected])
        if target_values.size != 1:
            raise RuntimeError("Nonunique target in online trial.")
        labels = labels_for_mapping(
            command[selected], target[selected], groups
        )
        counts = (
            int(np.sum(labels == "T")),
            int(np.sum(labels == "S")),
            int(np.sum(labels == "U")),
        )
        if counts != (3, 27, 60):
            raise RuntimeError(f"Unexpected T/S/U counts: {counts}")
        trial_values.append(
            float(np.mean(scores[selected][labels == "S"]))
            - float(np.mean(scores[selected][labels == "U"]))
        )
    return float(np.mean(trial_values)), trial_values


def safe_cosine(left: np.ndarray, right: np.ndarray) -> float:
    denominator = float(np.linalg.norm(left) * np.linalg.norm(right))
    if not np.isfinite(denominator) or denominator <= 1e-12:
        raise RuntimeError("Degenerate neural direction.")
    return float(np.vdot(left, right) / denominator)


def crossphase_directions(
    train_x: np.ndarray,
    train_labels: np.ndarray,
    test_x: np.ndarray,
    test_command: np.ndarray,
    test_target: np.ndarray,
    selections: list[np.ndarray],
    groups,
) -> tuple[np.ndarray, np.ndarray, float]:
    keep = np.isin(train_labels, ("S", "U"))
    train_outer = np.asarray(train_x[keep], dtype=np.float64)
    labels_outer = train_labels[keep]
    center = np.mean(train_outer, axis=0)
    scale = np.std(train_outer, axis=0, ddof=1)
    scale[~np.isfinite(scale) | (scale <= 1e-12)] = 1.0
    train_z = (train_outer - center) / scale
    offline = np.mean(train_z[labels_outer == "S"], axis=0) - np.mean(
        train_z[labels_outer == "U"], axis=0
    )
    test_z = (np.asarray(test_x, dtype=np.float64) - center) / scale
    online_trial_directions = []
    for selected in selections:
        labels = labels_for_mapping(
            test_command[selected], test_target[selected], groups
        )
        if (
            int(np.sum(labels == "T")),
            int(np.sum(labels == "S")),
            int(np.sum(labels == "U")),
        ) != (3, 27, 60):
            raise RuntimeError("Direction computation violated target removal.")
        online_trial_directions.append(
            np.mean(test_z[selected][labels == "S"], axis=0)
            - np.mean(test_z[selected][labels == "U"], axis=0)
        )
    online = np.mean(np.stack(online_trial_directions), axis=0)
    return offline, online, safe_cosine(offline, online)


def analyze_mapping(
    *,
    train_full: np.ndarray,
    test_full: np.ndarray,
    train_bins: np.ndarray,
    test_bins: np.ndarray,
    train_command: np.ndarray,
    train_target: np.ndarray,
    test_command: np.ndarray,
    test_target: np.ndarray,
    selections: list[np.ndarray],
    groups,
) -> dict:
    train_labels = labels_for_mapping(
        train_command, train_target, groups
    )
    full_scores, full_fit = fit_standardized_scores(
        train_full, train_labels, test_full
    )
    full_effect, trial_values = trial_balanced_score_effect(
        full_scores,
        test_command,
        test_target,
        selections,
        groups,
    )
    offline_direction, online_direction, alignment = crossphase_directions(
        train_full,
        train_labels,
        test_full,
        test_command,
        test_target,
        selections,
        groups,
    )
    time_effects = []
    time_fits = []
    for bin_index in range(10):
        scores, fit = fit_standardized_scores(
            train_bins[:, :, bin_index],
            train_labels,
            test_bins[:, :, bin_index],
        )
        effect, _ = trial_balanced_score_effect(
            scores,
            test_command,
            test_target,
            selections,
            groups,
        )
        time_effects.append(effect)
        time_fits.append(fit)
    return {
        "full_effect": full_effect,
        "trial_effects": trial_values,
        "time_effects": time_effects,
        "alignment": alignment,
        "offline_direction": offline_direction,
        "online_direction": online_direction,
        "full_fit": full_fit,
        "time_fits": time_fits,
    }


def analyze_subject(arrays: dict, subject: int) -> dict:
    train_mask = (
        (arrays["subject"] == subject)
        & (arrays["task"] == "offline")
        & arrays["accepted"].astype(bool)
    )
    test_mask = (
        (arrays["subject"] == subject)
        & (arrays["task"] == "online")
        & arrays["accepted"].astype(bool)
    )
    train_bins = arrays["eeg_bins"][train_mask, :, 1:]
    test_bins = arrays["eeg_bins"][test_mask, :, 1:]
    train_full = train_bins.reshape(train_bins.shape[0], -1)
    test_full = test_bins.reshape(test_bins.shape[0], -1)
    train_eog = arrays["eog_bins"][train_mask, :, 1:].reshape(
        int(np.sum(train_mask)), -1
    )
    test_eog = arrays["eog_bins"][test_mask, :, 1:].reshape(
        int(np.sum(test_mask)), -1
    )
    train_command = arrays["command"][train_mask].astype(int)
    train_target = arrays["target"][train_mask].astype(int)
    test_command = arrays["command"][test_mask].astype(int)
    test_target = arrays["target"][test_mask].astype(int)
    test_trial = arrays["trial"][test_mask].astype(int)
    train_selections, train_incomplete = trial_horizon_indices(
        train_command, arrays["trial"][train_mask].astype(int)
    )
    test_selections, test_incomplete = trial_horizon_indices(
        test_command, test_trial
    )
    if train_incomplete or test_incomplete:
        raise RuntimeError(
            f"Subject {subject} has incomplete dataset-native trials."
        )
    if len(train_selections) != 30 or len(test_selections) != 15:
        raise RuntimeError(
            f"Subject {subject} coverage is "
            f"{len(train_selections)}/{len(test_selections)}, expected 30/15."
        )

    physical = analyze_mapping(
        train_full=train_full,
        test_full=test_full,
        train_bins=train_bins,
        test_bins=test_bins,
        train_command=train_command,
        train_target=train_target,
        test_command=test_command,
        test_target=test_target,
        selections=test_selections,
        groups=PHYSICAL_GROUPS,
    )
    wrong = [
        analyze_mapping(
            train_full=train_full,
            test_full=test_full,
            train_bins=train_bins,
            test_bins=test_bins,
            train_command=train_command,
            train_target=train_target,
            test_command=test_command,
            test_target=test_target,
            selections=test_selections,
            groups=groups,
        )
        for groups in WRONG_GROUPS
    ]
    train_labels_physical = labels_for_mapping(
        train_command, train_target, PHYSICAL_GROUPS
    )
    eog_scores, eog_fit = fit_standardized_scores(
        train_eog, train_labels_physical, test_eog
    )
    eog_effect, eog_trial_values = trial_balanced_score_effect(
        eog_scores,
        test_command,
        test_target,
        test_selections,
        PHYSICAL_GROUPS,
    )
    return {
        "subject": subject,
        "n_offline_trials": len(train_selections),
        "n_online_trials": len(test_selections),
        "n_offline_events": int(np.sum(train_mask)),
        "n_online_events": int(np.sum(test_mask)),
        "physical": physical,
        "wrong": wrong,
        "eog_effect": eog_effect,
        "eog_trial_effects": eog_trial_values,
        "eog_fit": eog_fit,
    }


def bootstrap_ci(values: np.ndarray) -> list[float]:
    rng = np.random.default_rng(BOOTSTRAP_SEED)
    indices = rng.integers(
        0, values.size, size=(BOOTSTRAP_N, values.size)
    )
    return [
        float(value)
        for value in np.quantile(values[indices].mean(axis=1), [0.025, 0.975])
    ]


def exact_signflip_p(values: np.ndarray) -> float:
    observed = float(np.mean(values))
    permutations = [
        float(np.mean(values * np.asarray(signs)))
        for signs in itertools.product((-1.0, 1.0), repeat=values.size)
    ]
    return float(np.mean(np.asarray(permutations) >= observed - 1e-15))


def summarize(values: np.ndarray) -> dict:
    values = np.asarray(values, dtype=float)
    return {
        "mean": float(np.mean(values)),
        "bootstrap_95_CI_mean": bootstrap_ci(values),
        "n_positive": int(np.sum(values > 0)),
        "n_total": int(values.size),
        "one_sided_exact_signflip_p": exact_signflip_p(values),
        "participant_values": [float(value) for value in values],
    }


def t_stat(values: np.ndarray) -> np.ndarray:
    denominator = np.std(values, axis=0, ddof=1) / math.sqrt(values.shape[0])
    return np.divide(
        np.mean(values, axis=0),
        denominator,
        out=np.zeros(values.shape[1], dtype=float),
        where=denominator > 1e-12,
    )


def exact_time_max_t(
    values: np.ndarray,
) -> tuple[np.ndarray, np.ndarray, np.ndarray]:
    observed = t_stat(values)
    maxima = []
    for signs in itertools.product((-1.0, 1.0), repeat=values.shape[0]):
        signed = values * np.asarray(signs)[:, None]
        maxima.append(float(np.max(np.abs(t_stat(signed)))))
    maxima = np.asarray(maxima)
    p_fwer = np.mean(
        maxima[:, None] >= np.abs(observed)[None, :] - 1e-15, axis=0
    )
    return observed, p_fwer, maxima


def run_analysis(arrays: dict) -> dict:
    records = [
        analyze_subject(arrays, subject)
        for subject in sorted(int(x) for x in np.unique(arrays["subject"]))
    ]
    offline_physical = [
        item["physical"]["offline_direction"] for item in records
    ]
    rows = []
    time_specificity = []
    for index, item in enumerate(records):
        physical = item["physical"]
        wrong_1, wrong_2 = item["wrong"]
        between = float(
            np.mean(
                [
                    safe_cosine(
                        physical["online_direction"],
                        offline_physical[other],
                    )
                    for other in range(len(records))
                    if other != index
                ]
            )
        )
        best_wrong_effect = max(
            wrong_1["full_effect"], wrong_2["full_effect"]
        )
        best_wrong_alignment = max(
            wrong_1["alignment"], wrong_2["alignment"]
        )
        row = {
            "subject": item["subject"],
            "n_offline_trials": item["n_offline_trials"],
            "n_online_trials": item["n_online_trials"],
            "physical_full_effect": physical["full_effect"],
            "wrong_1_full_effect": wrong_1["full_effect"],
            "wrong_2_full_effect": wrong_2["full_effect"],
            "best_wrong_full_effect": best_wrong_effect,
            "binding_specificity": physical["full_effect"]
            - best_wrong_effect,
            "physical_eog_effect": item["eog_effect"],
            "nonocular_specificity": physical["full_effect"]
            - item["eog_effect"],
            "physical_within_alignment": physical["alignment"],
            "wrong_1_within_alignment": wrong_1["alignment"],
            "wrong_2_within_alignment": wrong_2["alignment"],
            "best_wrong_within_alignment": best_wrong_alignment,
            "mapping_alignment_specificity": physical["alignment"]
            - best_wrong_alignment,
            "between_person_substitution": between,
            "identity_advantage": physical["alignment"] - between,
        }
        rows.append(row)
        time_specificity.append(
            np.asarray(physical["time_effects"])
            - np.maximum(
                np.asarray(wrong_1["time_effects"]),
                np.asarray(wrong_2["time_effects"]),
            )
        )
    time_specificity = np.stack(time_specificity)
    time_t, time_p, time_null = exact_time_max_t(time_specificity)
    time_rows = [
        {
            "bin_index": bin_index,
            "time_start_s": bin_index / 10,
            "time_stop_s": (bin_index + 1) / 10,
            "mean_physical_minus_best_wrong": float(
                np.mean(time_specificity[:, bin_index])
            ),
            "t_value": float(time_t[bin_index]),
            "max_t_fwer_p": float(time_p[bin_index]),
            "fwer_significant_0_05": bool(time_p[bin_index] <= 0.05),
            "participant_values": [
                float(value) for value in time_specificity[:, bin_index]
            ],
        }
        for bin_index in range(10)
    ]

    summaries = {
        "physical_crossphase": summarize(
            np.asarray([row["physical_full_effect"] for row in rows])
        ),
        "binding_specificity": summarize(
            np.asarray([row["binding_specificity"] for row in rows])
        ),
        "nonocular_specificity": summarize(
            np.asarray([row["nonocular_specificity"] for row in rows])
        ),
        "identity_advantage": summarize(
            np.asarray([row["identity_advantage"] for row in rows])
        ),
        "mapping_alignment_specificity": summarize(
            np.asarray(
                [row["mapping_alignment_specificity"] for row in rows]
            )
        ),
        "physical_within_alignment": summarize(
            np.asarray(
                [row["physical_within_alignment"] for row in rows]
            )
        ),
        "between_person_substitution": summarize(
            np.asarray(
                [row["between_person_substitution"] for row in rows]
            )
        ),
    }
    n = len(rows)

    def scalar_gate(name: str, required: int) -> bool:
        item = summaries[name]
        return (
            item["mean"] > 0
            and item["bootstrap_95_CI_mean"][0] > 0
            and item["n_positive"] >= required
            and item["one_sided_exact_signflip_p"] <= 0.05
        )

    gates = {
        "G0_coverage": n == 10
        and all(
            row["n_offline_trials"] == 30
            and row["n_online_trials"] == 15
            for row in rows
        ),
        "G1_physical_crossphase": scalar_gate(
            "physical_crossphase", math.ceil(0.9 * n)
        ),
        "G2_binding_specificity": scalar_gate(
            "binding_specificity", math.ceil(0.8 * n)
        ),
        "G3_nonocular_specificity": scalar_gate(
            "nonocular_specificity", math.ceil(0.8 * n)
        ),
        "G4_identity_advantage": scalar_gate(
            "identity_advantage", math.ceil(0.8 * n)
        ),
        "G5_mapping_alignment_specificity": scalar_gate(
            "mapping_alignment_specificity", math.ceil(0.8 * n)
        ),
        "G6_time_localization": any(
            row["mean_physical_minus_best_wrong"] > 0
            and row["max_t_fwer_p"] <= 0.05
            for row in time_rows
        ),
    }
    return {
        "records": records,
        "participant_rows": rows,
        "time_rows": time_rows,
        "summaries": summaries,
        "time_max_t_null": time_null,
        "gates": gates,
        "supported": all(gates.values()),
    }


def main() -> None:
    validate_frozen_inputs()
    if RESULT.exists() or PARTICIPANT_CSV.exists() or TIME_CSV.exists():
        raise FileExistsError("Immutable v1 outcome already exists.")
    with np.load(FEATURES, allow_pickle=True) as source:
        arrays = {key: source[key] for key in source.files}
    analysis = run_analysis(arrays)
    output = {
        "route": "WZ-HIC-ASME30-crossphase-individual-geometry-v1",
        "created_utc": datetime.now(timezone.utc).isoformat(),
        "status": (
            "crossphase_individual_geometry_supported"
            if analysis["supported"]
            else "crossphase_individual_geometry_not_supported_no_retuning"
        ),
        "individual_geometry_supported": analysis["supported"],
        "gates": analysis["gates"],
        "summaries": analysis["summaries"],
        "participant_geometry": analysis["participant_rows"],
        "time_resolved_binding_specificity": analysis["time_rows"],
        "time_max_t": {
            "exact_sign_patterns": int(
                2 ** len(analysis["participant_rows"])
            ),
            "null_95_percentile": float(
                np.quantile(analysis["time_max_t_null"], 0.95)
            ),
        },
        "fit_audit": {
            str(item["subject"]): {
                "physical_full": item["physical"]["full_fit"],
                "wrong_1_full": item["wrong"][0]["full_fit"],
                "wrong_2_full": item["wrong"][1]["full_fit"],
                "eog_full": item["eog_fit"],
            }
            for item in analysis["records"]
        },
        "provenance": {
            "protocol_sha256": sha256(PROTOCOL),
            "implementation_lock_sha256": sha256(LOCK),
            "features_sha256": sha256(FEATURES),
            "runner_sha256": sha256(Path(__file__)),
        },
        "interpretation_boundary": (
            "Post-primary target-free mechanistic development result. "
            "Support would demonstrate participant-specific offline-to-"
            "online transport and physical-binding specificity in sensor "
            "space; it would not be a cortical source, an independent "
            "confirmation, or a direct use of the Kakeya theorem."
        ),
    }
    OUTDIR.mkdir(parents=True, exist_ok=True)
    RESULT.write_text(
        json.dumps(output, ensure_ascii=False, indent=2) + "\n",
        encoding="utf-8",
    )
    with PARTICIPANT_CSV.open("x", newline="", encoding="utf-8-sig") as handle:
        writer = csv.DictWriter(
            handle, fieldnames=list(analysis["participant_rows"][0])
        )
        writer.writeheader()
        writer.writerows(analysis["participant_rows"])
    with TIME_CSV.open("x", newline="", encoding="utf-8-sig") as handle:
        fieldnames = [
            key
            for key in analysis["time_rows"][0]
            if key != "participant_values"
        ]
        writer = csv.DictWriter(handle, fieldnames=fieldnames)
        writer.writeheader()
        writer.writerows(
            [
                {key: row[key] for key in fieldnames}
                for row in analysis["time_rows"]
            ]
        )
    print(
        json.dumps(
            {
                "result": RESULT.relative_to(ROOT).as_posix(),
                "result_sha256": sha256(RESULT),
                "participant_csv_sha256": sha256(PARTICIPANT_CSV),
                "time_csv_sha256": sha256(TIME_CSV),
                "status": output["status"],
                "gates": output["gates"],
                "summaries": output["summaries"],
                "time": output["time_resolved_binding_specificity"],
            },
            ensure_ascii=False,
            indent=2,
        )
    )


if __name__ == "__main__":
    main()
```
