## Supplementary material for "Falsifiable substitution tests reveal task-structured neural evidence for auditory attention": Article_Analysis_Source_Code: run_wz_hic_asme30_dataset_native_v2.py

```
#!/usr/bin/env python3
"""Run the frozen, dataset-native WZ-HIC ASME30 robustness analysis."""

from __future__ import annotations

import hashlib
import json
from datetime import datetime, timezone
from pathlib import Path

import numpy as np

from run_wz_hic_asme30_external_v1 import (
    absolute_gate,
    effect,
    mapping_key,
    paired_gate,
    select_bins,
)
from wz_hic_asme30_core_v1 import (
    PHYSICAL_GROUPS,
    WRONG_GROUPS,
    certificate_metrics,
    fit_primary_event_scores,
    fit_wrong_mapping_score,
    trial_rows,
)


ROOT = Path(__file__).resolve().parents[1]
FEATURES = (
    ROOT / "data" / "derived" / "asme30_wz_hic_dataset_native_features_v2.npz"
)
FEATURE_AUDIT = (
    ROOT
    / "data"
    / "inventory"
    / "asme30_wz_hic_dataset_native_feature_audit_v2.json"
)
PROTOCOL = ROOT / "docs" / "wz_hic_asme30_dataset_native_protocol_v2.json"
LOCK = ROOT / "docs" / "wz_hic_asme30_dataset_native_implementation_lock_v2a.json"
RESULT = (
    ROOT
    / "results"
    / "wz_hic_asme30"
    / "dataset_native_v2"
    / "dataset_native_decision_v2.json"
)


def sha256(path: Path) -> str:
    h = hashlib.sha256()
    with path.open("rb") as f:
        for block in iter(lambda: f.read(1024 * 1024), b""):
            h.update(block)
    return h.hexdigest()


def validate_lock() -> dict:
    lock = json.loads(LOCK.read_text(encoding="utf-8"))
    files = {
        "protocol": PROTOCOL,
        "primary_external_runner_dependency": (
            ROOT / "analysis" / "run_wz_hic_asme30_external_v1.py"
        ),
        "primary_feature_builder_dependency": (
            ROOT / "analysis" / "build_asme30_wz_hic_features_v1.py"
        ),
        "primary_core": ROOT / "analysis" / "wz_hic_asme30_core_v1.py",
        "lda_dependency": ROOT / "analysis" / "wz_hif_core_v1.py",
        "feature_builder": (
            ROOT / "analysis" / "build_asme30_wz_hic_dataset_native_features_v2.py"
        ),
        "runner": Path(__file__),
        "synthetic_test": ROOT / "analysis" / "test_wz_hic_asme30_synthetic_v1.py",
    }
    observed = {name: sha256(path) for name, path in files.items()}
    if observed != lock["locked_sha256"]:
        raise RuntimeError(f"Dataset-native lock mismatch: {observed}")
    return lock


def participant_analysis(z, subject: int) -> dict:
    accepted = z["accepted"]
    train = (z["subject"] == subject) & (z["task"] == "offline") & accepted
    test = (z["subject"] == subject) & (z["task"] == "online") & accepted
    train_eeg_outer = select_bins(z["eeg_bins"][train], (0.0, 1.0))
    test_eeg_outer = select_bins(z["eeg_bins"][test], (0.0, 1.0))
    train_eog_outer = select_bins(z["eog_bins"][train], (0.0, 1.0))
    test_eog_outer = select_bins(z["eog_bins"][test], (0.0, 1.0))
    train_flat = select_bins(z["eeg_bins"][train], (0.3, 0.8))
    test_flat = select_bins(z["eeg_bins"][test], (0.3, 0.8))
    scores, fit_audit = fit_primary_event_scores(
        train_eeg_outer=train_eeg_outer,
        train_eog_outer=train_eog_outer,
        train_prestim=z["prestim_raw"][train],
        train_flat=train_flat,
        train_command=z["command"][train],
        train_target=z["target"][train],
        test_eeg_outer=test_eeg_outer,
        test_eog_outer=test_eog_outer,
        test_prestim=z["prestim_raw"][test],
        test_flat=test_flat,
    )
    common = {
        "command": z["command"][test],
        "trial": z["trial"][test],
        "target": z["target"][test],
        "horizon": 3,
    }
    rows = {
        "EEG_physical": trial_rows(
            flat_score=scores["flat"], outer_score=scores["eeg_outer"], **common
        ),
        "EOG_physical": trial_rows(
            flat_score=scores["flat"], outer_score=scores["eog_outer"], **common
        ),
        "prestim_physical": trial_rows(
            flat_score=scores["flat"],
            outer_score=scores["prestim_outer"],
            **common,
        ),
    }
    wrong_audit = {}
    for groups in WRONG_GROUPS:
        key = mapping_key(groups)
        wrong_score, wrong_audit[key] = fit_wrong_mapping_score(
            train_eeg_outer,
            z["command"][train],
            z["target"][train],
            test_eeg_outer,
            groups,
        )
        rows[f"wrong_{key}"] = trial_rows(
            flat_score=scores["flat"],
            outer_score=wrong_score,
            command=common["command"],
            trial=common["trial"],
            target=common["target"],
            horizon=3,
            pooling_groups=groups,
            correctness_groups=PHYSICAL_GROUPS,
        )
    return {
        "metrics": {name: certificate_metrics(value) for name, value in rows.items()},
        "trial_rows": rows,
        "fit_audit": fit_audit,
        "wrong_fit_audit": wrong_audit,
    }


def main() -> None:
    if RESULT.exists():
        raise FileExistsError(RESULT)
    lock = validate_lock()
    feature_audit = json.loads(FEATURE_AUDIT.read_text(encoding="utf-8"))
    if feature_audit["output_sha256"] != sha256(FEATURES):
        raise RuntimeError("Dataset-native feature hash mismatch.")
    z = np.load(FEATURES, allow_pickle=False)
    participants = {}
    for subject in range(1, 11):
        participants[str(subject)] = participant_analysis(z, subject)
        item = participants[str(subject)]["metrics"]["EEG_physical"]
        print(
            f"sub-{subject}: parent={item['n_parent_correct']}/"
            f"{item['n_trials']} AUC={item['certificate_AUC']}",
            flush=True,
        )

    def metric(subject, name, field):
        return participants[str(subject)]["metrics"][name][field]

    evaluable = [
        s
        for s in range(1, 11)
        if metric(s, "EEG_physical", "n_trials") >= 12
        and metric(s, "EEG_physical", "n_parent_correct") > 0
        and metric(s, "EEG_physical", "n_parent_error") > 0
    ]
    aucs = [metric(s, "EEG_physical", "certificate_AUC") for s in evaluable]
    flat = [
        metric(s, "EEG_physical", "certificate_AUC")
        - metric(s, "EEG_physical", "flat_margin_AUC")
        for s in evaluable
    ]
    eog = [
        metric(s, "EEG_physical", "certificate_AUC")
        - metric(s, "EOG_physical", "certificate_AUC")
        for s in evaluable
    ]
    mapping = []
    for s in evaluable:
        wrong = [
            value["certificate_AUC"]
            for name, value in participants[str(s)]["metrics"].items()
            if name.startswith("wrong_")
        ]
        mapping.append(metric(s, "EEG_physical", "certificate_AUC") - max(wrong))
    agreement = [
        metric(s, "EEG_physical", "agreement_accuracy_gain") for s in range(1, 11)
    ]
    coverage = [
        metric(s, "EEG_physical", "agreement_coverage") for s in range(1, 11)
    ]
    prestim = [
        metric(s, "prestim_physical", "certificate_AUC") for s in evaluable
    ]
    summaries = {
        "E1": effect(aucs, null=0.5),
        "E2": effect(flat, signflip=True),
        "E3": effect(eog, signflip=True),
        "E4": effect(mapping, signflip=True),
        "E5": effect(agreement, signflip=True),
        "prestim": effect(prestim, null=0.5),
    }
    gates = {
        "E1": absolute_gate(summaries["E1"]),
        "E2": paired_gate(summaries["E2"], 0.08),
        "E3": paired_gate(summaries["E3"], 0.05),
        "E4": paired_gate(summaries["E4"], 0.10),
        "E5": (
            all(metric(s, "EEG_physical", "n_trials") >= 12 for s in range(1, 11))
            and summaries["E5"]["n_total"] == 10
            and summaries["E5"]["mean"] >= 0.03
            and float(np.mean(coverage)) >= 0.50
            and summaries["E5"]["n_positive"] >= 8
            and summaries["E5"]["one_sided_exact_p"] <= 0.05
        ),
        "prestim_negative": not absolute_gate(summaries["prestim"]),
    }
    passed = all(gates.values())
    result = {
        "route": "WZ-HIC-ASME30-dataset-native-v2",
        "status": (
            "dataset_native_robustness_supported"
            if passed
            else "dataset_native_v2_not_supported_no_retuning"
        ),
        "epistemic_label": "secondary_input_QC_prespecified_not_primary_rescue",
        "created_utc": datetime.now(timezone.utc).isoformat(),
        "all_gates_pass": passed,
        "gates": gates,
        "evaluable_participants": evaluable,
        "summaries": summaries,
        "agreement_coverage_mean": float(np.mean(coverage)),
        "participants": participants,
        "provenance": {
            "protocol_sha256": sha256(PROTOCOL),
            "implementation_lock_sha256": sha256(LOCK),
            "features_sha256": sha256(FEATURES),
            "feature_audit_sha256": sha256(FEATURE_AUDIT),
        },
        "implementation_lock": lock,
    }
    RESULT.parent.mkdir(parents=True, exist_ok=True)
    RESULT.write_text(json.dumps(result, indent=2) + "\n", encoding="utf-8")
    print(json.dumps({"status": result["status"], "gates": gates}, indent=2))
    print(f"result={RESULT} sha256={sha256(RESULT)}")


if __name__ == "__main__":
    main()
```
