## Supplementary material for "Falsifiable substitution tests reveal task-structured neural evidence for auditory attention": Article_Analysis_Source_Code: run_wz_hic_asme30_external_v1.py

```
#!/usr/bin/env python3
"""Run the single frozen WZ-HIC-ASME30 external validation decision."""

from __future__ import annotations

import hashlib
import json
from datetime import datetime, timezone
from pathlib import Path

import numpy as np

from wz_hic_asme30_core_v1 import (
    BIN_EDGES_S,
    PHYSICAL_GROUPS,
    WRONG_GROUPS,
    bootstrap_mean_ci,
    certificate_metrics,
    exact_sign_p,
    exact_signflip_mean_p,
    fit_primary_event_scores,
    fit_wrong_mapping_score,
    trial_rows,
)


ROOT = Path(__file__).resolve().parents[1]
FEATURES = ROOT / "data" / "derived" / "asme30_wz_hic_features_v1.npz"
FEATURE_AUDIT = (
    ROOT / "data" / "inventory" / "asme30_wz_hic_feature_audit_v1.json"
)
PROTOCOL = ROOT / "docs" / "wz_hic_asme30_external_protocol_v1.json"
LOCK = ROOT / "docs" / "wz_hic_asme30_external_implementation_lock_v1.json"
RESULT = (
    ROOT
    / "results"
    / "wz_hic_asme30"
    / "external_v1"
    / "external_decision_v1.json"
)


def sha256(path: Path) -> str:
    h = hashlib.sha256()
    with path.open("rb") as f:
        for block in iter(lambda: f.read(1024 * 1024), b""):
            h.update(block)
    return h.hexdigest()


def mapping_key(groups) -> str:
    return "-".join(".".join(str(x) for x in group) for group in groups)


def validate_lock() -> dict:
    lock = json.loads(LOCK.read_text(encoding="utf-8"))
    files = {
        "protocol": PROTOCOL,
        "structure_audit": (
            ROOT / "docs" / "asme30_parent_child_identifiability_audit_v1.json"
        ),
        "core": ROOT / "analysis" / "wz_hic_asme30_core_v1.py",
        "feature_builder": ROOT / "analysis" / "build_asme30_wz_hic_features_v1.py",
        "runner": Path(__file__),
        "synthetic_test": ROOT / "analysis" / "test_wz_hic_asme30_synthetic_v1.py",
    }
    observed = {name: sha256(path) for name, path in files.items()}
    if observed != lock["locked_sha256"]:
        raise RuntimeError(f"Implementation lock mismatch: {observed}")
    return lock


def select_bins(x: np.ndarray, window: tuple[float, float]) -> np.ndarray:
    starts, ends = BIN_EDGES_S[:-1], BIN_EDGES_S[1:]
    ids = np.flatnonzero(
        (starts >= window[0] - 1e-9) & (ends <= window[1] + 1e-9)
    )
    if ids.size == 0:
        raise ValueError(window)
    return x[:, :, ids].reshape(x.shape[0], -1)


def effect(
    values: list[float], *, null: float = 0.0, signflip: bool = False
) -> dict:
    a = np.asarray(values, dtype=float)
    a = a[np.isfinite(a)]
    differences = a - null
    return {
        "mean": float(np.mean(a)) if a.size else float("nan"),
        "mean_difference_from_null": (
            float(np.mean(differences)) if a.size else float("nan")
        ),
        "bootstrap_95_CI_mean": (
            bootstrap_mean_ci([float(x) for x in a])
            if a.size
            else [float("nan"), float("nan")]
        ),
        "n_positive": int(np.sum(differences > 0)),
        "n_total": int(a.size),
        "fraction_positive": (
            float(np.mean(differences > 0)) if a.size else float("nan")
        ),
        "one_sided_exact_p": (
            exact_signflip_mean_p([float(x) for x in differences])
            if signflip and a.size
            else exact_sign_p([float(x) for x in a], null)
            if a.size
            else float("nan")
        ),
        "test": "exact_signflip_mean" if signflip else "exact_sign",
        "participant_values": [float(x) for x in a],
    }


def absolute_gate(summary: dict) -> bool:
    return (
        summary["n_total"] >= 5
        and summary["mean"] >= 0.75
        and summary["n_positive"] == summary["n_total"]
        and summary["one_sided_exact_p"] <= 0.03125
    )


def paired_gate(summary: dict, threshold: float) -> bool:
    return (
        summary["n_total"] >= 5
        and summary["mean"] >= threshold
        and summary["fraction_positive"] >= 0.8
        and summary["one_sided_exact_p"] <= 0.05
    )


def participant_analysis(z, subject: int) -> dict:
    accepted = z["accepted"]
    train_mask = (
        (z["subject"] == subject) & (z["task"] == "offline") & accepted
    )
    test_mask = (
        (z["subject"] == subject) & (z["task"] == "online") & accepted
    )
    train_eeg_outer = select_bins(z["eeg_bins"][train_mask], (-0.1, 1.0))
    test_eeg_outer = select_bins(z["eeg_bins"][test_mask], (-0.1, 1.0))
    train_eog_outer = select_bins(z["eog_bins"][train_mask], (-0.1, 1.0))
    test_eog_outer = select_bins(z["eog_bins"][test_mask], (-0.1, 1.0))
    train_flat = select_bins(z["eeg_bins"][train_mask], (0.3, 0.8))
    test_flat = select_bins(z["eeg_bins"][test_mask], (0.3, 0.8))
    scores, fit_audit = fit_primary_event_scores(
        train_eeg_outer=train_eeg_outer,
        train_eog_outer=train_eog_outer,
        train_prestim=z["prestim_raw"][train_mask],
        train_flat=train_flat,
        train_command=z["command"][train_mask],
        train_target=z["target"][train_mask],
        test_eeg_outer=test_eeg_outer,
        test_eog_outer=test_eog_outer,
        test_prestim=z["prestim_raw"][test_mask],
        test_flat=test_flat,
    )
    common = {
        "command": z["command"][test_mask],
        "trial": z["trial"][test_mask],
        "target": z["target"][test_mask],
        "horizon": 3,
    }
    rows = {
        "EEG_physical": trial_rows(
            flat_score=scores["flat"],
            outer_score=scores["eeg_outer"],
            **common,
        ),
        "EOG_physical": trial_rows(
            flat_score=scores["flat"],
            outer_score=scores["eog_outer"],
            **common,
        ),
        "prestim_physical": trial_rows(
            flat_score=scores["flat"],
            outer_score=scores["prestim_outer"],
            **common,
        ),
    }
    wrong_fit_audit = {}
    for groups in WRONG_GROUPS:
        key = mapping_key(groups)
        wrong_score, wrong_fit_audit[key] = fit_wrong_mapping_score(
            train_eeg_outer,
            z["command"][train_mask],
            z["target"][train_mask],
            test_eeg_outer,
            groups,
        )
        rows[f"wrong_{key}"] = trial_rows(
            flat_score=scores["flat"],
            outer_score=wrong_score,
            command=common["command"],
            trial=common["trial"],
            target=common["target"],
            horizon=3,
            pooling_groups=groups,
            correctness_groups=PHYSICAL_GROUPS,
        )

    metrics = {name: certificate_metrics(item) for name, item in rows.items()}
    horizon_sensitivity = {}
    for horizon in (1, 2, 5, 10, 15):
        horizon_sensitivity[str(horizon)] = certificate_metrics(
            trial_rows(
                flat_score=scores["flat"],
                outer_score=scores["eeg_outer"],
                command=common["command"],
                trial=common["trial"],
                target=common["target"],
                horizon=horizon,
            )
        )
    return {
        "n_offline_events_accepted": int(np.sum(train_mask)),
        "n_online_events_accepted": int(np.sum(test_mask)),
        "primary_metrics": metrics,
        "primary_trial_rows": rows,
        "fit_audit": fit_audit,
        "wrong_mapping_fit_audit": wrong_fit_audit,
        "non_gating_horizon_sensitivity": horizon_sensitivity,
    }


def main() -> None:
    if RESULT.exists():
        raise FileExistsError(f"Single-run result already exists: {RESULT}")
    lock = validate_lock()
    feature_audit = json.loads(FEATURE_AUDIT.read_text(encoding="utf-8"))
    if feature_audit["status"] != "PASS":
        raise RuntimeError("Feature audit failed.")
    if feature_audit["output_sha256"] != sha256(FEATURES):
        raise RuntimeError("Feature hash mismatch.")
    if feature_audit["protocol_sha256"] != sha256(PROTOCOL):
        raise RuntimeError("Protocol/feature hash mismatch.")
    z = np.load(FEATURES, allow_pickle=False)
    participants = {}
    for subject in range(1, 11):
        participants[str(subject)] = participant_analysis(z, subject)
        m = participants[str(subject)]["primary_metrics"]["EEG_physical"]
        print(
            f"sub-{subject}: trials={m['n_trials']} "
            f"parent={m['n_parent_correct']}/{m['n_trials']} "
            f"AUC={m['certificate_AUC']}",
            flush=True,
        )

    def metric(subject: int, name: str, field: str) -> float:
        return participants[str(subject)]["primary_metrics"][name][field]

    evaluable = [
        subject
        for subject in range(1, 11)
        if metric(subject, "EEG_physical", "n_trials") >= 12
        and metric(subject, "EEG_physical", "n_parent_correct") > 0
        and metric(subject, "EEG_physical", "n_parent_error") > 0
    ]
    ceiling = [
        subject
        for subject in range(1, 11)
        if metric(subject, "EEG_physical", "n_trials") >= 12
        and metric(subject, "EEG_physical", "n_parent_error") == 0
    ]
    aucs = [
        metric(subject, "EEG_physical", "certificate_AUC")
        for subject in evaluable
    ]
    flat_gains = [
        metric(subject, "EEG_physical", "certificate_AUC")
        - metric(subject, "EEG_physical", "flat_margin_AUC")
        for subject in evaluable
    ]
    eog_gains = [
        metric(subject, "EEG_physical", "certificate_AUC")
        - metric(subject, "EOG_physical", "certificate_AUC")
        for subject in evaluable
    ]
    mapping_gains = []
    for subject in evaluable:
        wrong_aucs = [
            value["certificate_AUC"]
            for key, value in participants[str(subject)]["primary_metrics"].items()
            if key.startswith("wrong_")
        ]
        mapping_gains.append(
            metric(subject, "EEG_physical", "certificate_AUC") - max(wrong_aucs)
        )
    agreement_gains = [
        metric(subject, "EEG_physical", "agreement_accuracy_gain")
        for subject in range(1, 11)
    ]
    agreement_coverages = [
        metric(subject, "EEG_physical", "agreement_coverage")
        for subject in range(1, 11)
    ]
    prestim_aucs = [
        metric(subject, "prestim_physical", "certificate_AUC")
        for subject in evaluable
    ]

    E1 = effect(aucs, null=0.5)
    E2 = effect(flat_gains, signflip=True)
    E3 = effect(eog_gains, signflip=True)
    E4 = effect(mapping_gains, signflip=True)
    E5 = effect(agreement_gains, signflip=True)
    prestim = effect(prestim_aucs, null=0.5)
    all_trial_counts_ok = all(
        metric(subject, "EEG_physical", "n_trials") >= 12
        for subject in range(1, 11)
    )
    gates = {
        "E1_external_certificate": absolute_gate(E1),
        "E2_beyond_flat_confidence": paired_gate(E2, 0.08),
        "E3_beyond_ocular_certificate": paired_gate(E3, 0.05),
        "E4_mapping_specificity": paired_gate(E4, 0.10),
        "E5_selective_reliability": (
            all_trial_counts_ok
            and E5["n_total"] == 10
            and E5["mean"] >= 0.03
            and float(np.mean(agreement_coverages)) >= 0.50
            and E5["n_positive"] >= 8
            and E5["one_sided_exact_p"] <= 0.05
        ),
        "prestimulus_negative_control": not absolute_gate(prestim),
    }
    pass_all = all(gates.values())

    horizon_summary = {}
    for horizon in (1, 2, 5, 10, 15):
        values = [
            participants[str(subject)]["non_gating_horizon_sensitivity"][
                str(horizon)
            ]["certificate_AUC"]
            for subject in evaluable
        ]
        horizon_summary[str(horizon)] = effect(values, null=0.5)

    result = {
        "route": "WZ-HIC-ASME30-external-v1",
        "status": (
            "externally_confirmed_on_temporally_later_online_sessions"
            if pass_all
            else "external_v1_not_confirmed_no_retuning"
        ),
        "created_utc": datetime.now(timezone.utc).isoformat(),
        "all_gates_pass": pass_all,
        "gates": gates,
        "evaluable_participants": evaluable,
        "flat_parent_ceiling_participants": ceiling,
        "primary_outcomes": {
            "E1_external_certificate": E1,
            "E2_beyond_flat_confidence": E2,
            "E3_beyond_ocular_certificate": E3,
            "E4_mapping_specificity": E4,
            "E5_selective_reliability": E5,
            "agreement_coverage_mean": float(np.mean(agreement_coverages)),
        },
        "negative_controls": {"prestimulus": prestim},
        "participants": participants,
        "non_gating_horizon_sensitivity": horizon_summary,
        "provenance": {
            "protocol_sha256": sha256(PROTOCOL),
            "implementation_lock_sha256": sha256(LOCK),
            "features_sha256": sha256(FEATURES),
            "feature_audit_sha256": sha256(FEATURE_AUDIT),
        },
        "implementation_lock": lock,
    }
    RESULT.parent.mkdir(parents=True, exist_ok=True)
    RESULT.write_text(json.dumps(result, indent=2) + "\n", encoding="utf-8")
    print(json.dumps({"status": result["status"], "gates": gates}, indent=2))
    print(f"result={RESULT} sha256={sha256(RESULT)}")


if __name__ == "__main__":
    main()
```
