## Supplementary material for "Falsifiable substitution tests reveal task-structured neural evidence for auditory attention": Article_Analysis_Source_Code: run_wz_hic_asme30_target_free_spatiotemporal_v1.py

```
#!/usr/bin/env python3
"""Frozen target-free scalp-time mechanism localization for ASME30."""

from __future__ import annotations

import csv
import hashlib
import itertools
import json
import math
from datetime import datetime, timezone
from pathlib import Path

import numpy as np

from wz_hic_asme30_core_v1 import (
    PHYSICAL_GROUPS,
    WRONG_GROUPS,
    labels_for_mapping,
)


ROOT = Path(__file__).resolve().parents[1]
PROTOCOL = (
    ROOT
    / "docs"
    / "wz_hic_asme30_target_free_spatiotemporal_protocol_v1.json"
)
LOCK = (
    ROOT
    / "docs"
    / "wz_hic_asme30_target_free_spatiotemporal_implementation_lock_v1.json"
)
FEATURES = ROOT / "data" / "derived" / "asme30_wz_hic_features_v1.npz"
OUTDIR = (
    ROOT / "results" / "wz_hic_asme30" / "target_free_spatiotemporal_v1"
)
RESULT = OUTDIR / "spatiotemporal_decision_v1.json"
PARTICIPANT_CSV = OUTDIR / "participant_projections_v1.csv"
CELL_CSV = OUTDIR / "scalp_time_cells_v1.csv"

PROTOCOL_SHA256 = (
    "69a1278c6e5be9d529c97c37a95d4c1df13eb5064322099d7d1856a1fa4699f2"
)
FEATURES_SHA256 = (
    "0159ba568625845731789e54402fbed98c93edd2db3afd0bd41e1507e1c7f756"
)
HORIZON = 3
MIN_TRIALS = 10
BOOTSTRAP_N = 20_000
BOOTSTRAP_SEED = 20260730


def sha256(path: Path) -> str:
    digest = hashlib.sha256()
    with path.open("rb") as handle:
        for block in iter(lambda: handle.read(1024 * 1024), b""):
            digest.update(block)
    return digest.hexdigest()


def validate_frozen_inputs() -> None:
    observed = {
        "protocol": sha256(PROTOCOL),
        "features": sha256(FEATURES),
    }
    expected = {
        "protocol": PROTOCOL_SHA256,
        "features": FEATURES_SHA256,
    }
    if observed != expected:
        raise RuntimeError(f"Frozen input mismatch: {observed!r} != {expected!r}")
    if not LOCK.exists():
        raise FileNotFoundError("Implementation lock must exist before outcomes.")
    lock = json.loads(LOCK.read_text(encoding="utf-8"))
    if lock.get("synthetic_test_status") != "PASS":
        raise RuntimeError("Synthetic test was not locked as PASS.")
    if lock.get("scientific_outcomes_available_at_lock") is not False:
        raise RuntimeError("Lock does not certify outcome unavailability.")


def group_labels(
    commands: np.ndarray, target: int, groups
) -> np.ndarray:
    targets = np.full(commands.size, int(target), dtype=int)
    return labels_for_mapping(commands, targets, groups)


def trial_map(
    eeg: np.ndarray,
    commands: np.ndarray,
    target: int,
    groups,
) -> np.ndarray:
    """Return target-free same-parent minus other-parent scalp-time map."""

    labels = group_labels(commands, target, groups)
    if int(np.sum(labels == "T")) != HORIZON:
        raise RuntimeError("Target removal count is not the frozen horizon.")
    if int(np.sum(labels == "S")) != 9 * HORIZON:
        raise RuntimeError("Same-parent non-target count is not 27.")
    if int(np.sum(labels == "U")) != 20 * HORIZON:
        raise RuntimeError("Other-parent count is not 60.")
    return np.mean(eeg[labels == "S"], axis=0) - np.mean(
        eeg[labels == "U"], axis=0
    )


def participant_maps(
    *,
    eeg_bins: np.ndarray,
    command: np.ndarray,
    target: np.ndarray,
    trial: np.ndarray,
) -> dict:
    physical_trials = []
    wrong_trials = [[] for _ in WRONG_GROUPS]
    eligible_trials = []
    ineligible_trials = []
    for trial_id in sorted(np.unique(trial)):
        ids = np.flatnonzero(trial == trial_id)
        target_values = np.unique(target[ids])
        if target_values.size != 1:
            raise RuntimeError(f"Trial {trial_id} has nonunique target.")
        by_command = {}
        for value in range(1, 31):
            value_ids = ids[command[ids] == value]
            if value_ids.size >= HORIZON:
                by_command[value] = value_ids[:HORIZON]
        if len(by_command) != 30:
            ineligible_trials.append(
                {
                    "trial": int(trial_id),
                    "minimum_accepted_repetitions": int(
                        min(
                            np.sum(command[ids] == value)
                            for value in range(1, 31)
                        )
                    ),
                }
            )
            continue
        selected = np.concatenate([by_command[value] for value in range(1, 31)])
        selected_commands = command[selected]
        selected_eeg = eeg_bins[selected, :, 1:]
        target_value = int(target_values[0])
        physical_trials.append(
            trial_map(
                selected_eeg,
                selected_commands,
                target_value,
                PHYSICAL_GROUPS,
            )
        )
        for index, groups in enumerate(WRONG_GROUPS):
            wrong_trials[index].append(
                trial_map(
                    selected_eeg,
                    selected_commands,
                    target_value,
                    groups,
                )
            )
        eligible_trials.append(int(trial_id))
    result = {
        "n_eligible_trials": len(eligible_trials),
        "eligible_trials": eligible_trials,
        "ineligible_trials": ineligible_trials,
        "evaluable": len(eligible_trials) >= MIN_TRIALS,
    }
    if result["evaluable"]:
        result["physical"] = np.mean(np.stack(physical_trials), axis=0)
        result["wrong"] = [
            np.mean(np.stack(values), axis=0) for values in wrong_trials
        ]
    return result


def lopo_projections(
    physical_maps: np.ndarray, wrong_maps: np.ndarray
) -> list[dict]:
    rows = []
    for index in range(physical_maps.shape[0]):
        others = np.delete(physical_maps, index, axis=0)
        template = np.mean(others, axis=0)
        norm = float(np.linalg.norm(template))
        if not np.isfinite(norm) or norm <= 1e-12:
            raise RuntimeError("Degenerate LOPO template.")
        template /= norm
        physical = float(np.sum(physical_maps[index] * template))
        wrong = [
            float(np.sum(wrong_maps[index, mapping] * template))
            for mapping in range(wrong_maps.shape[1])
        ]
        rows.append(
            {
                "physical_projection": physical,
                "wrong_1_projection": wrong[0],
                "wrong_2_projection": wrong[1],
                "best_wrong_projection": max(wrong),
                "physical_minus_wrong_1": physical - wrong[0],
                "physical_minus_wrong_2": physical - wrong[1],
                "physical_minus_best_wrong": physical - max(wrong),
                "template_norm_before_normalization": norm,
            }
        )
    return rows


def bootstrap_ci(values: np.ndarray) -> list[float]:
    rng = np.random.default_rng(BOOTSTRAP_SEED)
    draws = values[
        rng.integers(
            0,
            values.size,
            size=(BOOTSTRAP_N, values.size),
        )
    ].mean(axis=1)
    return [float(value) for value in np.quantile(draws, [0.025, 0.975])]


def exact_signflip_p(values: np.ndarray) -> float:
    observed = float(np.mean(values))
    permuted = [
        float(np.mean(values * np.asarray(signs, dtype=float)))
        for signs in itertools.product((-1.0, 1.0), repeat=values.size)
    ]
    return float(np.mean(np.asarray(permuted) >= observed - 1e-15))


def t_map(values: np.ndarray) -> np.ndarray:
    n = values.shape[0]
    mean = np.mean(values, axis=0)
    sd = np.std(values, axis=0, ddof=1)
    denominator = sd / math.sqrt(n)
    return np.divide(
        mean,
        denominator,
        out=np.zeros_like(mean, dtype=float),
        where=denominator > 1e-12,
    )


def exact_max_t(physical_maps: np.ndarray) -> tuple[np.ndarray, np.ndarray, np.ndarray]:
    observed = t_map(physical_maps)
    maxima = []
    for signs in itertools.product(
        (-1.0, 1.0), repeat=physical_maps.shape[0]
    ):
        signed = physical_maps * np.asarray(signs)[:, None, None]
        maxima.append(float(np.max(np.abs(t_map(signed)))))
    maxima_array = np.asarray(maxima, dtype=float)
    p_fwer = np.mean(
        maxima_array[:, None, None]
        >= np.abs(observed)[None, :, :] - 1e-15,
        axis=0,
    )
    return observed, p_fwer, maxima_array


def run_analysis(arrays: dict) -> dict:
    all_participants = []
    map_records = {}
    for subject in sorted(int(value) for value in np.unique(arrays["subject"])):
        mask = (
            (arrays["subject"] == subject)
            & (arrays["task"] == "online")
            & arrays["accepted"]
        )
        item = participant_maps(
            eeg_bins=arrays["eeg_bins"][mask],
            command=arrays["command"][mask],
            target=arrays["target"][mask],
            trial=arrays["trial"][mask],
        )
        item["subject"] = subject
        item["accepted_online_events"] = int(np.sum(mask))
        all_participants.append(item)
        if item["evaluable"]:
            map_records[subject] = item

    evaluable_subjects = sorted(map_records)
    physical_maps = np.stack(
        [map_records[subject]["physical"] for subject in evaluable_subjects]
    )
    wrong_maps = np.stack(
        [map_records[subject]["wrong"] for subject in evaluable_subjects]
    )
    projections = lopo_projections(physical_maps, wrong_maps)
    for subject, row in zip(evaluable_subjects, projections):
        row["subject"] = subject
        row["n_eligible_trials"] = map_records[subject]["n_eligible_trials"]

    specificity = np.asarray(
        [row["physical_minus_best_wrong"] for row in projections]
    )
    required_positive = int(math.ceil(0.75 * specificity.size))
    mapping_summary = {
        "mean": float(np.mean(specificity)),
        "bootstrap_95_CI_mean": bootstrap_ci(specificity),
        "n_positive": int(np.sum(specificity > 0)),
        "n_total": int(specificity.size),
        "required_positive": required_positive,
        "one_sided_exact_signflip_p": exact_signflip_p(specificity),
        "participant_values": [float(value) for value in specificity],
    }
    for key in ("physical_minus_wrong_1", "physical_minus_wrong_2"):
        values = np.asarray([row[key] for row in projections])
        mapping_summary[key] = {
            "mean": float(np.mean(values)),
            "bootstrap_95_CI_mean": bootstrap_ci(values),
            "n_positive": int(np.sum(values > 0)),
            "one_sided_exact_signflip_p": exact_signflip_p(values),
        }

    t_values, p_fwer, max_null = exact_max_t(physical_maps)
    significant = np.argwhere(p_fwer <= 0.05 + 1e-15)
    strongest_flat = int(np.argmax(np.abs(t_values)))
    strongest_channel, strongest_bin = np.unravel_index(
        strongest_flat, t_values.shape
    )

    m0 = (
        len(evaluable_subjects) >= 8
        and all(
            map_records[subject]["n_eligible_trials"] >= MIN_TRIALS
            for subject in evaluable_subjects
        )
    )
    m1 = (
        mapping_summary["mean"] > 0
        and mapping_summary["bootstrap_95_CI_mean"][0] > 0
        and mapping_summary["n_positive"] >= required_positive
        and mapping_summary["one_sided_exact_signflip_p"] <= 0.05
    )
    m2 = bool(significant.size)
    return {
        "participants_structurally_screened": all_participants,
        "evaluable_subjects": evaluable_subjects,
        "physical_maps": physical_maps,
        "wrong_maps": wrong_maps,
        "projections": projections,
        "mapping_summary": mapping_summary,
        "t_values": t_values,
        "p_fwer": p_fwer,
        "max_t_null": max_null,
        "strongest_channel_index": int(strongest_channel),
        "strongest_bin_index": int(strongest_bin),
        "significant_indices": significant,
        "gates": {
            "M0_coverage": m0,
            "M1_binding_specific_lopo": m1,
            "M2_spatiotemporal_fwer": m2,
        },
        "supported": m0 and m1 and m2,
    }


def main() -> None:
    validate_frozen_inputs()
    if RESULT.exists() or PARTICIPANT_CSV.exists() or CELL_CSV.exists():
        raise FileExistsError("Immutable v1 output already exists.")
    with np.load(FEATURES, allow_pickle=True) as source:
        arrays = {key: source[key] for key in source.files}
    analysis = run_analysis(arrays)
    channel_names = [str(value) for value in arrays["channel_names"]]
    bin_edges = [float(value) for value in arrays["bin_edges_s"][1:]]

    cell_rows = []
    for channel_index, channel in enumerate(channel_names):
        for bin_index in range(10):
            start = bin_edges[bin_index]
            stop = float(arrays["bin_edges_s"][bin_index + 2])
            values = analysis["physical_maps"][:, channel_index, bin_index]
            cell_rows.append(
                {
                    "channel": channel,
                    "channel_index": channel_index,
                    "time_start_s": start,
                    "time_stop_s": stop,
                    "mean_s_minus_u_microvolts": float(np.mean(values)),
                    "t_value": float(
                        analysis["t_values"][channel_index, bin_index]
                    ),
                    "max_t_fwer_p": float(
                        analysis["p_fwer"][channel_index, bin_index]
                    ),
                    "fwer_significant_0_05": bool(
                        analysis["p_fwer"][channel_index, bin_index] <= 0.05
                    ),
                }
            )

    strongest = next(
        row
        for row in cell_rows
        if row["channel_index"] == analysis["strongest_channel_index"]
        and abs(row["time_start_s"] - bin_edges[analysis["strongest_bin_index"]])
        < 1e-12
    )
    output = {
        "route": "WZ-HIC-ASME30-target-free-spatiotemporal-v1",
        "created_utc": datetime.now(timezone.utc).isoformat(),
        "status": (
            "post_outcome_mechanism_localization_supported"
            if analysis["supported"]
            else "post_outcome_mechanism_localization_not_supported_no_retuning"
        ),
        "mechanism_localization_supported": analysis["supported"],
        "gates": analysis["gates"],
        "coverage": {
            "subjects_screened": 10,
            "evaluable_subjects": analysis["evaluable_subjects"],
            "n_evaluable": len(analysis["evaluable_subjects"]),
            "participant_trial_counts": [
                {
                    key: value
                    for key, value in item.items()
                    if key
                    in (
                        "subject",
                        "accepted_online_events",
                        "n_eligible_trials",
                        "evaluable",
                    )
                }
                for item in analysis["participants_structurally_screened"]
            ],
        },
        "mapping_specificity": analysis["mapping_summary"],
        "spatiotemporal": {
            "cells_tested": len(cell_rows),
            "n_fwer_significant_cells": int(
                sum(row["fwer_significant_0_05"] for row in cell_rows)
            ),
            "strongest_absolute_t_cell": strongest,
            "exact_sign_patterns": int(2 ** len(analysis["evaluable_subjects"])),
            "max_t_null_95_percentile": float(
                np.quantile(analysis["max_t_null"], 0.95)
            ),
            "physical_participant_maps": {
                str(subject): analysis["physical_maps"][index].tolist()
                for index, subject in enumerate(analysis["evaluable_subjects"])
            },
        },
        "participant_projections": analysis["projections"],
        "provenance": {
            "protocol_sha256": sha256(PROTOCOL),
            "implementation_lock_sha256": sha256(LOCK),
            "features_sha256": sha256(FEATURES),
            "runner_sha256": sha256(Path(__file__)),
        },
        "interpretation_boundary": (
            "Post-primary-outcome scalp-sensor localization on ASME30. "
            "A passing result identifies a target-free, physical-binding-"
            "specific sensor-time pattern; it is not a cortical source, an "
            "independent confirmation, or a deployable online score."
        ),
    }

    OUTDIR.mkdir(parents=True, exist_ok=True)
    RESULT.write_text(
        json.dumps(output, ensure_ascii=False, indent=2) + "\n",
        encoding="utf-8",
    )
    with PARTICIPANT_CSV.open("x", newline="", encoding="utf-8-sig") as handle:
        fieldnames = list(analysis["projections"][0])
        writer = csv.DictWriter(handle, fieldnames=fieldnames)
        writer.writeheader()
        writer.writerows(analysis["projections"])
    with CELL_CSV.open("x", newline="", encoding="utf-8-sig") as handle:
        writer = csv.DictWriter(handle, fieldnames=list(cell_rows[0]))
        writer.writeheader()
        writer.writerows(cell_rows)

    print(
        json.dumps(
            {
                "result": RESULT.relative_to(ROOT).as_posix(),
                "result_sha256": sha256(RESULT),
                "participant_csv_sha256": sha256(PARTICIPANT_CSV),
                "cell_csv_sha256": sha256(CELL_CSV),
                "status": output["status"],
                "gates": output["gates"],
                "mapping_specificity": output["mapping_specificity"],
                "n_fwer_significant_cells": output["spatiotemporal"][
                    "n_fwer_significant_cells"
                ],
                "strongest_cell": strongest,
            },
            indent=2,
        )
    )


if __name__ == "__main__":
    main()
```
