## Supplementary material for "Falsifiable substitution tests reveal task-structured neural evidence for auditory attention": Article_Analysis_Source_Code: run_wz_hic_continuous_listener_transport_v1.py

```
#!/usr/bin/env python3
"""Frozen continuous-speech listener-specific direction-family transport test."""

from __future__ import annotations

import csv
import hashlib
import itertools
import json
from pathlib import Path
from typing import Iterable

import numpy as np
from scipy.io import loadmat
from scipy.signal import resample_poly
from scipy.stats import rankdata

from kite_kul_akb_v1 import biosemi64_geometry, fit_ridge as kul_fit_ridge
from kite_kul_akb_v1 import make_windows as kul_make_windows
from kite_sat_adaptive_broad_v1 import family_weights, reliability
from kite_sat_backward_v1 import block_design as kite_block_design
from kite_sat_direction_delay_controls_v1 import TRANSFORMS, transform_vector
from kite_sat_geometry_v1 import cell_delta_contributions, partition_ids
from kite_st_hd_core_v1 import load_public_geometry
from run_kite_sat_ear_saad_external_subject_v1 import (
    BLOCKS_PER_TRIAL as EAR_BLOCKS_PER_TRIAL,
    N_CHANNELS as EAR_N_CHANNELS,
    N_FEATURES as EAR_N_FEATURES,
    SAMPLES_PER_BLOCK as EAR_SAMPLES_PER_BLOCK,
    USABLE as EAR_USABLE,
    block_design as ear_block_design,
    cell_contributions as ear_cell_contributions,
    fit_from_blocks as ear_fit_from_blocks,
    sufficient_statistics as ear_sufficient_statistics,
)


ROOT = Path(__file__).resolve().parents[1]
PROTOCOL = ROOT / "docs" / "wz_hic_continuous_listener_transport_protocol_v1.json"
PROTOCOL_SHA256 = "b5f80cc7c275724de5e823cc01aa2047e1483a0dd851eacab3018fe5440d2534"
LOCK = ROOT / "docs" / "wz_hic_continuous_listener_transport_implementation_lock_v1.json"
OUTPUT_ROOT = (
    ROOT / "results" / "wz_hic_cross_dataset"
    / "continuous_listener_transport_v1"
)
DECISION = OUTPUT_ROOT / "continuous_listener_transport_decision_v1.json"
PARTICIPANT_CSV = OUTPUT_ROOT / "participant_endpoints_v1.csv"
DIRECTION_CSV = OUTPUT_ROOT / "direction_endpoints_v1.csv"

KITE_SUBJECTS = (
    "S0", "S10", "S2", "S21", "S28", "S22",
    "S9", "S15", "S16", "S18", "S13", "S1",
)
KUL_SUBJECTS = tuple(f"S{value}" for value in range(1, 17))
EAR_SUBJECTS = tuple(f"S{value}" for value in range(1, 16))

KITE_PREPROCESSED = (
    ROOT / "data" / "derived" / "kite_st_hd_zenodo4518754"
    / "confirmation_preprocessed_v1"
)
KITE_AUDIO = (
    ROOT / "data" / "derived" / "kite_st_hd_zenodo4518754"
    / "kite_sat_audio_features_v1.npz"
)
KITE_MODELS = (
    ROOT / "data" / "derived" / "kite_st_hd_zenodo4518754"
    / "kite_sat_confirmation_complete_case_models_v2"
)
KITE_RESULTS = (
    ROOT / "results" / "kite_sat" / "akb_confirmation_complete_case_v2"
    / "subjects"
)
KITE_COORDINATES = (
    ROOT / "downloads" / "candidate_audit" / "zenodo4518754_support"
    / "misc" / "misc" / "eeg255ch_locs.csv"
)
KITE_PHYSICAL = {
    "1L": ("part1_track1", "part1_track2"),
    "1R": ("part1_track1", "part1_track2"),
    "2L": ("part2_track2", "part2_track1"),
    "2R": ("part2_track2", "part2_track1"),
}

KUL_ODD = ROOT / "data" / "derived" / "kul_discovery_32hz"
KUL_EVEN = ROOT / "data" / "derived" / "kul_confirmation_32hz"
KUL_AUDIO = KUL_ODD / "audio_powerlaw_subbands.npz"
KUL_HALVES = {"A": (1, 4), "B": (2, 3)}

EAR_ROOT = ROOT / "data" / "raw" / "ear_saad" / "preprocessedData"
EAR_GEOMETRY = ROOT / "docs" / "kite_sat_ear_saad_scalp_geometry_v1.json"
EAR_ORIGINAL_PROTOCOL = ROOT / "docs" / "kite_sat_ear_saad_external_protocol_v1.json"

ENDPOINTS = (
    "F1_fingerprint_listener_rank_advantage",
    "F2_fingerprint_physical_mapping_rank_advantage",
    "P1_predictive_listener_rank_advantage",
    "P2_predictive_physical_mapping_rank_advantage",
)
BOOTSTRAP_DRAWS = 100_000
BOOTSTRAP_SEED = 2026073001
RANDOMIZATION_DRAWS = 1_000_000
RANDOMIZATION_SEED = 2026073002


def sha256(path: Path) -> str:
    digest = hashlib.sha256()
    with path.open("rb") as handle:
        for block in iter(lambda: handle.read(8 * 1024 * 1024), b""):
            digest.update(block)
    return digest.hexdigest()


def validate_lock() -> tuple[dict, str]:
    if sha256(PROTOCOL) != PROTOCOL_SHA256:
        raise RuntimeError("Continuous-listener protocol hash mismatch.")
    if not LOCK.exists():
        raise RuntimeError("Implementation lock is missing.")
    payload = json.loads(LOCK.read_text(encoding="utf-8"))
    lock_hash = sha256(LOCK)
    if (
        payload["status"]
        != "frozen_before_computing_any_continuous_listener_transport_outcome"
        or payload["protocol_sha256"] != PROTOCOL_SHA256
        or payload["runner"]["path"] != Path(__file__).relative_to(ROOT).as_posix()
        or payload["runner"]["sha256"] != sha256(Path(__file__))
    ):
        raise RuntimeError("Continuous-listener implementation lock mismatch.")
    for record in payload["immutable_dependencies"]:
        if sha256(ROOT / record["path"]) != record["sha256"]:
            raise RuntimeError(f"Dependency hash mismatch: {record['path']}")
    return payload, lock_hash


def pearson(a: np.ndarray, b: np.ndarray) -> float:
    left = np.asarray(a, dtype=np.float64)
    right = np.asarray(b, dtype=np.float64)
    if left.shape != (16,) or right.shape != (16,):
        raise ValueError("Fingerprint correlation expects two 16-cell profiles.")
    left = left - np.mean(left)
    right = right - np.mean(right)
    denominator = float(np.sqrt(np.sum(left * left) * np.sum(right * right)))
    if denominator <= 1e-12:
        raise RuntimeError("Degenerate continuous-speech reliability profile.")
    return float(np.sum(left * right) / denominator)


def rank_probability(value: np.ndarray | float, comparators: np.ndarray) -> float:
    reference = np.asarray(value, dtype=np.float64)
    other = np.asarray(comparators, dtype=np.float64)
    return float(np.mean((reference[..., None] > other) + 0.5 * (reference[..., None] == other)))


def binary_auc(scores: np.ndarray, labels: np.ndarray) -> float:
    values = np.asarray(scores, dtype=np.float64)
    outcome = np.asarray(labels, dtype=np.int64)
    positive = values[outcome == 1]
    negative = values[outcome == 0]
    if not len(positive) or not len(negative):
        return float("nan")
    joined = np.r_[positive, negative]
    ranks = rankdata(joined, method="average")
    u = float(np.sum(ranks[: len(positive)]) - len(positive) * (len(positive) + 1) / 2)
    return u / (len(positive) * len(negative))


def model_from_archive(saved: np.lib.npyio.NpzFile, prefix: str) -> dict[str, np.ndarray]:
    return {
        "x_mean": np.asarray(saved[f"{prefix}_x_mean"], dtype=np.float64),
        "x_scale": np.asarray(saved[f"{prefix}_x_scale"], dtype=np.float64),
        "weights": np.asarray(saved[f"{prefix}_weights"], dtype=np.float64),
    }


def kite_offset(record: dict) -> float:
    return (
        record["crop"]["padded_start"]
        + 3000
        - record["trigger_train"]["first_sample"]
    ) / 1000.0


def load_kite_dataset() -> tuple[list[dict], list[dict]]:
    with np.load(KITE_AUDIO, allow_pickle=False) as saved:
        audio = {name: np.asarray(saved[name]) for name in saved.files}
    geometry = load_public_geometry(KITE_COORDINATES)
    ids = partition_ids(geometry.xy)["S16"]
    participants: list[dict] = []
    inputs: list[dict] = [
        {"path": KITE_AUDIO.relative_to(ROOT).as_posix(), "sha256": sha256(KITE_AUDIO)},
        {
            "path": KITE_COORDINATES.relative_to(ROOT).as_posix(),
            "sha256": sha256(KITE_COORDINATES),
        },
    ]
    for subject in KITE_SUBJECTS:
        result_path = KITE_RESULTS / f"{subject}_AKB_confirmation_v1.json"
        model_path = KITE_MODELS / f"{subject}_AKB_confirmation_models_v1.npz"
        result = json.loads(result_path.read_text(encoding="utf-8"))
        manifest_path = ROOT / result["preprocessing_manifest"]["path"]
        manifest = json.loads(manifest_path.read_text(encoding="utf-8"))
        preprocessing = {
            row["recording"].rsplit("_", 1)[-1]: row
            for row in manifest["preprocessing_records"]
        }
        inputs.extend(
            [
                {"path": result_path.relative_to(ROOT).as_posix(), "sha256": sha256(result_path)},
                {"path": model_path.relative_to(ROOT).as_posix(), "sha256": sha256(model_path)},
                {"path": manifest_path.relative_to(ROOT).as_posix(), "sha256": sha256(manifest_path)},
            ]
        )
        condition_blocks: dict[str, list[dict]] = {}
        for condition in ("1L", "1R", "2L", "2R"):
            input_path = (
                KITE_PREPROCESSED / subject
                / f"{subject}_AAD_{condition}_preprocessed_v1.npz"
            )
            observed_hash = sha256(input_path)
            if observed_hash != preprocessing[condition]["output"]["sha256"]:
                raise RuntimeError(f"KITE input hash mismatch: {subject} {condition}")
            inputs.append({"path": input_path.relative_to(ROOT).as_posix(), "sha256": observed_hash})
            with np.load(input_path, allow_pickle=False) as saved:
                eeg = np.asarray(saved["eeg"])
            offset = int(round(kite_offset(preprocessing[condition]) * 100))
            left_name, right_name = KITE_PHYSICAL[condition]
            rows = []
            for block_index in range(12):
                first = offset + block_index * 3000
                stop = first + 3000
                left100 = audio[left_name][first:stop]
                right100 = audio[right_name][first:stop]
                design, left_features = kite_block_design(
                    eeg[block_index * 3000 : (block_index + 1) * 3000],
                    left100,
                )
                right_features = resample_poly(right100, 1, 5, axis=0)[:595]
                rows.append(
                    {
                        "design": design,
                        "left": left_features[:, 0],
                        "right": right_features[:, 0],
                        "label": 1 if condition.endswith("L") else 0,
                        "condition": condition,
                        "block": block_index + 1,
                    }
                )
            condition_blocks[condition] = rows
        with np.load(model_path, allow_pickle=False) as saved:
            profiles = {
                "A": np.asarray(result["fold_reliabilities"]["train_story_1"], dtype=np.float64),
                "B": np.asarray(result["fold_reliabilities"]["train_story_2"], dtype=np.float64),
            }
            models = {
                "A": model_from_archive(saved, "train_story_1"),
                "B": model_from_archive(saved, "train_story_2"),
            }
        directions = {}
        for source, target_conditions in (
            ("A", ("2L", "2R")),
            ("B", ("1L", "1R")),
        ):
            rows = [row for condition in target_conditions for row in condition_blocks[condition]]
            deltas, labels, metadata = [], [], []
            for row in rows:
                delta, _ = cell_delta_contributions(
                    row["design"], models[source], row["left"], row["right"], ids
                )
                deltas.append(delta)
                labels.append(row["label"])
                metadata.append({"condition": row["condition"], "block": row["block"]})
            directions[f"{source}_to_{'B' if source == 'A' else 'A'}"] = {
                "source": source,
                "target": "B" if source == "A" else "A",
                "deltas": np.asarray(deltas),
                "labels": np.asarray(labels, dtype=np.int64),
                "metadata": metadata,
            }
        participants.append(
            {"subject": subject, "profiles": profiles, "directions": directions}
        )
        print(f"KITE-ST {subject} complete", flush=True)
    return participants, inputs


def kul_profile(windows: list, parts: tuple[int, int], ids: np.ndarray) -> np.ndarray:
    aligned = []
    for held_part in parts:
        training = [window for window in windows if window.part in parts and window.part != held_part]
        held = [window for window in windows if window.part == held_part]
        model = kul_fit_ridge(training)
        for window in held:
            delta, _ = cell_delta_contributions(
                window.design, model, window.left, window.right, ids
            )
            aligned.append((1.0 if window.attended_ear == "L" else -1.0) * delta)
    return reliability(np.asarray(aligned))


def load_kul_dataset() -> tuple[list[dict], list[dict]]:
    with np.load(KUL_AUDIO, allow_pickle=False) as saved:
        audio = {
            name.removeprefix("audio::"): np.asarray(saved[name])
            for name in saved.files
            if name.startswith("audio::")
        }
    _, _, ids = biosemi64_geometry()
    participants: list[dict] = []
    inputs = [{"path": KUL_AUDIO.relative_to(ROOT).as_posix(), "sha256": sha256(KUL_AUDIO)}]
    for subject_text in KUL_SUBJECTS:
        subject = int(subject_text[1:])
        root = KUL_ODD if subject % 2 else KUL_EVEN
        input_path = root / f"S{subject}.npz"
        inputs.append({"path": input_path.relative_to(ROOT).as_posix(), "sha256": sha256(input_path)})
        with np.load(input_path, allow_pickle=False) as saved:
            metadata = json.loads(str(saved["metadata_json"]))
            eeg = [np.asarray(saved[f"eeg_{index:02d}"]) for index in range(1, 21)]
        windows = kul_make_windows(eeg, metadata, audio)
        profiles = {
            name: kul_profile(windows, parts, ids)
            for name, parts in KUL_HALVES.items()
        }
        directions = {}
        for source, target in (("A", "B"), ("B", "A")):
            source_windows = [window for window in windows if window.part in KUL_HALVES[source]]
            target_windows = [window for window in windows if window.part in KUL_HALVES[target]]
            model = kul_fit_ridge(source_windows)
            deltas, labels, metadata_rows = [], [], []
            for window in target_windows:
                delta, _ = cell_delta_contributions(
                    window.design, model, window.left, window.right, ids
                )
                deltas.append(delta)
                labels.append(1 if window.attended_ear == "L" else 0)
                metadata_rows.append(
                    {
                        "trial": int(window.trial),
                        "part": int(window.part),
                        "window_index": int(window.window_index),
                        "condition": str(window.condition),
                    }
                )
            directions[f"{source}_to_{target}"] = {
                "source": source,
                "target": target,
                "deltas": np.asarray(deltas),
                "labels": np.asarray(labels, dtype=np.int64),
                "metadata": metadata_rows,
            }
        participants.append(
            {"subject": subject_text, "profiles": profiles, "directions": directions}
        )
        print(f"KUL {subject_text} complete", flush=True)
    return participants, inputs


def ear_split(attended: np.ndarray, video: np.ndarray) -> dict[str, tuple[int, ...]]:
    candidates = []
    for first in itertools.combinations(range(6), 3):
        if 0 not in first:
            continue
        second = tuple(index for index in range(6) if index not in first)
        if any(
            len(set(values[list(indices)].tolist())) != 2
            for values in (attended, video)
            for indices in (first, second)
        ):
            continue
        imbalance = (
            abs(int(np.sum(attended[list(first)] == 1)) - int(np.sum(attended[list(second)] == 1)))
            + abs(int(np.sum(video[list(first)] == 1)) - int(np.sum(video[list(second)] == 1)))
        )
        candidates.append((imbalance, first, second))
    if not candidates:
        raise RuntimeError("No Ear-SAAD split satisfies the locked balance rule.")
    _, first, second = sorted(candidates)[0]
    return {"A": first, "B": second}


def ear_profile(records: list[dict], trials: tuple[int, ...], ids: np.ndarray) -> np.ndarray:
    aligned = []
    for held_trial in trials:
        training = [
            index for index, row in enumerate(records)
            if row["trial"] in trials and row["trial"] != held_trial
        ]
        held = [
            index for index, row in enumerate(records)
            if row["trial"] == held_trial
        ]
        model = ear_fit_from_blocks(records, training)
        for index in held:
            row = records[index]
            delta, _ = ear_cell_contributions(
                row["design"], model, row["speaker1"], row["speaker2"], ids
            )
            aligned.append((1.0 if row["label"] == 1 else -1.0) * delta)
    return reliability(np.asarray(aligned))


def load_ear_dataset() -> tuple[list[dict], list[dict]]:
    original_protocol = json.loads(EAR_ORIGINAL_PROTOCOL.read_text(encoding="utf-8"))
    declared = {
        int(row["subject"]): row
        for row in original_protocol["dataset"]["participants"]
    }
    geometry = json.loads(EAR_GEOMETRY.read_text(encoding="utf-8"))
    sectors = np.asarray(geometry["angular_sector_id_0_to_7"], dtype=np.int64)
    lag_index = np.repeat(np.arange(6), EAR_N_CHANNELS)
    ids = np.tile(sectors, 6) + 8 * (lag_index >= 3)
    if ids.shape != (EAR_N_FEATURES,) or np.any(np.bincount(ids, minlength=16) == 0):
        raise RuntimeError("Degenerate Ear-SAAD geometry.")
    inputs = [
        {"path": EAR_GEOMETRY.relative_to(ROOT).as_posix(), "sha256": sha256(EAR_GEOMETRY)},
        {
            "path": EAR_ORIGINAL_PROTOCOL.relative_to(ROOT).as_posix(),
            "sha256": sha256(EAR_ORIGINAL_PROTOCOL),
        },
    ]
    participants: list[dict] = []
    for subject_text in EAR_SUBJECTS:
        subject = int(subject_text[1:])
        input_path = EAR_ROOT / f"dataSubject{subject}.mat"
        observed_hash = sha256(input_path)
        if observed_hash != declared[subject]["sha256"]:
            raise RuntimeError(f"Ear-SAAD input hash mismatch: {subject_text}")
        inputs.append({"path": input_path.relative_to(ROOT).as_posix(), "sha256": observed_hash})
        data = loadmat(input_path, simplify_cells=True)
        channels = [str(value) for value in np.asarray(data["channels"]).ravel()]
        if channels[:EAR_N_CHANNELS] != geometry["labels_in_data_order"]:
            raise RuntimeError(f"Ear-SAAD channel mismatch: {subject_text}")
        eeg_trials = [
            np.nan_to_num(np.asarray(value, dtype=np.float64), nan=0.0, posinf=0.0, neginf=0.0)
            for value in np.asarray(data["eegTrials"], dtype=object).ravel()
        ]
        envelope_trials = [
            np.asarray(value, dtype=np.float64)
            for value in np.asarray(data["envelopes"], dtype=object).ravel()
        ]
        attended = np.asarray(data["attSpeaker"]).ravel().astype(np.int64)
        video = np.asarray(data["videoCondition"]).ravel().astype(np.int64)
        split = ear_split(attended, video)
        records = []
        for trial in range(6):
            for block in range(EAR_BLOCKS_PER_TRIAL):
                start = block * EAR_SAMPLES_PER_BLOCK
                stop = start + EAR_SAMPLES_PER_BLOCK
                design = ear_block_design(eeg_trials[trial][start:stop, :EAR_N_CHANNELS])
                envelopes = envelope_trials[trial][start:stop, :][:EAR_USABLE]
                target = envelopes[:, int(attended[trial]) - 1]
                records.append(
                    {
                        "trial": trial,
                        "block": block,
                        "design": design,
                        "speaker1": envelopes[:, 0],
                        "speaker2": envelopes[:, 1],
                        "label": 1 if attended[trial] == 1 else 0,
                        "video": int(video[trial]),
                        "stats": ear_sufficient_statistics(design, target),
                    }
                )
        profiles = {
            name: ear_profile(records, trials, ids)
            for name, trials in split.items()
        }
        directions = {}
        for source, target in (("A", "B"), ("B", "A")):
            training = [
                index for index, row in enumerate(records)
                if row["trial"] in split[source]
            ]
            testing = [
                index for index, row in enumerate(records)
                if row["trial"] in split[target]
            ]
            model = ear_fit_from_blocks(records, training)
            deltas, labels, metadata_rows = [], [], []
            for index in testing:
                row = records[index]
                delta, _ = ear_cell_contributions(
                    row["design"], model, row["speaker1"], row["speaker2"], ids
                )
                deltas.append(delta)
                labels.append(row["label"])
                metadata_rows.append(
                    {
                        "trial": int(row["trial"] + 1),
                        "block": int(row["block"] + 1),
                        "video": int(row["video"]),
                    }
                )
            directions[f"{source}_to_{target}"] = {
                "source": source,
                "target": target,
                "deltas": np.asarray(deltas),
                "labels": np.asarray(labels, dtype=np.int64),
                "metadata": metadata_rows,
            }
        participants.append(
            {
                "subject": subject_text,
                "profiles": profiles,
                "directions": directions,
                "split": {key: [value + 1 for value in values] for key, values in split.items()},
            }
        )
        print(f"Ear-SAAD {subject_text} complete", flush=True)
    return participants, inputs


def directional_endpoints(
    receiver_index: int,
    participants: list[dict],
    direction_name: str,
) -> dict[str, float | int | str]:
    receiver = participants[receiver_index]
    direction = receiver["directions"][direction_name]
    source = direction["source"]
    target = direction["target"]
    source_profile = receiver["profiles"][source]
    target_profile = receiver["profiles"][target]
    donor_profiles = np.row_stack(
        [
            participant["profiles"][source]
            for index, participant in enumerate(participants)
            if index != receiver_index
        ]
    )
    same_correlation = pearson(source_profile, target_profile)
    donor_correlations = np.asarray(
        [pearson(profile, target_profile) for profile in donor_profiles]
    )
    wrong_correlations = np.asarray(
        [
            pearson(transform_vector(source_profile, shift, swap), target_profile)
            for shift, swap in TRANSFORMS[1:]
        ]
    )
    fingerprint_listener_rank = rank_probability(same_correlation, donor_correlations)
    fingerprint_mapping_rank = rank_probability(same_correlation, wrong_correlations)

    delta = np.asarray(direction["deltas"], dtype=np.float64)
    labels = np.asarray(direction["labels"], dtype=np.int64)
    alignment = np.where(labels == 1, 1.0, -1.0)
    own_weights = family_weights(source_profile, "softmax2")
    donor_weights = np.row_stack(
        [family_weights(profile, "softmax2") for profile in donor_profiles]
    )
    wrong_weights = np.row_stack(
        [
            transform_vector(own_weights, shift, swap)
            for shift, swap in TRANSFORMS[1:]
        ]
    )
    own_scores = delta @ own_weights
    donor_scores = delta @ donor_weights.T
    wrong_scores = delta @ wrong_weights.T
    aligned_own = alignment * own_scores
    aligned_donor = alignment[:, None] * donor_scores
    aligned_wrong = alignment[:, None] * wrong_scores
    predictive_listener_rank = float(
        np.mean((aligned_own[:, None] > aligned_donor) + 0.5 * (aligned_own[:, None] == aligned_donor))
    )
    predictive_mapping_rank = float(
        np.mean((aligned_own[:, None] > aligned_wrong) + 0.5 * (aligned_own[:, None] == aligned_wrong))
    )
    donor_scale = np.sqrt(np.mean(np.column_stack([aligned_own, aligned_donor]) ** 2, axis=1))
    wrong_scale = np.sqrt(np.mean(np.column_stack([aligned_own, aligned_wrong]) ** 2, axis=1))
    donor_scale = np.maximum(donor_scale, 1e-12)
    wrong_scale = np.maximum(wrong_scale, 1e-12)
    return {
        "subject": receiver["subject"],
        "direction": direction_name,
        "source_partition": source,
        "target_partition": target,
        "target_observations": int(len(labels)),
        "same_profile_correlation": same_correlation,
        "mean_donor_profile_correlation": float(np.mean(donor_correlations)),
        "mean_wrong_mapping_correlation": float(np.mean(wrong_correlations)),
        "fingerprint_listener_rank_probability": fingerprint_listener_rank,
        "fingerprint_physical_mapping_rank_probability": fingerprint_mapping_rank,
        "predictive_listener_rank_probability": predictive_listener_rank,
        "predictive_physical_mapping_rank_probability": predictive_mapping_rank,
        "normalized_own_minus_donor_score": float(
            np.mean((aligned_own - np.mean(aligned_donor, axis=1)) / donor_scale)
        ),
        "normalized_physical_minus_wrong_score": float(
            np.mean((aligned_own - np.mean(aligned_wrong, axis=1)) / wrong_scale)
        ),
        "global_auc": binary_auc(np.sum(delta, axis=1), labels),
        "own_profile_auc": binary_auc(own_scores, labels),
    }


def participant_endpoints(
    dataset: str, participants: list[dict]
) -> tuple[list[dict], list[dict]]:
    direction_rows = []
    participant_rows = []
    for index, participant in enumerate(participants):
        rows = [
            directional_endpoints(index, participants, direction)
            for direction in ("A_to_B", "B_to_A")
        ]
        for row in rows:
            row["dataset"] = dataset
            direction_rows.append(row)
        participant_rows.append(
            {
                "dataset": dataset,
                "subject": participant["subject"],
                "F1_fingerprint_listener_rank_advantage": float(
                    np.mean([row["fingerprint_listener_rank_probability"] for row in rows]) - 0.5
                ),
                "F2_fingerprint_physical_mapping_rank_advantage": float(
                    np.mean([row["fingerprint_physical_mapping_rank_probability"] for row in rows]) - 0.5
                ),
                "P1_predictive_listener_rank_advantage": float(
                    np.mean([row["predictive_listener_rank_probability"] for row in rows]) - 0.5
                ),
                "P2_predictive_physical_mapping_rank_advantage": float(
                    np.mean([row["predictive_physical_mapping_rank_probability"] for row in rows]) - 0.5
                ),
                "same_minus_mean_donor_correlation": float(
                    np.mean(
                        [
                            row["same_profile_correlation"] - row["mean_donor_profile_correlation"]
                            for row in rows
                        ]
                    )
                ),
                "physical_minus_mean_wrong_correlation": float(
                    np.mean(
                        [
                            row["same_profile_correlation"] - row["mean_wrong_mapping_correlation"]
                            for row in rows
                        ]
                    )
                ),
                "normalized_own_minus_donor_score": float(
                    np.mean([row["normalized_own_minus_donor_score"] for row in rows])
                ),
                "normalized_physical_minus_wrong_score": float(
                    np.mean([row["normalized_physical_minus_wrong_score"] for row in rows])
                ),
                "global_auc": float(np.mean([row["global_auc"] for row in rows])),
                "own_profile_auc": float(np.mean([row["own_profile_auc"] for row in rows])),
            }
        )
    return participant_rows, direction_rows


def exact_sign_flip(values: np.ndarray) -> float:
    array = np.asarray(values, dtype=np.float64)
    observed = float(np.mean(array))
    count = 0
    total = 1 << len(array)
    for mask in range(total):
        signs = np.asarray(
            [1.0 if (mask >> index) & 1 else -1.0 for index in range(len(array))]
        )
        if float(np.mean(signs * array)) >= observed - 1e-15:
            count += 1
    return count / total


def dataset_summary(
    values: np.ndarray, seed: int
) -> dict[str, float | int | list[float]]:
    array = np.asarray(values, dtype=np.float64)
    rng = np.random.default_rng(seed)
    sampled = array[rng.integers(0, len(array), size=(BOOTSTRAP_DRAWS, len(array)))]
    boot = np.mean(sampled, axis=1)
    return {
        "n": int(len(array)),
        "mean": float(np.mean(array)),
        "bootstrap_95": [
            float(np.quantile(boot, 0.025)),
            float(np.quantile(boot, 0.975)),
        ],
        "exact_one_sided_signflip_p": exact_sign_flip(array),
        "positive_participants": int(np.sum(array > 0)),
        "zero_participants": int(np.sum(array == 0)),
    }


def cross_dataset_summary(
    arrays: list[np.ndarray], bootstrap_seed: int, randomization_seed: int
) -> dict[str, float | int | list[float]]:
    observed = float(np.mean([np.mean(values) for values in arrays]))
    rng = np.random.default_rng(bootstrap_seed)
    boot_means = []
    for values in arrays:
        indices = rng.integers(0, len(values), size=(BOOTSTRAP_DRAWS, len(values)))
        boot_means.append(np.mean(values[indices], axis=1))
    boot = np.mean(np.row_stack(boot_means), axis=0)
    rng = np.random.default_rng(randomization_seed)
    extreme = 0
    remaining = RANDOMIZATION_DRAWS
    chunk = 10_000
    weights = [1.0 / (len(arrays) * len(values)) for values in arrays]
    while remaining:
        current = min(chunk, remaining)
        null = np.zeros(current, dtype=np.float64)
        for values, weight in zip(arrays, weights):
            signs = rng.integers(0, 2, size=(current, len(values)), dtype=np.int8)
            signs = signs.astype(np.float64) * 2.0 - 1.0
            null += weight * (signs @ values)
        extreme += int(np.sum(null >= observed - 1e-15))
        remaining -= current
    return {
        "dataset_count": len(arrays),
        "participant_count": int(sum(len(values) for values in arrays)),
        "equal_dataset_mean": observed,
        "stratified_bootstrap_95": [
            float(np.quantile(boot, 0.025)),
            float(np.quantile(boot, 0.975)),
        ],
        "participant_signflip_randomization_p": float(
            (1 + extreme) / (1 + RANDOMIZATION_DRAWS)
        ),
        "bootstrap_draws": BOOTSTRAP_DRAWS,
        "randomization_draws": RANDOMIZATION_DRAWS,
    }


def write_csv_exclusive(path: Path, rows: list[dict]) -> None:
    if not rows:
        raise ValueError(f"No rows for {path.name}.")
    with path.open("x", encoding="utf-8", newline="") as handle:
        writer = csv.DictWriter(handle, fieldnames=list(rows[0]))
        writer.writeheader()
        writer.writerows(rows)


def unique_inputs(records: Iterable[dict]) -> list[dict]:
    output = {}
    for record in records:
        prior = output.get(record["path"])
        if prior is not None and prior != record["sha256"]:
            raise RuntimeError(f"Conflicting input hash: {record['path']}")
        output[record["path"]] = record["sha256"]
    return [{"path": path, "sha256": digest} for path, digest in sorted(output.items())]


def main() -> None:
    if DECISION.exists() or PARTICIPANT_CSV.exists() or DIRECTION_CSV.exists():
        raise RuntimeError("Refusing to overwrite continuous-listener transport output.")
    lock, lock_hash = validate_lock()
    datasets = {}
    all_inputs = []
    for dataset, loader in (
        ("KITE_ST_confirmation_complete_case_v2", load_kite_dataset),
        ("KUL_64ch", load_kul_dataset),
        ("Ear_SAAD_29ch", load_ear_dataset),
    ):
        participants, inputs = loader()
        datasets[dataset] = participants
        all_inputs.extend(inputs)

    participant_rows, direction_rows = [], []
    for dataset, participants in datasets.items():
        rows, directions = participant_endpoints(dataset, participants)
        participant_rows.extend(rows)
        direction_rows.extend(directions)

    expected_counts = {
        "KITE_ST_confirmation_complete_case_v2": 12,
        "KUL_64ch": 16,
        "Ear_SAAD_29ch": 15,
    }
    integrity_checks = {
        "dataset_counts_exact": all(
            len(datasets[name]) == count for name, count in expected_counts.items()
        ),
        "participant_rows_exact": len(participant_rows) == 43,
        "direction_rows_exact": len(direction_rows) == 86,
        "all_profiles_16_finite": all(
            profile.shape == (16,) and np.isfinite(profile).all()
            for participants in datasets.values()
            for participant in participants
            for profile in participant["profiles"].values()
        ),
        "all_target_contributions_16_finite": all(
            direction["deltas"].ndim == 2
            and direction["deltas"].shape[1] == 16
            and np.isfinite(direction["deltas"]).all()
            for participants in datasets.values()
            for participant in participants
            for direction in participant["directions"].values()
        ),
        "two_directions_per_participant": all(
            set(participant["directions"]) == {"A_to_B", "B_to_A"}
            for participants in datasets.values()
            for participant in participants
        ),
        "ear_splits_disjoint_and_complete": all(
            set(participant["split"]["A"]).isdisjoint(participant["split"]["B"])
            and set(participant["split"]["A"]) | set(participant["split"]["B"])
            == set(range(1, 7))
            for participant in datasets["Ear_SAAD_29ch"]
        ),
        "paper_mutated": False,
    }
    c0 = all(value is True for key, value in integrity_checks.items() if key != "paper_mutated")
    summaries = {}
    cross_summaries = {}
    dataset_names = list(datasets)
    for endpoint_index, endpoint in enumerate(ENDPOINTS):
        summaries[endpoint] = {}
        arrays = []
        for dataset_index, dataset in enumerate(dataset_names):
            values = np.asarray(
                [
                    row[endpoint]
                    for row in participant_rows
                    if row["dataset"] == dataset
                ],
                dtype=np.float64,
            )
            arrays.append(values)
            summaries[endpoint][dataset] = dataset_summary(
                values, BOOTSTRAP_SEED + 100 * endpoint_index + dataset_index
            )
        cross = cross_dataset_summary(
            arrays,
            BOOTSTRAP_SEED + 10_000 * endpoint_index,
            RANDOMIZATION_SEED + 10_000 * endpoint_index,
        )
        dataset_summaries = summaries[endpoint]
        cross["positive_dataset_means"] = int(
            sum(dataset_summaries[name]["mean"] > 0 for name in dataset_names)
        )
        cross["datasets_with_bootstrap_upper_below_zero"] = [
            name
            for name in dataset_names
            if dataset_summaries[name]["bootstrap_95"][1] < 0
        ]
        cross_summaries[endpoint] = cross

    def endpoint_gate(endpoint: str) -> dict:
        row = cross_summaries[endpoint]
        checks = {
            "equal_dataset_mean_gt_zero": row["equal_dataset_mean"] > 0,
            "bootstrap_lower_gt_zero": row["stratified_bootstrap_95"][0] > 0,
            "randomization_p_le_0_05": row["participant_signflip_randomization_p"] <= 0.05,
            "at_least_two_positive_dataset_means": row["positive_dataset_means"] >= 2,
            "no_dataset_bootstrap_upper_below_zero": not row[
                "datasets_with_bootstrap_upper_below_zero"
            ],
        }
        return {"pass": all(checks.values()), "checks": checks}

    gates = {
        "C0_integrity": {"pass": c0, "checks": integrity_checks},
        "C1_fingerprint_listener": endpoint_gate(ENDPOINTS[0]),
        "C2_fingerprint_mapping": endpoint_gate(ENDPOINTS[1]),
        "C3_predictive_listener": endpoint_gate(ENDPOINTS[2]),
        "C4_predictive_mapping": endpoint_gate(ENDPOINTS[3]),
    }
    conjunction = all(row["pass"] for row in gates.values())
    output = {
        "schema": "WZ-HIC-continuous-listener-transport-decision-v1",
        "status": (
            "continuous_listener_transport_supported"
            if conjunction
            else "continuous_listener_transport_conjunction_not_supported"
        ),
        "protocol": {
            "path": PROTOCOL.relative_to(ROOT).as_posix(),
            "sha256": PROTOCOL_SHA256,
        },
        "implementation_lock": {
            "path": LOCK.relative_to(ROOT).as_posix(),
            "sha256": lock_hash,
        },
        "runner": {
            "path": Path(__file__).relative_to(ROOT).as_posix(),
            "sha256": sha256(Path(__file__)),
        },
        "frozen_inputs": unique_inputs(all_inputs),
        "dataset_counts": expected_counts,
        "participant_count": len(participant_rows),
        "direction_count": len(direction_rows),
        "endpoint_definitions": json.loads(PROTOCOL.read_text(encoding="utf-8"))[
            "participant_endpoints"
        ],
        "dataset_summaries": summaries,
        "cross_dataset_summaries": cross_summaries,
        "gates": gates,
        "conjunction_pass": conjunction,
        "ear_splits": {
            participant["subject"]: participant["split"]
            for participant in datasets["Ear_SAAD_29ch"]
        },
        "artifacts": {
            "participant_csv": PARTICIPANT_CSV.relative_to(ROOT).as_posix(),
            "direction_csv": DIRECTION_CSV.relative_to(ROOT).as_posix(),
        },
        "claim_boundary": {
            "supported_if_conjunction_passes": (
                "A listener-matched continuous-speech direction-by-delay family recurs "
                "across nonoverlapping material and improves held-partition evidence "
                "relative to other listeners and capacity-matched wrong mappings."
            ),
            "never_supported_by_this_experiment": [
                "theorem-to-neuroscience deduction",
                "speaker-identity decoding",
                "generic donor ridge-model transport",
                "universal direction family across tasks",
            ],
        },
        "paper_mutated": False,
    }
    OUTPUT_ROOT.mkdir(parents=True, exist_ok=True)
    write_csv_exclusive(PARTICIPANT_CSV, participant_rows)
    write_csv_exclusive(DIRECTION_CSV, direction_rows)
    with DECISION.open("x", encoding="utf-8") as handle:
        json.dump(output, handle, ensure_ascii=False, indent=2, allow_nan=False)
        handle.write("\n")
    for path in (PARTICIPANT_CSV, DIRECTION_CSV, DECISION):
        sidecar = path.with_name(path.name + ".sha256")
        with sidecar.open("x", encoding="ascii") as handle:
            handle.write(f"{sha256(path)}  {path.name}\n")
    print(json.dumps({"status": output["status"], "gates": gates}, indent=2), flush=True)


if __name__ == "__main__":
    main()
```
