## Supplementary material for "Falsifiable substitution tests reveal task-structured neural evidence for auditory attention": Article_Analysis_Source_Code: run_wz_hic_continuous_listener_transport_v2.py

```
#!/usr/bin/env python3
"""Metadata-path correction for the frozen continuous-listener v1 runner.

The v1 execution stopped before any profile or endpoint was computed because
the complete-case subject JSON uses ``upstream_manifest`` rather than the
adapter name ``preprocessing_manifest``.  This wrapper replaces only the
KITE-ST loader and the implementation-lock validator; all scientific
functions, endpoints, thresholds, seeds, and output serialization remain the
locked v1 implementation.
"""

from __future__ import annotations

import json
from pathlib import Path

import numpy as np
from scipy.signal import resample_poly

import run_wz_hic_continuous_listener_transport_v1 as v1


LOCK_V3 = (
    v1.ROOT / "docs"
    / "wz_hic_continuous_listener_transport_implementation_lock_v3.json"
)


def validate_lock_v3() -> tuple[dict, str]:
    if v1.sha256(v1.PROTOCOL) != v1.PROTOCOL_SHA256:
        raise RuntimeError("Continuous-listener protocol hash mismatch.")
    if not LOCK_V3.exists():
        raise RuntimeError("Public-reproduction implementation lock is missing.")
    payload = json.loads(LOCK_V3.read_text(encoding="utf-8"))
    lock_hash = v1.sha256(LOCK_V3)
    wrapper_path = Path(__file__).resolve()
    if (
        payload["status"]
        != "outcome_aware_public_reproduction_repair"
        or payload["protocol_sha256"] != v1.PROTOCOL_SHA256
        or payload["runner"]["path"] != wrapper_path.relative_to(v1.ROOT).as_posix()
        or payload["runner"]["sha256"] != v1.sha256(wrapper_path)
    ):
        raise RuntimeError("Corrected continuous-listener lock mismatch.")
    for record in payload["immutable_dependencies"]:
        if v1.sha256(v1.ROOT / record["path"]) != record["sha256"]:
            raise RuntimeError(f"Dependency hash mismatch: {record['path']}")
    return payload, lock_hash


def load_kite_dataset_corrected() -> tuple[list[dict], list[dict]]:
    """Byte-equivalent v1 loader except for the corrected manifest key."""
    with np.load(v1.KITE_AUDIO, allow_pickle=False) as saved:
        audio = {name: np.asarray(saved[name]) for name in saved.files}
    geometry = v1.load_public_geometry(v1.KITE_COORDINATES)
    ids = v1.partition_ids(geometry.xy)["S16"]
    participants: list[dict] = []
    inputs: list[dict] = [
        {
            "path": v1.KITE_AUDIO.relative_to(v1.ROOT).as_posix(),
            "sha256": v1.sha256(v1.KITE_AUDIO),
        },
        {
            "path": v1.KITE_COORDINATES.relative_to(v1.ROOT).as_posix(),
            "sha256": v1.sha256(v1.KITE_COORDINATES),
        },
    ]
    for subject in v1.KITE_SUBJECTS:
        result_path = v1.KITE_RESULTS / f"{subject}_AKB_confirmation_v1.json"
        model_path = v1.KITE_MODELS / f"{subject}_AKB_confirmation_models_v1.npz"
        result = json.loads(result_path.read_text(encoding="utf-8"))
        manifest_path = v1.ROOT / result["upstream_manifest"]["path"]
        if v1.sha256(manifest_path) != result["upstream_manifest"]["sha256"]:
            raise RuntimeError(f"KITE upstream manifest hash mismatch: {subject}")
        manifest = json.loads(manifest_path.read_text(encoding="utf-8"))
        preprocessing = {
            row["recording"].rsplit("_", 1)[-1]: row
            for row in manifest["preprocessing_records"]
        }
        inputs.extend(
            [
                {
                    "path": result_path.relative_to(v1.ROOT).as_posix(),
                    "sha256": v1.sha256(result_path),
                },
                {
                    "path": model_path.relative_to(v1.ROOT).as_posix(),
                    "sha256": v1.sha256(model_path),
                },
                {
                    "path": manifest_path.relative_to(v1.ROOT).as_posix(),
                    "sha256": v1.sha256(manifest_path),
                },
            ]
        )
        condition_blocks: dict[str, list[dict]] = {}
        for condition in ("1L", "1R", "2L", "2R"):
            input_path = (
                v1.KITE_PREPROCESSED / subject
                / f"{subject}_AAD_{condition}_preprocessed_v1.npz"
            )
            observed_hash = v1.sha256(input_path)
            if observed_hash != preprocessing[condition]["output"]["sha256"]:
                raise RuntimeError(f"KITE input hash mismatch: {subject} {condition}")
            inputs.append(
                {
                    "path": input_path.relative_to(v1.ROOT).as_posix(),
                    "sha256": observed_hash,
                }
            )
            with np.load(input_path, allow_pickle=False) as saved:
                eeg = np.asarray(saved["eeg"])
            offset = int(round(v1.kite_offset(preprocessing[condition]) * 100))
            left_name, right_name = v1.KITE_PHYSICAL[condition]
            rows = []
            for block_index in range(12):
                first = offset + block_index * 3000
                stop = first + 3000
                left100 = audio[left_name][first:stop]
                right100 = audio[right_name][first:stop]
                design, left_features = v1.kite_block_design(
                    eeg[block_index * 3000 : (block_index + 1) * 3000],
                    left100,
                )
                right_features = resample_poly(right100, 1, 5, axis=0)[:595]
                rows.append(
                    {
                        "design": design,
                        "left": left_features[:, 0],
                        "right": right_features[:, 0],
                        "label": 1 if condition.endswith("L") else 0,
                        "condition": condition,
                        "block": block_index + 1,
                    }
                )
            condition_blocks[condition] = rows
        with np.load(model_path, allow_pickle=False) as saved:
            profiles = {
                "A": np.asarray(
                    result["fold_reliabilities"]["train_story_1"],
                    dtype=np.float64,
                ),
                "B": np.asarray(
                    result["fold_reliabilities"]["train_story_2"],
                    dtype=np.float64,
                ),
            }
            models = {
                "A": v1.model_from_archive(saved, "train_story_1"),
                "B": v1.model_from_archive(saved, "train_story_2"),
            }
        directions = {}
        for source, target_conditions in (
            ("A", ("2L", "2R")),
            ("B", ("1L", "1R")),
        ):
            rows = [
                row
                for condition in target_conditions
                for row in condition_blocks[condition]
            ]
            deltas, labels, metadata = [], [], []
            for row in rows:
                delta, _ = v1.cell_delta_contributions(
                    row["design"],
                    models[source],
                    row["left"],
                    row["right"],
                    ids,
                )
                deltas.append(delta)
                labels.append(row["label"])
                metadata.append(
                    {"condition": row["condition"], "block": row["block"]}
                )
            target = "B" if source == "A" else "A"
            directions[f"{source}_to_{target}"] = {
                "source": source,
                "target": target,
                "deltas": np.asarray(deltas),
                "labels": np.asarray(labels, dtype=np.int64),
                "metadata": metadata,
            }
        participants.append(
            {"subject": subject, "profiles": profiles, "directions": directions}
        )
        print(f"KITE-ST {subject} complete", flush=True)
    return participants, inputs


def main() -> None:
    v1.LOCK = LOCK_V3
    v1.validate_lock = validate_lock_v3
    v1.load_kite_dataset = load_kite_dataset_corrected
    v1.__file__ = str(Path(__file__).resolve())
    v1.main()


if __name__ == "__main__":
    main()
```
