## Supplementary material for "Falsifiable substitution tests reveal task-structured neural evidence for auditory attention": Article_Analysis_Source_Code: run_wz_hic_continuous_shared_template_v1.py

```
#!/usr/bin/env python3
"""Frozen leave-one-listener shared-template comparison."""

from __future__ import annotations

import csv
import hashlib
import itertools
import json
from pathlib import Path

import numpy as np

from audit_wz_hic_continuous_listener_transport_source_v1 import (
    auc,
    load_ear,
    load_kite,
    load_kul,
)
from kite_sat_adaptive_broad_v1 import family_weights


ROOT = Path(__file__).resolve().parents[1]
PROTOCOL = ROOT / "docs" / "wz_hic_continuous_shared_template_protocol_v1.json"
PROTOCOL_SHA256 = "3087f17acea6b7a9fcb1fc5c1c92d715aeabb959c61f04abb17c75ef91deb22e"
LOCK = ROOT / "docs" / "wz_hic_continuous_shared_template_implementation_lock_v2.json"
OUTPUT_ROOT = (
    ROOT / "results" / "wz_hic_cross_dataset"
    / "continuous_shared_template_v1"
)
DECISION = OUTPUT_ROOT / "continuous_shared_template_decision_v1.json"
PARTICIPANT_CSV = OUTPUT_ROOT / "participant_endpoints_v1.csv"
DIRECTION_CSV = OUTPUT_ROOT / "direction_endpoints_v1.csv"
DATASETS = (
    "KITE_ST_confirmation_complete_case_v2",
    "KUL_64ch",
    "Ear_SAAD_29ch",
)
ENDPOINTS = (
    "G1_own_vs_mean_profile_rank_advantage",
    "G2_own_vs_mean_weight_rank_advantage",
)
BOOTSTRAP_DRAWS = 100_000
RANDOMIZATION_DRAWS = 1_000_000
BOOTSTRAP_SEED = 2026073011
RANDOMIZATION_SEED = 2026073012


def sha256(path: Path) -> str:
    digest = hashlib.sha256()
    with path.open("rb") as handle:
        for block in iter(lambda: handle.read(8 * 1024 * 1024), b""):
            digest.update(block)
    return digest.hexdigest()


def semantic_json_sha256(path: Path, excluded_paths: set[str]) -> str:
    """Hash scientific JSON content with explicitly listed provenance removed."""

    def clean(value, prefix=""):
        if isinstance(value, dict):
            output = {}
            for key, item in value.items():
                current = f"{prefix}/{key}" if prefix else key
                if current in excluded_paths:
                    continue
                output[key] = clean(item, current)
            return output
        if isinstance(value, list):
            return [
                clean(item, f"{prefix}/{index}")
                for index, item in enumerate(value)
            ]
        return value

    payload = json.loads(path.read_text(encoding="utf-8"))
    canonical = json.dumps(
        clean(payload), sort_keys=True, separators=(",", ":"), allow_nan=True
    ).encode("utf-8")
    return hashlib.sha256(canonical).hexdigest()


def validate_lock() -> tuple[dict, str]:
    if sha256(PROTOCOL) != PROTOCOL_SHA256:
        raise RuntimeError("Shared-template protocol hash mismatch.")
    if not LOCK.exists():
        raise RuntimeError("Shared-template implementation lock missing.")
    lock = json.loads(LOCK.read_text(encoding="utf-8"))
    lock_hash = sha256(LOCK)
    if (
        lock["status"] != "outcome_aware_public_reproduction_repair"
        or lock["protocol_sha256"] != PROTOCOL_SHA256
        or lock["runner"]["path"] != Path(__file__).relative_to(ROOT).as_posix()
        or lock["runner"]["sha256"] != sha256(Path(__file__))
    ):
        raise RuntimeError("Shared-template implementation lock mismatch.")
    for record in lock["byte_dependencies"]:
        if sha256(ROOT / record["path"]) != record["sha256"]:
            raise RuntimeError(f"Dependency mismatch: {record['path']}")
    for record in lock["semantic_json_dependencies"]:
        observed = semantic_json_sha256(
            ROOT / record["path"], set(record["excluded_paths"])
        )
        if observed != record["sha256"]:
            raise RuntimeError(
                f"Scientific dependency mismatch: {record['path']}"
            )
    return lock, lock_hash


def exact_sign_flip(values: np.ndarray) -> float:
    values = np.asarray(values, dtype=np.float64)
    observed = float(np.mean(values))
    return float(
        np.mean(
            [
                np.mean(values * np.asarray(signs)) >= observed - 1e-15
                for signs in itertools.product((-1.0, 1.0), repeat=len(values))
            ]
        )
    )


def dataset_summary(values: np.ndarray, seed: int) -> dict:
    values = np.asarray(values, dtype=np.float64)
    rng = np.random.default_rng(seed)
    samples = values[
        rng.integers(0, len(values), size=(BOOTSTRAP_DRAWS, len(values)))
    ]
    boot = np.mean(samples, axis=1)
    return {
        "n": len(values),
        "mean": float(np.mean(values)),
        "bootstrap_95": [
            float(np.quantile(boot, 0.025)),
            float(np.quantile(boot, 0.975)),
        ],
        "exact_one_sided_signflip_p": exact_sign_flip(values),
        "positive_participants": int(np.sum(values > 0)),
        "zero_participants": int(np.sum(values == 0)),
    }


def cross_summary(
    arrays: list[np.ndarray], bootstrap_seed: int, randomization_seed: int
) -> dict:
    observed = float(np.mean([np.mean(values) for values in arrays]))
    rng = np.random.default_rng(bootstrap_seed)
    boot = []
    for values in arrays:
        indices = rng.integers(
            0, len(values), size=(BOOTSTRAP_DRAWS, len(values))
        )
        boot.append(np.mean(values[indices], axis=1))
    bootstrap = np.mean(np.vstack(boot), axis=0)
    rng = np.random.default_rng(randomization_seed)
    extreme = 0
    remaining = RANDOMIZATION_DRAWS
    while remaining:
        current = min(10_000, remaining)
        null = np.zeros(current, dtype=np.float64)
        for values in arrays:
            signs = rng.integers(
                0, 2, size=(current, len(values)), dtype=np.int8
            )
            signs = signs.astype(np.float64) * 2.0 - 1.0
            null += signs @ values / (len(arrays) * len(values))
        extreme += int(np.sum(null >= observed - 1e-15))
        remaining -= current
    return {
        "dataset_count": 3,
        "participant_count": int(sum(len(values) for values in arrays)),
        "equal_dataset_mean": observed,
        "stratified_bootstrap_95": [
            float(np.quantile(bootstrap, 0.025)),
            float(np.quantile(bootstrap, 0.975)),
        ],
        "participant_signflip_randomization_p": float(
            (1 + extreme) / (1 + RANDOMIZATION_DRAWS)
        ),
        "bootstrap_draws": BOOTSTRAP_DRAWS,
        "randomization_draws": RANDOMIZATION_DRAWS,
    }


def direction_endpoint(index: int, people: list[dict], direction: str) -> dict:
    receiver = people[index]
    test = receiver["tests"][direction]
    source = test["source"]
    own_profile = np.asarray(receiver["profiles"][source], dtype=np.float64)
    donor_profiles = np.asarray(
        [
            person["profiles"][source]
            for donor_index, person in enumerate(people)
            if donor_index != index
        ],
        dtype=np.float64,
    )
    if len(donor_profiles) != len(people) - 1:
        raise RuntimeError("Receiver exclusion failed.")
    own_weights = family_weights(own_profile, "softmax2")
    mean_profile = np.mean(donor_profiles, axis=0)
    template_profile_weights = family_weights(mean_profile, "softmax2")
    donor_weights = np.asarray(
        [family_weights(profile, "softmax2") for profile in donor_profiles]
    )
    template_mean_weights = np.mean(donor_weights, axis=0)
    delta = np.asarray(test["delta"], dtype=np.float64)
    labels = np.asarray(test["labels"], dtype=np.int64)
    sign = np.where(labels == 1, 1.0, -1.0)
    own_score = delta @ own_weights
    profile_score = delta @ template_profile_weights
    weight_score = delta @ template_mean_weights
    aligned_own = sign * own_score
    aligned_profile = sign * profile_score
    aligned_weight = sign * weight_score

    def comparison(template: np.ndarray) -> tuple[float, float]:
        rank = float(
            np.mean(
                (aligned_own > template)
                + 0.5 * (aligned_own == template)
            )
        )
        scale = np.maximum(
            np.sqrt(np.mean(np.column_stack((aligned_own, template)) ** 2, axis=1)),
            1e-12,
        )
        normalized = float(np.mean((aligned_own - template) / scale))
        return rank, normalized

    profile_rank, profile_normalized = comparison(aligned_profile)
    weight_rank, weight_normalized = comparison(aligned_weight)
    return {
        "subject": receiver["subject"],
        "direction": direction,
        "source_partition": source,
        "target_partition": test["target"],
        "donor_count": len(donor_profiles),
        "target_observations": len(labels),
        "own_vs_mean_profile_rank_probability": profile_rank,
        "own_vs_mean_weight_rank_probability": weight_rank,
        "own_minus_mean_profile_auc": (
            auc(own_score, labels) - auc(profile_score, labels)
        ),
        "own_minus_mean_weight_auc": (
            auc(own_score, labels) - auc(weight_score, labels)
        ),
        "normalized_own_minus_mean_profile_score": profile_normalized,
        "normalized_own_minus_mean_weight_score": weight_normalized,
    }


def write_csv(path: Path, records: list[dict]) -> None:
    with path.open("x", encoding="utf-8", newline="") as handle:
        writer = csv.DictWriter(handle, fieldnames=list(records[0]))
        writer.writeheader()
        writer.writerows(records)


def main() -> None:
    if DECISION.exists() or PARTICIPANT_CSV.exists() or DIRECTION_CSV.exists():
        raise RuntimeError("Refusing to overwrite shared-template result.")
    lock, lock_hash = validate_lock()
    loaded = {
        dataset: loader()
        for dataset, loader in (
            (DATASETS[0], load_kite),
            (DATASETS[1], load_kul),
            (DATASETS[2], load_ear),
        )
    }
    direction_rows = []
    participant_rows = []
    for dataset, people in loaded.items():
        for index, person in enumerate(people):
            directions = [
                direction_endpoint(index, people, direction)
                for direction in ("A_to_B", "B_to_A")
            ]
            for row in directions:
                row["dataset"] = dataset
                direction_rows.append(row)
            participant_rows.append(
                {
                    "dataset": dataset,
                    "subject": person["subject"],
                    "G1_own_vs_mean_profile_rank_advantage": float(
                        np.mean(
                            [
                                row["own_vs_mean_profile_rank_probability"]
                                for row in directions
                            ]
                        )
                        - 0.5
                    ),
                    "G2_own_vs_mean_weight_rank_advantage": float(
                        np.mean(
                            [
                                row["own_vs_mean_weight_rank_probability"]
                                for row in directions
                            ]
                        )
                        - 0.5
                    ),
                    "own_minus_mean_profile_auc": float(
                        np.mean(
                            [
                                row["own_minus_mean_profile_auc"]
                                for row in directions
                            ]
                        )
                    ),
                    "own_minus_mean_weight_auc": float(
                        np.mean(
                            [
                                row["own_minus_mean_weight_auc"]
                                for row in directions
                            ]
                        )
                    ),
                    "normalized_own_minus_mean_profile_score": float(
                        np.mean(
                            [
                                row[
                                    "normalized_own_minus_mean_profile_score"
                                ]
                                for row in directions
                            ]
                        )
                    ),
                    "normalized_own_minus_mean_weight_score": float(
                        np.mean(
                            [
                                row[
                                    "normalized_own_minus_mean_weight_score"
                                ]
                                for row in directions
                            ]
                        )
                    ),
                }
            )
    integrity = {
        "participant_rows_43": len(participant_rows) == 43,
        "direction_rows_86": len(direction_rows) == 86,
        "donor_counts_exact": all(
            int(row["donor_count"])
            == (
                11
                if row["dataset"] == DATASETS[0]
                else 15
                if row["dataset"] == DATASETS[1]
                else 14
            )
            for row in direction_rows
        ),
        "all_finite": all(
            np.isfinite(float(value))
            for row in participant_rows
            for key, value in row.items()
            if key not in ("dataset", "subject")
        ),
        "paper_mutated": False,
    }
    summaries = {}
    crosses = {}
    for endpoint_index, endpoint in enumerate(ENDPOINTS):
        summaries[endpoint] = {}
        arrays = []
        for dataset_index, dataset in enumerate(DATASETS):
            values = np.asarray(
                [
                    row[endpoint]
                    for row in participant_rows
                    if row["dataset"] == dataset
                ]
            )
            arrays.append(values)
            summaries[endpoint][dataset] = dataset_summary(
                values, BOOTSTRAP_SEED + 100 * endpoint_index + dataset_index
            )
        cross = cross_summary(
            arrays,
            BOOTSTRAP_SEED + 10_000 * endpoint_index,
            RANDOMIZATION_SEED + 10_000 * endpoint_index,
        )
        cross["positive_dataset_means"] = int(
            sum(summaries[endpoint][dataset]["mean"] > 0 for dataset in DATASETS)
        )
        cross["datasets_with_bootstrap_upper_below_zero"] = [
            dataset
            for dataset in DATASETS
            if summaries[endpoint][dataset]["bootstrap_95"][1] < 0
        ]
        crosses[endpoint] = cross

    def endpoint_gate(endpoint: str) -> dict:
        result = crosses[endpoint]
        checks = {
            "equal_dataset_mean_gt_zero": result["equal_dataset_mean"] > 0,
            "bootstrap_lower_gt_zero": result["stratified_bootstrap_95"][0] > 0,
            "randomization_p_le_0_05": (
                result["participant_signflip_randomization_p"] <= 0.05
            ),
            "at_least_two_positive_dataset_means": (
                result["positive_dataset_means"] >= 2
            ),
            "no_dataset_bootstrap_upper_below_zero": not result[
                "datasets_with_bootstrap_upper_below_zero"
            ],
        }
        return {"pass": all(checks.values()), "checks": checks}

    gates = {
        "S0_integrity": {
            "pass": all(
                value is True
                for key, value in integrity.items()
                if key != "paper_mutated"
            ),
            "checks": integrity,
        },
        "S1_mean_profile": endpoint_gate(ENDPOINTS[0]),
        "S2_mean_weight": endpoint_gate(ENDPOINTS[1]),
    }
    conjunction = all(gate["pass"] for gate in gates.values())
    output = {
        "schema": "WZ-HIC-continuous-shared-template-decision-v1",
        "status": (
            "individual_benefit_over_shared_templates_supported"
            if conjunction
            else "individual_benefit_over_shared_templates_not_supported"
        ),
        "protocol": {
            "path": PROTOCOL.relative_to(ROOT).as_posix(),
            "sha256": PROTOCOL_SHA256,
        },
        "implementation_lock": {
            "path": LOCK.relative_to(ROOT).as_posix(),
            "sha256": lock_hash,
        },
        "runner": {
            "path": Path(__file__).relative_to(ROOT).as_posix(),
            "sha256": sha256(Path(__file__)),
        },
        "source_loader": {
            "path": (
                "analysis/"
                "audit_wz_hic_continuous_listener_transport_source_v1.py"
            ),
            "sha256": sha256(
                ROOT
                / "analysis"
                / "audit_wz_hic_continuous_listener_transport_source_v1.py"
            ),
            "prior_source_audit_status": "pass",
        },
        "dataset_counts": {
            dataset: len(people) for dataset, people in loaded.items()
        },
        "participant_count": len(participant_rows),
        "direction_count": len(direction_rows),
        "dataset_summaries": summaries,
        "cross_dataset_summaries": crosses,
        "gates": gates,
        "conjunction_pass": conjunction,
        "artifacts": {
            "participant_csv": PARTICIPANT_CSV.relative_to(ROOT).as_posix(),
            "direction_csv": DIRECTION_CSV.relative_to(ROOT).as_posix(),
        },
        "paper_mutated": False,
    }
    OUTPUT_ROOT.mkdir(parents=True, exist_ok=True)
    write_csv(PARTICIPANT_CSV, participant_rows)
    write_csv(DIRECTION_CSV, direction_rows)
    with DECISION.open("x", encoding="utf-8") as handle:
        json.dump(output, handle, ensure_ascii=False, indent=2, allow_nan=False)
        handle.write("\n")
    for path in (PARTICIPANT_CSV, DIRECTION_CSV, DECISION):
        with path.with_name(path.name + ".sha256").open(
            "x", encoding="ascii"
        ) as handle:
            handle.write(f"{sha256(path)}  {path.name}\n")
    print(json.dumps({"status": output["status"], "gates": gates}, indent=2))


if __name__ == "__main__":
    main()
```
