## Supplementary material for "Falsifiable substitution tests reveal task-structured neural evidence for auditory attention": Article_Analysis_Source_Code: run_wz_hic_cross_dataset_command_identity_residual_v1.py

```
#!/usr/bin/env python3
"""Frozen cross-dataset command-identity residual control for WZ-HIC."""

from __future__ import annotations

import hashlib
import itertools
import json
from datetime import datetime, timezone
from pathlib import Path

import numpy as np

from wz_hif_core_v1 import fit_shrinkage_lda


ROOT = Path(__file__).resolve().parents[1]
PROTOCOL = (
    ROOT
    / "docs"
    / "wz_hic_cross_dataset_command_identity_residual_protocol_v1.json"
)
LOCK = (
    ROOT
    / "docs"
    / "wz_hic_cross_dataset_command_identity_residual_implementation_lock_v1.json"
)
ASME_FEATURES = (
    ROOT
    / "data"
    / "derived"
    / "asme30_wz_hic_dataset_native_features_v2.npz"
)
KOJIMA_FEATURES = (
    ROOT
    / "data"
    / "derived"
    / "kojima2024b_wz_hic_confirmation_features_v1.npz"
)
ASME_RAW_RESULT = (
    ROOT
    / "results"
    / "wz_hic_asme30"
    / "crossphase_individual_geometry_v1"
    / "crossphase_decision_v1.json"
)
KOJIMA_RAW_RESULT = (
    ROOT
    / "results"
    / "wz_hic_kojima2024b"
    / "crossrun_individual_geometry_v1"
    / "crossrun_decision_v1.json"
)
OUTDIR = (
    ROOT
    / "results"
    / "wz_hic_cross_dataset"
    / "command_identity_residual_v1"
)
RESULT = OUTDIR / "residual_decision_v1.json"

PROTOCOL_SHA256 = (
    "36aa2a43ad55d4597473bfdda9aa4406541b65047f42ea8789833085704a68a3"
)
ASME_SHA256 = (
    "e767e08dce8e46d1589203e76289a5acf64284166efc54325b945108e9559017"
)
KOJIMA_SHA256 = (
    "a3874f1b8dfb8af1d9329f95e4a6da9a0cb86935d1b17f294e7601c092cc32c9"
)
ASME_PHYSICAL = (
    (17, 23, 5, 18, 20, 25, 21, 9, 15, 16),
    (1, 19, 4, 6, 7, 8, 10, 11, 12, 29),
    (26, 24, 3, 22, 2, 14, 13, 27, 28, 30),
)
ASME_WRONG = (
    (
        tuple(range(1, 31, 3)),
        tuple(range(2, 31, 3)),
        tuple(range(3, 31, 3)),
    ),
    (
        tuple(range(1, 11)),
        tuple(range(11, 21)),
        tuple(range(21, 31)),
    ),
)
KOJIMA_PHYSICAL = ((1, 2), (3, 4))
KOJIMA_WRONG = (((1, 3), (2, 4)), ((1, 4), (2, 3)))
KOJIMA_SUBJECTS = tuple("BDFHJLN")
HORIZON = 3
BOOTSTRAP_N = 20_000
BOOTSTRAP_SEED = 20260730


def sha256(path: Path) -> str:
    digest = hashlib.sha256()
    with path.open("rb") as handle:
        for block in iter(lambda: handle.read(1024 * 1024), b""):
            digest.update(block)
    return digest.hexdigest()


def validate_lock() -> None:
    observed = {
        "protocol": sha256(PROTOCOL),
        "asme_features": sha256(ASME_FEATURES),
        "kojima_features": sha256(KOJIMA_FEATURES),
    }
    expected = {
        "protocol": PROTOCOL_SHA256,
        "asme_features": ASME_SHA256,
        "kojima_features": KOJIMA_SHA256,
    }
    if observed != expected:
        raise RuntimeError(f"Frozen input mismatch: {observed} != {expected}")
    if not LOCK.exists():
        raise FileNotFoundError("Implementation lock required.")
    lock = json.loads(LOCK.read_text(encoding="utf-8"))
    if lock.get("synthetic_test_status") != "PASS":
        raise RuntimeError("Synthetic test not locked as PASS.")
    if lock.get("scientific_outcomes_available_at_lock") is not False:
        raise RuntimeError("Lock is not pre-outcome.")


def event_labels(
    command: np.ndarray,
    target: np.ndarray,
    groups,
    n_commands: int,
) -> np.ndarray:
    lookup = np.full(n_commands + 1, -1, dtype=int)
    flattened = []
    for group_index, group in enumerate(groups):
        lookup[list(group)] = group_index
        flattened.extend(int(value) for value in group)
    if sorted(flattened) != list(range(1, n_commands + 1)):
        raise RuntimeError("Invalid mapping.")
    result = np.full(command.size, "U", dtype="U1")
    target_mask = command == target
    result[target_mask] = "T"
    result[(lookup[command] == lookup[target]) & ~target_mask] = "S"
    return result


def command_residualize(
    train_x: np.ndarray,
    test_x: np.ndarray,
    train_command: np.ndarray,
    train_target: np.ndarray,
    test_command: np.ndarray,
    n_commands: int,
) -> tuple[np.ndarray, np.ndarray, dict]:
    residual_train = np.asarray(train_x, dtype=float).copy()
    residual_test = np.asarray(test_x, dtype=float).copy()
    counts = {}
    non_target = train_command != train_target
    for command_value in range(1, n_commands + 1):
        baseline_ids = (train_command == command_value) & non_target
        if not np.any(baseline_ids):
            raise RuntimeError(f"No non-target baseline for {command_value}.")
        command_mean = np.mean(train_x[baseline_ids], axis=0)
        residual_train[train_command == command_value] -= command_mean
        residual_test[test_command == command_value] -= command_mean
        counts[str(command_value)] = int(np.sum(baseline_ids))
    return residual_train, residual_test, counts


def complete_trials(
    command: np.ndarray, trial: np.ndarray, n_commands: int
) -> list[np.ndarray]:
    selections = []
    for trial_id in sorted(int(value) for value in np.unique(trial)):
        ids = np.flatnonzero(trial == trial_id)
        chunks = []
        for command_value in range(1, n_commands + 1):
            command_ids = ids[command[ids] == command_value]
            if command_ids.size < HORIZON:
                chunks = []
                break
            chunks.append(command_ids[:HORIZON])
        if len(chunks) == n_commands:
            selections.append(np.concatenate(chunks))
    return selections


def fit_scores(
    train_x: np.ndarray,
    train_labels: np.ndarray,
    test_x: np.ndarray,
    train_command: np.ndarray,
    n_commands: int,
) -> tuple[np.ndarray, dict]:
    outer = np.isin(train_labels, ("S", "U"))
    y = (train_labels[outer] == "S").astype(int)
    model = fit_shrinkage_lda(train_x[outer], y)
    train_score = model.decision_function(train_x[outer])
    mean = float(train_score.mean())
    sd = float(train_score.std(ddof=1))
    score = (model.decision_function(test_x) - mean) / sd
    per_command_balance = {}
    for command_value in range(1, n_commands + 1):
        ids = train_command == command_value
        per_command_balance[str(command_value)] = {
            "S": int(np.sum(ids & (train_labels == "S"))),
            "U": int(np.sum(ids & (train_labels == "U"))),
        }
    return score, {
        "n_train_S": int(np.sum(train_labels == "S")),
        "n_train_U": int(np.sum(train_labels == "U")),
        "n_train_T_excluded": int(np.sum(train_labels == "T")),
        "minimum_command_S": min(
            item["S"] for item in per_command_balance.values()
        ),
        "minimum_command_U": min(
            item["U"] for item in per_command_balance.values()
        ),
        "per_command_balance": per_command_balance,
    }


def heldout_effects(
    score: np.ndarray,
    command: np.ndarray,
    target: np.ndarray,
    selections: list[np.ndarray],
    groups,
    n_commands: int,
) -> list[float]:
    expected_s = (len(groups[0]) - 1) * HORIZON
    expected_u = (n_commands - len(groups[0])) * HORIZON
    values = []
    for selected in selections:
        labels = event_labels(
            command[selected], target[selected], groups, n_commands
        )
        counts = tuple(
            int(np.sum(labels == value)) for value in ("T", "S", "U")
        )
        if counts != (HORIZON, expected_s, expected_u):
            raise RuntimeError(f"Unexpected target-free counts: {counts}")
        values.append(
            float(np.mean(score[selected][labels == "S"]))
            - float(np.mean(score[selected][labels == "U"]))
        )
    return values


def cosine(left: np.ndarray, right: np.ndarray) -> float:
    denominator = float(np.linalg.norm(left) * np.linalg.norm(right))
    if denominator <= 1e-12 or not np.isfinite(denominator):
        raise RuntimeError("Degenerate residual direction.")
    return float(np.vdot(left, right) / denominator)


def residual_directions(
    train_x: np.ndarray,
    train_labels: np.ndarray,
    test_x: np.ndarray,
    test_command: np.ndarray,
    test_target: np.ndarray,
    selections: list[np.ndarray],
    groups,
    n_commands: int,
) -> tuple[np.ndarray, np.ndarray, float]:
    outer = np.isin(train_labels, ("S", "U"))
    x = np.asarray(train_x[outer], dtype=float)
    labels = train_labels[outer]
    center = x.mean(axis=0)
    scale = x.std(axis=0, ddof=1)
    scale[~np.isfinite(scale) | (scale <= 1e-12)] = 1.0
    train_z = (x - center) / scale
    training = train_z[labels == "S"].mean(axis=0) - train_z[
        labels == "U"
    ].mean(axis=0)
    test_z = (np.asarray(test_x, dtype=float) - center) / scale
    test_trial_directions = []
    for selected in selections:
        test_labels = event_labels(
            test_command[selected],
            test_target[selected],
            groups,
            n_commands,
        )
        test_trial_directions.append(
            test_z[selected][test_labels == "S"].mean(axis=0)
            - test_z[selected][test_labels == "U"].mean(axis=0)
        )
    heldout = np.stack(test_trial_directions).mean(axis=0)
    return training, heldout, cosine(training, heldout)


def mapping_analysis(
    *,
    train_x: np.ndarray,
    test_x: np.ndarray,
    train_command: np.ndarray,
    train_target: np.ndarray,
    test_command: np.ndarray,
    test_target: np.ndarray,
    selections: list[np.ndarray],
    groups,
    n_commands: int,
) -> dict:
    labels = event_labels(
        train_command, train_target, groups, n_commands
    )
    score, fit = fit_scores(
        train_x, labels, test_x, train_command, n_commands
    )
    effects = heldout_effects(
        score,
        test_command,
        test_target,
        selections,
        groups,
        n_commands,
    )
    training, heldout, alignment = residual_directions(
        train_x,
        labels,
        test_x,
        test_command,
        test_target,
        selections,
        groups,
        n_commands,
    )
    return {
        "trial_effects": effects,
        "effect": float(np.mean(effects)),
        "training_direction": training,
        "heldout_direction": heldout,
        "alignment": alignment,
        "fit": fit,
    }


def analyze_asme_subject(arrays: dict, subject: int) -> dict:
    train = (
        (arrays["subject"] == subject)
        & (arrays["task"] == "offline")
        & arrays["accepted"].astype(bool)
    )
    test = (
        (arrays["subject"] == subject)
        & (arrays["task"] == "online")
        & arrays["accepted"].astype(bool)
    )
    train_x = arrays["eeg_bins"][train, :, 1:].reshape(
        int(np.sum(train)), -1
    )
    test_x = arrays["eeg_bins"][test, :, 1:].reshape(
        int(np.sum(test)), -1
    )
    train_command = arrays["command"][train].astype(int)
    train_target = arrays["target"][train].astype(int)
    test_command = arrays["command"][test].astype(int)
    test_target = arrays["target"][test].astype(int)
    train_residual, test_residual, baseline_counts = command_residualize(
        train_x,
        test_x,
        train_command,
        train_target,
        test_command,
        30,
    )
    train_trials = complete_trials(
        train_command, arrays["trial"][train].astype(int), 30
    )
    test_trials = complete_trials(
        test_command, arrays["trial"][test].astype(int), 30
    )
    physical = mapping_analysis(
        train_x=train_residual,
        test_x=test_residual,
        train_command=train_command,
        train_target=train_target,
        test_command=test_command,
        test_target=test_target,
        selections=test_trials,
        groups=ASME_PHYSICAL,
        n_commands=30,
    )
    wrong = [
        mapping_analysis(
            train_x=train_residual,
            test_x=test_residual,
            train_command=train_command,
            train_target=train_target,
            test_command=test_command,
            test_target=test_target,
            selections=test_trials,
            groups=groups,
            n_commands=30,
        )
        for groups in ASME_WRONG
    ]
    return {
        "subject": subject,
        "n_train_trials": len(train_trials),
        "n_test_trials": len(test_trials),
        "baseline_counts": baseline_counts,
        "physical": physical,
        "wrong": wrong,
    }


def analyze_kojima_subject(arrays: dict, subject: str) -> dict:
    base = (
        (arrays["subject"] == subject)
        & (arrays["task"] == "2stream")
        & arrays["accepted"].astype(bool)
    )
    runs = sorted(int(value) for value in np.unique(arrays["run"][base]))
    folds = []
    for heldout_run in runs:
        train = base & (arrays["run"] != heldout_run)
        test = base & (arrays["run"] == heldout_run)
        train_x = arrays["eeg_bins"][train, :, 1:].reshape(
            int(np.sum(train)), -1
        )
        test_x = arrays["eeg_bins"][test, :, 1:].reshape(
            int(np.sum(test)), -1
        )
        train_command = arrays["command"][train].astype(int)
        train_target = arrays["target"][train].astype(int)
        test_command = arrays["command"][test].astype(int)
        test_target = arrays["target"][test].astype(int)
        train_residual, test_residual, baseline_counts = command_residualize(
            train_x,
            test_x,
            train_command,
            train_target,
            test_command,
            4,
        )
        selections = complete_trials(
            test_command, arrays["trial"][test].astype(int), 4
        )
        physical = mapping_analysis(
            train_x=train_residual,
            test_x=test_residual,
            train_command=train_command,
            train_target=train_target,
            test_command=test_command,
            test_target=test_target,
            selections=selections,
            groups=KOJIMA_PHYSICAL,
            n_commands=4,
        )
        wrong = [
            mapping_analysis(
                train_x=train_residual,
                test_x=test_residual,
                train_command=train_command,
                train_target=train_target,
                test_command=test_command,
                test_target=test_target,
                selections=selections,
                groups=groups,
                n_commands=4,
            )
            for groups in KOJIMA_WRONG
        ]
        folds.append(
            {
                "heldout_run": heldout_run,
                "n_test_trials": len(selections),
                "baseline_counts": baseline_counts,
                "physical": physical,
                "wrong": wrong,
            }
        )
    return {
        "subject": subject,
        "runs": runs,
        "n_test_trials": sum(fold["n_test_trials"] for fold in folds),
        "folds": folds,
        "physical_effect": float(
            np.mean(
                [
                    value
                    for fold in folds
                    for value in fold["physical"]["trial_effects"]
                ]
            )
        ),
        "wrong_effects": [
            float(
                np.mean(
                    [
                        value
                        for fold in folds
                        for value in fold["wrong"][mapping_index][
                            "trial_effects"
                        ]
                    ]
                )
            )
            for mapping_index in range(2)
        ],
        "physical_alignment": float(
            np.mean([fold["physical"]["alignment"] for fold in folds])
        ),
        "wrong_alignments": [
            float(
                np.mean(
                    [
                        fold["wrong"][mapping_index]["alignment"]
                        for fold in folds
                    ]
                )
            )
            for mapping_index in range(2)
        ],
    }


def bootstrap_ci(values: np.ndarray) -> list[float]:
    rng = np.random.default_rng(BOOTSTRAP_SEED)
    ids = rng.integers(0, len(values), size=(BOOTSTRAP_N, len(values)))
    return [
        float(value)
        for value in np.quantile(values[ids].mean(axis=1), [0.025, 0.975])
    ]


def exact_p(values: np.ndarray) -> float:
    observed = float(values.mean())
    permutations = [
        float(np.mean(values * np.asarray(signs)))
        for signs in itertools.product((-1.0, 1.0), repeat=len(values))
    ]
    return float(np.mean(np.asarray(permutations) >= observed - 1e-15))


def summarize(values: np.ndarray) -> dict:
    return {
        "mean": float(values.mean()),
        "bootstrap_95_CI_mean": bootstrap_ci(values),
        "n_positive": int(np.sum(values > 0)),
        "n_total": int(len(values)),
        "one_sided_exact_signflip_p": exact_p(values),
        "participant_values": [float(value) for value in values],
    }


def gate(summary: dict, required: int) -> bool:
    return (
        summary["mean"] > 0
        and summary["bootstrap_95_CI_mean"][0] > 0
        and summary["n_positive"] >= required
        and summary["one_sided_exact_signflip_p"] <= 0.05
    )


def run_analysis(
    asme_arrays: dict,
    kojima_arrays: dict,
    asme_raw: dict,
    kojima_raw: dict,
) -> dict:
    asme_records = [
        analyze_asme_subject(asme_arrays, subject)
        for subject in range(1, 11)
    ]
    asme_raw_lookup = {
        int(row["subject"]): row for row in asme_raw["participant_geometry"]
    }
    asme_rows = []
    asme_offline = [
        record["physical"]["training_direction"] for record in asme_records
    ]
    for index, record in enumerate(asme_records):
        physical = record["physical"]
        wrong_1, wrong_2 = record["wrong"]
        between = float(
            np.mean(
                [
                    cosine(
                        physical["heldout_direction"],
                        asme_offline[donor],
                    )
                    for donor in range(len(asme_records))
                    if donor != index
                ]
            )
        )
        best_wrong_effect = max(wrong_1["effect"], wrong_2["effect"])
        best_wrong_alignment = max(
            wrong_1["alignment"], wrong_2["alignment"]
        )
        raw = asme_raw_lookup[record["subject"]]
        asme_rows.append(
            {
                "subject": record["subject"],
                "n_train_trials": record["n_train_trials"],
                "n_test_trials": record["n_test_trials"],
                "physical_effect": physical["effect"],
                "wrong_1_effect": wrong_1["effect"],
                "wrong_2_effect": wrong_2["effect"],
                "binding_specificity": physical["effect"]
                - best_wrong_effect,
                "physical_alignment": physical["alignment"],
                "wrong_1_alignment": wrong_1["alignment"],
                "wrong_2_alignment": wrong_2["alignment"],
                "mapping_alignment_specificity": physical["alignment"]
                - best_wrong_alignment,
                "between_person_substitution": between,
                "identity_advantage": physical["alignment"] - between,
                "residual_to_raw_physical_ratio": physical["effect"]
                / raw["physical_full_effect"],
                "residual_to_raw_binding_ratio": (
                    physical["effect"] - best_wrong_effect
                )
                / raw["binding_specificity"],
            }
        )

    kojima_records = [
        analyze_kojima_subject(kojima_arrays, subject)
        for subject in KOJIMA_SUBJECTS
    ]
    kojima_raw_lookup = {
        row["subject"]: row
        for row in kojima_raw["participant_geometry"]
    }
    kojima_rows = []
    for participant_index, record in enumerate(kojima_records):
        between_fold = []
        for fold_index, heldout_run in enumerate(record["runs"]):
            heldout = record["folds"][fold_index]["physical"][
                "heldout_direction"
            ]
            donors = []
            for donor_index, donor in enumerate(kojima_records):
                if donor_index == participant_index:
                    continue
                donor_fold_index = donor["runs"].index(heldout_run)
                donors.append(
                    cosine(
                        heldout,
                        donor["folds"][donor_fold_index]["physical"][
                            "training_direction"
                        ],
                    )
                )
            between_fold.append(float(np.mean(donors)))
        between = float(np.mean(between_fold))
        best_wrong_effect = max(record["wrong_effects"])
        best_wrong_alignment = max(record["wrong_alignments"])
        raw = kojima_raw_lookup[record["subject"]]
        kojima_rows.append(
            {
                "subject": record["subject"],
                "n_runs": len(record["runs"]),
                "n_test_trials": record["n_test_trials"],
                "physical_effect": record["physical_effect"],
                "wrong_1_effect": record["wrong_effects"][0],
                "wrong_2_effect": record["wrong_effects"][1],
                "binding_specificity": record["physical_effect"]
                - best_wrong_effect,
                "physical_alignment": record["physical_alignment"],
                "wrong_1_alignment": record["wrong_alignments"][0],
                "wrong_2_alignment": record["wrong_alignments"][1],
                "mapping_alignment_specificity": record[
                    "physical_alignment"
                ]
                - best_wrong_alignment,
                "between_person_substitution": between,
                "identity_advantage": record["physical_alignment"] - between,
                "residual_to_raw_physical_ratio": record["physical_effect"]
                / raw["physical_full_effect"],
                "residual_to_raw_binding_ratio": (
                    record["physical_effect"] - best_wrong_effect
                )
                / raw["binding_specificity"],
            }
        )

    def dataset_summaries(rows: list[dict]) -> dict:
        return {
            name: summarize(np.asarray([row[key] for row in rows]))
            for name, key in (
                ("physical", "physical_effect"),
                ("binding", "binding_specificity"),
                ("identity", "identity_advantage"),
                ("mapping_alignment", "mapping_alignment_specificity"),
                ("physical_retention_ratio", "residual_to_raw_physical_ratio"),
                ("binding_retention_ratio", "residual_to_raw_binding_ratio"),
            )
        }

    summaries = {
        "ASME30": dataset_summaries(asme_rows),
        "Kojima2024B": dataset_summaries(kojima_rows),
    }
    balance_ok = all(
        min(record["baseline_counts"].values()) > 0
        and all(
            mapping["fit"]["minimum_command_S"] > 0
            and mapping["fit"]["minimum_command_U"] > 0
            for mapping in [record["physical"], *record["wrong"]]
        )
        for record in asme_records
    ) and all(
        min(fold["baseline_counts"].values()) > 0
        and all(
            mapping["fit"]["minimum_command_S"] > 0
            and mapping["fit"]["minimum_command_U"] > 0
            for mapping in [fold["physical"], *fold["wrong"]]
        )
        for record in kojima_records
        for fold in record["folds"]
    )
    gates = {
        "C0_balance_and_coverage": balance_ok
        and all(
            row["n_train_trials"] == 30 and row["n_test_trials"] == 15
            for row in asme_rows
        )
        and all(
            row["n_runs"] == 6 and row["n_test_trials"] >= 23
            for row in kojima_rows
        ),
        "C1_ASME_physical": gate(summaries["ASME30"]["physical"], 10),
        "C2_ASME_binding": gate(summaries["ASME30"]["binding"], 8),
        "C3_ASME_identity": gate(summaries["ASME30"]["identity"], 8),
        "C4_ASME_mapping_alignment": gate(
            summaries["ASME30"]["mapping_alignment"], 8
        ),
        "C5_Kojima_physical": gate(
            summaries["Kojima2024B"]["physical"], 7
        ),
        "C6_Kojima_binding": gate(
            summaries["Kojima2024B"]["binding"], 6
        ),
        "C7_Kojima_identity": gate(
            summaries["Kojima2024B"]["identity"], 6
        ),
        "C8_Kojima_mapping_alignment": gate(
            summaries["Kojima2024B"]["mapping_alignment"], 6
        ),
    }
    return {
        "asme_records": asme_records,
        "kojima_records": kojima_records,
        "participant_rows": {
            "ASME30": asme_rows,
            "Kojima2024B": kojima_rows,
        },
        "summaries": summaries,
        "gates": gates,
        "supported": all(gates.values()),
    }


def main() -> None:
    validate_lock()
    if RESULT.exists():
        raise FileExistsError("Immutable v1 result already exists.")
    with np.load(ASME_FEATURES, allow_pickle=True) as source:
        asme_arrays = {key: source[key] for key in source.files}
    with np.load(KOJIMA_FEATURES, allow_pickle=True) as source:
        kojima_arrays = {key: source[key] for key in source.files}
    asme_raw = json.loads(ASME_RAW_RESULT.read_text(encoding="utf-8"))
    kojima_raw = json.loads(KOJIMA_RAW_RESULT.read_text(encoding="utf-8"))
    analysis = run_analysis(asme_arrays, kojima_arrays, asme_raw, kojima_raw)
    output = {
        "route": "WZ-HIC-cross-dataset-command-identity-residual-v1",
        "created_utc": datetime.now(timezone.utc).isoformat(),
        "status": (
            "command_identity_explanation_disfavored"
            if analysis["supported"]
            else "command_identity_control_not_supported_no_retuning"
        ),
        "command_identity_explanation_disfavored": analysis["supported"],
        "gates": analysis["gates"],
        "summaries": analysis["summaries"],
        "participant_results": analysis["participant_rows"],
        "balance_audit": {
            "ASME30": {
                str(record["subject"]): {
                    "command_baseline_counts": record["baseline_counts"],
                    "physical_fit": record["physical"]["fit"],
                    "wrong_fits": [
                        mapping["fit"] for mapping in record["wrong"]
                    ],
                }
                for record in analysis["asme_records"]
            },
            "Kojima2024B": {
                record["subject"]: [
                    {
                        "heldout_run": fold["heldout_run"],
                        "command_baseline_counts": fold["baseline_counts"],
                        "physical_fit": fold["physical"]["fit"],
                        "wrong_fits": [
                            mapping["fit"] for mapping in fold["wrong"]
                        ],
                    }
                    for fold in record["folds"]
                ]
                for record in analysis["kojima_records"]
            },
        },
        "provenance": {
            "protocol_sha256": sha256(PROTOCOL),
            "implementation_lock_sha256": sha256(LOCK),
            "asme_features_sha256": sha256(ASME_FEATURES),
            "kojima_features_sha256": sha256(KOJIMA_FEATURES),
            "asme_raw_result_sha256": sha256(ASME_RAW_RESULT),
            "kojima_raw_result_sha256": sha256(KOJIMA_RAW_RESULT),
            "runner_sha256": sha256(Path(__file__)),
        },
        "interpretation_boundary": (
            "Training-only command-mean residualization removes stable "
            "command identity patterns while preserving attention-dependent "
            "deviations. A passing result disfavors, but cannot eliminate "
            "every possible, acoustic or task confound."
        ),
    }
    OUTDIR.mkdir(parents=True, exist_ok=True)
    RESULT.write_text(
        json.dumps(output, ensure_ascii=False, indent=2) + "\n",
        encoding="utf-8",
    )
    print(
        json.dumps(
            {
                "result": RESULT.relative_to(ROOT).as_posix(),
                "result_sha256": sha256(RESULT),
                "status": output["status"],
                "gates": output["gates"],
                "summaries": output["summaries"],
            },
            ensure_ascii=False,
            indent=2,
        )
    )


if __name__ == "__main__":
    main()
```
