## Supplementary material for "Falsifiable substitution tests reveal task-structured neural evidence for auditory attention": Article_Analysis_Source_Code: run_wz_hic_cross_dataset_wearable_calibration_v1.py

```
#!/usr/bin/env python3
"""Frozen WZ-HIC cross-dataset wearable/calibration robustness analysis."""

from __future__ import annotations

import csv
import hashlib
import itertools
import json
import math
from datetime import datetime, timezone
from pathlib import Path

import numpy as np

from run_wz_hic_cross_dataset_command_identity_residual_v1 import (
    command_residualize,
    complete_trials,
    cosine,
    event_labels,
    mapping_analysis,
)


ROOT = Path(__file__).resolve().parents[1]
PROTOCOL = (
    ROOT
    / "docs"
    / "wz_hic_cross_dataset_wearable_calibration_protocol_v1.json"
)
LOCK = (
    ROOT
    / "docs"
    / "wz_hic_cross_dataset_wearable_calibration_implementation_lock_v1.json"
)
ASME_FEATURES = (
    ROOT
    / "data"
    / "derived"
    / "asme30_wz_hic_dataset_native_features_v2.npz"
)
KOJIMA_FEATURES = (
    ROOT
    / "data"
    / "derived"
    / "kojima2024b_wz_hic_confirmation_features_v1.npz"
)
REFERENCE_RUNNER = (
    ROOT
    / "analysis"
    / "run_wz_hic_cross_dataset_command_identity_residual_v1.py"
)
LDA_CORE = ROOT / "analysis" / "wz_hif_core_v1.py"
OUTDIR = (
    ROOT
    / "results"
    / "wz_hic_cross_dataset"
    / "wearable_calibration_v1"
)
RESULT = OUTDIR / "wearable_calibration_decision_v1.json"
PARTICIPANT_CSV = OUTDIR / "participant_metrics_v1.csv"
MONTAGE_CSV = OUTDIR / "montage_curve_v1.csv"
CALIBRATION_CSV = OUTDIR / "calibration_curve_v1.csv"

PROTOCOL_SHA256 = (
    "ea2e64d24720ff2a02871572fb77977f1a8024c7858fe216a28febd280671977"
)
ASME_SHA256 = (
    "e767e08dce8e46d1589203e76289a5acf64284166efc54325b945108e9559017"
)
KOJIMA_SHA256 = (
    "a3874f1b8dfb8af1d9329f95e4a6da9a0cb86935d1b17f294e7601c092cc32c9"
)
REFERENCE_RUNNER_SHA256 = (
    "f5149e0f4bf6f01679641275b0c4d871855ff0cc85739ba4a87f61fcab37a405"
)
LDA_CORE_SHA256 = (
    "06b32980cd02b7a186c90cd13c84f9abb76377b7837e49563190a28bf2bcd7d9"
)
BOOTSTRAP_N = 20_000
BOOTSTRAP_SEED = 20260730
ASME_SUBJECTS = tuple(range(1, 11))
KOJIMA_SUBJECTS = tuple("BDFHJLN")

ASME_PHYSICAL = (
    (17, 23, 5, 18, 20, 25, 21, 9, 15, 16),
    (1, 19, 4, 6, 7, 8, 10, 11, 12, 29),
    (26, 24, 3, 22, 2, 14, 13, 27, 28, 30),
)
ASME_WRONG = (
    (
        tuple(range(1, 31, 3)),
        tuple(range(2, 31, 3)),
        tuple(range(3, 31, 3)),
    ),
    (
        tuple(range(1, 11)),
        tuple(range(11, 21)),
        tuple(range(21, 31)),
    ),
)
KOJIMA_PHYSICAL = ((1, 2), (3, 4))
KOJIMA_WRONG = (((1, 3), (2, 4)), ((1, 4), (2, 3)))

MONTAGE_8 = ("Fz", "FCz", "C3", "Cz", "C4", "CPz", "Pz", "Oz")
MONTAGE_16 = MONTAGE_8 + (
    "F3",
    "F4",
    "FC3",
    "FC4",
    "CP3",
    "CP4",
    "P3",
    "P4",
)
MONTAGE_32 = MONTAGE_16 + (
    "Fp1",
    "Fp2",
    "AF3",
    "AF4",
    "F7",
    "F8",
    "FC1",
    "FC2",
    "T7",
    "T8",
    "CP1",
    "CP2",
    "P7",
    "P8",
    "PO3",
    "PO4",
)
EXPECTED_CHANNELS = (
    "Fp1",
    "Fp2",
    "AF7",
    "AF3",
    "AFz",
    "AF4",
    "AF8",
    "F7",
    "F5",
    "F3",
    "F1",
    "Fz",
    "F2",
    "F4",
    "F6",
    "F8",
    "FT9",
    "FT7",
    "FC5",
    "FC3",
    "FC1",
    "FCz",
    "FC2",
    "FC4",
    "FC6",
    "FT8",
    "FT10",
    "T7",
    "C5",
    "C3",
    "C1",
    "Cz",
    "C2",
    "C4",
    "C6",
    "T8",
    "TP9",
    "TP7",
    "CP5",
    "CP3",
    "CP1",
    "CPz",
    "CP2",
    "CP4",
    "CP6",
    "TP8",
    "TP10",
    "P7",
    "P5",
    "P3",
    "P1",
    "Pz",
    "P2",
    "P4",
    "P6",
    "P8",
    "PO7",
    "PO3",
    "POz",
    "PO4",
    "PO8",
    "O1",
    "Oz",
    "O2",
)
MONTAGES = {
    8: MONTAGE_8,
    16: MONTAGE_16,
    32: MONTAGE_32,
    64: EXPECTED_CHANNELS,
}


def sha256(path: Path) -> str:
    digest = hashlib.sha256()
    with path.open("rb") as handle:
        for block in iter(lambda: handle.read(1024 * 1024), b""):
            digest.update(block)
    return digest.hexdigest()


def validate_lock() -> None:
    observed = {
        "protocol": sha256(PROTOCOL),
        "asme_features": sha256(ASME_FEATURES),
        "kojima_features": sha256(KOJIMA_FEATURES),
        "reference_runner": sha256(REFERENCE_RUNNER),
        "lda_core": sha256(LDA_CORE),
    }
    expected = {
        "protocol": PROTOCOL_SHA256,
        "asme_features": ASME_SHA256,
        "kojima_features": KOJIMA_SHA256,
        "reference_runner": REFERENCE_RUNNER_SHA256,
        "lda_core": LDA_CORE_SHA256,
    }
    if observed != expected:
        raise RuntimeError(f"Frozen-input mismatch: {observed} != {expected}")
    if not LOCK.exists():
        raise FileNotFoundError("Implementation lock is required.")
    lock = json.loads(LOCK.read_text(encoding="utf-8"))
    if lock.get("scientific_outcomes_available_at_lock") is not False:
        raise RuntimeError("Implementation was not locked before new outcomes.")
    if lock.get("synthetic_test_status") != "PASS":
        raise RuntimeError("Synthetic test was not locked as PASS.")
    if lock.get("runner_sha256") != sha256(Path(__file__)):
        raise RuntimeError("Runner differs from implementation lock.")


def channel_indices(arrays: dict, montage_n: int) -> np.ndarray:
    observed = tuple(str(value) for value in arrays["channel_names"])
    if observed != EXPECTED_CHANNELS:
        raise RuntimeError("Frozen channel order differs from expectation.")
    names = MONTAGES[montage_n]
    if len(names) != montage_n or len(set(names)) != montage_n:
        raise RuntimeError(f"Invalid montage {montage_n}.")
    lookup = {name: index for index, name in enumerate(observed)}
    return np.asarray([lookup[name] for name in names], dtype=int)


def eeg_matrix(arrays: dict, mask: np.ndarray, montage_n: int) -> np.ndarray:
    indices = channel_indices(arrays, montage_n)
    bins = arrays["eeg_bins"][mask][:, indices, 1:]
    if bins.shape[1:] != (montage_n, 10):
        raise RuntimeError(f"Unexpected EEG shape {bins.shape}.")
    return bins.reshape(bins.shape[0], -1)


def eog_matrix(arrays: dict, mask: np.ndarray) -> np.ndarray:
    bins = arrays["eog_bins"][mask, :, 1:]
    if bins.shape[1:] != (2, 10):
        raise RuntimeError(f"Unexpected EOG shape {bins.shape}.")
    return bins.reshape(bins.shape[0], -1)


def analyze_xy_partition(
    *,
    train_x: np.ndarray,
    test_x: np.ndarray,
    train_eog: np.ndarray,
    test_eog: np.ndarray,
    train_command: np.ndarray,
    train_target: np.ndarray,
    test_command: np.ndarray,
    test_target: np.ndarray,
    test_trial: np.ndarray,
    physical_groups,
    wrong_groups,
    n_commands: int,
) -> dict:
    train_x, test_x, baseline_counts = command_residualize(
        train_x,
        test_x,
        train_command,
        train_target,
        test_command,
        n_commands,
    )
    train_eog, test_eog, eog_baseline_counts = command_residualize(
        train_eog,
        test_eog,
        train_command,
        train_target,
        test_command,
        n_commands,
    )
    selections = complete_trials(test_command, test_trial, n_commands)
    physical = mapping_analysis(
        train_x=train_x,
        test_x=test_x,
        train_command=train_command,
        train_target=train_target,
        test_command=test_command,
        test_target=test_target,
        selections=selections,
        groups=physical_groups,
        n_commands=n_commands,
    )
    wrong = [
        mapping_analysis(
            train_x=train_x,
            test_x=test_x,
            train_command=train_command,
            train_target=train_target,
            test_command=test_command,
            test_target=test_target,
            selections=selections,
            groups=groups,
            n_commands=n_commands,
        )
        for groups in wrong_groups
    ]
    eog = mapping_analysis(
        train_x=train_eog,
        test_x=test_eog,
        train_command=train_command,
        train_target=train_target,
        test_command=test_command,
        test_target=test_target,
        selections=selections,
        groups=physical_groups,
        n_commands=n_commands,
    )
    return {
        "n_train_events": int(len(train_command)),
        "n_test_events": int(len(test_command)),
        "n_test_trials": int(len(selections)),
        "baseline_counts": baseline_counts,
        "eog_baseline_counts": eog_baseline_counts,
        "physical": physical,
        "wrong": wrong,
        "eog": eog,
    }


def analyze_mask_partition(
    *,
    arrays: dict,
    train: np.ndarray,
    test: np.ndarray,
    montage_n: int,
    physical_groups,
    wrong_groups,
    n_commands: int,
) -> dict:
    return analyze_xy_partition(
        train_x=eeg_matrix(arrays, train, montage_n),
        test_x=eeg_matrix(arrays, test, montage_n),
        train_eog=eog_matrix(arrays, train),
        test_eog=eog_matrix(arrays, test),
        train_command=arrays["command"][train].astype(int),
        train_target=arrays["target"][train].astype(int),
        test_command=arrays["command"][test].astype(int),
        test_target=arrays["target"][test].astype(int),
        test_trial=arrays["trial"][test].astype(int),
        physical_groups=physical_groups,
        wrong_groups=wrong_groups,
        n_commands=n_commands,
    )


def collapse_partition(partition: dict) -> dict:
    wrong_effects = [item["effect"] for item in partition["wrong"]]
    wrong_alignments = [item["alignment"] for item in partition["wrong"]]
    return {
        "physical_effect": float(partition["physical"]["effect"]),
        "best_wrong_effect": float(max(wrong_effects)),
        "binding_specificity": float(
            partition["physical"]["effect"] - max(wrong_effects)
        ),
        "eog_effect": float(partition["eog"]["effect"]),
        "nonocular_specificity": float(
            partition["physical"]["effect"] - partition["eog"]["effect"]
        ),
        "physical_alignment": float(partition["physical"]["alignment"]),
        "best_wrong_alignment": float(max(wrong_alignments)),
        "mapping_alignment_specificity": float(
            partition["physical"]["alignment"] - max(wrong_alignments)
        ),
    }


def asme_full_records(arrays: dict, montage_n: int) -> list[dict]:
    records = []
    for subject in ASME_SUBJECTS:
        accepted = arrays["accepted"].astype(bool)
        train = (
            (arrays["subject"] == subject)
            & (arrays["task"] == "offline")
            & accepted
        )
        test = (
            (arrays["subject"] == subject)
            & (arrays["task"] == "online")
            & accepted
        )
        partition = analyze_mask_partition(
            arrays=arrays,
            train=train,
            test=test,
            montage_n=montage_n,
            physical_groups=ASME_PHYSICAL,
            wrong_groups=ASME_WRONG,
            n_commands=30,
        )
        training_trials = complete_trials(
            arrays["command"][train].astype(int),
            arrays["trial"][train].astype(int),
            30,
        )
        records.append(
            {
                "participant": str(subject),
                "n_train_trials": len(training_trials),
                "n_test_trials": partition["n_test_trials"],
                "partition": partition,
                **collapse_partition(partition),
            }
        )
    for index, record in enumerate(records):
        heldout = record["partition"]["physical"]["heldout_direction"]
        between = np.mean(
            [
                cosine(
                    heldout,
                    donor["partition"]["physical"]["training_direction"],
                )
                for donor_index, donor in enumerate(records)
                if donor_index != index
            ]
        )
        record["between_person_substitution"] = float(between)
        record["identity_advantage"] = float(
            record["physical_alignment"] - between
        )
    return records


def kojima_full_records(arrays: dict, montage_n: int) -> list[dict]:
    records = []
    accepted = arrays["accepted"].astype(bool)
    for subject in KOJIMA_SUBJECTS:
        base = (
            (arrays["subject"] == subject)
            & (arrays["task"] == "2stream")
            & accepted
        )
        runs = sorted(int(value) for value in np.unique(arrays["run"][base]))
        if len(runs) != 6:
            raise RuntimeError(f"Subject {subject}: expected six runs.")
        folds = []
        for heldout_run in runs:
            train = base & (arrays["run"] != heldout_run)
            test = base & (arrays["run"] == heldout_run)
            partition = analyze_mask_partition(
                arrays=arrays,
                train=train,
                test=test,
                montage_n=montage_n,
                physical_groups=KOJIMA_PHYSICAL,
                wrong_groups=KOJIMA_WRONG,
                n_commands=4,
            )
            folds.append(
                {
                    "heldout_run": heldout_run,
                    "partition": partition,
                    **collapse_partition(partition),
                }
            )
        physical_trial_effects = [
            value
            for fold in folds
            for value in fold["partition"]["physical"]["trial_effects"]
        ]
        wrong_trial_effects = [
            [
                value
                for fold in folds
                for value in fold["partition"]["wrong"][wrong_index][
                    "trial_effects"
                ]
            ]
            for wrong_index in range(2)
        ]
        eog_trial_effects = [
            value
            for fold in folds
            for value in fold["partition"]["eog"]["trial_effects"]
        ]
        physical_effect = float(np.mean(physical_trial_effects))
        wrong_effects = [
            float(np.mean(values)) for values in wrong_trial_effects
        ]
        eog_effect = float(np.mean(eog_trial_effects))
        physical_alignment = float(
            np.mean([fold["physical_alignment"] for fold in folds])
        )
        wrong_alignments = [
            float(
                np.mean(
                    [
                        fold["partition"]["wrong"][wrong_index]["alignment"]
                        for fold in folds
                    ]
                )
            )
            for wrong_index in range(2)
        ]
        records.append(
            {
                "participant": subject,
                "runs": runs,
                "n_train_trials_per_fold": [
                    20 for _ in folds
                ],
                "n_test_trials": int(
                    sum(
                        fold["partition"]["n_test_trials"] for fold in folds
                    )
                ),
                "folds": folds,
                "physical_effect": physical_effect,
                "best_wrong_effect": max(wrong_effects),
                "binding_specificity": physical_effect - max(wrong_effects),
                "eog_effect": eog_effect,
                "nonocular_specificity": physical_effect - eog_effect,
                "physical_alignment": physical_alignment,
                "best_wrong_alignment": max(wrong_alignments),
                "mapping_alignment_specificity": (
                    physical_alignment - max(wrong_alignments)
                ),
            }
        )
    for subject_index, record in enumerate(records):
        between_folds = []
        for fold_index, fold in enumerate(record["folds"]):
            heldout = fold["partition"]["physical"]["heldout_direction"]
            donor_values = [
                cosine(
                    heldout,
                    donor["folds"][fold_index]["partition"]["physical"][
                        "training_direction"
                    ],
                )
                for donor_index, donor in enumerate(records)
                if donor_index != subject_index
            ]
            between_folds.append(float(np.mean(donor_values)))
        between = float(np.mean(between_folds))
        record["between_person_substitution"] = between
        record["identity_advantage"] = (
            record["physical_alignment"] - between
        )
    return records


def bootstrap_ci(values: np.ndarray) -> list[float]:
    values = np.asarray(values, dtype=float)
    rng = np.random.default_rng(BOOTSTRAP_SEED)
    ids = rng.integers(0, len(values), size=(BOOTSTRAP_N, len(values)))
    return [
        float(value)
        for value in np.quantile(values[ids].mean(axis=1), [0.025, 0.975])
    ]


def exact_p(values: np.ndarray) -> float:
    values = np.asarray(values, dtype=float)
    observed = float(np.mean(values))
    permuted = [
        float(np.mean(values * np.asarray(signs)))
        for signs in itertools.product((-1.0, 1.0), repeat=len(values))
    ]
    return float(np.mean(np.asarray(permuted) >= observed - 1e-15))


def summarize(values) -> dict:
    values = np.asarray(values, dtype=float)
    return {
        "mean": float(np.mean(values)),
        "bootstrap_95_CI_mean": bootstrap_ci(values),
        "n_positive": int(np.sum(values > 0)),
        "n_total": int(len(values)),
        "one_sided_exact_signflip_p": exact_p(values),
        "participant_values": [float(value) for value in values],
    }


def summarize_records(records: list[dict]) -> dict:
    metrics = (
        "physical_effect",
        "binding_specificity",
        "nonocular_specificity",
        "identity_advantage",
        "mapping_alignment_specificity",
        "physical_alignment",
        "between_person_substitution",
    )
    return {
        metric: summarize([record[metric] for record in records])
        for metric in metrics
    }


def scalar_support(item: dict, required_positive: int) -> bool:
    return (
        item["mean"] > 0
        and item["bootstrap_95_CI_mean"][0] > 0
        and item["n_positive"] >= required_positive
        and item["one_sided_exact_signflip_p"] <= 0.05
    )


def asme_calibration_records(arrays: dict) -> tuple[list[dict], dict]:
    records = []
    integrity = {}
    accepted = arrays["accepted"].astype(bool)
    for subject in ASME_SUBJECTS:
        offline = (
            (arrays["subject"] == subject)
            & (arrays["task"] == "offline")
            & accepted
        )
        test = (
            (arrays["subject"] == subject)
            & (arrays["task"] == "online")
            & accepted
        )
        runs = sorted(int(value) for value in np.unique(arrays["run"][offline]))
        if len(runs) != 6:
            raise RuntimeError(f"ASME {subject}: expected six offline runs.")
        integrity[str(subject)] = {}
        for k in range(1, 7):
            cells = []
            combinations = list(itertools.combinations(runs, k))
            for selected_runs in combinations:
                train = offline & np.isin(arrays["run"], selected_runs)
                partition = analyze_mask_partition(
                    arrays=arrays,
                    train=train,
                    test=test,
                    montage_n=8,
                    physical_groups=ASME_PHYSICAL,
                    wrong_groups=ASME_WRONG,
                    n_commands=30,
                )
                collapsed = collapse_partition(partition)
                cells.append(collapsed)
            integrity[str(subject)][str(k)] = len(cells)
            records.append(
                {
                    "dataset": "ASME30",
                    "participant": str(subject),
                    "training_runs": k,
                    "subset_cells": len(cells),
                    "physical_effect": float(
                        np.mean([cell["physical_effect"] for cell in cells])
                    ),
                    "binding_specificity": float(
                        np.mean(
                            [cell["binding_specificity"] for cell in cells]
                        )
                    ),
                }
            )
    return records, integrity


def kojima_calibration_records(arrays: dict) -> tuple[list[dict], dict]:
    records = []
    integrity = {}
    accepted = arrays["accepted"].astype(bool)
    for subject in KOJIMA_SUBJECTS:
        base = (
            (arrays["subject"] == subject)
            & (arrays["task"] == "2stream")
            & accepted
        )
        runs = sorted(int(value) for value in np.unique(arrays["run"][base]))
        if len(runs) != 6:
            raise RuntimeError(f"Kojima {subject}: expected six runs.")
        integrity[subject] = {}
        for k in range(1, 6):
            cells = []
            for heldout_run in runs:
                available = [run for run in runs if run != heldout_run]
                test = base & (arrays["run"] == heldout_run)
                for selected_runs in itertools.combinations(available, k):
                    train = base & np.isin(arrays["run"], selected_runs)
                    partition = analyze_mask_partition(
                        arrays=arrays,
                        train=train,
                        test=test,
                        montage_n=8,
                        physical_groups=KOJIMA_PHYSICAL,
                        wrong_groups=KOJIMA_WRONG,
                        n_commands=4,
                    )
                    cells.append(collapse_partition(partition))
            expected = 6 * math.comb(5, k)
            if len(cells) != expected:
                raise RuntimeError("Kojima subset enumeration failed.")
            integrity[subject][str(k)] = len(cells)
            records.append(
                {
                    "dataset": "Kojima2024B",
                    "participant": subject,
                    "training_runs": k,
                    "subset_cells": len(cells),
                    "physical_effect": float(
                        np.mean([cell["physical_effect"] for cell in cells])
                    ),
                    "binding_specificity": float(
                        np.mean(
                            [cell["binding_specificity"] for cell in cells]
                        )
                    ),
                }
            )
    return records, integrity


def calibration_summaries(records: list[dict]) -> dict:
    output = {}
    for dataset in ("ASME30", "Kojima2024B"):
        dataset_records = [
            record for record in records if record["dataset"] == dataset
        ]
        output[dataset] = {}
        for k in sorted(
            int(value)
            for value in set(
                record["training_runs"] for record in dataset_records
            )
        ):
            rows = [
                record
                for record in dataset_records
                if record["training_runs"] == k
            ]
            output[dataset][str(k)] = {
                "physical_effect": summarize(
                    [record["physical_effect"] for record in rows]
                ),
                "binding_specificity": summarize(
                    [record["binding_specificity"] for record in rows]
                ),
            }
    return output


def public_participant_row(
    dataset: str, montage_n: int, record: dict
) -> dict:
    keys = (
        "physical_effect",
        "best_wrong_effect",
        "binding_specificity",
        "eog_effect",
        "nonocular_specificity",
        "physical_alignment",
        "best_wrong_alignment",
        "mapping_alignment_specificity",
        "between_person_substitution",
        "identity_advantage",
    )
    return {
        "dataset": dataset,
        "montage_channels": montage_n,
        "participant": record["participant"],
        "n_test_trials": record["n_test_trials"],
        **{key: float(record[key]) for key in keys},
    }


def summary_rows(montage_summaries: dict) -> list[dict]:
    rows = []
    for dataset, budgets in montage_summaries.items():
        for montage_n, summaries in budgets.items():
            for metric, item in summaries.items():
                rows.append(
                    {
                        "dataset": dataset,
                        "montage_channels": int(montage_n),
                        "metric": metric,
                        "mean": item["mean"],
                        "ci_low": item["bootstrap_95_CI_mean"][0],
                        "ci_high": item["bootstrap_95_CI_mean"][1],
                        "n_positive": item["n_positive"],
                        "n_total": item["n_total"],
                        "p_one_sided_exact": item[
                            "one_sided_exact_signflip_p"
                        ],
                    }
                )
    return rows


def calibration_summary_rows(summaries: dict) -> list[dict]:
    rows = []
    for dataset, points in summaries.items():
        for k, metrics in points.items():
            for metric, item in metrics.items():
                rows.append(
                    {
                        "dataset": dataset,
                        "montage_channels": 8,
                        "training_runs": int(k),
                        "metric": metric,
                        "mean": item["mean"],
                        "ci_low": item["bootstrap_95_CI_mean"][0],
                        "ci_high": item["bootstrap_95_CI_mean"][1],
                        "n_positive": item["n_positive"],
                        "n_total": item["n_total"],
                        "p_one_sided_exact": item[
                            "one_sided_exact_signflip_p"
                        ],
                    }
                )
    return rows


def run_analysis(asme: dict, kojima: dict) -> dict:
    montage_records = {"ASME30": {}, "Kojima2024B": {}}
    montage_summaries = {"ASME30": {}, "Kojima2024B": {}}
    participant_rows = []
    for montage_n in (8, 16, 32, 64):
        asme_records = asme_full_records(asme, montage_n)
        kojima_records = kojima_full_records(kojima, montage_n)
        montage_records["ASME30"][str(montage_n)] = asme_records
        montage_records["Kojima2024B"][str(montage_n)] = kojima_records
        montage_summaries["ASME30"][str(montage_n)] = summarize_records(
            asme_records
        )
        montage_summaries["Kojima2024B"][str(montage_n)] = summarize_records(
            kojima_records
        )
        participant_rows.extend(
            public_participant_row("ASME30", montage_n, record)
            for record in asme_records
        )
        participant_rows.extend(
            public_participant_row("Kojima2024B", montage_n, record)
            for record in kojima_records
        )

    asme_calibration, asme_calibration_integrity = (
        asme_calibration_records(asme)
    )
    kojima_calibration, kojima_calibration_integrity = (
        kojima_calibration_records(kojima)
    )
    calibration_records = asme_calibration + kojima_calibration
    calibration_summary = calibration_summaries(calibration_records)

    required = {"ASME30": 8, "Kojima2024B": 6}

    def both_montage(metric: str) -> bool:
        return all(
            scalar_support(
                montage_summaries[dataset]["8"][metric],
                required[dataset],
            )
            for dataset in required
        )

    def both_calibration(metric: str) -> bool:
        return all(
            scalar_support(
                calibration_summary[dataset]["3"][metric],
                required[dataset],
            )
            for dataset in required
        )

    retention = {
        dataset: (
            montage_summaries[dataset]["8"]["binding_specificity"]["mean"]
            / montage_summaries[dataset]["64"]["binding_specificity"]["mean"]
        )
        for dataset in required
    }
    integrity = {
        "asme_participants": len(
            montage_records["ASME30"]["8"]
        ) == len(ASME_SUBJECTS),
        "kojima_participants": len(
            montage_records["Kojima2024B"]["8"]
        ) == len(KOJIMA_SUBJECTS),
        "all_montage_rows": len(participant_rows)
        == 4 * (len(ASME_SUBJECTS) + len(KOJIMA_SUBJECTS)),
        "asme_trial_coverage": all(
            record["n_train_trials"] == 30
            and record["n_test_trials"] == 15
            for budget in montage_records["ASME30"].values()
            for record in budget
        ),
        "kojima_trial_coverage": all(
            record["n_test_trials"] >= 23
            for budget in montage_records["Kojima2024B"].values()
            for record in budget
        ),
        "asme_subset_enumeration": all(
            count == math.comb(6, int(k))
            for participant in asme_calibration_integrity.values()
            for k, count in participant.items()
        ),
        "kojima_subset_enumeration": all(
            count == 6 * math.comb(5, int(k))
            for participant in kojima_calibration_integrity.values()
            for k, count in participant.items()
        ),
        "channel_archives_identical": tuple(
            str(value) for value in asme["channel_names"]
        )
        == tuple(str(value) for value in kojima["channel_names"])
        == EXPECTED_CHANNELS,
    }
    gates = {
        "W0_integrity": all(integrity.values()),
        "W1_eight_channel_physical": both_montage("physical_effect"),
        "W2_eight_channel_binding": both_montage("binding_specificity"),
        "W3_eight_channel_nonocular": both_montage(
            "nonocular_specificity"
        ),
        "W4_eight_channel_identity": both_montage("identity_advantage"),
        "W5_eight_channel_retention": all(
            np.isfinite(value) and value >= 0.50
            for value in retention.values()
        )
        and all(
            montage_summaries[dataset]["64"]["binding_specificity"]["mean"]
            > 0
            for dataset in required
        ),
        "W6_half_calibration_physical": both_calibration(
            "physical_effect"
        ),
        "W7_half_calibration_binding": both_calibration(
            "binding_specificity"
        ),
    }
    return {
        "montage_summaries": montage_summaries,
        "participant_rows": participant_rows,
        "calibration_records": calibration_records,
        "calibration_summaries": calibration_summary,
        "retention": retention,
        "integrity": integrity,
        "asme_subset_counts": asme_calibration_integrity,
        "kojima_subset_counts": kojima_calibration_integrity,
        "gates": gates,
        "supported": all(gates.values()),
    }


def write_csv(path: Path, rows: list[dict]) -> None:
    if not rows:
        raise RuntimeError(f"No rows for {path}.")
    with path.open("x", newline="", encoding="utf-8-sig") as handle:
        writer = csv.DictWriter(handle, fieldnames=list(rows[0]))
        writer.writeheader()
        writer.writerows(rows)


def main() -> None:
    validate_lock()
    for path in (RESULT, PARTICIPANT_CSV, MONTAGE_CSV, CALIBRATION_CSV):
        if path.exists():
            raise FileExistsError(f"Immutable v1 output exists: {path}")
    with np.load(ASME_FEATURES, allow_pickle=True) as source:
        asme = {key: source[key] for key in source.files}
    with np.load(KOJIMA_FEATURES, allow_pickle=True) as source:
        kojima = {key: source[key] for key in source.files}
    analysis = run_analysis(asme, kojima)
    output = {
        "route": "WZ-HIC-cross-dataset-wearable-calibration-v1",
        "created_utc": datetime.now(timezone.utc).isoformat(),
        "status": (
            "wearable_calibration_supported"
            if analysis["supported"]
            else "wearable_calibration_not_supported_no_retuning"
        ),
        "wearable_calibration_supported": analysis["supported"],
        "gates": analysis["gates"],
        "integrity": analysis["integrity"],
        "eight_channel_binding_retention": analysis["retention"],
        "montage_summaries": analysis["montage_summaries"],
        "calibration_summaries": analysis["calibration_summaries"],
        "subset_count_audit": {
            "asme": analysis["asme_subset_counts"],
            "kojima": analysis["kojima_subset_counts"],
        },
        "provenance": {
            "protocol_sha256": sha256(PROTOCOL),
            "implementation_lock_sha256": sha256(LOCK),
            "asme_features_sha256": sha256(ASME_FEATURES),
            "kojima_features_sha256": sha256(KOJIMA_FEATURES),
            "reference_runner_sha256": sha256(REFERENCE_RUNNER),
            "lda_core_sha256": sha256(LDA_CORE),
            "runner_sha256": sha256(Path(__file__)),
        },
        "interpretation_boundary": (
            "Post-primary robustness analysis. Support means the locked "
            "target-free, command-residual physical-binding and identity "
            "effects survive an eight-electrode montage and three-run "
            "calibration in two public event-related datasets. It does not "
            "demonstrate online benefit, continuous-speech generalization, "
            "source localization, or a direct application of the Kakeya "
            "theorem."
        ),
    }
    OUTDIR.mkdir(parents=True, exist_ok=True)
    RESULT.write_text(
        json.dumps(output, ensure_ascii=False, indent=2) + "\n",
        encoding="utf-8",
    )
    write_csv(PARTICIPANT_CSV, analysis["participant_rows"])
    write_csv(MONTAGE_CSV, summary_rows(analysis["montage_summaries"]))
    write_csv(
        CALIBRATION_CSV,
        calibration_summary_rows(analysis["calibration_summaries"]),
    )
    print(
        json.dumps(
            {
                "status": output["status"],
                "gates": output["gates"],
                "retention": output["eight_channel_binding_retention"],
                "eight_channel": {
                    dataset: output["montage_summaries"][dataset]["8"]
                    for dataset in ("ASME30", "Kojima2024B")
                },
                "three_run": {
                    dataset: output["calibration_summaries"][dataset]["3"]
                    for dataset in ("ASME30", "Kojima2024B")
                },
                "files": {
                    str(path.relative_to(ROOT)): sha256(path)
                    for path in (
                        RESULT,
                        PARTICIPANT_CSV,
                        MONTAGE_CSV,
                        CALIBRATION_CSV,
                    )
                },
            },
            ensure_ascii=False,
            indent=2,
        )
    )


if __name__ == "__main__":
    main()
```
