## Supplementary material for "Falsifiable substitution tests reveal task-structured neural evidence for auditory attention": Article_Analysis_Source_Code: run_wz_hic_kojima2024b_confirmation_v1.py

```
#!/usr/bin/env python
"""Run the single frozen WZ-HIC-v1 confirmation decision."""

from __future__ import annotations

import hashlib
import json
import math
from datetime import datetime, timezone
from pathlib import Path

import numpy as np

from wz_hic_confirmation_core_v1 import (
    TRUE_PAIRING,
    WRONG_PAIRINGS,
    bootstrap_mean_ci,
    certificate_metrics,
    exact_sign_p,
    exact_signflip_mean_p,
    loro_event_scores,
    pseudo_labels,
    trial_rows,
)
from wz_hif_core_v1 import select_window_features


ROOT = Path(__file__).resolve().parents[1]
FEATURES = (
    ROOT / "data" / "derived" / "kojima2024b_wz_hic_confirmation_features_v1.npz"
)
FEATURE_AUDIT = (
    ROOT
    / "data"
    / "inventory"
    / "kojima2024b_wz_hic_confirmation_feature_audit_v1.json"
)
PROTOCOL = ROOT / "docs" / "wz_hic_kojima2024b_confirmation_protocol_v1.json"
LOCK = ROOT / "docs" / "wz_hic_kojima2024b_confirmation_implementation_lock_v1.json"
RESULT = (
    ROOT
    / "results"
    / "wz_hic_kojima2024b"
    / "confirmation_v1"
    / "confirmation_decision_v1.json"
)
DATA_ROOT = ROOT / "data" / "raw" / "kojima2024b_structure_all"
CONFIRMATION = tuple("BDFHJLN")
ALL_PAIRINGS = (TRUE_PAIRING,) + WRONG_PAIRINGS


def sha256(path: Path) -> str:
    h = hashlib.sha256()
    with path.open("rb") as f:
        for block in iter(lambda: f.read(1024 * 1024), b""):
            h.update(block)
    return h.hexdigest()


def pairing_key(pairing) -> str:
    return "-".join("".join(str(x) for x in pair) for pair in pairing)


def validate_lock() -> dict:
    lock = json.loads(LOCK.read_text(encoding="utf-8"))
    files = {
        "protocol": PROTOCOL,
        "core": ROOT / "analysis" / "wz_hic_confirmation_core_v1.py",
        "feature_builder": ROOT
        / "analysis"
        / "build_kojima2024b_wz_hic_confirmation_features_v1.py",
        "runner": Path(__file__),
        "synthetic_test": ROOT
        / "analysis"
        / "test_wz_hic_confirmation_synthetic_v1.py",
    }
    observed = {name: sha256(path) for name, path in files.items()}
    if observed != lock["locked_sha256"]:
        raise RuntimeError(f"Implementation lock mismatch: {observed}")
    return lock


def effect(values: list[float], *, null: float = 0.0, signflip=False) -> dict:
    a = np.asarray(values, dtype=float)
    a = a[np.isfinite(a)]
    differences = a - null
    return {
        "mean": float(a.mean()) if a.size else float("nan"),
        "mean_difference_from_null": (
            float(differences.mean()) if a.size else float("nan")
        ),
        "bootstrap_95_CI_mean": (
            bootstrap_mean_ci([float(x) for x in a])
            if a.size
            else [float("nan"), float("nan")]
        ),
        "n_positive": int(np.sum(differences > 0)),
        "n_total": int(a.size),
        "fraction_positive": (
            float(np.mean(differences > 0)) if a.size else float("nan")
        ),
        "one_sided_exact_p": (
            exact_signflip_mean_p([float(x) for x in differences])
            if signflip and a.size
            else exact_sign_p([float(x) for x in a], null)
            if a.size
            else float("nan")
        ),
        "test": "exact_signflip_mean" if signflip else "exact_sign",
        "participant_values": [float(x) for x in a],
    }


def absolute_gate(s: dict) -> bool:
    return (
        s["n_total"] >= 5
        and s["mean"] >= 0.75
        and s["n_positive"] == s["n_total"]
        and s["one_sided_exact_p"] <= 0.03125
    )


def paired_gate(s: dict, threshold: float) -> bool:
    return (
        s["n_total"] >= 5
        and s["mean"] >= threshold
        and s["fraction_positive"] >= 0.8
        and s["one_sided_exact_p"] <= 0.05
    )


def make_scores_and_metrics(a: dict, mask: np.ndarray, labels: np.ndarray) -> tuple:
    scores = loro_event_scores(
        select_window_features(a["eeg_bins"][mask], (-0.1, 1.0)),
        select_window_features(a["eog_bins"][mask], (-0.1, 1.0)),
        select_window_features(a["eeg_bins"][mask], (-0.1, 0.0)),
        select_window_features(a["eeg_bins"][mask], (0.3, 0.8)),
        labels,
        a["run"][mask],
    )
    common = (
        a["command"][mask],
        a["trial"][mask],
        a["target"][mask],
    )
    true_rows = trial_rows(
        scores,
        *common,
        horizon=3,
        certificate_name="eeg_outer",
    )
    eog_rows = trial_rows(
        scores,
        *common,
        horizon=3,
        certificate_name="eog_outer",
    )
    prestim_rows = trial_rows(
        scores,
        *common,
        horizon=3,
        certificate_name="prestim_outer",
    )
    wrong_rows = [
        trial_rows(
            scores,
            *common,
            horizon=3,
            certificate_name="eeg_outer",
            pooling_pairing=pairing,
            correctness_pairing=TRUE_PAIRING,
        )
        for pairing in WRONG_PAIRINGS
    ]
    return {
        "EEG_true": certificate_metrics(true_rows),
        "EOG_true": certificate_metrics(eog_rows),
        "prestim_true": certificate_metrics(prestim_rows),
        "wrong_pairings": [certificate_metrics(rows) for rows in wrong_rows],
    }, scores


def main() -> None:
    if RESULT.exists():
        raise FileExistsError(f"Single-run decision already exists: {RESULT}")
    lock = validate_lock()
    feature_audit = json.loads(FEATURE_AUDIT.read_text(encoding="utf-8"))
    if feature_audit["status"] != "PASS":
        raise RuntimeError("Feature audit failed.")
    if feature_audit["output_sha256"] != sha256(FEATURES):
        raise RuntimeError("Feature hash mismatch.")
    if feature_audit["protocol_sha256"] != sha256(PROTOCOL):
        raise RuntimeError("Protocol hash mismatch.")
    if list(DATA_ROOT.glob("sub-*/nasatlx/*")):
        raise RuntimeError("NASA-TLX seal broken.")

    z = np.load(FEATURES, allow_pickle=False)
    accepted = z["accepted"]
    a = {
        key: z[key][accepted]
        for key in (
            "eeg_bins",
            "eog_bins",
            "subject",
            "task",
            "run",
            "trial",
            "target",
            "command",
            "label",
        )
    }
    if set(a["subject"]) != set(CONFIRMATION):
        raise RuntimeError("Unexpected confirmation participant set.")

    participants = {}
    for subject in CONFIRMATION:
        mask = (a["subject"] == subject) & (a["task"] == "2stream")
        participants[subject], _ = make_scores_and_metrics(
            a, mask, a["label"][mask]
        )
        print("2STREAM", subject, flush=True)

    evaluable = [
        subject
        for subject in CONFIRMATION
        if participants[subject]["EEG_true"]["n_trials"] >= 12
        and participants[subject]["EEG_true"]["n_parent_correct"] > 0
        and participants[subject]["EEG_true"]["n_parent_error"] > 0
    ]
    ceiling = [
        subject
        for subject in CONFIRMATION
        if participants[subject]["EEG_true"]["n_trials"] >= 12
        and participants[subject]["EEG_true"]["n_parent_error"] == 0
    ]
    absolute_values = [
        participants[s]["EEG_true"]["certificate_AUC"] for s in evaluable
    ]
    flat_gains = [
        participants[s]["EEG_true"]["certificate_AUC"]
        - participants[s]["EEG_true"]["flat_margin_AUC"]
        for s in evaluable
    ]
    eog_gains = [
        participants[s]["EEG_true"]["certificate_AUC"]
        - participants[s]["EOG_true"]["certificate_AUC"]
        for s in evaluable
    ]
    mapping_gains = [
        participants[s]["EEG_true"]["certificate_AUC"]
        - max(
            item["certificate_AUC"]
            for item in participants[s]["wrong_pairings"]
        )
        for s in evaluable
    ]
    agreement_gains = [
        participants[s]["EEG_true"]["agreement_accuracy_gain"]
        for s in CONFIRMATION
    ]
    agreement_coverages = [
        participants[s]["EEG_true"]["agreement_coverage"]
        for s in CONFIRMATION
    ]
    prestim_values = [
        participants[s]["prestim_true"]["certificate_AUC"] for s in evaluable
    ]

    H1 = effect(absolute_values, null=0.5)
    H2 = effect(flat_gains, signflip=True)
    H3 = effect(eog_gains, signflip=True)
    H4 = effect(mapping_gains, signflip=True)
    H5 = effect(agreement_gains, signflip=True)
    prestim_summary = effect(prestim_values, null=0.5)

    # Four-stream pseudo-parent control: fit each arbitrary pairing separately.
    pseudo4 = {}
    for subject in CONFIRMATION:
        mask = (a["subject"] == subject) & (a["task"] == "4stream")
        pseudo4[subject] = {}
        for pairing in ALL_PAIRINGS:
            labels = pseudo_labels(
                a["command"][mask], a["target"][mask], pairing
            )
            scores = loro_event_scores(
                select_window_features(a["eeg_bins"][mask], (-0.1, 1.0)),
                select_window_features(a["eog_bins"][mask], (-0.1, 1.0)),
                select_window_features(a["eeg_bins"][mask], (-0.1, 0.0)),
                select_window_features(a["eeg_bins"][mask], (0.3, 0.8)),
                labels,
                a["run"][mask],
            )
            rows = trial_rows(
                scores,
                a["command"][mask],
                a["trial"][mask],
                a["target"][mask],
                horizon=3,
                certificate_name="eeg_outer",
                pooling_pairing=pairing,
                correctness_pairing=pairing,
            )
            pseudo4[subject][pairing_key(pairing)] = certificate_metrics(rows)
        print("4STREAM", subject, flush=True)
    pseudo_valid = [
        s
        for s in CONFIRMATION
        if all(
            np.isfinite(pseudo4[s][pairing_key(p)]["certificate_AUC"])
            for p in ALL_PAIRINGS
        )
    ]
    pseudo_advantage = [
        pseudo4[s][pairing_key(TRUE_PAIRING)]["certificate_AUC"]
        - max(
            pseudo4[s][pairing_key(p)]["certificate_AUC"]
            for p in WRONG_PAIRINGS
        )
        for s in pseudo_valid
    ]
    pseudo_summary = effect(pseudo_advantage, signflip=True)
    pseudo_privileged = paired_gate(pseudo_summary, 0.10)

    gates = {
        "H1_absolute_certificate": absolute_gate(H1),
        "H2_beyond_flat_confidence": paired_gate(H2, 0.08),
        "H3_beyond_ocular_certificate": paired_gate(H3, 0.05),
        "H4_physical_mapping_specificity": paired_gate(H4, 0.10),
        "H5_selective_reliability": (
            H5["n_total"] == 7
            and H5["mean"] >= 0.03
            and float(np.mean(agreement_coverages)) >= 0.5
            and H5["n_positive"] >= 6
            and H5["one_sided_exact_p"] <= 0.05
        ),
        "H6_negative_controls": (
            not absolute_gate(prestim_summary) and not pseudo_privileged
        ),
    }
    pass_all = all(gates.values())
    result = {
        "route": "WZ-HIC-v1",
        "status": (
            "confirmed_advance_to_behavior_and_external_protocol"
            if pass_all
            else "stop_WZ_HIC_v1_without_behavior_or_external_rescue"
        ),
        "created_utc": datetime.now(timezone.utc).isoformat(),
        "provenance": {
            "protocol_sha256": sha256(PROTOCOL),
            "implementation_lock_sha256": sha256(LOCK),
            "features_sha256": sha256(FEATURES),
            "feature_audit_sha256": sha256(FEATURE_AUDIT),
            "NASA_TLX_accessed": False,
        },
        "evaluable_participants": evaluable,
        "flat_parent_ceiling_participants": ceiling,
        "participant_results_2stream": participants,
        "primary_outcomes": {
            "H1_absolute_certificate": H1,
            "H2_beyond_flat_confidence": H2,
            "H3_beyond_ocular_certificate": H3,
            "H4_physical_mapping_specificity": H4,
            "H5_selective_reliability": H5,
            "agreement_coverage_mean": float(np.mean(agreement_coverages)),
        },
        "negative_controls": {
            "prestimulus": prestim_summary,
            "four_stream_pseudo_parent_participants": pseudo4,
            "four_stream_true_pairing_advantage": pseudo_summary,
            "four_stream_true_pairing_privileged": pseudo_privileged,
        },
        "gates": gates,
        "all_gates_pass": pass_all,
        "implementation_lock": lock,
    }
    RESULT.parent.mkdir(parents=True, exist_ok=True)
    RESULT.write_text(
        json.dumps(result, ensure_ascii=False, indent=2) + "\n", encoding="utf-8"
    )
    print(json.dumps({"status": result["status"], "gates": gates}, indent=2))
    print(RESULT, sha256(RESULT))


if __name__ == "__main__":
    main()
```
