## Supplementary material for "Falsifiable substitution tests reveal task-structured neural evidence for auditory attention": Article_Analysis_Source_Code: run_wz_hic_kojima2024b_crossrun_individual_geometry_v1.py

```
#!/usr/bin/env python3
"""Frozen Kojima2024B target-free cross-run individualized geometry."""

from __future__ import annotations

import csv
import hashlib
import itertools
import json
import math
from datetime import datetime, timezone
from pathlib import Path

import numpy as np

from wz_hif_core_v1 import fit_shrinkage_lda


ROOT = Path(__file__).resolve().parents[1]
PROTOCOL = (
    ROOT
    / "docs"
    / "wz_hic_kojima2024b_crossrun_individual_geometry_protocol_v1.json"
)
LOCK = (
    ROOT
    / "docs"
    / "wz_hic_kojima2024b_crossrun_individual_geometry_implementation_lock_v1.json"
)
FEATURES = (
    ROOT
    / "data"
    / "derived"
    / "kojima2024b_wz_hic_confirmation_features_v1.npz"
)
OUTDIR = (
    ROOT
    / "results"
    / "wz_hic_kojima2024b"
    / "crossrun_individual_geometry_v1"
)
RESULT = OUTDIR / "crossrun_decision_v1.json"
PARTICIPANT_CSV = OUTDIR / "participant_geometry_v1.csv"
TIME_CSV = OUTDIR / "time_resolved_v1.csv"

PROTOCOL_SHA256 = (
    "00db57d1d3ab9f8116001b0c5200005aeafe8828d97eaa70fe576da6d0ba9b59"
)
FEATURES_SHA256 = (
    "a3874f1b8dfb8af1d9329f95e4a6da9a0cb86935d1b17f294e7601c092cc32c9"
)
SUBJECTS = tuple("BDFHJLN")
PHYSICAL = ((1, 2), (3, 4))
WRONG = (((1, 3), (2, 4)), ((1, 4), (2, 3)))
HORIZON = 3
BOOTSTRAP_N = 20_000
BOOTSTRAP_SEED = 20260730


def sha256(path: Path) -> str:
    digest = hashlib.sha256()
    with path.open("rb") as handle:
        for block in iter(lambda: handle.read(1024 * 1024), b""):
            digest.update(block)
    return digest.hexdigest()


def validate_frozen_inputs() -> None:
    observed = {
        "protocol": sha256(PROTOCOL),
        "features": sha256(FEATURES),
    }
    expected = {
        "protocol": PROTOCOL_SHA256,
        "features": FEATURES_SHA256,
    }
    if observed != expected:
        raise RuntimeError(f"Frozen input mismatch: {observed} != {expected}")
    if not LOCK.exists():
        raise FileNotFoundError("Implementation lock must exist before outcomes.")
    lock = json.loads(LOCK.read_text(encoding="utf-8"))
    if lock.get("synthetic_test_status") != "PASS":
        raise RuntimeError("Synthetic test was not locked as PASS.")
    if lock.get("scientific_outcomes_available_at_lock") is not False:
        raise RuntimeError("Implementation lock is not pre-outcome.")


def mapping_lookup(groups) -> np.ndarray:
    lookup = np.full(5, -1, dtype=int)
    flattened = []
    for index, group in enumerate(groups):
        lookup[list(group)] = index
        flattened.extend(int(value) for value in group)
    if sorted(flattened) != [1, 2, 3, 4]:
        raise RuntimeError("Invalid four-command pairing.")
    return lookup


def labels_for_pairing(
    command: np.ndarray, target: np.ndarray, groups
) -> np.ndarray:
    lookup = mapping_lookup(groups)
    labels = np.full(command.size, "U", dtype="U1")
    is_target = command == target
    labels[is_target] = "T"
    labels[(lookup[command] == lookup[target]) & ~is_target] = "S"
    return labels


def trial_selections(
    command: np.ndarray, trial: np.ndarray
) -> tuple[list[np.ndarray], list[int]]:
    selections = []
    trial_ids = []
    for trial_id in sorted(int(value) for value in np.unique(trial)):
        ids = np.flatnonzero(trial == trial_id)
        chunks = []
        for value in range(1, 5):
            value_ids = ids[command[ids] == value]
            if value_ids.size < HORIZON:
                chunks = []
                break
            chunks.append(value_ids[:HORIZON])
        if len(chunks) == 4:
            selections.append(np.concatenate(chunks))
            trial_ids.append(trial_id)
    return selections, trial_ids


def fit_scores(
    train_x: np.ndarray,
    train_labels: np.ndarray,
    test_x: np.ndarray,
) -> tuple[np.ndarray, dict]:
    outer = np.isin(train_labels, ("S", "U"))
    y = (train_labels[outer] == "S").astype(int)
    model = fit_shrinkage_lda(train_x[outer], y)
    train_score = model.decision_function(train_x[outer])
    mean = float(train_score.mean())
    sd = float(train_score.std(ddof=1))
    if not np.isfinite(sd) or sd <= 1e-12:
        raise RuntimeError("Degenerate training-fold score scale.")
    test_score = (model.decision_function(test_x) - mean) / sd
    return test_score, {
        "n_train_S": int(np.sum(train_labels == "S")),
        "n_train_U": int(np.sum(train_labels == "U")),
        "n_train_T_excluded": int(np.sum(train_labels == "T")),
        "training_score_mean": mean,
        "training_score_sd": sd,
    }


def trial_effects(
    score: np.ndarray,
    command: np.ndarray,
    target: np.ndarray,
    selections: list[np.ndarray],
    groups,
) -> list[float]:
    values = []
    for selected in selections:
        labels = labels_for_pairing(
            command[selected], target[selected], groups
        )
        counts = tuple(
            int(np.sum(labels == value)) for value in ("T", "S", "U")
        )
        if counts != (3, 3, 6):
            raise RuntimeError(f"Unexpected T/S/U held-out counts: {counts}")
        values.append(
            float(np.mean(score[selected][labels == "S"]))
            - float(np.mean(score[selected][labels == "U"]))
        )
    return values


def cosine(left: np.ndarray, right: np.ndarray) -> float:
    denominator = float(np.linalg.norm(left) * np.linalg.norm(right))
    if not np.isfinite(denominator) or denominator <= 1e-12:
        raise RuntimeError("Degenerate fold direction.")
    return float(np.vdot(left, right) / denominator)


def fold_directions(
    train_x: np.ndarray,
    train_labels: np.ndarray,
    test_x: np.ndarray,
    test_command: np.ndarray,
    test_target: np.ndarray,
    selections: list[np.ndarray],
    groups,
) -> tuple[np.ndarray, np.ndarray, float]:
    outer = np.isin(train_labels, ("S", "U"))
    x = np.asarray(train_x[outer], dtype=float)
    labels = train_labels[outer]
    center = x.mean(axis=0)
    scale = x.std(axis=0, ddof=1)
    scale[~np.isfinite(scale) | (scale <= 1e-12)] = 1.0
    train_z = (x - center) / scale
    training = train_z[labels == "S"].mean(axis=0) - train_z[
        labels == "U"
    ].mean(axis=0)
    test_z = (np.asarray(test_x, dtype=float) - center) / scale
    heldout_trial = []
    for selected in selections:
        test_labels = labels_for_pairing(
            test_command[selected], test_target[selected], groups
        )
        heldout_trial.append(
            test_z[selected][test_labels == "S"].mean(axis=0)
            - test_z[selected][test_labels == "U"].mean(axis=0)
        )
    heldout = np.stack(heldout_trial).mean(axis=0)
    return training, heldout, cosine(training, heldout)


def analyze_mapping_fold(
    *,
    train_bins: np.ndarray,
    test_bins: np.ndarray,
    train_command: np.ndarray,
    train_target: np.ndarray,
    test_command: np.ndarray,
    test_target: np.ndarray,
    selections: list[np.ndarray],
    groups,
) -> dict:
    train_full = train_bins.reshape(train_bins.shape[0], -1)
    test_full = test_bins.reshape(test_bins.shape[0], -1)
    labels = labels_for_pairing(train_command, train_target, groups)
    score, fit = fit_scores(train_full, labels, test_full)
    full_trial_effects = trial_effects(
        score, test_command, test_target, selections, groups
    )
    training_direction, heldout_direction, alignment = fold_directions(
        train_full,
        labels,
        test_full,
        test_command,
        test_target,
        selections,
        groups,
    )
    time_trial_effects = []
    for bin_index in range(10):
        time_score, _ = fit_scores(
            train_bins[:, :, bin_index],
            labels,
            test_bins[:, :, bin_index],
        )
        time_trial_effects.append(
            trial_effects(
                time_score,
                test_command,
                test_target,
                selections,
                groups,
            )
        )
    return {
        "full_trial_effects": full_trial_effects,
        "time_trial_effects": time_trial_effects,
        "training_direction": training_direction,
        "heldout_direction": heldout_direction,
        "alignment": alignment,
        "fit": fit,
    }


def analyze_subject(arrays: dict, subject: str) -> dict:
    subject_mask = (
        (arrays["subject"] == subject)
        & (arrays["task"] == "2stream")
        & arrays["accepted"].astype(bool)
    )
    runs = sorted(int(value) for value in np.unique(arrays["run"][subject_mask]))
    if len(runs) != 6:
        raise RuntimeError(f"Subject {subject} has {len(runs)} runs.")
    folds = []
    for heldout_run in runs:
        train_mask = subject_mask & (arrays["run"] != heldout_run)
        test_mask = subject_mask & (arrays["run"] == heldout_run)
        train_bins = arrays["eeg_bins"][train_mask, :, 1:]
        test_bins = arrays["eeg_bins"][test_mask, :, 1:]
        train_command = arrays["command"][train_mask].astype(int)
        train_target = arrays["target"][train_mask].astype(int)
        test_command = arrays["command"][test_mask].astype(int)
        test_target = arrays["target"][test_mask].astype(int)
        selections, trial_ids = trial_selections(
            test_command, arrays["trial"][test_mask].astype(int)
        )
        physical = analyze_mapping_fold(
            train_bins=train_bins,
            test_bins=test_bins,
            train_command=train_command,
            train_target=train_target,
            test_command=test_command,
            test_target=test_target,
            selections=selections,
            groups=PHYSICAL,
        )
        wrong = [
            analyze_mapping_fold(
                train_bins=train_bins,
                test_bins=test_bins,
                train_command=train_command,
                train_target=train_target,
                test_command=test_command,
                test_target=test_target,
                selections=selections,
                groups=groups,
            )
            for groups in WRONG
        ]
        train_eog = arrays["eog_bins"][train_mask, :, 1:].reshape(
            int(np.sum(train_mask)), -1
        )
        test_eog = arrays["eog_bins"][test_mask, :, 1:].reshape(
            int(np.sum(test_mask)), -1
        )
        physical_labels = labels_for_pairing(
            train_command, train_target, PHYSICAL
        )
        eog_score, eog_fit = fit_scores(
            train_eog, physical_labels, test_eog
        )
        eog_trial_effects = trial_effects(
            eog_score,
            test_command,
            test_target,
            selections,
            PHYSICAL,
        )
        folds.append(
            {
                "heldout_run": heldout_run,
                "n_eligible_trials": len(selections),
                "trial_ids": trial_ids,
                "physical": physical,
                "wrong": wrong,
                "eog_trial_effects": eog_trial_effects,
                "eog_fit": eog_fit,
            }
        )

    def concatenate(mapping_key: str, mapping_index: int | None = None):
        if mapping_index is None:
            return np.asarray(
                [
                    value
                    for fold in folds
                    for value in fold[mapping_key]["full_trial_effects"]
                ]
            )
        return np.asarray(
            [
                value
                for fold in folds
                for value in fold[mapping_key][mapping_index][
                    "full_trial_effects"
                ]
            ]
        )

    physical_values = concatenate("physical")
    wrong_values = [
        concatenate("wrong", index) for index in range(2)
    ]
    eog_values = np.asarray(
        [
            value
            for fold in folds
            for value in fold["eog_trial_effects"]
        ]
    )
    time_effects = []
    for key, mapping_index in (("physical", None), ("wrong", 0), ("wrong", 1)):
        per_bin = []
        for bin_index in range(10):
            per_bin.append(
                np.mean(
                    [
                        value
                        for fold in folds
                        for value in (
                            fold[key]["time_trial_effects"][bin_index]
                            if mapping_index is None
                            else fold[key][mapping_index][
                                "time_trial_effects"
                            ][bin_index]
                        )
                    ]
                )
            )
        time_effects.append(np.asarray(per_bin))
    return {
        "subject": subject,
        "runs": runs,
        "n_eligible_trials": int(len(physical_values)),
        "folds": folds,
        "physical_effect": float(np.mean(physical_values)),
        "wrong_effects": [float(np.mean(value)) for value in wrong_values],
        "eog_effect": float(np.mean(eog_values)),
        "physical_time": time_effects[0],
        "wrong_time": time_effects[1:],
        "physical_alignment": float(
            np.mean([fold["physical"]["alignment"] for fold in folds])
        ),
        "wrong_alignments": [
            float(
                np.mean(
                    [fold["wrong"][index]["alignment"] for fold in folds]
                )
            )
            for index in range(2)
        ],
    }


def bootstrap_ci(values: np.ndarray) -> list[float]:
    rng = np.random.default_rng(BOOTSTRAP_SEED)
    ids = rng.integers(0, len(values), size=(BOOTSTRAP_N, len(values)))
    return [
        float(value)
        for value in np.quantile(values[ids].mean(axis=1), [0.025, 0.975])
    ]


def exact_p(values: np.ndarray) -> float:
    observed = float(np.mean(values))
    permuted = [
        float(np.mean(values * np.asarray(signs)))
        for signs in itertools.product((-1.0, 1.0), repeat=len(values))
    ]
    return float(np.mean(np.asarray(permuted) >= observed - 1e-15))


def summarize(values: np.ndarray) -> dict:
    return {
        "mean": float(np.mean(values)),
        "bootstrap_95_CI_mean": bootstrap_ci(values),
        "n_positive": int(np.sum(values > 0)),
        "n_total": int(len(values)),
        "one_sided_exact_signflip_p": exact_p(values),
        "participant_values": [float(value) for value in values],
    }


def t_stat(values: np.ndarray) -> np.ndarray:
    denominator = values.std(axis=0, ddof=1) / math.sqrt(len(values))
    return np.divide(
        values.mean(axis=0),
        denominator,
        out=np.zeros(values.shape[1]),
        where=denominator > 1e-12,
    )


def exact_max_t(
    values: np.ndarray,
) -> tuple[np.ndarray, np.ndarray, np.ndarray]:
    observed = t_stat(values)
    maxima = []
    for signs in itertools.product((-1.0, 1.0), repeat=len(values)):
        maxima.append(
            float(
                np.max(
                    np.abs(
                        t_stat(values * np.asarray(signs)[:, None])
                    )
                )
            )
        )
    maxima = np.asarray(maxima)
    p = (
        maxima[:, None] >= np.abs(observed)[None, :] - 1e-15
    ).mean(axis=0)
    return observed, p, maxima


def run_analysis(arrays: dict) -> dict:
    records = [analyze_subject(arrays, subject) for subject in SUBJECTS]
    rows = []
    time_specificity = []
    for index, record in enumerate(records):
        between_fold_values = []
        for fold_index, heldout_run in enumerate(record["runs"]):
            heldout_direction = record["folds"][fold_index]["physical"][
                "heldout_direction"
            ]
            donor_values = []
            for donor_index, donor in enumerate(records):
                if donor_index == index:
                    continue
                donor_fold_index = donor["runs"].index(heldout_run)
                donor_training = donor["folds"][donor_fold_index][
                    "physical"
                ]["training_direction"]
                donor_values.append(cosine(heldout_direction, donor_training))
            between_fold_values.append(float(np.mean(donor_values)))
        between = float(np.mean(between_fold_values))
        best_wrong_effect = max(record["wrong_effects"])
        best_wrong_alignment = max(record["wrong_alignments"])
        rows.append(
            {
                "subject": record["subject"],
                "n_runs": len(record["runs"]),
                "n_eligible_trials": record["n_eligible_trials"],
                "physical_full_effect": record["physical_effect"],
                "wrong_1_full_effect": record["wrong_effects"][0],
                "wrong_2_full_effect": record["wrong_effects"][1],
                "best_wrong_full_effect": best_wrong_effect,
                "binding_specificity": record["physical_effect"]
                - best_wrong_effect,
                "physical_eog_effect": record["eog_effect"],
                "nonocular_specificity": record["physical_effect"]
                - record["eog_effect"],
                "physical_within_alignment": record["physical_alignment"],
                "wrong_1_within_alignment": record["wrong_alignments"][0],
                "wrong_2_within_alignment": record["wrong_alignments"][1],
                "best_wrong_within_alignment": best_wrong_alignment,
                "mapping_alignment_specificity": record[
                    "physical_alignment"
                ]
                - best_wrong_alignment,
                "between_person_substitution": between,
                "identity_advantage": record["physical_alignment"] - between,
            }
        )
        time_specificity.append(
            record["physical_time"]
            - np.maximum(record["wrong_time"][0], record["wrong_time"][1])
        )
    time_specificity = np.stack(time_specificity)
    time_t, time_p, time_null = exact_max_t(time_specificity)
    time_rows = [
        {
            "bin_index": bin_index,
            "time_start_s": bin_index / 10,
            "time_stop_s": (bin_index + 1) / 10,
            "mean_physical_minus_best_wrong": float(
                time_specificity[:, bin_index].mean()
            ),
            "t_value": float(time_t[bin_index]),
            "max_t_fwer_p": float(time_p[bin_index]),
            "fwer_significant_0_05": bool(time_p[bin_index] <= 0.05),
            "participant_values": [
                float(value) for value in time_specificity[:, bin_index]
            ],
        }
        for bin_index in range(10)
    ]
    summaries = {
        "physical_crossrun": summarize(
            np.asarray([row["physical_full_effect"] for row in rows])
        ),
        "binding_specificity": summarize(
            np.asarray([row["binding_specificity"] for row in rows])
        ),
        "nonocular_specificity": summarize(
            np.asarray([row["nonocular_specificity"] for row in rows])
        ),
        "identity_advantage": summarize(
            np.asarray([row["identity_advantage"] for row in rows])
        ),
        "mapping_alignment_specificity": summarize(
            np.asarray(
                [row["mapping_alignment_specificity"] for row in rows]
            )
        ),
        "physical_within_alignment": summarize(
            np.asarray(
                [row["physical_within_alignment"] for row in rows]
            )
        ),
        "between_person_substitution": summarize(
            np.asarray(
                [row["between_person_substitution"] for row in rows]
            )
        ),
    }

    def gate(name: str, required: int) -> bool:
        item = summaries[name]
        return (
            item["mean"] > 0
            and item["bootstrap_95_CI_mean"][0] > 0
            and item["n_positive"] >= required
            and item["one_sided_exact_signflip_p"] <= 0.05
        )

    gates = {
        "K0_coverage": len(rows) == 7
        and all(
            row["n_runs"] == 6 and row["n_eligible_trials"] >= 23
            for row in rows
        ),
        "K1_physical_crossrun": gate("physical_crossrun", 7),
        "K2_binding_specificity": gate("binding_specificity", 6),
        "K3_nonocular_specificity": gate("nonocular_specificity", 6),
        "K4_identity_advantage": gate("identity_advantage", 6),
        "K5_mapping_alignment_specificity": gate(
            "mapping_alignment_specificity", 6
        ),
        "K6_time_localization": any(
            row["mean_physical_minus_best_wrong"] > 0
            and row["max_t_fwer_p"] <= 0.05
            for row in time_rows
        ),
    }
    return {
        "records": records,
        "participant_rows": rows,
        "time_rows": time_rows,
        "summaries": summaries,
        "gates": gates,
        "time_null": time_null,
        "supported": all(gates.values()),
    }


def main() -> None:
    validate_frozen_inputs()
    if RESULT.exists() or PARTICIPANT_CSV.exists() or TIME_CSV.exists():
        raise FileExistsError("Immutable v1 outcome already exists.")
    with np.load(FEATURES, allow_pickle=True) as source:
        arrays = {key: source[key] for key in source.files}
    analysis = run_analysis(arrays)
    output = {
        "route": "WZ-HIC-Kojima2024B-crossrun-individual-geometry-v1",
        "created_utc": datetime.now(timezone.utc).isoformat(),
        "status": (
            "cross_dataset_individual_geometry_supported"
            if analysis["supported"]
            else "cross_dataset_individual_geometry_not_supported_no_retuning"
        ),
        "cross_dataset_geometry_supported": analysis["supported"],
        "gates": analysis["gates"],
        "summaries": analysis["summaries"],
        "participant_geometry": analysis["participant_rows"],
        "time_resolved_binding_specificity": analysis["time_rows"],
        "time_max_t": {
            "exact_sign_patterns": 2 ** len(analysis["participant_rows"]),
            "null_95_percentile": float(
                np.quantile(analysis["time_null"], 0.95)
            ),
        },
        "fold_coverage_and_fit_audit": {
            record["subject"]: [
                {
                    "heldout_run": fold["heldout_run"],
                    "n_eligible_trials": fold["n_eligible_trials"],
                    "physical_full": fold["physical"]["fit"],
                    "wrong_1_full": fold["wrong"][0]["fit"],
                    "wrong_2_full": fold["wrong"][1]["fit"],
                    "eog_full": fold["eog_fit"],
                }
                for fold in record["folds"]
            ]
            for record in analysis["records"]
        },
        "provenance": {
            "protocol_sha256": sha256(PROTOCOL),
            "implementation_lock_sha256": sha256(LOCK),
            "features_sha256": sha256(FEATURES),
            "runner_sha256": sha256(Path(__file__)),
        },
        "interpretation_boundary": (
            "Frozen target-free mechanism transfer in seven confirmation "
            "participants under leave-one-run-out validation. A positive "
            "result is distinct-public-dataset support for participant-"
            "specific task-native binding geometry, not a new untouched "
            "primary test or a direct use of the Kakeya theorem."
        ),
    }
    OUTDIR.mkdir(parents=True, exist_ok=True)
    RESULT.write_text(
        json.dumps(output, ensure_ascii=False, indent=2) + "\n",
        encoding="utf-8",
    )
    with PARTICIPANT_CSV.open("x", newline="", encoding="utf-8-sig") as handle:
        writer = csv.DictWriter(
            handle, fieldnames=list(analysis["participant_rows"][0])
        )
        writer.writeheader()
        writer.writerows(analysis["participant_rows"])
    with TIME_CSV.open("x", newline="", encoding="utf-8-sig") as handle:
        keys = [
            key
            for key in analysis["time_rows"][0]
            if key != "participant_values"
        ]
        writer = csv.DictWriter(handle, fieldnames=keys)
        writer.writeheader()
        writer.writerows(
            [{key: row[key] for key in keys} for row in analysis["time_rows"]]
        )
    print(
        json.dumps(
            {
                "result": RESULT.relative_to(ROOT).as_posix(),
                "result_sha256": sha256(RESULT),
                "participant_csv_sha256": sha256(PARTICIPANT_CSV),
                "time_csv_sha256": sha256(TIME_CSV),
                "status": output["status"],
                "gates": output["gates"],
                "summaries": output["summaries"],
                "time": output["time_resolved_binding_specificity"],
            },
            ensure_ascii=False,
            indent=2,
        )
    )


if __name__ == "__main__":
    main()
```
