## Supplementary material for "Falsifiable substitution tests reveal task-structured neural evidence for auditory attention": Article_Analysis_Source_Code: run_wz_hic_target_free_mechanism_v1.py

```
#!/usr/bin/env python3
"""Run the frozen target-event-free WZ-HIC mechanism audit."""

from __future__ import annotations

import hashlib
import itertools
import json
from datetime import datetime, timezone
from pathlib import Path

import numpy as np
from sklearn.metrics import roc_auc_score

from run_wz_hic_asme30_external_v1 import select_bins
from wz_hic_asme30_core_v1 import (
    PHYSICAL_GROUPS,
    WRONG_GROUPS,
    fit_primary_event_scores,
    fit_wrong_mapping_score,
)
from wz_hic_confirmation_core_v1 import (
    TRUE_PAIRING,
    WRONG_PAIRINGS,
    loro_event_scores,
)
from wz_hif_core_v1 import select_window_features


ROOT = Path(__file__).resolve().parents[1]
PROTOCOL = ROOT / "docs" / "wz_hic_target_free_mechanism_protocol_v1.json"
LOCK = (
    ROOT
    / "docs"
    / "wz_hic_target_free_mechanism_implementation_lock_v1c.json"
)
KOJIMA_FEATURES = (
    ROOT
    / "data"
    / "derived"
    / "kojima2024b_wz_hic_confirmation_features_v1.npz"
)
KOJIMA_PRIMARY = (
    ROOT
    / "results"
    / "wz_hic_kojima2024b"
    / "confirmation_v1"
    / "confirmation_decision_v1.json"
)
ASME_FEATURES = (
    ROOT / "data" / "derived" / "asme30_wz_hic_dataset_native_features_v2.npz"
)
ASME_PRIMARY = (
    ROOT
    / "results"
    / "wz_hic_asme30"
    / "dataset_native_v2"
    / "dataset_native_decision_v2.json"
)
RESULT = (
    ROOT
    / "results"
    / "wz_hic_cross_dataset"
    / "target_free_mechanism_v1"
    / "target_free_decision_v1.json"
)

KOJIMA_SUBJECTS = tuple("BDFHJLN")
HORIZON = 3
BOOTSTRAP_SEED = 20260730
BOOTSTRAP_DRAWS = 20_000


def sha256(path: Path) -> str:
    h = hashlib.sha256()
    with path.open("rb") as f:
        for block in iter(lambda: f.read(1024 * 1024), b""):
            h.update(block)
    return h.hexdigest()


def semantic_json_sha256(path: Path) -> str:
    """Hash scientific JSON content while excluding rerun-time metadata.

    Primary result files are regenerated from the locked feature archives.  Their
    ``created_utc`` values and feature-audit byte hashes legitimately change on a
    new run even when every scientific value is identical.  Those fields must not
    be used as scientific input locks.
    """

    def clean(value):
        if isinstance(value, dict):
            return {
                key: clean(item)
                for key, item in value.items()
                if key != "created_utc" and not key.endswith("_audit_sha256")
            }
        if isinstance(value, list):
            return [clean(item) for item in value]
        return value

    payload = json.loads(path.read_text(encoding="utf-8"))
    canonical = json.dumps(
        clean(payload), sort_keys=True, separators=(",", ":"), allow_nan=True
    ).encode("utf-8")
    return hashlib.sha256(canonical).hexdigest()


def validate_lock() -> dict:
    lock = json.loads(LOCK.read_text(encoding="utf-8"))
    files = {
        "protocol": PROTOCOL,
        "runner": Path(__file__),
        "kojima_core": ROOT / "analysis" / "wz_hic_confirmation_core_v1.py",
        "asme_core": ROOT / "analysis" / "wz_hic_asme30_core_v1.py",
        "window_dependency": (
            ROOT / "analysis" / "run_wz_hic_asme30_external_v1.py"
        ),
        "lda_dependency": ROOT / "analysis" / "wz_hif_core_v1.py",
        "synthetic_test": (
            ROOT
            / "analysis"
            / "test_wz_hic_target_free_mechanism_synthetic_v1.py"
        ),
        "engineering_amendment_01": (
            ROOT
            / "docs"
            / "wz_hic_target_free_mechanism_engineering_amendment_01.json"
        ),
        "engineering_amendment_02": (
            ROOT
            / "docs"
            / "wz_hic_target_free_mechanism_engineering_amendment_02.json"
        ),
    }
    observed = {name: sha256(path) for name, path in files.items()}
    if observed != lock["locked_sha256"]:
        raise RuntimeError(
            f"Target-free mechanism implementation lock mismatch: {observed}"
        )
    return lock


def group_index(command: int, groups) -> int:
    matches = [
        index for index, group in enumerate(groups) if int(command) in group
    ]
    if len(matches) != 1:
        raise RuntimeError(
            f"Command {command} belongs to {len(matches)} groups, expected one."
        )
    return matches[0]


def trial_rows(
    *,
    flat_score: np.ndarray,
    outer_score: np.ndarray,
    command: np.ndarray,
    trial: np.ndarray,
    target: np.ndarray,
    commands: tuple[int, ...],
    pooling_groups,
    correctness_groups,
    exclude_true_target: bool,
) -> list[dict]:
    rows = []
    for trial_id in sorted(np.unique(trial)):
        ids = np.flatnonzero(trial == trial_id)
        target_values = np.unique(target[ids])
        if target_values.size != 1:
            raise RuntimeError("Trial target is nonunique.")
        true_target = int(target_values[0])
        if any(
            np.sum(command[ids] == value) < HORIZON
            for value in commands
        ):
            continue
        by_command = {}
        for value in commands:
            value_ids = ids[command[ids] == value]
            by_command[value] = value_ids[:HORIZON]
        child_score = {
            value: float(np.mean(flat_score[by_command[value]]))
            for value in commands
        }
        order = sorted(child_score, key=child_score.get, reverse=True)
        prediction = int(order[0])
        parent_scores = []
        parent_counts = []
        for group in pooling_groups:
            kept_commands = [
                int(value)
                for value in group
                if not (exclude_true_target and int(value) == true_target)
            ]
            if not kept_commands:
                raise RuntimeError("Target exclusion emptied a parent group.")
            event_ids = np.concatenate(
                [by_command[value] for value in kept_commands]
            )
            parent_scores.append(float(np.mean(outer_score[event_ids])))
            parent_counts.append(int(event_ids.size))
        predicted_pooling_parent = group_index(prediction, pooling_groups)
        competitors = [
            value
            for index, value in enumerate(parent_scores)
            if index != predicted_pooling_parent
        ]
        certificate = (
            parent_scores[predicted_pooling_parent] - max(competitors)
        )
        rows.append(
            {
                "trial": int(trial_id),
                "target": true_target,
                "flat_prediction": prediction,
                "parent_correct": (
                    group_index(prediction, correctness_groups)
                    == group_index(true_target, correctness_groups)
                ),
                "certificate": float(certificate),
                "parent_event_counts": parent_counts,
            }
        )
    return rows


def row_metrics(rows: list[dict]) -> dict:
    y = np.asarray([row["parent_correct"] for row in rows], dtype=int)
    score = np.asarray([row["certificate"] for row in rows], dtype=float)
    auc = (
        float(roc_auc_score(y, score))
        if y.size and np.unique(y).size == 2
        else float("nan")
    )
    return {
        "n_trials": int(y.size),
        "n_parent_correct": int(np.sum(y)),
        "n_parent_error": int(np.sum(1 - y)),
        "certificate_AUC": auc,
        "certificate_mean_parent_correct": (
            float(np.mean(score[y == 1])) if np.any(y == 1) else float("nan")
        ),
        "certificate_mean_parent_error": (
            float(np.mean(score[y == 0])) if np.any(y == 0) else float("nan")
        ),
        "minimum_parent_event_count": int(
            min(min(row["parent_event_counts"]) for row in rows)
        ),
        "maximum_parent_event_count": int(
            max(max(row["parent_event_counts"]) for row in rows)
        ),
    }


def exact_signflip_mean_p(values, null=0.0) -> float:
    x = np.asarray(values, dtype=float) - float(null)
    x = x[np.isfinite(x)]
    if x.size == 0:
        return float("nan")
    observed = float(np.mean(x))
    count = 0
    total = 2 ** int(x.size)
    for signs in itertools.product((-1.0, 1.0), repeat=int(x.size)):
        if float(np.mean(x * np.asarray(signs))) >= observed - 1e-15:
            count += 1
    return float(count / total)


def summarize(values, null=0.0) -> dict:
    x = np.asarray(values, dtype=float)
    x = x[np.isfinite(x)]
    delta = x - float(null)
    if not x.size:
        return {
            "mean": float("nan"),
            "bootstrap_95_CI_mean": [float("nan"), float("nan")],
            "n_positive": 0,
            "n_total": 0,
            "one_sided_exact_signflip_p": float("nan"),
            "participant_values": [],
        }
    rng = np.random.default_rng(BOOTSTRAP_SEED)
    index = rng.integers(0, x.size, size=(BOOTSTRAP_DRAWS, x.size))
    means = np.mean(x[index], axis=1)
    return {
        "mean": float(np.mean(x)),
        "bootstrap_95_CI_mean": [
            float(np.quantile(means, 0.025)),
            float(np.quantile(means, 0.975)),
        ],
        "n_positive": int(np.sum(delta > 0)),
        "n_total": int(x.size),
        "one_sided_exact_signflip_p": exact_signflip_mean_p(x, null),
        "participant_values": [float(value) for value in x],
    }


def kojima_scores(z, subject: str) -> tuple[dict, dict]:
    accepted = z["accepted"]
    mask = (
        (z["subject"] == subject)
        & (z["task"] == "2stream")
        & accepted
    )
    labels = z["label"][mask]
    scores = loro_event_scores(
        select_window_features(z["eeg_bins"][mask], (-0.1, 1.0)),
        select_window_features(z["eog_bins"][mask], (-0.1, 1.0)),
        select_window_features(z["eeg_bins"][mask], (-0.1, 0.0)),
        select_window_features(z["eeg_bins"][mask], (0.3, 0.8)),
        labels,
        z["run"][mask],
    )
    arrays = {
        "command": z["command"][mask],
        "trial": z["trial"][mask],
        "target": z["target"][mask],
    }
    return scores, arrays


def analyze_kojima(z, subject: str) -> dict:
    scores, a = kojima_scores(z, subject)
    common = {
        **a,
        "flat_score": scores["flat"],
        "commands": (1, 2, 3, 4),
        "correctness_groups": TRUE_PAIRING,
    }
    inclusive = row_metrics(
        trial_rows(
            **common,
            outer_score=scores["eeg_outer"],
            pooling_groups=TRUE_PAIRING,
            exclude_true_target=False,
        )
    )
    target_free = row_metrics(
        trial_rows(
            **common,
            outer_score=scores["eeg_outer"],
            pooling_groups=TRUE_PAIRING,
            exclude_true_target=True,
        )
    )
    eog = row_metrics(
        trial_rows(
            **common,
            outer_score=scores["eog_outer"],
            pooling_groups=TRUE_PAIRING,
            exclude_true_target=True,
        )
    )
    wrong = {}
    for pairing in WRONG_PAIRINGS:
        key = "-".join("".join(str(x) for x in group) for group in pairing)
        wrong[key] = row_metrics(
            trial_rows(
                **common,
                outer_score=scores["eeg_outer"],
                pooling_groups=pairing,
                exclude_true_target=True,
            )
        )
    best_wrong = max(value["certificate_AUC"] for value in wrong.values())
    return {
        "target_inclusive_EEG": inclusive,
        "target_free_EEG": target_free,
        "target_free_EOG": eog,
        "target_free_wrong_pairings": wrong,
        "contrasts": {
            "EEG_minus_EOG": (
                target_free["certificate_AUC"] - eog["certificate_AUC"]
            ),
            "EEG_minus_best_wrong": (
                target_free["certificate_AUC"] - best_wrong
            ),
            "target_free_minus_inclusive": (
                target_free["certificate_AUC"]
                - inclusive["certificate_AUC"]
            ),
        },
    }


def asme_scores(z, subject: int) -> tuple[dict, dict, dict]:
    accepted = z["accepted"]
    train = (
        (z["subject"] == subject)
        & (z["task"] == "offline")
        & accepted
    )
    test = (
        (z["subject"] == subject)
        & (z["task"] == "online")
        & accepted
    )
    train_eeg = select_bins(z["eeg_bins"][train], (0.0, 1.0))
    test_eeg = select_bins(z["eeg_bins"][test], (0.0, 1.0))
    train_eog = select_bins(z["eog_bins"][train], (0.0, 1.0))
    test_eog = select_bins(z["eog_bins"][test], (0.0, 1.0))
    train_flat = select_bins(z["eeg_bins"][train], (0.3, 0.8))
    test_flat = select_bins(z["eeg_bins"][test], (0.3, 0.8))
    scores, fit_audit = fit_primary_event_scores(
        train_eeg_outer=train_eeg,
        train_eog_outer=train_eog,
        train_prestim=z["prestim_raw"][train],
        train_flat=train_flat,
        train_command=z["command"][train],
        train_target=z["target"][train],
        test_eeg_outer=test_eeg,
        test_eog_outer=test_eog,
        test_prestim=z["prestim_raw"][test],
        test_flat=test_flat,
    )
    wrong = {}
    wrong_audit = {}
    for index, groups in enumerate(WRONG_GROUPS, start=1):
        wrong[str(index)], wrong_audit[str(index)] = fit_wrong_mapping_score(
            train_eeg,
            z["command"][train],
            z["target"][train],
            test_eeg,
            groups,
        )
    arrays = {
        "command": z["command"][test],
        "trial": z["trial"][test],
        "target": z["target"][test],
    }
    return scores, wrong, {
        "arrays": arrays,
        "fit_audit": fit_audit,
        "wrong_fit_audit": wrong_audit,
    }


def analyze_asme(z, subject: int) -> dict:
    scores, wrong_scores, extra = asme_scores(z, subject)
    common = {
        **extra["arrays"],
        "flat_score": scores["flat"],
        "commands": tuple(range(1, 31)),
        "correctness_groups": PHYSICAL_GROUPS,
    }
    inclusive = row_metrics(
        trial_rows(
            **common,
            outer_score=scores["eeg_outer"],
            pooling_groups=PHYSICAL_GROUPS,
            exclude_true_target=False,
        )
    )
    target_free = row_metrics(
        trial_rows(
            **common,
            outer_score=scores["eeg_outer"],
            pooling_groups=PHYSICAL_GROUPS,
            exclude_true_target=True,
        )
    )
    eog = row_metrics(
        trial_rows(
            **common,
            outer_score=scores["eog_outer"],
            pooling_groups=PHYSICAL_GROUPS,
            exclude_true_target=True,
        )
    )
    wrong = {}
    for index, groups in enumerate(WRONG_GROUPS, start=1):
        wrong[str(index)] = row_metrics(
            trial_rows(
                **common,
                outer_score=wrong_scores[str(index)],
                pooling_groups=groups,
                exclude_true_target=True,
            )
        )
    best_wrong = max(value["certificate_AUC"] for value in wrong.values())
    return {
        "target_inclusive_EEG": inclusive,
        "target_free_EEG": target_free,
        "target_free_EOG": eog,
        "target_free_wrong_mappings": wrong,
        "contrasts": {
            "EEG_minus_EOG": (
                target_free["certificate_AUC"] - eog["certificate_AUC"]
            ),
            "EEG_minus_best_wrong": (
                target_free["certificate_AUC"] - best_wrong
            ),
            "target_free_minus_inclusive": (
                target_free["certificate_AUC"]
                - inclusive["certificate_AUC"]
            ),
        },
        "fit_audit": extra["fit_audit"],
        "wrong_fit_audit": extra["wrong_fit_audit"],
    }


def evaluable(participant: dict) -> bool:
    metric = participant["target_free_EEG"]
    return (
        metric["n_trials"] >= 12
        and metric["n_parent_correct"] > 0
        and metric["n_parent_error"] > 0
        and np.isfinite(metric["certificate_AUC"])
    )


def dataset_summaries(participants: dict, ids: list[str]) -> dict:
    return {
        "target_free_absolute": summarize(
            [
                participants[s]["target_free_EEG"]["certificate_AUC"]
                for s in ids
            ],
            null=0.5,
        ),
        "mapping_specificity": summarize(
            [
                participants[s]["contrasts"]["EEG_minus_best_wrong"]
                for s in ids
            ]
        ),
        "nonocular": summarize(
            [
                participants[s]["contrasts"]["EEG_minus_EOG"]
                for s in ids
            ]
        ),
        "target_free_minus_inclusive": summarize(
            [
                participants[s]["contrasts"][
                    "target_free_minus_inclusive"
                ]
                for s in ids
            ]
        ),
    }


def main() -> None:
    if RESULT.exists():
        raise FileExistsError(RESULT)
    lock = validate_lock()
    expected_hashes = {
        KOJIMA_FEATURES: lock["input_sha256"]["kojima_features"],
        ASME_FEATURES: lock["input_sha256"]["asme_features"],
    }
    for path, expected in expected_hashes.items():
        observed = sha256(path)
        if observed != expected:
            raise RuntimeError(
                f"Input hash mismatch for {path}: {observed} != {expected}"
            )
    semantic_inputs = {
        KOJIMA_PRIMARY: lock["input_semantic_sha256"][
            "kojima_primary_result"
        ],
        ASME_PRIMARY: lock["input_semantic_sha256"]["asme_primary_result"],
    }
    for path, expected in semantic_inputs.items():
        observed = semantic_json_sha256(path)
        if observed != expected:
            raise RuntimeError(
                f"Scientific input mismatch for {path}: {observed} != {expected}"
            )

    z_kojima = np.load(KOJIMA_FEATURES, allow_pickle=False)
    kojima = {}
    for subject in KOJIMA_SUBJECTS:
        kojima[subject] = analyze_kojima(z_kojima, subject)
        print(
            f"Kojima {subject}: inclusive="
            f"{kojima[subject]['target_inclusive_EEG']['certificate_AUC']:.6f} "
            f"target_free="
            f"{kojima[subject]['target_free_EEG']['certificate_AUC']:.6f}",
            flush=True,
        )

    z_asme = np.load(ASME_FEATURES, allow_pickle=False)
    asme = {}
    for subject in range(1, 11):
        key = str(subject)
        asme[key] = analyze_asme(z_asme, subject)
        print(
            f"ASME {subject}: inclusive="
            f"{asme[key]['target_inclusive_EEG']['certificate_AUC']:.6f} "
            f"target_free="
            f"{asme[key]['target_free_EEG']['certificate_AUC']:.6f}",
            flush=True,
        )

    kojima_ids = [s for s in KOJIMA_SUBJECTS if evaluable(kojima[s])]
    asme_ids = [str(s) for s in range(1, 11) if evaluable(asme[str(s)])]
    ks = dataset_summaries(kojima, kojima_ids)
    ass = dataset_summaries(asme, asme_ids)

    kojima_primary = json.loads(KOJIMA_PRIMARY.read_text(encoding="utf-8"))
    asme_primary = json.loads(ASME_PRIMARY.read_text(encoding="utf-8"))
    equivalence_errors = {
        "Kojima2024B": {
            subject: abs(
                kojima[subject]["target_inclusive_EEG"]["certificate_AUC"]
                - kojima_primary["participant_results_2stream"][subject][
                    "EEG_true"
                ]["certificate_AUC"]
            )
            for subject in KOJIMA_SUBJECTS
        },
        "ASME30": {
            subject: abs(
                asme[subject]["target_inclusive_EEG"]["certificate_AUC"]
                - asme_primary["participants"][subject]["metrics"][
                    "EEG_physical"
                ]["certificate_AUC"]
            )
            for subject in map(str, range(1, 11))
        },
    }
    max_equivalence_error = max(
        value
        for dataset in equivalence_errors.values()
        for value in dataset.values()
        if np.isfinite(value)
    )

    gates = {
        "T1_Kojima_target_free_absolute": (
            ks["target_free_absolute"]["mean"] >= 0.75
            and ks["target_free_absolute"]["n_positive"] >= 6
            and ks["target_free_absolute"]["n_total"] == 7
            and ks["target_free_absolute"]["one_sided_exact_signflip_p"]
            <= 0.05
        ),
        "T2_Kojima_mapping_specificity": (
            ks["mapping_specificity"]["mean"] >= 0.15
            and ks["mapping_specificity"]["n_positive"] >= 6
            and ks["mapping_specificity"]["one_sided_exact_signflip_p"]
            <= 0.05
        ),
        "T3_Kojima_nonocular": (
            ks["nonocular"]["mean"] >= 0.05
            and ks["nonocular"]["n_positive"] >= 6
            and ks["nonocular"]["one_sided_exact_signflip_p"] <= 0.05
        ),
        "T4_ASME_target_free_absolute": (
            ass["target_free_absolute"]["mean"] >= 0.80
            and ass["target_free_absolute"]["n_positive"] >= 8
            and ass["target_free_absolute"]["n_total"] == 9
            and ass["target_free_absolute"]["one_sided_exact_signflip_p"]
            <= 0.05
        ),
        "T5_ASME_mapping_specificity": (
            ass["mapping_specificity"]["mean"] >= 0.15
            and ass["mapping_specificity"]["n_positive"] >= 8
            and ass["mapping_specificity"]["one_sided_exact_signflip_p"]
            <= 0.05
        ),
        "T6_ASME_nonocular": (
            ass["nonocular"]["mean"] >= 0.10
            and ass["nonocular"]["n_positive"] >= 8
            and ass["nonocular"]["one_sided_exact_signflip_p"] <= 0.05
        ),
        "implementation_equivalence": max_equivalence_error <= 1e-10,
    }
    absolute_pass = (
        gates["T1_Kojima_target_free_absolute"]
        and gates["T4_ASME_target_free_absolute"]
    )
    all_mechanism_gates = all(gates.values())
    if all_mechanism_gates:
        status = "target_free_cross_dataset_mechanism_supported"
    elif absolute_pass:
        status = "target_free_absolute_supported_specificity_incomplete"
    else:
        status = "target_free_mechanism_not_supported_no_retuning"

    result = {
        "route": "WZ-HIC-target-free-mechanism-v1",
        "status": status,
        "created_utc": datetime.now(timezone.utc).isoformat(),
        "all_mechanism_gates_pass": all_mechanism_gates,
        "gates": gates,
        "evaluable_participants": {
            "Kojima2024B": kojima_ids,
            "ASME30": asme_ids,
        },
        "summaries": {
            "Kojima2024B": ks,
            "ASME30": ass,
        },
        "participants": {
            "Kojima2024B": kojima,
            "ASME30": asme,
        },
        "implementation_equivalence": {
            "maximum_absolute_AUC_error": max_equivalence_error,
            "participant_errors": equivalence_errors,
        },
        "provenance": {
            "protocol_sha256": sha256(PROTOCOL),
            "implementation_lock_sha256": sha256(LOCK),
            "kojima_features_sha256": sha256(KOJIMA_FEATURES),
            "asme_features_sha256": sha256(ASME_FEATURES),
            "kojima_primary_result_sha256": sha256(KOJIMA_PRIMARY),
            "asme_primary_result_sha256": sha256(ASME_PRIMARY),
        },
        "implementation_lock": lock,
        "execution_incidents": {
            "first_locked_run": (
                "Stopped on an under-horizon trial; partial B/D/F/H output "
                "was recorded in engineering amendment 01."
            ),
            "second_locked_run": (
                "Completed all scientific AUCs, then stopped on an archived "
                "JSON lookup key; all visible output was recorded in "
                "engineering amendment 02."
            ),
            "final_execution_label": (
                "All scientific AUC outcomes were visible before the v1b "
                "archived-key repair; no scientific rule changed."
            ),
        },
        "interpretation_boundary": (
            "The true target was removed with oracle labels only to isolate "
            "non-target stream-selection evidence. This is a post-primary "
            "frozen mechanism audit, not a deployable online score or an "
            "independent new-dataset confirmation."
        ),
    }
    RESULT.parent.mkdir(parents=True, exist_ok=True)
    RESULT.write_text(json.dumps(result, indent=2) + "\n", encoding="utf-8")
    print(json.dumps({"status": status, "gates": gates}, indent=2))
    print(f"result={RESULT} sha256={sha256(RESULT)}")


if __name__ == "__main__":
    main()
```
