## Supplementary material for "Falsifiable substitution tests reveal task-structured neural evidence for auditory attention": Article_Analysis_Source_Code: summarize_kite_st_hd_confirmation_complete_case_preprocessing_v2.py

```
"""Freeze the label-blind 12-pass/2-technical-failure confirmation disposition."""

from __future__ import annotations

import json
from pathlib import Path

from kite_sat_confirmation_lock_utils_v1 import ROOT, sha256, validate_confirmation_lock


ORIGINAL_SUBJECT_SCRIPT = (
    ROOT / "analysis" / "run_kite_sat_akb_confirmation_subject_v1.py"
)
ORIGINAL_ORDER = (
    "S0", "S10", "S26", "S2", "S21", "S28", "S22",
    "S9", "S3", "S15", "S16", "S18", "S13", "S1",
)
PASS_ORDER = (
    "S0", "S10", "S2", "S21", "S28", "S22",
    "S9", "S15", "S16", "S18", "S13", "S1",
)
FAILURES = {"S26": 16, "S3": 19}
SUBJECT_ROOT = ROOT / "results" / "kite_st_hd" / "confirmation_v1" / "subjects"
OUTPUT = (
    ROOT / "results" / "kite_st_hd" / "confirmation_complete_case_v2"
    / "preprocessing_disposition_v2.json"
)


def validate_sidecar(path: Path) -> str:
    fields = path.with_name(path.name + ".sha256").read_text(
        encoding="ascii"
    ).strip().split()
    observed = sha256(path)
    if len(fields) != 2 or fields[0].lower() != observed or fields[1] != path.name:
        raise RuntimeError(f"Invalid SHA256 sidecar: {path}")
    return observed


def main() -> None:
    if OUTPUT.exists():
        raise RuntimeError("Refusing to overwrite complete-case preprocessing disposition.")
    lock, lock_hash = validate_confirmation_lock(ORIGINAL_SUBJECT_SCRIPT)
    if tuple(lock["confirmation_subject_order"]) != ORIGINAL_ORDER:
        raise RuntimeError("Original confirmation order mismatch.")
    if set(PASS_ORDER) | set(FAILURES) != set(ORIGINAL_ORDER):
        raise RuntimeError("Complete-case disposition does not cover the original cohort.")
    forbidden_subject_root = (
        ROOT / "results" / "kite_sat" / "akb_confirmation_v1" / "subjects"
    )
    forbidden_group = (
        ROOT / "results" / "kite_sat" / "akb_confirmation_v1"
        / "confirmation_decision_v1.json"
    )
    if (
        forbidden_subject_root.exists()
        and any(forbidden_subject_root.glob("*.json"))
    ) or forbidden_group.exists():
        raise RuntimeError("A confirmation attention outcome exists before disposition freeze.")
    pass_records = []
    for subject in PASS_ORDER:
        path = SUBJECT_ROOT / f"{subject}_production_manifest_v1.json"
        manifest_hash = validate_sidecar(path)
        manifest = json.loads(path.read_text(encoding="utf-8"))
        if (
            manifest["schema"] != "KITE-ST-HD-confirmation-subject-preprocessing-v1"
            or manifest["status"] != "pass"
            or manifest["subject"] != subject
            or manifest["maximum_bad_channel_count"] > 15
            or manifest["confirmation_implementation_lock_sha256"] != lock_hash
            or manifest["firewall"]["attention_contrast_computed"]
            or manifest["firewall"]["AKB_score_computed"]
        ):
            raise RuntimeError(f"Invalid complete-case pass manifest: {subject}")
        output_hashes = {}
        for record in manifest["preprocessing_records"]:
            output_path = ROOT / record["output"]["path"]
            if (
                output_path.stat().st_size != record["output"]["bytes"]
                or sha256(output_path) != record["output"]["sha256"]
            ):
                raise RuntimeError(f"Preprocessed output mismatch: {output_path}")
            output_hashes[record["recording"]] = record["output"]["sha256"]
        feature = manifest["feature_archive"]
        feature_path = ROOT / feature["path"]
        if (
            feature_path.stat().st_size != feature["bytes"]
            or sha256(feature_path) != feature["sha256"]
            or len(output_hashes) != 4
        ):
            raise RuntimeError(f"Feature/output mismatch: {subject}")
        pass_records.append(
            {
                "subject": subject,
                "maximum_bad_channel_count": manifest["maximum_bad_channel_count"],
                "manifest_path": path.relative_to(ROOT).as_posix(),
                "manifest_sha256": manifest_hash,
                "preprocessed_sha256": output_hashes,
                "feature_path": feature["path"],
                "feature_sha256": feature["sha256"],
            }
        )
    failure_records = []
    for subject, bad_count in FAILURES.items():
        path = SUBJECT_ROOT / f"{subject}_technical_failure_v1.json"
        failure_hash = validate_sidecar(path)
        record = json.loads(path.read_text(encoding="utf-8"))
        observed_bad_count = record.get("bad_channel_count")
        observed_maximum = record.get("maximum_allowed")
        if observed_bad_count is None:
            qc_events = [
                event
                for event in record.get("events", [])
                if "bad_channel_count" in event
            ]
            if len(qc_events) != 1:
                raise RuntimeError(f"Missing unique QC event: {subject}")
            observed_bad_count = qc_events[0]["bad_channel_count"]
            observed_maximum = qc_events[0]["maximum_allowed"]
        if (
            record["status"] != "technical_failure"
            or record["subject"] != subject
            or observed_bad_count != bad_count
            or observed_maximum != 15
            or record["scientific_outputs"]["attention_contrast_computed"]
            or record["scientific_outputs"]["global_AUC_computed"]
            or record["scientific_outputs"]["AKB_AUC_computed"]
        ):
            raise RuntimeError(f"Invalid technical-failure record: {subject}")
        failure_records.append(
            {
                "subject": subject,
                "bad_channel_count": bad_count,
                "failure_path": path.relative_to(ROOT).as_posix(),
                "failure_sha256": failure_hash,
            }
        )
    payload = {
        "schema": "KITE-ST-HD-confirmation-complete-case-preprocessing-disposition-v2",
        "status": "pass",
        "timing": "after_all_label_blind_QC_and_before_any_confirmation_attention_or_AKB_score",
        "original_confirmation_implementation_lock_sha256": lock_hash,
        "original_subject_order": list(ORIGINAL_ORDER),
        "pass_subject_order": list(PASS_ORDER),
        "pass_count": len(PASS_ORDER),
        "technical_failures": failure_records,
        "technical_failure_count": len(failure_records),
        "pass_records": pass_records,
        "replacement_subjects": [],
        "QC_threshold_changed": False,
        "attention_outcome_computed": False,
        "AKB_score_computed": False,
        "original_v1_decision_remains": "technical_failure",
        "paper_mutated": False,
    }
    OUTPUT.parent.mkdir(parents=True, exist_ok=True)
    OUTPUT.write_text(json.dumps(payload, indent=2) + "\n", encoding="utf-8")
    observed = sha256(OUTPUT)
    OUTPUT.with_name(OUTPUT.name + ".sha256").write_text(
        f"{observed}  {OUTPUT.name}\n", encoding="ascii"
    )
    print(
        json.dumps(
            {
                "status": "pass",
                "pass_count": len(PASS_ORDER),
                "failure_count": len(failure_records),
                "sha256": observed,
            }
        )
    )


if __name__ == "__main__":
    main()
```
