## Supplementary material for "Falsifiable substitution tests reveal task-structured neural evidence for auditory attention": Article_Analysis_Source_Code: summarize_kite_st_hd_confirmation_preprocessing_v1.py

```
"""Verify that all 14 frozen confirmation preprocessing outputs are ready."""

from __future__ import annotations

import json
from pathlib import Path

from download_kite_st_hd_confirmation_v1 import SUBJECTS
from kite_sat_confirmation_lock_utils_v1 import ROOT, sha256, validate_confirmation_lock


RESULT_ROOT = ROOT / "results" / "kite_st_hd" / "confirmation_v1"
OUTPUT = RESULT_ROOT / "confirmation_preprocessing_ready_v1.json"


def main() -> None:
    if OUTPUT.exists():
        raise RuntimeError("Refusing to overwrite confirmation preprocessing-ready record.")
    lock, lock_hash = validate_confirmation_lock(Path(__file__))
    if tuple(lock["confirmation_subject_order"]) != SUBJECTS:
        raise RuntimeError("Confirmation subjects differ from lock.")
    subjects = []
    for subject in SUBJECTS:
        manifest_path = RESULT_ROOT / "subjects" / f"{subject}_production_manifest_v1.json"
        manifest = json.loads(manifest_path.read_text(encoding="utf-8"))
        if (
            manifest["schema"] != "KITE-ST-HD-confirmation-subject-preprocessing-v1"
            or manifest["status"] != "pass"
            or manifest["subject"] != subject
            or manifest["maximum_bad_channel_count"] > 15
            or manifest["confirmation_implementation_lock_sha256"] != lock_hash
            or manifest["firewall"]["attention_contrast_computed"]
            or manifest["firewall"]["AKB_score_computed"]
        ):
            raise RuntimeError(f"Invalid confirmation preprocessing manifest: {subject}")
        outputs = {}
        for record in manifest["preprocessing_records"]:
            path = ROOT / record["output"]["path"]
            if (
                path.stat().st_size != record["output"]["bytes"]
                or sha256(path) != record["output"]["sha256"]
            ):
                raise RuntimeError(f"Preprocessed output mismatch: {path}")
            outputs[record["recording"]] = record["output"]["sha256"]
        feature = manifest["feature_archive"]
        feature_path = ROOT / feature["path"]
        if (
            feature_path.stat().st_size != feature["bytes"]
            or sha256(feature_path) != feature["sha256"]
        ):
            raise RuntimeError(f"Feature archive mismatch: {subject}")
        subjects.append(
            {
                "subject": subject,
                "manifest_path": manifest_path.relative_to(ROOT).as_posix(),
                "manifest_sha256": sha256(manifest_path),
                "maximum_bad_channel_count": manifest["maximum_bad_channel_count"],
                "preprocessed_sha256": outputs,
                "feature_archive_sha256": feature["sha256"],
            }
        )
    payload = {
        "schema": "KITE-ST-HD-confirmation-preprocessing-ready-v1",
        "status": "pass",
        "confirmation_implementation_lock_sha256": lock_hash,
        "subject_order": list(SUBJECTS),
        "subject_count": len(subjects),
        "subjects": subjects,
        "all_four_recordings_present": all(
            len(record["preprocessed_sha256"]) == 4 for record in subjects
        ),
        "attention_outcome_computed": False,
        "AKB_score_computed": False,
        "paper_mutated": False,
    }
    OUTPUT.write_text(json.dumps(payload, indent=2) + "\n", encoding="utf-8")
    observed = sha256(OUTPUT)
    OUTPUT.with_name(OUTPUT.name + ".sha256").write_text(
        f"{observed}  {OUTPUT.name}\n", encoding="ascii"
    )
    print(json.dumps({"status": "pass", "subjects": len(subjects), "sha256": observed}))


if __name__ == "__main__":
    main()
```
