## Supplementary material for "Falsifiable substitution tests reveal task-structured neural evidence for auditory attention": Article_Analysis_Source_Code: synthesize_multimodal_locked_results.py

```
"""Synthesize three independent, hash-locked projection-breadth tests.

The Ear-SAAD primary endpoint is one value per participant, averaged equally
across scalp, around-ear, and in-ear EEG.  Sensor modalities are therefore not
treated as independent replications.
"""

from __future__ import annotations

import argparse
import csv
import json
from pathlib import Path

import numpy as np
from scipy.stats import combine_pvalues


SEED = 20260728
BOOTSTRAPS = 100_000


def read_csv(path: Path) -> list[dict[str, str]]:
    with path.open("r", encoding="utf-8", newline="") as handle:
        return list(csv.DictReader(handle))


def write_csv(path: Path, rows: list[dict[str, object]]) -> None:
    with path.open("w", encoding="utf-8", newline="") as handle:
        writer = csv.DictWriter(handle, fieldnames=list(rows[0]))
        writer.writeheader()
        writer.writerows(rows)


def bootstrap_study_means(
    rng: np.random.Generator, values: np.ndarray
) -> np.ndarray:
    indices = rng.integers(
        0, values.size, size=(BOOTSTRAPS, values.size)
    )
    return values[indices].mean(axis=1)


def interval(values: np.ndarray) -> list[float]:
    return [
        float(np.quantile(values, 0.025)),
        float(np.quantile(values, 0.975)),
    ]


def main() -> None:
    parser = argparse.ArgumentParser()
    parser.add_argument("--results-root", type=Path, default=Path("results"))
    parser.add_argument(
        "--output-dir",
        type=Path,
        default=Path("results/multimodal_projection_breadth_synthesis"),
    )
    args = parser.parse_args()

    kul_json = json.loads(
        (
            args.results_root
            / "kul_aad/locked_confirmation/locked_confirmation.json"
        ).read_text(encoding="utf-8")
    )
    dtu_json = json.loads(
        (
            args.results_root
            / "dtu_aad/locked_external/locked_external.json"
        ).read_text(encoding="utf-8")
    )
    ear_json = json.loads(
        (
            args.results_root
            / "ear_saad/locked_multimodal_external/locked_external.json"
        ).read_text(encoding="utf-8")
    )

    study_specs = [
        {
            "study": "KUL even-ID locked confirmation",
            "dataset": "KUL",
            "measurement_geometry": "scalp EEG",
            "path": (
                args.results_root
                / "kul_aad/locked_confirmation/participants.csv"
            ),
            "gain_column": "equal_k4_gain_vs_dominant_k1",
            "candidate_column": "equal_k4_accuracy",
            "baseline_column": "dominant_k1_accuracy",
            "p": float(
                kul_json["primary_result"][
                    "exact_one_sided_sign_flip_p_vs_dominant_k1"
                ]
            ),
            "lock_sha256": kul_json["candidate_lock_sha256"],
        },
        {
            "study": "DTU locked external validation",
            "dataset": "DTU",
            "measurement_geometry": "scalp EEG",
            "path": (
                args.results_root
                / "dtu_aad/locked_external/participants.csv"
            ),
            "gain_column": "equal_k4_gain_vs_dominant_k1",
            "candidate_column": "equal_k4_accuracy",
            "baseline_column": "dominant_k1_accuracy",
            "p": float(
                dtu_json["locked_primary_inference"][
                    "exact_one_sided_sign_flip_p"
                ]
            ),
            "lock_sha256": dtu_json["lock_sha256"],
        },
        {
            "study": "Ear-SAAD locked multimodal external validation",
            "dataset": "Ear-SAAD",
            "measurement_geometry": (
                "participant-equal mean over simultaneous scalp, "
                "around-ear, and in-ear EEG"
            ),
            "path": (
                args.results_root
                / "ear_saad/locked_multimodal_external/"
                "primary_participants.csv"
            ),
            "gain_column": "gain_equal_minus_k1",
            "candidate_column": "equal_k4_accuracy",
            "baseline_column": "k1_accuracy",
            "p": float(
                ear_json["primary_equal_modality_average"][
                    "exact_one_sided_p"
                ]
            ),
            "lock_sha256": ear_json["lock_sha256"],
        },
    ]

    rng = np.random.default_rng(SEED)
    gains: list[np.ndarray] = []
    boot_means: list[np.ndarray] = []
    study_rows: list[dict[str, object]] = []
    p_values: list[float] = []
    sample_sizes: list[int] = []

    for spec in study_specs:
        rows = read_csv(spec["path"])
        gain = np.asarray(
            [float(row[spec["gain_column"]]) for row in rows]
        )
        candidate = np.asarray(
            [float(row[spec["candidate_column"]]) for row in rows]
        )
        baseline = np.asarray(
            [float(row[spec["baseline_column"]]) for row in rows]
        )
        gains.append(gain)
        boot_means.append(bootstrap_study_means(rng, gain))
        p_values.append(float(spec["p"]))
        sample_sizes.append(int(gain.size))
        study_rows.append(
            {
                "study": spec["study"],
                "dataset": spec["dataset"],
                "measurement_geometry": spec["measurement_geometry"],
                "participants": int(gain.size),
                "baseline_k1_accuracy": float(baseline.mean()),
                "equal_k4_accuracy": float(candidate.mean()),
                "equal_k4_minus_k1": float(gain.mean()),
                "participants_improved": int(np.sum(gain > 0)),
                "exact_one_sided_sign_flip_p": float(spec["p"]),
                "lock_sha256": spec["lock_sha256"],
            }
        )

    sizes = np.asarray(sample_sizes)
    study_equal_boot = np.mean(np.vstack(boot_means), axis=0)
    participant_weighted_boot = np.average(
        np.vstack(boot_means), axis=0, weights=sizes
    )
    all_gains = np.concatenate(gains)
    study_effects = np.asarray([gain.mean() for gain in gains])
    fisher_statistic, fisher_p = combine_pvalues(
        p_values, method="fisher"
    )
    stouffer_statistic, stouffer_p = combine_pvalues(
        p_values, method="stouffer", weights=np.sqrt(sizes)
    )

    report = {
        "claim": (
            "Using four neural-speech projections rather than only the "
            "dominant projection improves five-second auditory-attention "
            "decoding across independent scalp and ear-EEG datasets."
        ),
        "unit_of_replication": (
            "Participant within independently hash-locked datasets. "
            "Ear-SAAD contributes one participant-level endpoint obtained "
            "by equal averaging across its three simultaneous modalities."
        ),
        "studies": study_rows,
        "independent_locked_synthesis": {
            "number_of_studies": len(study_rows),
            "participants_total": int(all_gains.size),
            "participants_improved_total": int(np.sum(all_gains > 0)),
            "study_equal_mean_gain": float(study_effects.mean()),
            "study_equal_stratified_bootstrap_95": interval(
                study_equal_boot
            ),
            "participant_weighted_mean_gain": float(all_gains.mean()),
            "participant_weighted_stratified_bootstrap_95": interval(
                participant_weighted_boot
            ),
            "fisher_statistic": float(fisher_statistic),
            "fisher_combined_one_sided_p": float(fisher_p),
            "sample_size_weighted_stouffer_z": float(
                stouffer_statistic
            ),
            "sample_size_weighted_stouffer_one_sided_p": float(
                stouffer_p
            ),
            "replicability_conjunction_p": float(max(p_values)),
            "all_three_study_effects_positive": bool(
                np.all(study_effects > 0)
            ),
        },
        "interpretive_guardrails": [
            (
                "The synthesis is secondary because it was specified only "
                "after completion of the Ear-SAAD external test."
            ),
            (
                "The three Ear-SAAD sensor modalities are repeated "
                "measurements, not independent samples."
            ),
            (
                "The claim concerns projection breadth; equal versus "
                "canonical weighting is not claimed to be universal."
            ),
            (
                "Separate locked tests in feedback and attention-switching "
                "settings did not support a universal dynamic extension."
            ),
        ],
        "bootstrap_seed": SEED,
        "bootstrap_replicates": BOOTSTRAPS,
    }

    args.output_dir.mkdir(parents=True, exist_ok=True)
    write_csv(args.output_dir / "studies.csv", study_rows)
    (args.output_dir / "synthesis.json").write_text(
        json.dumps(report, indent=2), encoding="utf-8"
    )
    print(json.dumps(report, indent=2))


if __name__ == "__main__":
    main()
```
