## Supplementary material for "Falsifiable substitution tests reveal task-structured neural evidence for auditory attention": Article_Analysis_Source_Code: test_kite_sat_direction_delay_controls_v1.py

```
"""Unit tests for the fixed AKB direction-delay control group."""

from __future__ import annotations

import numpy as np

from kite_sat_direction_delay_controls_v1 import TRANSFORMS, transformed_matrix


def main() -> None:
    values = np.arange(16, dtype=np.float64)
    transformed = transformed_matrix(values)
    if transformed.shape != (16, 16):
        raise RuntimeError("Transformation matrix shape failed.")
    if not np.array_equal(transformed[0], values) or TRANSFORMS[0] != (0, 0):
        raise RuntimeError("Identity transformation is not first.")
    if len({tuple(row) for row in transformed}) != 16:
        raise RuntimeError("Direction-delay transformations are not unique.")
    if not all(np.array_equal(np.sort(row), values) for row in transformed):
        raise RuntimeError("A transformation did not preserve the value histogram.")
    # Two angular groups crossed with the delay swap must be closed.
    observed = {tuple(row) for row in transformed}
    for first in transformed:
        nested = transformed_matrix(first)
        if not all(tuple(row) in observed for row in nested):
            raise RuntimeError("Direction-delay transformations are not closed.")
    print("KITE-SAT direction-delay control tests passed.")


if __name__ == "__main__":
    main()
```
