## Supplementary material for "Falsifiable substitution tests reveal task-structured neural evidence for auditory attention": Article_Analysis_Source_Code: test_kite_sat_ear_saad_external_v1.py

```
#!/usr/bin/env python3
"""Synthetic unit tests for the Ear-SAAD KITE-SAT external engine."""

from __future__ import annotations

import numpy as np

import run_kite_sat_ear_saad_external_subject_v1 as engine


def direct_fit(designs: list[np.ndarray], targets: list[np.ndarray]) -> dict:
    x = np.vstack(designs)
    y = np.concatenate(targets)
    x_mean = np.mean(x, axis=0)
    x_scale = np.std(x, axis=0)
    x_scale = np.where(x_scale > 1e-12, x_scale, 1.0)
    y_mean = float(np.mean(y))
    z = (x - x_mean) / x_scale
    gram = z.T @ z
    cross = z.T @ (y - y_mean)
    alpha = float(np.mean(np.diag(gram)))
    weights = np.linalg.solve(gram + alpha * np.eye(len(gram)), cross)
    return {
        "x_mean": x_mean,
        "x_scale": x_scale,
        "y_mean": y_mean,
        "weights": weights,
        "alpha": alpha,
    }


def main() -> None:
    rng = np.random.default_rng(20260729)
    records = []
    designs = []
    targets = []
    for _ in range(4):
        design = rng.normal(size=(engine.USABLE, engine.N_FEATURES))
        target = rng.normal(size=engine.USABLE)
        designs.append(design)
        targets.append(target)
        records.append(
            {"stats": engine.sufficient_statistics(design, target)}
        )
    efficient = engine.fit_from_blocks(records, list(range(4)))
    direct = direct_fit(designs, targets)
    for name in ("x_mean", "x_scale", "weights"):
        if not np.allclose(efficient[name], direct[name], atol=1e-10, rtol=0):
            raise AssertionError(f"Sufficient-statistic fit mismatch: {name}")
    if not np.isclose(efficient["y_mean"], direct["y_mean"], atol=1e-12):
        raise AssertionError("Target mean mismatch.")

    sectors = np.arange(engine.N_CHANNELS) % 8
    lag_index = np.repeat(np.arange(6), engine.N_CHANNELS)
    ids = np.tile(sectors, 6) + 8 * (lag_index >= 3)
    speaker1 = rng.normal(size=engine.USABLE)
    speaker2 = rng.normal(size=engine.USABLE)
    delta, margin = engine.cell_contributions(
        designs[0], efficient, speaker1, speaker2, ids
    )
    if delta.shape != (16,) or not np.isclose(np.sum(delta), margin, atol=1e-10):
        raise AssertionError("Cell additivity test failed.")

    positive = np.asarray([3.0, 2.0, 1.0])
    negative = np.asarray([-1.0, -2.0, -3.0])
    if engine.binary_auc(positive, negative) != 1.0:
        raise AssertionError("AUC orientation test failed.")
    labels = np.asarray([1, 1, 0, 0])
    scores = np.asarray([1.0, 2.0, -1.0, -2.0])
    if engine.balanced_accuracy(scores, labels) != 1.0:
        raise AssertionError("Balanced-accuracy orientation test failed.")

    print("Ear-SAAD KITE-SAT synthetic unit tests: PASS")


if __name__ == "__main__":
    main()
```
