## Supplementary material for "Falsifiable substitution tests reveal task-structured neural evidence for auditory attention": Article_Analysis_Source_Code: test_kite_sat_patch_erasure_v1.py

```
"""Synthetic equivalence tests for the KITE-SAT patch-erasure primitive."""

from __future__ import annotations

from pathlib import Path

import numpy as np

from kite_sat_geometry_v1 import cell_delta_contributions, partition_ids
from kite_sat_patch_erasure_v1 import erased_cell_delta_contributions
from kite_st_hd_core_v1 import load_public_geometry


ROOT = Path(__file__).resolve().parents[1]
COORDINATES = (
    ROOT
    / "downloads"
    / "candidate_audit"
    / "zenodo4518754_support"
    / "misc"
    / "misc"
    / "eeg255ch_locs.csv"
)


def main() -> None:
    rng = np.random.default_rng(20260729)
    geometry = load_public_geometry(COORDINATES)
    ids = partition_ids(geometry.xy)["S16"]
    design = rng.normal(size=(595, 1518))
    model = {
        "x_mean": rng.normal(scale=0.1, size=1518),
        "x_scale": rng.uniform(0.7, 1.4, size=1518),
        "weights": rng.normal(scale=0.03, size=1518),
        "y_mean": 0.0,
    }
    left = rng.normal(size=595)
    right = rng.normal(size=595)
    none = np.zeros((1, 253), dtype=bool)
    vectorized_delta, vectorized_margin = erased_cell_delta_contributions(
        design, model, left, right, ids, none
    )
    reference_delta, reference_margin = cell_delta_contributions(
        design, model, left, right, ids
    )
    if not np.allclose(vectorized_delta[0], reference_delta, atol=1e-12, rtol=0.0):
        raise RuntimeError("No-erasure cell equivalence failed.")
    if abs(vectorized_margin[0] - reference_margin) > 1e-12:
        raise RuntimeError("No-erasure global equivalence failed.")

    selected = np.sort(rng.choice(253, size=51, replace=False))
    patch = np.zeros((1, 253), dtype=bool)
    patch[0, selected] = True
    vectorized_delta, vectorized_margin = erased_cell_delta_contributions(
        design, model, left, right, ids, patch
    )
    brute_design = design.copy()
    feature_indices = np.concatenate([selected + lag * 253 for lag in range(6)])
    brute_design[:, feature_indices] = np.asarray(model["x_mean"])[feature_indices]
    reference_delta, reference_margin = cell_delta_contributions(
        brute_design, model, left, right, ids
    )
    if not np.allclose(vectorized_delta[0], reference_delta, atol=1e-12, rtol=0.0):
        raise RuntimeError("Patch-erasure cell equivalence failed.")
    if abs(vectorized_margin[0] - reference_margin) > 1e-12:
        raise RuntimeError("Patch-erasure global equivalence failed.")
    print("KITE-SAT patch-erasure equivalence tests passed.")


if __name__ == "__main__":
    main()
```
