## Supplementary material for "Falsifiable substitution tests reveal task-structured neural evidence for auditory attention": Article_Analysis_Source_Code: test_wz_hic_asme30_crossphase_individual_geometry_synthetic_v1.py

```
#!/usr/bin/env python3
"""Synthetic tests for target removal and cross-phase direction transport."""

from __future__ import annotations

import copy

import numpy as np

from run_wz_hic_asme30_crossphase_individual_geometry_v1 import (
    analyze_subject,
)
from wz_hic_asme30_core_v1 import PHYSICAL_GROUPS, group_index


def synthetic_arrays(target_spike: float) -> dict:
    rng = np.random.default_rng(20260730)
    rows = []
    for phase, n_trials, offset in (
        ("offline", 30, 0),
        ("online", 15, 30),
    ):
        for local_trial in range(n_trials):
            trial = offset + local_trial + 1
            target = (3 * local_trial + (0 if phase == "offline" else 1)) % 30 + 1
            target_parent = group_index(target, PHYSICAL_GROUPS)
            for command in range(1, 31):
                for repetition in range(3):
                    rows.append(
                        (phase, trial, target, command, repetition, target_parent)
                    )
    n = len(rows)
    n_channels = 6
    eeg = rng.normal(0, 0.7, size=(n, n_channels, 11))
    eog = rng.normal(0, 0.7, size=(n, 2, 11))
    direction = np.asarray([1.0, -0.7, 0.5, -0.3, 0.8, -0.4])
    time = np.asarray([0.0, 0.2, 0.5, 0.9, 1.2, 1.4, 1.2, 0.8, 0.4, 0.1])
    command = np.asarray([row[3] for row in rows], dtype=np.int8)
    target = np.asarray([row[2] for row in rows], dtype=np.int8)
    for index, row in enumerate(rows):
        command_value = row[3]
        target_value = row[2]
        same_parent = (
            group_index(command_value, PHYSICAL_GROUPS) == row[5]
            and command_value != target_value
        )
        if same_parent:
            eeg[index, :, 1:] += 1.2 * direction[:, None] * time[None, :]
            eog[index, :, 1:] += 0.08 * time[None, :]
        if command_value == target_value:
            eeg[index, :, 1:] += target_spike
            eog[index, :, 1:] += target_spike
    return {
        "eeg_bins": eeg.astype(np.float32),
        "eog_bins": eog.astype(np.float32),
        "subject": np.ones(n, dtype=np.int8),
        "task": np.asarray([row[0] for row in rows]),
        "trial": np.asarray([row[1] for row in rows], dtype=np.int16),
        "target": target,
        "command": command,
        "accepted": np.ones(n, dtype=bool),
    }


def outcome_vector(result: dict) -> np.ndarray:
    return np.asarray(
        [
            result["physical"]["full_effect"],
            result["wrong"][0]["full_effect"],
            result["wrong"][1]["full_effect"],
            result["eog_effect"],
            result["physical"]["alignment"],
            result["wrong"][0]["alignment"],
            result["wrong"][1]["alignment"],
            *result["physical"]["time_effects"],
            *result["wrong"][0]["time_effects"],
            *result["wrong"][1]["time_effects"],
        ],
        dtype=float,
    )


def main() -> None:
    baseline = synthetic_arrays(0.0)
    spiked = copy.deepcopy(baseline)
    is_target = spiked["command"] == spiked["target"]
    spiked["eeg_bins"][is_target, :, 1:] += 1_000_000.0
    spiked["eog_bins"][is_target, :, 1:] += 1_000_000.0
    result_baseline = analyze_subject(baseline, 1)
    result_spiked = analyze_subject(spiked, 1)
    influence = float(
        np.max(
            np.abs(outcome_vector(result_baseline) - outcome_vector(result_spiked))
        )
    )
    physical = result_baseline["physical"]["full_effect"]
    best_wrong = max(
        result_baseline["wrong"][0]["full_effect"],
        result_baseline["wrong"][1]["full_effect"],
    )
    if influence != 0.0:
        raise AssertionError(f"Target spike influenced analysis by {influence}.")
    if physical <= 0 or physical <= best_wrong:
        raise AssertionError(
            f"Synthetic physical effect {physical} did not exceed {best_wrong}."
        )
    if result_baseline["physical"]["alignment"] <= 0:
        raise AssertionError("Synthetic physical direction did not transport.")
    print(
        "PASS synthetic cross-phase individualized geometry; "
        "target-event influence is exactly zero."
    )


if __name__ == "__main__":
    main()
```
