## Supplementary material for "Falsifiable substitution tests reveal task-structured neural evidence for auditory attention": Article_Analysis_Source_Code: test_wz_hic_asme30_synthetic_v1.py

```
#!/usr/bin/env python3
"""Outcome-independent synthetic tests for the frozen ASME30 core."""

from __future__ import annotations

import json
from pathlib import Path

import numpy as np

from wz_hic_asme30_core_v1 import (
    COMMANDS,
    PHYSICAL_GROUPS,
    WRONG_GROUPS,
    certificate_metrics,
    fit_primary_event_scores,
    fit_wrong_mapping_score,
    group_index,
    labels_for_mapping,
    trial_rows,
)


ROOT = Path(__file__).resolve().parents[1]
OUT = ROOT / "results" / "wz_hic_asme30" / "synthetic_v1.json"


def make_events(rng, n_trials: int, signal: bool):
    command, target, trial, eeg, eog, prestim, flat = [], [], [], [], [], [], []
    for trial_id in range(n_trials):
        true = int(rng.integers(1, 31))
        true_parent = group_index(true)
        for repetition in range(5):
            for cmd in rng.permutation(COMMANDS):
                same_parent = group_index(int(cmd)) == true_parent
                is_target = int(cmd) == true
                # EEG feature 0 is the parent-level response; feature 1 is child.
                parent_mu = 1.5 if (signal and same_parent and not is_target) else 0.0
                # Deliberately make 30-way child decoding imperfect so that the
                # reliability AUC has both parent-correct and parent-error trials.
                child_mu = 1.1 if (signal and is_target) else 0.0
                eeg.append(
                    [
                        parent_mu + rng.normal(scale=0.7),
                        child_mu + rng.normal(scale=1.0),
                        rng.normal(),
                        rng.normal(),
                    ]
                )
                eog.append(rng.normal(size=2))
                prestim.append(rng.normal(size=3))
                flat.append(
                    [
                        child_mu + rng.normal(scale=1.0),
                        rng.normal(),
                        rng.normal(),
                    ]
                )
                command.append(cmd)
                target.append(true)
                trial.append(trial_id)
    return {
        "command": np.asarray(command),
        "target": np.asarray(target),
        "trial": np.asarray(trial),
        "eeg": np.asarray(eeg),
        "eog": np.asarray(eog),
        "prestim": np.asarray(prestim),
        "flat": np.asarray(flat),
    }


def main() -> None:
    rng = np.random.default_rng(20260729)
    train = make_events(rng, 40, True)
    test = make_events(rng, 30, True)
    labels = labels_for_mapping(train["command"], train["target"])
    assert dict(zip(*np.unique(labels, return_counts=True))) == {
        "S": 1800,
        "T": 200,
        "U": 4000,
    }
    scores, fits = fit_primary_event_scores(
        train_eeg_outer=train["eeg"],
        train_eog_outer=train["eog"],
        train_prestim=train["prestim"],
        train_flat=train["flat"],
        train_command=train["command"],
        train_target=train["target"],
        test_eeg_outer=test["eeg"],
        test_eog_outer=test["eog"],
        test_prestim=test["prestim"],
        test_flat=test["flat"],
    )
    true_rows = trial_rows(
        flat_score=scores["flat"],
        outer_score=scores["eeg_outer"],
        command=test["command"],
        trial=test["trial"],
        target=test["target"],
    )
    eog_rows = trial_rows(
        flat_score=scores["flat"],
        outer_score=scores["eog_outer"],
        command=test["command"],
        trial=test["trial"],
        target=test["target"],
    )
    wrong_metrics = []
    for groups in WRONG_GROUPS:
        wrong_score, _ = fit_wrong_mapping_score(
            train["eeg"],
            train["command"],
            train["target"],
            test["eeg"],
            groups,
        )
        wrong_metrics.append(
            certificate_metrics(
                trial_rows(
                    flat_score=scores["flat"],
                    outer_score=wrong_score,
                    command=test["command"],
                    trial=test["trial"],
                    target=test["target"],
                    pooling_groups=groups,
                    correctness_groups=PHYSICAL_GROUPS,
                )
            )
        )
    true_metrics = certificate_metrics(true_rows)
    eog_metrics = certificate_metrics(eog_rows)
    checks = {
        "30_trials": len(true_rows) == 30,
        "both_parent_outcomes": (
            true_metrics["n_parent_correct"] > 0
            and true_metrics["n_parent_error"] > 0
        ),
        "physical_certificate_high": true_metrics["certificate_AUC"] > 0.85,
        "physical_beats_eog": (
            true_metrics["certificate_AUC"] - eog_metrics["certificate_AUC"] > 0.2
        ),
        "physical_beats_wrong_maps": (
            true_metrics["certificate_AUC"]
            - max(item["certificate_AUC"] for item in wrong_metrics)
            > 0.2
        ),
        "training_only_scaling_recorded": all(
            item["n_train"] > 0 and item["training_score_sd"] > 0
            for item in fits.values()
        ),
    }
    result = {
        "status": "PASS" if all(checks.values()) else "FAIL",
        "checks": checks,
        "physical": true_metrics,
        "EOG": eog_metrics,
        "wrong_maps": wrong_metrics,
    }
    OUT.parent.mkdir(parents=True, exist_ok=True)
    OUT.write_text(json.dumps(result, indent=2) + "\n", encoding="utf-8")
    print(json.dumps(result, indent=2))
    if result["status"] != "PASS":
        raise SystemExit(1)


if __name__ == "__main__":
    main()
```
