## Supplementary material for "Falsifiable substitution tests reveal task-structured neural evidence for auditory attention": Article_Analysis_Source_Code: test_wz_hic_asme30_target_free_spatiotemporal_synthetic_v1.py

```
#!/usr/bin/env python3
"""Synthetic engineering test for the frozen scalp-time localization."""

from __future__ import annotations

import numpy as np

from run_wz_hic_asme30_target_free_spatiotemporal_v1 import run_analysis
from wz_hic_asme30_core_v1 import PHYSICAL_GROUPS, labels_for_mapping


def synthetic_arrays(target_spike: float) -> dict:
    rng = np.random.default_rng(20260730)
    rows = {
        key: []
        for key in (
            "eeg_bins",
            "subject",
            "task",
            "accepted",
            "command",
            "target",
            "trial",
        )
    }
    template = np.asarray(
        [
            [0.50, 1.10, 0.25],
            [-0.25, -0.70, -0.15],
            [0.15, 0.45, 0.10],
            [-0.10, -0.30, -0.05],
        ]
    )
    for subject in range(1, 9):
        participant_scale = 0.85 + 0.04 * subject
        for trial_index in range(12):
            target = 1 + ((trial_index * 7 + subject) % 30)
            commands = np.repeat(np.arange(1, 31), 3)
            targets = np.full(commands.size, target)
            labels = labels_for_mapping(commands, targets, PHYSICAL_GROUPS)
            for event_index, (command, label) in enumerate(
                zip(commands, labels)
            ):
                if label == "S":
                    signal = participant_scale * template
                elif label == "U":
                    signal = -0.20 * participant_scale * template
                else:
                    signal = target_spike * template
                noise = rng.normal(0, 0.08, size=template.shape)
                epoch = np.concatenate(
                    [np.zeros((template.shape[0], 1)), signal + noise],
                    axis=1,
                )
                rows["eeg_bins"].append(epoch)
                rows["subject"].append(subject)
                rows["task"].append("online")
                rows["accepted"].append(True)
                rows["command"].append(int(command))
                rows["target"].append(target)
                rows["trial"].append(subject * 100 + trial_index)
    return {
        "eeg_bins": np.asarray(rows["eeg_bins"], dtype=np.float32),
        "subject": np.asarray(rows["subject"], dtype=np.int16),
        "task": np.asarray(rows["task"], dtype="U7"),
        "accepted": np.asarray(rows["accepted"], dtype=bool),
        "command": np.asarray(rows["command"], dtype=np.int16),
        "target": np.asarray(rows["target"], dtype=np.int16),
        "trial": np.asarray(rows["trial"], dtype=np.int32),
    }


def main() -> None:
    base = run_analysis(synthetic_arrays(target_spike=0.0))
    spike = run_analysis(synthetic_arrays(target_spike=1000.0))
    assert base["gates"]["M0_coverage"]
    assert base["gates"]["M1_binding_specific_lopo"]
    assert base["gates"]["M2_spatiotemporal_fwer"]
    assert base["supported"]
    np.testing.assert_allclose(
        base["physical_maps"], spike["physical_maps"], atol=0, rtol=0
    )
    assert len(base["evaluable_subjects"]) == 8
    assert base["mapping_summary"]["n_positive"] == 8
    assert base["mapping_summary"]["bootstrap_95_CI_mean"][0] > 0
    assert np.min(base["p_fwer"]) <= 0.05
    print(
        "PASS synthetic target-free scalp-time localization; "
        "target spike has exactly zero influence."
    )


if __name__ == "__main__":
    main()
```
