## Supplementary material for "Falsifiable substitution tests reveal task-structured neural evidence for auditory attention": Article_Analysis_Source_Code: test_wz_hic_confirmation_synthetic_v1.py

```
#!/usr/bin/env python
"""Engineering-only synthetic test of the frozen WZ-HIC confirmation core."""

from __future__ import annotations

import hashlib
import json
from datetime import datetime, timezone
from pathlib import Path

import numpy as np

from wz_hic_confirmation_core_v1 import (
    TRUE_PAIRING,
    WRONG_PAIRINGS,
    certificate_metrics,
    loro_event_scores,
    trial_rows,
)


ROOT = Path(__file__).resolve().parents[1]
OUTPUT = ROOT / "results" / "wz_hic_kojima2024b" / "synthetic_confirmation_v1.json"


def sha256(path: Path) -> str:
    h = hashlib.sha256()
    h.update(path.read_bytes())
    return h.hexdigest()


def parent(command: int) -> int:
    return 1 if command in (1, 2) else 2


def main() -> None:
    rng = np.random.default_rng(7292026)
    rows = []
    for run in range(1, 7):
        for trial_order, target in enumerate(rng.permutation([1, 2, 3, 4]), start=1):
            trial = run * 10 + trial_order
            for command in (1, 2, 3, 4):
                for _ in range(15):
                    attended = parent(command) == parent(int(target))
                    label = (
                        "T"
                        if command == target
                        else "S"
                        if attended
                        else "U"
                    )
                    eeg_parent = (1.0 if attended else -1.0) + rng.normal(0, 1.2)
                    eog_parent = (0.22 if attended else -0.22) + rng.normal(0, 1.5)
                    flat_target = (
                        (0.5 if command == target else -0.1)
                        + rng.normal(0, 1.8)
                    )
                    nuisance = rng.normal(size=4)
                    rows.append(
                        (
                            run,
                            trial,
                            int(target),
                            command,
                            label,
                            np.r_[eeg_parent, nuisance],
                            np.r_[eog_parent, nuisance[:1]],
                            rng.normal(size=3),
                            np.r_[flat_target, nuisance],
                        )
                    )
    runs = np.asarray([r[0] for r in rows])
    trial = np.asarray([r[1] for r in rows])
    target = np.asarray([r[2] for r in rows])
    command = np.asarray([r[3] for r in rows])
    labels = np.asarray([r[4] for r in rows])
    score = loro_event_scores(
        np.stack([r[5] for r in rows]),
        np.stack([r[6] for r in rows]),
        np.stack([r[7] for r in rows]),
        np.stack([r[8] for r in rows]),
        labels,
        runs,
    )
    true = certificate_metrics(
        trial_rows(
            score,
            command,
            trial,
            target,
            horizon=3,
            certificate_name="eeg_outer",
        )
    )
    eog = certificate_metrics(
        trial_rows(
            score,
            command,
            trial,
            target,
            horizon=3,
            certificate_name="eog_outer",
        )
    )
    pre = certificate_metrics(
        trial_rows(
            score,
            command,
            trial,
            target,
            horizon=3,
            certificate_name="prestim_outer",
        )
    )
    wrong = [
        certificate_metrics(
            trial_rows(
                score,
                command,
                trial,
                target,
                horizon=3,
                certificate_name="eeg_outer",
                pooling_pairing=pairing,
                correctness_pairing=TRUE_PAIRING,
            )
        )
        for pairing in WRONG_PAIRINGS
    ]
    checks = {
        "all_24_trials": true["n_trials"] == 24,
        "both_parent_outcomes": (
            true["n_parent_correct"] > 0 and true["n_parent_error"] > 0
        ),
        "certificate_AUC_above_0.80": true["certificate_AUC"] > 0.80,
        "beats_flat_margin_by_0.10": (
            true["certificate_AUC"] - true["flat_margin_AUC"] > 0.10
        ),
        "beats_EOG_by_0.10": (
            true["certificate_AUC"] - eog["certificate_AUC"] > 0.10
        ),
        "beats_wrong_mapping_by_0.10": (
            true["certificate_AUC"]
            - max(item["certificate_AUC"] for item in wrong)
            > 0.10
        ),
        "prestim_not_strong": pre["certificate_AUC"] < 0.70,
    }
    result = {
        "status": "PASS" if all(checks.values()) else "FAIL",
        "created_utc": datetime.now(timezone.utc).isoformat(),
        "scope": "synthetic engineering test; no empirical EEG or behavior",
        "seed": 7292026,
        "true": true,
        "EOG": eog,
        "prestimulus": pre,
        "wrong_pairings": wrong,
        "checks": checks,
        "core_sha256": sha256(ROOT / "analysis" / "wz_hic_confirmation_core_v1.py"),
    }
    OUTPUT.parent.mkdir(parents=True, exist_ok=True)
    OUTPUT.write_text(
        json.dumps(result, ensure_ascii=False, indent=2) + "\n", encoding="utf-8"
    )
    print(json.dumps(result, ensure_ascii=False, indent=2))
    if result["status"] != "PASS":
        raise RuntimeError("Synthetic confirmation test failed.")


if __name__ == "__main__":
    main()
```
