## Supplementary material for "Falsifiable substitution tests reveal task-structured neural evidence for auditory attention": Article_Analysis_Source_Code: test_wz_hic_continuous_listener_transport_synthetic_v1.py

```
#!/usr/bin/env python3
"""Outcome-free synthetic checks for the frozen continuous-listener runner."""

from __future__ import annotations

import numpy as np

from run_wz_hic_continuous_listener_transport_v1 import (
    directional_endpoints,
    ear_split,
    pearson,
    rank_probability,
)


def main() -> None:
    rng = np.random.default_rng(20260730)
    participants = []
    for subject in range(5):
        base = rng.normal(size=16)
        base -= np.mean(base)
        profile_a = base + rng.normal(scale=0.01, size=16)
        profile_b = base + rng.normal(scale=0.01, size=16)
        weights = np.exp(2 * profile_a - np.max(2 * profile_a))
        weights = weights / np.sum(weights) * 16
        labels = np.tile(np.asarray([0, 1], dtype=np.int64), 30)
        alignment = np.where(labels == 1, 1.0, -1.0)
        deltas = alignment[:, None] * weights[None, :]
        participants.append(
            {
                "subject": f"S{subject + 1}",
                "profiles": {"A": profile_a, "B": profile_b},
                "directions": {
                    "A_to_B": {
                        "source": "A",
                        "target": "B",
                        "deltas": deltas,
                        "labels": labels,
                        "metadata": [],
                    },
                    "B_to_A": {
                        "source": "B",
                        "target": "A",
                        "deltas": deltas,
                        "labels": labels,
                        "metadata": [],
                    },
                },
            }
        )
    row = directional_endpoints(0, participants, "A_to_B")
    if row["fingerprint_listener_rank_probability"] != 1.0:
        raise RuntimeError("Planted listener fingerprint was not identified.")
    if row["predictive_listener_rank_probability"] <= 0.5:
        raise RuntimeError("Planted listener profile did not improve prediction rank.")
    if abs(pearson(participants[0]["profiles"]["A"], participants[0]["profiles"]["A"]) - 1.0) > 1e-12:
        raise RuntimeError("Profile self-correlation failed.")
    if rank_probability(2.0, np.asarray([1.0, 2.0, 3.0])) != 0.5:
        raise RuntimeError("Tie-aware rank probability failed.")

    attended = np.asarray([1, 1, 2, 2, 1, 2])
    video = np.asarray([0, 1, 0, 1, 0, 1])
    split = ear_split(attended, video)
    if set(split["A"]) & set(split["B"]) or set(split["A"]) | set(split["B"]) != set(range(6)):
        raise RuntimeError("Ear-SAAD split integrity failed.")
    for values in (attended, video):
        for indices in split.values():
            if len(set(values[list(indices)].tolist())) != 2:
                raise RuntimeError("Ear-SAAD balance constraint failed.")
    print(
        {
            "status": "pass",
            "fingerprint_listener_rank_probability": row[
                "fingerprint_listener_rank_probability"
            ],
            "predictive_listener_rank_probability": row[
                "predictive_listener_rank_probability"
            ],
            "ear_split": split,
        }
    )


if __name__ == "__main__":
    main()
```
