## Supplementary material for "Falsifiable substitution tests reveal task-structured neural evidence for auditory attention": Article_Analysis_Source_Code: test_wz_hic_continuous_shared_template_synthetic_v1.py

```
#!/usr/bin/env python3
"""Synthetic-only checks for shared-template comparison."""

from __future__ import annotations

import numpy as np

from run_wz_hic_continuous_shared_template_v1 import direction_endpoint


def main() -> None:
    rng = np.random.default_rng(2026073010)
    people = []
    for index in range(6):
        profile = rng.normal(size=16)
        profile -= np.mean(profile)
        labels = np.tile(np.asarray([0, 1]), 40)
        sign = np.where(labels == 1, 1.0, -1.0)
        planted = np.exp(2 * profile - np.max(2 * profile))
        planted = planted / np.sum(planted) * 16
        delta = sign[:, None] * planted[None, :]
        people.append(
            {
                "subject": f"S{index + 1}",
                "profiles": {
                    "A": profile + rng.normal(scale=0.005, size=16),
                    "B": profile + rng.normal(scale=0.005, size=16),
                },
                "tests": {
                    "A_to_B": {
                        "source": "A",
                        "target": "B",
                        "delta": delta,
                        "labels": labels,
                    },
                    "B_to_A": {
                        "source": "B",
                        "target": "A",
                        "delta": delta,
                        "labels": labels,
                    },
                },
            }
        )
    row = direction_endpoint(0, people, "A_to_B")
    if row["donor_count"] != 5:
        raise RuntimeError("Receiver exclusion failed in synthetic test.")
    if row["own_vs_mean_profile_rank_probability"] <= 0.5:
        raise RuntimeError("Planted own profile did not beat mean-profile template.")
    if row["own_vs_mean_weight_rank_probability"] <= 0.5:
        raise RuntimeError("Planted own profile did not beat mean-weight template.")
    print({"status": "pass", **row})


if __name__ == "__main__":
    main()
```
