## Supplementary material for "Falsifiable substitution tests reveal task-structured neural evidence for auditory attention": Article_Analysis_Source_Code: test_wz_hic_cross_dataset_command_identity_residual_synthetic_v1.py

```
#!/usr/bin/env python3
"""Synthetic target-removal tests for command-identity residual control."""

from __future__ import annotations

import copy

import numpy as np

from run_wz_hic_cross_dataset_command_identity_residual_v1 import (
    analyze_asme_subject,
    analyze_kojima_subject,
)
from test_wz_hic_asme30_crossphase_individual_geometry_synthetic_v1 import (
    synthetic_arrays,
)
from test_wz_hic_kojima2024b_crossrun_individual_geometry_synthetic_v1 import (
    make_arrays,
)


def asme_vector(result: dict) -> np.ndarray:
    return np.asarray(
        [
            result["physical"]["effect"],
            result["wrong"][0]["effect"],
            result["wrong"][1]["effect"],
            result["physical"]["alignment"],
            result["wrong"][0]["alignment"],
            result["wrong"][1]["alignment"],
        ],
        dtype=float,
    )


def kojima_vector(result: dict) -> np.ndarray:
    values = [
        result["physical_effect"],
        *result["wrong_effects"],
        result["physical_alignment"],
        *result["wrong_alignments"],
    ]
    for fold in result["folds"]:
        values.extend(fold["physical"]["trial_effects"])
        values.extend(fold["wrong"][0]["trial_effects"])
        values.extend(fold["wrong"][1]["trial_effects"])
    return np.asarray(values, dtype=float)


def spike_targets(arrays: dict) -> dict:
    spiked = copy.deepcopy(arrays)
    target_events = spiked["command"] == spiked["target"]
    spiked["eeg_bins"][target_events, :, 1:] += 1_000_000.0
    return spiked


def main() -> None:
    asme = synthetic_arrays(0.0)
    asme_spiked = spike_targets(asme)
    asme_result = analyze_asme_subject(asme, 1)
    asme_spiked_result = analyze_asme_subject(asme_spiked, 1)
    asme_influence = float(
        np.max(
            np.abs(
                asme_vector(asme_result) - asme_vector(asme_spiked_result)
            )
        )
    )
    if asme_influence != 0.0:
        raise AssertionError(f"ASME target influence {asme_influence}.")
    if asme_result["physical"]["effect"] <= max(
        mapping["effect"] for mapping in asme_result["wrong"]
    ):
        raise AssertionError("ASME residual physical effect did not win.")

    kojima = make_arrays()
    kojima_spiked = spike_targets(kojima)
    kojima_result = analyze_kojima_subject(kojima, "B")
    kojima_spiked_result = analyze_kojima_subject(kojima_spiked, "B")
    kojima_influence = float(
        np.max(
            np.abs(
                kojima_vector(kojima_result)
                - kojima_vector(kojima_spiked_result)
            )
        )
    )
    if kojima_influence != 0.0:
        raise AssertionError(f"Kojima target influence {kojima_influence}.")
    if kojima_result["physical_effect"] <= max(
        kojima_result["wrong_effects"]
    ):
        raise AssertionError("Kojima residual physical effect did not win.")

    print(
        "PASS synthetic cross-dataset command residual control; "
        "target-event influence is exactly zero. The real-data balance "
        "gate is evaluated from the complete target schedule."
    )


if __name__ == "__main__":
    main()
```
