## Supplementary material for "Falsifiable substitution tests reveal task-structured neural evidence for auditory attention": Article_Analysis_Source_Code: test_wz_hic_cross_dataset_wearable_calibration_synthetic_v1.py

```
#!/usr/bin/env python3
"""Construct-validity tests for the frozen wearable/calibration runner."""

from __future__ import annotations

import json

import numpy as np

from run_wz_hic_cross_dataset_command_identity_residual_v1 import (
    command_residualize,
    event_labels,
)
from run_wz_hic_cross_dataset_wearable_calibration_v1 import (
    KOJIMA_PHYSICAL,
    KOJIMA_WRONG,
    MONTAGES,
    analyze_xy_partition,
    collapse_partition,
)


def make_events(
    n_trials: int, seed: int
) -> tuple[np.ndarray, np.ndarray, np.ndarray, np.ndarray, np.ndarray]:
    rng = np.random.default_rng(seed)
    command = []
    target = []
    trial = []
    for trial_id in range(n_trials):
        target_value = 1 + (trial_id % 4)
        for command_value in range(1, 5):
            for _ in range(3):
                command.append(command_value)
                target.append(target_value)
                trial.append(trial_id)
    command = np.asarray(command, dtype=int)
    target = np.asarray(target, dtype=int)
    trial = np.asarray(trial, dtype=int)
    labels = event_labels(command, target, KOJIMA_PHYSICAL, 4)
    direction = np.linspace(-1.0, 1.0, 80)
    direction /= np.linalg.norm(direction)
    command_offsets = rng.normal(0, 0.7, size=(4, 80))
    amplitude = np.zeros(len(command))
    amplitude[labels == "S"] = 1.5
    amplitude[labels == "U"] = -0.75
    x = (
        amplitude[:, None] * direction[None, :]
        + command_offsets[command - 1]
        + rng.normal(0, 0.25, size=(len(command), 80))
    )
    eog = (
        0.05 * amplitude[:, None]
        + rng.normal(0, 0.5, size=(len(command), 20))
    )
    return x, eog, command, target, trial


def analyze(target_spike: float) -> dict:
    train_x, train_eog, train_command, train_target, _ = make_events(
        16, 20260730
    )
    test_x, test_eog, test_command, test_target, test_trial = make_events(
        8, 20260731
    )
    train_target_mask = train_command == train_target
    test_target_mask = test_command == test_target
    train_x[train_target_mask] += target_spike
    test_x[test_target_mask] -= target_spike
    partition = analyze_xy_partition(
        train_x=train_x,
        test_x=test_x,
        train_eog=train_eog,
        test_eog=test_eog,
        train_command=train_command,
        train_target=train_target,
        test_command=test_command,
        test_target=test_target,
        test_trial=test_trial,
        physical_groups=KOJIMA_PHYSICAL,
        wrong_groups=KOJIMA_WRONG,
        n_commands=4,
    )
    return collapse_partition(partition)


def main() -> None:
    base = analyze(0.0)
    spike = analyze(1_000_000.0)
    compared = (
        "physical_effect",
        "best_wrong_effect",
        "binding_specificity",
        "physical_alignment",
        "best_wrong_alignment",
        "mapping_alignment_specificity",
    )
    target_spike_max_abs_error = max(
        abs(base[key] - spike[key]) for key in compared
    )

    train_x, _, command, target, _ = make_events(16, 20260730)
    residual, _, _ = command_residualize(
        train_x, train_x.copy(), command, target, command, 4
    )
    non_target = command != target
    residual_command_means = [
        float(
            np.max(
                np.abs(
                    residual[(command == value) & non_target].mean(axis=0)
                )
            )
        )
        for value in range(1, 5)
    ]
    nested = (
        set(MONTAGES[8]).issubset(MONTAGES[16])
        and set(MONTAGES[16]).issubset(MONTAGES[32])
        and set(MONTAGES[32]).issubset(MONTAGES[64])
        and all(len(MONTAGES[n]) == n for n in (8, 16, 32, 64))
    )
    checks = {
        "physical_exceeds_best_wrong": (
            base["physical_effect"] > base["best_wrong_effect"]
        ),
        "binding_specificity_positive": base["binding_specificity"] > 0,
        "million_amplitude_target_spike_invariant": (
            target_spike_max_abs_error <= 1e-9
        ),
        "training_non_target_command_means_removed": (
            max(residual_command_means) <= 1e-12
        ),
        "montages_are_strictly_nested_and_sized": nested,
    }
    output = {
        "status": "PASS" if all(checks.values()) else "FAIL",
        "checks": checks,
        "base_metrics": base,
        "target_spike_max_abs_error": target_spike_max_abs_error,
        "maximum_residual_command_mean": max(residual_command_means),
    }
    print(json.dumps(output, indent=2))
    if output["status"] != "PASS":
        raise SystemExit(1)


if __name__ == "__main__":
    main()
```
