## Supplementary material for "Falsifiable substitution tests reveal task-structured neural evidence for auditory attention": Article_Analysis_Source_Code: test_wz_hic_kojima2024b_crossrun_individual_geometry_synthetic_v1.py

```
#!/usr/bin/env python3
"""Synthetic target-removal test for Kojima cross-run geometry."""

from __future__ import annotations

import copy

import numpy as np

from run_wz_hic_kojima2024b_crossrun_individual_geometry_v1 import (
    PHYSICAL,
    analyze_subject,
    mapping_lookup,
)


def make_arrays() -> dict:
    rng = np.random.default_rng(20260730)
    rows = []
    runs = (1, 3, 5, 8, 10, 12)
    for run_index, run in enumerate(runs):
        for local_trial in range(4):
            target = (run_index + local_trial) % 4 + 1
            trial = run * 10 + local_trial + 1
            for command in range(1, 5):
                for repetition in range(3):
                    rows.append((run, trial, target, command, repetition))
    n = len(rows)
    eeg = rng.normal(0, 0.65, size=(n, 6, 11))
    eog = rng.normal(0, 0.65, size=(n, 2, 11))
    direction = np.asarray([1.0, -0.8, 0.6, -0.4, 0.9, -0.5])
    time = np.asarray([0.0, 0.2, 0.5, 0.8, 1.1, 1.3, 1.1, 0.7, 0.4, 0.1])
    lookup = mapping_lookup(PHYSICAL)
    command = np.asarray([row[3] for row in rows], dtype=np.int8)
    target = np.asarray([row[2] for row in rows], dtype=np.int8)
    for index, row in enumerate(rows):
        same = (
            row[3] != row[2] and lookup[row[3]] == lookup[row[2]]
        )
        if same:
            eeg[index, :, 1:] += 1.3 * direction[:, None] * time[None, :]
            eog[index, :, 1:] += 0.05 * time[None, :]
    return {
        "eeg_bins": eeg.astype(np.float32),
        "eog_bins": eog.astype(np.float32),
        "subject": np.full(n, "B"),
        "task": np.full(n, "2stream"),
        "run": np.asarray([row[0] for row in rows], dtype=np.int16),
        "trial": np.asarray([row[1] for row in rows], dtype=np.int16),
        "target": target,
        "command": command,
        "accepted": np.ones(n, dtype=bool),
    }


def vector(result: dict) -> np.ndarray:
    values = [
        result["physical_effect"],
        *result["wrong_effects"],
        result["eog_effect"],
        result["physical_alignment"],
        *result["wrong_alignments"],
        *result["physical_time"],
        *result["wrong_time"][0],
        *result["wrong_time"][1],
    ]
    for fold in result["folds"]:
        values.extend(fold["physical"]["full_trial_effects"])
        values.extend(fold["wrong"][0]["full_trial_effects"])
        values.extend(fold["wrong"][1]["full_trial_effects"])
    return np.asarray(values, dtype=float)


def main() -> None:
    baseline = make_arrays()
    spiked = copy.deepcopy(baseline)
    target_events = spiked["command"] == spiked["target"]
    spiked["eeg_bins"][target_events, :, 1:] += 1_000_000.0
    spiked["eog_bins"][target_events, :, 1:] += 1_000_000.0
    result = analyze_subject(baseline, "B")
    result_spiked = analyze_subject(spiked, "B")
    influence = float(np.max(np.abs(vector(result) - vector(result_spiked))))
    if influence != 0.0:
        raise AssertionError(f"Target spike influence was {influence}.")
    if result["physical_effect"] <= max(result["wrong_effects"]):
        raise AssertionError("Physical synthetic effect did not exceed wrong.")
    if result["physical_alignment"] <= max(result["wrong_alignments"]):
        raise AssertionError("Physical synthetic alignment did not exceed wrong.")
    if result["n_eligible_trials"] != 24:
        raise AssertionError("Synthetic coverage was not 24 trials.")
    print(
        "PASS synthetic Kojima cross-run geometry; target-event influence "
        "is exactly zero."
    )


if __name__ == "__main__":
    main()
```
