## Supplementary material for "Falsifiable substitution tests reveal task-structured neural evidence for auditory attention": Article_Analysis_Source_Code: test_wz_hic_target_free_mechanism_synthetic_v1.py

```
#!/usr/bin/env python3
"""Synthetic engineering tests for target-free parent pooling."""

from __future__ import annotations

import numpy as np

from run_wz_hic_target_free_mechanism_v1 import row_metrics, trial_rows


def main() -> None:
    commands = (1, 2, 3, 4)
    groups = ((1, 2), (3, 4))
    horizon = 3
    command = np.tile(np.repeat(np.asarray(commands), horizon), 4)
    trial = np.repeat(np.arange(4), len(commands) * horizon)
    target = np.repeat(np.asarray([1, 3, 2, 4]), len(commands) * horizon)

    flat = np.zeros(command.size, dtype=float)
    outer = np.zeros(command.size, dtype=float)
    for trial_id in range(4):
        ids = trial == trial_id
        true_parent = 0 if target[ids][0] in (1, 2) else 1
        prediction = (1, 3, 3, 2)[trial_id]
        flat[ids & (command == prediction)] = 5.0
        for value in commands:
            parent = 0 if value in (1, 2) else 1
            outer[ids & (command == value)] = (
                2.0 if parent == true_parent else -2.0
            )
        outer[ids & (command == target[ids][0])] = 100.0

    rows = trial_rows(
        flat_score=flat,
        outer_score=outer,
        command=command,
        trial=trial,
        target=target,
        commands=commands,
        pooling_groups=groups,
        correctness_groups=groups,
        exclude_true_target=True,
    )
    metrics = row_metrics(rows)
    assert metrics["n_trials"] == 4
    assert metrics["n_parent_correct"] == 2
    assert metrics["n_parent_error"] == 2
    assert metrics["certificate_AUC"] == 1.0
    assert metrics["minimum_parent_event_count"] == 3
    assert metrics["maximum_parent_event_count"] == 6
    assert all(
        abs(row["certificate"]) == 4.0 for row in rows
    ), "Target score leaked into the target-free certificate."

    inclusive = trial_rows(
        flat_score=flat,
        outer_score=outer,
        command=command,
        trial=trial,
        target=target,
        commands=commands,
        pooling_groups=groups,
        correctness_groups=groups,
        exclude_true_target=False,
    )
    assert any(
        abs(row["certificate"]) > 4.0 for row in inclusive
    ), "Synthetic target spike should affect the inclusive score."
    print(
        {
            "status": "PASS",
            "target_free_auc": metrics["certificate_AUC"],
            "target_free_rows": len(rows),
        }
    )


if __name__ == "__main__":
    main()
```
