## Supplementary material for "Falsifiable substitution tests reveal task-structured neural evidence for auditory attention": Article_Analysis_Source_Code: validate_dtu_locked_external.py

```
"""Run the hash-locked DTU external evaluation exactly once."""

from __future__ import annotations

import argparse
import csv
import hashlib
import json
from collections import defaultdict
from itertools import product
from pathlib import Path

import numpy as np

import discover_kul_wang_cca as base
from audit_dtu_aad import SUBJECTS
from validate_kul_discovery_candidate import (
    RANDOM_SEED,
    bootstrap_interval,
    correlation_columns,
)


base.AUDIO_LAG_SAMPLES = 16
EXPECTED_LOCK_SHA256 = (
    "f87d8bf4ac291d2c1f2bb57138f0175335dc58a9624a205fbf5c5777dcf89440"
)
WINDOW_SECONDS = 5
NULL_REPLICATES = 99


def sha256(path: Path) -> str:
    digest = hashlib.sha256()
    with path.open("rb") as handle:
        while block := handle.read(1024 * 1024):
            digest.update(block)
    return digest.hexdigest()


def trial_margins(
    x: np.ndarray,
    attended: np.ndarray,
    unattended: np.ndarray,
    canonical: np.ndarray,
) -> dict[str, np.ndarray]:
    trim = base.TRIM_SECONDS * base.FS
    window = WINDOW_SECONDS * base.FS
    output = defaultdict(list)
    weights = canonical[:4] / canonical[:4].sum()
    for start in range(
        trim, x.shape[0] - trim - window + 1, window
    ):
        stop = start + window
        component = correlation_columns(
            x[start:stop], attended[start:stop]
        ) - correlation_columns(x[start:stop], unattended[start:stop])
        output["dominant_k1"].append(float(component[0]))
        output["equal_k4"].append(float(np.mean(component)))
        output["canonical_k4"].append(float(np.dot(weights, component)))
    return {name: np.asarray(value) for name, value in output.items()}


def participant_accuracies(
    records: list[dict[str, object]],
    shifts: dict[int, int] | None = None,
) -> dict[str, dict[int, float]]:
    trials = defaultdict(list)
    for index, record in enumerate(records):
        attended = np.asarray(record["attended"])
        unattended = np.asarray(record["unattended"])
        if shifts is not None:
            attended = np.roll(attended, shifts[index], axis=0)
            unattended = np.roll(unattended, shifts[index], axis=0)
        margins = trial_margins(
            np.asarray(record["x"]),
            attended,
            unattended,
            np.asarray(record["canonical"]),
        )
        for method, value in margins.items():
            trials[(method, int(record["subject"]))].append(
                float(np.mean(value > 0))
            )
    output = defaultdict(dict)
    for (method, subject), values in trials.items():
        output[method][subject] = float(np.mean(values))
    return dict(output)


def vector(
    accuracies: dict[str, dict[int, float]], method: str
) -> np.ndarray:
    return np.asarray(
        [accuracies[method][subject] for subject in SUBJECTS]
    )


def exact_sign_flip(differences: np.ndarray) -> float:
    observed = float(differences.mean())
    exceedances = 0
    total = 0
    for signs in product((-1, 1), repeat=differences.size):
        total += 1
        if (
            np.mean(differences * np.asarray(signs))
            >= observed - 1e-15
        ):
            exceedances += 1
    return exceedances / total


def write_csv(path: Path, rows: list[dict[str, object]]) -> None:
    with path.open("w", encoding="utf-8", newline="") as handle:
        writer = csv.DictWriter(handle, fieldnames=list(rows[0]))
        writer.writeheader()
        writer.writerows(rows)


def main() -> None:
    parser = argparse.ArgumentParser()
    parser.add_argument(
        "--input-dir",
        type=Path,
        default=Path("data/derived/dtu_aad_32hz"),
    )
    parser.add_argument(
        "--lock-file",
        type=Path,
        default=Path("docs/dtu_external_lock.json"),
    )
    parser.add_argument(
        "--output-dir",
        type=Path,
        default=Path("results/dtu_aad/locked_external"),
    )
    args = parser.parse_args()
    lock_hash = sha256(args.lock_file)
    if lock_hash != EXPECTED_LOCK_SHA256:
        raise RuntimeError(f"DTU lock hash mismatch: {lock_hash}")
    manifest = json.loads(
        (args.input_dir / "manifest.json").read_text(encoding="utf-8")
    )
    if (
        tuple(manifest["participants"]) != SUBJECTS
        or manifest["lock_sha256"] != EXPECTED_LOCK_SHA256
    ):
        raise RuntimeError("DTU manifest or lock mismatch")
    audio_archive = np.load(
        args.input_dir / str(manifest["audio_cache"])
    )
    audio = {
        name.removeprefix("audio::"): audio_archive[name]
        for name in audio_archive.files
        if name.startswith("audio::")
    }

    records = []
    fold_rows = []
    trial_rows = []
    for subject in SUBJECTS:
        archive = np.load(args.input_dir / f"S{subject}.npz")
        metadata = json.loads(str(archive["metadata_json"]))
        eeg = [
            archive[f"eeg_{index:02d}"] for index in range(1, 61)
        ]
        attended = [
            audio[str(row["attended_audio"])][: int(row["samples"])]
            for row in metadata
        ]
        pair_groups = sorted(
            {str(row["unordered_story_pair_group"]) for row in metadata}
        )
        for pair_group in pair_groups:
            held_out_stories = set(pair_group.split("|"))
            testing_indices = [
                index
                for index, row in enumerate(metadata)
                if str(row["unordered_story_pair_group"]) == pair_group
            ]
            training_indices = [
                index
                for index, row in enumerate(metadata)
                if {
                    str(row["attended_story_identity"]),
                    str(row["unattended_story_identity"]),
                }.isdisjoint(held_out_stories)
            ]
            if not training_indices or not testing_indices:
                raise RuntimeError("DTU strict identity fold is empty")
            basis, pca_scale = base.pca_basis(
                [eeg[index] for index in training_indices]
            )
            statistics = base.sufficient_statistics(
                eeg,
                attended,
                training_indices,
                basis,
                pca_scale,
            )
            a, b, canonical = base.fit_cca(*statistics["full"])
            fold_rows.append(
                {
                    "subject": subject,
                    "held_out_story_pair": pair_group,
                    "training_trials": len(training_indices),
                    "testing_trials": len(testing_indices),
                    "excluded_shared_story_trials": (
                        60 - len(training_indices) - len(testing_indices)
                    ),
                    **{
                        f"canonical_{component + 1}": float(
                            canonical[component]
                        )
                        for component in range(4)
                    },
                }
            )
            for trial_index in testing_indices:
                row = metadata[trial_index]
                projected = (
                    base.standardize_columns(eeg[trial_index]) @ basis
                ) / pca_scale
                x = base.lag_matrix(projected) @ a[:, :4]
                attended_y = (
                    base.audio_lag_matrix(
                        base.standardize_columns(attended[trial_index])
                    )
                    @ b[:, :4]
                )
                unattended_audio = audio[str(row["unattended_audio"])][
                    : int(row["samples"])
                ]
                unattended_y = (
                    base.audio_lag_matrix(
                        base.standardize_columns(unattended_audio)
                    )
                    @ b[:, :4]
                )
                usable = min(
                    x.shape[0],
                    attended_y.shape[0],
                    unattended_y.shape[0],
                )
                record = {
                    "subject": subject,
                    "trial": int(row["trial"]),
                    "held_out_story_pair": pair_group,
                    "x": x[:usable],
                    "attended": attended_y[:usable],
                    "unattended": unattended_y[:usable],
                    "canonical": canonical[:4],
                }
                records.append(record)
                margins = trial_margins(
                    record["x"],
                    record["attended"],
                    record["unattended"],
                    record["canonical"],
                )
                for method, value in margins.items():
                    trial_rows.append(
                        {
                            "subject": subject,
                            "trial": int(row["trial"]),
                            "held_out_story_pair": pair_group,
                            "method": method,
                            "windows": int(value.size),
                            "accuracy": float(np.mean(value > 0)),
                            "mean_margin": float(value.mean()),
                        }
                    )
            print(
                f"DTU S{subject:02d}, held out {pair_group}: complete",
                flush=True,
            )
        del archive, eeg, attended

    observed = participant_accuracies(records)
    dominant = vector(observed, "dominant_k1")
    equal = vector(observed, "equal_k4")
    canonical = vector(observed, "canonical_k4")
    gain = equal - dominant
    capacity_gain = equal - canonical
    rng = np.random.default_rng(RANDOM_SEED)
    null_accuracy = []
    null_gain = []
    for replicate in range(NULL_REPLICATES):
        shifts = {}
        for index, record in enumerate(records):
            length = np.asarray(record["x"]).shape[0]
            minimum = 10 * base.FS
            shifts[index] = int(rng.integers(minimum, length - minimum))
        shifted = participant_accuracies(records, shifts)
        shifted_equal = vector(shifted, "equal_k4")
        shifted_dominant = vector(shifted, "dominant_k1")
        null_accuracy.append(float(shifted_equal.mean()))
        null_gain.append(
            float((shifted_equal - shifted_dominant).mean())
        )
        if (replicate + 1) % 10 == 0:
            print(
                f"DTU time-shift null {replicate + 1}/{NULL_REPLICATES}",
                flush=True,
            )
    null_accuracy_array = np.asarray(null_accuracy)
    null_gain_array = np.asarray(null_gain)
    inference_rng = np.random.default_rng(RANDOM_SEED + 2)
    primary_p = exact_sign_flip(gain)
    success = bool(gain.mean() > 0 and primary_p < 0.05)
    participant_rows = [
        {
            "subject": subject,
            "dominant_k1_accuracy": float(dominant[index]),
            "equal_k4_accuracy": float(equal[index]),
            "canonical_k4_accuracy": float(canonical[index]),
            "equal_k4_gain_vs_dominant_k1": float(gain[index]),
            "equal_k4_gain_vs_canonical_k4": float(
                capacity_gain[index]
            ),
        }
        for index, subject in enumerate(SUBJECTS)
    ]
    report = {
        "status": "hash-locked one-shot DTU external evaluation",
        "lock_sha256": lock_hash,
        "design": {
            "participants": list(SUBJECTS),
            "story_pair_groups": 7,
            "strict_shared_story_exclusion": True,
            "window_seconds": WINDOW_SECONDS,
            "candidate": "equal-four-projection dual-lag CCA",
            "primary_baseline": "dominant first-projection CCA",
            "capacity_matched_control": (
                "canonical-correlation-weighted four-projection CCA"
            ),
        },
        "observed": {
            "dominant_k1_participant_mean_accuracy": float(
                dominant.mean()
            ),
            "equal_k4_participant_mean_accuracy": float(equal.mean()),
            "canonical_k4_participant_mean_accuracy": float(
                canonical.mean()
            ),
            "equal_k4_mean_gain_vs_dominant_k1": float(gain.mean()),
            "equal_k4_mean_gain_vs_canonical_k4": float(
                capacity_gain.mean()
            ),
            "participants_improved_vs_dominant_k1": int(
                np.sum(gain > 0)
            ),
            "participants_improved_vs_canonical_k4": int(
                np.sum(capacity_gain > 0)
            ),
        },
        "locked_primary_inference": {
            "estimand": "equal_k4 minus dominant_k1 participant accuracy",
            "mean_gain": float(gain.mean()),
            "bootstrap_95_interval": bootstrap_interval(
                gain, inference_rng
            ),
            "exact_one_sided_sign_flip_p": primary_p,
            "success_criterion_met": success,
        },
        "capacity_control_inference": {
            "mean_gain": float(capacity_gain.mean()),
            "bootstrap_95_interval": bootstrap_interval(
                capacity_gain, inference_rng
            ),
            "exact_one_sided_sign_flip_p": exact_sign_flip(
                capacity_gain
            ),
        },
        "time_shift_null": {
            "replicates": NULL_REPLICATES,
            "equal_k4_accuracy_null_mean": float(
                null_accuracy_array.mean()
            ),
            "equal_k4_accuracy_null_95_interval": [
                float(np.quantile(null_accuracy_array, 0.025)),
                float(np.quantile(null_accuracy_array, 0.975)),
            ],
            "equal_k4_accuracy_empirical_p": float(
                (
                    1
                    + np.sum(
                        null_accuracy_array >= float(equal.mean())
                    )
                )
                / (NULL_REPLICATES + 1)
            ),
            "equal_k4_gain_null_mean": float(null_gain_array.mean()),
            "equal_k4_gain_null_95_interval": [
                float(np.quantile(null_gain_array, 0.025)),
                float(np.quantile(null_gain_array, 0.975)),
            ],
            "equal_k4_gain_empirical_p": float(
                (
                    1
                    + np.sum(null_gain_array >= float(gain.mean()))
                )
                / (NULL_REPLICATES + 1)
            ),
        },
        "participant_results": participant_rows,
        "interpretation_guardrail": (
            "The locked claim concerns breadth versus the conventional "
            "dominant projection. Equal weighting is not claimed superior "
            "unless the capacity-control interval excludes zero."
        ),
    }
    args.output_dir.mkdir(parents=True, exist_ok=True)
    write_csv(args.output_dir / "trial_results.csv", trial_rows)
    write_csv(args.output_dir / "folds.csv", fold_rows)
    write_csv(args.output_dir / "participants.csv", participant_rows)
    (args.output_dir / "locked_external.json").write_text(
        json.dumps(report, indent=2), encoding="utf-8"
    )
    print(json.dumps(report, indent=2))


if __name__ == "__main__":
    main()
```
