## Supplementary material for "Falsifiable substitution tests reveal task-structured neural evidence for auditory attention": Article_Analysis_Source_Code: validate_ear_saad_multimodal_external.py

```
"""Locked multimodal projection-breadth test in the Ear-SAAD dataset."""

from __future__ import annotations

import argparse
import csv
import hashlib
import json
from itertools import product
from pathlib import Path

import numpy as np
import pandas as pd
from scipy.io import loadmat
from scipy.linalg import eigh, svd


EXPECTED_LOCK_SHA256 = (
    "2afcc6451d06625a521f10abd967d8618babc02dd1c649a8c97aee14b5926827"
)
SUBJECTS = tuple(range(1, 16))
MODALITIES = ("scalp", "around_ear", "in_ear")
FS = 20
LAG_SAMPLES = 10
TRIM = 5 * FS
WINDOW = 5 * FS
REGULARIZATION = 0.1
BOOTSTRAP_REPLICATES = 20_000
BOOTSTRAP_SEED = 26_072_521
NULL_REPLICATES = 99
NULL_SEED = 26_072_522


def sha256(path: Path) -> str:
    digest = hashlib.sha256()
    with path.open("rb") as handle:
        while block := handle.read(8 * 1024 * 1024):
            digest.update(block)
    return digest.hexdigest()


def write_csv(path: Path, rows: list[dict[str, object]]) -> None:
    if not rows:
        raise RuntimeError(f"Empty Ear-SAAD output: {path}")
    with path.open("w", encoding="utf-8", newline="") as handle:
        writer = csv.DictWriter(handle, fieldnames=list(rows[0]))
        writer.writeheader()
        writer.writerows(rows)


def standardize_columns(value: np.ndarray) -> np.ndarray:
    centered = value - value.mean(axis=0, keepdims=True)
    scale = centered.std(axis=0, keepdims=True)
    scale[scale < 1e-8] = 1.0
    return centered / scale


def lag_matrix(value: np.ndarray) -> np.ndarray:
    length = value.shape[0] - LAG_SAMPLES
    return np.concatenate(
        [value[lag : lag + length] for lag in range(LAG_SAMPLES + 1)],
        axis=1,
    )


def pca_basis(
    trials: list[np.ndarray], components: int
) -> tuple[np.ndarray, np.ndarray]:
    covariance = np.zeros((trials[0].shape[1], trials[0].shape[1]))
    count = 0
    for trial in trials:
        segment = standardize_columns(trial)[TRIM:-TRIM]
        covariance += segment.T @ segment
        count += segment.shape[0]
    covariance /= count
    eigenvalues, eigenvectors = eigh(covariance)
    order = np.argsort(eigenvalues)[::-1][:components]
    scale = np.sqrt(np.maximum(eigenvalues[order], 1e-8))
    return eigenvectors[:, order], scale


def inverse_square_root(value: np.ndarray) -> np.ndarray:
    scale = np.trace(value) / value.shape[0]
    eigenvalues, eigenvectors = eigh(
        value + REGULARIZATION * scale * np.eye(value.shape[0])
    )
    eigenvalues = np.maximum(eigenvalues, 1e-10)
    return (eigenvectors / np.sqrt(eigenvalues)) @ eigenvectors.T


def fit_cca(
    cxx: np.ndarray, cyy: np.ndarray, cxy: np.ndarray
) -> tuple[np.ndarray, np.ndarray, np.ndarray]:
    wx = inverse_square_root(cxx)
    wy = inverse_square_root(cyy)
    left, canonical, right_t = svd(
        wx @ cxy @ wy, full_matrices=False, check_finite=False
    )
    return wx @ left, wy @ right_t.T, canonical


def sufficient_statistics(
    eeg: list[np.ndarray],
    target_lagged: list[np.ndarray],
    training: list[int],
    basis: np.ndarray,
    pca_scale: np.ndarray,
) -> tuple[np.ndarray, np.ndarray, np.ndarray]:
    components = basis.shape[1]
    x_dim = components * (LAG_SAMPLES + 1)
    y_dim = LAG_SAMPLES + 1
    cxx = np.zeros((x_dim, x_dim))
    cyy = np.zeros((y_dim, y_dim))
    cxy = np.zeros((x_dim, y_dim))
    count = 0
    for index in training:
        projected = (
            standardize_columns(eeg[index]) @ basis
        ) / pca_scale
        x = lag_matrix(projected)
        y = target_lagged[index]
        usable = min(x.shape[0], y.shape[0])
        xs = x[TRIM : usable - TRIM]
        ys = y[TRIM : usable - TRIM]
        cxx += xs.T @ xs
        cyy += ys.T @ ys
        cxy += xs.T @ ys
        count += xs.shape[0]
    if count == 0:
        raise RuntimeError("Empty Ear-SAAD training statistic")
    return cxx / count, cyy / count, cxy / count


def correlations_by_window(
    neural: np.ndarray,
    speech: np.ndarray,
    shift: int | None = None,
) -> np.ndarray:
    usable = min(neural.shape[0], speech.shape[0])
    windows = (usable - 2 * TRIM) // WINDOW
    length = windows * WINDOW
    neural_window = neural[TRIM : TRIM + length].reshape(
        windows, WINDOW, neural.shape[1]
    )
    if shift is None:
        speech_segment = speech[TRIM : TRIM + length]
    else:
        indices = (
            np.arange(TRIM, TRIM + length) - shift
        ) % speech.shape[0]
        speech_segment = speech[indices]
    speech_window = speech_segment.reshape(
        windows, WINDOW, speech.shape[1]
    )
    neural_centered = neural_window - neural_window.mean(axis=1, keepdims=True)
    speech_centered = speech_window - speech_window.mean(axis=1, keepdims=True)
    numerator = np.sum(neural_centered * speech_centered, axis=1)
    denominator = np.linalg.norm(neural_centered, axis=1) * np.linalg.norm(
        speech_centered, axis=1
    )
    return np.divide(
        numerator,
        denominator,
        out=np.zeros_like(numerator),
        where=denominator > 1e-12,
    )


def record_decisions(
    record: dict[str, object],
    shift: int | None = None,
) -> dict[str, np.ndarray]:
    x = np.asarray(record["x"])
    first = np.asarray(record["first"])
    second = np.asarray(record["second"])
    first_r = correlations_by_window(x, first, shift)
    second_r = correlations_by_window(x, second, shift)
    attended = int(record["attended_speaker"])
    component = first_r - second_r if attended == 1 else second_r - first_r
    canonical = np.asarray(record["canonical"], dtype=float)[:4]
    canonical /= canonical.sum()
    margins = {
        "k1": component[:, 0],
        "equal_k4": component[:, :4].mean(axis=1),
        "canonical_k4": component[:, :4] @ canonical,
    }
    return {
        **margins,
        **{f"correct_{name}": value > 0 for name, value in margins.items()},
    }


def exact_sign_flip(values: np.ndarray) -> float:
    values = np.asarray(values, dtype=float)
    observed = float(values.mean())
    count = 0
    total = 0
    for signs in product((-1, 1), repeat=values.size):
        statistic = float(np.mean(values * np.asarray(signs)))
        count += int(statistic >= observed - 1e-15)
        total += 1
    return count / total


def bootstrap_interval(
    values: np.ndarray, rng: np.random.Generator
) -> list[float]:
    values = np.asarray(values, dtype=float)
    indices = rng.integers(
        0,
        values.size,
        size=(BOOTSTRAP_REPLICATES, values.size),
    )
    means = values[indices].mean(axis=1)
    return [
        float(np.quantile(means, 0.025)),
        float(np.quantile(means, 0.975)),
    ]


def load_subject(
    root: Path, subject: int
) -> tuple[
    dict[str, list[np.ndarray]],
    list[np.ndarray],
    np.ndarray,
    np.ndarray,
    np.ndarray,
    tuple[str, ...],
]:
    data = loadmat(
        root / f"dataSubject{subject}.mat",
        squeeze_me=True,
        struct_as_record=False,
    )
    if int(data["fs"]) != FS:
        raise RuntimeError(f"Ear-SAAD S{subject}: sample rate mismatch")
    channels = tuple(str(value) for value in np.asarray(data["channels"]).ravel())
    if len(channels) != 61:
        raise RuntimeError(f"Ear-SAAD S{subject}: channel count mismatch")
    channel_indices = {
        "scalp": np.arange(29),
        "around_ear": np.asarray(
            [index for index, name in enumerate(channels) if name.startswith("cE")]
        ),
        "in_ear": np.asarray(
            [index for index, name in enumerate(channels) if name.startswith("E")]
        ),
    }
    expected_counts = {"scalp": 29, "around_ear": 19, "in_ear": 12}
    if any(
        channel_indices[name].size != expected_counts[name]
        for name in MODALITIES
    ):
        raise RuntimeError(f"Ear-SAAD S{subject}: modality assignment mismatch")
    raw_trials = np.asarray(data["eegTrials"], dtype=object).ravel()
    speech = [
        np.asarray(value, dtype=float)
        for value in np.asarray(data["envelopes"], dtype=object).ravel()
    ]
    attended = np.asarray(data["attSpeaker"], dtype=int).ravel()
    ears = np.asarray(data["attendedEar"], dtype=int).ravel()
    videos = np.asarray(data["videoCondition"], dtype=int).ravel()
    if (
        raw_trials.size != 6
        or len(speech) != 6
        or attended.shape != (6,)
        or any(value.shape != (12000, 2) for value in speech)
    ):
        raise RuntimeError(f"Ear-SAAD S{subject}: trial structure mismatch")
    modalities: dict[str, list[np.ndarray]] = {
        name: [] for name in MODALITIES
    }
    for raw in raw_trials:
        matrix = np.asarray(raw, dtype=float)
        if matrix.shape != (12000, 61):
            raise RuntimeError(f"Ear-SAAD S{subject}: EEG shape mismatch")
        matrix = np.nan_to_num(matrix, nan=0.0, posinf=0.0, neginf=0.0)
        for name in MODALITIES:
            modalities[name].append(matrix[:, channel_indices[name]])
    return modalities, speech, attended, ears, videos, channels


def trial_rows_from_records(
    records: list[dict[str, object]],
) -> tuple[list[dict[str, object]], list[dict[str, object]]]:
    trial_rows = []
    window_rows = []
    for record in records:
        values = record_decisions(record)
        trial_rows.append(
            {
                "subject": int(record["subject"]),
                "trial": int(record["trial"]),
                "modality": str(record["modality"]),
                "attended_speaker": int(record["attended_speaker"]),
                "attended_ear": int(record["attended_ear"]),
                "video_condition": int(record["video_condition"]),
                "windows": int(values["k1"].size),
                "k1_accuracy": float(values["correct_k1"].mean()),
                "equal_k4_accuracy": float(
                    values["correct_equal_k4"].mean()
                ),
                "canonical_k4_accuracy": float(
                    values["correct_canonical_k4"].mean()
                ),
                "mean_k1_margin": float(values["k1"].mean()),
                "mean_equal_k4_margin": float(values["equal_k4"].mean()),
            }
        )
        for window in range(values["k1"].size):
            window_rows.append(
                {
                    "subject": int(record["subject"]),
                    "trial": int(record["trial"]),
                    "modality": str(record["modality"]),
                    "window": window + 1,
                    "margin_k1": float(values["k1"][window]),
                    "margin_equal_k4": float(values["equal_k4"][window]),
                    "margin_canonical_k4": float(
                        values["canonical_k4"][window]
                    ),
                    "correct_k1": int(values["correct_k1"][window]),
                    "correct_equal_k4": int(
                        values["correct_equal_k4"][window]
                    ),
                    "correct_canonical_k4": int(
                        values["correct_canonical_k4"][window]
                    ),
                }
            )
    return trial_rows, window_rows


def participant_modality_table(trials: pd.DataFrame) -> pd.DataFrame:
    columns = ["k1_accuracy", "equal_k4_accuracy", "canonical_k4_accuracy"]
    table = trials.groupby(["subject", "modality"], as_index=False)[columns].mean()
    table["gain_equal_minus_k1"] = (
        table["equal_k4_accuracy"] - table["k1_accuracy"]
    )
    table["gain_canonical_minus_k1"] = (
        table["canonical_k4_accuracy"] - table["k1_accuracy"]
    )
    return table


def main() -> None:
    parser = argparse.ArgumentParser()
    parser.add_argument(
        "--root",
        type=Path,
        default=Path("data/raw/ear_saad/preprocessedData"),
    )
    parser.add_argument(
        "--output-dir",
        type=Path,
        default=Path("results/ear_saad/locked_multimodal_external"),
    )
    parser.add_argument(
        "--lock",
        type=Path,
        default=Path("docs/ear_saad_multimodal_external_lock.json"),
    )
    args = parser.parse_args()
    if sha256(args.lock) != EXPECTED_LOCK_SHA256:
        raise RuntimeError("Ear-SAAD lock hash mismatch")
    args.output_dir.mkdir(parents=True, exist_ok=True)

    all_records: list[dict[str, object]] = []
    fold_rows: list[dict[str, object]] = []
    null_subjects = np.empty((len(SUBJECTS), NULL_REPLICATES, 2))
    null_rng = np.random.default_rng(NULL_SEED)
    reference_channels: tuple[str, ...] | None = None

    for subject_position, subject in enumerate(SUBJECTS):
        modalities, speech, attended, ears, videos, channels = load_subject(
            args.root, subject
        )
        if reference_channels is None:
            reference_channels = channels
        elif channels != reference_channels:
            raise RuntimeError("Ear-SAAD channel labels vary across participants")
        speech_lagged = [
            [
                lag_matrix(
                    standardize_columns(value[:, stream : stream + 1])
                )
                for stream in range(2)
            ]
            for value in speech
        ]
        target_lagged = [
            speech_lagged[trial][int(attended[trial]) - 1]
            for trial in range(6)
        ]
        subject_records: list[dict[str, object]] = []
        for modality in MODALITIES:
            eeg = modalities[modality]
            components = min(16, eeg[0].shape[1])
            for test in range(6):
                training = [index for index in range(6) if index != test]
                basis, pca_scale = pca_basis(
                    [eeg[index] for index in training], components
                )
                cxx, cyy, cxy = sufficient_statistics(
                    eeg,
                    target_lagged,
                    training,
                    basis,
                    pca_scale,
                )
                neural_weights, speech_weights, canonical = fit_cca(
                    cxx, cyy, cxy
                )
                projected_eeg = (
                    lag_matrix(
                        (
                            standardize_columns(eeg[test]) @ basis
                        )
                        / pca_scale
                    )
                    @ neural_weights[:, :4]
                )
                projected_first = (
                    speech_lagged[test][0] @ speech_weights[:, :4]
                )
                projected_second = (
                    speech_lagged[test][1] @ speech_weights[:, :4]
                )
                usable = min(
                    projected_eeg.shape[0],
                    projected_first.shape[0],
                    projected_second.shape[0],
                )
                record = {
                    "subject": subject,
                    "trial": test + 1,
                    "modality": modality,
                    "attended_speaker": int(attended[test]),
                    "attended_ear": int(ears[test]),
                    "video_condition": int(videos[test]),
                    "x": projected_eeg[:usable],
                    "first": projected_first[:usable],
                    "second": projected_second[:usable],
                    "canonical": canonical[:4],
                }
                subject_records.append(record)
                fold_rows.append(
                    {
                        "subject": subject,
                        "modality": modality,
                        "held_out_trial": test + 1,
                        "training_trials": 5,
                        "testing_trials": 1,
                        "pca_components": components,
                        "trial_overlap": 0,
                        **{
                            f"canonical_{component + 1}": float(
                                canonical[component]
                            )
                            for component in range(4)
                        },
                    }
                )
        if len(subject_records) != 18:
            raise RuntimeError(f"Ear-SAAD S{subject}: record count mismatch")
        all_records.extend(subject_records)

        for replicate in range(NULL_REPLICATES):
            modality_values = []
            for modality in MODALITIES:
                trial_values = []
                for record in subject_records:
                    if record["modality"] != modality:
                        continue
                    usable = np.asarray(record["x"]).shape[0]
                    shift = int(
                        null_rng.integers(10 * FS, usable - 10 * FS)
                    )
                    decisions = record_decisions(record, shift)
                    trial_values.append(
                        (
                            float(decisions["correct_k1"].mean()),
                            float(decisions["correct_equal_k4"].mean()),
                        )
                    )
                modality_values.append(np.mean(trial_values, axis=0))
            null_subjects[subject_position, replicate] = np.mean(
                modality_values, axis=0
            )
        print(
            f"Ear-SAAD locked multimodal {subject_position + 1:02d}/15: "
            f"S{subject:02d}",
            flush=True,
        )

    trial_rows, window_rows = trial_rows_from_records(all_records)
    trials = pd.DataFrame(trial_rows)
    modality_table = participant_modality_table(trials)
    primary = (
        modality_table.groupby("subject", as_index=False)[
            [
                "k1_accuracy",
                "equal_k4_accuracy",
                "canonical_k4_accuracy",
                "gain_equal_minus_k1",
                "gain_canonical_minus_k1",
            ]
        ]
        .mean()
        .sort_values("subject")
    )
    if tuple(primary["subject"]) != SUBJECTS:
        raise RuntimeError("Ear-SAAD primary participant set mismatch")
    gains = primary["gain_equal_minus_k1"].to_numpy()
    rng = np.random.default_rng(BOOTSTRAP_SEED)
    primary_p = exact_sign_flip(gains)
    primary_success = bool(gains.mean() > 0 and primary_p < 0.05)

    modality_summary = {}
    for modality in MODALITIES:
        table = modality_table[modality_table["modality"] == modality]
        modality_gain = table["gain_equal_minus_k1"].to_numpy()
        modality_summary[modality] = {
            "k1_accuracy_mean": float(table["k1_accuracy"].mean()),
            "equal_k4_accuracy_mean": float(
                table["equal_k4_accuracy"].mean()
            ),
            "canonical_k4_accuracy_mean": float(
                table["canonical_k4_accuracy"].mean()
            ),
            "equal_k4_minus_k1_mean": float(modality_gain.mean()),
            "participants_positive": int((modality_gain > 0).sum()),
            "bootstrap_95": bootstrap_interval(modality_gain, rng),
            "descriptive_exact_one_sided_p": exact_sign_flip(modality_gain),
        }

    condition_summary = {}
    for condition, column, value in (
        ("video", "video_condition", 1),
        ("fixation", "video_condition", 0),
        ("left_attention", "attended_ear", 1),
        ("right_attention", "attended_ear", 2),
    ):
        selected = trials[trials[column] == value]
        table = participant_modality_table(selected)
        participant = table.groupby("subject")["gain_equal_minus_k1"].mean()
        condition_summary[condition] = {
            "participants": int(participant.size),
            "equal_k4_minus_k1_mean": float(participant.mean()),
            "participants_positive": int((participant > 0).sum()),
        }

    pivot = modality_table.pivot(
        index="subject", columns="modality", values="gain_equal_minus_k1"
    )
    sensor_gradient = (
        pivot["in_ear"].to_numpy() - pivot["scalp"].to_numpy()
    ) / 2.0
    null_k1 = null_subjects[:, :, 0].mean(axis=0)
    null_k4 = null_subjects[:, :, 1].mean(axis=0)
    null_gain = null_k4 - null_k1
    observed_k4 = float(primary["equal_k4_accuracy"].mean())
    observed_gain = float(gains.mean())
    null_rows = [
        {
            "replicate": replicate + 1,
            "k1_accuracy": float(null_k1[replicate]),
            "equal_k4_accuracy": float(null_k4[replicate]),
            "gain_equal_minus_k1": float(null_gain[replicate]),
        }
        for replicate in range(NULL_REPLICATES)
    ]

    scalp_k1 = modality_table[
        modality_table["modality"] == "scalp"
    ]["k1_accuracy"].to_numpy()
    payload = {
        "dataset": "Ear-SAAD",
        "source_doi": "10.5281/zenodo.16536441",
        "role": "locked multimodal external projection-breadth test",
        "lock_sha256": EXPECTED_LOCK_SHA256,
        "participants": 15,
        "modalities": list(MODALITIES),
        "primary_equal_modality_average": {
            "k1_accuracy_mean": float(primary["k1_accuracy"].mean()),
            "equal_k4_accuracy_mean": observed_k4,
            "canonical_k4_accuracy_mean": float(
                primary["canonical_k4_accuracy"].mean()
            ),
            "equal_k4_minus_k1_mean": observed_gain,
            "participants_positive": int((gains > 0).sum()),
            "bootstrap_95": bootstrap_interval(gains, rng),
            "exact_one_sided_p": primary_p,
            "success_rule_met": primary_success,
        },
        "modality_specific": modality_summary,
        "sensor_gradient_in_ear_minus_scalp_half_contrast": {
            "mean": float(sensor_gradient.mean()),
            "bootstrap_95": bootstrap_interval(sensor_gradient, rng),
            "descriptive_exact_one_sided_p_greater_than_zero": exact_sign_flip(
                sensor_gradient
            ),
        },
        "condition_strata": condition_summary,
        "published_baseline_audit": {
            "scalp_k1_accuracy_mean": float(scalp_k1.mean()),
            "scalp_k1_vs_chance_exact_one_sided_p": exact_sign_flip(
                scalp_k1 - 0.5
            ),
            "passed": bool(
                scalp_k1.mean() > 0.5
                and exact_sign_flip(scalp_k1 - 0.5) < 0.05
            ),
        },
        "temporal_alignment_null": {
            "replicates": NULL_REPLICATES,
            "observed_equal_k4_accuracy": observed_k4,
            "accuracy_null_mean": float(null_k4.mean()),
            "accuracy_null_95": [
                float(np.quantile(null_k4, 0.025)),
                float(np.quantile(null_k4, 0.975)),
            ],
            "accuracy_empirical_p": float(
                (1 + np.sum(null_k4 >= observed_k4))
                / (NULL_REPLICATES + 1)
            ),
            "observed_gain": observed_gain,
            "gain_null_mean": float(null_gain.mean()),
            "gain_null_95": [
                float(np.quantile(null_gain, 0.025)),
                float(np.quantile(null_gain, 0.975)),
            ],
            "gain_empirical_p": float(
                (1 + np.sum(null_gain >= observed_gain))
                / (NULL_REPLICATES + 1)
            ),
        },
        "decision": (
            "advance as multimodal external support"
            if primary_success
            else "retain as failed external test; do not substitute a modality"
        ),
    }
    write_csv(args.output_dir / "trial_results.csv", trial_rows)
    write_csv(args.output_dir / "window_results.csv", window_rows)
    write_csv(args.output_dir / "folds.csv", fold_rows)
    write_csv(
        args.output_dir / "participant_modalities.csv",
        modality_table.to_dict(orient="records"),
    )
    write_csv(
        args.output_dir / "primary_participants.csv",
        primary.to_dict(orient="records"),
    )
    write_csv(args.output_dir / "temporal_shift_null.csv", null_rows)
    (args.output_dir / "locked_external.json").write_text(
        json.dumps(payload, indent=2),
        encoding="utf-8",
    )
    print(json.dumps(payload, indent=2))


if __name__ == "__main__":
    main()
```
