## Supplementary material for "Falsifiable substitution tests reveal task-structured neural evidence for auditory attention": Article_Analysis_Source_Code: validate_kul_discovery_candidate.py

```
"""Stress-test the selected KUL discovery candidate before any unsealing.

Selected discovery candidate:
  * dual 0--500 ms lag embeddings for EEG and 15-subband speech;
  * regularized CCA (0.1), participant-specific and training-fold only;
  * equal mean of the first four canonical correlation margins;
  * five-second decisions;
  * leave-one-complete-base-part-identity-out folds.

The script recomputes the candidate, dominant-projection baseline, and
canonical-weighted four-projection capacity control.  It then constructs 99
independent circular time-shift null datasets with at least 10 seconds of
misalignment per trial.  Participant-level exact sign-flip and bootstrap
intervals are descriptive because the candidate was selected on these odd-ID
participants; they are not represented as confirmation tests.
"""

from __future__ import annotations

import argparse
import json
from collections import defaultdict
from itertools import product
from pathlib import Path

import numpy as np

import discover_kul_wang_cca as base
from audit_kul_discovery import DISCOVERY_SUBJECTS


base.AUDIO_LAG_SAMPLES = 16
WINDOW_SECONDS = 5
NULL_REPLICATES = 99
RANDOM_SEED = 20260725


def correlation_columns(first: np.ndarray, second: np.ndarray) -> np.ndarray:
    a = first - first.mean(axis=0, keepdims=True)
    b = second - second.mean(axis=0, keepdims=True)
    denominator = np.linalg.norm(a, axis=0) * np.linalg.norm(b, axis=0)
    return np.divide(
        np.sum(a * b, axis=0),
        denominator,
        out=np.zeros(a.shape[1]),
        where=denominator > 1e-12,
    )


def trial_margins(
    x: np.ndarray,
    attended: np.ndarray,
    unattended: np.ndarray,
    canonical: np.ndarray,
) -> dict[str, np.ndarray]:
    trim = base.TRIM_SECONDS * base.FS
    window = WINDOW_SECONDS * base.FS
    output = defaultdict(list)
    for start in range(
        trim, x.shape[0] - trim - window + 1, window
    ):
        stop = start + window
        r_attended = correlation_columns(x[start:stop], attended[start:stop])
        r_unattended = correlation_columns(
            x[start:stop], unattended[start:stop]
        )
        component = r_attended - r_unattended
        output["dominant_k1"].append(float(component[0]))
        output["equal_k4"].append(float(np.mean(component[:4])))
        weights = canonical[:4] / canonical[:4].sum()
        output["canonical_k4"].append(
            float(np.dot(weights, component[:4]))
        )
    return {name: np.asarray(value) for name, value in output.items()}


def participant_accuracies(
    records: list[dict[str, object]],
    *,
    shifts: dict[int, int] | None = None,
) -> dict[str, dict[int, float]]:
    trial_values = defaultdict(list)
    for record_index, record in enumerate(records):
        attended = np.asarray(record["attended"])
        unattended = np.asarray(record["unattended"])
        if shifts is not None:
            shift = shifts[record_index]
            attended = np.roll(attended, shift, axis=0)
            unattended = np.roll(unattended, shift, axis=0)
        margins = trial_margins(
            np.asarray(record["x"]),
            attended,
            unattended,
            np.asarray(record["canonical"]),
        )
        for method, values in margins.items():
            trial_values[(method, int(record["subject"]))].append(
                float(np.mean(values > 0))
            )
    output = defaultdict(dict)
    for (method, subject), values in trial_values.items():
        output[method][subject] = float(np.mean(values))
    return dict(output)


def exact_sign_flip(
    differences: np.ndarray, alternative: str = "greater"
) -> float:
    observed = float(np.mean(differences))
    null = np.asarray(
        [
            np.mean(differences * np.asarray(signs))
            for signs in product((-1, 1), repeat=differences.size)
        ]
    )
    if alternative == "greater":
        return float(np.mean(null >= observed - 1e-15))
    if alternative == "two-sided":
        return float(np.mean(np.abs(null) >= abs(observed) - 1e-15))
    raise ValueError(alternative)


def bootstrap_interval(
    values: np.ndarray, rng: np.random.Generator
) -> list[float]:
    indices = rng.integers(
        0, values.size, size=(20000, values.size)
    )
    means = values[indices].mean(axis=1)
    return [
        float(np.quantile(means, 0.025)),
        float(np.quantile(means, 0.975)),
    ]


def summary_vector(
    values: dict[str, dict[int, float]], method: str
) -> np.ndarray:
    return np.asarray(
        [values[method][subject] for subject in DISCOVERY_SUBJECTS]
    )


def load_existing_robustness(root: Path) -> dict[str, object]:
    output = {}
    for label, folder in (
        ("all_trials", "discovery_wang_cca"),
        ("full_length_trials_only", "discovery_wang_cca_full_only"),
    ):
        payload = json.loads(
            (root / folder / "summary.json").read_text(encoding="utf-8")
        )["models"]
        baseline = payload["full_k1|w=5"]
        candidate = payload["full_equal_k4|w=5"]
        control = payload["full_canonical_k4|w=5"]
        output[label] = {
            "dominant_k1_accuracy": baseline[
                "participant_mean_accuracy"
            ],
            "equal_k4_accuracy": candidate[
                "participant_mean_accuracy"
            ],
            "canonical_k4_accuracy": control[
                "participant_mean_accuracy"
            ],
            "equal_k4_gain_vs_dominant_k1": candidate[
                "participant_mean_gain_vs_full_k1"
            ],
            "participants_improved_vs_dominant_k1": candidate[
                "participants_improved_vs_full_k1"
            ],
            "participant_gains_vs_dominant_k1": candidate[
                "participant_gains_vs_full_k1"
            ],
        }
    return output


def main() -> None:
    parser = argparse.ArgumentParser()
    parser.add_argument(
        "--input-dir",
        type=Path,
        default=Path("data/derived/kul_discovery_32hz"),
    )
    parser.add_argument(
        "--results-root",
        type=Path,
        default=Path("results/kul_aad"),
    )
    parser.add_argument(
        "--output",
        type=Path,
        default=Path(
            "results/kul_aad/discovery_candidate_stress_test.json"
        ),
    )
    args = parser.parse_args()
    manifest = json.loads(
        (args.input_dir / "manifest.json").read_text(encoding="utf-8")
    )
    if tuple(manifest["participants"]) != DISCOVERY_SUBJECTS:
        raise RuntimeError("KUL discovery participant firewall violation")
    if manifest["confirmation_participants_loaded"]:
        raise RuntimeError("KUL confirmation partition is not sealed")
    audio_archive = np.load(
        args.input_dir / str(manifest["audio_cache"])
    )
    audio = {
        name.removeprefix("audio::"): audio_archive[name]
        for name in audio_archive.files
        if name.startswith("audio::")
    }

    records = []
    trial_metadata = []
    for subject in DISCOVERY_SUBJECTS:
        eeg, metadata, attended = base.load_subject(
            args.input_dir, subject, audio
        )
        for held_out_part in (1, 2, 3, 4):
            training_indices = [
                index
                for index, row in enumerate(metadata)
                if int(row["base_part_identity_group"]) != held_out_part
            ]
            testing_indices = [
                index
                for index, row in enumerate(metadata)
                if int(row["base_part_identity_group"]) == held_out_part
            ]
            basis, pca_scale = base.pca_basis(
                [eeg[index] for index in training_indices]
            )
            statistics = base.sufficient_statistics(
                eeg,
                attended,
                training_indices,
                basis,
                pca_scale,
            )
            a, b, canonical = base.fit_cca(*statistics["full"])
            for trial_index in testing_indices:
                row = metadata[trial_index]
                projected = (
                    base.standardize_columns(eeg[trial_index]) @ basis
                ) / pca_scale
                x = base.lag_matrix(projected) @ a[:, :4]
                attended_y = (
                    base.audio_lag_matrix(
                        base.standardize_columns(attended[trial_index])
                    )
                    @ b[:, :4]
                )
                unattended_audio = audio[str(row["unattended_audio"])][
                    : int(row["samples"])
                ]
                unattended_y = (
                    base.audio_lag_matrix(
                        base.standardize_columns(unattended_audio)
                    )
                    @ b[:, :4]
                )
                usable = min(
                    x.shape[0],
                    attended_y.shape[0],
                    unattended_y.shape[0],
                )
                records.append(
                    {
                        "subject": subject,
                        "trial": int(row["trial"]),
                        "held_out_part": held_out_part,
                        "condition": str(row["condition"]),
                        "repetition": bool(row["repetition"]),
                        "attended_ear": str(row["attended_ear"]),
                        "x": x[:usable],
                        "attended": attended_y[:usable],
                        "unattended": unattended_y[:usable],
                        "canonical": canonical[:4],
                    }
                )
                trial_metadata.append(
                    {
                        "subject": subject,
                        "trial": int(row["trial"]),
                        "samples": usable,
                    }
                )
            print(
                f"S{subject:02d}, held-out identity part "
                f"{held_out_part}: projected",
                flush=True,
            )

    observed = participant_accuracies(records)
    rng = np.random.default_rng(RANDOM_SEED)
    null_accuracy = {
        "dominant_k1": [],
        "equal_k4": [],
        "canonical_k4": [],
    }
    null_equal_gain = []
    for replicate in range(NULL_REPLICATES):
        shifts = {}
        for index, record in enumerate(records):
            length = np.asarray(record["x"]).shape[0]
            minimum = 10 * base.FS
            shifts[index] = int(rng.integers(minimum, length - minimum))
        shifted = participant_accuracies(records, shifts=shifts)
        for method in null_accuracy:
            null_accuracy[method].append(
                float(summary_vector(shifted, method).mean())
            )
        null_equal_gain.append(
            float(
                (
                    summary_vector(shifted, "equal_k4")
                    - summary_vector(shifted, "dominant_k1")
                ).mean()
            )
        )
        if (replicate + 1) % 10 == 0:
            print(
                f"Time-shift null {replicate + 1}/{NULL_REPLICATES}",
                flush=True,
            )

    dominant = summary_vector(observed, "dominant_k1")
    equal = summary_vector(observed, "equal_k4")
    canonical = summary_vector(observed, "canonical_k4")
    equal_gain = equal - dominant
    capacity_gain = equal - canonical
    chance_difference = equal - 0.5
    inference_rng = np.random.default_rng(RANDOM_SEED + 1)
    equal_null = np.asarray(null_accuracy["equal_k4"])
    gain_null = np.asarray(null_equal_gain)
    report = {
        "status": (
            "discovery stress test; all participant-level p-values are "
            "descriptive because methods were screened on these participants"
        ),
        "firewall": {
            "participants_loaded": list(DISCOVERY_SUBJECTS),
            "confirmation_participants_loaded": [],
            "confirmation_partition_remains_sealed": True,
        },
        "candidate": {
            "name": "equal-four-projection dual-lag CCA",
            "window_seconds": WINDOW_SECONDS,
            "eeg_lags_ms": [0, 500],
            "audio_lags_ms": [0, 500],
            "cca_regularization": base.REGULARIZATION,
            "aggregation": "equal mean of first four component margins",
        },
        "recomputed_observed": {
            "dominant_k1_participant_mean_accuracy": float(
                dominant.mean()
            ),
            "equal_k4_participant_mean_accuracy": float(equal.mean()),
            "canonical_k4_participant_mean_accuracy": float(
                canonical.mean()
            ),
            "equal_k4_gain_vs_dominant_k1": float(equal_gain.mean()),
            "equal_k4_gain_vs_canonical_k4": float(
                capacity_gain.mean()
            ),
            "participants_improved_vs_dominant_k1": int(
                np.sum(equal_gain > 0)
            ),
            "participants_improved_vs_canonical_k4": int(
                np.sum(capacity_gain > 0)
            ),
            "participant_equal_k4_accuracies": equal.tolist(),
            "participant_gains_vs_dominant_k1": equal_gain.tolist(),
            "participant_gains_vs_canonical_k4": capacity_gain.tolist(),
        },
        "descriptive_participant_inference": {
            "equal_k4_vs_dominant_k1": {
                "mean_gain": float(equal_gain.mean()),
                "bootstrap_95_interval": bootstrap_interval(
                    equal_gain, inference_rng
                ),
                "exact_one_sided_sign_flip_p": exact_sign_flip(
                    equal_gain, "greater"
                ),
                "exact_two_sided_sign_flip_p": exact_sign_flip(
                    equal_gain, "two-sided"
                ),
            },
            "equal_k4_vs_canonical_k4": {
                "mean_gain": float(capacity_gain.mean()),
                "bootstrap_95_interval": bootstrap_interval(
                    capacity_gain, inference_rng
                ),
                "exact_one_sided_sign_flip_p": exact_sign_flip(
                    capacity_gain, "greater"
                ),
            },
            "equal_k4_vs_chance": {
                "mean_difference": float(chance_difference.mean()),
                "bootstrap_95_interval": bootstrap_interval(
                    chance_difference, inference_rng
                ),
                "exact_one_sided_sign_flip_p": exact_sign_flip(
                    chance_difference, "greater"
                ),
            },
        },
        "time_shift_null": {
            "replicates": NULL_REPLICATES,
            "minimum_absolute_shift_seconds": 10,
            "shift_scope": (
                "same independently sampled circular shift applied to both "
                "candidate speech projections within each trial"
            ),
            "equal_k4_accuracy_null_mean": float(equal_null.mean()),
            "equal_k4_accuracy_null_95_interval": [
                float(np.quantile(equal_null, 0.025)),
                float(np.quantile(equal_null, 0.975)),
            ],
            "equal_k4_accuracy_empirical_p": float(
                (1 + np.sum(equal_null >= equal.mean()))
                / (NULL_REPLICATES + 1)
            ),
            "equal_k4_gain_null_mean": float(gain_null.mean()),
            "equal_k4_gain_null_95_interval": [
                float(np.quantile(gain_null, 0.025)),
                float(np.quantile(gain_null, 0.975)),
            ],
            "equal_k4_gain_empirical_p": float(
                (1 + np.sum(gain_null >= equal_gain.mean()))
                / (NULL_REPLICATES + 1)
            ),
        },
        "material_robustness": load_existing_robustness(
            args.results_root
        ),
        "selection_disclosure": {
            "fixed_before_this_stress_test": True,
            "selected_after_screening_multiple_discovery_variants": True,
            "valid_confirmation_requires_untouched_even_participants": True,
        },
    }
    args.output.parent.mkdir(parents=True, exist_ok=True)
    args.output.write_text(json.dumps(report, indent=2), encoding="utf-8")
    print(json.dumps(report, indent=2))


if __name__ == "__main__":
    main()
```
