## Supplementary material for "Falsifiable substitution tests reveal task-structured neural evidence for auditory attention": Article_Analysis_Source_Code: validate_kul_locked_confirmation.py

```
"""Run the one-shot, hash-locked KUL even-participant confirmation."""

from __future__ import annotations

import argparse
import csv
import hashlib
import json
from collections import defaultdict
from pathlib import Path

import numpy as np

import discover_kul_wang_cca as base
from audit_kul_confirmation import (
    CONFIRMATION_SUBJECTS,
    EXPECTED_LOCK_SHA256,
)
from validate_kul_discovery_candidate import (
    bootstrap_interval,
    exact_sign_flip,
    participant_accuracies,
    trial_margins,
)


base.AUDIO_LAG_SAMPLES = 16
BOOTSTRAP_SEED = 20260726
NULL_REPLICATES = 99


def sha256(path: Path) -> str:
    digest = hashlib.sha256()
    with path.open("rb") as handle:
        while block := handle.read(1024 * 1024):
            digest.update(block)
    return digest.hexdigest()


def vector(
    accuracy: dict[str, dict[int, float]], method: str
) -> np.ndarray:
    return np.asarray(
        [accuracy[method][subject] for subject in CONFIRMATION_SUBJECTS]
    )


def write_csv(path: Path, rows: list[dict[str, object]]) -> None:
    with path.open("w", encoding="utf-8", newline="") as handle:
        writer = csv.DictWriter(handle, fieldnames=list(rows[0]))
        writer.writeheader()
        writer.writerows(rows)


def run_evaluation(
    input_dir: Path,
    audio: dict[str, np.ndarray],
    *,
    full_only: bool,
) -> tuple[
    list[dict[str, object]],
    list[dict[str, object]],
    list[dict[str, object]],
]:
    records = []
    trial_rows = []
    fold_rows = []
    for subject in CONFIRMATION_SUBJECTS:
        archive = np.load(input_dir / f"S{subject}.npz")
        metadata = json.loads(str(archive["metadata_json"]))
        eeg = [
            archive[f"eeg_{index:02d}"] for index in range(1, 21)
        ]
        attended = [
            audio[str(row["attended_audio"])][: int(row["samples"])]
            for row in metadata
        ]
        for held_out_part in (1, 2, 3, 4):
            training_indices = [
                index
                for index, row in enumerate(metadata)
                if int(row["base_part_identity_group"]) != held_out_part
                and (not full_only or not bool(row["repetition"]))
            ]
            testing_indices = [
                index
                for index, row in enumerate(metadata)
                if int(row["base_part_identity_group"]) == held_out_part
                and (not full_only or not bool(row["repetition"]))
            ]
            expected = (6, 2) if full_only else (15, 5)
            if (len(training_indices), len(testing_indices)) != expected:
                raise RuntimeError(
                    "KUL confirmation fold cardinality mismatch"
                )
            basis, pca_scale = base.pca_basis(
                [eeg[index] for index in training_indices]
            )
            statistics = base.sufficient_statistics(
                eeg,
                attended,
                training_indices,
                basis,
                pca_scale,
            )
            a, b, canonical = base.fit_cca(*statistics["full"])
            fold_rows.append(
                {
                    "analysis": (
                        "full_only" if full_only else "primary_all_trials"
                    ),
                    "subject": subject,
                    "held_out_part": held_out_part,
                    "training_trials": len(training_indices),
                    "testing_trials": len(testing_indices),
                    **{
                        f"canonical_{component + 1}": float(
                            canonical[component]
                        )
                        for component in range(4)
                    },
                }
            )
            for trial_index in testing_indices:
                row = metadata[trial_index]
                projected = (
                    base.standardize_columns(eeg[trial_index]) @ basis
                ) / pca_scale
                x = base.lag_matrix(projected) @ a[:, :4]
                attended_y = (
                    base.audio_lag_matrix(
                        base.standardize_columns(attended[trial_index])
                    )
                    @ b[:, :4]
                )
                unattended_audio = audio[str(row["unattended_audio"])][
                    : int(row["samples"])
                ]
                unattended_y = (
                    base.audio_lag_matrix(
                        base.standardize_columns(unattended_audio)
                    )
                    @ b[:, :4]
                )
                usable = min(
                    x.shape[0],
                    attended_y.shape[0],
                    unattended_y.shape[0],
                )
                record = {
                    "subject": subject,
                    "trial": int(row["trial"]),
                    "held_out_part": held_out_part,
                    "condition": str(row["condition"]),
                    "repetition": bool(row["repetition"]),
                    "attended_ear": str(row["attended_ear"]),
                    "x": x[:usable],
                    "attended": attended_y[:usable],
                    "unattended": unattended_y[:usable],
                    "canonical": canonical[:4],
                }
                records.append(record)
                margins = trial_margins(
                    record["x"],
                    record["attended"],
                    record["unattended"],
                    record["canonical"],
                )
                for method, values in margins.items():
                    trial_rows.append(
                        {
                            "analysis": (
                                "full_only"
                                if full_only
                                else "primary_all_trials"
                            ),
                            "subject": subject,
                            "trial": int(row["trial"]),
                            "held_out_part": held_out_part,
                            "condition": str(row["condition"]),
                            "repetition": int(bool(row["repetition"])),
                            "attended_ear": str(row["attended_ear"]),
                            "method": method,
                            "windows": int(values.size),
                            "accuracy": float(np.mean(values > 0)),
                            "mean_margin": float(values.mean()),
                        }
                    )
            print(
                f"KUL confirmation S{subject:02d}, part {held_out_part}, "
                f"{'full-only' if full_only else 'primary'}: complete",
                flush=True,
            )
        del archive, eeg, attended
    return records, trial_rows, fold_rows


def strata_summary(
    trial_rows: list[dict[str, object]]
) -> dict[str, object]:
    output = {}
    for field in (
        "condition",
        "repetition",
        "attended_ear",
        "held_out_part",
    ):
        field_output = {}
        levels = sorted({str(row[field]) for row in trial_rows})
        for level in levels:
            method_subject = defaultdict(list)
            for row in trial_rows:
                if str(row[field]) == level:
                    method_subject[
                        (str(row["method"]), int(row["subject"]))
                    ].append(float(row["accuracy"]))
            methods = {}
            for method in ("dominant_k1", "equal_k4", "canonical_k4"):
                participant = np.asarray(
                    [
                        np.mean(method_subject[(method, subject)])
                        for subject in CONFIRMATION_SUBJECTS
                    ]
                )
                methods[method] = float(participant.mean())
            field_output[level] = {
                **methods,
                "equal_k4_gain_vs_dominant_k1": (
                    methods["equal_k4"] - methods["dominant_k1"]
                ),
                "equal_k4_gain_vs_canonical_k4": (
                    methods["equal_k4"] - methods["canonical_k4"]
                ),
            }
        output[field] = field_output
    return output


def main() -> None:
    parser = argparse.ArgumentParser()
    parser.add_argument(
        "--input-dir",
        type=Path,
        default=Path("data/derived/kul_confirmation_32hz"),
    )
    parser.add_argument(
        "--lock-file",
        type=Path,
        default=Path("docs/kul_candidate_lock.json"),
    )
    parser.add_argument(
        "--output-dir",
        type=Path,
        default=Path("results/kul_aad/locked_confirmation"),
    )
    args = parser.parse_args()
    lock_hash = sha256(args.lock_file)
    if lock_hash != EXPECTED_LOCK_SHA256:
        raise RuntimeError(f"KUL candidate lock mismatch: {lock_hash}")
    manifest = json.loads(
        (args.input_dir / "manifest.json").read_text(encoding="utf-8")
    )
    if (
        tuple(manifest["participants"]) != CONFIRMATION_SUBJECTS
        or manifest["discovery_participants_loaded"]
        or manifest["candidate_lock_sha256"] != EXPECTED_LOCK_SHA256
    ):
        raise RuntimeError("KUL confirmation manifest mismatch")
    audio_archive = np.load(Path(manifest["audio_cache"]))
    audio = {
        name.removeprefix("audio::"): audio_archive[name]
        for name in audio_archive.files
        if name.startswith("audio::")
    }

    primary_records, primary_trials, primary_folds = run_evaluation(
        args.input_dir, audio, full_only=False
    )
    full_records, full_trials, full_folds = run_evaluation(
        args.input_dir, audio, full_only=True
    )
    observed = participant_accuracies(primary_records)
    full_observed = participant_accuracies(full_records)
    dominant = vector(observed, "dominant_k1")
    equal = vector(observed, "equal_k4")
    canonical = vector(observed, "canonical_k4")
    gain = equal - dominant
    capacity_gain = equal - canonical
    full_dominant = vector(full_observed, "dominant_k1")
    full_equal = vector(full_observed, "equal_k4")
    full_gain = full_equal - full_dominant

    rng = np.random.default_rng(BOOTSTRAP_SEED)
    primary_p = exact_sign_flip(gain, "greater")
    success = bool(gain.mean() > 0 and primary_p < 0.05)
    null_accuracy = []
    null_gain = []
    for replicate in range(NULL_REPLICATES):
        shifts = {}
        for index, record in enumerate(primary_records):
            length = np.asarray(record["x"]).shape[0]
            minimum = 10 * base.FS
            shifts[index] = int(rng.integers(minimum, length - minimum))
        shifted = participant_accuracies(
            primary_records, shifts=shifts
        )
        shifted_equal = vector(shifted, "equal_k4")
        shifted_dominant = vector(shifted, "dominant_k1")
        null_accuracy.append(float(shifted_equal.mean()))
        null_gain.append(
            float((shifted_equal - shifted_dominant).mean())
        )
        if (replicate + 1) % 10 == 0:
            print(
                f"KUL confirmation null "
                f"{replicate + 1}/{NULL_REPLICATES}",
                flush=True,
            )
    null_accuracy_array = np.asarray(null_accuracy)
    null_gain_array = np.asarray(null_gain)

    participant_rows = [
        {
            "subject": subject,
            "dominant_k1_accuracy": float(dominant[index]),
            "equal_k4_accuracy": float(equal[index]),
            "canonical_k4_accuracy": float(canonical[index]),
            "equal_k4_gain_vs_dominant_k1": float(gain[index]),
            "equal_k4_gain_vs_canonical_k4": float(
                capacity_gain[index]
            ),
            "full_only_dominant_k1_accuracy": float(
                full_dominant[index]
            ),
            "full_only_equal_k4_accuracy": float(full_equal[index]),
            "full_only_gain_vs_dominant_k1": float(full_gain[index]),
        }
        for index, subject in enumerate(CONFIRMATION_SUBJECTS)
    ]
    report = {
        "status": "one-shot hash-locked KUL even-ID confirmation",
        "candidate_lock_sha256": lock_hash,
        "participants": list(CONFIRMATION_SUBJECTS),
        "primary_result": {
            "dominant_k1_participant_mean_accuracy": float(
                dominant.mean()
            ),
            "equal_k4_participant_mean_accuracy": float(equal.mean()),
            "canonical_k4_participant_mean_accuracy": float(
                canonical.mean()
            ),
            "equal_k4_mean_gain_vs_dominant_k1": float(gain.mean()),
            "equal_k4_mean_gain_vs_canonical_k4": float(
                capacity_gain.mean()
            ),
            "participants_improved_vs_dominant_k1": int(
                np.sum(gain > 0)
            ),
            "participants_improved_vs_canonical_k4": int(
                np.sum(capacity_gain > 0)
            ),
            "bootstrap_95_interval_gain_vs_dominant_k1": (
                bootstrap_interval(gain, rng)
            ),
            "exact_one_sided_sign_flip_p_vs_dominant_k1": primary_p,
            "exact_two_sided_sign_flip_p_vs_dominant_k1": (
                exact_sign_flip(gain, "two-sided")
            ),
            "success_criterion_met": success,
        },
        "capacity_control": {
            "mean_gain": float(capacity_gain.mean()),
            "bootstrap_95_interval": bootstrap_interval(
                capacity_gain, rng
            ),
            "exact_one_sided_sign_flip_p": exact_sign_flip(
                capacity_gain, "greater"
            ),
        },
        "accuracy_vs_chance": {
            "mean_difference": float((equal - 0.5).mean()),
            "bootstrap_95_interval": bootstrap_interval(
                equal - 0.5, rng
            ),
            "exact_one_sided_sign_flip_p": exact_sign_flip(
                equal - 0.5, "greater"
            ),
        },
        "full_length_trial_sensitivity": {
            "dominant_k1_accuracy": float(full_dominant.mean()),
            "equal_k4_accuracy": float(full_equal.mean()),
            "mean_gain": float(full_gain.mean()),
            "participants_improved": int(np.sum(full_gain > 0)),
            "bootstrap_95_interval": bootstrap_interval(full_gain, rng),
            "exact_one_sided_sign_flip_p": exact_sign_flip(
                full_gain, "greater"
            ),
        },
        "strata": strata_summary(primary_trials),
        "time_shift_null": {
            "replicates": NULL_REPLICATES,
            "accuracy_null_mean": float(null_accuracy_array.mean()),
            "accuracy_null_95_interval": [
                float(np.quantile(null_accuracy_array, 0.025)),
                float(np.quantile(null_accuracy_array, 0.975)),
            ],
            "accuracy_empirical_p": float(
                (
                    1
                    + np.sum(null_accuracy_array >= float(equal.mean()))
                )
                / (NULL_REPLICATES + 1)
            ),
            "gain_null_mean": float(null_gain_array.mean()),
            "gain_null_95_interval": [
                float(np.quantile(null_gain_array, 0.025)),
                float(np.quantile(null_gain_array, 0.975)),
            ],
            "gain_empirical_p": float(
                (
                    1
                    + np.sum(null_gain_array >= float(gain.mean()))
                )
                / (NULL_REPLICATES + 1)
            ),
        },
        "participant_results": participant_rows,
        "guardrail": (
            "No tuning or alternative primary candidate is permitted after "
            "this result, regardless of success or failure."
        ),
    }
    args.output_dir.mkdir(parents=True, exist_ok=True)
    write_csv(
        args.output_dir / "trial_results.csv",
        primary_trials + full_trials,
    )
    write_csv(
        args.output_dir / "folds.csv", primary_folds + full_folds
    )
    write_csv(args.output_dir / "participants.csv", participant_rows)
    (args.output_dir / "locked_confirmation.json").write_text(
        json.dumps(report, indent=2), encoding="utf-8"
    )
    print(json.dumps(report, indent=2))


if __name__ == "__main__":
    main()
```
