## Supplementary material for "Falsifiable substitution tests reveal task-structured neural evidence for auditory attention": Article_Analysis_Source_Code: validate_projection_analysis_v1.py

```
#!/usr/bin/env python3
"""Validate the public projection-breadth analysis outputs only.

This validator deliberately excludes manuscripts, release documents, internal
logs, exploratory boundary datasets, and formatted supplementary copies.  It
is suitable for the raw-to-result workflow distributed with the article.
"""

from __future__ import annotations

import csv
import hashlib
import json
from pathlib import Path

import numpy as np


ROOT = Path(__file__).resolve().parents[1]


def digest(path: Path) -> str:
    value = hashlib.sha256()
    with path.open("rb") as handle:
        for block in iter(lambda: handle.read(1024 * 1024), b""):
            value.update(block)
    return value.hexdigest()


def load_json(relative: str) -> dict:
    return json.loads((ROOT / relative).read_text(encoding="utf-8"))


def load_csv(relative: str) -> list[dict[str, str]]:
    with (ROOT / relative).open("r", encoding="utf-8", newline="") as handle:
        return list(csv.DictReader(handle))


def assert_close(observed: float, expected: float, label: str) -> None:
    if not np.isclose(observed, expected, atol=1e-12, rtol=0):
        raise AssertionError(f"{label}: {observed} != {expected}")


def verify_locks() -> None:
    expected = {
        "docs/dtu_external_lock.json":
            "f87d8bf4ac291d2c1f2bb57138f0175335dc58a9624a205fbf5c5777dcf89440",
        "docs/kul_candidate_lock.json":
            "0e152a85fc86620d325a2414c683049ede0ccac0a29ab536bd73c899885886fc",
        "docs/ear_saad_multimodal_external_lock.json":
            "2afcc6451d06625a521f10abd967d8618babc02dd1c649a8c97aee14b5926827",
    }
    for relative, expected_hash in expected.items():
        observed = digest(ROOT / relative)
        if observed != expected_hash:
            raise AssertionError(f"lock hash changed: {relative}: {observed}")


def verify_discovery_outputs() -> None:
    standard = load_json("results/kul_aad/discovery_wang_cca/summary.json")
    full_only = load_json(
        "results/kul_aad/discovery_wang_cca_full_only/summary.json"
    )
    stress = load_json("results/kul_aad/discovery_candidate_stress_test.json")
    for label, payload in (("standard", standard), ("full-only", full_only)):
        if "design" not in payload or "models" not in payload:
            raise AssertionError(f"incomplete KUL {label} discovery summary")
    if not stress.get("status") or not stress.get("candidate"):
        raise AssertionError("KUL discovery candidate stress test is incomplete")
    firewall = stress.get("firewall", {})
    if firewall.get("confirmation_participants_loaded"):
        raise AssertionError("KUL confirmation data entered discovery validation")
    if not firewall.get("confirmation_partition_remains_sealed"):
        raise AssertionError("KUL confirmation partition was not sealed")


def verify_locked_outputs() -> None:
    kul = load_json(
        "results/kul_aad/locked_confirmation/locked_confirmation.json"
    )
    dtu = load_json("results/dtu_aad/locked_external/locked_external.json")
    ear = load_json(
        "results/ear_saad/locked_multimodal_external/locked_external.json"
    )
    synthesis = load_json(
        "results/multimodal_projection_breadth_synthesis/synthesis.json"
    )
    kul_rows = load_csv("results/kul_aad/locked_confirmation/participants.csv")
    dtu_rows = load_csv("results/dtu_aad/locked_external/participants.csv")
    ear_rows = load_csv(
        "results/ear_saad/locked_multimodal_external/primary_participants.csv"
    )

    if (len(kul_rows), len(dtu_rows), len(ear_rows)) != (8, 18, 15):
        raise AssertionError("unexpected locked participant count")
    if {int(row["subject"]) for row in kul_rows} != set(range(2, 17, 2)):
        raise AssertionError("KUL confirmation is not the even-ID partition")
    if {int(row["subject"]) for row in dtu_rows} != set(range(1, 19)):
        raise AssertionError("DTU participant mismatch")
    if {int(row["subject"]) for row in ear_rows} != set(range(1, 16)):
        raise AssertionError("Ear-SAAD participant mismatch")

    kul_gain = np.asarray(
        [float(row["equal_k4_gain_vs_dominant_k1"]) for row in kul_rows]
    )
    dtu_gain = np.asarray(
        [float(row["equal_k4_gain_vs_dominant_k1"]) for row in dtu_rows]
    )
    ear_gain = np.asarray([float(row["gain_equal_minus_k1"]) for row in ear_rows])
    assert_close(
        kul_gain.mean(),
        kul["primary_result"]["equal_k4_mean_gain_vs_dominant_k1"],
        "KUL mean gain",
    )
    assert_close(
        dtu_gain.mean(),
        dtu["locked_primary_inference"]["mean_gain"],
        "DTU mean gain",
    )
    assert_close(
        ear_gain.mean(),
        ear["primary_equal_modality_average"]["equal_k4_minus_k1_mean"],
        "Ear-SAAD mean gain",
    )

    locked = synthesis["independent_locked_synthesis"]
    all_gains = np.concatenate([kul_gain, dtu_gain, ear_gain])
    if locked["number_of_studies"] != 3 or locked["participants_total"] != 41:
        raise AssertionError("projection synthesis sample-size mismatch")
    if locked["participants_improved_total"] != 32:
        raise AssertionError("projection synthesis positive-participant mismatch")
    assert_close(
        all_gains.mean(),
        locked["participant_weighted_mean_gain"],
        "participant-weighted synthesis",
    )
    assert_close(
        np.mean([kul_gain.mean(), dtu_gain.mean(), ear_gain.mean()]),
        locked["study_equal_mean_gain"],
        "study-equal synthesis",
    )


def main() -> None:
    verify_locks()
    verify_discovery_outputs()
    verify_locked_outputs()
    print("Projection-breadth analysis audit: PASS")


if __name__ == "__main__":
    main()
```
