## Supplementary material for "Falsifiable substitution tests reveal task-structured neural evidence for auditory attention": Article_Analysis_Source_Code: wz_hic_asme30_core_v1.py

```
"""Frozen computational core for WZ-HIC-ASME30 external validation v1."""

from __future__ import annotations

import itertools
import math

import numpy as np
from sklearn.metrics import roc_auc_score

from wz_hif_core_v1 import fit_shrinkage_lda


COMMANDS = tuple(range(1, 31))
PHYSICAL_GROUPS = (
    (17, 23, 5, 18, 20, 25, 21, 9, 15, 16),
    (1, 19, 4, 6, 7, 8, 10, 11, 12, 29),
    (26, 24, 3, 22, 2, 14, 13, 27, 28, 30),
)
WRONG_GROUPS = (
    (
        tuple(range(1, 31, 3)),
        tuple(range(2, 31, 3)),
        tuple(range(3, 31, 3)),
    ),
    (
        tuple(range(1, 11)),
        tuple(range(11, 21)),
        tuple(range(21, 31)),
    ),
)
BIN_EDGES_S = np.round(np.arange(-0.1, 1.0001, 0.1), 10)


def validate_groups(groups) -> tuple[tuple[int, ...], ...]:
    normalized = tuple(tuple(int(x) for x in group) for group in groups)
    flattened = [x for group in normalized for x in group]
    if len(normalized) != 3 or any(len(group) != 10 for group in normalized):
        raise ValueError("Each mapping must contain three groups of ten.")
    if sorted(flattened) != list(COMMANDS):
        raise ValueError("Mapping must cover commands 1..30 exactly once.")
    return normalized


def group_index(command: int, groups=PHYSICAL_GROUPS) -> int:
    groups = validate_groups(groups)
    for index, group in enumerate(groups):
        if int(command) in group:
            return index
    raise ValueError(command)


def labels_for_mapping(
    command: np.ndarray, target: np.ndarray, groups=PHYSICAL_GROUPS
) -> np.ndarray:
    """Return T/S/U labels for an incidence mapping."""

    groups = validate_groups(groups)
    command = np.asarray(command, dtype=int)
    target = np.asarray(target, dtype=int)
    labels = np.full(command.size, "U", dtype="U1")
    labels[command == target] = "T"
    lookup = np.empty(31, dtype=int)
    lookup.fill(-1)
    for i, group in enumerate(groups):
        lookup[list(group)] = i
    same = lookup[command] == lookup[target]
    labels[same & (command != target)] = "S"
    if np.any(lookup[command] < 0) or np.any(lookup[target] < 0):
        raise ValueError("Unknown command.")
    return labels


def moments(values: np.ndarray) -> tuple[float, float]:
    values = np.asarray(values, dtype=float)
    mu = float(np.mean(values))
    sd = float(np.std(values, ddof=1))
    if not np.isfinite(sd) or sd < 1e-12:
        sd = 1.0
    return mu, sd


def fit_zscore_apply(
    train_x: np.ndarray,
    train_y: np.ndarray,
    test_x: np.ndarray,
) -> tuple[np.ndarray, dict]:
    model = fit_shrinkage_lda(train_x, train_y)
    train_score = model.decision_function(train_x)
    mu, sd = moments(train_score)
    test_score = (model.decision_function(test_x) - mu) / sd
    if not np.isfinite(test_score).all():
        raise RuntimeError("Nonfinite external event score.")
    return test_score, {
        "n_train": int(train_y.size),
        "n_train_negative": int(np.sum(train_y == 0)),
        "n_train_positive": int(np.sum(train_y == 1)),
        "training_score_mean": mu,
        "training_score_sd": sd,
    }


def fit_primary_event_scores(
    *,
    train_eeg_outer: np.ndarray,
    train_eog_outer: np.ndarray,
    train_prestim: np.ndarray,
    train_flat: np.ndarray,
    train_command: np.ndarray,
    train_target: np.ndarray,
    test_eeg_outer: np.ndarray,
    test_eog_outer: np.ndarray,
    test_prestim: np.ndarray,
    test_flat: np.ndarray,
) -> tuple[dict[str, np.ndarray], dict]:
    labels = labels_for_mapping(train_command, train_target, PHYSICAL_GROUPS)
    outer = np.isin(labels, ["S", "U"])
    y_outer = (labels[outer] == "S").astype(int)
    y_flat = (labels == "T").astype(int)
    inputs = {
        "eeg_outer": (train_eeg_outer[outer], y_outer, test_eeg_outer),
        "eog_outer": (train_eog_outer[outer], y_outer, test_eog_outer),
        "prestim_outer": (train_prestim[outer], y_outer, test_prestim),
        "flat": (train_flat, y_flat, test_flat),
    }
    scores, fits = {}, {}
    for name, (train_x, y, test_x) in inputs.items():
        scores[name], fits[name] = fit_zscore_apply(train_x, y, test_x)
    return scores, fits


def fit_wrong_mapping_score(
    train_eeg_outer: np.ndarray,
    train_command: np.ndarray,
    train_target: np.ndarray,
    test_eeg_outer: np.ndarray,
    groups,
) -> tuple[np.ndarray, dict]:
    labels = labels_for_mapping(train_command, train_target, groups)
    outer = np.isin(labels, ["S", "U"])
    y = (labels[outer] == "S").astype(int)
    return fit_zscore_apply(train_eeg_outer[outer], y, test_eeg_outer)


def trial_rows(
    *,
    flat_score: np.ndarray,
    outer_score: np.ndarray,
    command: np.ndarray,
    trial: np.ndarray,
    target: np.ndarray,
    horizon: int = 3,
    pooling_groups=PHYSICAL_GROUPS,
    correctness_groups=PHYSICAL_GROUPS,
) -> list[dict]:
    pooling_groups = validate_groups(pooling_groups)
    correctness_groups = validate_groups(correctness_groups)
    command = np.asarray(command, dtype=int)
    trial = np.asarray(trial)
    target = np.asarray(target, dtype=int)
    rows = []
    for trial_id in sorted(np.unique(trial)):
        ids = np.flatnonzero(trial == trial_id)
        target_values = np.unique(target[ids])
        if target_values.size != 1:
            raise RuntimeError(f"Trial {trial_id} has nonunique targets.")
        if any(np.sum(command[ids] == c) < horizon for c in COMMANDS):
            continue
        command_ids = {
            c: ids[command[ids] == c][:horizon] for c in COMMANDS
        }
        child_scores = {
            c: float(np.mean(flat_score[command_ids[c]])) for c in COMMANDS
        }
        ordered = sorted(
            COMMANDS, key=lambda c: (child_scores[c], -c), reverse=True
        )
        prediction = int(ordered[0])
        margin = child_scores[ordered[0]] - child_scores[ordered[1]]
        parent_scores = []
        for group in pooling_groups:
            group_ids = np.concatenate([command_ids[c] for c in group])
            parent_scores.append(float(np.mean(outer_score[group_ids])))
        predicted_parent = group_index(prediction, pooling_groups)
        other_max = max(
            score
            for i, score in enumerate(parent_scores)
            if i != predicted_parent
        )
        certificate = parent_scores[predicted_parent] - other_max
        outer_parent = int(np.argmax(parent_scores))
        true_target = int(target_values[0])
        rows.append(
            {
                "trial": int(trial_id),
                "target": true_target,
                "flat_prediction": prediction,
                "flat_correct": prediction == true_target,
                "parent_correct": (
                    group_index(prediction, correctness_groups)
                    == group_index(true_target, correctness_groups)
                ),
                "agreement": predicted_parent == outer_parent,
                "flat_margin": float(margin),
                "signed_certificate": float(certificate),
                "predicted_pooling_parent": predicted_parent,
                "outer_pooling_parent": outer_parent,
            }
        )
    return rows


def auc_or_nan(y: np.ndarray, score: np.ndarray) -> float:
    if np.unique(y).size != 2:
        return float("nan")
    return float(roc_auc_score(y.astype(int), score))


def certificate_metrics(rows: list[dict]) -> dict:
    if not rows:
        return {
            "n_trials": 0,
            "n_parent_correct": 0,
            "n_parent_error": 0,
            "certificate_AUC": float("nan"),
            "flat_margin_AUC": float("nan"),
            "flat_accuracy": float("nan"),
            "parent_accuracy": float("nan"),
            "agreement_coverage": float("nan"),
            "agreement_accuracy": float("nan"),
            "agreement_accuracy_gain": float("nan"),
        }
    y = np.asarray([row["parent_correct"] for row in rows], dtype=int)
    certificate = np.asarray(
        [row["signed_certificate"] for row in rows], dtype=float
    )
    margin = np.asarray([row["flat_margin"] for row in rows], dtype=float)
    correct = np.asarray([row["flat_correct"] for row in rows], dtype=bool)
    agreement = np.asarray([row["agreement"] for row in rows], dtype=bool)
    flat_accuracy = float(np.mean(correct))
    agreement_accuracy = (
        float(np.mean(correct[agreement])) if np.any(agreement) else float("nan")
    )
    return {
        "n_trials": len(rows),
        "n_parent_correct": int(np.sum(y)),
        "n_parent_error": int(np.sum(1 - y)),
        "certificate_AUC": auc_or_nan(y, certificate),
        "flat_margin_AUC": auc_or_nan(y, margin),
        "flat_accuracy": flat_accuracy,
        "parent_accuracy": float(np.mean(y)),
        "agreement_coverage": float(np.mean(agreement)),
        "agreement_accuracy": agreement_accuracy,
        "agreement_accuracy_gain": agreement_accuracy - flat_accuracy,
    }


def exact_sign_p(values: list[float], null: float = 0.0) -> float:
    a = np.asarray(values, dtype=float)
    k = int(np.sum(a > null))
    n = int(a.size)
    return float(sum(math.comb(n, j) for j in range(k, n + 1)) / 2**n)


def exact_signflip_mean_p(values: list[float]) -> float:
    a = np.asarray(values, dtype=float)
    observed = float(np.mean(a))
    permuted = [
        float(np.mean(a * np.asarray(signs)))
        for signs in itertools.product((-1.0, 1.0), repeat=a.size)
    ]
    return float(np.mean(np.asarray(permuted) >= observed - 1e-15))


def bootstrap_mean_ci(
    values: list[float], *, seed: int = 20260729, n_boot: int = 20_000
) -> list[float]:
    a = np.asarray(values, dtype=float)
    rng = np.random.default_rng(seed)
    means = a[rng.integers(0, a.size, size=(n_boot, a.size))].mean(axis=1)
    return [float(x) for x in np.quantile(means, [0.025, 0.975])]
```
