## Supplementary material for "Falsifiable substitution tests reveal task-structured neural evidence for auditory attention": Article_Analysis_Source_Code: wz_hic_confirmation_core_v1.py

```
"""Frozen computational core for WZ-HIC-v1 confirmation."""

from __future__ import annotations

import itertools
import math

import numpy as np
from sklearn.metrics import roc_auc_score

from wz_hif_core_v1 import fit_shrinkage_lda


TRUE_PAIRING = ((1, 2), (3, 4))
WRONG_PAIRINGS = (((1, 3), (2, 4)), ((1, 4), (2, 3)))


def pair_for(command: int, pairing) -> tuple[int, int]:
    return next(pair for pair in pairing if command in pair)


def moments(x: np.ndarray) -> tuple[float, float]:
    mu = float(np.mean(x))
    sd = float(np.std(x, ddof=1))
    return mu, sd if np.isfinite(sd) and sd > 1e-12 else 1.0


def loro_event_scores(
    eeg_outer: np.ndarray,
    eog_outer: np.ndarray,
    eeg_prestim: np.ndarray,
    flat_features: np.ndarray,
    labels: np.ndarray,
    runs: np.ndarray,
) -> dict[str, np.ndarray]:
    score = {
        "eeg_outer": np.full(labels.size, np.nan),
        "eog_outer": np.full(labels.size, np.nan),
        "prestim_outer": np.full(labels.size, np.nan),
        "flat": np.full(labels.size, np.nan),
    }
    for held_out in sorted(np.unique(runs)):
        train = runs != held_out
        test = ~train
        outer_train = train & np.isin(labels, ["S", "U"])
        y_outer = (labels[outer_train] == "S").astype(int)
        y_flat = (labels[train] == "T").astype(int)
        models = {
            "eeg_outer": (
                fit_shrinkage_lda(eeg_outer[outer_train], y_outer),
                eeg_outer,
                outer_train,
            ),
            "eog_outer": (
                fit_shrinkage_lda(eog_outer[outer_train], y_outer),
                eog_outer,
                outer_train,
            ),
            "prestim_outer": (
                fit_shrinkage_lda(eeg_prestim[outer_train], y_outer),
                eeg_prestim,
                outer_train,
            ),
            "flat": (
                fit_shrinkage_lda(flat_features[train], y_flat),
                flat_features,
                train,
            ),
        }
        for name, (model, x, train_mask) in models.items():
            train_score = model.decision_function(x[train_mask])
            mu, sd = moments(train_score)
            score[name][test] = (model.decision_function(x[test]) - mu) / sd
    if any(not np.isfinite(v).all() for v in score.values()):
        raise RuntimeError("Nonfinite LORO event score.")
    return score


def pseudo_labels(
    command: np.ndarray, target: np.ndarray, pairing
) -> np.ndarray:
    labels = np.full(command.size, "U", dtype="U1")
    labels[command == target] = "T"
    for pair in pairing:
        same_parent = np.isin(target, pair) & np.isin(command, pair)
        labels[same_parent & (command != target)] = "S"
    return labels


def trial_rows(
    scores: dict[str, np.ndarray],
    command: np.ndarray,
    trial: np.ndarray,
    target: np.ndarray,
    *,
    horizon: int,
    certificate_name: str,
    pooling_pairing=TRUE_PAIRING,
    correctness_pairing=TRUE_PAIRING,
) -> list[dict]:
    rows = []
    for trial_id in sorted(np.unique(trial)):
        ids = np.flatnonzero(trial == trial_id)
        if any(np.sum(command[ids] == c) < horizon for c in (1, 2, 3, 4)):
            continue
        command_ids = {
            c: ids[command[ids] == c][:horizon] for c in (1, 2, 3, 4)
        }
        flat_score = {
            c: float(np.mean(scores["flat"][command_ids[c]]))
            for c in (1, 2, 3, 4)
        }
        order = sorted(flat_score, key=flat_score.get, reverse=True)
        prediction = order[0]
        flat_margin = flat_score[order[0]] - flat_score[order[1]]
        parent_score = {}
        for pair in pooling_pairing:
            ids_pair = np.concatenate([command_ids[c] for c in pair])
            parent_score[pair] = float(
                np.mean(scores[certificate_name][ids_pair])
            )
        predicted_pooling_parent = pair_for(prediction, pooling_pairing)
        other_pooling_parent = next(
            p for p in pooling_pairing if p != predicted_pooling_parent
        )
        signed_certificate = (
            parent_score[predicted_pooling_parent]
            - parent_score[other_pooling_parent]
        )
        outer_parent = max(parent_score, key=parent_score.get)
        true_target = int(np.unique(target[ids])[0])
        rows.append(
            {
                "trial": int(trial_id),
                "target": true_target,
                "flat_prediction": prediction,
                "flat_correct": prediction == true_target,
                "parent_correct": (
                    pair_for(prediction, correctness_pairing)
                    == pair_for(true_target, correctness_pairing)
                ),
                "agreement": predicted_pooling_parent == outer_parent,
                "flat_margin": flat_margin,
                "signed_certificate": signed_certificate,
            }
        )
    return rows


def auc_or_nan(y: np.ndarray, score: np.ndarray) -> float:
    if np.unique(y).size != 2:
        return float("nan")
    return float(roc_auc_score(y.astype(int), score))


def certificate_metrics(rows: list[dict]) -> dict:
    if not rows:
        return {
            "n_trials": 0,
            "n_parent_correct": 0,
            "n_parent_error": 0,
            "certificate_AUC": float("nan"),
            "flat_margin_AUC": float("nan"),
            "flat_accuracy": float("nan"),
            "agreement_coverage": float("nan"),
            "agreement_accuracy": float("nan"),
            "agreement_accuracy_gain": float("nan"),
        }
    y = np.asarray([r["parent_correct"] for r in rows], dtype=int)
    cert = np.asarray([r["signed_certificate"] for r in rows], dtype=float)
    margin = np.asarray([r["flat_margin"] for r in rows], dtype=float)
    correct = np.asarray([r["flat_correct"] for r in rows], dtype=int)
    agreement = np.asarray([r["agreement"] for r in rows], dtype=bool)
    flat_accuracy = float(correct.mean())
    agreement_accuracy = (
        float(correct[agreement].mean()) if np.any(agreement) else float("nan")
    )
    return {
        "n_trials": int(len(rows)),
        "n_parent_correct": int(y.sum()),
        "n_parent_error": int((1 - y).sum()),
        "certificate_AUC": auc_or_nan(y, cert),
        "flat_margin_AUC": auc_or_nan(y, margin),
        "flat_accuracy": flat_accuracy,
        "agreement_coverage": float(agreement.mean()),
        "agreement_accuracy": agreement_accuracy,
        "agreement_accuracy_gain": agreement_accuracy - flat_accuracy,
    }


def exact_sign_p(values: list[float], null: float = 0.0) -> float:
    a = np.asarray(values, dtype=float)
    k = int(np.sum(a > null))
    n = int(a.size)
    return float(sum(math.comb(n, j) for j in range(k, n + 1)) / 2**n)


def exact_signflip_mean_p(values: list[float]) -> float:
    """One-sided exhaustive randomization P(mean(sign*d) >= observed mean)."""
    a = np.asarray(values, dtype=float)
    observed = float(a.mean())
    permuted = []
    for signs in itertools.product((-1.0, 1.0), repeat=a.size):
        permuted.append(float(np.mean(a * np.asarray(signs))))
    return float(np.mean(np.asarray(permuted) >= observed - 1e-15))


def bootstrap_mean_ci(
    values: list[float], *, seed: int = 20260729, n_boot: int = 20000
) -> list[float]:
    a = np.asarray(values, dtype=float)
    rng = np.random.default_rng(seed)
    means = a[rng.integers(0, a.size, size=(n_boot, a.size))].mean(axis=1)
    return [float(x) for x in np.quantile(means, [0.025, 0.975])]
```
