## Supplementary material for "Falsifiable substitution tests reveal task-structured neural evidence for auditory attention": Article_Analysis_Source_Code: wz_hif_core_v1.py

```
"""Core, outcome-independent implementation for WZ-HIF-v1."""

from __future__ import annotations

import math
from dataclasses import dataclass
from itertools import product
from typing import Iterable

import numpy as np
from scipy.special import expit
from sklearn.covariance import ledoit_wolf
from sklearn.metrics import roc_auc_score


BIN_EDGES_S = np.round(np.arange(-0.1, 1.0001, 0.1), 10)
TRUE_PAIRING = ((1, 2), (3, 4))
WRONG_PAIRINGS = (((1, 3), (2, 4)), ((1, 4), (2, 3)))


def exact_one_sided_sign_p(differences: Iterable[float]) -> float:
    """P[X >= observed positives], X~Binom(nonzero, .5); zeros are failures."""
    x = np.asarray(list(differences), dtype=float)
    k = int(np.sum(x > 0))
    n = int(x.size)
    return float(sum(math.comb(n, j) for j in range(k, n + 1)) / (2**n))


def bootstrap_mean_ci(
    values: Iterable[float], seed: int = 20260729, n_boot: int = 20000
) -> tuple[float, float]:
    x = np.asarray(list(values), dtype=float)
    rng = np.random.default_rng(seed)
    means = x[rng.integers(0, x.size, size=(n_boot, x.size))].mean(axis=1)
    return tuple(float(v) for v in np.quantile(means, [0.025, 0.975]))


def window_bin_indices(window_s: tuple[float, float]) -> np.ndarray:
    lo, hi = window_s
    starts = BIN_EDGES_S[:-1]
    ends = BIN_EDGES_S[1:]
    idx = np.flatnonzero((starts >= lo - 1e-9) & (ends <= hi + 1e-9))
    if idx.size == 0:
        raise ValueError(f"No complete 100-ms bin in {window_s}.")
    return idx


def select_window_features(
    x_bins: np.ndarray, window_s: tuple[float, float]
) -> np.ndarray:
    """Select bins from [event, channel, bin], then flatten channel-major."""
    idx = window_bin_indices(window_s)
    return x_bins[:, :, idx].reshape(x_bins.shape[0], -1)


@dataclass
class ShrinkageLDA:
    mean: np.ndarray
    scale: np.ndarray
    direction: np.ndarray
    intercept: float

    def decision_function(self, x: np.ndarray) -> np.ndarray:
        z = (np.asarray(x, dtype=float) - self.mean) / self.scale
        return z @ self.direction + self.intercept


def fit_shrinkage_lda(
    x: np.ndarray, y: np.ndarray, *, max_train_per_class: int | None = None
) -> ShrinkageLDA:
    """Equal-prior binary Ledoit-Wolf LDA with deterministic optional thinning."""
    x = np.asarray(x, dtype=float)
    y = np.asarray(y, dtype=int)
    if set(np.unique(y)) != {0, 1}:
        raise ValueError("Both binary classes are required.")

    if max_train_per_class is not None:
        keep = []
        for label in (0, 1):
            ids = np.flatnonzero(y == label)
            if ids.size > max_train_per_class:
                positions = np.linspace(
                    0, ids.size - 1, max_train_per_class, dtype=int
                )
                ids = ids[positions]
            keep.append(ids)
        keep_idx = np.sort(np.concatenate(keep))
        x, y = x[keep_idx], y[keep_idx]

    mean = x.mean(axis=0)
    scale = x.std(axis=0, ddof=1)
    scale[~np.isfinite(scale) | (scale < 1e-12)] = 1.0
    z = (x - mean) / scale
    m0 = z[y == 0].mean(axis=0)
    m1 = z[y == 1].mean(axis=0)
    centered = np.vstack((z[y == 0] - m0, z[y == 1] - m1))
    covariance, _ = ledoit_wolf(centered, assume_centered=True)
    delta = m1 - m0
    try:
        direction = np.linalg.solve(covariance, delta)
    except np.linalg.LinAlgError:
        direction = np.linalg.pinv(covariance, hermitian=True) @ delta
    intercept = float(-0.5 * (m1 + m0) @ direction)
    return ShrinkageLDA(mean, scale, direction, intercept)


def score_binary_loro(
    x: np.ndarray,
    y: np.ndarray,
    runs: np.ndarray,
    *,
    max_train_per_class: int | None = None,
) -> np.ndarray:
    scores = np.full(len(y), np.nan, dtype=float)
    for held_out in sorted(np.unique(runs)):
        train = runs != held_out
        test = ~train
        model = fit_shrinkage_lda(
            x[train], y[train], max_train_per_class=max_train_per_class
        )
        scores[test] = model.decision_function(x[test])
    if not np.isfinite(scores).all():
        raise RuntimeError("Nonfinite LORO score.")
    return scores


def participant_certificate_auc(
    x_bins: np.ndarray,
    labels: np.ndarray,
    runs: np.ndarray,
    windows: list[tuple[float, float]],
    *,
    positive_label: str,
    negative_label: str,
    max_train_per_class: int | None = None,
) -> dict[str, float]:
    keep = np.isin(labels, [negative_label, positive_label])
    y = (labels[keep] == positive_label).astype(int)
    out: dict[str, float] = {}
    for window in windows:
        x = select_window_features(x_bins[keep], window)
        score = score_binary_loro(
            x, y, runs[keep], max_train_per_class=max_train_per_class
        )
        out[f"{window[0]:.1f}_{window[1]:.1f}"] = float(roc_auc_score(y, score))
    return out


def _training_score_moments(scores: np.ndarray) -> tuple[float, float]:
    mean = float(np.mean(scores))
    sd = float(np.std(scores, ddof=1))
    return mean, sd if np.isfinite(sd) and sd > 1e-12 else 1.0


def _pair_for_command(
    command: int, pairing: tuple[tuple[int, int], tuple[int, int]]
) -> tuple[int, int]:
    for pair in pairing:
        if command in pair:
            return pair
    raise ValueError(command)


def _trial_predictions(
    event_command: np.ndarray,
    outer_z: np.ndarray,
    inner_z: np.ndarray,
    flat_z: np.ndarray,
    trial_id: np.ndarray,
    target_command: np.ndarray,
    horizon: int,
    lam: float,
    pairing: tuple[tuple[int, int], tuple[int, int]],
) -> tuple[np.ndarray, np.ndarray, np.ndarray]:
    true, factorized, flat = [], [], []
    for trial in sorted(np.unique(trial_id)):
        ids = np.flatnonzero(trial_id == trial)
        target_values = np.unique(target_command[ids])
        if target_values.size != 1:
            raise RuntimeError("Trial has nonunique target.")
        if any(
            np.sum(event_command[ids] == command) < horizon
            for command in (1, 2, 3, 4)
        ):
            # Artifact rejection can make a trial ineligible at a long horizon.
            # The whole trial is removed for both methods and every pairing.
            continue
        f_scores, b_scores = [], []
        for command in (1, 2, 3, 4):
            cmd_ids = ids[event_command[ids] == command][:horizon]
            pair = _pair_for_command(command, pairing)
            parent_ids = np.concatenate(
                [ids[event_command[ids] == member][:horizon] for member in pair]
            )
            f_scores.append(
                float(np.mean(outer_z[parent_ids]) + lam * np.mean(inner_z[cmd_ids]))
            )
            b_scores.append(float(np.mean(flat_z[cmd_ids])))
        true.append(int(target_values[0]))
        factorized.append(int(np.argmax(f_scores) + 1))
        flat.append(int(np.argmax(b_scores) + 1))
    return np.asarray(true), np.asarray(factorized), np.asarray(flat)


def participant_decoder_loro(
    x_outer: np.ndarray,
    x_inner: np.ndarray,
    x_flat: np.ndarray,
    labels: np.ndarray,
    event_command: np.ndarray,
    runs: np.ndarray,
    trial_id: np.ndarray,
    target_command: np.ndarray,
    *,
    horizons: list[int],
    lambdas: list[float],
    pairings: tuple[
        tuple[tuple[int, int], tuple[int, int]], ...
    ] = (TRUE_PAIRING,) + WRONG_PAIRINGS,
    max_train_per_class: int | None = None,
) -> dict:
    """Cross-validated event scores and trial-level four-command predictions."""
    fold_scores: dict[int, dict[str, np.ndarray]] = {}
    for held_out in sorted(np.unique(runs)):
        train = runs != held_out
        test = ~train

        outer_train = train & np.isin(labels, ["S", "U"])
        inner_train = train & np.isin(labels, ["T", "S"])
        flat_train = train
        outer_y = (labels[outer_train] == "S").astype(int)
        inner_y = (labels[inner_train] == "T").astype(int)
        flat_y = (labels[flat_train] == "T").astype(int)

        outer_model = fit_shrinkage_lda(
            x_outer[outer_train],
            outer_y,
            max_train_per_class=max_train_per_class,
        )
        inner_model = fit_shrinkage_lda(
            x_inner[inner_train],
            inner_y,
            max_train_per_class=max_train_per_class,
        )
        flat_model = fit_shrinkage_lda(
            x_flat[flat_train], flat_y, max_train_per_class=max_train_per_class
        )

        # Fold normalization moments come exclusively from training observations.
        outer_mu, outer_sd = _training_score_moments(
            outer_model.decision_function(x_outer[outer_train])
        )
        inner_mu, inner_sd = _training_score_moments(
            inner_model.decision_function(x_inner[inner_train])
        )
        flat_mu, flat_sd = _training_score_moments(
            flat_model.decision_function(x_flat[flat_train])
        )
        fold_scores[int(held_out)] = {
            "test": test,
            "outer": (
                outer_model.decision_function(x_outer[test]) - outer_mu
            )
            / outer_sd,
            "inner": (
                inner_model.decision_function(x_inner[test]) - inner_mu
            )
            / inner_sd,
            "flat": (flat_model.decision_function(x_flat[test]) - flat_mu) / flat_sd,
        }

    results: dict = {"pairings": {}}
    for pairing in pairings:
        pairing_key = "-".join("".join(str(v) for v in p) for p in pairing)
        results["pairings"][pairing_key] = {}
        for lam, horizon in product(lambdas, horizons):
            correct_f, correct_b, n_trials = 0, 0, 0
            for held_out, item in fold_scores.items():
                test = item["test"]
                true, pred_f, pred_b = _trial_predictions(
                    event_command[test],
                    item["outer"],
                    item["inner"],
                    item["flat"],
                    trial_id[test],
                    target_command[test],
                    horizon,
                    lam,
                    pairing,
                )
                correct_f += int(np.sum(pred_f == true))
                correct_b += int(np.sum(pred_b == true))
                n_trials += int(true.size)
            key = f"lambda_{lam:g}_m_{horizon}"
            results["pairings"][pairing_key][key] = {
                "factorized_accuracy": (
                    correct_f / n_trials if n_trials else float("nan")
                ),
                "flat_accuracy": correct_b / n_trials if n_trials else float("nan"),
                "gain": (
                    (correct_f - correct_b) / n_trials
                    if n_trials
                    else float("nan")
                ),
                "n_trials": n_trials,
            }
    return results


def sigmoid_accuracy_surrogate(scores: np.ndarray) -> np.ndarray:
    """Only used by synthetic tests for monotonicity diagnostics."""
    return expit(np.asarray(scores))
```
