## Supplementary material for "Falsifiable substitution tests reveal task-structured neural evidence for auditory attention": Derived_Result_Verification_Archive: 00_README.md

# Derived-result verification archive

## Scope

This archive checks 33 ledger-indexed quantitative claims, runs a broader arithmetic audit implemented separately from the claim-ledger checker, recomputes selected participant-level summaries, and supplies the final inputs and code that render all six article figures. "Ledger-indexed" describes the public claim-to-file crosswalk; it does not mean that the study or claims were publicly preregistered. This is not a raw-EEG-to-result workflow. Raw EEG, audio, third-party papers, historical drafts, internal logs, and local machine paths are not included. Upstream feature extraction and model fitting are provided in a separate source-code archive and require downloading the six source datasets under their original terms.

## Run order

1. Extract the ZIP to a local folder. Do not rename files inside it.
2. Open a terminal in the extracted package root.
3. Before generating any new outputs, run `python 07_Analysis_Code/Validate_Package_Integrity.py`. Expected final line: `PASS: package contents, sizes, hashes, strict JSON, and claim-file verification passed.`
4. Run `python 07_Analysis_Code/Check_33_Ledger_Indexed_Claims.py`. Expected final line: `PASS: 33/33 ledger-indexed claims matched their referenced result files.` The report is written to `08_Generated_Outputs/Ledger_Indexed_Claim_Check.json`.
5. Run `python 07_Analysis_Code/Independent_Derived_Results_Audit.py`. This separately implemented checker recomputes the wider set of arithmetic relations documented in its JSON output; "independent" here refers to software implementation, not an external audit or an independent research team.
6. Optional limited check in R: set the working directory to the archive root and run `source("07_Analysis_Code/Check_Core_Means_R.R")`. R covers the projection-breadth means and four continuous-speech study-equal endpoints only; it does not implement Results 3–5.
7. Optional limited check in MATLAB: set Current Folder to the archive root and run `run('07_Analysis_Code/Check_Core_Means_MATLAB.m')`. The script resolves the archive root from its own file location and therefore remains valid when MATLAB temporarily changes the working directory while executing `run`. MATLAB covers the projection-breadth means only; it does not implement Results 2–5.
8. To render all six article figures from the included final artifacts, run `python 07_Analysis_Code/Generate_Article_Figures.py`. Figure 5 is drawn from the participant-level geometry rows and the fixed-bin maximum-t results in `04_Final_Results/Individual_Event_Geometry_Transport_Results.json`; no completed figure is copied into the output directory. Under the pinned environment, all six regenerated PNGs are expected to match the packaged PNGs byte-for-byte. PDF/SVG files are also rendered, but their byte hashes can contain backend metadata and are not the byte-identity criterion.

The integrity validator intentionally rejects files not listed in the manifest. Run it on the pristine extraction before R, MATLAB, or figure scripts create local output files.

## File map

- `01_Result_Map.csv`: manuscript/result/code crosswalk.
- `02_Dataset_Citations_and_Licenses.csv`: source locations, required or recommended citations, and source licenses.
- `03_Derived_Data`: deidentified participant- or direction-level tables.
- `04_Final_Results`: final JSON sources for the 33 ledger-indexed claims.
- `05_Audit_Results`: public audit summaries and final analysis protocols.
- `06_Current_Figures`: publication figure files (PNG, PDF, SVG).
- `07_Analysis_Code`: complete Python verification and plotting code, plus explicitly limited R and MATLAB mean checks.
- `08_Generated_Outputs`: deterministic baseline outputs of the claim check, independent arithmetic audit, and package audit.
- `09_Claim_Evidence_Ledger.json`: 33 claim-to-source bindings.
- `10_File_Manifest.json`: byte size and SHA-256 for every member except the JSON manifest itself.
- `11_Licenses`: separate notices for original code, original documentation, and derived data.

## Licensing boundary

The file-specific license structure is intentional and is separate from the distribution license selected for the manuscript or preprint: original analysis code is MIT licensed, and original explanatory documentation is CC BY 4.0. Derived data are not blanket-relicensed; each artifact remains subject to the applicable license and attribution requirements of its source dataset, including noncommercial or share-alike terms where applicable. See `11_Licenses` and the dataset inventory.

## Dataset-identifier clarification

The Kojima dataset used by these analyses is the four-class ASME BCI dataset at DOI `10.7910/DVN/1UJDV6`. The identifier and recommended citation are recorded in the dataset inventory.
