## Supplementary material for "Falsifiable substitution tests reveal task-structured neural evidence for auditory attention": Derived_Result_Verification_Archive: Protocol_1_Projection_Breadth.md

# Protocol 1: projection breadth

The discovery rule averaged the first four held-window CCA margins. It was selected in odd-ID KUL participants and applied without retuning to even-ID KUL, DTU, and Ear-SAAD. All PCA, lag, covariance, and CCA transformations were estimated inside the outer training fold. The participant is the inferential unit. The three evaluation studies are summarized separately and with a study-equal mean.
