## Supplementary material for "Falsifiable substitution tests reveal task-structured neural evidence for auditory attention": Derived_Result_Verification_Archive: Protocol_2_High_Density_Direction_Delay.md

# Protocol 2: high-density direction-delay structure

A development cohort fixed 16 cells formed by eight scalp sectors and two response-delay ranges. Confirmation used participant-disjoint complete cases. The physical mapping was compared with 15 remappings that preserved cell count, fitted quantities, and the fold-specific weight distribution. Sensor-density analyses retrained each model independently.
