## Supplementary material for "Falsifiable substitution tests reveal task-structured neural evidence for auditory attention": Derived_Result_Verification_Archive: Protocol_3_Continuous_Cross_Listener_Transport.md

# Protocol 3: continuous-speech transport

KITE-ST, KUL, and Ear-SAAD were split into nonoverlapping source and target material. Source data estimated 16 reliability weights. Target data evaluated matched-listener weights against other-listener, leave-one-listener pooled, and capacity-matched remapped alternatives while holding receiver data and decoder quantities fixed. Each participant contributed both directed transfers.
