## Supplementary material for "Falsifiable substitution tests reveal task-structured neural evidence for auditory attention": Derived_Result_Verification_Archive: Protocol_4_Event_Hierarchy_and_Target_Exclusion.md

# Protocol 4: event hierarchy and target-event exclusion

The flat decoder chose a child command. A separate parent-stream score pooled sibling-versus-unattended evidence. The mechanism check used the known target label to remove every target-command event before pooling, so it is not an online confidence algorithm. Physical command-stream binding was compared with capacity-matched wrong bindings and with an identically pooled EOG-only comparator. The latter does not imply ocular activity was residualized from EEG.
