## Supplementary material for "Falsifiable substitution tests reveal task-structured neural evidence for auditory attention": Derived_Result_Verification_Archive: Protocol_5_Individual_Event_Geometry_and_Command_Residual.md

# Protocol 5: individual event geometry and command residualization

Sensor-by-time directions were trained and tested across disjoint phases or runs with target-command events excluded. Same-listener transport was compared with other-listener and wrong-mapping alternatives. In the command-identity control, command means were estimated from non-target training events only and removed from both training and held-out features before refitting.
