## Supplementary material for "Falsifiable substitution tests reveal task-structured neural evidence for auditory attention": Derived_Result_Verification_Archive: Protocol_6_Wearable_Channels_and_Calibration.md

# Protocol 6: wearable channels and calibration

Models for 8, 16, 32, and 64 electrodes were retrained for each montage. Calibration subsets were enumerated within participant and equal-sized subsets were averaged before group inference. The joint criterion required physical binding, an EEG-over-EOG-only contrast, listener transport, minimum binding retention, and three-run physical and binding effects. The eight-channel ASME30 listener-consistency criterion failed.
