## Supplementary material for "Falsifiable substitution tests reveal task-structured neural evidence for auditory attention": Derived_Result_Verification_Archive: Figure2_Continuous_Speech_Direction_Delay.pdf

### Continuous-speech information extends beyond the leading projection and depends on direction–delay structure

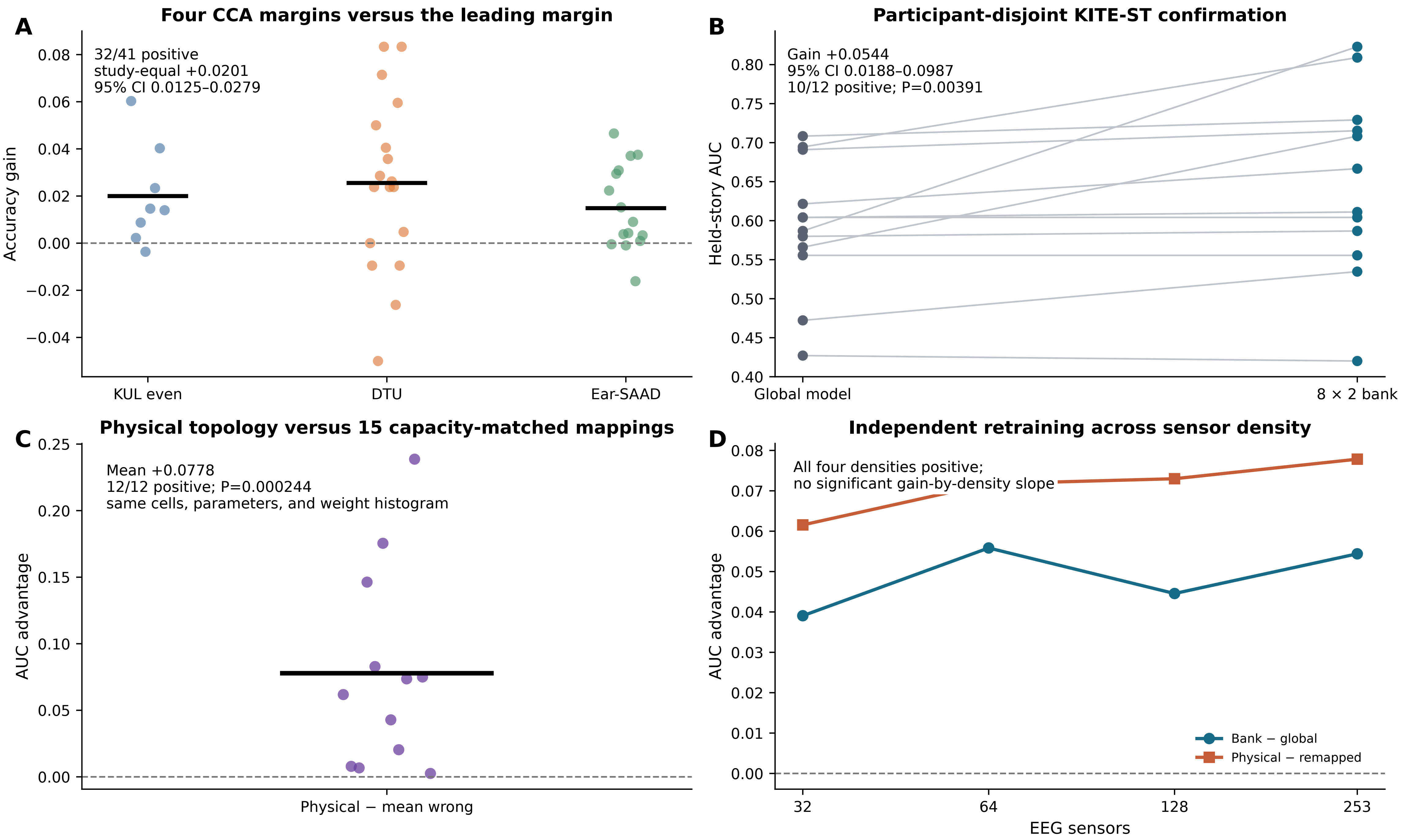
