## Supplementary material for "Falsifiable substitution tests reveal task-structured neural evidence for auditory attention": Derived_Result_Verification_Archive: Figure3_Continuous_Speech_Transport.pdf

### Structure-dependent held-speech evidence across nonoverlapping continuous speech

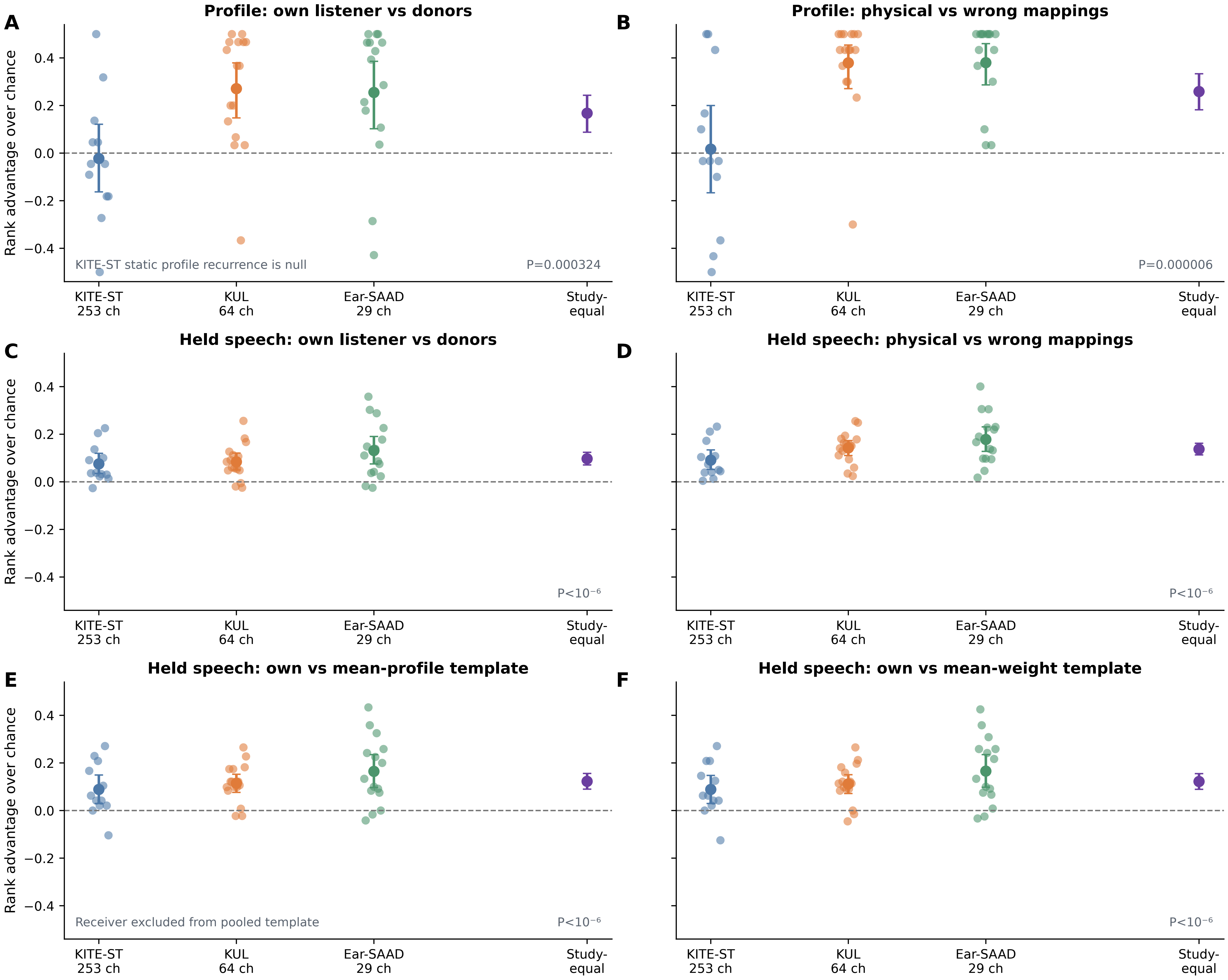


Rank advantage is the probability of winning a matched comparison minus 0.5; it is not an accuracy gain.
