## Supplementary material for "Falsifiable substitution tests reveal task-structured neural evidence for auditory attention": Derived_Result_Verification_Archive: Figure5_Event_Geometry_Transport.pdf

### Task-defined event directions transport within listeners across phases or runs

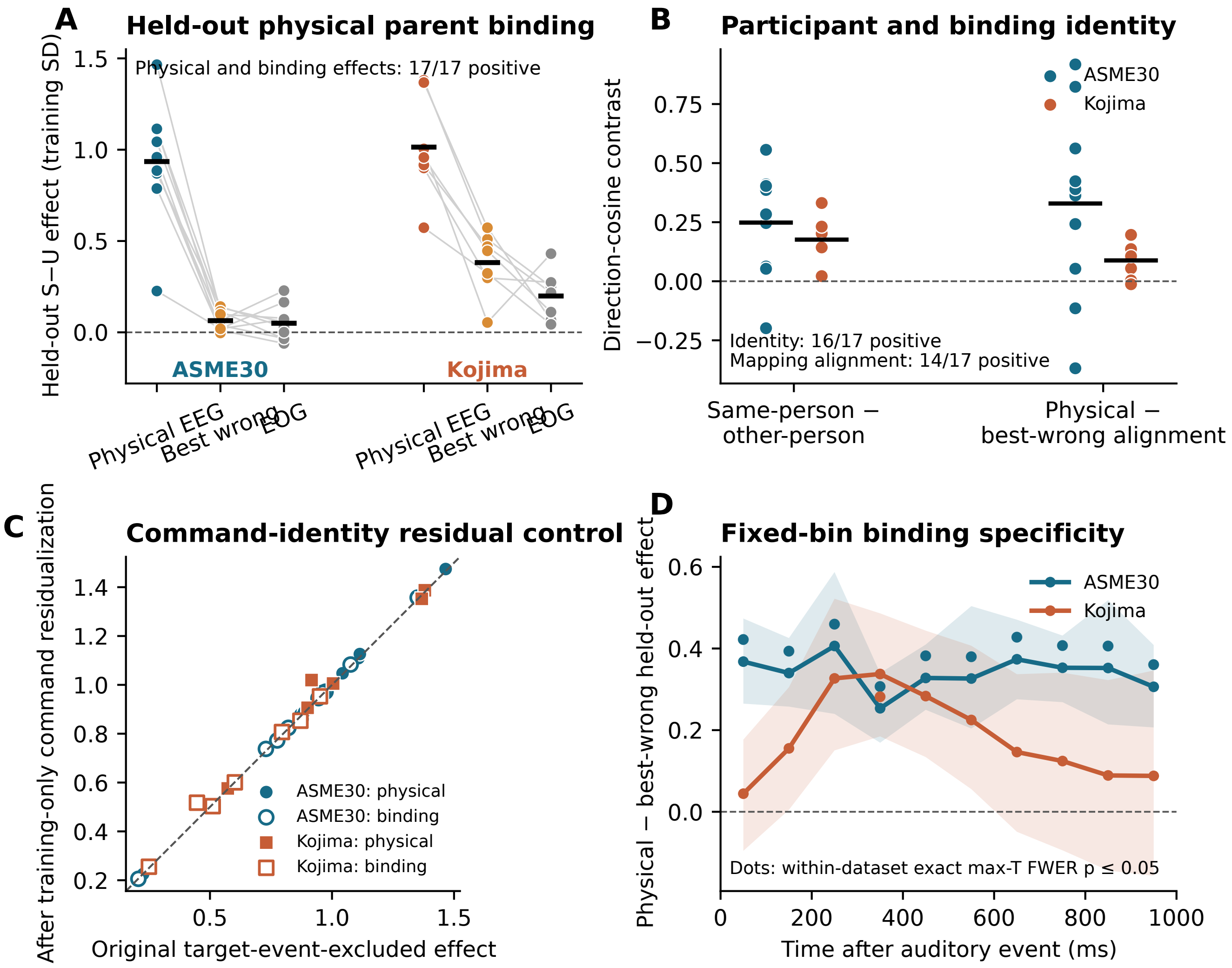


Known target-command events excluded with oracle labels; models evaluated on held-out phases or runs. No cross-dataset model training; inference remains dataset specific.
