## Supplementary material for "Falsifiable substitution tests reveal task-structured neural evidence for auditory attention": Derived_Result_Verification_Archive: Check_33_Ledger_Indexed_Claims.py

```
#!/usr/bin/env python3
"""Check 33 ledger-indexed claims and recompute core participant summaries."""
from __future__ import annotations
import csv, hashlib, json, math
from pathlib import Path

ROOT = Path(__file__).resolve().parents[1]

def strict_load(path):
    def reject(value): raise ValueError(f"Non-standard JSON constant {value} in {path}")
    return json.loads(path.read_text(encoding="utf-8"), parse_constant=reject)

def digest(path):
    h=hashlib.sha256()
    with path.open("rb") as stream:
        for block in iter(lambda:stream.read(1024*1024),b""): h.update(block)
    return h.hexdigest()

def get(value, path):
    for part in path.split("."): value=value[int(part)] if isinstance(value,list) else value[part]
    return value

def equal(a,b):
    if isinstance(a,bool) or isinstance(b,bool): return a is b
    if isinstance(a,(int,float)) and isinstance(b,(int,float)): return math.isclose(float(a),float(b),rel_tol=0,abs_tol=1e-12)
    if isinstance(a,list) and isinstance(b,list) and len(a)==len(b): return all(equal(x,y) for x,y in zip(a,b))
    if isinstance(a,dict) and isinstance(b,dict) and set(a)==set(b): return all(equal(a[k],b[k]) for k in a)
    return a==b

def table(name):
    with (ROOT/"03_Derived_Data"/name).open(encoding="utf-8-sig",newline="") as stream: return list(csv.DictReader(stream))

def mean(rows, field): return sum(float(row[field]) for row in rows)/len(rows)

ledger=strict_load(ROOT/"09_Claim_Evidence_Ledger.json")
records=[]
for row in ledger["claims"]:
    source=ROOT/row["evidence_file"]
    exists=source.is_file()
    hash_ok=exists and digest(source)==row["evidence_file_sha256"]
    value_ok=False
    if exists:
        value_ok=equal(get(strict_load(source),row["value_path"]),row["value"])
    records.append({"claim_id":row["claim_id"],"source_exists":exists,"hash_matches":hash_ok,"value_matches":value_ok})

projection=[
 ("KUL",table("Projection_Breadth_KUL_Participants.csv"),"equal_k4_gain_vs_dominant_k1"),
 ("DTU",table("Projection_Breadth_DTU_Participants.csv"),"equal_k4_gain_vs_dominant_k1"),
 ("Ear-SAAD",table("Projection_Breadth_EarSAAD_Participants.csv"),"gain_equal_minus_k1")]
study_means={name:mean(rows,field) for name,rows,field in projection}
values=[float(row[field]) for _,rows,field in projection for row in rows]
continuous=table("Continuous_Speech_Transport_Participant_Endpoints.csv")
datasets=sorted({row["dataset"] for row in continuous})
fields=["F1_fingerprint_listener_rank_advantage","F2_fingerprint_physical_mapping_rank_advantage","P1_predictive_listener_rank_advantage","P2_predictive_physical_mapping_rank_advantage"]
continuous_equal={field:sum(mean([r for r in continuous if r["dataset"]==dataset],field) for dataset in datasets)/len(datasets) for field in fields}
passed=all(r["source_exists"] and r["hash_matches"] and r["value_matches"] for r in records) and len(records)==33
output={"status":"PASS" if passed else "FAIL","ledger_indexed_claims":len(records),"claims_verified":sum(r["source_exists"] and r["hash_matches"] and r["value_matches"] for r in records),"claim_checks":records,"recomputed_core_summaries":{"projection_breadth":{"participants":len(values),"positive_participants":sum(v>0 for v in values),"study_means":study_means,"study_equal_mean":sum(study_means.values())/3},"continuous_speech_study_equal_means":continuous_equal}}
out=ROOT/"08_Generated_Outputs"/"Ledger_Indexed_Claim_Check.json"
out.parent.mkdir(exist_ok=True)
out.write_text(json.dumps(output,indent=2,allow_nan=False)+"\n",encoding="utf-8")
print(f"{output['status']}: {output['claims_verified']}/{output['ledger_indexed_claims']} ledger-indexed claims matched their referenced result files.")
if not passed: raise SystemExit(1)
```
