## Supplementary material for "Falsifiable substitution tests reveal task-structured neural evidence for auditory attention": Derived_Result_Verification_Archive: Check_Core_Means_MATLAB.m

```
% Independent recomputation of core participant-level means.
script_path = mfilename('fullpath');
if isempty(script_path)
    error('Unable to resolve the MATLAB verification script location.');
end
root = fileparts(fileparts(script_path));
data_dir = fullfile(root,'03_Derived_Data');
kul = readtable(fullfile(data_dir,'Projection_Breadth_KUL_Participants.csv'));
dtu = readtable(fullfile(data_dir,'Projection_Breadth_DTU_Participants.csv'));
ear = readtable(fullfile(data_dir,'Projection_Breadth_EarSAAD_Participants.csv'));
names = {'KUL';'DTU';'Ear-SAAD'};
values = [mean(kul.equal_k4_gain_vs_dominant_k1); mean(dtu.equal_k4_gain_vs_dominant_k1); mean(ear.gain_equal_minus_k1)];
disp(table(names,values));
fprintf('Study-equal projection-breadth mean: %.12f\n',mean(values));
out = fullfile(root,'08_Generated_Outputs','Core_Means_Check_MATLAB.txt');
fid=fopen(out,'w');
fprintf(fid,'KUL %.12f\nDTU %.12f\nEar-SAAD %.12f\nStudy-equal %.12f\n',values(1),values(2),values(3),mean(values));
fclose(fid);
```
