## Supplementary material for "Falsifiable substitution tests reveal task-structured neural evidence for auditory attention": Derived_Result_Verification_Archive: Check_Core_Means_R.R

```
# Independent recomputation of core participant-level means using base R.
root <- normalizePath(getwd(), winslash="/", mustWork=TRUE)
read_table <- function(name) read.csv(file.path(root,"03_Derived_Data",name),check.names=FALSE)
mean_field <- function(tab,field) mean(as.numeric(tab[[field]]))
projection <- c(
 KUL=mean_field(read_table("Projection_Breadth_KUL_Participants.csv"),"equal_k4_gain_vs_dominant_k1"),
 DTU=mean_field(read_table("Projection_Breadth_DTU_Participants.csv"),"equal_k4_gain_vs_dominant_k1"),
 Ear_SAAD=mean_field(read_table("Projection_Breadth_EarSAAD_Participants.csv"),"gain_equal_minus_k1"))
continuous <- read_table("Continuous_Speech_Transport_Participant_Endpoints.csv")
fields <- c("F1_fingerprint_listener_rank_advantage","F2_fingerprint_physical_mapping_rank_advantage","P1_predictive_listener_rank_advantage","P2_predictive_physical_mapping_rank_advantage")
study_equal <- sapply(fields,function(field) mean(tapply(as.numeric(continuous[[field]]),continuous$dataset,mean)))
result <- data.frame(metric=c(names(projection),"projection_study_equal",names(study_equal)),value=c(projection,mean(projection),study_equal))
print(result,row.names=FALSE,digits=12)
write.csv(result,file.path(root,"08_Generated_Outputs","Core_Means_Check_R.csv"),row.names=FALSE)
```
