## Supplementary material for "Falsifiable substitution tests reveal task-structured neural evidence for auditory attention": Derived_Result_Verification_Archive: Independent_Derived_Results_Audit.py

```
"""Independent arithmetic audit of the supplied *derived* participant tables.

This intentionally does not import the package's own checking code.  It verifies
what can be recomputed from Supplementary Data 1, while recording the important
boundary that raw-EEG feature extraction and model fitting are not present.
"""

from __future__ import annotations

import csv
import itertools
import json
import math
from pathlib import Path
from statistics import fmean


ROOT = Path(__file__).resolve().parents[1]
DERIVED = ROOT / "03_Derived_Data"
FINAL = ROOT / "04_Final_Results"
OUT = ROOT / "08_Generated_Outputs" / "Independent_Derived_Results_Audit.json"
TOL = 1e-11


def read_csv(name: str) -> list[dict[str, str]]:
    with (DERIVED / name).open(encoding="utf-8-sig", newline="") as handle:
        return list(csv.DictReader(handle))


def read_json(folder: Path, name: str):
    return json.loads((folder / name).read_text(encoding="utf-8"))


def number(value: str) -> float:
    return float(value)


def close(left: float, right: float, tol: float = TOL) -> bool:
    return math.isclose(float(left), float(right), rel_tol=tol, abs_tol=tol)


checks: list[dict] = []


def record(name: str, passed: bool, **detail) -> None:
    checks.append({"name": name, "passed": bool(passed), **detail})


def exact_signflip_p(values: list[float], null: float = 0.0) -> float | None:
    """Enumerate the exact one-sided randomization distribution for small n."""
    centered = [float(v) - null for v in values]
    n = len(centered)
    if n > 22:
        return None
    observed = sum(centered)
    extreme = 0
    for signs in itertools.product((-1.0, 1.0), repeat=n):
        if sum(sign * value for sign, value in zip(signs, centered)) >= observed - 1e-14:
            extreme += 1
    return extreme / (2**n)


def audit_summary_nodes(document_name: str, obj, path: str = "") -> None:
    if isinstance(obj, dict):
        if {"participant_values", "mean", "n_positive", "n_total"} <= set(obj):
            values = [float(value) for value in obj["participant_values"]]
            reported_positive = int(obj["n_positive"])
            possible_nulls = [
                null for null in (0.0, 0.5)
                if sum(value > null for value in values) == reported_positive
            ]
            null = possible_nulls[0] if possible_nulls else None
            record(
                f"{document_name}:{path}:mean_count",
                len(values) == int(obj["n_total"])
                and close(fmean(values), obj["mean"])
                and null is not None,
                n=len(values),
                recomputed_mean=fmean(values),
                reported_mean=obj["mean"],
                inferred_null=null,
            )
            if "one_sided_exact_signflip_p" in obj and null is not None:
                recomputed_p = exact_signflip_p(values, null)
                if recomputed_p is not None:
                    record(
                        f"{document_name}:{path}:exact_signflip",
                        close(recomputed_p, obj["one_sided_exact_signflip_p"]),
                        recomputed_p=recomputed_p,
                        reported_p=obj["one_sided_exact_signflip_p"],
                        n=len(values),
                    )
        for key, value in obj.items():
            audit_summary_nodes(document_name, value, f"{path}.{key}" if path else key)
    elif isinstance(obj, list):
        for index, value in enumerate(obj):
            audit_summary_nodes(document_name, value, f"{path}[{index}]")


# 1. Recompute every final-result summary that exposes participant values.
for final_file in sorted(FINAL.glob("*.json")):
    audit_summary_nodes(final_file.name, read_json(FINAL, final_file.name))


# 2. Projection breadth: participant tables -> three study rows -> synthesis.
projection_specs = [
    ("Projection_Breadth_KUL_Participants.csv", "equal_k4_gain_vs_dominant_k1", "KUL"),
    ("Projection_Breadth_DTU_Participants.csv", "equal_k4_gain_vs_dominant_k1", "DTU"),
    ("Projection_Breadth_EarSAAD_Participants.csv", "gain_equal_minus_k1", "Ear-SAAD"),
]
projection = read_json(FINAL, "Projection_Breadth_Synthesis_Results.json")
study_means = []
participant_values = []
for csv_name, column, dataset in projection_specs:
    values = [number(row[column]) for row in read_csv(csv_name)]
    participant_values.extend(values)
    study_means.append(fmean(values))
    target = next(item for item in projection["studies"] if item["dataset"] == dataset)
    record(
        f"projection:{dataset}:participant_table_to_final",
        close(fmean(values), target["equal_k4_minus_k1"])
        and sum(value > 0 for value in values) == target["participants_improved"]
        and len(values) == target["participants"],
        recomputed_mean=fmean(values),
        reported_mean=target["equal_k4_minus_k1"],
    )
record(
    "projection:study_equal_synthesis",
    close(fmean(study_means), projection["independent_analysis_synthesis"]["study_equal_mean_gain"]),
    recomputed=fmean(study_means),
    reported=projection["independent_analysis_synthesis"]["study_equal_mean_gain"],
)
record(
    "projection:participant_weighted_synthesis",
    close(fmean(participant_values), projection["independent_analysis_synthesis"]["participant_weighted_mean_gain"]),
    recomputed=fmean(participant_values),
    reported=projection["independent_analysis_synthesis"]["participant_weighted_mean_gain"],
)


# 3. Direction rows -> participant rows for both continuous-speech analyses.
continuous_specs = [
    (
        "Continuous_Speech_Transport_Direction_Endpoints.csv",
        "Continuous_Speech_Transport_Participant_Endpoints.csv",
        [
            ("fingerprint_listener_rank_probability", "F1_fingerprint_listener_rank_advantage", -0.5),
            ("fingerprint_physical_mapping_rank_probability", "F2_fingerprint_physical_mapping_rank_advantage", -0.5),
            ("predictive_listener_rank_probability", "P1_predictive_listener_rank_advantage", -0.5),
            ("predictive_physical_mapping_rank_probability", "P2_predictive_physical_mapping_rank_advantage", -0.5),
            ("normalized_own_minus_donor_score", "normalized_own_minus_donor_score", 0.0),
            ("normalized_physical_minus_wrong_score", "normalized_physical_minus_wrong_score", 0.0),
            ("global_auc", "global_auc", 0.0),
            ("own_profile_auc", "own_profile_auc", 0.0),
        ],
    ),
    (
        "Continuous_Speech_Shared_Template_Direction_Endpoints.csv",
        "Continuous_Speech_Shared_Template_Participant_Endpoints.csv",
        [
            ("own_vs_mean_profile_rank_probability", "G1_own_vs_mean_profile_rank_advantage", -0.5),
            ("own_vs_mean_weight_rank_probability", "G2_own_vs_mean_weight_rank_advantage", -0.5),
            ("own_minus_mean_profile_auc", "own_minus_mean_profile_auc", 0.0),
            ("own_minus_mean_weight_auc", "own_minus_mean_weight_auc", 0.0),
            ("normalized_own_minus_mean_profile_score", "normalized_own_minus_mean_profile_score", 0.0),
            ("normalized_own_minus_mean_weight_score", "normalized_own_minus_mean_weight_score", 0.0),
        ],
    ),
]
for direction_file, participant_file, columns in continuous_specs:
    direction_rows = read_csv(direction_file)
    participant_rows = read_csv(participant_file)
    by_key: dict[tuple[str, str], list[dict[str, str]]] = {}
    for row in direction_rows:
        by_key.setdefault((row["dataset"], row["subject"]), []).append(row)
    failures = []
    for row in participant_rows:
        pair = by_key[(row["dataset"], row["subject"])]
        if len(pair) != 2:
            failures.append({"key": [row["dataset"], row["subject"]], "reason": "direction_count"})
            continue
        for source_column, target_column, adjustment in columns:
            expected = fmean(number(item[source_column]) for item in pair) + adjustment
            if not close(expected, number(row[target_column])):
                failures.append(
                    {"key": [row["dataset"], row["subject"]], "column": target_column,
                     "expected": expected, "reported": number(row[target_column])}
                )
    record(
        f"continuous:{direction_file}:direction_to_participant",
        not failures and len(by_key) == len(participant_rows),
        participant_rows=len(participant_rows),
        direction_rows=len(direction_rows),
        failures=failures[:10],
    )


# 4. Participant CSVs -> final summary participant vectors for named analyses.
def compare_summary_vectors(
    check_prefix: str,
    rows: list[dict[str, str]],
    final_doc: dict,
    mappings: dict[str, str],
    summary_root: str = "summaries",
) -> None:
    for dataset in sorted({row["dataset"] for row in rows}):
        subset = [row for row in rows if row["dataset"] == dataset]
        for summary_name, csv_column in mappings.items():
            expected = sorted(number(row[csv_column]) for row in subset)
            actual = sorted(float(value) for value in final_doc[summary_root][dataset][summary_name]["participant_values"])
            record(
                f"{check_prefix}:{dataset}:{summary_name}:csv_to_final_vector",
                len(expected) == len(actual) and all(close(a, b) for a, b in zip(expected, actual)),
                n=len(expected),
            )


residual_rows = read_csv("Command_Identity_Residual_Participant_Results.csv")
residual_final = read_json(FINAL, "Command_Identity_Residual_Control_Results.json")
compare_summary_vectors(
    "command_residual",
    residual_rows,
    residual_final,
    {
        "physical": "physical_effect",
        "binding": "binding_specificity",
        "identity": "identity_advantage",
        "mapping_alignment": "mapping_alignment_specificity",
        "physical_retention_ratio": "residual_to_raw_physical_ratio",
        "binding_retention_ratio": "residual_to_raw_binding_ratio",
    },
)

target_rows = [row for row in read_csv("Target_Event_Exclusion_Participant_Results.csv") if row["evaluable"] == "True"]
target_final = read_json(FINAL, "Target_Event_Exclusion_and_EOG_Comparator_Results.json")
compare_summary_vectors(
    "target_exclusion",
    target_rows,
    target_final,
    {
        "target_event_excluded_absolute": "target_event_excluded_eeg_auc",
        "mapping_specificity": "target_event_excluded_minus_best_wrong_auc",
        "eeg_minus_eog": "target_event_excluded_minus_eog_auc",
        "target_event_excluded_minus_inclusive": "target_event_excluded_minus_inclusive_auc",
    },
)


# 5. Wearable participant table -> channel-count aggregate table and final vectors.
wear_rows = read_csv("Wearable_Channels_Participant_Metrics.csv")
wear_curve = read_csv("Wearable_Channels_Channel_Count_Curve.csv")
curve_failures = []
for summary_row in wear_curve:
    subset = [
        row for row in wear_rows
        if row["dataset"] == summary_row["dataset"]
        and row["montage_channels"] == summary_row["montage_channels"]
    ]
    values = [number(row[summary_row["metric"]]) for row in subset]
    if not (
        close(fmean(values), number(summary_row["mean"]))
        and len(values) == int(summary_row["n_total"])
        and sum(value > 0 for value in values) == int(summary_row["n_positive"])
    ):
        curve_failures.append({"dataset": summary_row["dataset"], "montage": summary_row["montage_channels"], "metric": summary_row["metric"]})
record(
    "wearable:participant_table_to_channel_count_curve",
    not curve_failures,
    aggregate_rows=len(wear_curve),
    failures=curve_failures,
)

wear_final = read_json(FINAL, "Wearable_Channels_and_Calibration_Results.json")
vector_failures = []
for dataset, montage_map in wear_final["montage_summaries"].items():
    for montage, metric_map in montage_map.items():
        subset = [
            row for row in wear_rows
            if row["dataset"] == dataset and row["montage_channels"] == str(montage)
        ]
        for metric, summary in metric_map.items():
            expected = sorted(number(row[metric]) for row in subset)
            actual = sorted(float(value) for value in summary["participant_values"])
            if len(expected) != len(actual) or any(not close(a, b) for a, b in zip(expected, actual)):
                vector_failures.append({"dataset": dataset, "montage": montage, "metric": metric})
record(
    "wearable:participant_table_to_final_vectors",
    not vector_failures,
    vectors_checked=sum(len(metrics) for montages in wear_final["montage_summaries"].values() for metrics in montages.values()),
    failures=vector_failures,
)


# 6. High-density participant JSON -> its own metrics and final ledger-indexed values.
hd_confirm = read_json(DERIVED, "High_Density_Direction_Delay_Confirmation_Participants.json")
hd_topology = read_json(DERIVED, "High_Density_Direction_Delay_Topology_Control_Participants.json")
hd_final = read_json(FINAL, "High_Density_Direction_Delay_Structure_Results.json")
confirm_values = [float(row["intact_gain_auc"]) for row in hd_confirm["participant_results"]]
topology_values = [float(row["intact_identity_advantage_auc"]) for row in hd_topology["participant_results"]]
record(
    "high_density:confirmation_participants_to_metrics",
    close(fmean(confirm_values), hd_confirm["metrics"]["mean_intact_gain_auc"])
    and sum(value > 0 for value in confirm_values) == hd_confirm["metrics"]["participants_positive_intact_gain"],
    n=len(confirm_values),
)
record(
    "high_density:topology_participants_to_metrics",
    close(fmean(topology_values), hd_topology["metrics"]["mean_intact_identity_advantage_auc"])
    and sum(value > 0 for value in topology_values) == hd_topology["metrics"]["participants_positive_intact_identity_advantage"],
    n=len(topology_values),
)
ledger_values = hd_final["ledger_indexed_claim_values"]
record(
    "high_density:derived_metrics_to_ledger_indexed_claims",
    close(hd_confirm["metrics"]["mean_intact_gain_auc"], ledger_values["KITE-CONF"]["value"])
    and close(hd_topology["metrics"]["mean_intact_identity_advantage_auc"], ledger_values["KITE-TOPO"]["value"]),
)


# 7. Explicitly record unavailable replication layers.
limitations = {
    "raw_eeg_to_preprocessed": "not present in this derived-result verification archive; use the separate source-code archive",
    "preprocessed_to_participant_endpoints": "not present in this derived-result verification archive; use the separate source-code archive",
    "model_fitting_and_upstream_feature_extraction": "not present in this derived-result verification archive; use the separate source-code archive",
    "bootstrap_ci_reproduction": "participant values are present for many endpoints, but endpoint-specific seeds/code are generally absent",
    "calibration_participant_rows": "aggregate curve and final participant vectors are present, but no calibration participant table is supplied",
}

result = {
    "scope": "arithmetic and cross-file audit from supplied derived participant-level files; not raw-data reproduction",
    "package_root": ".",
    "checks_total": len(checks),
    "checks_passed": sum(item["passed"] for item in checks),
    "checks_failed": sum(not item["passed"] for item in checks),
    "checks": checks,
    "unavailable_layers": limitations,
}
OUT.write_text(json.dumps(result, indent=2, ensure_ascii=False) + "\n", encoding="utf-8")
print(json.dumps({key: result[key] for key in ("checks_total", "checks_passed", "checks_failed")}, indent=2))
if result["checks_failed"]:
    for item in checks:
        if not item["passed"]:
            print("FAIL", item["name"], item)
    raise SystemExit(1)
```
