## Supplementary material for "Falsifiable substitution tests reveal task-structured neural evidence for auditory attention": Derived_Result_Verification_Archive: Validate_Package_Integrity.py

```
#!/usr/bin/env python3
"""Strictly validate package members, JSON, hashes, sizes, and claim-file references."""
from __future__ import annotations
import hashlib, json, math, re
from pathlib import Path

ROOT=Path(__file__).resolve().parents[1]
MANIFEST=ROOT/"10_File_Manifest.json"

def load(path):
    def reject(value): raise ValueError(f"Non-standard JSON constant {value} in {path}")
    return json.loads(path.read_text(encoding="utf-8"),parse_constant=reject)

def digest(path):
    h=hashlib.sha256()
    with path.open("rb") as stream:
        for block in iter(lambda:stream.read(1024*1024),b""): h.update(block)
    return h.hexdigest()

problems=[]
manifest=load(MANIFEST)
listed={row["file"] for row in manifest["files"]}
actual={p.relative_to(ROOT).as_posix() for p in ROOT.rglob("*") if p.is_file() and p!=MANIFEST}
if listed!=actual:
    for path in sorted(listed-actual): problems.append(f"missing manifest member: {path}")
    for path in sorted(actual-listed): problems.append(f"unlisted extra file: {path}")
for row in manifest["files"]:
    path=ROOT/row["file"]
    if not path.is_file(): continue
    if path.stat().st_size!=row["bytes"]: problems.append(f"size mismatch: {row['file']}")
    if digest(path)!=row["sha256"]: problems.append(f"hash mismatch: {row['file']}")
for path in sorted(ROOT.rglob("*.json")):
    try: load(path)
    except Exception as exc: problems.append(f"strict JSON failure: {path.relative_to(ROOT).as_posix()} ({exc})")

forbidden={"old package token":"preprint_"+"reproducibility_bundle","wrong DOI":"10.1002/"+"widm.1400","wrong Kojima dataset DOI":"10.7910/DVN/"+"MQOVEY","disabled draft":"DO_"+"NOT_USE","internal state":"project_"+"state","old target token":"target_"+"free","old ocular token":"non"+"ocular","old certificate token":"certifi"+"cate_","local generator metadata":"generated by python"+"-docx"}
text_ext={".md",".txt",".csv",".json",".py",".r",".m",".svg"}
local_path_pattern=r"(?i)(?<![A-Za-z])(?:[A-Z]:[\\/]|/"+r"home/|/"+r"Users/)"
for path in sorted(p for p in ROOT.rglob("*") if p.is_file()):
    rel=path.relative_to(ROOT).as_posix()
    if not rel.isascii(): problems.append(f"non-ASCII public path: {rel}")
    if path.suffix.lower() in {".zip",".7z",".rar",".tar",".gz"}: problems.append(f"nested archive: {rel}")
    if path.suffix.lower() not in text_ext: continue
    text=path.read_text(encoding="utf-8-sig",errors="replace")
    for label,token in forbidden.items():
        if token.lower() in text.lower(): problems.append(f"{label}: {rel}")
    if re.search(r"\b"+"regis"+r"tered\b",text,flags=re.IGNORECASE): problems.append(f"misleading claim-registration term: {rel}")
    if re.search(local_path_pattern,text): problems.append(f"absolute local path: {rel}")

ledger=load(ROOT/"09_Claim_Evidence_Ledger.json")
if ledger.get("ledger_indexed_claim_count")!=33 or len(ledger.get("claims",[]))!=33: problems.append("claim ledger does not contain exactly 33 ledger-indexed claims")
for row in ledger.get("claims",[]):
    path=ROOT/row["evidence_file"]
    if not path.is_file(): problems.append(f"ledger source missing: {row['claim_id']}")
    elif digest(path)!=row["evidence_file_sha256"]: problems.append(f"ledger source hash mismatch: {row['claim_id']}")
if problems:
    print("FAIL")
    for problem in problems: print("- "+problem)
    raise SystemExit(1)
print("PASS: package contents, sizes, hashes, strict JSON, and claim-file verification passed.")
```
