## Supplementary material for "Falsifiable substitution tests reveal task-structured neural evidence for auditory attention": Derived_Result_Verification_Archive: Derived_Data_License_Notice.txt

Derived tables and result artifacts are not blanket-relicensed. They remain subject to the applicable source-dataset license and attribution requirements listed in 02_Dataset_Citations_and_Licenses.csv, including CC BY-NC or CC BY-NC-SA terms where applicable. Source-license URLs are included in that inventory. No additional rights in source data are granted by this package, and the manuscript or preprint distribution setting does not supersede these source-license terms.
