## Supplementary material for "Falsifiable substitution tests reveal task-structured neural evidence for auditory attention": Derived_Result_Verification_Archive: LICENSE_Documentation_CC_BY_4.0.txt

Original explanatory documentation authored for this package is licensed under CC BY 4.0: https://creativecommons.org/licenses/by/4.0/. This intentional file-specific license is separate from the distribution license selected for the manuscript or preprint. It does not apply to derived data or third-party source material.
