## Supplementary figures and images for "Falsifiable substitution tests reveal task-structured neural evidence for auditory attention"

### Figure1_Evidence_Attribution_Framework.pdf

#### From a Kakeya proof strategy to falsifiable attribution of auditory-attention evidence

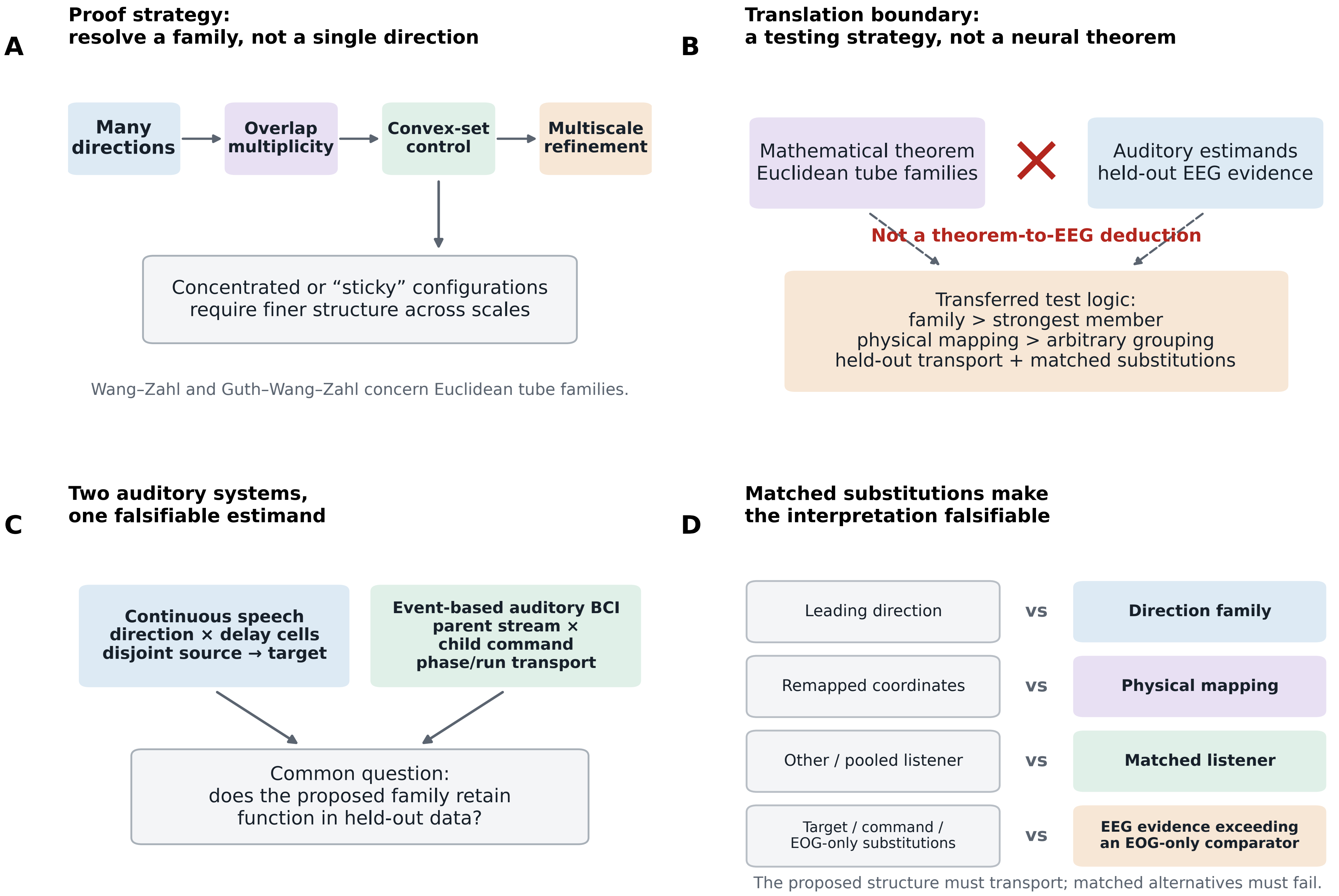

### Figure1_Evidence_Attribution_Framework.png

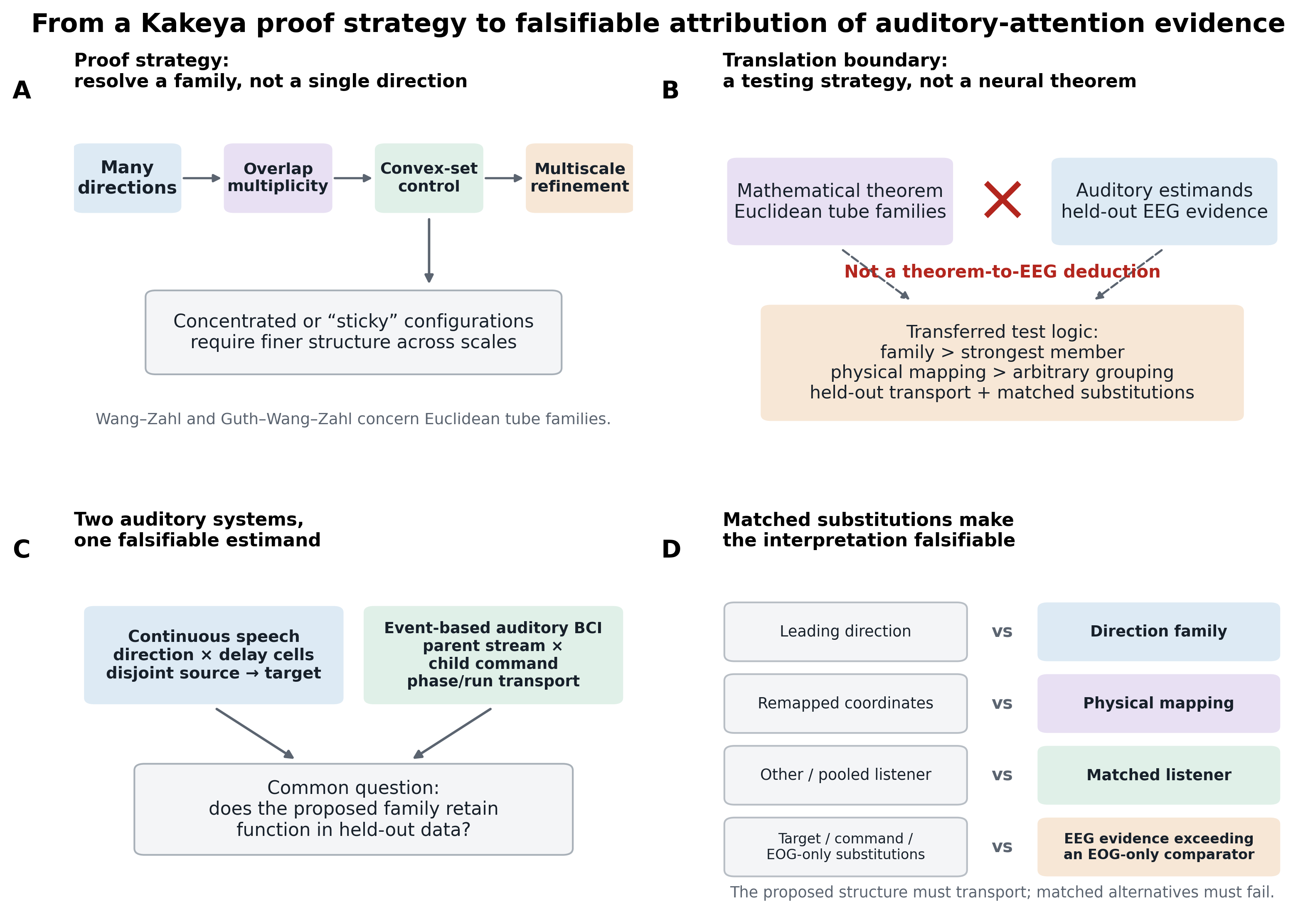

### Figure2_Continuous_Speech_Direction_Delay.png

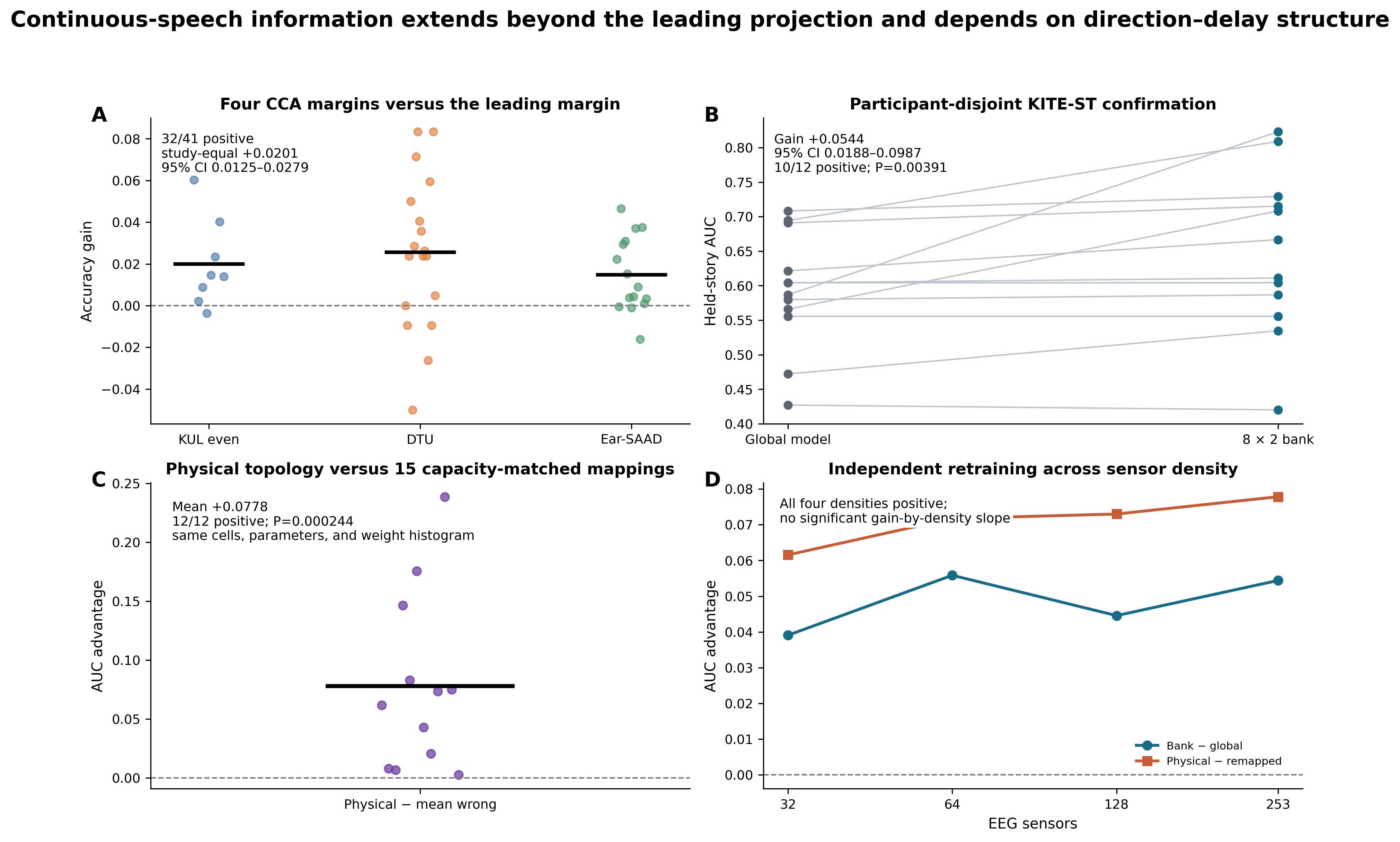

### Figure3_Continuous_Speech_Transport.png

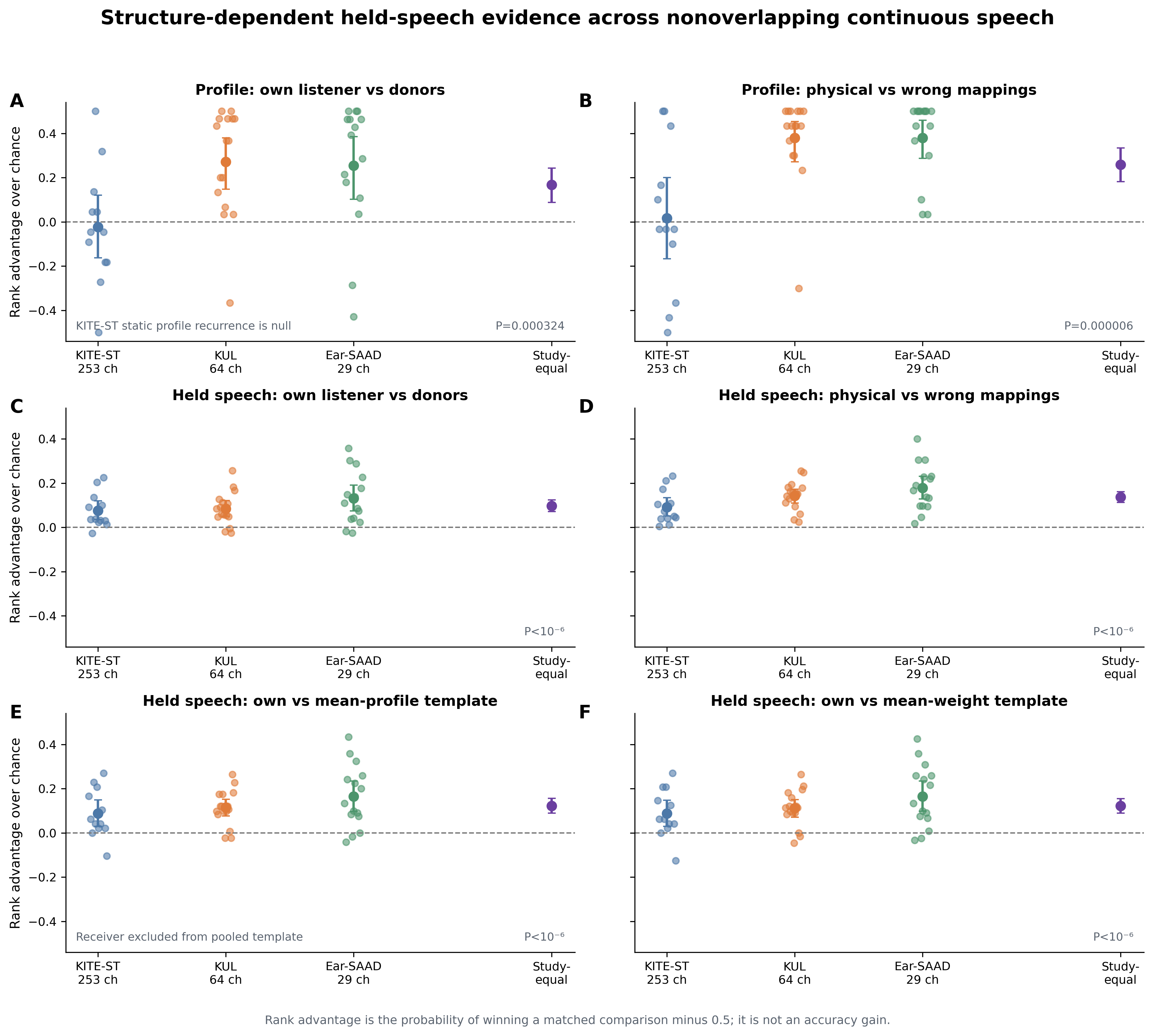

### Figure4_Target_Event_Excluded_Parent_Decoding.pdf

#### Non-target sibling events identify parent-stream errors hidden by a flat auditory BCI

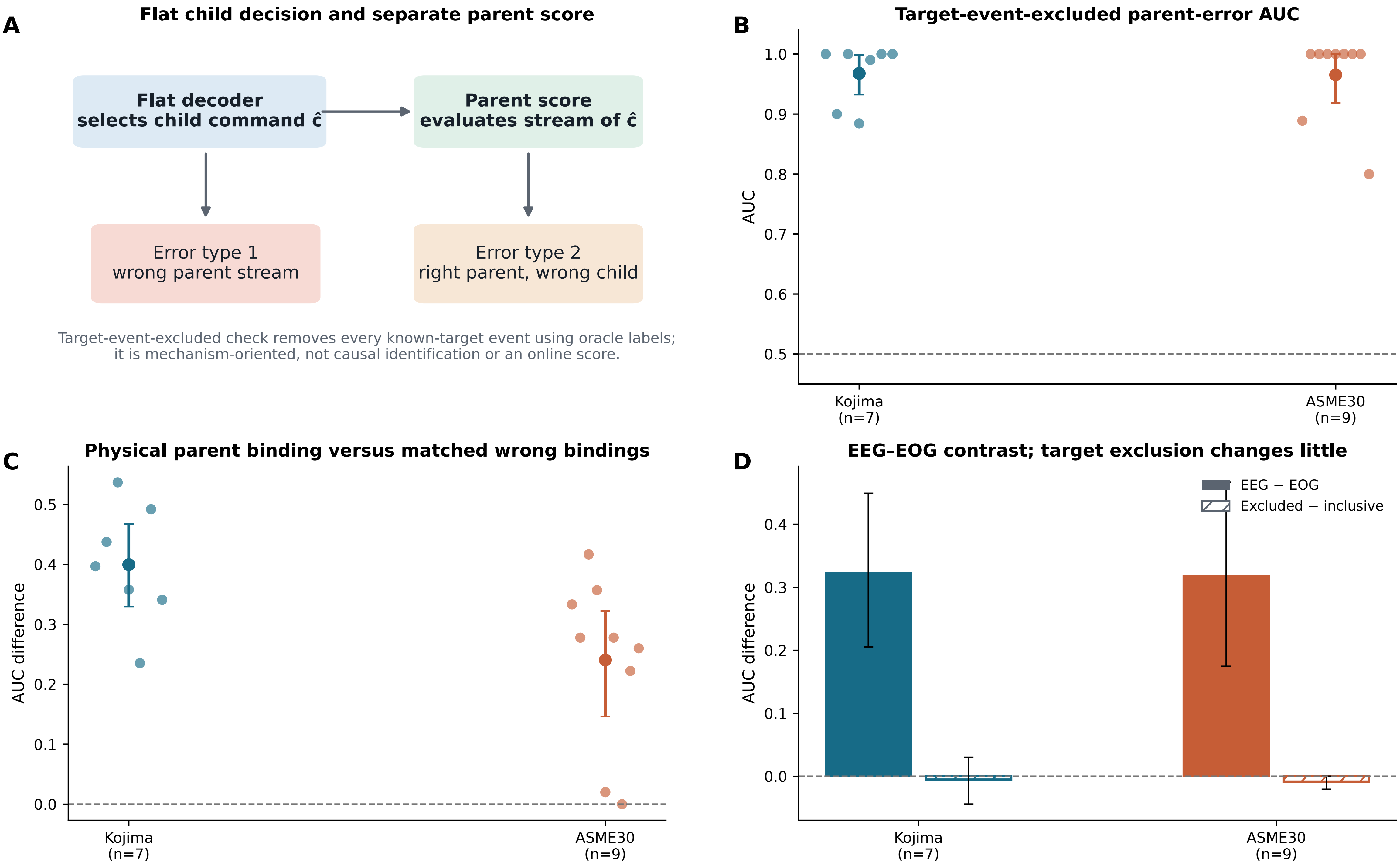

### Figure4_Target_Event_Excluded_Parent_Decoding.png

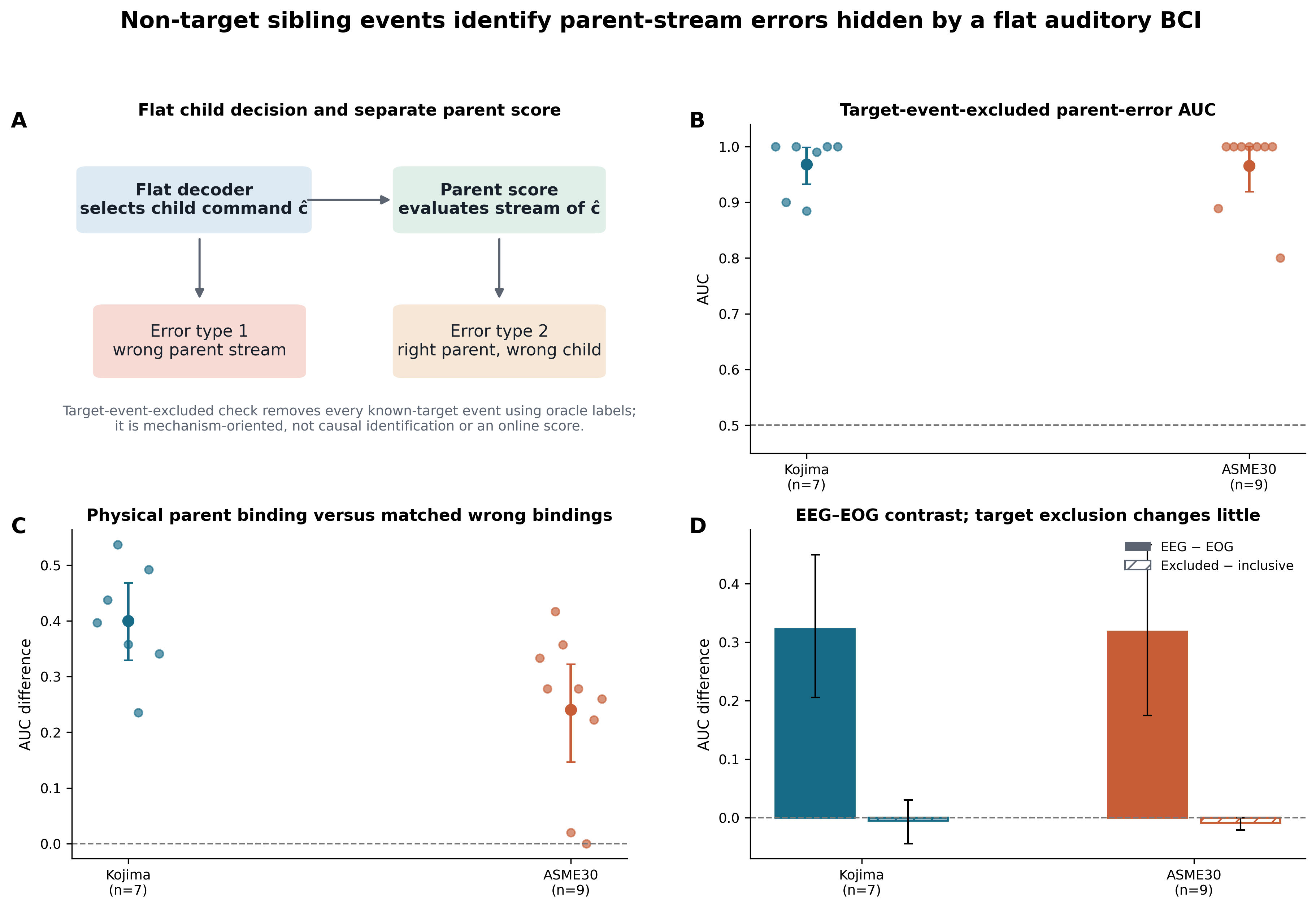

### Figure5_Event_Geometry_Transport.png

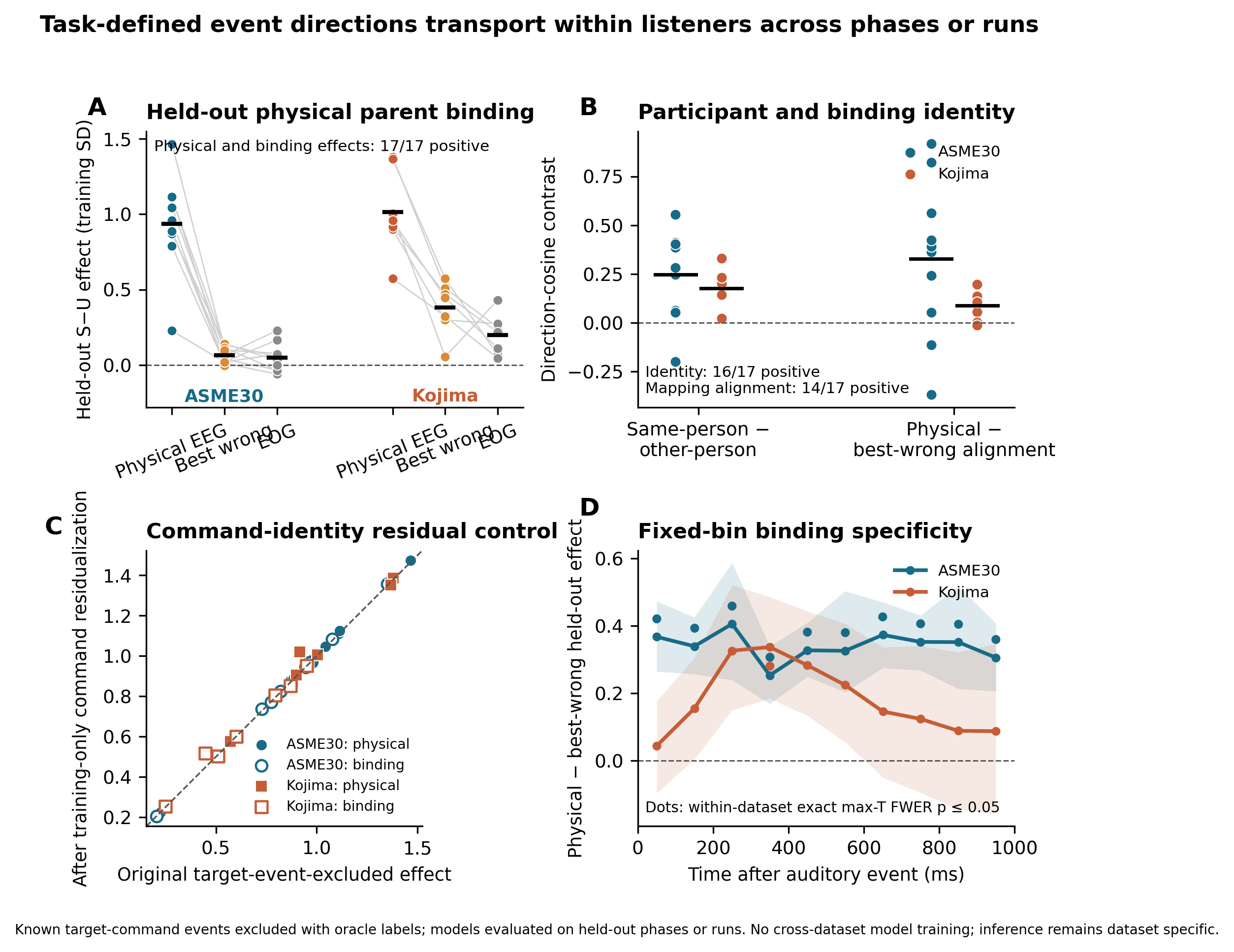

### Figure6_Channels_and_Calibration_Boundaries.pdf

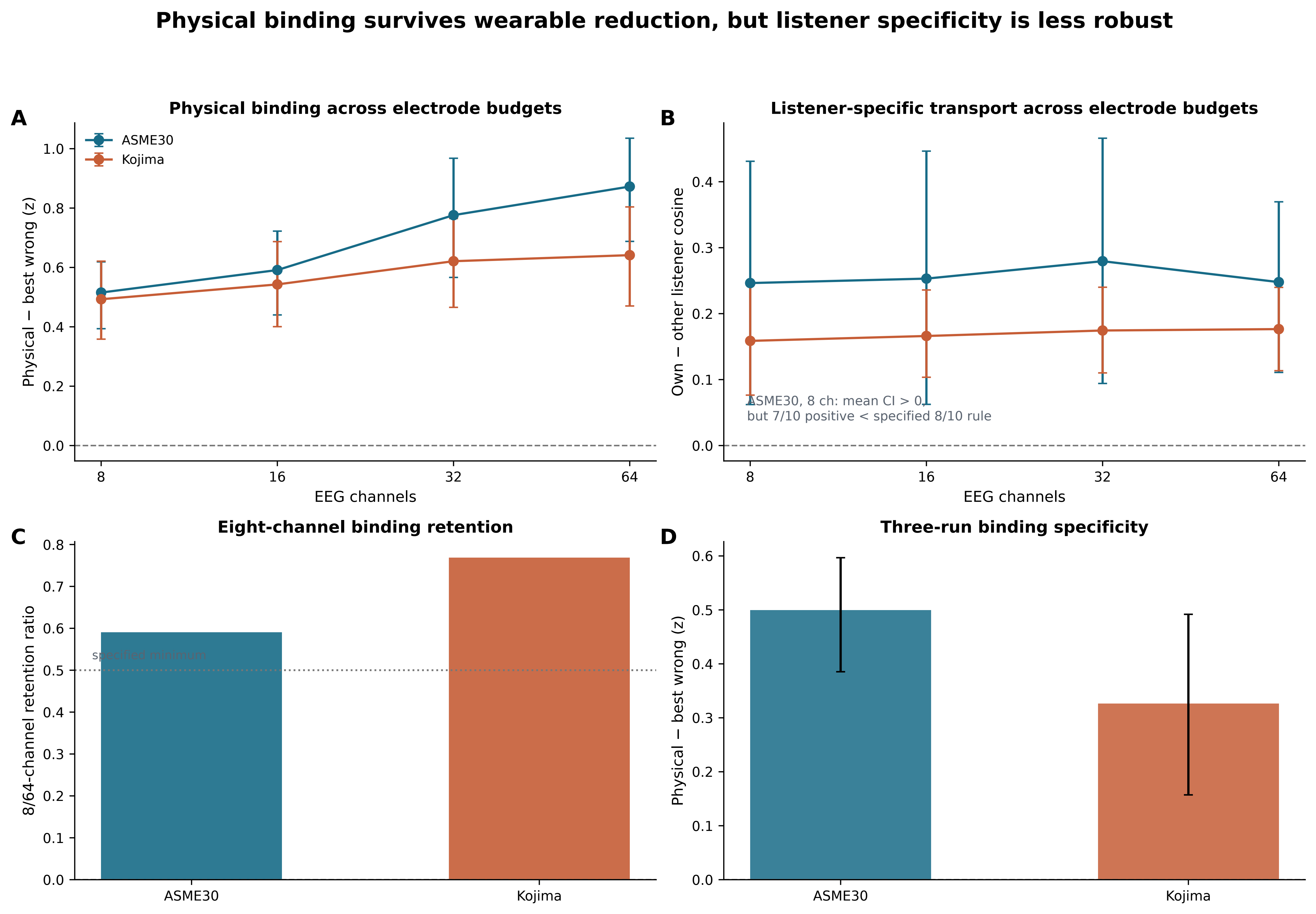

### Figure6_Channels_and_Calibration_Boundaries.png

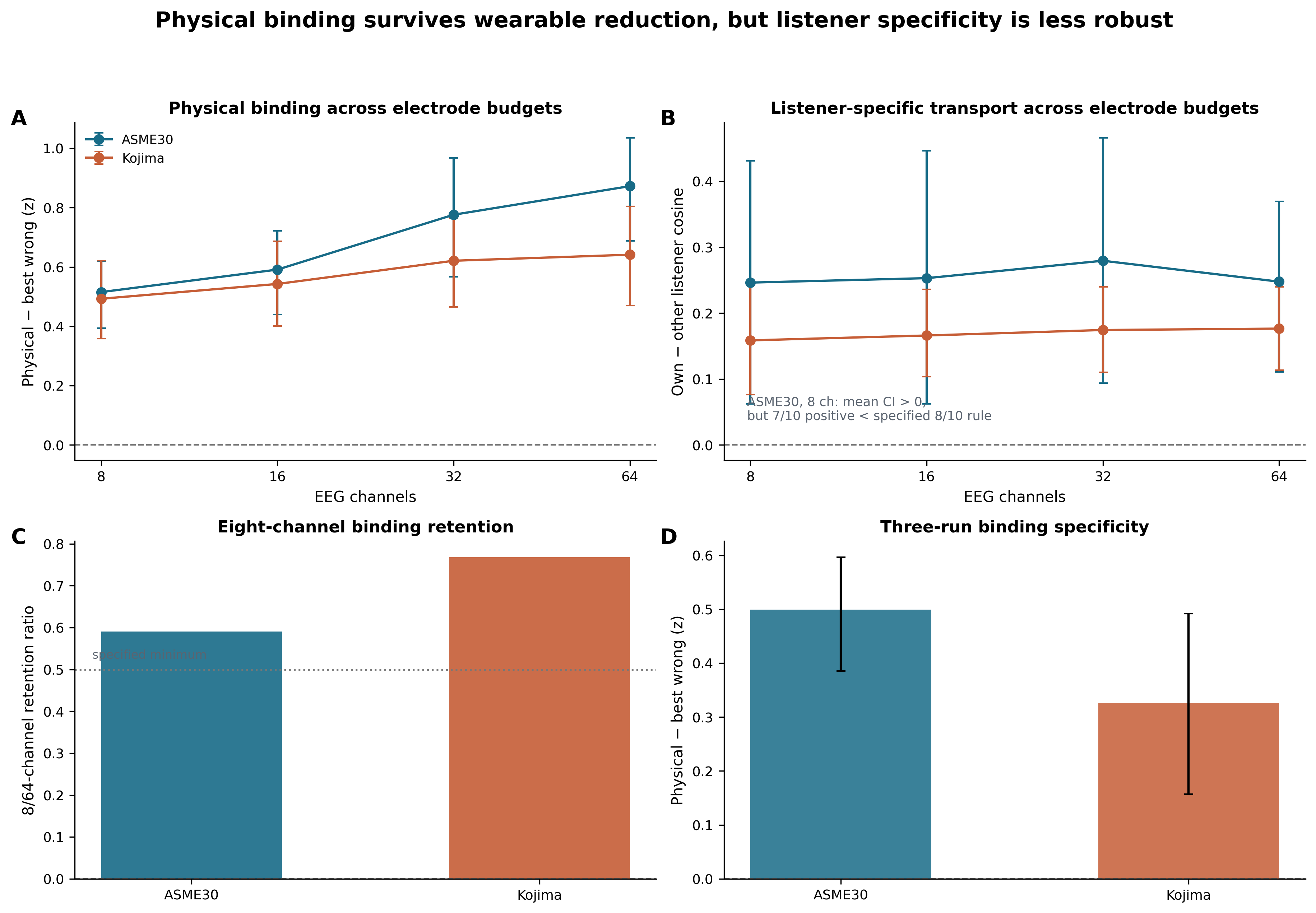
