## Supplementary_Information for "Falsifiable substitution tests reveal task-structured neural evidence for auditory attention"

Yu Ding

School of Psychology and Cognitive Sciences, Beijing Language and Culture University, Beijing 100083, China

### Contents

1. Scope, terminology, and evidence chronology
2. Public datasets and analysis populations
3. Mathematical inspiration and empirical translation
4. Projection-breadth analysis
5. High-density direction–delay bank
6. Continuous listener-transport experiment
7. Leave-one-listener pooled-template controls
8. Hierarchical target-event-excluded parent score
9. Cross-phase and cross-run individual geometry
10. Training-only command residualization
11. Wearable and calibration reductions
12. Statistical analysis
13. Negative results, stopped analyses, and implementation deviations
14. Analysis code, provenance, and data access
15. Archive contents supplied with this submission
16. SI References

### 1. Scope, terminology, and evidence chronology

This study is a public-data reanalysis. No participant was recruited and no new neural recording was made. Six unique public EEG resources, introduced in full in Table S2, supplied the primary analyses reported here. The same dataset can appear in more than one experiment; participant counts are therefore reported per estimand and are never added across analyses as though they were independent samples.

The mathematical and neural meanings of direction are related by experimental design, not by identity of mathematical objects. In Kakeya geometry, directions index the orientations of Euclidean line segments or tubes. Here a **neural direction family** is a fixed collection of neural–stimulus projections, scalp-direction-by-delay cells, or sensor-by-time contrasts indexed by an experimentally meaningful coordinate. **Physical organization** means that the coordinate relation is fixed by the recording or task arrangement rather than selected from the outcome—for example, which scalp sector is paired with which response delay, or which spoken command belongs to which pitch-defined stream. "Transport" means that a direction quantity estimated in a source partition retains a specified representational or held-score relationship in a disjoint target partition. It does not mean transporting another participant's raw EEG, high-dimensional decoder coefficients, feature means, feature scales, or target labels.

Four evidence labels are kept distinct:

- **Method-selecting discovery**: the candidate or hyperparameter was selected in the analyzed sample.
- **Rule recorded before outcome computation**: the candidate and primary endpoint were fixed in the local project before the relevant result was computed.
- **Post-primary mechanism test**: a new counterfactual endpoint was recorded after a related primary result was known but before the counterfactual result itself was computed.

- **Post-outcome robustness or descriptive boundary**: a later analysis that cannot upgrade an earlier inferential claim.

This study was not preregistered. Local protocol files and SHA-256 locks preserve the project's internal evidence chronology and are included in the supplied source-code archive; they should not be interpreted as a public preregistration.

#### Table S1. Evidence chronology

| Experiment | Timing of specification | Analysis population | Standing used in the manuscript |
|---|---|---|---|
| Four CCA margins versus the leading margin | Four-margin rule selected in odd-ID KUL | KUL even $n = 8$, DTU $n = 18$, Ear-SAAD $n = 15$ | Three tests with rules recorded before outcome computation; later study-equal synthesis is secondary |
| KITE-ST 16-cell bank | Bank selected in a 12-person development cohort | Participant-disjoint complete cases $n = 12$ | Independent-participant confirmation within one acquisition site |
| KITE-ST physical versus 15 wrong mappings | Specified before confirmation mapping scores | Same confirmation participants $n = 12$ | Post-primary mechanism test fixed before the remapping results |
| Continuous listener transport | Protocol recorded before F1–P2 outcomes | KITE-ST $n = 12$, KUL $n = 16$, Ear-SAAD $n = 15$ | Cross-dataset experiment fixed before endpoint computation |
| Leave-one-listener pooled templates | Specified after the individual-donor result but before pooled-template outcomes | Same 43 continuous-speech participants | Strong post-primary comparator |
| Target-event-excluded parent score | Fixed after parent-score development, before target-event-excluded outcomes | Kojima $n = 7$, ASME30 $n = 9$ evaluable for AUC | Oracle-label mechanism test fixed before the exclusion results |
| Event individual geometry | Dataset-specific protocols recorded before geometry outcomes | Kojima $n = 7$, ASME30 $n = 10$ | Cross-run/cross-phase transport fixed before endpoint computation |
| Command residualization | Specified before command-residual outcomes | Kojima $n = 7$, ASME30 $n = 10$ | Post-primary shortcut control fixed before its results |
| Wearable/calibration reduction | Specified after full-density results, before reduction outcomes | Kojima $n = 7$, ASME30 $n = 10$ | Post-primary joint robustness test |

### 2. Public datasets and analysis populations

#### Table S2. Public data resources

| Resource | Public locator | Recording/task | Use here |
|---|---|---|---|
| KU Leuven Auditory Attention Dataset (KUL) | Zenodo record 4004271; Biesmans et al. (1) | 64-channel two-talker continuous speech | Projection discovery/held participants; continuous transport |
| Technical University of Denmark auditory-attention dataset (DTU) | Zenodo record 1199011 | 64-channel two-talker continuous speech | Projection-breadth external test fixed before outcome computation |
| Ear-SAAD | doi:10.5281/zenodo.16536441; Geirnaert, Kappel, and Kidmose (2, 3) | Simultaneous 29-channel scalp, 19-channel around-ear, and 12-channel in-ear EEG | Multimodal projection test fixed before outcome computation; 29-channel scalp continuous transport |
| Ultra-high-density EEG-AAD | doi:10.5281/zenodo.4518754 | 255-channel acquisition; 253 scalp signals retained; two stories with left/right attention | Direction–delay bank, sensor density, continuous transport |
| Kojima ASME | doi:10.7910/DVN/1UJDV6; Kojima and Kanoh (4) | Four-class ASME auditory BCI | Target-event-excluded parent score; leave-one-run-out geometry; wearable analysis |
| ASME30 | doi:10.7910/DVN/TYRCWL | 30-command hierarchical auditory speller | Target-event-excluded parent score; offline-to-online geometry; wearable analysis |

All analyses used deidentified public releases under the source investigators' ethics approvals and repository terms. The supplied derived-result archive contains a six-row inventory of the public locator, repository-level license, recommended citation, sample used here, and role of each dataset. No attempt was made to reidentify participants.

#### KUL and DTU

The projection analysis used participant-specific regularized CCA. KUL odd-ID participants were the discovery set; even-ID participants remained held until the candidate rule had been specified. DTU included 18 participants and 60 trials per participant. For DTU, the first 64 released scalp channels were retained, the released 64-Hz EEG was modeled at 32 Hz, and signals were restricted to 1–9 Hz. The speech representation used 15 gammatone bands between 150 and 4,000 Hz, 0.6 power compression, a 1–9-Hz envelope, and 32-Hz output. EEG and speech lags spanned 0–500 ms in 31.25-ms steps. Outer test groups were unordered pairs of underlying speaker–story identities; every trial containing either held identity was excluded from training.

For continuous transport, all 16 KUL participants were used. Base-part identity groups $1, 4$ and $2, 3$ formed two balanced, nonoverlapping partitions. Each contained one left-attended and one right-attended part and one HRTF-filtered and one dry part.

#### Ear-SAAD

The prespecified projection test used all 15 participants and six 10-min trials. The preprocessed release is sampled at 20 Hz and band limited to 1–9 Hz. The released 19-band gammatone envelope with 0.6 compression was used. Modality-specific channel counts were 29 scalp, 19 around-ear, and 12 in-ear; Fp1-cr was excluded from pure ear modalities. Nonfinite released values were replaced by zero as specified by the source processing. Outer validation left out one entire trial. PCA was fitted on the remaining trials and retained at most 16 components. Neural and speech lags spanned 0–500 ms in 50-ms steps. The participant endpoint averaged the three simultaneous modalities equally, preventing repeated sensor measurements from being counted as separate participants.

For continuous transport, only the 29 scalp channels were used. We enumerated complementary 3-of-6 trial splits once by requiring trial 1 in the first half. Both halves had to include both attended-speaker labels and both video-condition labels. The chosen split minimized between-half imbalances and used lexicographic tie breaking.

#### KITE-ST high-density continuous speech

The source release contains 30 participants and two stories, each presented once with left and once with right attention. The project retained 253 scalp channels from the nominal 255-channel acquisition. Four participants (S20, S6, S8, and S7) were used only for engineering and quality-control work. Twelve participant-disjoint development cases (S4, S24, S11, S23, S14, S29, S19, S17, S12, S25, S27, and S5) selected the direction–delay bank. The remaining 14 participants formed the confirmation candidate list. S26 failed the specified first-recording signal-quality rule because 16 scalp channels exceeded the bad-channel threshold, above the allowed maximum of 15. The released archive for S3 lacked `S3_AAD_1L.ceo`; after applying the documented packaging exception, 19 scalp channels exceeded the same threshold. Both exclusions were made before either participant contributed an endpoint. The remaining 12 complete cases supplied the intact and patch analyses. Thus all 30 released participants are accounted for: 4 engineering/QC, 12 development, 2 confirmation exclusions, and 12 confirmation complete cases.

For the continuous transport analysis, partition A was story 1 (conditions 1L and 1R) and partition B was story 2 (2L and 2R). The confirmation complete cases were S0, S10, S2, S21, S28, S22, S9, S15, S16, S18, S13, and S1.

#### Kojima and ASME30 event data

Kojima analyses used the seven two-stream participants B, D, F, H, J, L, and N. Validation left one complete two-stream run out and fitted on the remaining runs. Commands 1–2 and 3–4 were the physical two-parent grouping; $1, 3/2, 4$ and $1, 4/2, 3$ were capacity-matched wrong groupings.

ASME30 used all ten participants for effect and geometry analyses. Six offline runs trained and three later online runs tested. The physical parents were the three pitch-defined streams corresponding to QWERTY rows. Two fixed wrong three-way partitions retained ten commands per parent. Nine participants had both parent-correct and parent-error trials and were evaluable for parent-error AUC. Participant 6 had 15 parent-correct and no parent-error flat predictions, so AUC was mathematically undefined rather than missing. This participant remained in effect, geometry, wearable, and the exploratory all-participant analyses for which both outcome classes were not required.

### 3. Mathematical inspiration and empirical translation

The three-dimensional Kakeya theorem concerns lower bounds on the volume of sets containing a unit segment in every direction. Central elements of Wang and Zahl's proof and the Guth–Wang–Zahl streamlined argument include family-level volume and multiplicity bounds, non-clustering conditions, organization through convex sets, induction across scales, and reduction to the sticky case. None of those theorems assumes neural data, and none proves a statement about EEG, attention, CCA, scalp topology, or event-related potentials.

The transfer is therefore an **experimental design map**, not a mathematical deduction:

#### Table S3. Proof-architecture concept and empirical test

| Proof-architecture concept | Auditory operationalization | Falsifying substitute |
|---|---|---|
| Control the family rather than one extremal tube | Use four ordered neural–speech margins or a 16-cell direction–delay family | Leading CCA margin or global bank |
| Track multiplicity/overlap rather than total energy alone | Repeated held contributions across cells or repeated non-target sibling events | A target-only or winner-only statistic |
| Rule out concentrated explanations | Require held-material transport and report failed static recurrence | Within-sample fit or one high-reliability cell |
| Challenge a proposed organization or factorization | Maintain physical scalp direction–delay or parent–child binding | Fifteen shift/swap mappings or two matched wrong parents |
| Refine across scale and partition | Transport across stories, trials, phases, runs, time bins, and sensor budgets | Same-partition evaluation |

This translation addresses a design problem in auditory decoding: common decoders often show that a feature is predictive but do not identify which part of its apparent attention sensitivity survives plausible substitutions. Each ingredient above has precedents in statistics, decoding, or cognitive neuroscience. The Kakeya-inspired contribution is their conjunction: a family-level statistic; repeated structured contributions; capacity-matched physical remappings; disjoint partitions or scales; and separate recurrence, held-score, and negative-control endpoints. That conjunction generated the 15 remappings, the separation of static recurrence from held-score ordering, and the negative scale tests. Capacity-matched remapping is our empirical translation of challenging an organization; it is not the same operation as mathematical factorization. The result is an experimental-design inspiration, not an application of the Kakeya theorem and not evidence that neural activity itself implements the proof.

### 4. Projection-breadth analysis

Within each outer-training fold, EEG normalization, PCA, lag construction, covariance estimates, and regularized CCA were fitted without the held stimulus identity. CCA provided paired neural and speech projections. For component $k$, a 5-s test window produced

$$M_k = \mathrm{corr}(E_k, S_{A,k}) - \mathrm{corr}(E_k, S_{U,k}),$$

where $A$ and $U$ denote attended and unattended speech. The leading-component baseline classified by the sign of $M_1$. The selected breadth candidate classified by

$$B_4 = \frac{1}{4}\sum_{k=1}^{4} M_k.$$

The fitted CCA was identical for both readouts. The comparison therefore isolated final-stage compression and did not add training parameters. Ties were scored as incorrect.

The four-margin rule was selected after a documented multi-method screen in KUL odd participants. For each of the following samples, the rule and endpoint were recorded before the result was computed: KUL even participants, DTU, and Ear-SAAD. Ear-SAAD contributed one participant value after equal averaging over its scalp, around-ear, and in-ear results.

#### Table S4. Prespecified projection-breadth tests

| Test | $n$ | $M_1$ accuracy | $B_4$ accuracy | Gain | Positive | Exact one-sided $P$ |
|---|---|---|---|---|---|---|
| KUL even | 8 | 0.600868 | 0.620845 | 0.019976 | 7/8 | 0.011719 |
| DTU | 18 | 0.616138 | 0.641667 | 0.025529 | 13/18 | 0.005043 |
| Ear-SAAD, modality-equal | 15 | 0.558721 | 0.573568 | 0.014846 | 12/15 | 0.003723 |
| Study-equal synthesis | 41 | — | — | 0.020117 | 32/41 | conservative replicability $P = 0.011719$ |

The study-equal stratified-bootstrap 95% interval was 0.012483–0.027905. Canonical-correlation-weighted four-margin scoring also exceeded $M_1$ in each study, but equal versus canonical weighting reversed sign across studies. The replicated result is breadth beyond the leading component, not a uniquely optimal weight formula.

### 5. High-density direction–delay bank

The KITE-ST decoder represented each training-story model with eight fixed scalp angular sectors crossed with two latency groups, producing 16 cells. At the full density, six nonnegative response lags between 0 and 250 ms were split into the first three and last three lags. Within a training story, even/odd window-index cross-fitting fitted on one parity and measured label-aligned held-parity cell contributions on the other, then reversed. Cell reliability was transformed by

$$w_j = 16\frac{\exp(2r_j)}{\sum_{\ell=1}^{16}\exp(2r_\ell)}.$$

The comparator pooled the same receiver information without the learned direction–delay weighting. Both were evaluated on the unseen story and averaged across the two story directions.

The complete-case confirmation produced a mean AUC gain of 0.054398 (95% interval, 0.018808–0.098669; 10/12 positive; exact one-sided $P = 0.003906$). The fixed 51-of-253 contiguous sensor-patch endpoint also confirmed (gain 0.050456, 95% interval 0.020580–0.085684; 10/12 positive; $P = 0.001709$).

#### Capacity-matched topology control

Cells were indexed $(s, b)$, with angular sector $s \in 0, \ldots, 7$ and latency $b \in 0, 1$. The 16 transformations

$$g(k, d) : (s, b) \mapsto ((s - k) \bmod 8,\ b \oplus d)$$

comprise eight circular shifts crossed with optional early/late exchange. $g(0, 0)$ is physical identity; the other 15 transformations preserve cell count, parameter count, and the exact fold-specific weight histogram.

Physical identity exceeded the mean of the 15 controls by 0.077836 AUC. All 12 participants were positive and the exact one-sided sign-flip value was 0.000244. Angular-only and delay-swapped advantages were 0.074694 and 0.080584 AUC, respectively. The learned distribution remained broad: effective cells 14.399 of 16, normalized entropy 0.9617, maximum cell mass 0.1272, and effective angular sectors 7.541 of 8.

The fixed-sequence reliability-stability endpoint failed (mean advantage 0.027780; 6/12 positive; $P = 0.376709$). Thus, held-story function required the physical topology, but a universally recurring static reliability profile was not supported.

Nested 32-, 64-, 128-, and 253-electrode subsets were chosen by outcome-independent farthest-point sampling beginning at Cz. The full model was retrained separately at every density. Bank gain and topology advantage remained positive at all four densities; neither the prespecified gain-by-density slope nor monotonicity prediction was supported. This is evidence against attributing the result simply to extreme electrode density.

### 6. Continuous listener-transport experiment

#### Common coordinate and source–target separation

Each source partition produced a 16-dimensional profile on the same eight-sector-by-two-delay coordinate. Reliability was the aligned held contribution mean divided by its root-mean-square and bounded in $[-1, 1]$. The fixed softmax-2 map converted the profile to positive weights with absolute sum 16. The physical map was zero angular shift without delay exchange; the 15 alternatives were all nonidentity elements of the transformation group above.

Source and target partitions shared no EEG window, trial, story, or base audio-part identity as applicable. Profiles, ridge coefficients, standardization, and reliability were obtained from source data only. The target supplied held contributions and labels to the fixed scoring rule. A donor supplied exactly 16 dimensionless profile values. Donor EEG, decoder coefficients, feature means, feature scales, and target information never entered the receiver.

Each participant contributed A-to-B and B-to-A transport. Participant endpoints averaged the two directions, so the 86 directions were not treated as 86 independent people.

#### Static endpoints

F1 ranked the correlation between a receiver target profile and its own source profile against correlations obtained after substituting every other listener's source profile within the same dataset. F2 ranked the physical own-profile correlation against the 15 transformed-own correlations.

#### Held-speech score endpoints

For P1, the receiver source-trained model and held-target cell contributions were fixed; only the 16 source weights were replaced by each donor's weights. For P2, only the physical association between weights and target cells was transformed. Scores were oriented to the attended side before comparison.

For every endpoint, ties counted 0.5 and rank advantage was

$$R = \Pr(\text{matched value exceeds substitute}) + \tfrac{1}{2}\Pr(\text{tie}) - 0.5.$$

Therefore $R = 0.10$ means a matched winning probability of 0.60; it is not a ten-percentage-point change in classification accuracy.

#### Table S5. Continuous transport, dataset-specific endpoints

| Endpoint | KITE-ST, $n = 12$ | KUL, $n = 16$ | Ear-SAAD, $n = 15$ | Study-equal mean (95% interval) |
|---|---|---|---|---|
| F1 own versus donor profile | −0.022727 [−0.162879, 0.121212] | 0.270833 [0.147917, 0.379167] | 0.254762 [0.102381, 0.385714] | 0.167623 [0.087861, 0.243065] |
| F2 physical versus wrong profile map | 0.016667 [−0.166667, 0.200000] | 0.379167 [0.270833, 0.454167] | 0.380000 [0.286667, 0.460000] | 0.258611 [0.182037, 0.333426] |
| P1 held speech, own versus donor weights | 0.075284 [0.035354, 0.119792] | 0.084091 [0.049715, 0.120833] | 0.131310 [0.075159, 0.190556] | 0.096895 [0.070975, 0.124063] |
| P2 held speech, physical versus wrong weights | 0.090972 [0.052083, 0.134144] | 0.142393 [0.110543, 0.173359] | 0.177963 [0.127852, 0.230963] | 0.137109 [0.113189, 0.162000] |

The fixed-seed cross-dataset randomization values were 0.000324, 0.000006, $< 10^{-6}$, and $< 10^{-6}$ for F1, F2, P1, and P2. All four conjunction criteria were satisfied. KITE-ST was null for both static endpoints but positive for both held-speech score endpoints. This dissociation prevents the stronger but unsupported claim of a universal static neural fingerprint. P1 and P2 concern matched ordering of fixed held-speech scores; they do not by themselves establish improved classification accuracy, calibration, or online BCI performance.

### 7. Leave-one-listener pooled-template controls

Individual donors might fail merely because each donor profile is noisy. We therefore froze two pooled controls after observing the individual-donor result:

- **Mean-profile template**: average every other listener's source reliability profile, then apply the prespecified softmax-2 transform.
- **Mean-weight template**: transform every other listener's profile, then average the resulting weights.

The receiver was excluded from both templates and all source–target separation rules were unchanged.

#### Table S6. Own weights versus pooled donor templates

| Endpoint | KITE-ST | KUL | Ear-SAAD | Study-equal mean (95% interval) |
|---|---|---|---|---|
| Own versus mean-profile template | 0.088542 | 0.114583 | 0.164444 | 0.122523 [0.089880, 0.155838] |
| Own versus mean-weight template | 0.088542 | 0.111742 | 0.165556 | 0.121947 [0.089156, 0.155072] |

Both fixed-seed randomization values were $< 10^{-6}$. However, changes in final classification AUC were heterogeneous: positive in KITE-ST, slightly negative in KUL, and approximately zero in Ear-SAAD. The result supports individualized ordering and scaling of held evidence, not universal accuracy, calibration, or confidence improvement.

### 8. Hierarchical target-event-excluded parent score

Let $T$ denote the target command, $S$ a non-target sibling in the attended physical parent, and $U$ a command in an unattended parent. An equal-prior Ledoit–Wolf LDA was fitted to $S$ versus $U$ events. Separately, the flat decoder chose a child command by averaging its first three accepted event scores and taking the maximum.

For a flat-predicted child $\hat{c}$, the signed parent score was

$$C(\hat{c}) = \mathrm{mean}_{e \in P(\hat{c})}\, q(e) - \max_{p \neq P(\hat{c})} \mathrm{mean}_{e \in p}\, q(e),$$

where $P(\hat{c})$ is the physical parent assigned to $\hat{c}$ and $q(e)$ is the outer $S - U$ score. AUC measured whether $C(\hat{c})$ distinguished flat predictions that crossed a parent boundary from those that remained in the correct parent.

The exclusion check deleted every event whose command equaled the known true target before parent pooling. The flat prediction, outer model, physical mapping, three-event horizon, and analysis windows remained fixed. Because deletion uses the known target label, this is an oracle-label, mechanism-oriented exclusion check rather than an online confidence algorithm or a causal identification strategy.

#### Table S7. Target-event-excluded parent score

| Dataset | Evaluable $n$ | Target-event-excluded AUC (95% interval) | Physical minus wrong binding | EEG minus EOG | Excluded minus inclusive |
|---|---|---|---|---|---|
| Kojima | 7 | 0.967772 [0.932434, 0.998599] | 0.399618 [0.329385, 0.467674] | 0.322057 [0.205649, 0.448983] | −0.005160 |
| ASME30 | 9 | 0.965432 [0.918519, 1.000000] | 0.240547 [0.146420, 0.322328] | 0.317937 [0.174443, 0.466490] | −0.008395 |

All evaluable participants exceeded AUC 0.5 for the absolute endpoint. The physical-minus-wrong and EEG-minus-EOG criteria were satisfied in both event datasets. EOG was formed and pooled identically to EEG. This comparison reduces a purely ocular account in these two datasets but cannot exclude every physiological or acquisition artifact, and it does not supply an ocular control for the continuous-speech datasets.

ASME30 participant 6 could not contribute to the AUC because all 15 flat predictions remained in the correct parent. To show that the principal mapping contrast did not depend on dropping this participant, we added a clearly post-audit, exploratory endpoint: for each participant, the mean target-event-excluded physical score across all 15 trials minus the better of the two wrong-mapping means. The effect was positive in 10/10 participants (mean 0.654161; exact one-sided sign-flip $P = 0.000977$); leave-one-participant-out means ranged from 0.584959 to 0.713882. Participant 6 contributed 1.276980. This analysis supports robustness of the mapping contrast, but it does not retroactively change the AUC population or its evidential standing.

The small AUC samples constrain attainable exact values and interval stability. ASME30 participant AUCs ranged from 0.800000 to 1.000000; leave-one-participant-out means ranged from 0.961111 to 0.986111, and the smallest attainable exhaustive one-sided value for $n = 9$ is $1/512 = 0.001953$. Kojima participant AUCs ranged from 0.884211 to 1.000000; leave-one-participant-out means ranged from 0.962401 to 0.981699, and the corresponding resolution for $n = 7$ is $1/128 = 0.007813$.

### 9. Cross-phase and cross-run individual geometry

Event EEG was represented as 64 electrodes by ten fixed 100-ms mean-amplitude bins from 0 to 1 s. ASME30 trained on six offline runs and tested on three later online runs. Kojima used leave-one-complete-run-out folds. Known target-command events were excluded from fitting, testing, and direction estimation.

For every physical or wrong mapping, a separate equal-prior Ledoit–Wolf $S - U$ model was fitted. Held scores were centered and scaled only with training-score statistics. The physical effect was the trial-balanced held mean standardized $S$ score minus $U$ score. Binding specificity subtracted the larger of the two independently trained wrong-mapping effects; EEG–EOG specificity subtracted the corresponding EOG effect.

For the direction analysis, training $S/U$ events standardized each feature. The source direction was

$$d_{\text{train}} = \mu_{S,\text{train}} - \mu_{U,\text{train}},$$

and the target direction was the trial-balanced held analogue. Within-listener alignment was their cosine. Listener specificity subtracted the mean cosine after replacing $d_{\text{train}}$ with another listener's training direction under the same partition design. Mapping alignment specificity subtracted the better wrong-mapping within-listener cosine.

#### Table S8. Individual event geometry

| Metric | ASME30, offline-to-online $n = 10$ | Kojima, cross-run $n = 7$ |
|---|---:|---:|
| Physical held effect | 0.935083 [0.737337, 1.107302] | 1.014133 [0.826552, 1.200715] |
| Binding specificity | 0.871092 [0.687419, 1.032255] | 0.632254 [0.456404, 0.803715] |
| EEG–EOG specificity | 0.885268 [0.671902, 1.083719] | 0.815221 [0.625419, 1.031600] |
| Listener-specific alignment | 0.248342 [0.111382, 0.369872] | 0.176172 [0.111639, 0.241200] |
| Physical-minus-wrong alignment | 0.329088 [0.093235, 0.555568] | 0.088070 [0.035193, 0.139904] |

Each dataset satisfied its seven internally defined coverage, physical, binding, EEG–EOG, listener, mapping-alignment, and fixed-time-bin criteria. Across the 17 participant rows, physical, binding, and EEG–EOG effects were descriptively positive in 17/17, listener specificity in 16/17, and mapping-alignment specificity in 14/17. Because the task designs and partitions are not exchangeable, no pooled 17-person $P$ value is reported.

Ten fixed time bins were tested with exhaustive sign flips and a two-sided maximum-$t$ statistic. This familywise procedure was defined before inspecting which latency was strongest and prevents selection of a favorable ERP interval.

### 10. Training-only command residualization

Stable command acoustics or labels could generate a repeatable $S - U$ direction even after target removal. Inside every outer-training partition, we estimated each command's mean from non-target events only. The corresponding training-derived command mean was subtracted from training and held events with that command. No held event or target-command event contributed to a residual mean.

All nine internally defined balance/coverage, physical, binding, listener, and mapping criteria were satisfied. Residual physical and binding effects were 0.939403 and 0.872515 in ASME30 and 1.028848 and 0.641402 in Kojima. Listener-specific alignment remained 0.247811 and 0.176570. Mapping-alignment specificity remained 0.304675 and 0.088267. Physical and binding retention ratios were approximately one. These results disfavor a stable command-mean explanation without establishing that all acoustic information was removed. In ASME30, the parent stream is itself defined by pitch; training-only command residualization cannot remove every parent-level acoustic cue. A prospective experiment that randomizes parent-first task instructions while holding the sound mixture fixed is required to separate task structure from parent acoustics causally.

### 11. Wearable and calibration reductions

The montage experiment used nested, geometry-defined 8-, 16-, 32-, and 64-electrode sets. The eight-electrode set was Fz, FCz, C3, Cz, C4, CPz, Pz, and Oz. Every montage was retrained; a full-density model was not merely evaluated after dropping channels.

ASME30 reduced calibration from all six offline runs to every combination of $k = 1, \ldots, 6$ runs. Kojima reduced the five training runs available inside each held-run fold to every $k$-run combination. Repeated subsets shared held observations, so they were averaged within participant before inference.

At eight electrodes, physical, binding, and EEG-minus-EOG criteria were satisfied for both datasets. Binding retention relative to 64 channels was 0.590735 in ASME30 and 0.768500 in Kojima, exceeding the defined 0.50 thresholds. Three-run physical and binding criteria were satisfied in both datasets.

The ASME30 eight-channel listener-specific mean was positive and its one-sided value was 0.0195, but only 7/10 participants were positive, below the specified 8/10 rule. Kojima satisfied its 6/7 rule. Hence W4 failed and the full W0–W7 wearable conjunction did not pass. Low-density EEG therefore preserved group-level physical binding more robustly than it preserved every participant's spatial direction.

### 12. Statistical analysis

The participant was the inferential unit. Scalar dataset summaries used participant percentile-bootstrap 95% intervals and exact exhaustive one-sided sign-flip tests where sample size permitted. Directional tests were used only for hypotheses whose sign was defined in advance by the matched-minus-substitute contrast: broader minus leading projection, physical minus remapped structure, own-listener minus donor, EEG minus EOG, or retained minus lost evidence. Two-sided maximum-$t$ tests were used for latency localization, where either sign could be selected across bins. Exact tests include zeros and use all $2^n$ sign patterns; their finite resolution is stated explicitly for the small event-dataset samples. Positive-count rules were specified where used.

Continuous cross-dataset summaries gave each dataset equal weight regardless of participant count. Confidence intervals used 100,000 participant resamples stratified by dataset. Randomization used 1,000,000 fixed-seed participant sign flips applied to the study-equal statistic and the add-one correction

$$P = \frac{1 + \#\{T_{\text{null}} \geq T_{\text{obs}}\}}{1 + N_{\text{draws}}}.$$

Projection synthesis similarly resampled participants within study and averaged the three study means equally. Its conservative replicability-conjunction value is the largest of the three one-sided values from the tests whose rules preceded outcome computation.

The F1–P2 transport endpoints formed an intersection requirement: one endpoint could not rescue another. The target-event-excluded analysis required all six absolute, mapping, and EEG–EOG criteria. Individual geometry and wearable tests also used conjunction rules. No pooled event inference was performed across ASME30 and Kojima. Time-localization tests used exact within-dataset maximum-$t$ familywise control across the ten bins fixed before analysis.

No a priori power analysis was possible because all samples were fixed by public availability and integrity criteria. Effect intervals and participant signs are therefore emphasized alongside $P$ values.

### 13. Negative results, stopped analyses, and implementation deviations

Negative results delimit the supported conclusions and are reported alongside successful tests.

#### Table S9. Material negative or nonpromoted results

| Branch | Result | Consequence |
|---|---|---|
| KITE-ST static reliability stability | Mean 0.027780; 6/12 positive; $P = 0.376709$ | No universal stable reliability-profile claim |
| Continuous F1/F2 in KITE-ST | −0.022727 and 0.016667; both intervals include zero | Static fingerprint not generalized to every dataset |
| Exact KITE bank transfer to KUL | Gain −0.019886 | High-density bank is not a universal external accuracy improvement |
| Exact KITE bank transfer to Ear-SAAD | Gain 0.005852; interval includes zero | Same boundary |
| Pooled-template AUC | Positive KITE-ST, slightly negative KUL, near zero Ear-SAAD | Held-score ordering/scaling conclusion only |
| Eight-channel listener consistency | ASME30 7/10 versus required 8/10 | One of eight wearable criteria failed |

Exploratory resources and candidate analyses outside these six datasets were not included in the present evidence set and are not reported as results. The dataset count, methods, tables, figures, and claim ledger therefore refer to the same six-resource scope.

### 14. Analysis code, provenance, and data access

The derived-result claim ledger binds 33 ledger-indexed quantitative claims to archive-internal JSON paths, source-file SHA-256 digests, evidence standing, and allowed interpretations. Every ledger source is checked automatically before packaging. A separate arithmetic audit recomputes all summary nodes supported by the included participant- and direction-level tables. The figure program reads final JSON/CSV artifacts and renders all six figures without refitting models. Figure 5 is drawn from the 17 participant-level individual-geometry rows and the two datasets' fixed-bin maximum-$t$ results; no completed figure is copied into the generated output.

The continuous source audit independently reconstructed 43 participant rows and 86 transport directions without importing the primary runner. Maximum discrepancies were $1.11 \times 10^{-16}$ at the participant level and $2.22 \times 10^{-16}$ at the direction level. A separate aggregate audit recomputed the fixed-seed randomizations and decision criteria with zero artifact-level discrepancy. Event, command-residual, and wearable branches have distinct source or artifact audits.

Two versioned archives accompany this submission. The source-code archive contains the preprocessing, feature extraction, model fitting, statistical, source-audit, and figure programs needed to run the reported workflows after the six source datasets have been downloaded. The derived-result verification archive contains deidentified tables and final summaries, the 33-claim ledger, a Python arithmetic audit, limited mean checks in R and MATLAB, figure inputs and code, protocols, audit summaries, dataset citation and license information, software licenses, and a member-level SHA-256 manifest. R checks the projection-breadth means and four continuous-speech study-equal endpoints; MATLAB R2015a was tested on Windows for the limited projection-breadth mean check. Neither is an independent implementation of the event, command-residual, or wearable analyses. These archives do not redistribute raw EEG and do not substitute for source-level model fitting. They are being prepared for permanent public deposition. Original code and documentation are licensed separately; derived data retain the applicable source-dataset terms.

### 15. Archive contents supplied with this submission

All paths and filenames in the supplied English archives use ASCII labels. `00_README.md` provides the run order, expected outputs, and a result-by-result map. `01_Result_Map.csv` links manuscript sections and figure panels to their inputs and checking code. `02_Dataset_Citations_and_Licenses.csv` records the six public resources. The directories `03_Derived_Data`, `04_Final_Results`, `05_Audit_Results`, `06_Current_Figures`, `07_Analysis_Code`, and `08_Generated_Outputs` separate data, evidence, audits, figures, executable checks, and generated outputs. `09_Claim_Evidence_Ledger.json` and its CSV companion provide human- and machine-readable bindings for the 33 ledger-indexed claims. `10_File_Manifest.json` records the byte size and SHA-256 digest of every archive member except itself; the companion CSV will be included in, and therefore checked by, the JSON manifest.
